# Stepwise origin and evolution of a transcriptional activator and repressor system integrating nutrient signaling in plants

**DOI:** 10.1101/2022.04.15.488190

**Authors:** Muhammed Jamsheer K, Rajesh Kumar Gazara, Sunita Jindal, Manoj Kumar

**Affiliations:** Amity Institute of Genome Engineering, Amity University Uttar Pradesh, Sector 125, Noida 201313, India; Department of Biotechnology, Indian Institute of Technology, Roorkee, Uttarakhand 247667, India; Department of Molecular Biology and Radiobiology, Faculty of AgriSciences, Mendel University in Brno, 61300, Brno, Czech Republic

**Keywords:** Nutrient signaling, Transcriptional Regulation, Gene duplication, Evolutionary Divergence, Molecular Evolution

## Abstract

Plants possess a unique transcriptional regulatory system in which two related MYB-related transcription factors (TFs) coordinate gene expression according to phosphate (Pi) and nitrogen (N) availability. The Phosphorus Starvation Response (PSR) type TFs are transcriptional activators integrating the cellular Pi sensing machinery and gene regulation majorly under Pi starvation. The Hypersensitivity To Low Pi-Elicited Primary Root Shortening (HRS) type TFs are transcriptional repressors integrating the Pi and N availability signals through different feedback loops. They are highly connected through multiple signaling loops to finetune the transcriptional responses according to nutrient availability. Molecular functions of these TFs are fairly uncovered in model systems; however, how plants evolved this activator-repressor system is currently unknown. In this study, using sensitive evolutionary analysis, we identified a stepwise origin of the PSR-HRS regulatory system in plants. The PSR TFs were originated before the split of Prasinodermophyta and Chlorophyta. The HRS TFs were originated later in the Streptrophycean algae. We also identified the asymmetric expansion of this TF repertoire in land plants majorly shaped by genome duplication and triplication events. The phylogenetic reconstruction coupled with motif analysis revealed that the origin of the specific accessory motifs is a major contributing factor in the functional divergence which led to the evolution of different sub-families preceding the angiosperm radiation. The spatiotemporal gene expression analysis in different developmental stages and nutrient availability conditions in angiosperms identified a critical role of expression divergence in shaping the functions of these TF families which is essential for adaptive plasticity of plants.

## Introduction

Inorganic phosphate (Pi) is an essential component of important biomolecules such as nucleotides and phospholipids. Organisms sense the Pi availability and modulate growth and development using specialized signaling pathways. However, Pi sensing and signaling mechanisms are fundamentally different in organisms such as bacteria, opisthokonts and plants (Bergwitz and Jüppner, 2011; Ham et al., 2018; Austin and Mayer, 2020). Nonetheless, studies suggest that the SPX (named after Suppressor of yeast *gpa1* (<u>S</u>yg1), the yeast Phosphatase 81 (<u>P</u>ho81), and the human Xenotropic and polytropic retrovirus receptor 1 (<u>X</u>pr1)) domain present in the proteins involved in Pi sensing and signaling machinery acts as a sensor of intracellular Pi level in eukaryotes (Wild et al. 2016; Azevedo and Saiardi 2017).

In eukaryotes, Pi starvation triggers a large-scale rearrangement of transcriptional, translational and metabolic networks directed at enhancing Pi uptake and internal recycling. These adjustments important for the survival of cells are orchestrated by specific transcription factors (TFs) in different eukaryotic lineages. In yeast, basic helix-loop-helix (bHLH) TF, Pho4 is a master regulator of Pi starvation responses (Bergwitz and Jüppner, 2011; Austin and Mayer, 2020). In Pi sufficiency, Pho80/85, a cyclin-dependent kinase (CDK) complex inactivates Pho4 by phosphorylation-mediated cytoplasmic retention. During Pi starvation, Pho80/85 activity is inhibited by the CDK inhibitor Pho81 and it allows the entry of Pho4 to the nucleus. Pho4 binds to specific motifs in the promoters leading to the expression of high-affinity Pi transporters, secreted acid phosphatases and other genes of Pi starvation responses (Austin and Mayer, 2020).

In plants, the transcriptional regulation according to Pi status is majorly orchestrated by the transcription activator Phosphorus Starvation Response (PSR) and repressor Hypersensitivity To Low Pi-Elicited Primary Root Shortening (HRS) TF subfamilies. They belong to the plant-specific <u>G</u>olden2, <u>AR</u>R-B, <u>P</u>SR1 (GARP) TF family with solitary MYB-related SHLQ(K/M)(Y/F) DNA binding domain (DBD) (Safi et al., 2017). Thus, the TFs regulating Pi starvation responses in plants are independently originated and are fundamentally different from other eukaryotes. Pi starvation regulates the expression of members of some other TF families (such as WRKY and AP2/ERF) (Jain et al., 2012). However, they are not directly linked with the central Pi sensing machinery (Wild et al. 2016; Safi et al. 2017; Ham et al. 2018; Ried et al. 2021).

A PSR type TF was initially identified from the green algae *Chlamydomonas reinhardtii* using a forward genetic screen (Wykoff et al., 1999). These TFs possess an N-terminal MYB-related SHLQ(K/M)(Y/F) domain followed by a coiled-coil (CC) domain. The transcription of *C. reinhardtii* PSR1 is induced during Pi starvation and it regulates the expression of Pi starvation responsive genes (Shimogawara et al., 1999; Wykoff et al., 1999). Later, PSR homologs were identified as master regulators of phosphate starvation responses in different angiosperms such as Arabidopsis, Brassica, rice, wheat, etc. (Rubio et al., 2001; Zhou et al., 2008; Ren et al., 2012; Wang et al., 2013). Thus, PSR subfamily TFs appear to be conserved master regulators of Pi starvation responses in the plant lineage. Intriguingly, the number of PSR TFs highly expanded in angiosperms. For example, Arabidopsis possesses 15 PSR subfamily TFs named Phosphate Starvation Response 1 (AthPHR1) and PHR1-LIKE (AthPHL1-14) (Bustos et al., 2010). Recent studies indicate that the AthPHR1 activity is under tight regulation of inositol pyrophosphate 8 (InsP8)-SPX domain-mediated Pi sensing mechanism and the CC domain of PHRs play an important role in this. In Pi sufficiency, AthPHR1 interacts with AthSPX1 which prevents its interaction with PHR1 binding site (P1BS) present in Pi starvation genes leading to the downregulation of AthPHR1 activity (Puga et al., 2014). Recently, the molecular mechanism of SPX domain-dependent regulation of AthPHR1 activity under different Pi availability was identified. During Pi sufficiency, InsP8 level is increased and InsP8-SPX complex binds to the CC domain preventing the AthPHR1 oligomerization and DNA binding (Ried et al., 2021). The SPX-dependent regulation of PSR/PHR TF activity is also found to be conserved in rice (Wang et al., 2014; Zhou et al., 2021). Taken together, these results suggest that the activity of PSR/PHR type TFs is tightly regulated according to the internal Pi availability through the InsP-SPX signaling module. Interestingly, PSR/PHR TF activity is also regulated by the nitrogen (N) availability. Nitrate signaling triggers the degradation of OsSPX4 in rice leading to the activation of OsPHR2 (Hu et al., 2019).

HRS1, the founding member of the HRS TF subfamily was identified from Arabidopsis (Liu et al., 2009). Arabidopsis possesses single HRS1 TF and six HRS1 Homologues (HHO1-6) and unlike PSR TFs, they function as transcriptional repressors (Sawaki et al., 2013; Medici et al., 2015; Kiba et al., 2018). Similar to PSR/PHR TFs, HRS/HHO TFs possess a conserved CC domain along with the solitary MYB-related SHLQ(K/M)(Y/F) DBD. The CC domain Arabidopsis HRS/HHO TFs was found to be important in dimerization which is essential for DNA binding (Ueda et al. 2020). Interestingly, HRS/HHO TFs work as integrators of Pi and N signaling (Sawaki et al., 2013; Medici et al., 2015; Maeda et al., 2018). Expression of these genes are rapidly induced by N sufficiency through NIN-Like Protein (NLP) TFs and therefore, they are also known as Nitrate-Inducible, GARP-type Transcriptional Repressors (NIGTs) (Liu et al., 2009; Sawaki et al., 2013; Medici et al., 2015; Maeda et al., 2018). AthHRS1 directly binds to the promoters of *SPX* genes and downregulates their expression (Ueda et al. 2020). Thus, HRS/HHO TFs work as activators of PSR/PHR TFs and phosphate starvation responses in plants. Interestingly, the expression of most of the AthHRS/HHOs is induced by PSR/PHR TFs while Pi starvation enhances the degradation of AthHRS1 (Sawaki et al., 2013; Medici et al., 2015; Maeda et al., 2018). Further, HRS/HHOs autoregulate their expression through a negative feedback loop (Sawaki et al., 2013; Maeda et al., 2018). These TFs are also involved in the negative regulation of nitrate (NO^-3^) uptake under Pi starvation by suppressing the expression of NO^-3^ transporter (Maeda et al., 2018; Wang et al., 2020). Similar regulatory functions of HRS/HHO TFs were also identified in rice and maize suggesting their role as important integrators of Pi and N uptake and signaling in land plants through multiple regulatory loops (Sawaki et al., 2013; Wang et al., 2020).

Pi and N are two major macronutrients and plant growth is dependent on the optimal availability of these nutrients. The coordinated activity of the transcriptional activator PSR/PHR and transcriptional repressor HRS/HHO subfamilies appear to be important for the regulation of gene expression according to the internal Pi and N levels. These TFs are majorly studied in angiosperms. However, how this transcriptional activator-repressor system is originated in plants is yet to be identified. Functional conservation of PSR TFs in green algae suggests an early origin of master regulators of phosphate starvation responses in the plant lineage (Wykoff et al., 1999). However, the evolutionary origin of the HRS/HHO regulatory system is completely unknown.

Angiosperms show dramatic expansion of TF gene families. The ancient and lineage-specific whole-genome duplications (WGD) are a major contributor to this expansion of the TF repertoire in angiosperms (Shiu et al., 2005; Lehti-Shiu et al., 2017). The PSR subfamily especially shows greater expansion in angiosperms (Bustos et al., 2010; Jain et al., 2012). Gene duplication may lead to increase in dosage if it is beneficial for the organism (Veitia, 2005). It can also relax the selection pressure leading to subfunctionalization or neofunctionalization. For example, neofunctionalization of TFs lead to the evolution of floral structures in different angiosperms (Rijpkema et al. 2006; Kramer et al. 2007; Mondragõn-Palomino and Theißen 2011). Divergence in the DNA binding site and/or other motifs/domains or origin of novel accessory motifs can alter the DNA binding specificity or important regulatory protein-protein interactions (Singh and Hannenhalli 2008; Shen et al. 2018; Brodsky et al. 2020). Further, changes in the promoter and regulatory regions can alter the spatiotemporal expression pattern of paralogs (Singh and Hannenhalli, 2008). The current knowledge indicates that PSR and HRS TF subfamilies are an important part of the adaptive mechanisms of plants to survive under different nutrient availability. However, nothing much is known about the evolution of these TFs in the plant lineage. Availability of new genomic and transcriptomic data sets of algae and early and late diverging land plants allow the in-depth analysis of the origin and evolution of gene families. In this study, using this information, we studied how the PSR and HRS TFs originated and diversified in the plant lineage.

## Results

### Identification of the origin of PSR and HRS TFs

We used a combination of sensitive BLAST and HMM-based searches to identify the PSR and HRS TFs from different basal genomes of Archaeplastida. In our analysis of Rhodophyta (*Cyanidioschyzon merolae*, *Porphyridium purpureum*, *Porphyra umbilicalis*, *Gracilariopsis chorda*, *Chondrus crispus*) and Glaucophyta (*Cyanophora paradoxa*) algae genomes, several proteins with MYB SHLQ(K/M)(Y/F) domain were identified. However, they lacked the typical CC domain characteristics of PSR and HRS TFs. A typical PSR TF with both MYB SHLQ(K/M)(Y/F) and CC domain was identified from *Prasinoderma coloniale*, a member of Prasinodermophyta, which diverged before the split of Chlorophyta and Streptophyta (Li et al. 2020) (Figure 1A, 2A, Supplementary Dataset 1). Further, typical PSRs were also found in all analyzed genomes of Chlorophyta and other aquatic and terrestrial algae (*Mesostigma viride*, *Chlorokybus atmophyticus*, *Chara braunii*, *Klebsormidium nitens*, *Mesotaenium endlicherianum*). However, HRS TFs were found to be absent in all analyzed genomes of Chlorophyta and they are present in early-diverging members of Streptophyta. This result suggests that the origin of HRS TFs coincides with the terrestrialization of plants (Figure 1A, 2A). Further, we also checked the presence of PSR and HRS-type TFs in other eukaryotic supergroups (Opisthokonta, Amoebozoa, Excavata and SAR)(Adl et al., 2012) using sensitive detection methods. However, we did not find any homologous proteins in the selected species of other eukaryotic supergroups suggesting that PSR and HRS TFs are specific to the plant lineage (Supplementary Figure 1).

**Figure 1.**
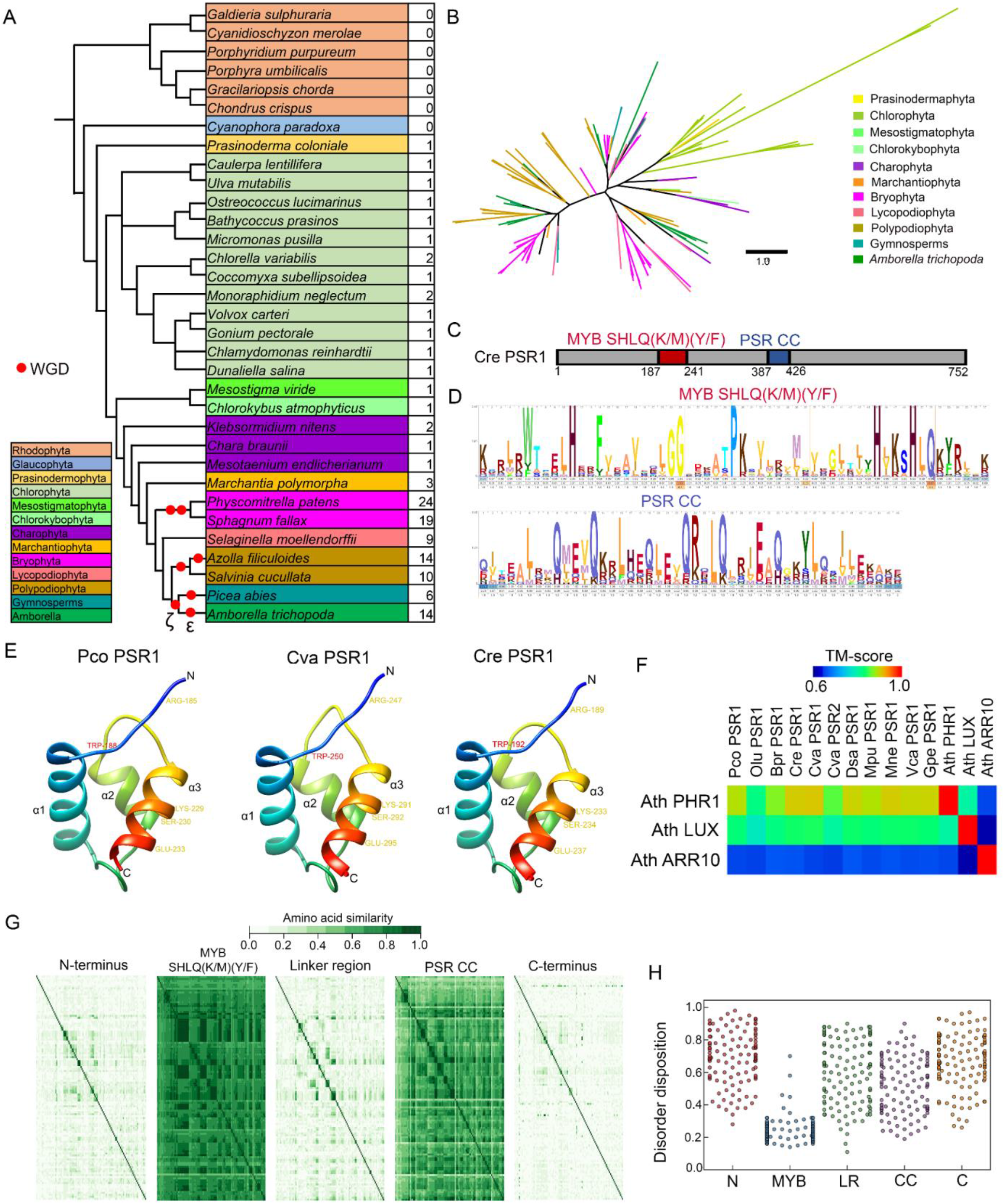
Origin and expansion of PSR transcription factor family in plants. (A) Number of PSR TFs in different species of Archaeplastida. (B) Simplified topology of the reconstructed phylogenetic tree of the PSR TF family across the plant lineage. Branches are colored according to taxonomic groups. The phylogenetic tree reconstruction was performed using the maximum likelihood estimation method based on the JTT+CAT substitution model with 1000 bootstrap replicates. The detailed visualization of the phylogenetic tree is given in Supplementary Figure S2. (C) The typical domain composition of PSR TFs. The *Chlamydomonas reinhar*dtii PSR1 is shown as the representative. (D) HMM profile of MYB SHLQ(K/M)(Y/F) and Coiled-Coil (CC) domains of PSR TF family. (E) Homology-based models of MYB SHLQ(K/M)(Y/F) from selected algal PSRs. The position of conserved tryptophan (red) and residues important for DNA recognition and binding (yellow) are indicated. (F) Structural similarity of MYB SHLQ(K/M)(Y/F) from different PSRs with MYB SHLQ(K/M)(Y/F) of Arabidopsis GARP TFs. (G) Sequence similarity of different regions of PSR proteins across the plant lineage. (H) Average disorder score of different regions of PSR proteins across the plant lineage.

**Figure 2.**
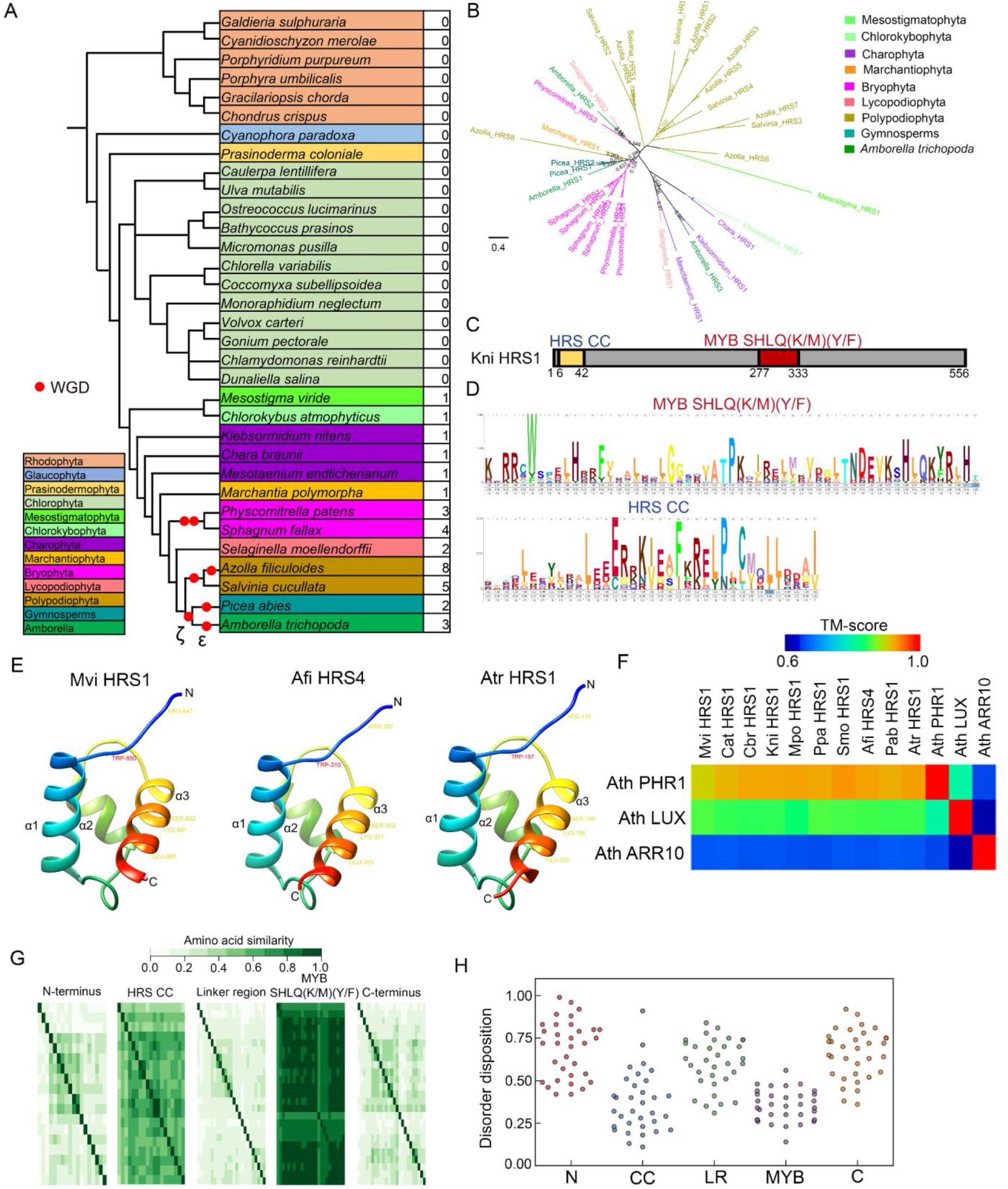
Origin and evolution of HRS transcription factor family in plants. (A) Number of HRS TFs in different species of Archaeplastida. (B) Phylogenetic reconstruction of HRS TF family across the plant lineage. Branches are colored according to taxonomic groups. The phylogenetic tree reconstruction was performed using the maximum likelihood estimation method based on the JTT+CAT substitution model with 1000 bootstrap replicates. (C) The typical domain composition of HRS TFs. The *Klebsormidium nitens* HRS1 is shown as the representative. (D) HMM profile of MYB SHLQ(K/M)(Y/F) and Coiled-Coil (CC) domains of HRS TF family. (E) Homology-based models of MYB SHLQ(K/M)(Y/F) from selected algal HRSs. The position of conserved tryptophan (red) and residues important for DNA recognition and binding (yellow) are indicated. (F) Structural similarity of MYB SHLQ(K/M)(Y/F) from different HRSs with MYB SHLQ(K/M)(Y/F) of Arabidopsis GARP TFs. (G) Sequence similarity of different regions of HRS proteins across the plant lineage. (H) Average disorder score of different regions of HRS proteins across the plant lineage.

To identify the early evolutionary pattern of these TFs, we analyzed the sequenced genomes of Marchantiophyta (*Marchantia polymorpha*), Bryophyta (*Physcomitrella patens*, *Sphagnum fallax*), Lycopodiophyta (Selaginella *moellendorffii*), Polypodiophyta (*Azolla filiculoides*, *Salvinia cucullata*), Pinophyta (*Picea abies*) and basal angiosperm *Amborella trichopoda*. Interestingly, in comparison to algal genomes, the number of both PSR and HRS TFs was found to be increased in these genomes (Figure 1A, 2A, Supplementary Dataset 1). However, the increase was more prominent in the PSR TF subfamily especially in genomes with lineage-specific WGDs such as *P. patens*, *S. fallax*, *A. filiculoides* and *S. cucullata* (Lang et al., 2018a; Li et al., 2018). Collectively, these results ascertain that the origin of PSR transcriptional activators precede the origin of HRS transcriptional repressors. Further, the emergence of HRS TFs coincides with the terrestrialization of the plant lineage.

### Analysis of the early evolution of PSR and HRS TFs in the plant lineage

To identify the early evolutionary patterns of PSR and HRS TFs, maximum likelihood phylogenetic reconstruction was performed using the identified protein sequences from aquatic and terrestrial algae (Prasinodermophyta, Chlorophyta, Mesostigmatophyta, Chlorokybophyta and Charophyta), Marchantiophyta, Bryophyta, Lycopodiophyta, Polypodiophyta, Pinophyta and basal angiosperm *A. trichopoda*. We included *A. trichopoda* sequences in our analysis as it is a reference point for angiosperm evolution and gene family size (Albert et al., 2013). In the phylogenetic reconstruction of PSRs, the PSR proteins from Chlorophyta were recovered along with *P. coloniale* PSR to a statistically supported clade (bootstrap value: 0.749), indicating the sequence divergence from PSR proteins from other species (Figure 1B, Supplementary Figure 2). Interestingly, most of the duplicated PSR proteins of Bryophyta and Polypodiophyta were recovered to specific clades with shorter branch lengths. The *A. trichopoda* PSR proteins also recovered to specific clades. However, in comparison, the branch length was higher. In the phylogenetic reconstruction of HRS TFs, duplicated HRS proteins of Bryophyta and Polypodiophyta were recovered to specific clades (Figure 2B). Interestingly, all three *A. trichopoda* HRS proteins were recovered to different clades, indicating the possible functional specialization of HRS proteins predating the angiosperm divergence.

We further analyzed potential domain gain in both PSR and HRS TFs. Domain pattern analysis revealed that the typical PSR (N-terminal MYB SHLQ(K/M)(Y/F) and C-terminal CC domains) and HRS (N-terminal CC and C-terminal MYB SHLQ(K/M)(Y/F)) pattern is highly conserved (Figure 1C, 2C). Further, the specific residues in both MYB SHLQ(K/M)(Y/F) and CC domain important for DNA recognition, binding and dimer/tetramerization, and interaction with SPX domain (in case of PSR) are fairly conserved across the plant kingdom (Figure 1D, 2D, Supplementary Figure 3 and 4). The structure of AthPHR1 MYB SHLQ(K/M)(Y/F) is deduced which shows the conserved MYB fold with three α-helices (α1, α2, and α3) and an N-terminal flexible arm region (Jiang et al. 2019). Homology modeling of MYB SHLQ(K/M)(Y/F) domains of PSR and HRS TFs identified that MYB fold and N-terminal arm region are conserved from algae to angiosperms (Figure 1E, 2E). Among the plant MYB SHLQ(K/M)(Y/F) TFs, structure of MYB SHLQ(K/M)(Y/F) of Arabidopsis Response Regulator 10 (ARR10) and LUX is also resolved and they also possess the conserved MYB fold with three α-helices. However, α3 of PHR1 MYB SHLQ(K/M)(Y/F) domain is larger than ARR10 and LUX MYB SHLQ(K/M)(Y/F) domains with different three-dimension orientation. These differences are important in DNA recognition and define the target genes (Jiang et al. 2019). Therefore, to understand the conservation of three-dimensional structural topology, the homology models of selected MYB SHLQ(K/M)(Y/F) domains of PSR and HRS TFs identified in this study was structurally aligned with PHR1, ARR10 and LUX MYB SHLQ(K/M)(Y/F) domains. In our analysis, PSR and HRS MYB SHLQ(K/M)(Y/F) domains selected from different taxonomic groups showed highest similarity with PHR1 MYB SHLQ(K/M)(Y/F) domain indicating the strong structural conservation in the DBD across the plant lineage (Figure 1F, 2F). Further, we also analyzed the sequence conservation of different regions of these TFs. In line with the structural conservation, the MYB SHLQ(K/M)(Y/F) domain showed highest sequence conservation in both PHR and HRS TF subfamilies (Figure 1G, 2G). The CC domain also showed a fair degree of sequence conservation. Interestingly, the N- and C-terminus and the linker region connecting MYB SHLQ(K/M)(Y/F) and CC domains showed very low sequence conservation. In many TF families including MYBs, these non-conserved regions possess intrinsically disordered regions (IDRs) which contribute to specific DNA and protein binding (Millard et al. 2019; Brodsky et al. 2020). Therefore, we tested the disorder propensity of different regions of these TFs using the meta-predictor PONDR-FIT that shows very high accuracy over individual predictors (Xue et al. 2010). In both PSR and HRS TFs, N and C termini and linker region connecting MYB SHLQ(K/M)(Y/F) and CC domains predicted to have high propensity for disorder (Figure 1H, 2H). In line with this, we found many short and long IDRs (SIDR and LIDR) in these regions (Supplementary Figure 5). Interestingly, the CC region of PSR TFs also showed relatively high propensity for disorder due to the presence of IDRs in the junction of CC domain with other regions (Figure 1H; Supplementary Figure 5A). Structural analysis of AthPHR1 CC domain revealed that the loop region is disordered that might be the reason for the prediction of IDRS in CC domain region in some cases (Ried et al., 2021).

### Analysis of the evolution of PSR and HRS TFs in angiosperms

Angiosperms show very high expansion of TF gene family size which led to functional divergence and origin of novel functions (Lehti-Shiu et al., 2017). Phylogenetic reconstruction with *A. trichopoda* PSR and HRS TFs suggests functional divergence early in the angiosperm evolution. Further, in comparison to algal genomes, angiosperm genomes such as Arabidopsis shows high expansion of PSR and HRS TFs. Therefore, to study the evolution of these TFs in angiosperms, we analyzed the genomes of selected eudicots and monocots. We found that the number of PSR and HRS TFs are highly expanded in both eudicot and monocot genomes (Supplementary Figure 6, Supplementary Dataset 1). This ancient and lineage-specific WGD and triplication (WGT) events in angiosperms can be a major factor for this expansion (Jiao et al., 2011; Qiao et al., 2019). In line with this, the polyploid species with lineage-specific WGD/WGT events possess more copies of these TFs in their genome than closely related species. For example, *Arabidopsis thaliana* has 15 PHR and 7 HRS TFs while *Brassica rapa* has 28 PHR and 14 HRS TFs. In addition to the two WGD (β and α) events common to Brassicaceae, *B. rapa* underwent an additional WGT event after the split of Arabidopsis and Brassica (Wang et al. 2011). Similarly, *Medicago truncatula* has 17 PHR and 5 HRS TFs while *Glycine max* genome that underwent a recent (~13 Mya) ago possesses 38 PHR and 15 HRS TFs (Schmutz et al. 2010). Although the number is considerably expanded, the domain composition analysis identified that the ancestral domain pattern of both PHR and HRS TFs is highly conserved in angiosperms. Further, the size and sequence similarity of MYB SHLQ(K/M)(Y/F) and CC domains were found to be highly conserved in angiosperms (Supplementary Figure 7-10). However, similar to ancestral PHR and HRS TFs, the N- and C-terminus and the linker region connecting MYB SHLQ(K/M)(Y/F) and CC domains possess very low sequence conservation in angiosperms (Supplementary Figure 7, 9).

Our analysis reveals a clear expansion in the PSR and HRS TF repertoire in the angiosperm genomes. Therefore, to identify the evolutionary patterns of the PSR and HRS TFs in angiosperms, a maximum likelihood phylogenetic reconstruction was performed. To aid the evolutionary analysis, we also included the ancestral sequences from aquatic and terrestrial algae, Marchantiophyta, Bryophyta, Lycopodiophyta, Polypodiophyta and Pinophyta. In the phylogenetic analysis, most of the PSR and HRS proteins were recovered into distinct clades (Figure 3, Figure 4A). We annotated the clade in which most of the proteins from basal plant genomes were recovered as subfamily I (SFI) in both phylogenetic trees. In the PSR phylogenetic tree, PSR SFI includes proteins from Chlorophyta and other aquatic and terrestrial algae along with proteins from Bryophyta, Lycopodiophyta, Polypodiophyta, Pinophyta and angiosperms (Supplementary Figure 11). Similarly, HRS SFI includes most of the proteins from aquatic and terrestrial algae (Mesostigmatophyta, Chlorokybophyta and Charophyta), all members of Polypodiophyta and few proteins from angiosperms (Supplementary Figure 12). Other distinct clades were named as SFs according to their relative closeness to SFI in both PSR and HRS phylogenetic tree. In the PSR phylogeny, we identified that except a few diverged members, all other members were recovered to eight distinct subfamilies including the ancestral PSR SFI (Figure 3; Supplementary Figure 11). In the case of HRS, four SFs were recovered, including the HRS SFI (Figure 4A; Supplementary Figure 12). Interestingly, the members from the *A. trichopoda* recovered to all SF clades in the PSR phylogenetic tree indicating the sequence divergence at the base of angiosperm evolution (Figure 3; Supplementary Figure 11). Supporting this hypothesis, the members from other angiosperms were found to be recovered in all SF clades with very few exceptions (for e.g., PSR SFII is absent in the analyzed legume genomes). Similar pattern was also observed in HRS TFs (Figure 4A; Supplementary Figure 12). For example, *A. trichopoda* possesses three HRS TFs and they were recovered to different clades (i.e., HRS SFII, III and IV). Further, most of the members from angiosperms were also recovered to clades corresponding to HRS SFII, III and IV (Figure 4A; Supplementary Figure 12). Collectively, our phylogenetic reconstruction suggests divergence of both PSR and HRS TF families before the angiosperm radiation. It could be possible that these SFs of PSR and HRS TFs might have specialized functions enhancing the TF repertoire available for responding to changes in nutrient availability in angiosperms.

**Figure 3.**
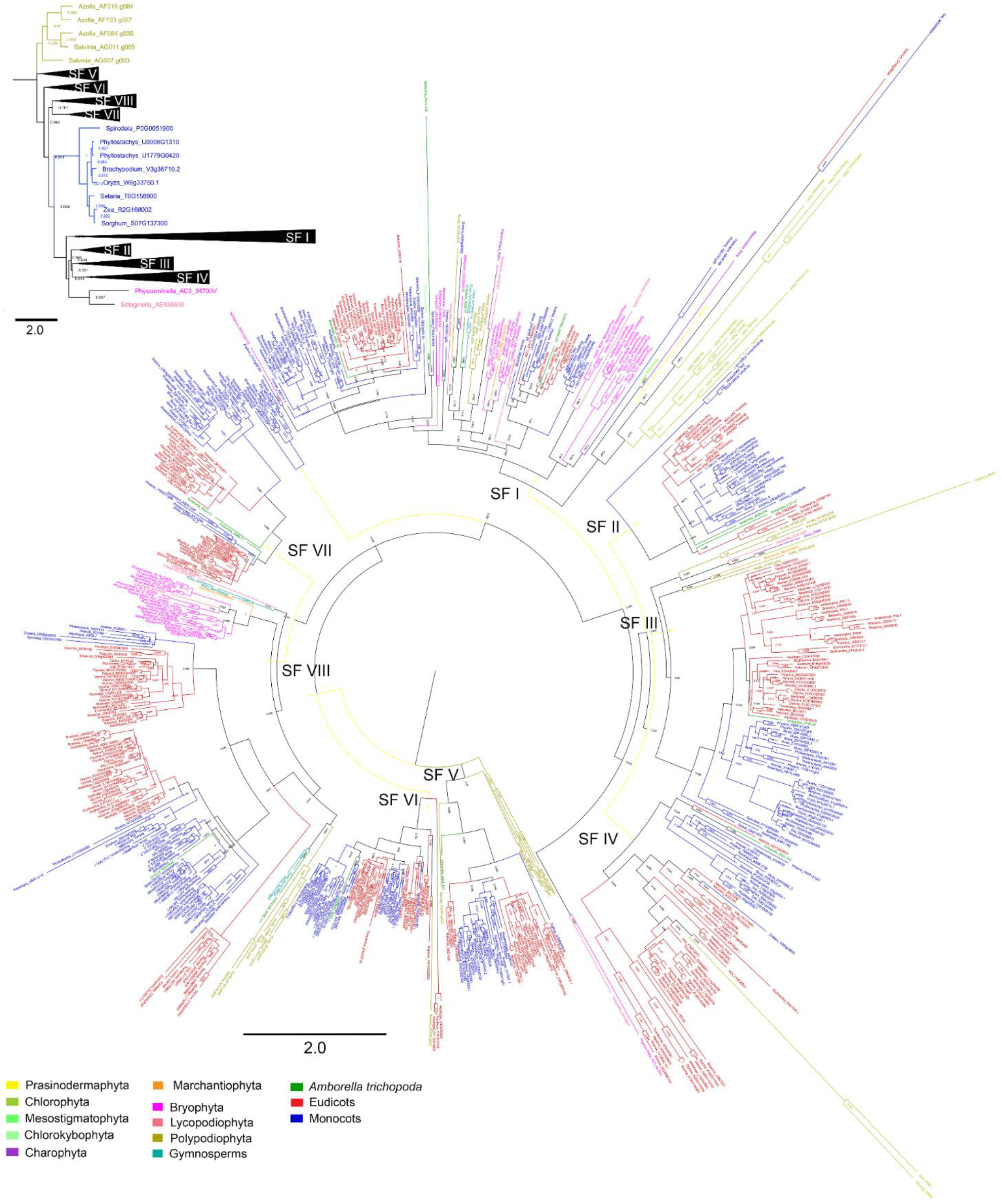
Phylogenetic reconstruction of PSR transcription factor family in the plant lineage. The simplified topology representation of the reconstructed phylogenetic tree showing different subfamilies (collapsed) is given at the top and the detailed phylogenetic tree is given below. The phylogenetic tree reconstruction was performed using the maximum likelihood estimation method based on the JTT+CAT substitution model with 1000 bootstrap replicates. Branches are colored according to taxonomic groups and different subfamilies are indicated.

**Figure 4.**
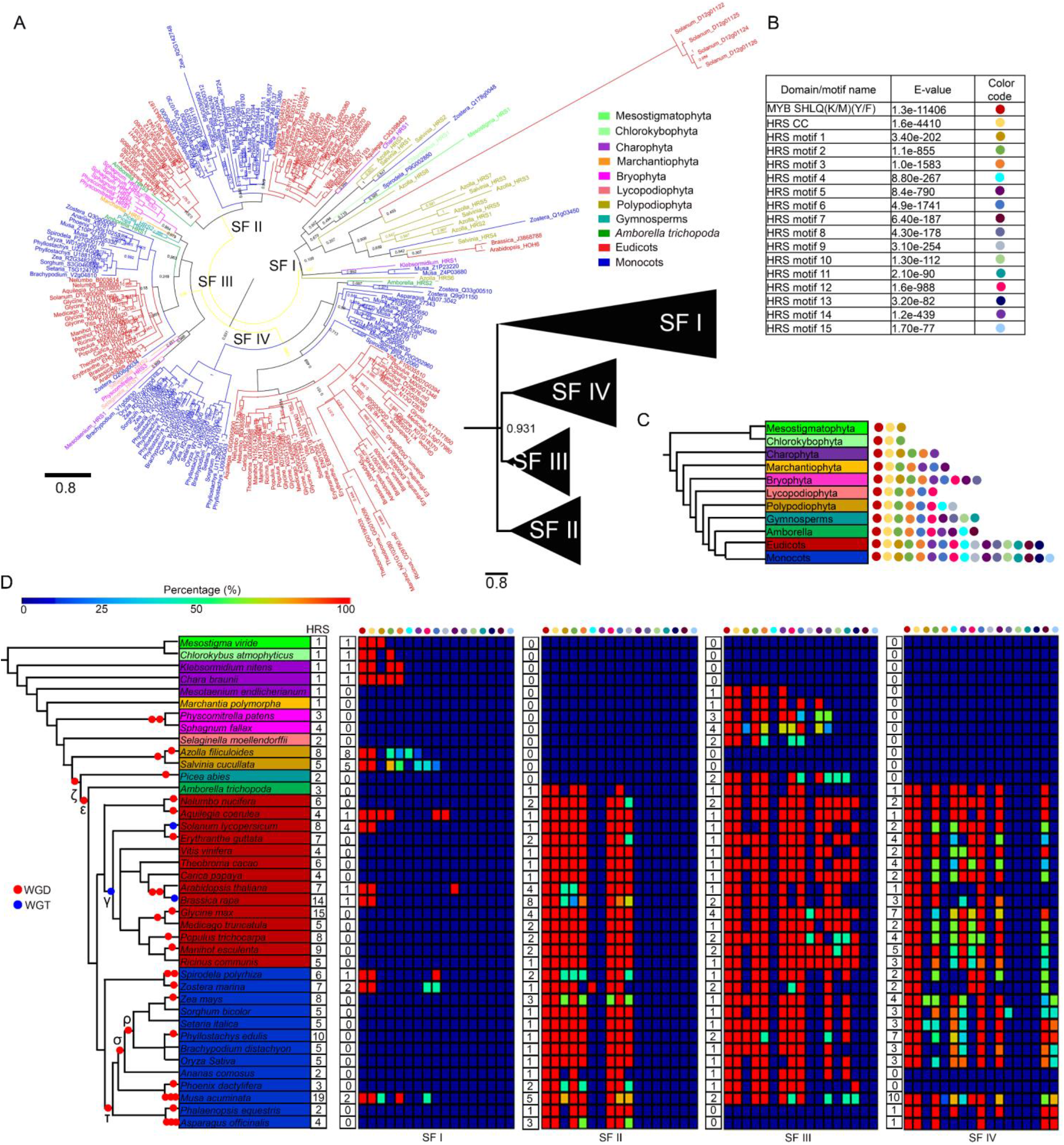
Phylogenetic reconstruction and motif analysis of HRS transcription factor family in the plant lineage. (A) The phylogenetic reconstruction of HRS TF family in the plant lineage. The detailed phylogenetic tree and simplified topology representation of the reconstructed phylogenetic tree showing different subfamilies (collapsed) are shown. The phylogenetic tree reconstruction was performed using the maximum likelihood estimation method based on the JTT+CAT substitution model with 1000 bootstrap replicates. Branches are colored according to taxonomic groups and different subfamilies are indicated. (B) Novel motifs identified from HRS TF family in the plant lineage. (C) Taxonomical categories of Archaeplastida showing the presence or absence of specific novel motifs. (D) Conservation of novel protein motifs in different subfamilies of HRS TF family. The detailed visualization of domain/motif arrangement in different HRS TF subfamilies is given Supplementary Figure S14.

### Origin and evolution of accessory motifs in PSR and HRS TFs

Recovery of PSR and HRS proteins to distinct clades suggests strong sequence divergence and possible functional specialization. Our analysis identified that the MYB SHLQ(K/M)(Y/F) and CC domains are highly conserved while the IDR enriched N, C and linker region connecting domains are non-conserved across the plant lineage (Figure 1G, 2G; Supplementary Figure 7, 9). Origin of novel motifs especially in the non-conserved regions was found to be a major factor in the protein divergence and functional specialization of TFs (Nguyen Ba et al. 2014; Cheatle Jarvela and Hinman 2015; Brodsky et al. 2020). These novel motifs may alter the protein-protein interaction and DNA recognition properties of TFs (Cheatle Jarvela and Hinman, 2015).

To identify whether the PSR and HRS TFs possess any novel motifs, we performed an in depth *de novo* motif discovery across the plant lineage. In our analysis, we identified 24 and 15 novel motifs with very strong statistical support in PSR and HRS TFs, respectively (Figure 4B, 5A, Supplementary Dataset 2). Interestingly, majority of these motifs were exclusive to land plants (Figure 4C, 5B). For example, the PSR TFs from aquatic and terrestrial algae were found to have only the MYB SHLQ(K/M)(Y/F) and CC domains. In case of HRS TFs, streptophyte algae also possess additional motifs such as HRS motif 1 and 2. Nevertheless, angiosperms PSR and HRS TFs have more diversity in motifs. We also identified monocot and eudicot-specific motifs in our analysis. For example, the PSR motif 15 is specific to eudicots while PSR motif 19 and 24 are specific to monocots. In HRS TFs, motif 15 was found to be specific to monocots (Figure 4C, 5B). We hypothesized that these novel motifs could be a major reason for the formation of PSR and HRS SFs in the phylogenetic reconstruction analysis. To test this, we analyzed the distribution of these motifs in SFs recovered in the phylogenetic reconstruction. We found that specific motifs and combinations are characteristics of specific subfamilies (Figure 4D, 5C, Supplementary Figure 13, 14). The SF I containing majority of the PSR and HRS TFs from algal and other lower plants possess relatively simple motif composition. In comparison, some SFs that are especially conserved in angiosperms such as PSR SF VIII and HRS SF III showed very complex motif composition. We also found some exceptions to this in PSR TFs, in which PSR SF IV which is highly conserved in angiosperms possesses simple motif composition. Interestingly, the conservation of specific motif composition was found to be varied in different SFs. For example, PSR SF II and HRS SF II showed very high conservation while PSR SF VIII and HRS SF IV showed high degree of divergence in motif composition among the members (Supplementary Figure 15, 16). Taken together, the phylogenetic reconstruction and *de novo* motif analysis identified that PSR and HRS TFs underwent a high degree of sequence divergence and specialization, especially in angiosperms. The high conservation of these motifs across the plant lineage suggests important biological functions of these motifs in the activity of PSR and HRS TFs.

**Figure 5.**
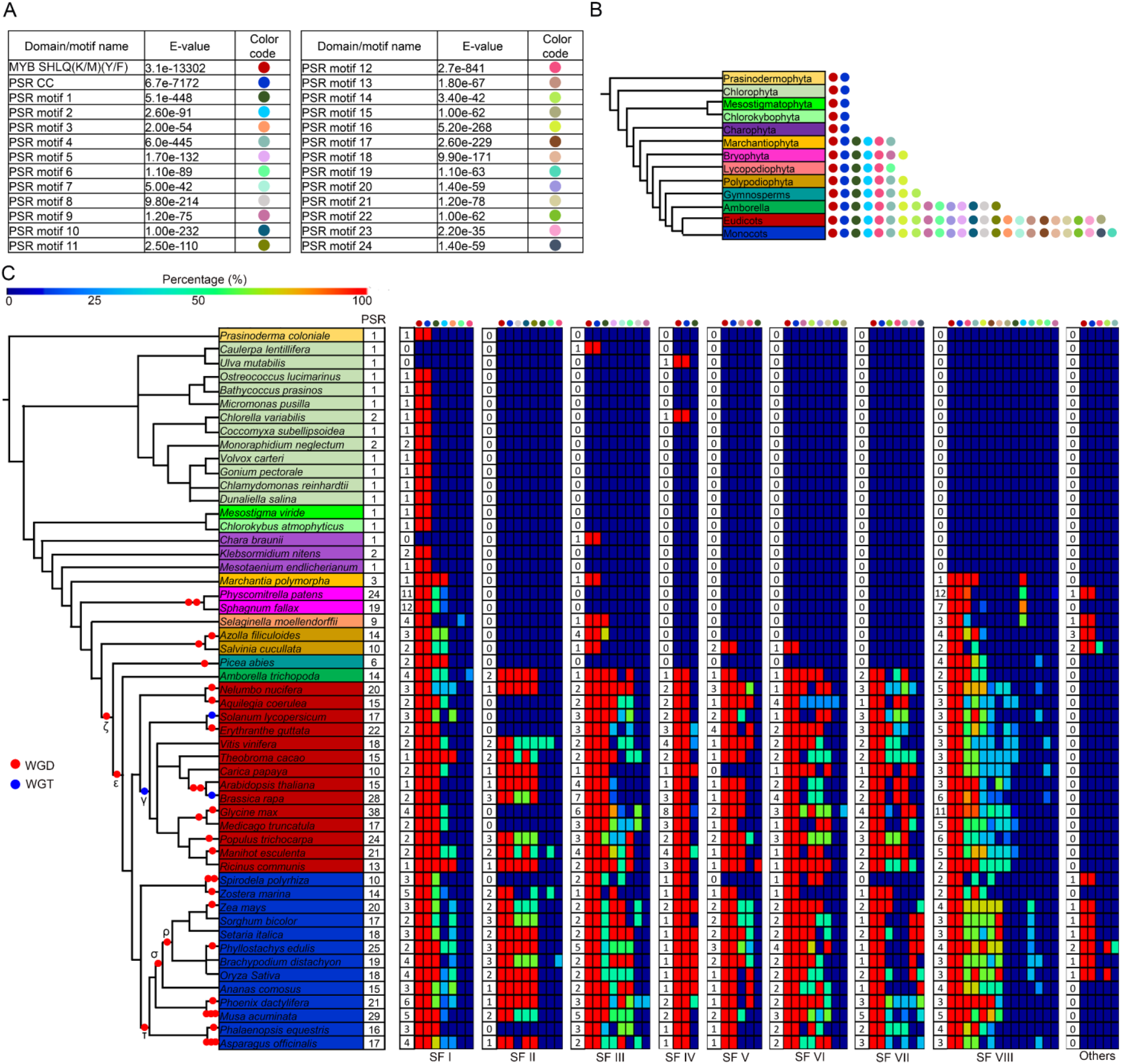
Origin and conservation of novel protein motifs in PSR transcription factor family in the plant lineage. (A) Novel motifs identified from PSR TF family in the plant lineage. (B) Taxonomical categories of Archaeplastida showing the presence or absence of specific novel motifs in PSR TF family. (C) Conservation of novel protein motifs in different subfamilies of PSR TF family. The detailed visualization of domain/motif arrangement in different PSR TF subfamilies is given Supplementary Figure S13.

### Analysis of the accessory motif divergence in the paralogous PSR and HRS TFs

In eukaryotes, divergence in the DBD and accessory motifs and expression domain were found to be the major determinants of functional specialization of paralogous TFs (Lehti-Shiu et al. 2015; Wang et al. 2016; Khor and Ettensohn 2017; Shen et al. 2018; Brodsky et al. 2020). Angiosperm genomes show high expansion in the PSR and HRS TF repertoire. We hypothesized that the divergence in motif composition and expression might have led to the specialization of these TFs. To test this hypothesis, we first identified the paralogous PSR and HRS gene pairs from angiosperm genomes. The sensitive detection based on homology in the genomic regions identified that a significant majority of PSR and HRS TFs from angiosperms are paralogs in nature (Supplementary Figure 17). Interestingly, motif comparison analysis identified that a large number of PSR and HRS paralogs show divergence in the motif composition (Figure 6A). In our analysis of 256 PSR paralogs from 21 angiosperm genomes, we observed motif divergence in approximately 69% of the paralogs. Similarly, analysis of 91 HRS paralogs from 16 angiosperms identified motif divergence in approximately 75% of paralogs. Therefore, we analyzed whether the motif divergence is correlated with sequence divergence. The analysis of the number of nonsynonymous substitutions per non-synonymous site (dN/dS) ratio identified that the PSR and HRS paralogs are under strong purifying selection (Supplementary Figure 18A). Nonetheless, at the global scale, we did not observe any clear correlation between motif divergence and sequence divergence (Supplementary Figure 18B). Collectively, our analysis suggests that motif divergence among paralogs is a major contributing factor in the divergence and functional specialization of PSR and HRS TF repertoire in angiosperms.

**Figure 6.**
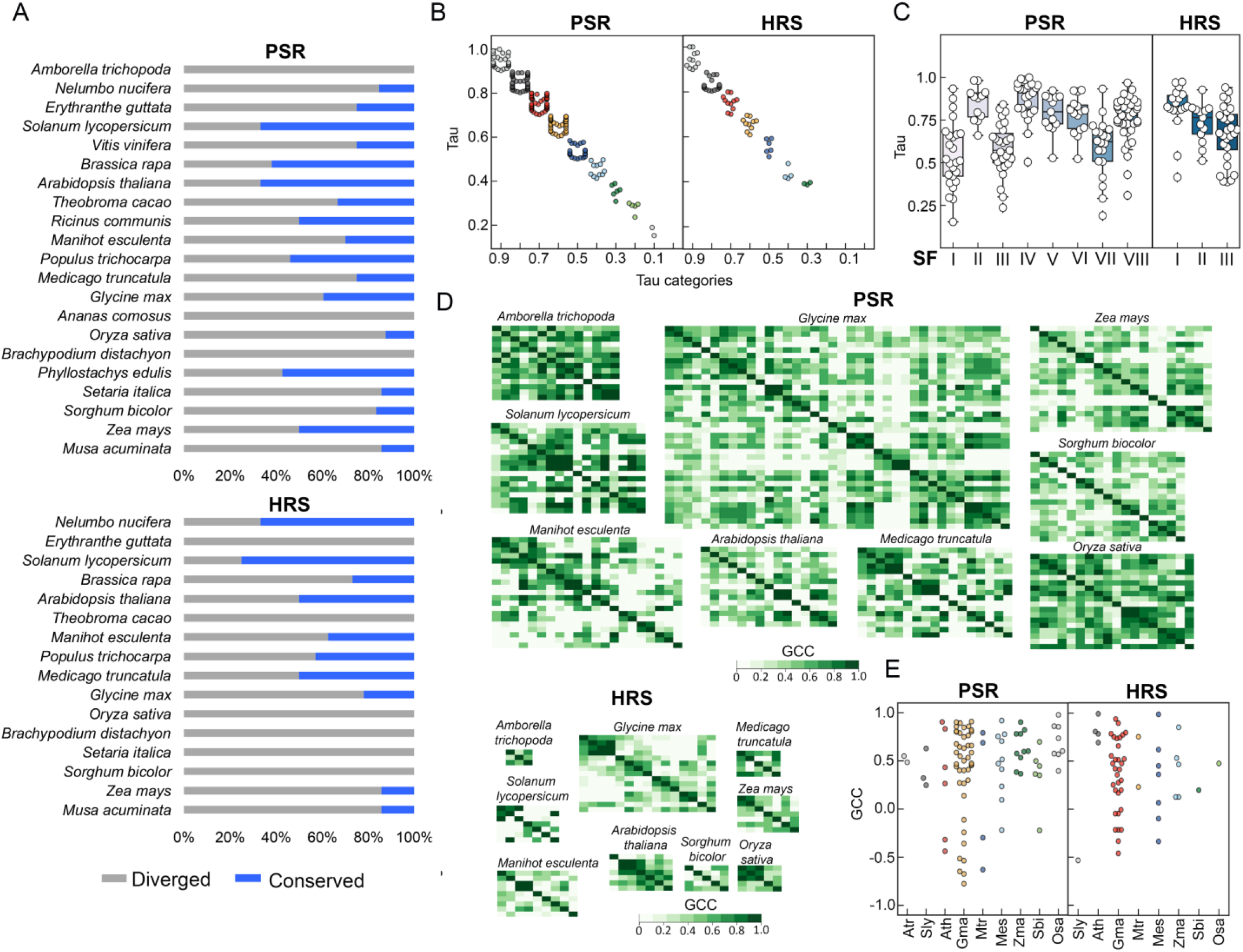
Motif evolution and expression patterns of PSR and HRS transcription factor families in angiosperms. (A) Motif conservation and divergence of paralogous PSR and HRS TFs across the angiosperm lineage. (B) Distribution of tissue-specificity score of PSR and HRS TFs. The tissue and developmental-stage specific expression data of basal angiosperm *Amborella trichopoda* and 5 eudicots (*Solanum lycopersicum*, *Arabidopsis thaliana*, *Glycine max*, *Medicago truncatula* and *Manihot esculenta*) and 3 monocots (*Zea mays*, *Sorghum bicolor* and *Oryza sativa*) were analyzed using Tau (τ) method. (C) Distribution of tissue-specificity score of different subfamilies of PSR and HRS TFs. (D) Correlation of expression pattern of PSR and HRS TFs analyzed using Gini Correlation Coefficient (GCC). The genes were arranged according to the evolutionary relationship (paralogs) and relative position in the phylogenetic reconstruction. (E) Correlation of expression pattern between paralogous PSR and HRS TFs.

### Analysis of gene expression divergence of PSR and HRS TFs in angiosperms

As expression divergence is an important determinant in the functional specialization of multigene families, we analyzed the conservation and divergence in tissue and developmental-specific expression of PSR and HRS gene families in different angiosperms. We analyzed the expression data from nine angiosperms which include the basal angiosperm *A. trichopoda* and five eudicots (*Solanum lycopersicum*, *A. thaliana*, *G. max*, *M. truncatula* and *Manihot esculenta*) and three monocots (*Zea mays*, *Sorghum bicolor* and *Oryza sativa*) (Supplementary Figure 19). Interestingly, we observed a great degree of divergence in the expression pattern of PSR and HRS TFs in different tissues and developmental stages. Therefore, we analyzed the tissue-specificity using Tau (τ), one of the best metrics to analyze tissue specificity of gene expression (Kryuchkova-Mostacci and Robinson-Rechavi, 2017). In our analysis of 176 PSR genes, approximately 65% (i.e., 114 genes) showed intermediate specificity (0.20 ≤ τ < 0.80) in expression (Figure 6B). In contrast, approximately 35% (i.e., 62 genes) PSR genes showed high specificity (τ ≥ 0.80) in expression. In 63 HRS genes, approximately 46% (i.e., 29 genes) genes showed high specificity while 54% (i.e., 34 genes) showed intermediatory specificity in expression. Further, we analyzed the relation between tissue-specificity with species and SFs. In our analysis, we did not observe major differences in the τ score among different species (Supplementary Figure 20). However, we found significant differences in the tissue-specificities of both PSR and HRS SFs (Figure 6C). For example, in PSR TFs, SFI and III showed high expression and low tissue-specificity (0.54 and 0.58 average τ respectively). The high expression levels and low-tissue specificity of these SFs suggest a predominant function of these members in Pi starvation responses. In line with this, Arabidopsis PHR1, the major PSR TF was found to be a member of SFIII with an intermediate specificity (τ: 0.65). Similarly, rice PHR1 and PHR2, which are crucial in Pi starvation responses were also found to be members of PSR SFIII with intermediate tissue-specificity in expression (OsaPHR1 τ: 0.53; OsaPHR2 τ: 0.47) (Zhou et al., 2008). Further, we tested the conservation and divergence in the expression domain at the gene family level using Gini Correlation Coefficient (GCC) (Ma and Wang, 2012). On the global level, we found a low degree of correlation indicating high divergence in tissue-specific expression domains of PSR and HRS TF families in angiosperms (Figure 6D). Nonetheless, we found small clusters with high or very low correlation in expression. In general, young duplicates tend to have more similar expression patterns while divergence in expression domains or suppression of the expression of one copy for maintaining dosage is also very common among paralogs. Therefore, we closely analyzed the expression pattern of paralogous genes from PSR and HRS TF families from nine angiosperms. We observed a nuanced pattern of correlation among paralogous genes from species with highly duplicated genomes such as *G. max*. Some paralogous genes showed positive correlation in expression while negative correlation was also evident among few paralogs (Figure 6E). Collectively, the in-depth global analysis of tissue-specific expression in angiosperms indicates that expression divergence also contributes to the functional specialization of PSR and HRS TF family in angiosperms.

The PSR and HRS TFs are majorly involved in Pi and N status dependent transcriptional coordination. Therefore, although angiosperm genomes possess large repertoire of these TF families, it is possible that they are involved in a spatiotemporal manner during different Pi and N nutrition regimes. In order to understand the global transcriptional regulation of these TF families in angiosperms, we studied their expression in rice under different Pi and N nutrition regimes in both shoots and roots. Rice genome contains 18 PSR and 5 HRS TF genes. Interestingly, our analysis revealed many interesting patterns of gene expression among related PSR and HRS genes. For example, among four PSR SF I genes in rice, expression of *LOC_Os08g25820* and *LOC_Os09g12770* were moderately increased in the late timepoints of Pi starvation in root (Figure 7A). Conversely, among the other two SFI genes, the expression of *LOC_Os05g41240* was repressed while no change in expression was observed in the case of *LOC_Os06g40710* in root. The same set of genes showed very different expression dynamics in shoot under Pi starvation. In the same time points, only the expression of *LOC_Os09g12770* is slightly induced in shoots. Another example is the paralogous gene pair *LOC_Os06g49040* and *LOC_Os02g04640* belonging to PSR SF II. The expression of *LOC_Os06g49040* is strongly induced in both root and shoot tissue at later stages of Pi starvation and Pi replenishment suppressed its expression. Conversely, expression of *LOC*_Os02g04640 is repressed in roots under Pi starvation and induced in response to Pi replenishment. Similar expression divergence was also observed in other PSR gene family members. Consequently, at a global scale we found very less correlation in the expression pattern under different Pi nutrition regimes (Figure 7B). High diversity in the expression dynamics was also observed in the case of rice HRS TF family under different Pi nutrition regimes (Figure 7C, D). For example, *OsaNIGT1* (*LOC_Os02g22020*) showed strong and time-dependent induction in expression under Pi starvation in both shoots and roots. The expression of *OsaNIGT1* is rapidly repressed by Pi re-addition. In comparison, other members showed more varied expression pattern under different Pi nutrition regimes. Interestingly, the heterogeneity in the expression pattern was also evident under N starvation conditions. Among PSR genes, some genes (such as *LOC_Os06g49040* and *LOC_Os02g04640*) showed highly similar expression patterns under N starvation (Figure 7E, F). It should be noted that these paralogs showed highly contrasting expression patterns under different Pi nutrition regimes (Figure 7A). Thus, closely related genes such as paralogs show condition-specific positive or negative correlation in the expression. Among HRS genes, in line with the previous reports indicating strong induction of these genes during N starvation (Liu et al., 2009; Sawaki et al., 2013; Medici et al., 2015; Maeda et al., 2018)transcript levels of most of rice HRS family members were found to be repressed during N starvation especially in roots (Figure 7G, H). However, we also found that transcript level of certain members induced during N starvation. For example, expression of *LOC_Os01g08160* was induced in root in most of the time points while the expression of *LOC_Os12g39640* was found to be induced in shoots. Collectively, the global analysis of tissue and developmental stage-specific transcriptional dynamics of PSR and HRS TF family in angiosperms indicate that the divergence in the expression domain is a major contributor to functional divergence. Further, expression analysis in rice under different Pi and N nutrition regimes indicates that specific members have probably acquired specialized roles in a spatiotemporal manner to provide a coordinated transcriptional response according to the nutrient availability.

**Figure 7.**
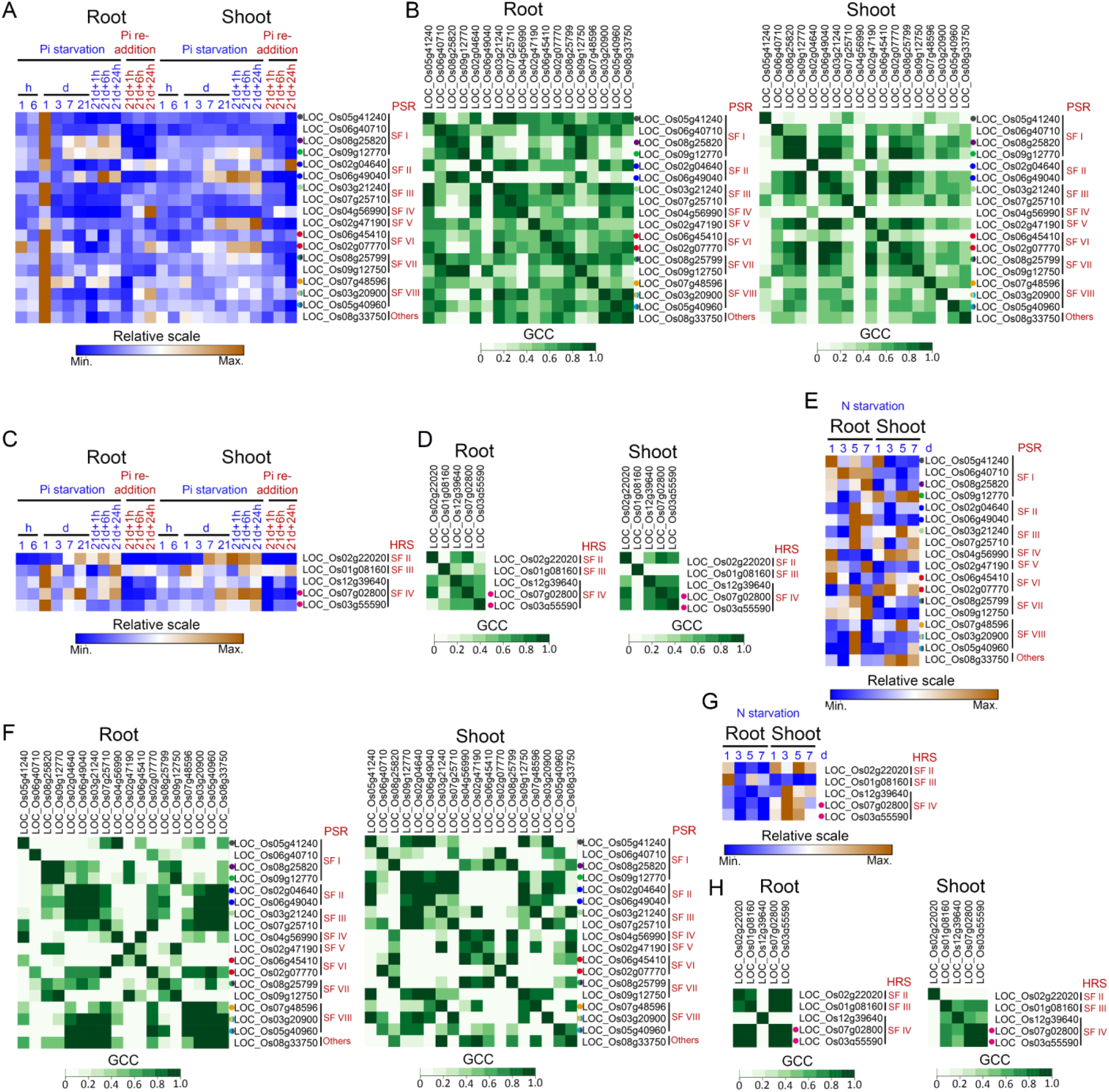
Expression analysis of rice PSR and HRS transcription factors under diverse phosphate and nitrogen regimes. (A) Root and shoot-specific expression pattern of rice PSR TFs in short-term (1 and 6 hours) and long-term (1, 3, 7, and 21 days) Pi starvation and replenishment after 21 days (1, 6 and 24 hours). (B) Correlation of expression pattern of PSR TFs in root and shoot under different Pi nutrition regimes analyzed using Gini Correlation Coefficient (GCC). (C) and (D) Root and shoot-specific expression pattern of rice HRS TFs and correlation of expression pattern under different Pi nutrition regimes. (E) and (F) Root and shoot-specific expression pattern of rice PSR TFs in different stages (1, 3, 5, 7 days) of N starvation and correlation of expression pattern. (G) and (H) Root and shoot-specific expression pattern of rice HRS TFs under different stages N starvation and correlation of expression pattern. Paralogs are indicated with dots of different colors.

## Discussion

Recent studies have identified an important role of the PSR-HRS transcriptional regulatory system in coordinating gene expression according to the Pi and N availability in plants. In this study, through sensitive evolutionary analysis, we identified interesting insights on the stepwise origin, expansion and diversification of this regulatory system in the plant lineage (Figure 8).

**Figure 8.**
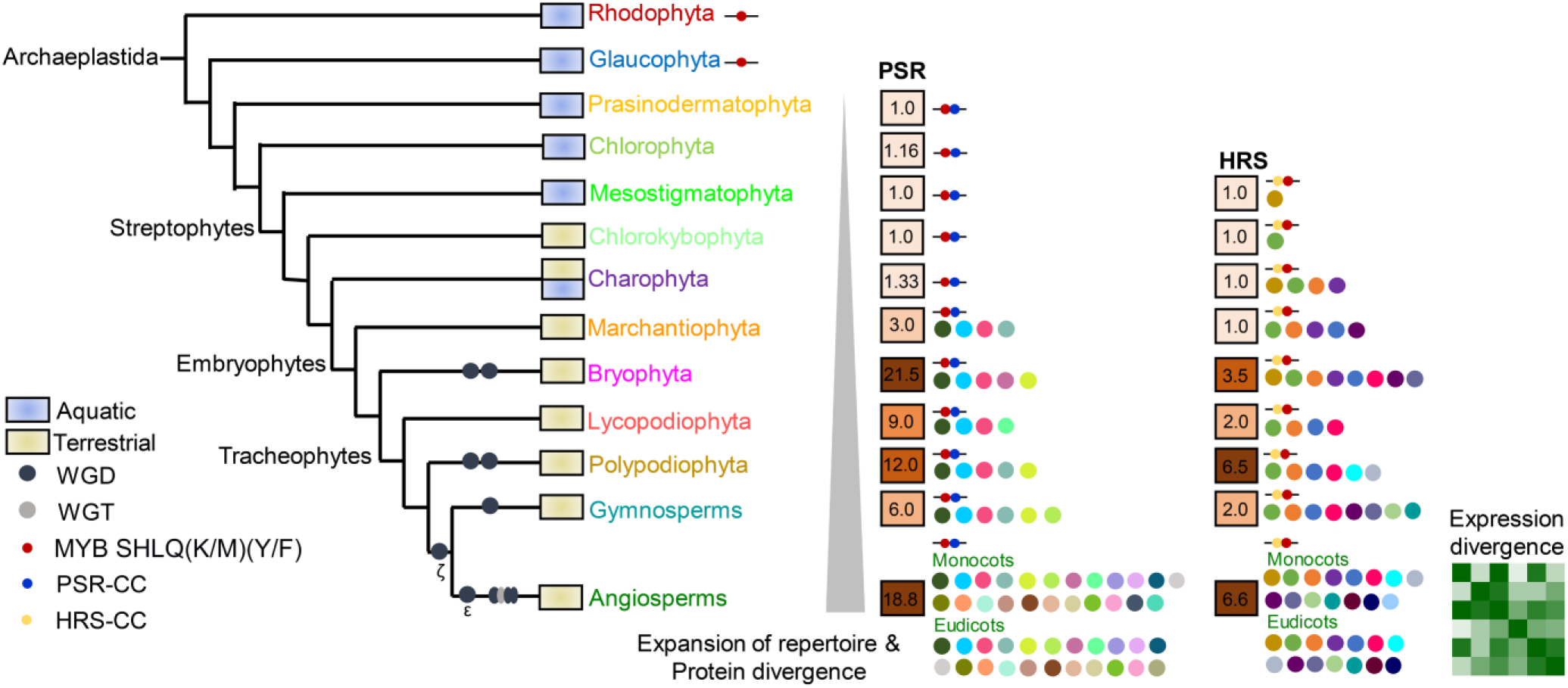
A model of the origin, expansion and divergence of PSR and HRS repertoire in the plant lineage. The PSR TF is more ancient and originated in aquatic algae. The origin of HRS TFs possibly coincides with the terrestrialization of plants. The origin of accessory motives is also indicated. In land plants, especially in angiosperms, the PSR-HRS repertoire is enhanced due to the ancient and lineage-specific whole-genome duplication and triplication events and the origin of novel motifs may have contributed to the functional specialization. Analysis of the expression dynamics in angiosperms suggests expression divergence also contributes to the functional specialization of these TFs in angiosperms.

In our analysis, the PSR TFs with the characteristic MYB SHLQ(K/M)(Y/F) DBD and CC domain were found in Prasinodermophyta and Chlorophyta genomes. In contrast, the typical HRS TFs were detected only in the Streptophyta genomes. Thus, the origin of PSR-type TFs precedes the origin of HRS TFs. Further, the origin of HRS TFs coincides with the terrestrialization of the plant lineage (de Vries and Archibald, 2018). HRS TFs are involved in finetuning the transcriptional responses through different feedback signaling loops (Sawaki et al., 2013; Medici et al., 2015; Safi et al., 2017; Maeda et al., 2018). Unlike the aquatic environment, nutrients are present in patches and in different gradients in the soil (Giehl and von Wirén, 2014). Therefore, HRS TFs must be an evolutionary innovation especially important in the land plants to optimize the transcriptional responses according to diverse nutrient availability conditions on the land.

Although genomes from Rhodophyta and Glaucophyta also possess MYB SHLQ(K/M)(Y/F) TFs, they lack the typical CC domain present in PSR and HRS TFs. It could be possible that some of these proteins represent the ancient PSR TF system in Archaeplastida. In line with this, a *Phaeodactylum tricornutum* (Stramenopile) MYB SHAQKYF TF without the CC domain was found to be involved in the transcriptional regulation of Pi starvation responses (Sharma et al. 2020). Therefore, further functional studies of the MYB SHLQ(K/M)(Y/F) TFs from Rhodophyta and Glaucophyta genomes will be needed to identify whether they represent the ancient PSR TF system of the plant lineage. The CC domain involved in the InsP-SPX-mediated regulation of PSR TF in Arabidopsis might be an evolutionary innovation that happened later to finetune transcriptional responses according to the cellular Pi availability.

In our analysis, we found that the number of these TFs especially PSR type is highly expanded in most of the land plants. On a relative scale, the number of HRS TFs is moderately expanded. This difference was consistent in all genomes we analyzed. The difference in the expansion of gene families can be due to several factors such as their dosage sensitivity, essential functionalities (e.g., mutants are lethal) and contribution to adaptive benefits (e.g., enhanced abiotic stress resilience or involvement in evolutionary arms-race with microbes) to a particular environment (Wang et al., 2018). The PSR and HRS TFs may have different functions in such processes and that might be contributing to the differences in gene family expansion. PSR TFs are involved in promoting the association with beneficial bacteria and mycorrhiza (Castrillo et al. 2017; Shi et al. 2021).The HRS TFs have an important role in finetuning the transcriptional responses according to Pi and N availability. However, further studies are required to identify their specific adaptive benefits shaping the gene family size in the plants. Nonetheless, the WGD/WGT events were found to be a major factor determining the size of these gene families. For example, the *M. polymorpha* genome contains three PSR and a single HRS TF gene while the *P. patens* genome contains 24 PSR and three HRS genes. The *M. polymorpha* genome shows low genetic redundancy especially in regulatory factors such as TFs due to the absence of ancient or lineage-specific WGD events (Bowman et al., 2017). In contrast, there is evidence for at least two WGD events specific to the moss lineage (Lang et al. 2018). The contribution of WGD/WGT in the expansion of these TFs is also evident in angiosperms and the gene family size is higher in genomes with recent WGD/WGT events.

Although there is strong conservation in MYB SHLQ(K/M)(Y/F) and CC domain in these TFs, we found that they were recovered into distinct clades in the phylogenetic reconstruction. This prompted us to investigate the highly disordered variable regions in these TFs. Our sensitive motif analysis identified accessory motifs conserved in these variable regions. Although there is very low sequence conservation, some of these identified motifs were found to be highly conserved and specific motifs or their combination largely define subfamilies. For example, the combination of PSR motif 10, 8 and 11 in the N-terminus is specific to PSR subfamily II. The combination of HRS motif 12, 6 and 2 in between the N-terminal CC domain and MYB SHLQ(K/M)(Y/F) largely defines the HRS subfamily II. Interestingly, the PSR and HRS TFs from algae largely possess simple domain composition with no or few accessory motifs. In contrast, land plants show very high diversity in the accessory motif composition. This is especially prominent in angiosperms where we identified a large number of accessory motifs specific to subfamilies. Thus, along with the expansion in the gene family size, these TFs underwent significant evolutionary divergence before the angiosperm radiation. Studies in eukaryotic TFs identified a huge impact of IDRs and accessory motifs in the TF function (Nguyen Ba et al. 2014; Cheatle Jarvela and Hinman 2015; Brodsky et al. 2020). Earlier the DBD was thought to be the major determinant in the recognition of target promoter regions. However, recent studies led to a paradigm shift in which the IDRs were found to be a major contributing factor defining the promoter recognition. For example, in Msn2 and Yap1 TFs from yeast, the partially redundant regions in the IDRs independent of the DBD recognize target DNA through a multitude of weak interactions with chromatin or histones (Brodsky et al. 2020). Thus, theAligned MSAs were subjecte daccessory motifs enriched in the IDRs in specific subfamilies of PSR and HRS TFs may potentially define the promoter specificities of individual subfamilies. IDRs and the motifs in IDRs may also serve as a site of protein-protein interaction (Jamsheer K et al. 2018). Therefore, it is also possible that these motifs in the IDRs may have a role in determining the recruitment of factors involved in the regulation of protein function and gene expression. For example, in plants, the divergence in the PHYB binding (APB) motif and motifs of unknown functions (MUFs) in Phytochrome Interacting Factor (PIF) TFs determine the functional specificity in plants (Possart et al., 2017). Another potential role of these accessory motifs in IDRs may be in the phase separation as the weak interaction of IDRs in TFs is a major factor determining the condensation of transcription apparatus to specific chromatin regions (Boija et al. 2018). These are the potential roles and more targeted studies will be required to determine the function of these accessory motifs in PSR and HRS TFs. Intriguingly, a large fraction of paralogs (69% in PSR and 75% in HRS on a global scale) showed divergence in accessory motif composition. Thus, even at the level of paralogs, the divergence in accessory motifs seems to be a major factor in the potential functional specialization of these TFs (Figure 8).

As divergence in the expression domain is a critical factor determining the functional specificities in multigene families, we also investigated tissue and developmental stage-specific expression pattern of PSR and HRS TFs from nine angiosperms. We found that a significant portion of these genes (65% in PSR and 54% in HRS) show intermediate specificity in the expression domain. The rest of the genes (35% in PSR and 46% in HRS) were found to be highly specific in the expression domain. These results indicate that specificities in the expression domain are also an important factor in the functional specialization of PSR and HRS TFs. Interestingly, we also found that subfamilies show differences in tissue-specificity. Some of the prominent members characterized as major regulators of Pi starvation in model plants such as Arabidopsis (AthPHR1) and rice (OsaPHR1 and OsaPHR2) were found to be expressed in most of the tissues and belong to subfamilies with intermediate specificity in the expression domain. We also analyzed the global correlation in the expression domain which further highlighted the differences in the expression domain. As these genes are majorly involved in the transcriptional regulation under different Pi and N availability conditions, we investigated the spatiotemporal expression pattern of rice PSR and HRS genes under different Pi and N conditions. Our analysis revealed spatiotemporal divergence in the expression domain even in closely related genes. Further, comparison of the expression pattern of closely related genes in Pi and N availability identified condition-specific positive or negative correlation in the expression. Taken together, these results suggest that although angiosperm genomes have a large repertoire of PSR and HRS TFs, different members might have specific functions under different environmental conditions.

Collectively, our comprehensive analysis of PSR and HRS TFs in the plant lineage identifies a stepwise evolution of a transcriptional activator-repressor coordinating gene expression according to Pi and N nutrient availability. We also found that the expansion of these genes in land plants is also correlated with the origin of novel accessory motifs and a high degree of expression divergence (Figure 8). Thus, along with redundant functions, individual members of these multigene families might also have unique functions. More targeted studies in this direction will be needed to reveal the functional complexities of these TFs in plants. Often the genetic analysis in controlled culture conditions overlooks the role of individual genes in multigene families in the fitness of the plants. However, in the natural conditions where plants face a large number of variabilities in environmental and biotic factors together, individual genes in multigene families seem to have a significant effect on the fitness of the plants. For example, genetic analysis of the three gibberellin receptor genes in tomato under a controlled environment suggests a highly redundant role of individual receptor genes. In contrast, under the field conditions, this genetic redundancy was minimal and the role of individual genes was found to be more prominent (Illouz-Eliaz et al., 2019). The current genome editing techniques can help to answer these questions as it enables large-scale, multiplexed genetic screening of multigene families. Thus, the evolutionary trajectory of the PSR and HRS TF system outlined in our study will be helpful in future functional studies in this important transcriptional regulatory system of plants.

## Methods

### Identification of PHR and HRS TF family members

The protein sequences of G2-Like TFs were identified from the selected species from PlantTFDB v5.0 (Jin et al., 2017). Further, using *A. thaliana* PHR/PHL and HRS/HHO sequences as queries, BLASTP (E-value < 1E–5) search was performed in Phytozome v11.0 (Goodstein et al., 2012)and PhycoCosm (Grigoriev et al., 2021)to identify the PHR and HRS sequences from different plant species. Further, a profile-HMM based search was performed using HMMER web server v2.41.1 (Potter et al., 2018) in the respective reference proteomes (Sequence E-value: 0.01, Hit E-value: 0.03). The dataset obtained was manually filtered and repeats were removed. The CDD v3.19 (E value: < 0.01, database CDD --5570 PSSMs) (Lu et al., 2020) and MEME v5.3.3 (Bailey et al., 2015)analysis was performed to identify the sequences with conserved MYB SHLQ(K/M)(Y/F) and CC domains characteristic of PHR and HRS TFs. The final curated dataset of PSR and HRS TFs is given as Supplementary Dataset 1.

### Sequence similarity analysis

The domain and interdomain regions were identified using CDD v3.19 (E value: < 0.01, database CDD --5570 PSSMs). The multiple sequence alignments (MSAs) of N-terminal, C-terminal, linker region and domains of HRS and PSR TFs were performed using MUSCLE v3.8.31 (Edgar, 2004). Aligned MSAs were subjected to CIAlign(Tumescheit et al., 2021) to obtain sequence similarity matrices and Skylign (https://skylign.org/) to create HMM logo. These sequence similarity matrices were used to generate heat map by using heamap.2 function of the gplots package in R.

### Phylogenetic reconstruction

The full-length protein sequences were aligned and phylogenetic trees were reconstructed using Maximum likelihood (ML) method using FastTreeMP v2.1.10 (JTT+CAT substitution model, 1000 bootstrap replicates) on CIPRES (https://www.phylo.org/). The phylogenetic trees were visualized and annotated using FigTree v1.4.4.

### Homology modeling and structural similarity analysis

The MYB SHLQ(K/M)(Y/F) domain boundary was identified using CDD v3.19 (E value: < 0.01, database CDD --5570 PSSMs) and homology models were created using I-TASSER server (Yang and Zhang, 2015). The model with the highest C-score was selected for further analysis. The energy minimization and refinement of the structure was performed using ModRefiner (D and Y, 2011)and quality was analyzed in SAVES v6.0 server using PROCHECK and Verify3D. The PDB files of MYB SHLQ(K/M)(Y/F) domain of AthPHR1, AthLUX and AthARR10 from RCSB PDB and the structural similarity analysis was performed using TM-score (Zhang and Skolnick, 2004).

### Disorder prediction

The disorder of PHR and HRS proteins were predicted using the meta-predictor PONDR-FIT (Xue et al., 2010). The average of disorder and standard deviation of each residue was obtained and used for further analysis. The boundaries of each domain/region (N-terminus, MYB SHLQ(K/M)(Y/F), Linker region, CC domain, C-terminus) were identified by CDD v3.19 (E value: < 0.01, database CDD --5570 PSSMs) and the average disorder score was calculated. The IDRs were categorized in two groups. As used in previous studies (Jamsheer K et al., 2018), stretches ranging from 10 to 29 protein residues with disorder score of ≥0.5 for each residue were categorized as short IDRs (SIDR). Amino acid stretches of ≥30 residues long with disorder score of ≥0.5 for each residue were categorized as long IDRs (LIDR). An IDR was considered as junction-IDR if at least five disordered (disorder score of ≥0.5 for each residue) residues are in the other domain/region. As used in previous studies (Jamsheer K et al., 2018), a tolerance limit of three tandem residues with less than 0.5 disorder score was set for this analysis.

### Novel motif identification and motif divergence analysis in paralogs

The full-length protein sequences were used for identifying novel motifs in PHR and HRS using MEME v5.3.3 (Bailey et al., 2015). The information regarding duplicated genes based on i-ADHoRe program, which detects homologous genomic regions were retrieved from Dicots PLAZA v4.0 and Monocots PLAZA v4.5 and used for the analysis of motif divergence among paralogs in angiosperms (Van Bel et al., 2018). Loss, gain or rearrangement of motifs were considered as divergence among paralogs. The data on conserved and diverged paralogs are in percentage in comparison with the total number of paralogs.

### Gene expression Gini Correlation coefficient analyses

The normalized RNA-seq based tissue and developmental stage-specific expression data of PHR/PHL and HRS/HHO genes from *Amborella trichocarpa* (Flores-Tornero et al., 2020), *Solanum lycopersicum*(Sato et al., 2012), *Arabidopsis thaliana* (Klepikova et al., 2016), *Glycine max* (Machado et al., 2020), *Medicago truncatula* (Dai et al., 2021), *Manihot esculenta* (Wilson et al., 2017), *Zea mays* (Stelpflug et al., 2016), *Sorghum bicolor* (McCormick et al., 2018), and *Oryza sativa* (Ouyang et al., 2007) were retrieved from previous studies. The tissue specificity of genes was calculated using the τ method (Yanai et al., 2005).

In order to analyze the expression pattern of *O. sativa* PSR and HRS genes during nitrogen starvation, RNA-seq reads with accession numbers SRR5713884, SRR5713902, SRR5713901, SRR5713900, SRR5713899, SRR5713898, SRR5713894, SRR5713907, SRR5713906, SRR5713905, SRR5713904 and SRR5713903 were downloaded from Gene Expression Omnibus (Shin et al., 2018). These RNA-seq reads were aligned to the rice reference genome (IRGSP-1.0 genome) (Kawahara et al., 2013)using STAR v2.7.7a (Dobin et al., 2013)with default parameters by using rice gene annotation (http://rapdb.dna.affrc.go.jp/). Normalized gene expression was calculated in terms of FPKM using StringTie v2.1.4 (Pertea et al., 2015). The publicly available normalized RNA-seq expression data of *O. sativa* Pi starvation and replenishment treatment was also used in our analysis (Secco et al., 2013). The heat map of the gene expression data was generated using Morpheus (https://software.broadinstitute.org/morpheus/). Gini correlation coefficient (GCC) between expressions was calculated using the cor.pair function of the ‘rsgcc’ package in R (Ma and Wang, 2012).

### Selection analysis

The synonymous (dS) and nonsynonymous (dN) nucleotide substitutions and dN/dS was calculated using PAL2NAL v14 (M et al., 2006) and KaKs Calculator (Zhang et al., 2006)using averaging (MA) method. The CDS and corresponding protein sequences were downloaded from PlantTFDB v5.0 (Jin et al., 2017), Phytozome v11.0 (Goodstein et al., 2012), PLAZA Dicots v4.0 and PLAZA Monocots v4.5 (Van Bel et al., 2018). Pair-wise protein sequence alignment of all paralogs was performed using MUSCLE v3.8.31 (Edgar, 2004). The aligned protein sequences were used to direct the conversion of their corresponding cDNAs into codon alignments.

## Acknowledgements and funding information

This work supported by the Department of Science and Technology, Ministry of Science and Technology (DST) (INSPIRE Faculty Programme, grant no. IFA18-LSPA110 to MJK). SJ is partially supported by the Ministry of Education, Youth and Sports of the Czech Republic, grant no. CZ.02.1.01/0.0/0.0/16_019/0000738 with support from the European Regional Development Fund - Project ‘Centre for Experimental Plant Biology’.

## Author contributions

MJK conceived the study and designed the analysis. MJK, RKG, SJ and MK performed the analysis. MJK and RKG wrote the first draft and SJ and MK reviewed the manuscript.

