## Supplementary Figures for "Stepwise origin and evolution of a transcriptional activator and repressor system integrating nutrient signaling in plants"

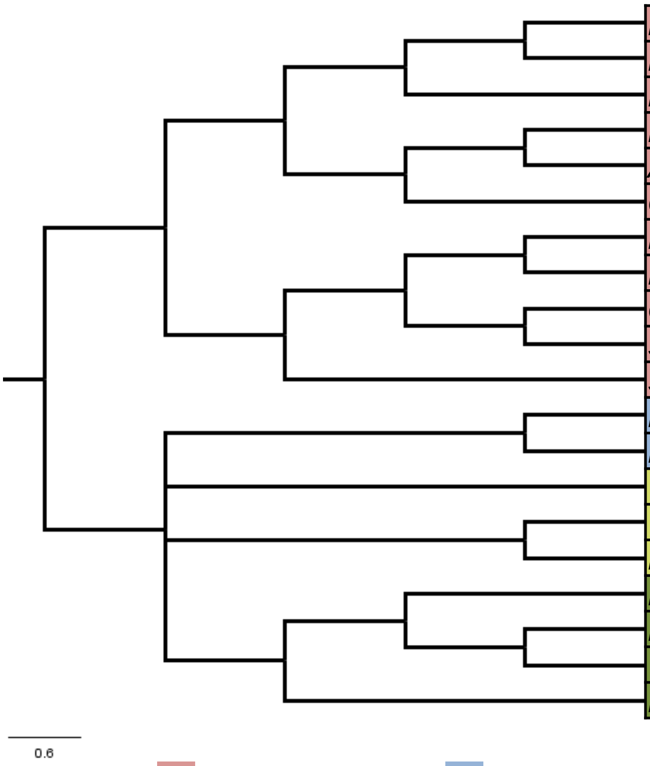


| Species | PSR/PHR/PHL | HRS/HHO |
|---|---|---|
| *Mus musculus* | 0 | 0 |
| *Homo sapiens* | 0 | 0 |
| *Danio rerio* | 0 | 0 |
| *Drosophila melanogaster* | 0 | 0 |
| *Aedes aegypti* | 0 | 0 |
| *Caenorhabditis elegans* | 0 | 0 |
| *Penicillium digitatum* | 0 | 0 |
| *Neurospora crassa* | 0 | 0 |
| *Candida albicans* | 0 | 0 |
| *Saccharomyces cerevisiae* | 0 | 0 |
| *Schizosaccharomyces pombe* | 0 | 0 |
| *Dictyostelium discoideum* | 0 | 0 |
| *Entamoeba histolytica* | 0 | 0 |
| *Trichomonas vaginalis* | 0 | 0 |
| *Trypanosoma cruzi* | 0 | 0 |
| *Leishmania braziliensis* | 0 | 0 |
| *Paramecium tetraurelia* | 0 | 0 |
| *Plasmodium falciparum* | 0 | 0 |
| *Toxoplasma gondii* | 0 | 0 |
| *Ectocarpus siliculosus* | 0 | 0 |

**Opisthokonta** **Amoebozoa** **Excavata** **SAR**

**Figure S1. Genome-wide identification of PSR and HRS family transcription factors from eukaryotic supergroups.**
A combination of BLAST and HMMER-based searches was used to identify PSR and HRS TFs from selected species from different eukaryotic supergroups.

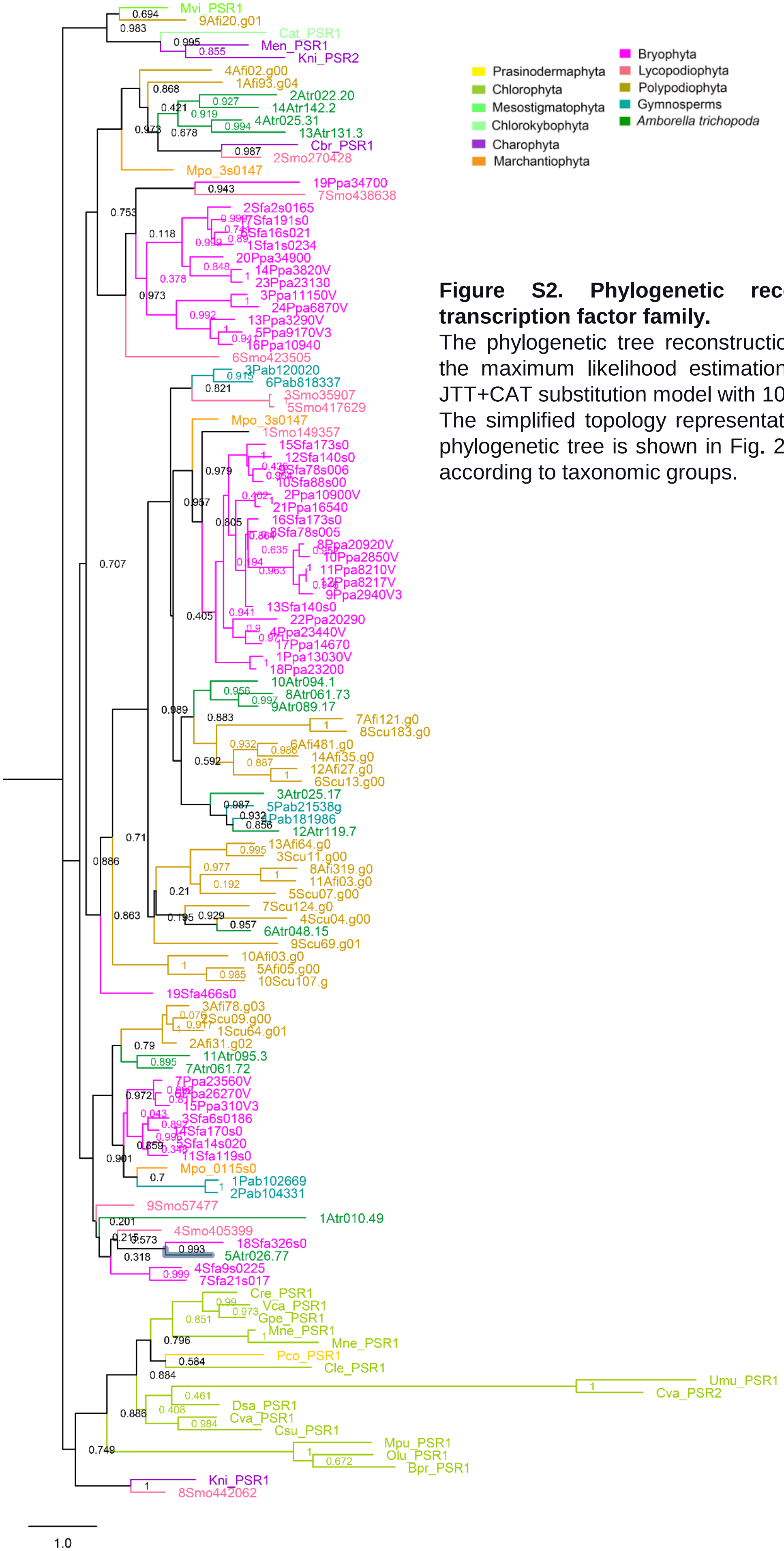


**Figure S2. Phylogenetic reconstruction of PSR transcription factor family.**

The phylogenetic tree reconstruction was performed using the maximum likelihood estimation method based on the JTT+CAT substitution model with 1000 bootstrap replicates. The simplified topology representation of the reconstructed phylogenetic tree is shown in Fig. 2b. Branches are colored according to taxonomic groups.

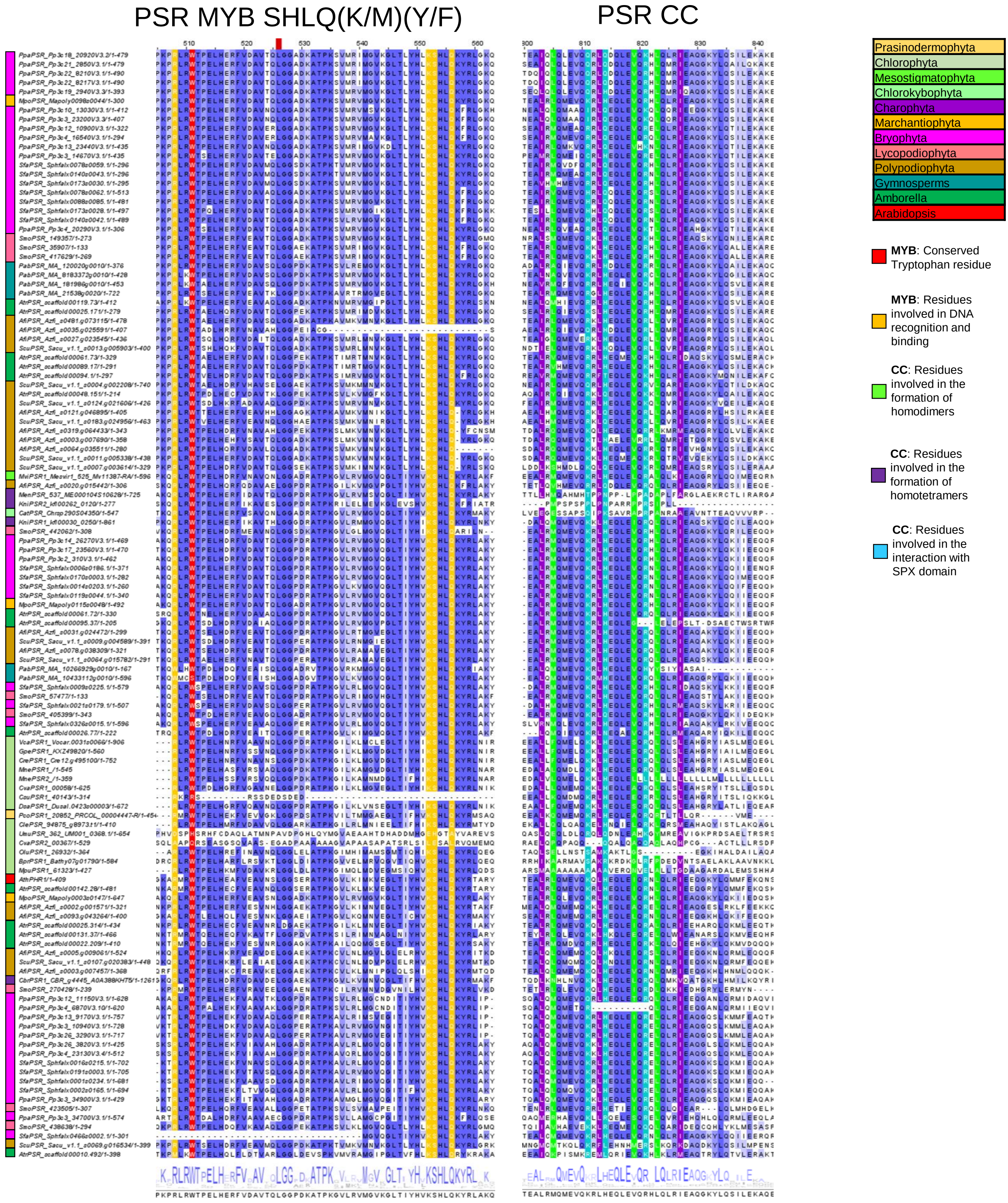


**Figure S3. Conservation of critical residues in MYB SHLQ(K/M)(Y/F) and CC domain of PSR TFs.** The full-length protein sequences were used for multiple sequence alignment and residue conservation analysis. The *Arabidopsis thaliana* PHR1 was used as a reference to mark the residues important for the function. The color key highlights different taxonomical groups of Archaeplastida.

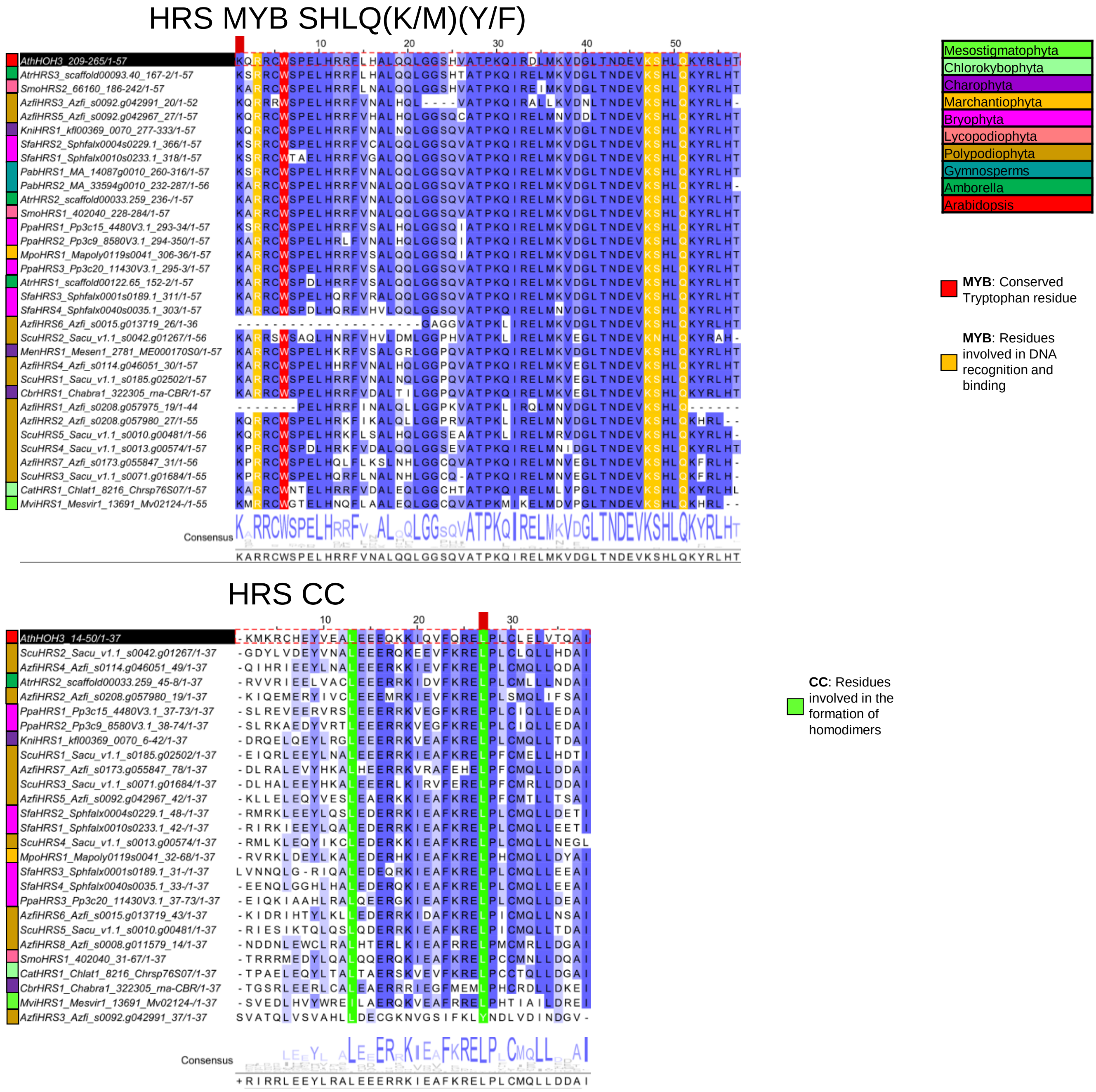


**Figure S4. Conservation of critical residues in MYB SHLQ(K/M)(Y/F) and CC domain of HRS TFs.** The full-length protein sequences were used for multiple sequence alignment and residue conservation analysis. The *Arabidopsis thaliana* HHO3 was used as a reference to mark the residues important for the function. The color key highlights different taxonomical groups of Archaeplastida.

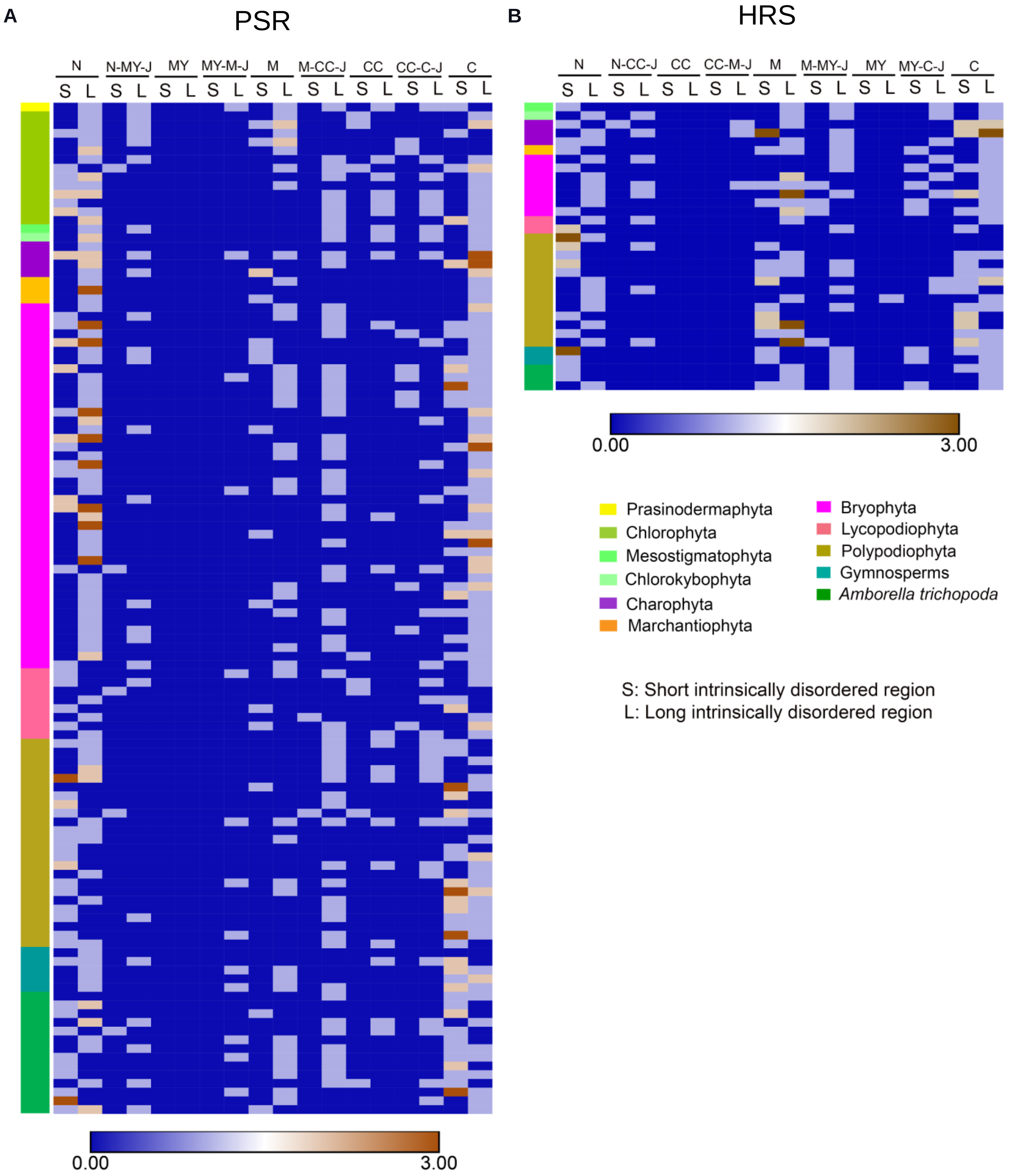


**Figure S5. Distribution of IDRs in different regions of PSR and HRS TFs.**
(A) and (B) The heatmaps represent the distribution of short and long IDRs in different regions of PSR and HRS TFs respectively. The IDRs were predicted using stringent parameters. The color key highlights different taxonomical groups of Archaeplastida. The details of the abbreviations used in the heatmap are as follows: N, N-terminus; MY, MYB SHLQ(K/M)(Y/F) domain; J, Junction region; M, Middle region; CC, Coiled coil domain; C, C-terminus.

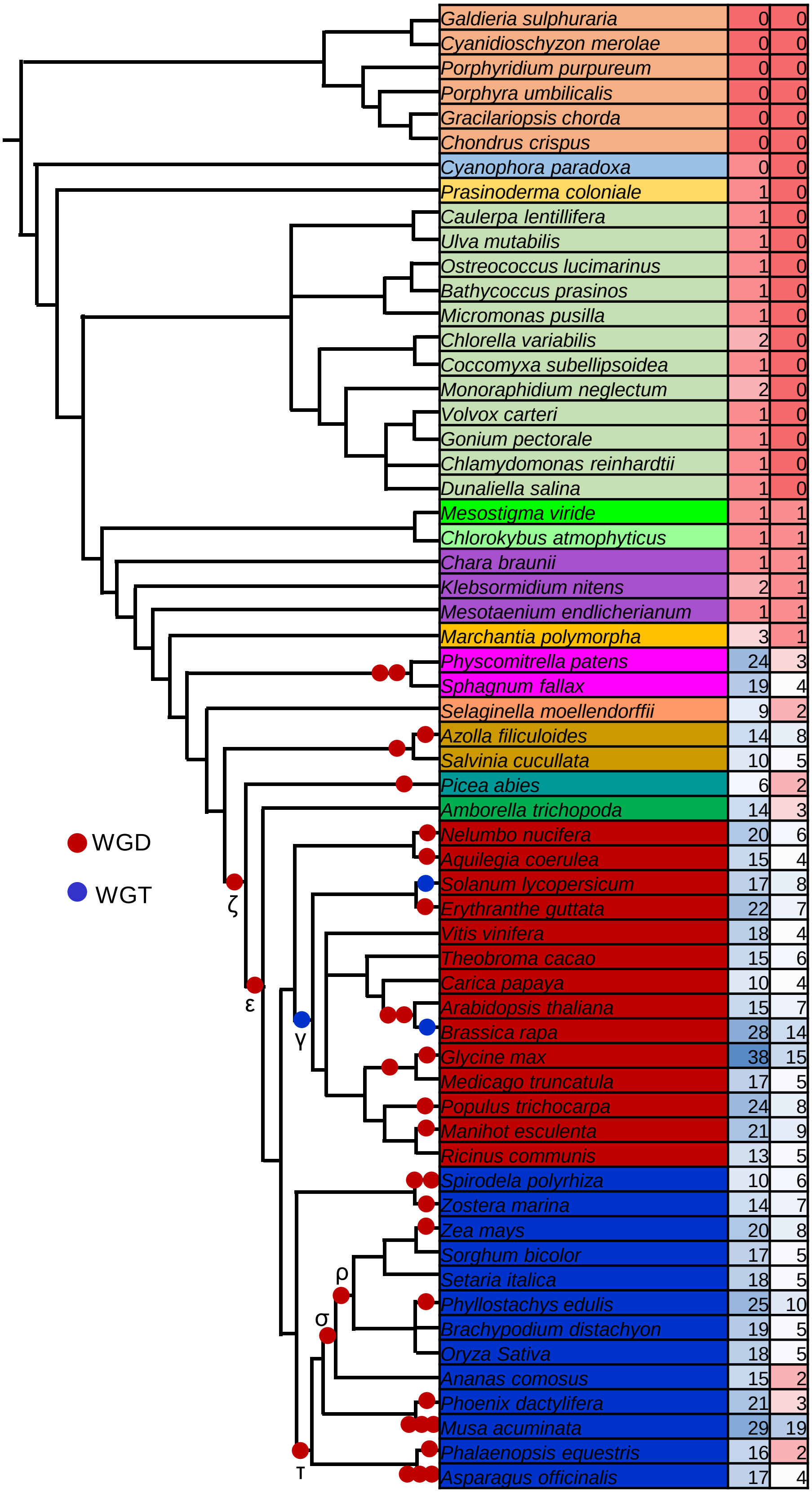


**Figure S6. Genome-wide identification of PSR and HRS family transcription factors from Archaeplastida.** A combination of BLAST and HMMER search was used to identify PSR and HRS TFs from selected species from Archaeplastida. WGD: Whole Genome Duplication; WGT: Whole Genome Triplication.

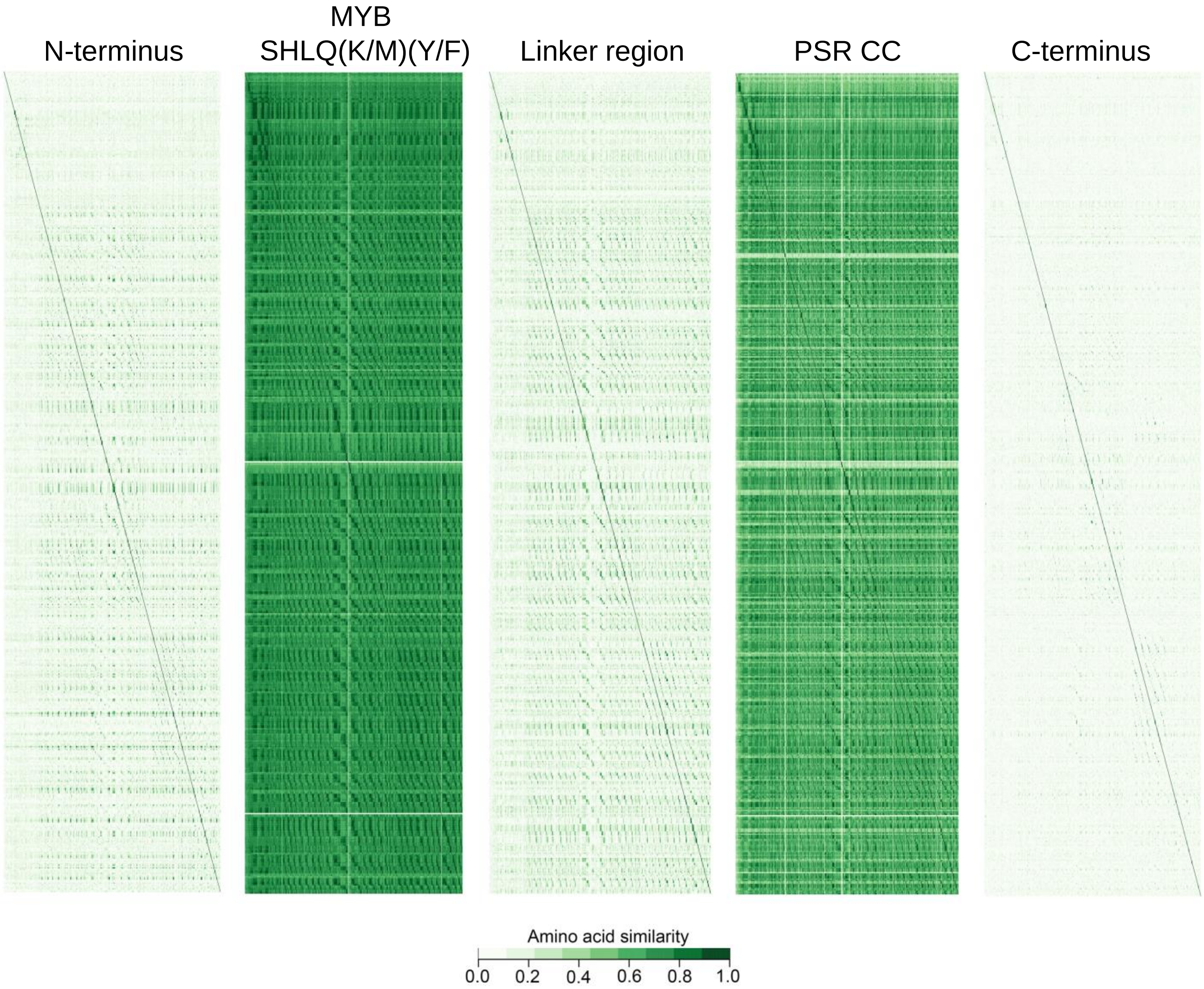


**Figure S7. Sequence similarity heatmap of different regions of PSR TFs across the plant lineage.** Different regions were identified through CDD v3.19-based domain analysis. The sequences were split accordingly and arranged as per the taxonomical position for the generation of sequence similarity heatmap.

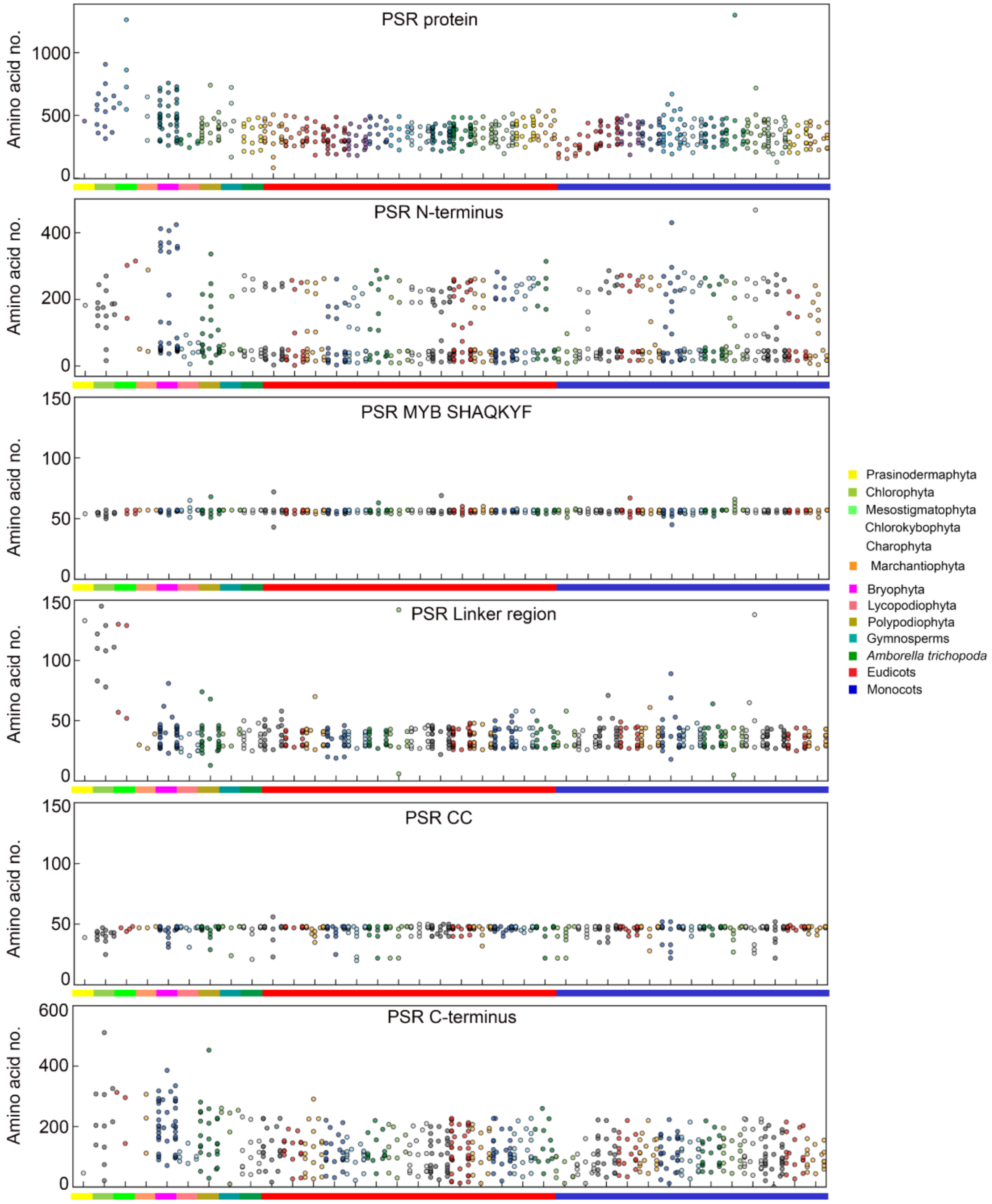


**Figure S8. Size distribution of PSR TFs and different regions across the plant lineage.**
Different regions were identified through CDD v3.19-based domain analysis. The color key highlights different taxonomical groups of Archaeplastida.

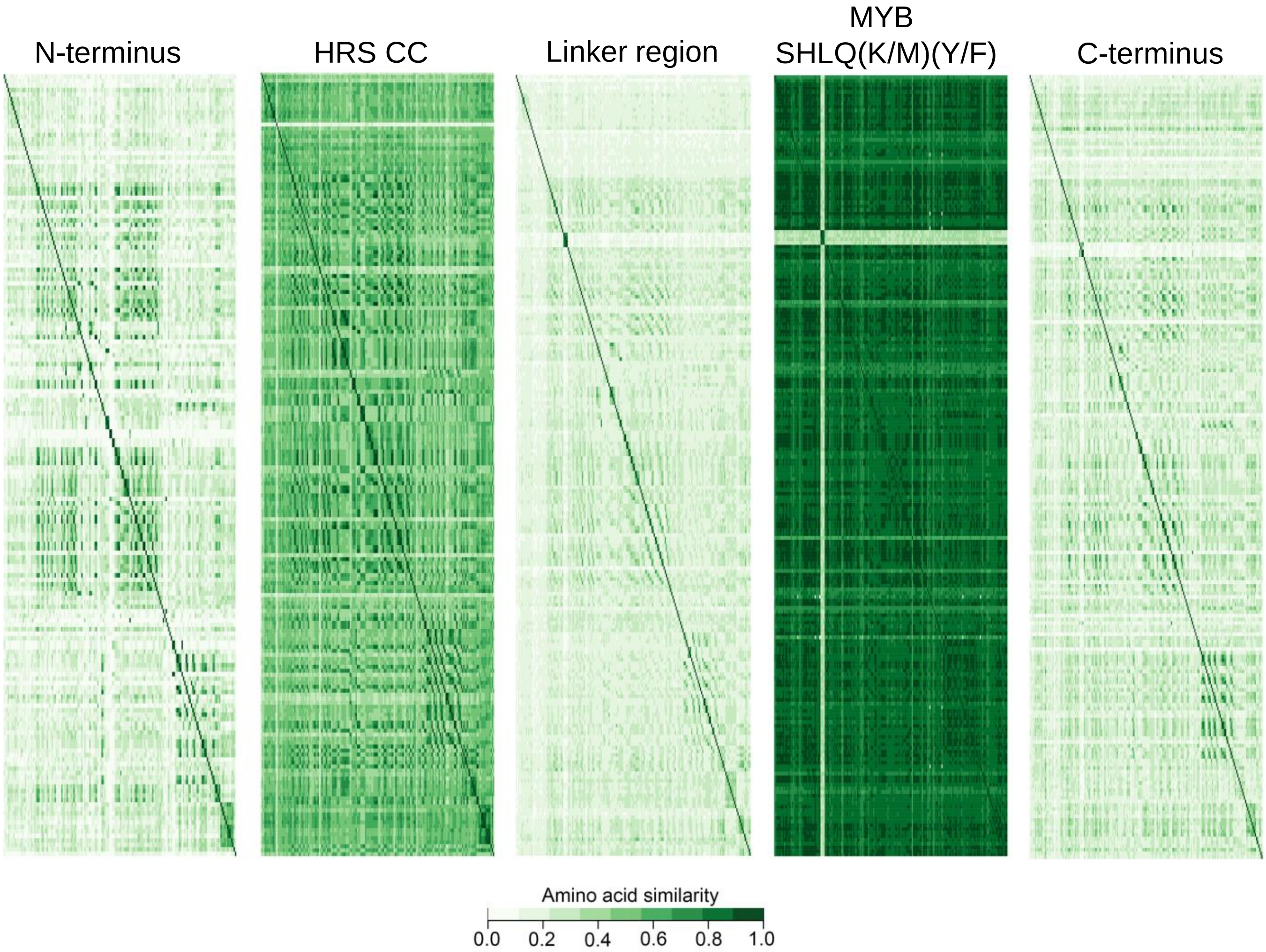


**Figure S9. Sequence similarity heatmap of different regions of HRS TFs across the plant lineage.** Different regions were identified through CDD v3.19-based domain analysis. The sequences were split accordingly and arranged as per the taxonomical position for the generation of sequence similarity heatmap.

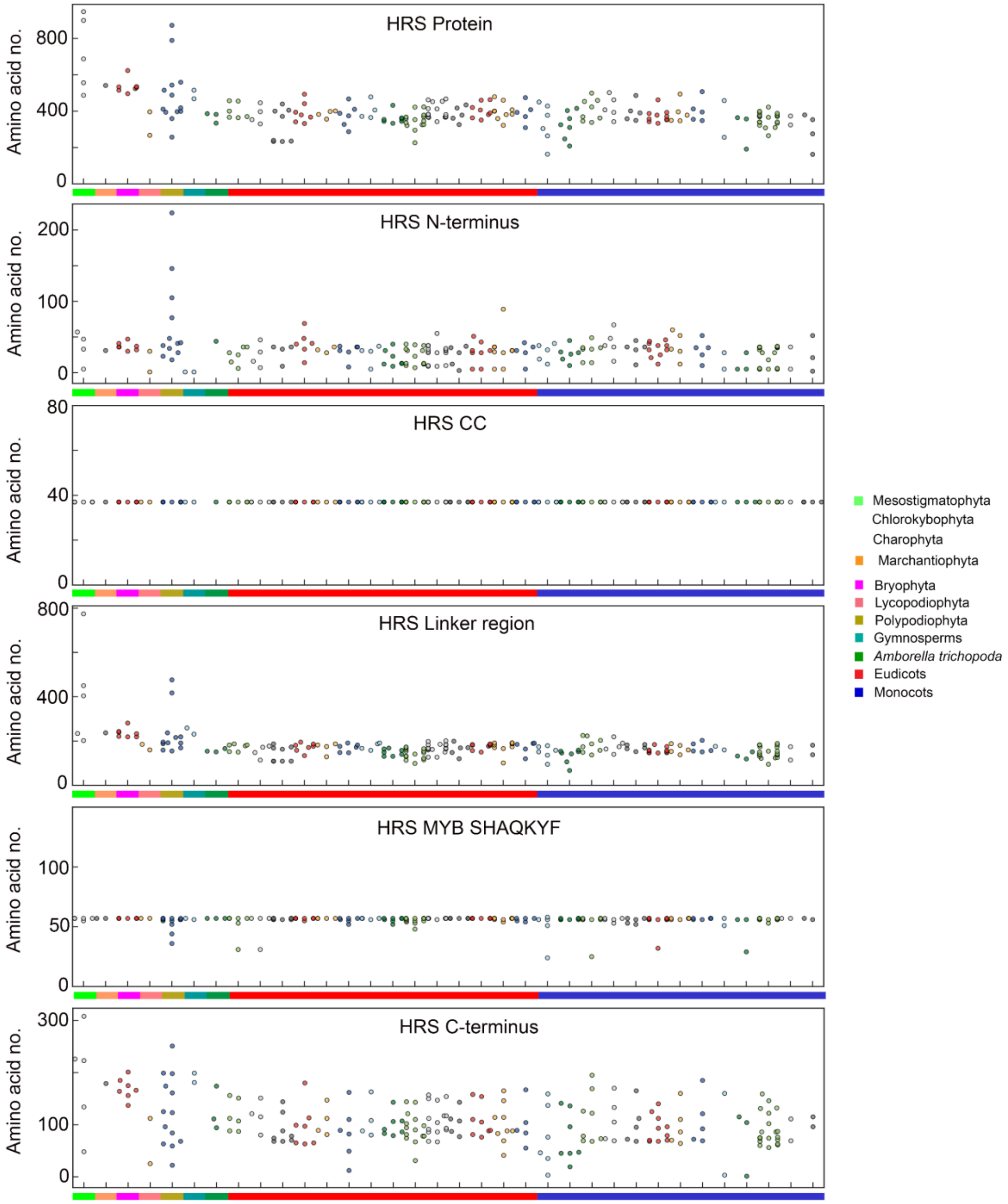


**Figure S10. Size distribution of HRS TFs and different regions across the plant lineage.**
Different regions were identified through CDD v3.19-based domain analysis. The color key highlights different taxonomical groups of Archaeplastida.

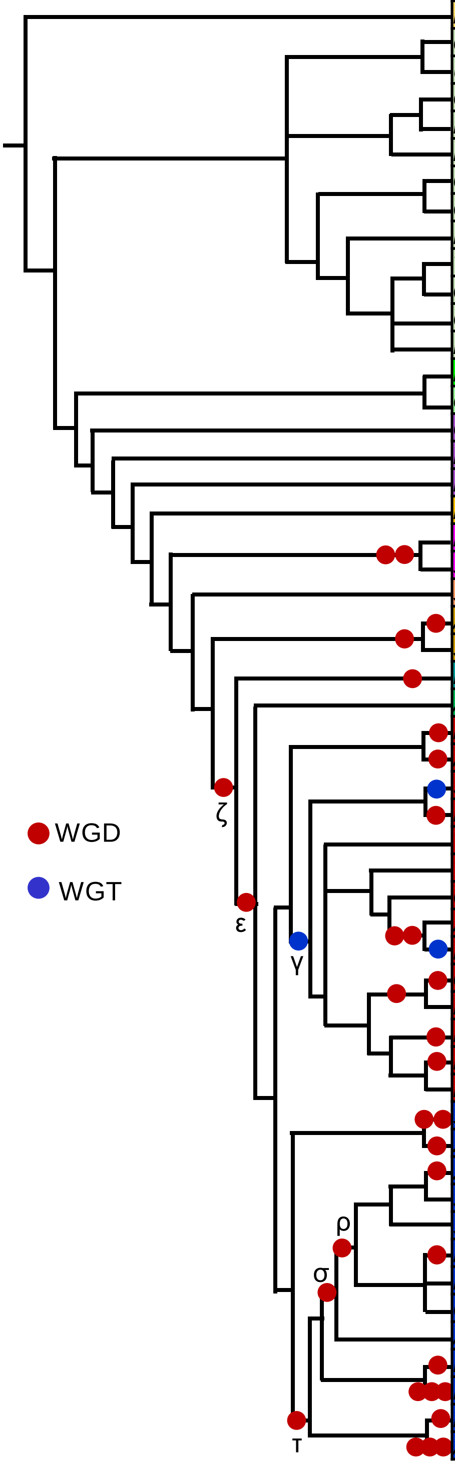


| Species | PSR | PSR SFI | PSR SFII | PSR SFIII | PSR SFIV | PSR SFV | PSR SFVI | PSR SFVII | PSR SFVIII | PSR other |
|---|---|---|---|---|---|---|---|---|---|---|
| *Prasinoderma coloniale* | 1 | 1 | 0 | 0 | 0 | 0 | 0 | 0 | 0 | 0 |
| *Caulerpa lentillifera* | 1 | 0 | 0 | 1 | 0 | 0 | 0 | 0 | 0 | 0 |
| *Ulva mutabilis* | 1 | 0 | 0 | 0 | 1 | 0 | 0 | 0 | 0 | 0 |
| *Ostreococcus lucimarinus* | 1 | 1 | 0 | 0 | 0 | 0 | 0 | 0 | 0 | 0 |
| *Bathycoccus prasinos* | 1 | 1 | 0 | 0 | 0 | 0 | 0 | 0 | 0 | 0 |
| *Micromonas pusilla* | 1 | 1 | 0 | 0 | 0 | 0 | 0 | 0 | 0 | 0 |
| *Chlorella variabilis* | 2 | 1 | 0 | 0 | 1 | 0 | 0 | 0 | 0 | 0 |
| *Coccomyxa subellipsoidea* | 1 | 1 | 0 | 0 | 0 | 0 | 0 | 0 | 0 | 0 |
| *Monoraphidium neglectum* | 2 | 2 | 0 | 0 | 0 | 0 | 0 | 0 | 0 | 0 |
| *Volvox carteri* | 1 | 1 | 0 | 0 | 0 | 0 | 0 | 0 | 0 | 0 |
| *Gonium pectorale* | 1 | 1 | 0 | 0 | 0 | 0 | 0 | 0 | 0 | 0 |
| *Chlamydomonas reinhardtii* | 1 | 1 | 0 | 0 | 0 | 0 | 0 | 0 | 0 | 0 |
| *Dunaliella salina* | 1 | 1 | 0 | 0 | 0 | 0 | 0 | 0 | 0 | 0 |
| *Mesostigma viride* | 1 | 1 | 0 | 0 | 0 | 0 | 0 | 0 | 0 | 0 |
| *Chlorokybus atmophyticus* | 1 | 1 | 0 | 0 | 0 | 0 | 0 | 0 | 0 | 0 |
| *Chara braunii* | 1 | 0 | 0 | 1 | 0 | 0 | 0 | 0 | 0 | 0 |
| *Klebsormidium nitens* | 2 | 2 | 0 | 0 | 0 | 0 | 0 | 0 | 0 | 0 |
| *Mesotaenium endlicherianum* | 1 | 1 | 0 | 0 | 0 | 0 | 0 | 0 | 0 | 0 |
| *Marchantia polymorpha* | 3 | 1 | 0 | 1 | 0 | 0 | 0 | 0 | 1 | 0 |
| *Physcomitrella patens* | 24 | 11 | 0 | 0 | 0 | 0 | 0 | 0 | 12 | 1 |
| *Sphagnum fallax* | 19 | 12 | 0 | 0 | 0 | 0 | 0 | 0 | 7 | 0 |
| *Selaginella moellendorffii* | 9 | 4 | 0 | 1 | 0 | 0 | 0 | 0 | 3 | 1 |
| *Azolla filiculoides* | 14 | 3 | 0 | 4 | 0 | 0 | 0 | 0 | 4 | 3 |
| *Salvinia cucullata* | 10 | 2 | 0 | 1 | 0 | 2 | 1 | 0 | 2 | 2 |
| *Picea abies* | 6 | 2 | 0 | 0 | 0 | 0 | 0 | 0 | 4 | 0 |
| *Amborella trichopoda* | 14 | 4 | 2 | 1 | 1 | 1 | 1 | 2 | 2 | 0 |
| *Nelumbo nucifera* | 20 | 3 | 1 | 2 | 2 | 3 | 1 | 3 | 5 | 0 |
| *Aquilegia coerulea* | 15 | 2 | 0 | 2 | 2 | 1 | 4 | 1 | 3 | 0 |
| *Solanum lycopersicum* | 17 | 3 | 0 | 3 | 2 | 2 | 1 | 3 | 3 | 0 |
| *Erythranthe guttata* | 22 | 2 | 0 | 3 | 3 | 4 | 2 | 3 | 5 | 0 |
| *Vitis vinifera* | 18 | 2 | 2 | 2 | 4 | 1 | 2 | 2 | 3 | 0 |
| *Theobroma cacao* | 15 | 1 | 2 | 2 | 2 | 1 | 2 | 2 | 3 | 0 |
| *Carica papaya* | 10 | 2 | 1 | 1 | 1 | 0 | 1 | 1 | 3 | 0 |
| *Arabidopsis thaliana* | 15 | 1 | 1 | 4 | 1 | 1 | 2 | 2 | 3 | 0 |
| *Brassica rapa* | 28 | 2 | 3 | 7 | 3 | 1 | 4 | 2 | 6 | 0 |
| *Glycine max* | 38 | 4 | 0 | 6 | 8 | 2 | 3 | 4 | 11 | 0 |
| *Medicago truncatula* | 17 | 2 | 0 | 3 | 3 | 1 | 1 | 2 | 5 | 0 |
| *Populus trichocarpa* | 24 | 2 | 3 | 3 | 2 | 2 | 3 | 3 | 6 | 0 |
| *Manihot esculenta* | 21 | 2 | 2 | 4 | 4 | 1 | 1 | 2 | 5 | 0 |
| *Ricinus communis* | 13 | 1 | 1 | 2 | 3 | 1 | 1 | 2 | 2 | 0 |
| *Spirodela polyrhiza* | 10 | 3 | 0 | 2 | 1 | 0 | 1 | 0 | 2 | 1 |
| *Zostera marina* | 14 | 5 | 2 | 1 | 1 | 1 | 1 | 1 | 2 | 0 |
| *Zea mays* | 20 | 3 | 2 | 2 | 2 | 2 | 2 | 2 | 4 | 1 |
| *Sorghum bicolor* | 17 | 2 | 3 | 2 | 1 | 2 | 2 | 1 | 3 | 1 |
| *Setaria italica* | 18 | 3 | 2 | 2 | 1 | 2 | 2 | 2 | 3 | 1 |
| *Phyllostachys edulis* | 25 | 2 | 3 | 5 | 1 | 3 | 4 | 1 | 4 | 2 |
| *Brachypodium distachyon* | 19 | 4 | 3 | 2 | 1 | 2 | 2 | 1 | 3 | 1 |
| *Oryza Sativa* | 18 | 4 | 2 | 2 | 1 | 1 | 2 | 2 | 3 | 1 |
| *Ananas comosus* | 15 | 3 | 1 | 2 | 1 | 1 | 2 | 1 | 4 | 0 |
| *Phoenix dactylifera* | 21 | 6 | 1 | 3 | 2 | 1 | 2 | 3 | 3 | 0 |
| *Musa acuminata* | 29 | 5 | 2 | 5 | 2 | 2 | 3 | 5 | 5 | 0 |
| *Phalaenopsis equestris* | 16 | 3 | 0 | 3 | 1 | 1 | 2 | 2 | 4 | 0 |
| *Asparagus officinalis* | 17 | 4 | 1 | 2 | 1 | 1 | 2 | 3 | 3 | 0 |

**Figure S11. Distribution of PSR TF subfamilies in Archaeplastida.**
The phylogenetic reconstruction analysis (Figure 3) was used for the classification of PSR TFs to different subfamilies. The major Whole Genome Duplication (WGD) and Whole Genome Triplication (WGT) events are indicated in the species tree.

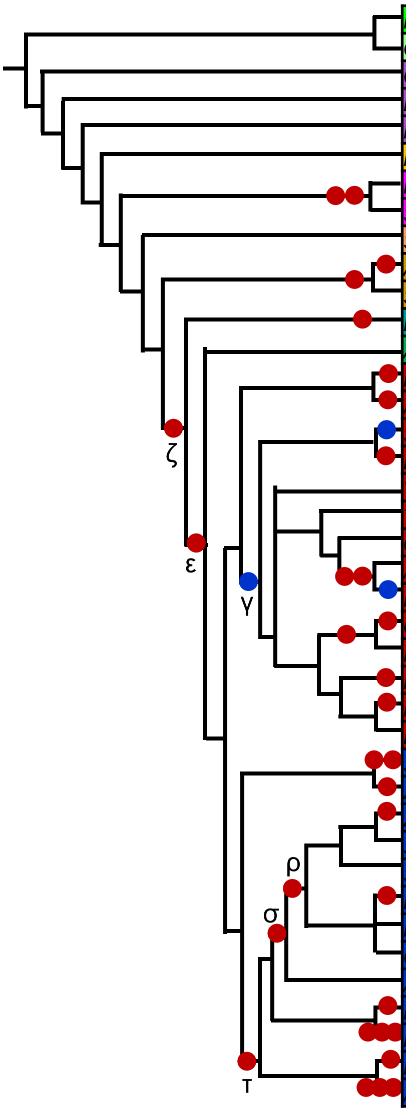


| Species | HRS | HRS SFI | HRS SFII | HRS SFIII | HRS SFIV |
|---|---|---|---|---|---|
| *Mesostigma viride* | 1 | 1 | 0 | 0 | 0 |
| *Chlorokybus atmophyticus* | 1 | 1 | 0 | 0 | 0 |
| *Chara braunii* | 1 | 1 | 0 | 0 | 0 |
| *Klebsormidium nitens* | 1 | 1 | 0 | 0 | 0 |
| *Mesotaenium endlicherianum* | 1 | 0 | 0 | 1 | 0 |
| *Marchantia polymorpha* | 1 | 0 | 0 | 1 | 0 |
| *Physcomitrella patens* | 3 | 0 | 0 | 3 | 0 |
| *Sphagnum fallax* | 4 | 0 | 0 | 4 | 0 |
| *Selaginella moellendorffii* | 2 | 0 | 0 | 2 | 0 |
| *Azolla filiculoides* | 8 | 8 | 0 | 0 | 0 |
| *Salvinia cucullata* | 5 | 5 | 0 | 0 | 0 |
| *Picea abies* | 2 | 0 | 0 | 2 | 0 |
| *Amborella trichopoda* | 3 | 0 | 1 | 1 | 1 |
| *Nelumbo nucifera* | 6 | 0 | 2 | 2 | 2 |
| *Aquilegia coerulea* | 4 | 1 | 1 | 1 | 1 |
| *Solanum lycopersicum* | 8 | 4 | 1 | 1 | 2 |
| *Erythranthe guttata* | 7 | 0 | 2 | 1 | 4 |
| *Vitis vinifera* | 4 | 0 | 1 | 1 | 2 |
| *Theobroma cacao* | 6 | 0 | 1 | 1 | 4 |
| *Carica papaya* | 4 | 0 | 1 | 1 | 2 |
| *Arabidopsis thaliana* | 7 | 1 | 4 | 1 | 1 |
| *Brassica rapa* | 14 | 1 | 8 | 2 | 3 |
| *Glycine max* | 15 | 0 | 4 | 4 | 7 |
| *Medicago truncatula* | 5 | 0 | 2 | 1 | 2 |
| *Populus trichocarpa* | 8 | 0 | 2 | 2 | 4 |
| *Manihot esculenta* | 9 | 0 | 2 | 2 | 5 |
| *Ricinus communis* | 5 | 0 | 1 | 1 | 3 |
| *Spirodela polyrhiza* | 6 | 1 | 2 | 1 | 2 |
| *Zostera marina* | 7 | 2 | 1 | 2 | 2 |
| *Zea mays* | 8 | 0 | 3 | 1 | 4 |
| *Sorghum bicolor* | 5 | 0 | 1 | 1 | 3 |
| *Setaria italica* | 5 | 0 | 1 | 1 | 3 |
| *Phyllostachys edulis* | 10 | 0 | 1 | 2 | 7 |
| *Brachypodium distachyon* | 5 | 0 | 1 | 1 | 3 |
| *Oryza Sativa* | 5 | 0 | 1 | 1 | 3 |
| *Ananas comosus* | 2 | 0 | 1 | 1 | 0 |
| *Phoenix dactylifera* | 3 | 0 | 2 | 1 | 0 |
| *Musa acuminata* | 19 | 2 | 5 | 2 | 10 |
| *Phalaenopsis equestris* | 2 | 0 | 1 | 0 | 1 |
| *Asparagus officinalis* | 4 | 0 | 3 | 0 | 1 |

**Figure S12. Distribution of HRS TF subfamilies in Archaeplastida.**
The phylogenetic reconstruction analysis (Figure 4A) was used for the classification of HRS TFs to different subfamilies. The major Whole Genome Duplication (WGD) and Whole Genome Triplication (WGT) events are indicated in the species tree.

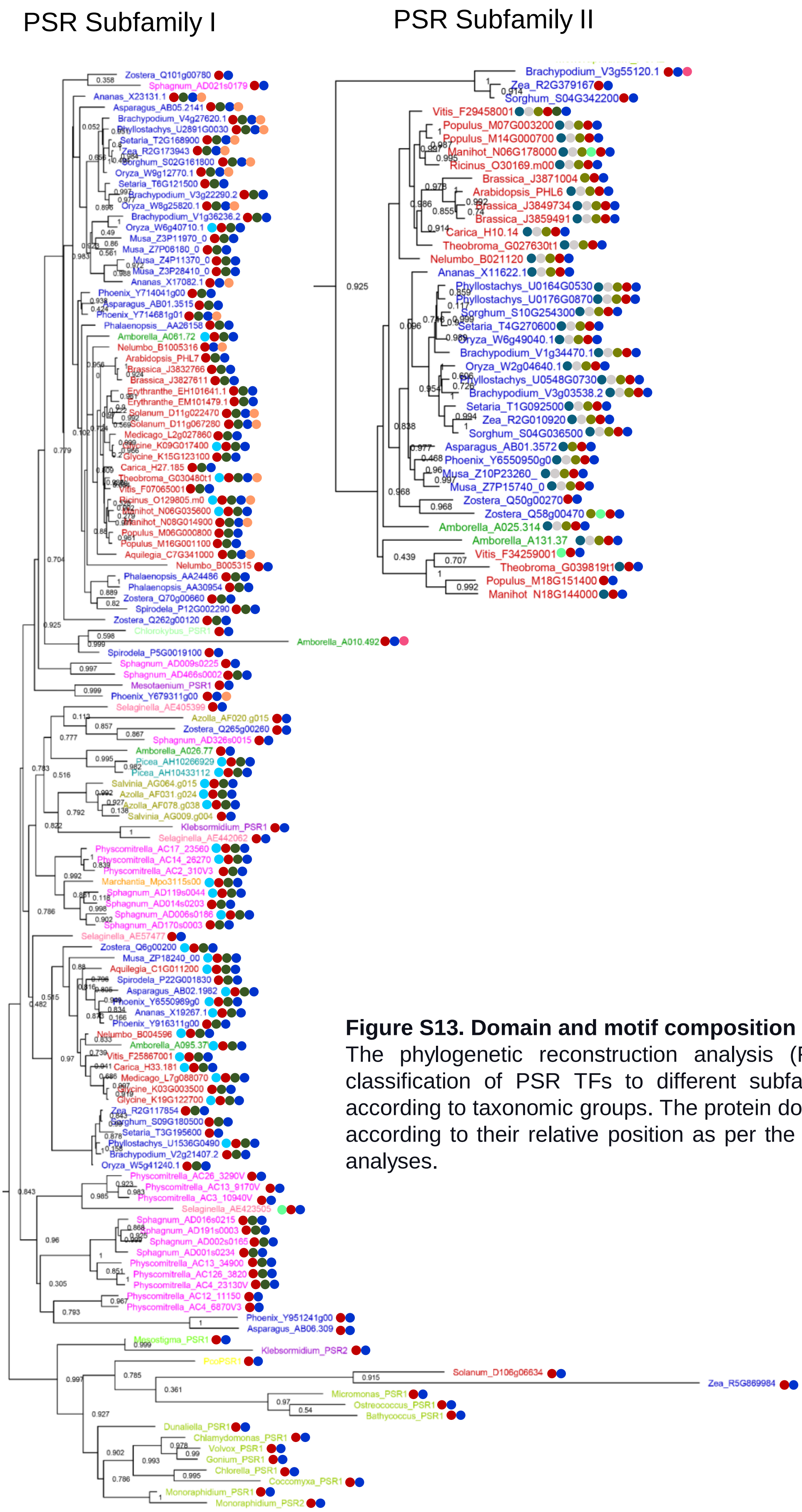


| Color code | Domain/motif |
|---|---|
| | MYB SHLQ(K/M)(Y/F) |
| | MYB_CC_LHEQLE |
| | PSR motif 1 |
| | PSR motif 2 |
| | PSR motif 3 |
| | PSR motif 4 |
| | PSR motif 5 |
| | PSR motif 6 |
| | PSR motif 7 |
| | PSR motif 8 |
| | PSR motif 9 |
| | PSR motif 10 |
| | PSR motif 11 |
| | PSR motif 12 |
| | PSR motif 13 |
| | PSR motif 14 |
| | PSR motif 15 |
| | PSR motif 16 |
| | PSR motif 17 |
| | PSR motif 18 |
| | PSR motif 19 |
| | PSR motif 20 |
| | PSR motif 21 |
| | PSR motif 22 |
| | PSR motif 23 |
| | PSR motif 24 |

**Figure S13. Domain and motif composition of PSR subfamilies.** The phylogenetic reconstruction analysis (Figure 3) was used for the classification of PSR TFs to different subfamilies. Branches are colored according to taxonomic groups. The protein domains and motifs are arranged according to their relative position as per the CDD v3.19 and MEME v5.3.3 analyses.

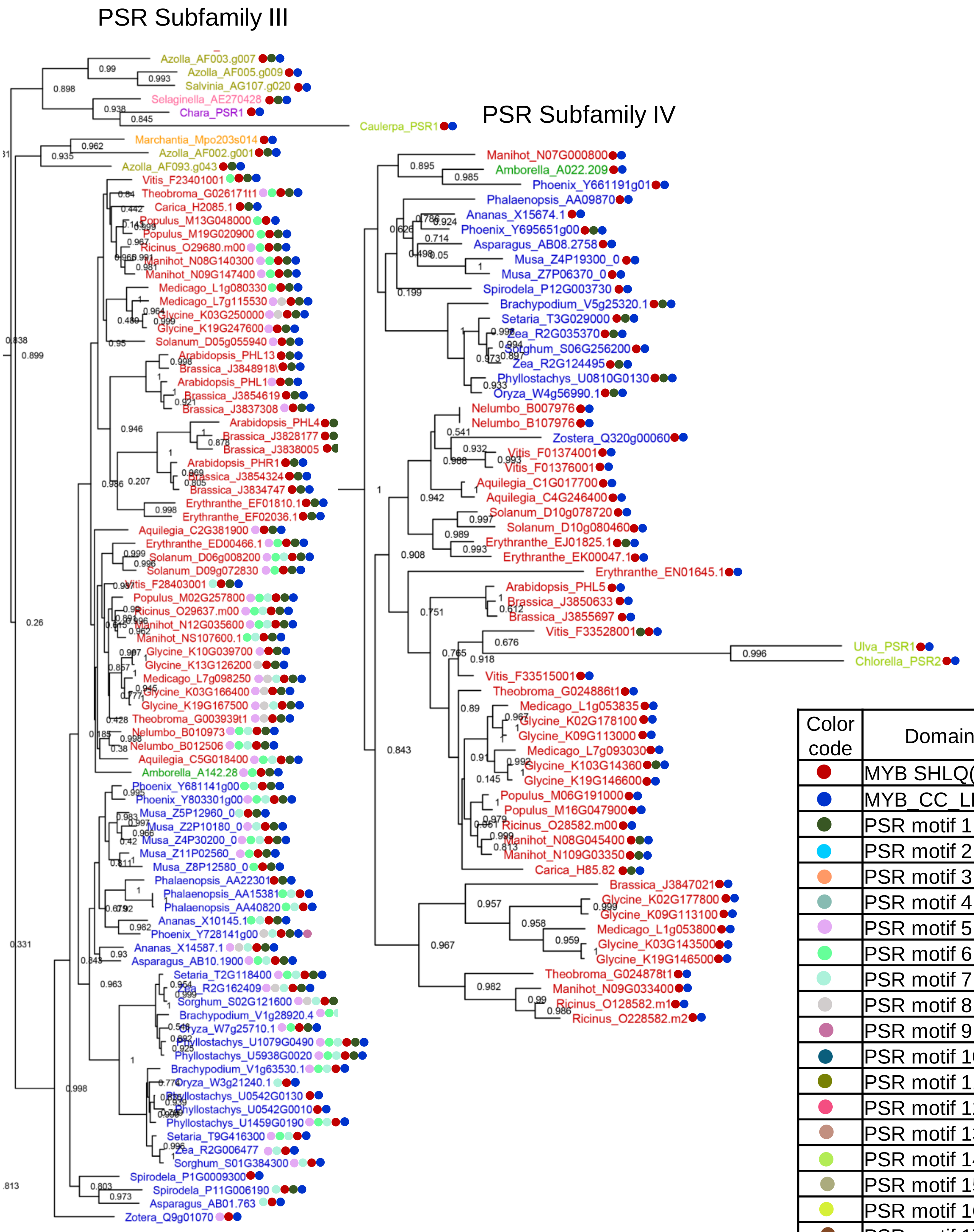


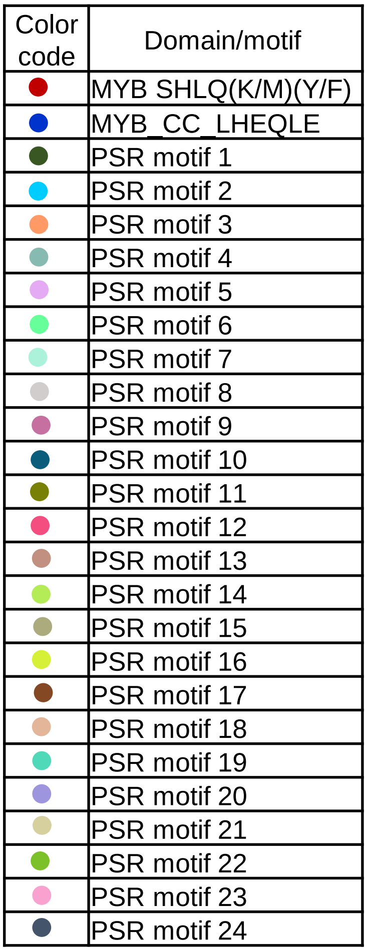


**Figure S13 (continued). Domain and motif composition of PSR subfamilies.**
The phylogenetic reconstruction analysis (Figure 3) was used for the classification of PSR TFs to different subfamilies. Branches are colored according to taxonomic groups. The protein domains and motifs are arranged according to their relative position as per the CDD v3.19 and MEME v5.3.3 analyses.

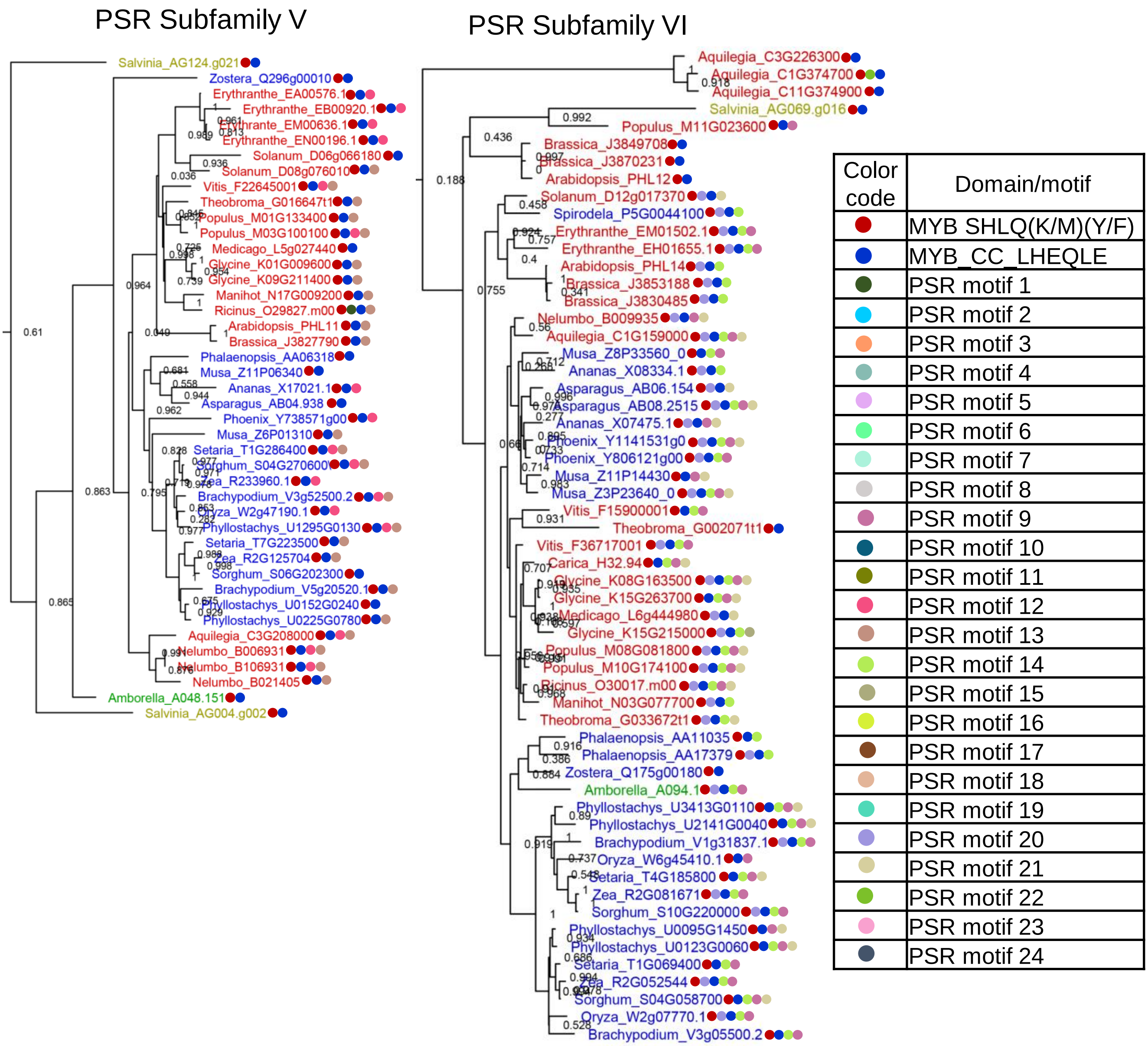


**Figure S13 (continued). Domain and motif composition of PSR subfamilies.**
The phylogenetic reconstruction analysis (Figure 3) was used for the classification of PSR TFs to different subfamilies. Branches are colored according to taxonomic groups. The protein domains and motifs are arranged according to their relative position as per the CDD v3.19 and MEME v5.3.3 analyses.

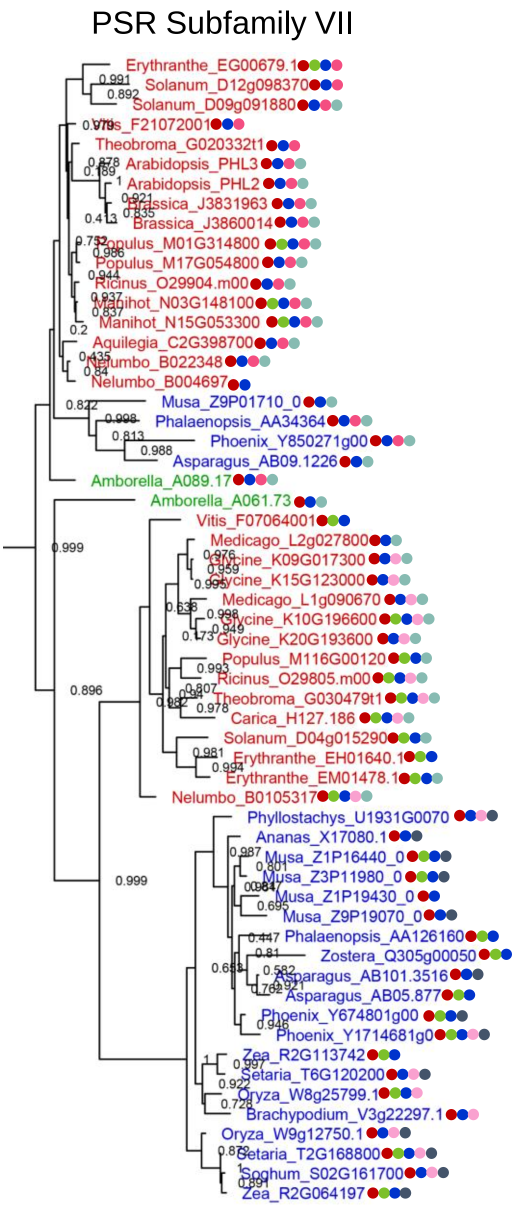


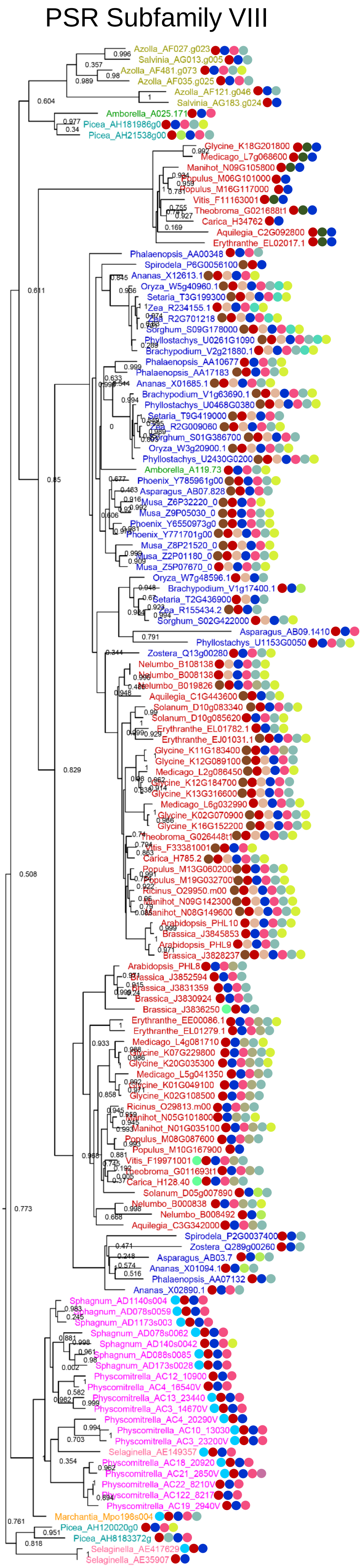


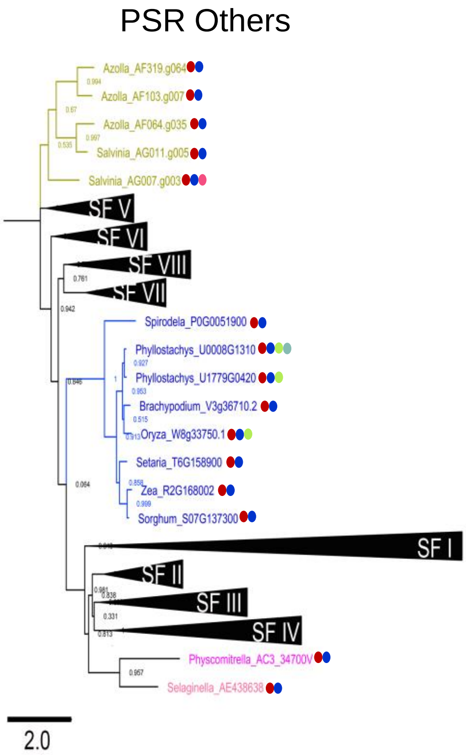


| Color code | Domain/motif |
|---|---|
| | MYB SHLQ(K/M)(Y/F) |
| | MYB_CC_LHEQLE |
| | PSR motif 1 |
| | PSR motif 2 |
| | PSR motif 3 |
| | PSR motif 4 |
| | PSR motif 5 |
| | PSR motif 6 |
| | PSR motif 7 |
| | PSR motif 8 |
| | PSR motif 9 |
| | PSR motif 10 |
| | PSR motif 11 |
| | PSR motif 12 |
| | PSR motif 13 |
| | PSR motif 14 |
| | PSR motif 15 |
| | PSR motif 16 |
| | PSR motif 17 |
| | PSR motif 18 |
| | PSR motif 19 |
| | PSR motif 20 |
| | PSR motif 21 |
| | PSR motif 22 |
| | PSR motif 23 |
| | PSR motif 24 |

**Figure S13 (continued). Domain and motif composition of PSR subfamilies.**
The phylogenetic reconstruction analysis (Figure 3) was used for the classification of PSR TFs to different subfamilies. Branches are colored according to taxonomic groups. The protein domains and motifs are arranged according to their relative position as per the CDD v3.19 and MEME v5.3.3 analyses.

HRS Subfamily I

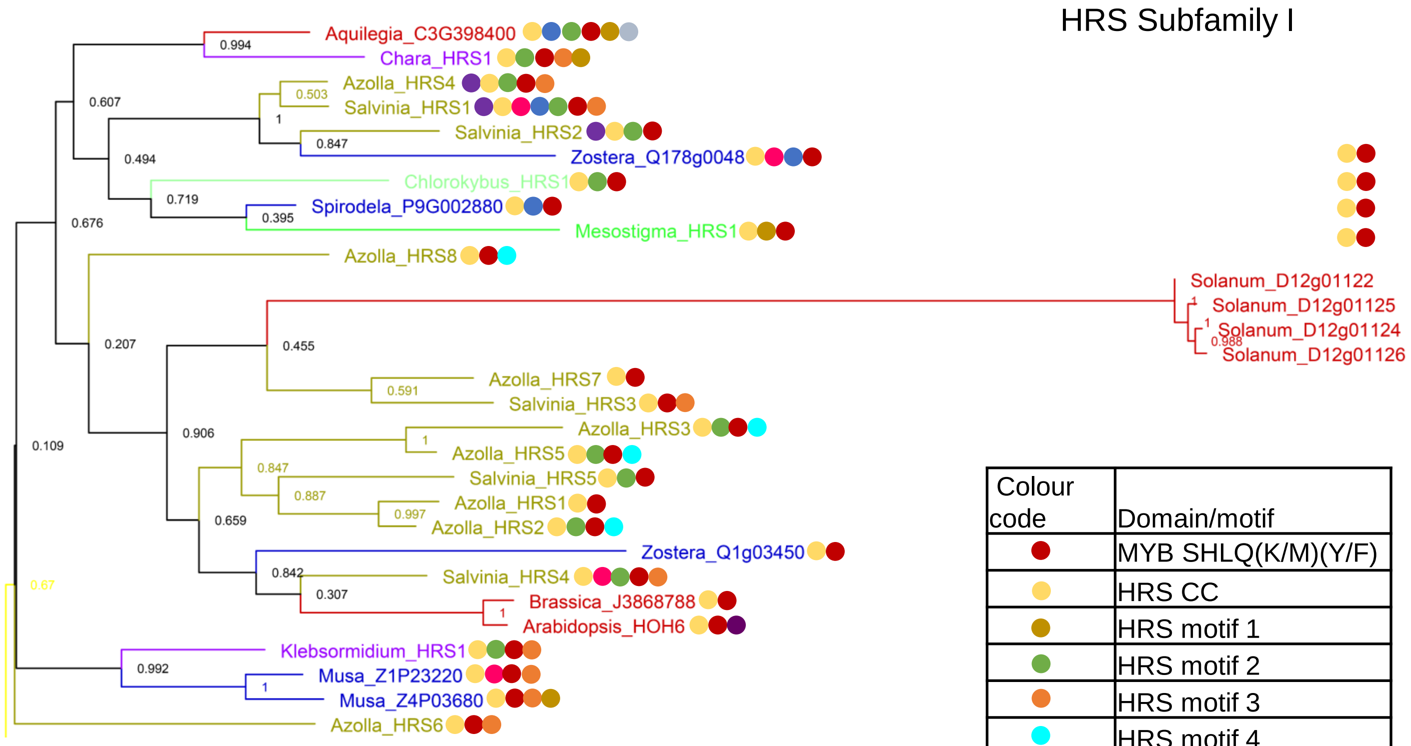


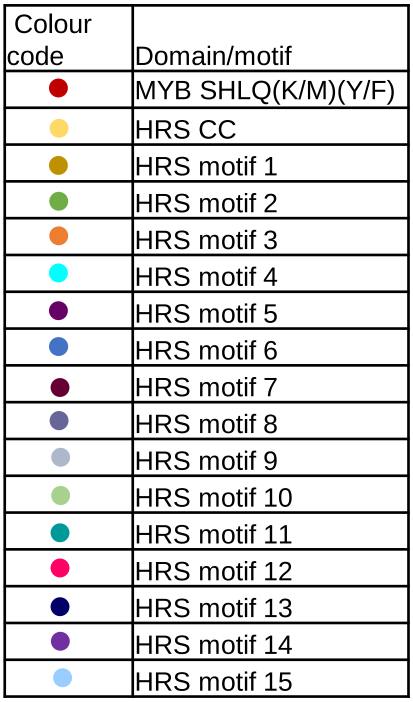


**Figure S14. Domain and motif composition of HRS subfamilies.**
The phylogenetic reconstruction analysis (Figure 3) was used for the classification of HRS TFs to different subfamilies. Branches are colored according to taxonomic groups. The protein domains and motifs are arranged according to their relative position as per the CDD v3.19 and MEME v5.3.3 analyses.

HRS Subfamily II

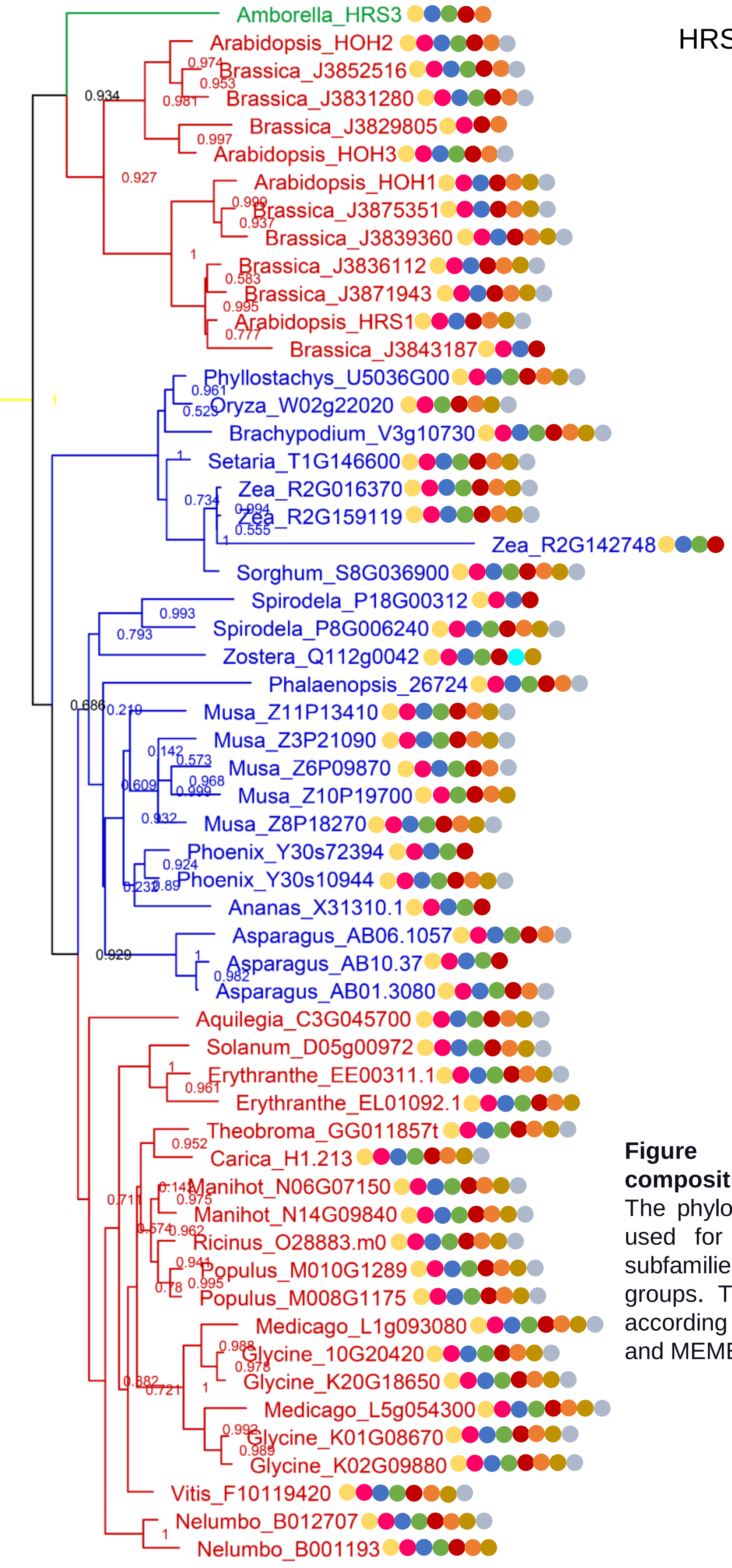


| Colour code | Domain/motif |
|---|---|
| | MYB SHLQ(K/M)(Y/F) |
| | HRS CC |
| | HRS motif 1 |
| | HRS motif 2 |
| | HRS motif 3 |
| | HRS motif 4 |
| | HRS motif 5 |
| | HRS motif 6 |
| | HRS motif 7 |
| | HRS motif 8 |
| | HRS motif 9 |
| | HRS motif 10 |
| | HRS motif 11 |
| | HRS motif 12 |
| | HRS motif 13 |
| | HRS motif 14 |
| | HRS motif 15 |

**Figure S14 (continued). Domain and motif composition of HRS subfamilies.**
The phylogenetic reconstruction analysis (Figure 3) was used for the classification of HRS TFs to different subfamilies. Branches are colored according to taxonomic groups. The protein domains and motifs are arranged according to their relative position as per the CDD v3.19 and MEME v5.3.3 analyses.

### HRS Subfamily III

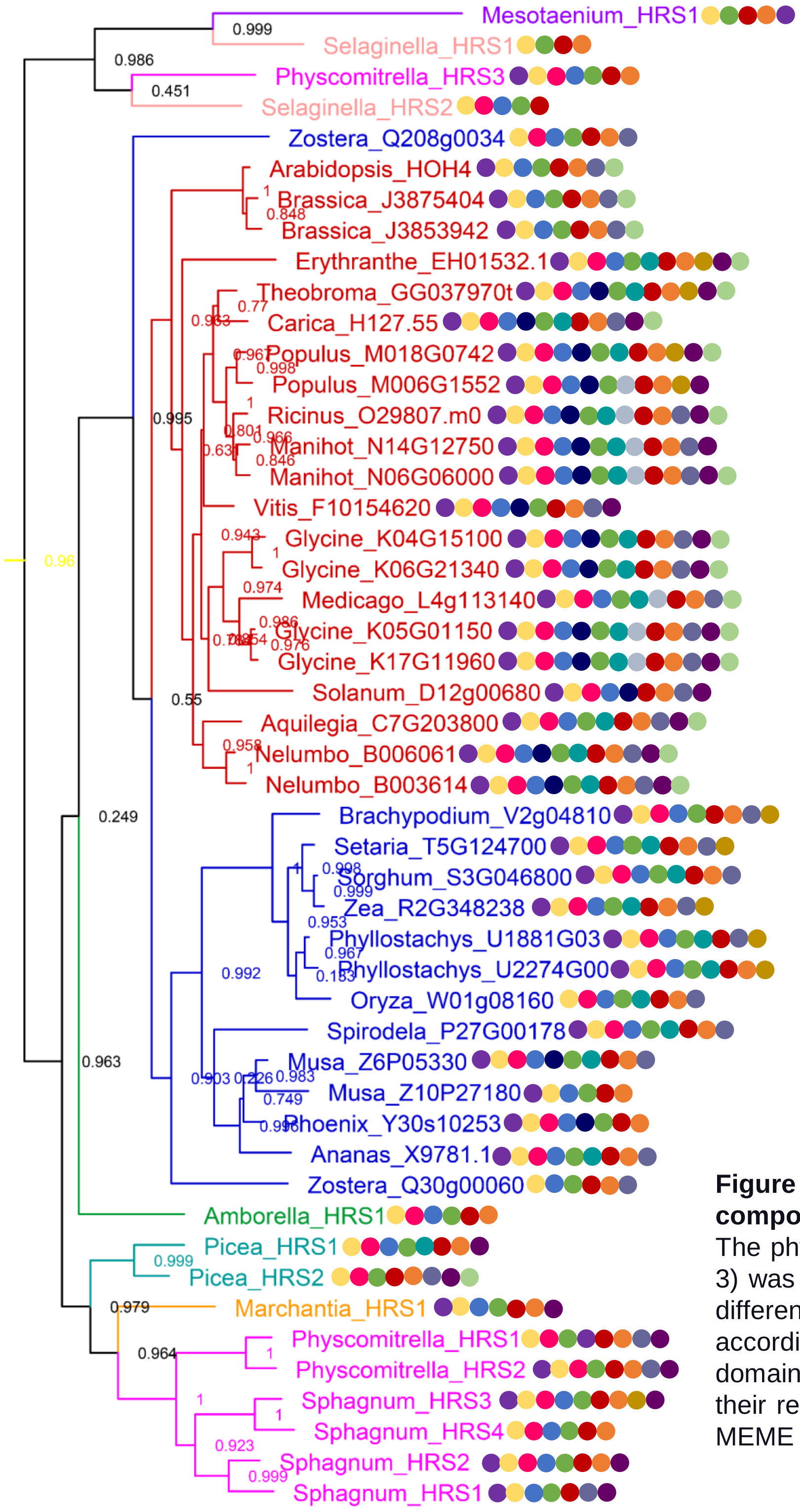


| Colour code | Domain/motif |
|---|---|
| | MYB SHLQ(K/M)(Y/F) |
| | HRS CC |
| | HRS motif 1 |
| | HRS motif 2 |
| | HRS motif 3 |
| | HRS motif 4 |
| | HRS motif 5 |
| | HRS motif 6 |
| | HRS motif 7 |
| | HRS motif 8 |
| | HRS motif 9 |
| | HRS motif 10 |
| | HRS motif 11 |
| | HRS motif 12 |
| | HRS motif 13 |
| | HRS motif 14 |
| | HRS motif 15 |

**Figure S14 (continued). Domain and motif composition of HRS subfamilies.**
The phylogenetic reconstruction analysis (Figure 3) was used for the classification of HRS TFs to different subfamilies. Branches are colored according to taxonomic groups. The protein domains and motifs are arranged according to their relative position as per the CDD v3.19 and MEME v5.3.3 analyses.

HRS Subfamily IV

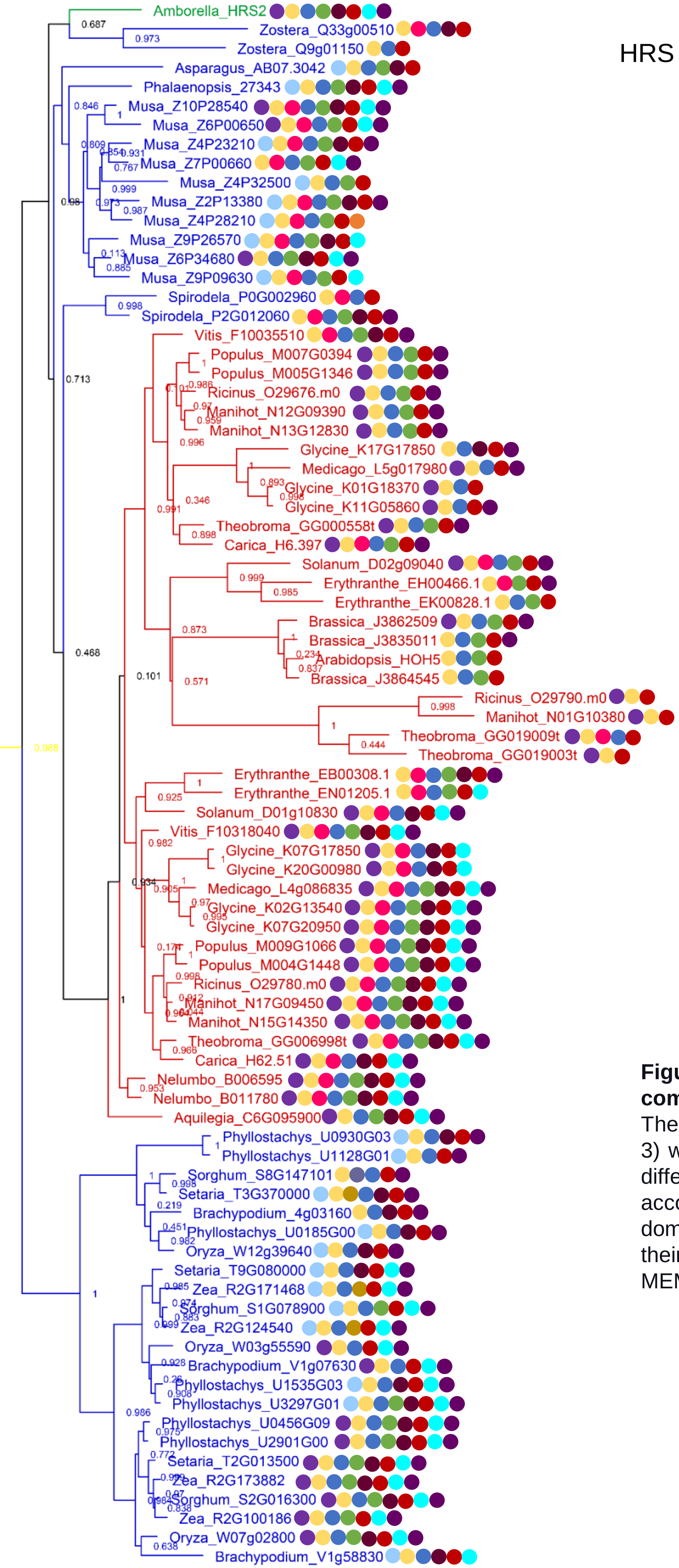


| Colour code | Domain/motif |
|---|---|
| | MYB SHLQ(K/M)(Y/F) |
| | HRS CC |
| | HRS motif 1 |
| | HRS motif 2 |
| | HRS motif 3 |
| | HRS motif 4 |
| | HRS motif 5 |
| | HRS motif 6 |
| | HRS motif 7 |
| | HRS motif 8 |
| | HRS motif 9 |
| | HRS motif 10 |
| | HRS motif 11 |
| | HRS motif 12 |
| | HRS motif 13 |
| | HRS motif 14 |
| | HRS motif 15 |

**Figure S14 (continued). Domain and motif composition of HRS subfamilies.**
The phylogenetic reconstruction analysis (Figure 3) was used for the classification of HRS TFs to different subfamilies. Branches are colored according to taxonomic groups. The protein domains and motifs are arranged according to their relative position as per the CDD v3.19 and MEME v5.3.3 analyses.

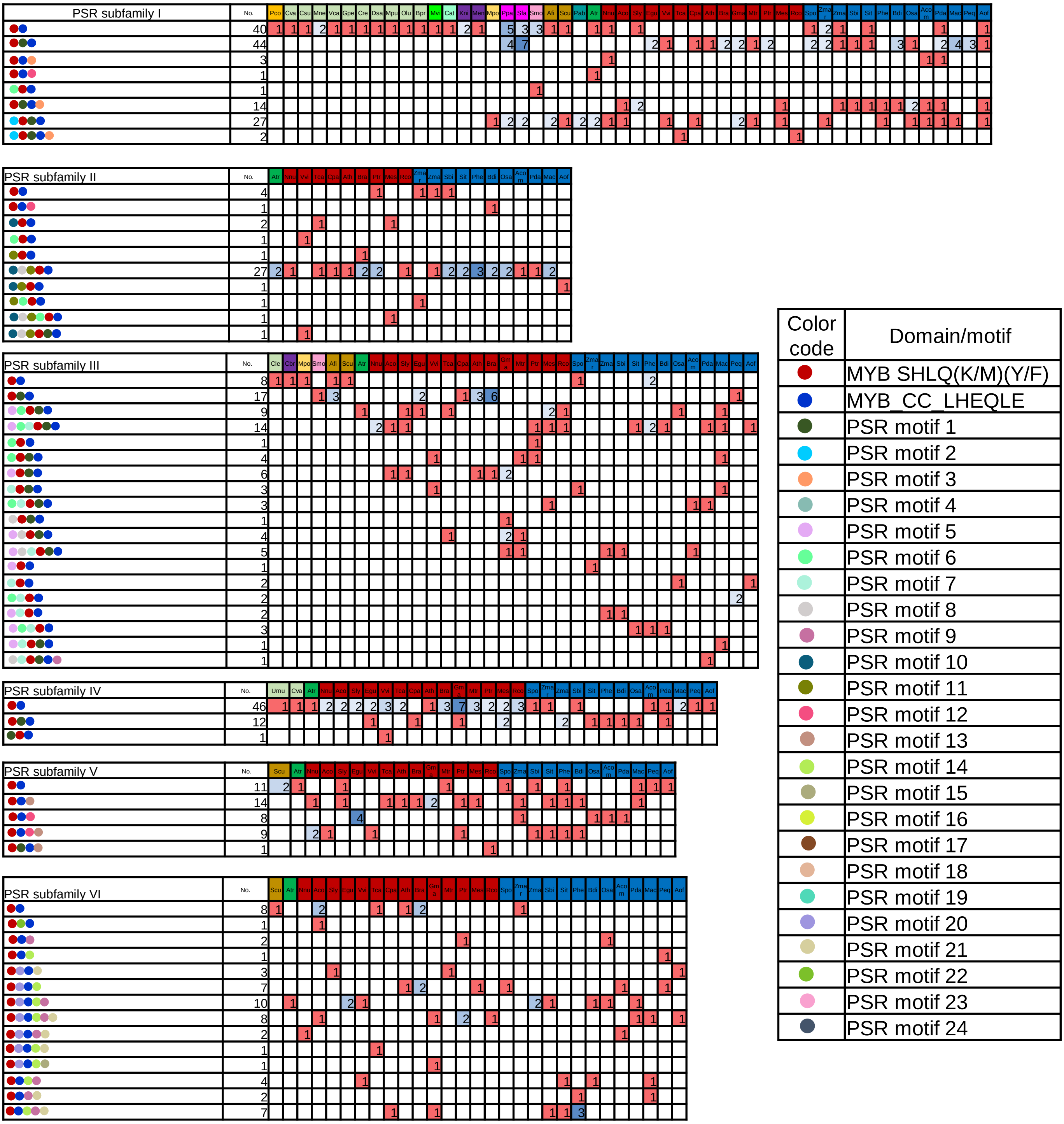


**Figure S15. Domain and motif organization in PSR subfamilies.**
The proteins were grouped in each subfamily according to the similarity/divergence in the protein domain and motif organization. The phylogenetic reconstruction analysis (Figure 3) was used for the classification of PSR TFs to different subfamilies. The protein domains and motifs are arranged according to their relative position as per the CDD v3.19 and MEME v5.3.3 analyses.

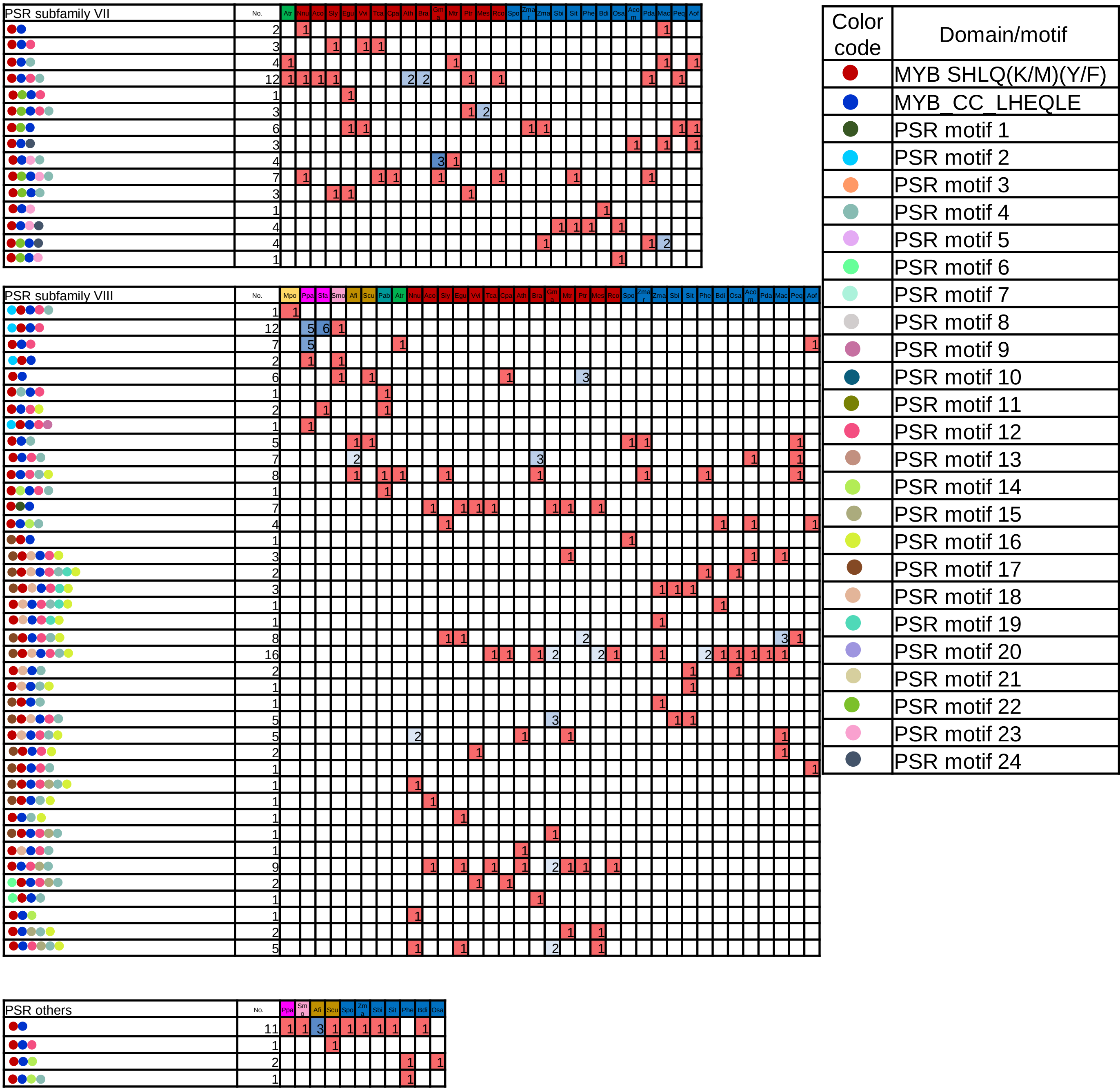


**Figure S15 (continued). Domain and motif organization in PSR subfamilies.**
The proteins were grouped in each subfamily according to the similarity/divergence in the protein domain and motif organization. The phylogenetic reconstruction analysis (Figure 3) was used for the classification of PSR TFs to different subfamilies. The protein domains and motifs are arranged according to their relative position as per the CDD v3.19 and MEME v5.3.3 analyses.

| Colour code | Domain/motif |
|---|---|
| | MYB SHLQ(K/M)(Y/F) |
| | HRS1 |
| | HRS motif 1 |
| | HRS motif 2 |
| | HRS motif 3 |
| | HRS motif 4 |
| | HRS motif 5 |
| | HRS motif 6 |
| | HRS motif 7 |
| | HRS motif 8 |
| | HRS motif 9 |
| | HRS motif 10 |
| | HRS motif 11 |
| | HRS motif 12 |
| | HRS motif 13 |
| | HRS motif 14 |
| | HRS motif 15 |

| HRS Subfamily I | No | Mvi | Cat | Kni | Cbr | Afi | Scu | Aco | Sly | Ath | Bra | Spo | Zmar | Mac |
|---|---|---|---|---|---|---|---|---|---|---|---|---|---|---|
| | 1 | 1 | | | | | | | | | | | | |
| | 2 | | 1 | | | | 1 | | | | | | | |
| | 1 | | | 1 | | | | | | | | | | |
| | 1 | | | | 1 | | | | | | | | | |
| | 3 | | | | | 3 | | | | | | | | |
| | 1 | | | | | 1 | | | | | | | | |
| | 1 | | | | | | 1 | | | | | | | |
| | 1 | | | | | | 1 | | | | | | | |
| | 8 | | | | | 2 | | | 4 | | 1 | | 1 | |
| | 2 | | | | | 1 | 1 | | | | | | | |
| | 1 | | | | | 1 | | | | | | | | |
| | 1 | | | | | | 1 | | | | | | | |
| | 1 | | | | | | | 1 | | | | | | |
| | 1 | | | | | | | | | 1 | | | | |
| | 1 | | | | | | | | | | | | 1 | |
| | 1 | | | | | | | | | | | 1 | | |
| | 1 | | | | | | | | | | | | | 1 |
| | 1 | | | | | | | | | | | | | 1 |
| Total | 29 | 1 | 1 | 1 | 1 | 8 | 5 | 1 | 4 | 1 | 1 | 1 | 2 | 2 |

| HRS Subfamily II | No. | Atr | Nnu | Aco | Sly | Egu | Vvi | Tca | Cpa | Ath | Bra | Gma | Mtr | Ptr | Mes | Rco | Spo | Zmar | Zma | Sbi | Svi | Phe | Bdi | Osa | Acom | Pda | Mac | Peq | Aof |
|---|---|---|---|---|---|---|---|---|---|---|---|---|---|---|---|---|---|---|---|---|---|---|---|---|---|---|---|---|---|
| | 1 | 1 | | | | | | | | | | | | | | | | | | | | | | | | | | | |
| | 2 | | 1 | | | 1 | | | | | | | | | | | | | | | | | | | | | | | |
| | 29 | | 1 | 1 | 1 | 1 | 1 | 1 | 1 | | | 4 | 2 | 2 | 2 | 1 | 1 | | 2 | 1 | 1 | 1 | 1 | | | 1 | 3 | | |
| | 1 | | | | | | | | | | 1 | | | | | | | | | | | | | | | | | | |
| | 2 | | | | | | | | | | 1 | | | | | | 1 | | | | | | | | | | | | |
| | 8 | | | | | | | | | 2 | 2 | | | | | | | | | | | | | | | | 1 | 1 | 2 |
| | 6 | | | | | | | | | 2 | 4 | | | | | | | | | | | | | | | | | | |
| | 1 | | | | | | | | | | | | | | | | | 1 | | | | | | | | | | | |
| | 1 | | | | | | | | | | | | | | | | | | 1 | | | | | | | | | | |
| | 1 | | | | | | | | | | | | | | | | | | | | | | | 1 | | | | | |
| | 3 | | | | | | | | | | | | | | | | | | | | | | | | 1 | 1 | | | 1 |
| | 1 | | | | | | | | | | | | | | | | | | | | | | | | | | 1 | | |
| Total | 56 | 1 | 2 | 1 | 1 | 2 | 1 | 1 | 1 | 4 | 8 | 4 | 2 | 2 | 2 | 1 | 2 | 1 | 3 | 1 | 1 | 1 | 1 | 1 | 1 | 2 | 5 | 1 | 3 |

**Figure S16. Domain and motif organization in HRS subfamilies.**

The proteins were grouped in each subfamily according to the similarity/divergence in the protein domain and motif organization. The phylogenetic reconstruction analysis (Figure 3) was used for the classification of PSR TFs to different subfamilies. The protein domains and motifs are arranged according to their relative position as per the CDD v3.19 and MEME v5.3.3 analyses.

| Colour code | Domain/motif |
|---|---|
| | MYB SHLQ(K/M)(Y/F) |
| | HRS1 |
| | HRS motif 1 |
| | HRS motif 2 |
| | HRS motif 3 |
| | HRS motif 4 |
| | HRS motif 5 |
| | HRS motif 6 |
| | HRS motif 7 |
| | HRS motif 8 |
| | HRS motif 9 |
| | HRS motif 10 |
| | HRS motif 11 |
| | HRS motif 12 |
| | HRS motif 13 |
| | HRS motif 14 |
| | HRS motif 15 |

| HRS Subfamily III | No. | Men | Mpo | Ppa | Sfa | Smo | Pab | Atr | Nnu | Aco | Sly | Egu | Vvi | Tca | Cpa | Ath | Bra | Gma | Mtr | Ptr | Mes | Rco | Spo | Zmar | Zma | Sbi | Sit | Phe | Bdi | Osa | Acom | Pda | Mac |
|---|---|---|---|---|---|---|---|---|---|---|---|---|---|---|---|---|---|---|---|---|---|---|---|---|---|---|---|---|---|---|---|---|---|
| | 1 | 1 | | | | | | | | | | | | | | | | | | | | | | | | | | | | | | | |
| | 1 | | 1 | | | | | | | | | | | | | | | | | | | | | | | | | | | | | | |
| | 1 | | | 1 | | | | | | | | | | | | | | | | | | | | | | | | | | | | | |
| | 1 | | | 1 | | | | | | | | | | | | | | | | | | | | | | | | | | | | | |
| | 1 | | | 1 | | | | | | | | | | | | | | | | | | | | | | | | | | | | | |
| | 1 | | | | 1 | | | | | | | | | | | | | | | | | | | | | | | | | | | | |
| | 1 | | | | 1 | | | | | | | | | | | | | | | | | | | | | | | | | | | | |
| | 1 | | | | 1 | | | | | | | | | | | | | | | | | | | | | | | | | | | | |
| | 1 | | | | 1 | | | | | | | | | | | | | | | | | | | | | | | | | | | | |
| | 1 | | | | | 1 | | | | | | | | | | | | | | | | | | | | | | | | | | | |
| | 1 | | | | | 1 | | | | | | | | | | | | | | | | | | | | | | | | | | | |
| | 1 | | | | | | 1 | | | | | | | | | | | | | | | | | | | | | | | | | | |
| | 1 | | | | | | 1 | | | | | | | | | | | | | | | | | | | | | | | | | | |
| | 1 | | | | | | | 1 | | | | | | | | | | | | | | | | | | | | | | | | | |
| | 5 | | | | | | | | 2 | | | | | | 1 | | | 2 | | | | | | | | | | | | | | | |
| | 1 | | | | | | | | | 1 | | | | | | | | | | | | | | | | | | | | | | | |
| | 1 | | | | | | | | | | 1 | | | | | | | | | | | | | | | | | | | | | | |
| | 1 | | | | | | | | | | 1 | | | | | | | | | | | | | | | | | | | | | | |
| | 1 | | | | | | | | | | | 1 | | | | | | | | | | | | | | | | | | | | | |
| | 1 | | | | | | | | | | | | 1 | | | | | | | | | | | | | | | | | | | | |
| | 2 | | | | | | | | | | | | | 1 | | | | | | 1 | | | | | | | | | | | | | |
| | 3 | | | | | | | | | | | | | | | 1 | 2 | | | | | | | | | | | | | | | | |
| | 4 | | | | | | | | | | | | | | | | | 2 | | | 1 | 1 | | | | | | | | | | | |
| | 1 | | | | | | | | | | | | | | | | | | 1 | | | | | | | | | | | | | | |
| | 1 | | | | | | | | | | | | | | | | | | | 1 | | | | | | | | | | | | | |
| | 1 | | | | | | | | | | | | | | | | | | | | 1 | | | | | | | | | | | | |
| | 3 | | | | | | | | | | | | | | | | | | | | | | 1 | | | 1 | | | | | 1 | | |
| | 1 | | | | | | | | | | | | | | | | | | | | | | | 1 | | | | | | | | | |
| | 1 | | | | | | | | | | | | | | | | | | | | | | | 1 | | | | | | | | | |
| | 2 | | | | | | | | | | | | | | | | | | | | | | | | 1 | | 1 | | | | | | |
| | 1 | | | | | | | | | | | | | | | | | | | | | | | | | | | 1 | | | | | |
| | 2 | | | | | | | | | | | | | | | | | | | | | | | | | | | 1 | 1 | | | | |
| | 1 | | | | | | | | | | | | | | | | | | | | | | | | | | | | | 1 | | | |
| | 1 | | | | | | | | | | | | | | | | | | | | | | | | | | | | | | | 1 | |
| | 1 | | | | | | | | | | | | | | | | | | | | | | | | | | | | | | | | 1 |
| | 1 | | | | | | | | | | | | | | | | | | | | | | | | | | | | | | | | 1 |
| Total | 50 | 1 | 1 | 3 | 4 | 2 | 2 | 1 | 2 | 1 | 2 | 1 | 1 | 1 | 1 | 1 | 2 | 4 | 1 | 2 | 2 | 1 | 1 | 2 | 1 | 1 | 1 | 2 | 1 | 1 | 1 | 1 | 2 |

**Figure S16 (continued). Domain and motif organization in HRS subfamilies.**
The proteins were grouped in each subfamily according to the similarity/divergence in the protein domain and motif organization. The phylogenetic reconstruction analysis (Figure 3) was used for the classification of PSR TFs to different subfamilies. The protein domains and motifs are arranged according to their relative position as per the CDD v3.19 and MEME v5.3.3 analyses.

| HRS subfamily IV | No. | Atr | Nnu | Aco | Sly | Egu | Vvi | Tca | Cpa | Ath | Bra | Gma | Mtr | Ptr | Mes | Rco | Spo | Zmar | Zma | Sbi | Sit | Phe | Bdi | Osa | Mac | Pea | Aof |
|---|---|---|---|---|---|---|---|---|---|---|---|---|---|---|---|---|---|---|---|---|---|---|---|---|---|---|---|
| | 1 | 1 | | | | | | | | | | | | | | | | | | | | | | | | | |
| | 1 | | | | | | | | | | | | | | | | | 1 | | | | | | | | | |
| | 1 | | | | | | | | | | | | | | | | | 1 | | | | | | | | | |
| | 1 | | | | | | | | | | | | | | | | | | | | | | | | | | 1 |
| | 2 | | | | | | | | | | | | | | | | | | | | | 1 | | | | 1 | |
| | 13 | | 2 | | | | 1 | 1 | | | | 2 | 1 | 2 | 2 | 1 | | | | | | | | | 1 | | |
| | 1 | | | | | | | | | | | | | | | | | | | | | | | | 1 | | |
| | 1 | | | | | | | | | | | | | | | | | | | | | | | | 1 | | |
| | 1 | | | | | | | | | | | | | | | | | | | | | | | | 1 | | |
| | 1 | | | | | | | | | | | | | | | | | | | | | | | | 1 | | |
| | 1 | | | | | | | | | | | | | | | | | | | | | | | | 1 | | |
| | 1 | | | | | | | | | | | | | | | | | | | | | | | | 1 | | |
| | 1 | | | | | | | | | | | | | | | | | | | | | | | | 1 | | |
| | 1 | | | | | | | | | | | | | | | | | | | | | | | | 1 | | |
| | 1 | | | | | | | | | | | | | | | | | | | | | | | | 1 | | |
| | 1 | | | | | | | | | | | | | | | | 1 | | | | | | | | | | |
| | 1 | | | | | | | | | | | | | | | | 1 | | | | | | | | | | |
| | 3 | | | | | | | | | | | | | | | | | | | | | 2 | | 1 | | | |
| | 1 | | | | | | | | | | | | | | | | | | | | | 1 | | | | | |
| | 1 | | | | | | | | | | | | | | | | | | | 1 | | | | | | | |
| | 1 | | | | | | | | | | | | | | | | | | | | 1 | | | | | | |
| | 1 | | | | | | | | | | | | | | | | | | | | | | 1 | | | | |
| | 2 | | | | | | | | | | | | | | | | | | | | 1 | 1 | | | | | |
| | 2 | | | | | | | | | | | | | | | | | | 2 | | | | | | | | |
| | 2 | | | | | | | | | | | | | | | | | | 1 | 1 | | | | | | | |
| | 2 | | | | | | | | | | | | | | | | | | | | | | 1 | 1 | | | |
| | 7 | | | 1 | | | | | | | | | | | | | | | 1 | 1 | 1 | 2 | | 1 | | | |
| | 1 | | | | | | | | | | | | | | | | | | | | | | 1 | | | | |
| | 2 | | | | | 1 | 1 | | | | | | | | | | | | | | | | | | | | |
| | 7 | | | | | | | 1 | | | 1 | | | 2 | 2 | 1 | | | | | | | | | | | |
| | 1 | | | | | | | | | | | 1 | | | | | | | | | | | | | | | |
| | 2 | | | | | | | | | | | 1 | 1 | | | | | | | | | | | | | | |
| | 1 | | | | | | | | | | | 1 | | | | | | | | | | | | | | | |
| | 2 | | | | 1 | | | | 1 | | | | | | | | | | | | | | | | | | |
| | 1 | | | | | 1 | | | | | | | | | | | | | | | | | | | | | |
| | 3 | | | | | 1 | | | | 1 | 1 | | | | | | | | | | | | | | | | |
| | 1 | | | | | | | | | | 1 | | | | | | | | | | | | | | | | |
| | 3 | | | | | | | 1 | | | | | | | 1 | 1 | | | | | | | | | | | |
| | 1 | | | | | | | 1 | | | | | | | | | | | | | | | | | | | |
| | 1 | | | | | 1 | | | | | | | | | | | | | | | | | | | | | |
| | 2 | | | | 1 | | | | 1 | | | | | | | | | | | | | | | | | | |
| | 2 | | | | | | | | | | | 2 | | | | | | | | | | | | | | | |
| Total | 82 | 1 | 2 | 1 | 2 | 4 | 2 | 4 | 2 | 1 | 3 | 7 | 2 | 4 | 5 | 3 | 2 | 2 | 4 | 3 | 3 | 7 | 3 | 3 | 10 | 1 | 1 |

| Colour code | Domain/motif |
|---|---|
| | MYB SHLQ(K/M)(Y/F) |
| | HRS1 |
| | HRS motif 1 |
| | HRS motif 2 |
| | HRS motif 3 |
| | HRS motif 4 |
| | HRS motif 5 |
| | HRS motif 6 |
| | HRS motif 7 |
| | HRS motif 8 |
| | HRS motif 9 |
| | HRS motif 10 |
| | HRS motif 11 |
| | HRS motif 12 |
| | HRS motif 13 |
| | HRS motif 14 |
| | HRS motif 15 |

**Figure S16 (continued). Domain and motif organization in HRS subfamilies.**
The proteins were grouped in each subfamily according to the similarity/divergence in the protein domain and motif organization. The phylogenetic reconstruction analysis (Figure 3) was used for the classification of PSR TFs to different subfamilies. The protein domains and motifs are arranged according to their relative position as per the CDD v3.19 and MEME v5.3.3 analyses.

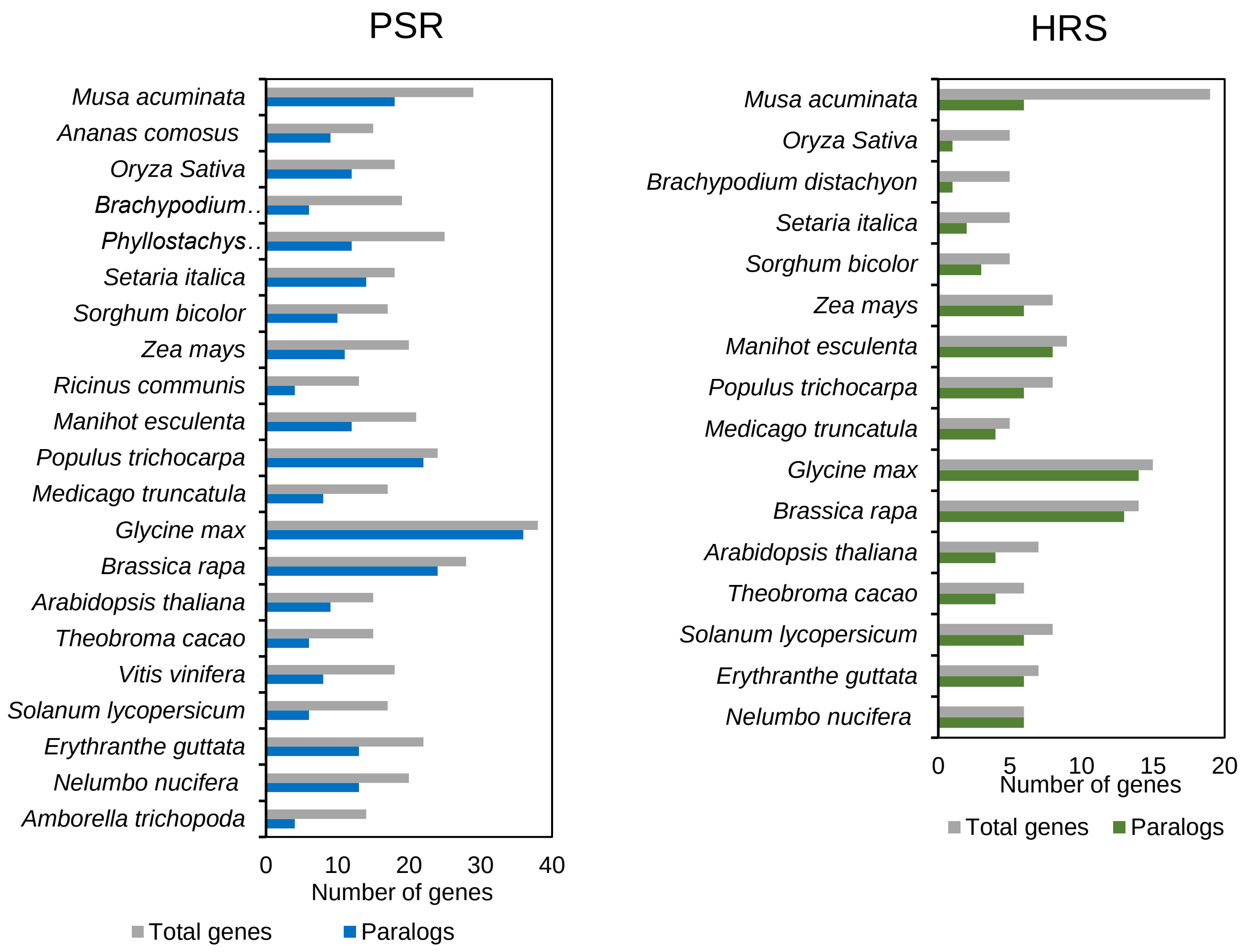


**Figure S17. Identification of paralogs in PSR and HRS TF families.**
The data of i-ADHoRe-based classification of paralogs was used for this analysis. In each species, the number of total genes and paralogous genes are represented.

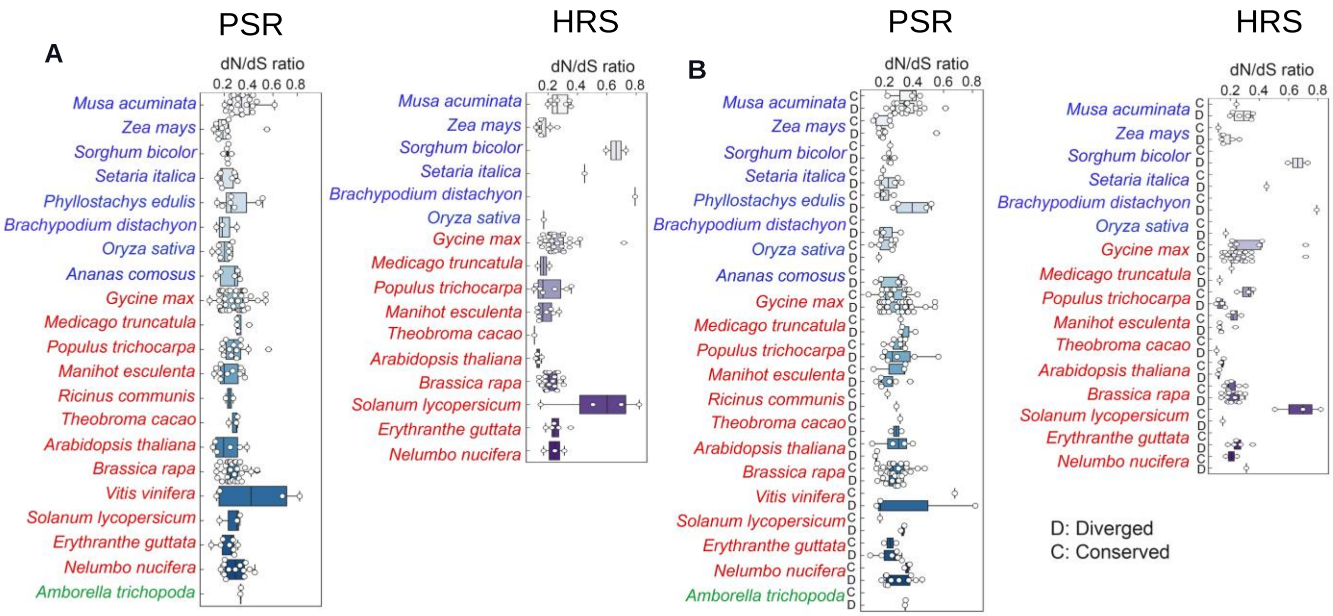


**Figure S18. Analysis of selection pressure in PSR and HRS TF families.**
(A) The dN/dS ratio between paralogs in PSR and HRS TF families in different angiosperms. (B) The dN/dS ratio between paralogs showing motif conservation and divergence.

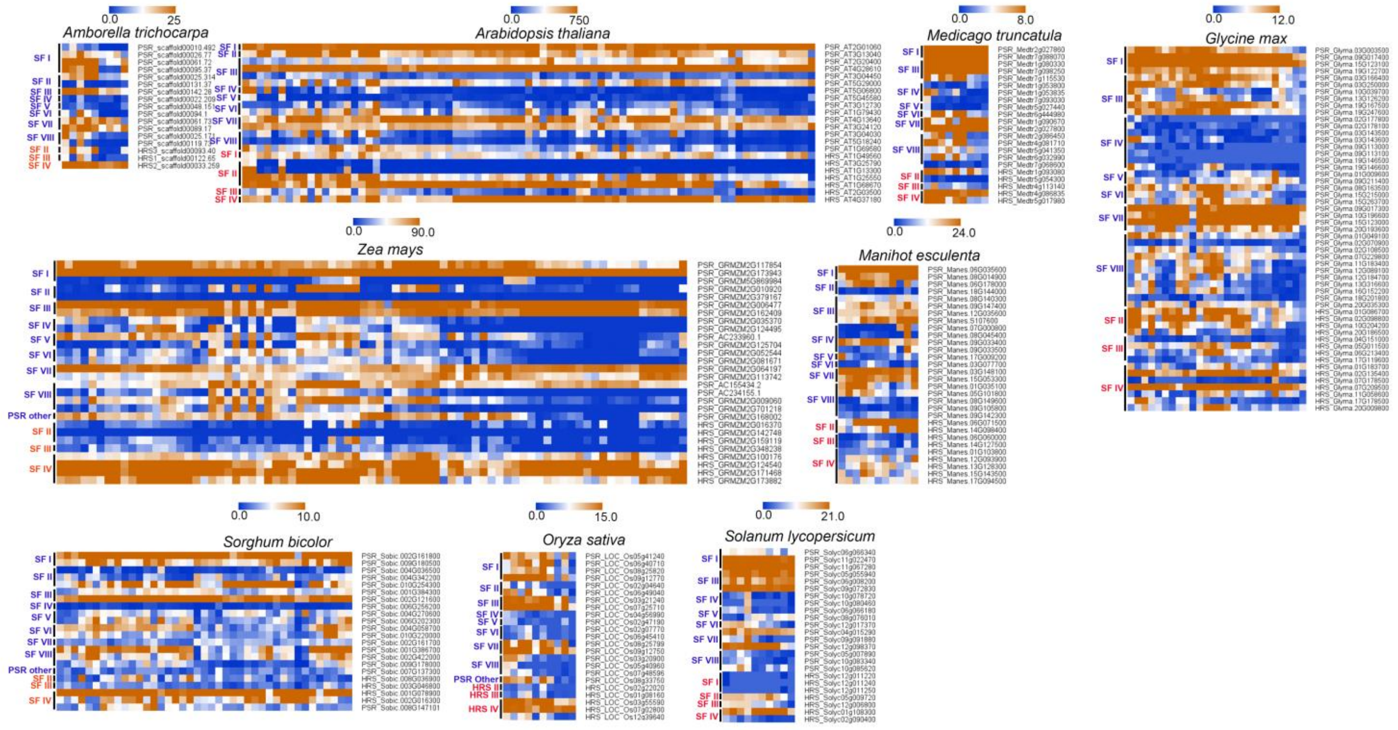


**Figure S19. Tissue and developmental stage-specific expression pattern of PSR and HRS genes.** The heatmap showing the expression pattern of members of PSR and HRS TFs in different developmental stages and tissues. The genes are arranged according to their evolutionary relation (belonging to same SF).

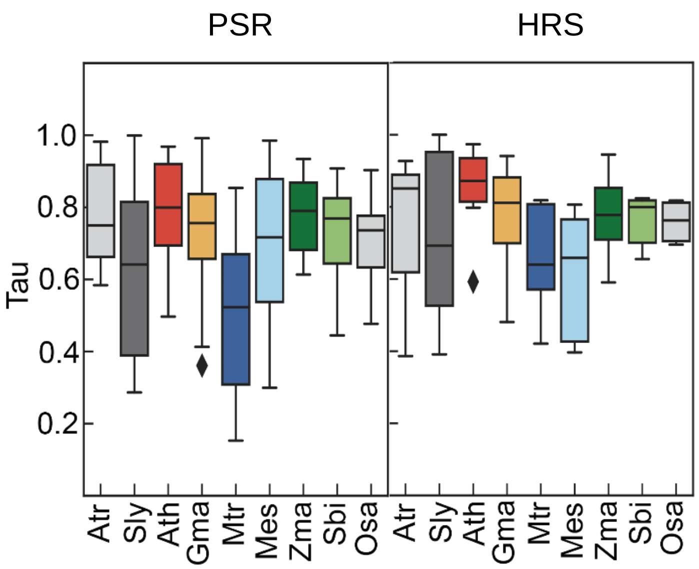


**Figure S20. Analysis of tissue and developmental stage-specificity in expression of PSR and HRS TFs in different angiosperms.**
The Tau (τ) method was used to analyze the tissue and developmental stage-specificity of individual PSR and HRS genes.
