## Supplementary Dataset 1 for "Stepwise origin and evolution of a transcriptional activator and repressor system integrating nutrient signaling in plants"

Supplementary Dataset 1. Protein sequence of PSR and HRS TFs used in this study

**PSR sequences**

>PcoPSR1_20852_PRCOL_00004447-RA
MQQVTRGVAVAASVGTGAHGSQHPSTAAPALSYEGQQTQAAVASATAAGGHLSPEQVQHLSALQWQMMRANSQ
DARSPYHGASQALHSPYSAGILSAQGAPQGASPVAGAAVDGVPPGMYMPPPQRPGEAGAHQPGMPAQLAQGGA
SSLAAPQPISGVAVAPGAAHQDAGAGRGTRPGPTAQKKRMRWTPELHEKFVEVVGKLGGPDSATPKVILTMMG
AEGLTIFHVKSHLQKYRMSAQANGGAGGGKASKGGQGGAAAKATKASGGRSGGASRSGAAKDRTGSRASGATK
RGSRRAAAAADKQASTPPEAAAAAAAAAAATATIASAAPAATSAPESGDALLQADLAATDEGILASAPAPTVA
DDVSVEEALKQQMEMQKRLHEQLEAQRQRTLTLLQRVMEEKRNKVAASSGGAVDVAAAAAAAVEDMDPATAEL
LAATGMGKYVEELKDR
>ClePSR_94875_g8973.t1
MNPYSSDNQDNALNSFFPPDSNLQYWPDYSIDDDWPEEMSGEPVNPEHKTEPLIPTDQQSVPIDSGSERNNSN
FFTEFRK
GDNNPEQNRTGFGLLSNEIFHPYQMELAQGIPVMQNLPYLAMPQTYDLHPSVGTRPIPPVHQQPRNVGPSTQS
NKKRLRW
TPELHMRFVDAVVQLGGARKATPKGILRLMNIEELTIFHIKSHLQKYRMTYDISGPEMSRRGSAESVTREDTV
QSQSAAS
NPQPLSLSLMSETSSQLAVRKRKLDSRIVDADDIMTPVLELDTFEHSEGCRQQLEKALLQQLQAQKELHNQIE
LQRKLQR
SMEAHAQYISTLAKQAGLHEKFPELASQLSDSYEIGTTQEDQKKAIKVEKMTEALENDDNTSIPVASAASTAA
ASAIPSP
STHASESKET
>UmuPSR_362_UM001_0368.1
MGDDVRDLDNLASFWLEEPMPDVLDILQHSETEPESRIPLKSEPDQRREAQQRRRRSASRYQSLAEPDDDLED
EFRTAAA
HNDADVQESKSQSGGKHSRQGAGGSSSKGESSSKPARLRWTVELHKNFTVAVNSLGGPDKATPKGILKLMNTP
GLHIYHI
KSHLQKYRTSVKAGIMDQSTGVMTMPFNVAASVAAPMAPIDLLATDLASGIPNLAARAQTDTKPPSTDAAAPR
NPRPLTP
SQAAGSAPEQGLQDPAGHPRSKRQRTDHDIVKRSDRCGRSDATLRHELLPSDVSDMFGDDDDIMGAMGFSGAI
TQDGASD
ACPVSEPHVDSPHSRHFCDAQLATMNPAVDPGHLQYMGVAEAAHTDHADDMHGEHGTAYVAREVSMEPTMAGV
AMHADSM
QPGLGQSAVRVVGLVAARPPGAAMAAGAGPQFWAAEGYVTAAGYVAAPMAMAGRHAAGPGPGGALRPASPPSL
INSDLWK
TALHKHHQLQASLQEQLDLQRQLQDNLEAHKGYMREAVIGKPRDSAELTRSRSLTRTAAPDALTFAEAARMPM
PLTLHPA
EAVDAARLNSHMHAPAVSAAVAPGVAHELDPLMQQHLPGDDLDAGPEGIGGGAHWDDNGELPDGKFEDDLADL
LDFDGAG
GLGDSPFDGVHDHG
>CatPSR_Chrsp290S04350
MPNNCLEGERASTAAVRKHATATVDEEEENGVKVGSCPSVGISNKRERESGGGESLDDLIANTEWRTWGEFAP
EDEAITTVWTDLLSVDEPVMQHNHGQLGSPGTSTNWPDTRVDSLFLSRQYGASPSHGSVLGGPGGLGAMGGVA
LPNGVSRPQPPAAGTLGASAAQATAATKQRLRWTPELHERFVSAVQNLGGADRATPKGVLRVMGVQGLTIYHV
KSHLQKYRMAKYLPEKQAGVATSSGGAADVTEERSNRPARSRSMKSLVEEGESSAPSQIPKSAVRAPRPANRA
AEAVNTTEAQVVVRPLPEAVAPAASSHAAALLPPAASLDISQLDPADMQFAMPEHLRTGAGQNAGIEEALRMQ
MEVQKRLHEQLEVQRQIQLRIEAQGKYLQRIIEEQRRAGQQAAASQAQQSHQQHSPHHSGQSMPSNSNSEQML
HEHATAAAALQTDEEAKQLEGLLAGEQGNLAQFESFLGRTAQSGPGLFATMLQDQGDILQGHSQLSQPTVGSP
LQSNTLDNRPNSSELDASGLASTWMWNSGQQPTHDS
>MviPSR1_Mesvir1_525_Mv11387-RA.1
MNRPPVGSASRTADQQLSSSAEVQPRTVQKLPTTVDELINQEWPIWGELAPNDDSITTCWTDLLTGPPPKNQD
MHRPQHA

TIQDDTSPGLYLARQQYLPGMGTLPPGGVPPLCAPPGLMDGGGMNLVPGMQASMAAAQSQQPPKQRLRWTPEL
HDRFVNA
VQNLGGADRATPKGVLRVMGVQGLTIYHVKSHLQKYRLAKFLPEEGGNSSKSLGGSKRDTDSDNDDASDGDPL
KMADLKA
GATELLTGEDGSVNIEEALRMQMEVQKRLHEQLELQRALQLKIEAQGRYLQQIMEEQRNAALARRAQAGGAAS
GATTQGQ
ATSAGQAASASSSRSSAGGGGKGPEGAAAPAAGEGGAGADSISHAADGARAGAEAGDAQRQAVASPSGAPVFA
ASGVHDA
DGAGATCPAVGAAGGHAPSPALVPKTESVACGGSLAMPDALASLPGGGGHHLGASGKLPGCELPLPSWSEPGA
ALLTANG
GILPFPSKVEGRNLPQLSLPSHLLLGVEDVDDGGGGGGGGGGGGGQSLQSGVGVGSKRAYDEMMGGGIAMEDGA
GDRLHTD
GSGLPTGGSLLPDDASLLAAQGGHASGGADPAPHML
>MenPSR_537_ME000104S10628
MSKLKAEPWQHVAPLESPRLAQSVEDTAVDGSAHPPAPPSTKTEPVHVSDALASIEDIGGTAISAELPPPLAP
PAPLPVV
PPPSPPRPASSDSILRKQDTDQGDLSTSFLESVDVDDLPPDDAADDGGLSSLLVAPEWPAWAEYAPEEEALGT
AWTEFLQ
VEEADADIAAKTKQVGQGTLMRNQEWHVGEPAGMHNTPPTSAPTPLPGGGSPGMSSQAAGHSSSSAKQRLRWT
PELHERF
IDAVTQLGGPDRATPKGVLRVMSVQGLTIYHVKSHLQKYRLAKYIPTPTASSSREQRAAATPTSSGPKVEKSK
KAPVDLV
AVPDSSAGYAHTRTCTTLLHMQAHMHTPPNPPLPPADAPLFARGLAEKRCTLIRARGAGAGMRAGTVGGVGVK
VQRTLQL
RIEAQGKYLQKIIEEQQKAGAVLASGSASAGGLTSSEAATSMGPSKDAAAGGAAAAAAGGGGGAFGAAQGTLP
SFAAPAE
PTKSGGERGRQSQVQAQVGSLTSSSSMSMRTTSMPTPGLAIPPPSAMPRTPSNGPQDMTGSLGPPSKRSRQDP
HGGMPSL
LHNNHQQQQPQPGGAYPGSAYQPSMYASSNHGYHPQQQQHPQMSMQPYPPHSGVASPVPAPPQPITNSLPFD
AEDMTLR
LGNVNSACRRYALAGPVLIDQEPLGAHPGQFGDPLQGLGGGASYTDSRWTPSSSTFTMMGNHQSFEQDTYQDQ
SSLFSME
GEMGE
>OluPSR1_26932
MADERFWPLFEGLAGAGDGALDGAVDDTLDWLDDVIVRGATSGRVEDEDALGEDAAEDDD
DARDGTRANDGRGGGRKRARASGDARASTSGQEKRAKANAGRGKEGNATRGRNGRGGARL
RWTPELHREFINAVNQLGGLELATPKGIMHIMAMSGMTIQHIKSHLQKYRLQEGAGGSRG
AELDAEADRERRAMIKRARVQQQAEMKQRASEADLMGLDVASAAPMGSGLSSDPPSTPVT
TAQLSELLNSTSAMKAKTLGSEGKIHALDAILAQALKPAATLSDEAAAAVVSAELIEDMP
HVGHALLKQLEMQKQLHDQLIAQRRLQTAIEEHGKYLASILAQEVSGKTKPPEAALGDDA
VDGA
>BprPSR1_Bathy07g01790
MDLREVFRERHFAAQKRALQREKRERERERKRRERERARERTLFTDGSFSPLLFLSAITA
RNYPPHGRRNIPFSRVRGKQKMFSLGNLHHEDEEEEEEPPTPMNASLWPSSNTNNNNNNK
IGSRENAKERSGAKKNNQNQSNNNGMSTFDFDVDVPGDLDWLDGFLRVEDDEDEEENENQ
NADQAWGVGKGAAWKTEAIKTTGGKSGVKRGRDVATASSAEKKKKKNAASNDDDESGENN
NNNNNNGKSSTSNTNNNNNNNNNNADSDRCRLRWTPELHARFLRSVKTLGGLDIATPKGV
VELMRVQGVTIQHVKSHLQKYRLQEQQMSKATSNARSKALSIGERSFLTSTFVGLPPIIA
NEEKVIAKVASIMNNQRPGATTSPPNATANKNTSSSPEGTENNTGNRNSTPGGDNSDGTI
MLGVPPLDEEEKRRHIKAARMAVSAKRKRDKGLRFPDEDVNTSAELAKLAAVNKKLILTN
DVPRDDNFENNNDTDDEDVFTAALPHISSPLKAPMMMKSALGGVGGGAKLPEDVSAVLLR
QIEMQKQLHEQLLKQRKLQTAIEEHGKYLQKIMEESATEKTSKS
>CrePSR1_Cre12.g495100

MDKAERAAGGPNAASEDDWLLEFWPEPAADFPAPVAPMLSQHQDAAQLPEAMPQQQGLALGGYGLTQQPSDFM
QTGMPGFDAFSSGKAATLGLPLLADPQRASTDGASALMNAAQQSSEYMLAPGMGGMPHLLAPSVGTALPGTGH
TGFADLSMGGMAGGIPGLGGPGIMHGQYFMQPQRAATGPAKSRLRWTPELHNRFVNAVNSLGGPDKATPKGIL
KLMGVDGLTIYHIKSHLQKYRLNIRLPGESGLAGDSADGSDGERSDGEGGVRRATSLERADTMSGMAGGAAAA
LGRAGGTPGGALISPGLAGGTSSTGGMAAGGGGGGGLVTEPSISRGTVLNAAGAVATAAPAAAAPAGGSAAVK
RPAGTSLSSGSTASATRRNLEEALLFQMELQKKLHEQLETQRQLQLSLEAHGRYIASLMEQEGLTSRLPELSG
GAPAAAPVAAGGAAGGMIAPPPPQQQLQHQPQLLQPQGSLPAGGSSEAHAAAGAGTMVVHQQQQQHVHHHHQQ
QQVQMQQHARHCDTCGAGGAGGAPSGGSSMQQLQAAEQQRTELVVAGRLGSMPAPASSSPLAGQAHQQQPLAG
GAAHLVHVHSHTPGGQPHVQHQDAFAGAATAAAHASPGLPQSHSHLLPADLSSNAGPDTSAGQIKPEPDMSQQ
QQQQEQQEAEQLAQGLLNDSSAGAGAVSGSDGGGLGDFDFGDFGDLDGGAQGGLLGPGDLIGIAELEAAAAHE
QQQEQEHDPLDADRAKRQRVEP
>CvaPSR1_00058
MGDTPKRSEPLENEAHSPLSLEFWPDAAPLNLDAMNNSLDWLGDLGHDQGGQGGAPPALP
PGDPLGGGAADTLPHGGGGSSSQLQHQQLLHSQLQSLHPMLAMGPAPGGDPLICGGSNGA
SGGDIYGGSGDLSGTLHLHLHPPMQPGGLPGGGDPALAHLPPLGPTGVAAPGGPLHAAGL
AFAHDLVGGHHQHLYQQPPMMHHQQQAFLSPYPGERSRPAGRAPTRPRRPAAARGVTLSA
AQSQKSRLRWTPDLHGRFVGAVNELGGPDRATPKGILKLMGVEGLTIYHIKSHLQKYRLN
IKLPGDQQEATEGERARRGRRRASRSKSESPADEDEEEGEEEEGGSRRESREQAPADSG
GGGGGAVSGGSGGSSRRQQLEDALLLQMEMQKKLHEQLEAQRQLQLSLEAHSRYITSLLE
QSDLRGSLPAHLVTGSSSGAQAALQQLQREGGAAAAPAAAPAALASAGSGAPVAAFSAAA
AAQAPSPALPRSEQQQQQQQQQQGGAAARLGKGGGDATAAPTAGTAGTGSLTAFGATATT
LASVSPLAPRGGVQPPSPGSALLDFEMAEAAAAWDAAAAGVGGPLPKLQLDPSAAAAAAA
QEAAPAVLAPAGEPGGAEPKRQRTT
>CvaPSR2_00367
MQHKQQQEGSSPQKRTRLRWSPALHAQFVAVVQQLGGAFQATPKRIQLAMNVPGLTLFHV
KSHLQKYREVTQGGRPAGNASKKRSGGGDAGEAAAVRSEQAAAPEHEHEHEHEQQQQQRD
LSAWDLCGAEDDGQCSPSDPFGDALLPPGWGTQPAPVVDAELAASLAPAGSLNNQQAAGT
ELTQQPPAAAEPRCAVCERAAQHQQLPRALPHHAAPQPATAKPDRSASQLAAPQRSEASG
SQVAASEGADPAAAAAAGVAPAASAPATSRLSILESALRVQMEMQRQLCCSMEAQRGLQM
QLEAHGQYIAGLLRCQARPPGHPTAAAGQATGSAKAGSRASSKLQLGVAGGGGGPPRGPP
PAGLHASLQQHVFQRAELQPQPAQQQQALQAQQAQLAQHPCGACTLLLRSDPPARSAAAP
AALEPAAPLSSGQPRPEPSCAGSRCSCDTSVVVACQQGGSWDAPRRQQQQVGSEPPAGGE
DAFHEAAALPNAGSLDDSLYPADVWDSLLIDSGVLSRLLEGDSDGGLLC
>CsuPSR1_40143
MAVEGLTIYHIKSHLQKYRLNVRLPGESGDMISGPDESEEPSRRKRRSRSHGQASSRRRS
SRQRKRRSRSSDEDSDEDDMEDEDMDDDENFEEGISRARPGTSVSGINGSSPHGGSPRGG
SPRGVEPLEVPDLDPEKQHSLEEALLKQMDMQKRLHEQLEEQRRLQLSLEAHGRYITSLI
QKKGLEGLPPQTKEALDAALVPPQGSGLSTLTHNTAPQWTPSVSEGSSLAQQVGHVMHHS
TAFMLGSASATDPESSLLLDTNMQAAAAVWDPSQAHGLDQSGSKQLYEEPKELYEEPGQL
YEERGGHVKPEEQL
>DsaPSR1_Dusal.0423s00003
MSSAPSDDLTDFLEYWPEPDMNSLLNAFAAPPPAPAAADAAAAAAASAQDNFQHQQATFL
PPPPPHHIPAFPPVDIHASTTSDAHHHSQGGSSGVAAAAAAVAVAGNTAMAGNFFPHALT
DPAASAAQGAPSIVSMEGSNMHSGSGPAFHGAAPLPALPGLPPPASMSIPGLAHQPFFSS
IPQVSGDQGIKSRLRWTPELHGRFVQAVNQLGGPDRATPKGILKLVNSEGLTIYHIKSHL
QKYRLNIKLPAESGGSGSRFLMGMGGGSGSMGGGSTGYMGGSGAGNSGRAASAAPAESLG
DSQEQGAAAGGTSAASTEVGSGFQGAGAGAPPPRMGPSLFSPHAFQHQPLTAQQQQQLLG
GSSWGAVVGGAGEQAGGTGGMLVMGQQGVVGTSSGGAGGVAGGAQPSSSIGAGGSAASGT
QSSMSLEAALLFQMEMQKKLHEQLESQRQLQLSLEAHGRYLATLIEQEARGGGSSQILAQ
HLVGPMGSIARAGSVSTHLALGGPTDSSRGPQVLPTTSCSQIVPTPSAAETMGATADEGL
QEGAQYQQQQPQQQQQQPQLESDPKEHEEQEGPCAPAHTATPQQLPACTSHSQPLSGGVH
VGGGSGEGGVGVRSSERSKRHRRGEDDAGMQQFQLYAQEGVEHLLQTKGVCEGHNLCLHQ
EEVPPVGKKHRS
>MpuPSR1_61323

MTRMSGIEDDSFSYSFLLISGQLRDPTRLTPRPQTSSAPLGAGGGAAAASGMNTSGTFWP
SDLDLDLESSDFLDSLLLGGEMMTTQQQHHHAGDERLGHGPVPLGARAAGEDHDPRVVLA
ERDAGDGLLPRALADERREPVARAVVASNNNKQRLRWTPELHKMFVDAVKRLGGLDLATP
KGIMQLMDVEGMSIQHVKSHLQKYRLQDSGGGASEFRVSPDASASGKRPRSEEDDAGGNG
KDGNNSAGKTRRRPSAAERSAARLRAAEEKAREERDAARSMAAAAAAAAAAAVERQNVEL
ALLTGDAAGARDALEMSSHHAASHVGGAYDDVLDLVAGPGESGDGDGAWGDVLHDHAIVG
AAGDDVGLVDDVGSDPEAAAAMLKQLELQKKLHEHLMSQRRLQQQVEAHGVYLETILDQQ
KRRRGVE
>MnePSR1_ MNEG_8773_XP_013898209
MDNGLNDAELGAWLEWLPQDDFHSTLEPAAGNGDAAQHQGAQMAVQPPSVGHQQQQQQHQ
QQGQGSGPSPQPAQAKTFQMVAMGAMDPLQAQAYALQYPVQHQAGAFDQLLTAMPLAGHS
MASSGTQQQVAMGMAGPQQVAPALYMPSGYSLQGAPSDASGAAAAAAAAAAGMGGNKSRL
RWTPELHASFVRSVAQLGGPDKATPKGILKAMGVDGLTIYHIKSHLQKYRLNARVPGGGG
SAAAGSVDGASDGAGAAGESWEAASAAGGDSPHSTGPGCGAPPTALGMPLAAAAPAAGGQ
AGAAASSALASAEQLLQQQRQVLPPVAQQQQQQQHQQQIQPPPRPPSAGPSTSAAAPTSS
AAAAGAAAVDAATAEQAASVTRRDLEDALALQMDLQKKLHEQLESQRQLQLSLEAHGRYI
ASLMEQEGLGHRLQDIAGITAPGPGTPAAPAAAAGAAFGGCSGGGGERQEVEAAAGAGVG
GGGAPRLPVQQQQQQQQQQQQQQLLMREPELGDLLGGLPLHAPSGFLPAAAGGGRLGAGV
GGGGA
>MnePSR2_ MNEG_0733_XP_013906239
MQQDLLGGPPGPMHHDEQHEMLQQQPQQQQAQPQAQQQQQQQHPQQQHQQHQHLPGKAAF
PPGMGVPGMDHFHGTPYGMQAVPMQPGHFEHLLNAMPVTGHSLSSSFATDNVHMSGAQPT
LYLAAGFNMAGSKPGVSAAGSAGGGGGGGSKTRLRWTPELHSSFVRSVQQLGGPDKATPK
GILKAMNMDGLTIFHIKSHLQKYRLNARVPGASSVDGGSDGCAAGDSAEGNSGSRPASAA
LDGLGSVPVSALTRKNLEDALVLQMELQKKLHEQLELLLLLLLLLLLLLLLMLLLLLLLLL
LLLLLLLLLLLLLLLLLLLLLLLLLLLLRLLSLEPRRSCDGSVACSCSCAEVFISGELY
>VcaPSR1_Vocar.0031s0066
MDKAERASSTAIGTEDDWLLEFWPEPAAADFLGPIPGTMQPQQQHHQQHLNASQLSELLP
QHGSLQMGQFNLNNTTAPNEYLPGLQFDAAYGGKTNALAGLSGLLPEPQRTSTDGASALL
QSSDFMLPMGGVAHPLMQPQNCLAGLPQPAFPDLPPLGGGLGGGSPFLMPGHFMAGQPQR
AAASGPPKSRLRWTPELHNRFVAAVNQLGGPDRATPKGILKLMCLEGLTIYHIKSHLQKY
RLNIRLPGEAMQGDSADTDASDGEGGAAVPSAPLDPSGYGGETQSALGGGGVGGAAAAAA
AAAAAAAAAPAEPTVSISAHGRSSARRVGGGGPAAAGGTSGSSGSAPSNTRRHLEEALLF
QMELQKKLHEQLESQRQLQLSLEAHGRYIASLMEQEGLTGKLPELGTAPQLGGGSAAAAA
AAAVQRRASGGLAPPSSSAHQHQHPTRSQSNGAAVLRADGGNLVGGGAVGQASLIGTGVG
GDASGGGGGGATGGGGAAMSLRPTSANVADLELQLQPSASSAVAVAATAAALTAHQSHAL
LQPHNHHNHLQQHHKQPTQQQSQEQQQQQQRQKHHNNQQQQQQHAMQPQPLSSMAGGSSG
GGGGDLGAGCGLSSRMTGGGVSSALQLQQLGQRMEAVSGVVPSAAPASIPTAAHDPVLQP
QRLSCFGGQQQQQQRLRQVSQTSSDLDDFACGDLDADVELIHEQHQQQGQQQQQQHHHQ
ARQIPGDARRLGIGLRNCLAEEEEEEEGAREEGGGGEGGGEGAGGGGLGGLPDLDFPDFP
DLDSAGLEQQGLMGVPGDLSLNVSDLDASSGGIAAAAATAAGGLAVAAGDGGSLAAAHHA
LLPLEGRQQGQQQKQRLLDGDSEEEQKQQQARAQLQQQQLVSEDEVGDEEEVDRIKRQKL
RHADDC
>GpePSR1_KXZ49820
MAPLLQPGVAGLHQAAFPDLGLGAAGLAPGLLPGHYLAHPQRAAAAGPAKSRLRWTPELH
NRFVSSVNQLGGPDKATPKGILKLMGVDGLTIYHIKSHLQKYRLNIRLPGEATQGDSADT
DASDGEGAVPSASLERTQSAVGCGGGGMGVPAGMGGAGGGGGGGATGGPAEPTVSINAGG
GAAAARAAMSRRPGTASGSSGSAPSATRRNLEEALLFQMELQKKLHEQLESQRQLQLSLE
AHGRYIAILMEQEGLTGKLTELSNPASGAGGGGADAGAGSAAAAAGPSSGGAGGHARRVS
SGLGPAAGTQHQSTQQQQLHQPLEMRRTGSMDQRSAPAAGGRDRGARGGAGSPMLSPSGS
GGGAGLLQPPPPQRTPSVGLARQRSNLAQQAAAAGPGSGSGVTGGSGVSSLQEQQQQHMY
ERTGRLGSLPTRGSLPGPDPGSLPGAGSGLAFADPMHRQGQAMGLPQPQERGLEPAGAVD
LHQAQEQQEQEQLQAPDGGMGALGEFDFGDFTDFTEQQNLLGGGDLLGMPNVFDTAAAAM
QQQQHQQQLQEGDEEGRRRG

>KniPSR2_kfl00262_0120
MDDHLMLTDPTSEEGPEGVLVAPEWPPWSEFNPDDDSLATGWNSLLQVGNENDGGIGGMQ
DQLLSEPALKDDLKPVSDSSEGMLPATHPPEAGPSQPFHHGAQSAPAPLQQQLEQQQQQE
AQNALHNSWPMTGPGPLRGHAGSSLLPGMSGMVSQAAAADSDERANLLWGAPSSKQRLRW
TPELHERFIKAVESLGGPDRATPKRILELMEVKGLEVSHVKSHLQKFRIATRIPELGTYG
MHMGSYQLTALPNSPMPSPSPPLPNPARRPPRSIPLL
>KniPSR1_kfl00030_0250
MDLIGRSKSRGGGGGEEPAQGFANLLHFGTNPATTGGQLQVEFPGLSGPLSNPVLAPTAL
IEQAEATDTQGGRLAKGGSSGSDDAANQRLPPGGRKARVGLPRQHDKAGPELSPDLRGLQ
NLPGLQNLPGLGPPNQYGDFSLGGMTSAVAALLRGSPDFMVEGGLGQGLAGGGTGGTVTP
NWPVWGENPEEDGIGPVWSELLVPPPAGEGENADGTGGSHQRQGGTVASLLGGGRELNLG
SLSRFQLENKPSQSSASLPAHLLIPQRQFRHAPGAHLVQSLPPTSSPLKPPGKEEDSSGG
HQSQGVRHSQLAPMPKQRLRWTPELHERFIKAVTHLGGGDRATPKGVLRMMGVQGLTIYH
IKSHLQKYRLNKYDGPESSHSSQPGPQHEAGTPDSHPPTAEESFKGPPSQDSPRVKPPSK
LVDSASTENPPPGKRPSKGKRKVKEGGELQRQNSGKSSELRRPLLTRKGSLDRPQKNMDV
MQTSSLNLAHQASALKQVQDAQMSDALQMQMEVQKKLHEQLEIQRQLQLRIEAQSKYLQC
ILEAQQKASLMLQQQQAAGGDFSGAREGQPLPPISAAGAFSLALEGRASASLKPFSLPDM
DLQAPMLGEGGQGFADVSSSERGFVPYEVGGAELPPRKRRSKELEATHLPLEEPLPLQPF
SEAVARILEHGPQGAELRQWSERVAAQSTGTTSSLSRTGLETEGLANLEPIANAWEVGGN
GLGDSGAAAALDLADEHAKEFGKVEGDPWPAPFAQNDVDPASQLDRQNARGFQTGGLSGG
EAGTRGGGRGNHVAPNSYSSMMPSELIELNVPPQESEVAWTPENSIIEDGLGGLESDPTF
GHRDNEKDGPDDGGDTQRPTT
>CbrPSR1_CBR_g4445_A0A388KHT5
MPQPESSRVSAADPLASARCADSSPSPPPSCWSPSPSAAAAAAGVASSDPDVGGRPRGGG
EVIGGGGEDGSSRNCGVDGEEFVPAASSAAAAAVSAGDASSPFRCASSPGCGGDRSQSLV
PIPPQHRCLLQTSAPDEAPGPVLGEEMASLADVSDSGHPALPTSSTQTSAAGNDDSASLD
TDVVAHSEWPTWWEFYPEAHALEPVWAELLNVSQMEADDGEEEEQRGVEGGGGATVNNVS
QHHSSNTSVFQESEKAMSGVDGATVTRDKTGGERHRVPGGGRSLIPGMPAGPLSPGAGVA
NGKQRLRWTPELHDRFVDAVDELGGAEKATPKGVLKVMNVQGLTIFHVKSHLQKYRMAKF
MPEQQVMPNDNEDSGAVCSSFGAAGSPGKGDGKVKGKTATELTAPQDPSRKCQTQDLKNH
LNVQKKLHEQLELQRQLQMKVQATGKHLHMILKQYRILCAQTERTQRELARSSNGDAPLP
KGSSDVLRPSGSADGSSSGGQGGSLSAMGGSMMPMNIVEEGDSVNAAMSLPVCCSHSQLS
SQLSPVDEQAGGQFVQASQNDYTSFINSAEPSYSQQNQAIPMLEDDLFTQEQRLPKQCHC
QQHESPPSVYYPTEQQSLRGQSQQQYENSQGGGPRTKAKYHHHNCGYQVPLQALSGKNQA
MGVTQQQQQQQQQQEQQQQQQQQQQQQQQQQQQQQVTENHRECEVVCLPSARAVISSAPG
RCDHVCVSQLKRLKHDNVQGMPSSVPSGQGLGSGLSRRTVYYQQQLTSSMMPTLDNSIPP
LYNGGASHHQQDGSNMQVPVGVEELDEDPSLLAYAGSGSFLGAAEHRICAMDAAATTMGV
GRVEMTGPMGLVPECQTITRGLENHATEGSGAGMGRVCVTCAPGCSLHHHHTQHVSAPIS
SCQTCRTSHIQHPPLHQPKGGVKRKPAPTMAVMNPSQPSSAQCSFPCRQHQSCLQHQQQQ
QQQQQQQQQQQQQLLQPNFGTGSALSTSERGGEGRGQFNRHSVPPVSSAQQRKQQHNQQS
QPQQQAQLRNEPLHDQRPQSRSSAGANLQQPQHVHAAGSYGLGSSPQLMASGITTFYRTH
SRGDDDRPCASPKESACQQFSSGTRSQRMMAQGDEHGHVNCSPASGSEVGELRGELRDFS
GSMDLSESMATKDIREAANDGIGLQRMDCTTGAAAAQSGDYSSPCFGENRWSVGRCADRA
RAPSLRDSSASSNERKFAGIVPVDCGCGAPRSFGRENDLPDRPSSQALVNAEVRYHKARL
A
>MpoPSR_Mapoly0098s0044
MYQMKKYSNTSLVPHRGQTSSQQERSMYGGSLPGDAGVVTSADPKPRLRWTPELHERFVD
AVTQLGGADKATPKSVMRVMGVKGLTLYHLKSHLQKYRLGKQLHKEVNVETIKDQGSSEG
QGPSSGTATDCVITQNPKESLQITEALRLQMEVQKRLHEQLEVQRHLQLRIEAQGKYLQS
ILEKARETLAGHTAASPGLEAAHAELSDLASKVTTENLSPAFSLTGVPSMSMPSLTVAEL
SVRGGEQQCGGVPHQLSSPNPQSRLSDCSSQSYLTSLACSGNPDGNEPSDQKQNTGKLPG

>MpoPSR_Mapoly0003s0147
MHWVTRTESQRPAGPPSDVMHRQKVQNWQPSQFNAPPIPPCGNPRFDHTVPGSSAPFIHQ

HYGGKQPVLGNPQSPPLLQVSAGYMVPTEAPSRQQVLHPQPLATSNPPSHSIHSQSISLV
PPSSSAPHSSLYSLPPQPQSSVNPPSLHPQQQYLDAAHSVHQAGHSKPDVDEEFNDQLQR
FLDSPDGASSDQMAESPCNEAFGATVEDGWPSWSEYVPDDSGIATCWTNFEPGCESGGDP
VVRTVYQSKDYPSLRLPEGGQASDQLQPGALLQPPSSCGPSGNSAASAKQRLRWTPELHE
RFVEAVSNLGGADKATPKGVLRVMGVKGLTIYHVKSHLQKYRLAKYIPDSVEGKTEKKRN
QDIIQQLDQTSGIQLTEALRLQLEVQKRLHEQLEVQRHLQLRIEAQGKYLQKIIEDQSKF
GGVLSYKLVPAGLSDSDPPGTVLLSGSHLSGDIPLPTTLVTLPATEKGQEQTASEPPLKR
SRTDDSNLSSETKVAMPEQVLSVTGSTEPQLLNDNPSSTVKQAVPQVLDTDQSCNPPLLI
SRENDSTASPAQPDTPAQARQQAGTAERQVLPGSPPQGIVVNPSSSQEAGQVASSTQPSI
TSDTGIRPVELMPDGGSIGKSSLVEAEADLIPPISQVSPPNGTVQAE
>MpoPSR_Mapoly0115s0048
MYAAKKFSTSSLVPQRPGQVPDPRGSNAAGGSTNSGGSPVASGGGGGGSAAKQRLRWTPE
LHERFVDAVTQLGGADRATPKGVLRVMGVQGLTIYHVKSHLQKYRLAKYIPESMSDGGKS
EKKKNPADIIPSLDATSGIQITEALRMQMEVQKRLHEQLEVQRHLQLRIEAQGKYLQKII
EEQQRHGGLLNGRAVPSEPGSATYPVPLAEMAPGQTPAGGVKYDSTSVSLPEGPGTSKPS
TESLAPGSSVQATTEAPALETSGPVDQASGLTSQPQETAQFLGYSSADKSGGQIAQPPQK
RIRLDESSVQSQPQSEAANNLSGQAASFPSADSMSVEGVSTPFHSNSQNYNPSAQEFAPL
SAFQSSTGGSFPQQSSTQQSENVQGRAQYQTGFQQQSMQPIDSSVGLVAPPQHSPRPQPP
SQPQLSSTPIRSQSPMRQEARVFSSSGVDNLEESDNGGVPRKPQFGANSLDVGGGQGRNL
PSSQTGVYEQWD
>PpaPSR_Pp3c10_13030V3.1
MYQMKKYVSPGLSSHMHQQIMLLEDRGPSMYASPMSGDLSPTDPKPRLRWTPELHERFVD
AVNQLGGADKATPKSVMRVMSVKGLTLYHLKSHLQKFRLGKHMQRESHEAIKNAAHGSSH
LKGSSSDSKLSPANLPNPQGKYVNVNEALQLQMAAQIRLQEQLEVQKQLQQRIEEQGKYL
QSILEKAKETLADHTSTSPVLKLAHEELTELASKVIDYEIPKPAFESIKLPGLNPPELSQ
KHGSADTQAVITPAAHNLRQQQQSHCIQSGASSQKSFLSNLPADQDDSGDEHQAVTERVS
NSSSQGAFMVEKGVSGSAFERPPASKTSILADHRDTTTTTTTSASTGGYTWPSAAACHGT
FQAAAQSRKVTSSACVKSEGSPLDLNGNNNNNTGGLDLNAFGWEQQQQQQQR
>PpaPSR_Pp3c12_10900V3.1
MFVISQEDLEAKDGKENITRMYQTLMKQYPSVGIVPASGAQSASSNQDMPSMYGASFSSN
GRASSPDPKPRLRWTPELHERFVDAVERLGGADKATPKSVMRVMGVKGLTLYHLKSHLQK
FRLGKQLHPENSGHEGGKGGSSDIQVTSNACSDEPSTSKALNQEGFQISEAIRMQMEAQR
RLQEQLEVQRELQLRIEAQGKYLQTILEKAKEALGRQIGESPGLETVHAKLTELVSKVNI
EPMNMSFPPFTSVEAPTHIADTNISTLPCQEPRISDSSSQKSHVTNVAANPEDSGDASAS
CENHPSIDTLLSPTQGLQSIAE
>PpaPSR_Pp3c12_11150V3.1
MAPILSTPPSHAPASQRDGAVASEDDIACPAHPVFSSSSKKPIPTQSMAEIMLRGRHQTP
PSLSDIHGPSPSFYVAPHGSHEDYDLQHVVDVAIAVPEHGSRQLAQILYDRSFELKAPQW
PRSNGLNSPQFQEPNRPPEWLRPSSATATRNLPVVAGPPNSASTQPKGQTINAPIKAEST
PLRNLAFPPYANPTAESALLSPIPTVPVPAPGLVNVPETTVNVLSATTTLHREPLQVLLA
GCTTSQQSSGSNASSQIAKSPTLTSDWETSNWPDIIDCMDIQNHSDVLRQQNNIPQMTAS
PSLLSIHHLKTDQPQQSTGTASEGSPGPSFETTHVYSAAEAAKARLRWTPELHEKFVAAV
TKLGGPDRATPKSVLRLMGCNDITIYHVKSHLQKYRLIPEMSTAESKCERRRHSQCQGGL
DAASTVKMSQALQMQMEVQQRLHEQLETQRQLQLRIEEQGANLQRMIDAQVIAGQALGIP
SDQIANGEFFARATGCALNPEDSTVFTGVTPPHITSWSSAAGTSSGPSLKRPRVEVTVPP
LVLIPKVLPTNINSNGSASSPNADASSPMSYTDTSGTSNGVMTSRGEASAYLETNAVTTA
IPRHASQPQGCSPKQHGSVPAATMQSSV
>PpaPSR_Pp3c13_23440V3.1
MAMYQMKKYSSLALAPHRHHQVSSYAPTLSTGEMNPVDPKPRLRWTPELHERFVDAVNQL
GGADKATPKSVMRIMGVKDLTLYHLKSHLQKYRLGKQLHRDSSVHEANKDVSHGSSDMLG
TSNGASDRPLTPTSQNSQDTMQITEAIRLQMKVQRRLQEQLEVHKNLQLRIEAQGKYLQT
ILEKAKETLAGHTSASPDLKAAHAELTELASKVIGDPGTFAASTSQQLAGLNPPELTNLA
SNLIQEAYNEVHNVATPTLTRQQSGRTSDTSSQKSHLTAIQEDSGGISGGCEQPSIPGVK
NNSCNQMQNSPGGYNSNNTSRNSDPFSPQGNGAAEMRPSPGTGASLERPTPRRGAIPTLF

TDTQTTTHNQKTYQSNIYMPHTLPELRISPTVKVEGGLDLNNHHLHPATGPNCSGNSIQP
ARGSDLDLNAFGWER
>PpaPSR_Pp3c13_9170V3.1
MPTQMPPLLPTSAARQRSHSNSSSPLASPGMLQSLPPLGAQCPSQLFLPTSVGGVRGIQP
LARDATPPEQPNYYSAVPGVRPSSWETSSPHVITSPSITAQYHPLNISKYTPPHVAAPTL
RNIHHGLAFVTLQEGSFPSFDIPSFHNLSATYAQQWPSTNGQWVVTESSQSAVPWPSSTI
SMGASSSSTGGSTAAAHAQLSALVPTNSCGLNFHCLRGGVTAATTDEGRGFSIIGRDDGT
GSGMRMELQEIGSGRGGDSTSLEQYANSRLGITPEINAANLCHPLQVLLEGASDQRILRD
SEGSSQTAEASVNIPEWPEPNWMEIFTDPYDESVPLDTINGRVFQAPANLPAPPPLIPAR
DLRTDWQHSGAVSEGSPGTSGTPTSVGASVGASVGVSAIASTTASTDEAASVKTRLRWTP
ELHEKFVDAVAQLGGPERATPKAVLRIMSVEGITIYHVKSHLQKYRLIPETSEDARNDRK
RNDSSFGGMDINSSVFRNFHISRRISHGFYFSTVWQYEKRLRCCDSSAWSLQMTQALQMQ
MEVQKRLHEQLEIQRELQLRIEAQGQSLKMMFEAQTKASGAFGSHPDLSSMEHSEGGSDA
PNSQDAPAAQSSHQVPETLSSPQDASGSSASQEPSTKRARVDVVPNPNLRMLHSKDTTTA
ATPKTELLHSCSAPPPPPPPDRTASCESSPQGSSPSRLAEGLSSSSLADGTPQTMAPMTT
DQGAGLTPQRHGTTSLQSCMPQQSGSMTLTSLQASVL
>PpaPSR_Pp3c14_26270V3.1
MYQNKKFSTTSLMPQRSASQAPEPRGSSAAGASTNSAGSPGGGSGGGGSAAKQRLRWTPE
LHDRFVDAVTQLGGPDRATPKGVLRVMGVQGLTIYHVKSHLQKYRLAKYIPESLSDGGKS
DKKKNQADLLPALDATSGIQITEALRMQMEVQKRLHEQLEVQRHLQLRIEAQGKYLQKII
EEQQRVGALNNGASTTADTTTAPTTTEATQGQTGSVVETKPALPLTNEAPTSNSVPTAPV
SSSSLGLTQATILTSQQPSQTQYGYETFTNPPVLEGSSVQGASKRTRMEDGAGQPVTGQL
ETQQVSGVTQPGGPFMQPGHIQGSCGVFHSNQSDFASGSTFSPRQGGGSFTQAPPLSQQA
YSQQQSLEGVFNTQSFLPANDADTSMPSQVQGTGAQPPPDSSTSGSTRQPPKLSGELDTG
SFLTNDKEATLPTSERGRGDGGPAPLPQVGMYEQWEHVGSGGHLFNQDG
>PpaPSR_Pp3c17_23560V3.1
MYQNKKFSTTSLMPQRSASQALELRGSSAAGASVNSAGSPGGGSGGGGSATKQRLRWTPE
LHDRFVDAVTQLGGPDRATPKGVLRVMGVQGLTIYHVKSHLQKYRLAKYIPESLSDGGKS
DKKNNPTDLLPTLDATSGIQITEALRMQMEVQKRLHEQLEVQRHLQLRIEAQGKYLQKII
EEQQRVGSLNNRPGVTTDSATPTVTDTTQGQVGLVNTKPALPLTSVPQSDTLASNSLPPV
PVSTSNTRLTQATILPPQQPFQTQYGFETFANPPALEGSPVQGASKRTRTEDRGGQSLAG
QSEPQQVSGVPQPGGPFLQPGQTQGSGGGAFHPSQSDFASGATFSPRQGGRSFSQAPLSQ
QAYSQQPLPGVFNSQSFHPANHANIYLPTQAQCTEAQAPPDNSTSGSTRPPQEVSGKLDT
GAFLTNEREVALPASEGDGGHRSAGPLSQAAIYEQWDHVGSDGQLCNEGG
>PpaPSR_Pp3c18_20920V3.2
MFSMKKYMNPGTFQQRHQSSGSLYIPDKLQFPYSSAMNGSDMVPLSPADPKPRLRWTPEL
HERFVDAVTQLGGADKATPKSVMRIMGVKGLTLYHLKSHLQKYRLGKQLNRDQHFHNKDN
AGSSDLQRSNSMSDGSQKSQNHQDGLQMTEAIQLQLEVQQRLQDQLEVQKHLQLRIEAQG
KYLQSILEKAKETLASHTSESPGLEAAHAELTELANKVTTVGMIPLGFSTLGMPLMAQPD
PLMTLHQLPRQPSRNSDTSSQKSFLTNLTANAEDSGGVSGSAEPHGATEDEDGVGQESSP
QFQEEMSANRLRTQGSNNARGPQARLAHAATTTALAGVVSSSSPHCNPVANFWSTSSFQN
MASSPMNGVLSEVVQDCSPKDNNANQLATSLAMERLTSQRVGIHHLSDIPHSSNGLGSIH
SGSTSTSYSSLQDLYCGSAATMHSGLVKAVINPMEQNHFLQCRGSELDLNFGWDSYISS
>PpaPSR_Pp3c19_2940V3.3
MFQMKSKYMNPGISSPGPYQNRAQPSVYAHDKLSPPYGITQTMNGGDMVSLSPADPKPRL
RWTPELHERFVDAVTQLGGADKATPKSVMRVMGVKGLTLYHLKSHLQKYRLGKQLNRDQH
LQNKDGSLQRSNSLSDGMQQLKPQNLQDGMQMSEQLQLQLEVQQRLHDQLEVQRHLQMRI
QAQGKYLQSILEKAKETLASHTMESPSLEAAHAELSELATKVTTLGMFPSGFSNINMPGM
AQPDPLMALHPQPRQPARNSDASPQKSFLNTNAEDNKGVSGSGDPQGASEEMSANCLHPD
GSYNAKSNQSRLAHAAVANSTTSDALAGITMSSSTYCKSVASFWRVSSYQSMLESSSTMF
RVLTEPNDNNANQFANAMVTEMLIQQHLLGSTS
>PpaPSR_Pp3c21_2850V3.1
MFPMKKYMSPGTFQQRPRSTGSLYIPEKLQSPYGSVMNGGDMVPLSPADPKPRLRWTPEL
HERFVDAVTQLGGADKATPKSVMRIMGVKGLTLYHLKSHLQKYRLGKQLTRDQHFHNKDG

NSDLQRSNSLSDGGMAQKSQNMQHGLQMSEAIQLQLEVQQRLQDQLEVQRHLQMRIEAQG
KYLQAILQKAKETLASHTSESPGLEAAHAELTELASKVTTVGYLSDFSNLGMPPMAQPDP
LMALHELPRQPSRNSDTSSQKSFLTNLTGNAEDSGAVSGSGEPRGLTEDEDGVGQESSAQ
FHEEMSANRLHPQGSGNSRGFQGGLAHAATTTALAGVVPSSSSHCNPITSFWHMNSFQNM
TSSPMSGSLCEVVQDYRTDNSNPNQLANSLIMERLSSSQGVGIHHLSGMHHTQNGLGFMH
GGSTSASYTSLQDLYCSSGDTLRGGLVKAEMNLAERNHIIQCRGSELDLNSGWERYISS
>PpaPSR_Pp3c22_8210V3.1
MFQMNSKYMSPGMSSPGPYQQRAQPSVYVHDKLSPSYGSSHGMNGGDMVPLSPADPKPRL
RWTPELHERFVDAVTQLGGADKATPKSVMRIMGVKGLTLYHLKSHLQKYRLGKQLNRDQH
LQNKDGSLQRSNSFSDGVQPLKSQNPQDGLQMTDQIQLQLEVQQRLQDQLEVQRHLQMRI
EAQGKYLQSILEKAKETLASHTNESPSLEAAHAELTKLATKVTTVGNMMPSGFSNLGMAG
MAQPDSLMNPHPLPRQPSRNSDTSSQKSFLTNLTPNAEDSGGITGSGEPQGATEDDDGVG
QESSTQFQEEMSANRLHAQGSSNGRGKPARLVHPAAVSTTSPALMRVNVSSSPHCSSVAS
FWPTSSFQNMPSSSMNIRVLSESNDSNANQLANGMVMERLTSQQRVGIPHPSVDSVAGDH
LGGLLQGGFASLHDMYGQEFERPNATMHGVAHDVKEEHSSLEQNYHGNSYLQCRGSELDL
NHGWERYISS
>PpaPSR_Pp3c22_8217V3.1
MFQMNSKYMSPGMSSPGPYQQRAQPSVYVHDKLSPSYGSSHGMNGGDMVPLSPADPKPRL
RWTPELHERFVDAVTQLGGADKATPKSVMRIMGVKGLTLYHLKSHLQKYRLGKQLNRDQH
LQNKDGSLQRSNSFSDGVQPLKSQNPQDGLQMTDQIQLQLEVQQRLQDQLEVQRHLQMRI
EAQGKYLQSILEKAKETLASHTNESPSLEAAHAELTKLATKVTTVGNMMPSGFSNLGMAG
MAQPDSLMNPHPLPRQPSRNSDTSSQKSFLTNLTPNAEDSGGITGSGEPQGATEDDDGVG
QESSTQFQEEMSANRLHAQGSSNGRGKPARLVHPAAVSTTSPALMRVNVSSSPHCSSVAS
FWPTSSFQNMPSSSMNIRVLSESNDSNANQLANGMVMERLTSQQRVGIPHPSVDSVAGDH
LGGLLQGGFASLHDMYGQEFERPNATMHGVAHDVKEEHSSLEQNYHGNSYLQCRGSELDL
NHGWERYISS
>PpaPSR_Pp3c26_3290V3.1
MVLSPLQIEQTGKCTIAAYNTWLPTFWRSFLSPSFLFLLSHAKLSMPSQMTPSLTTLSAH
QRGGSNMSPRASPGFPQTLASPDTRMPFISQGFAASSSIPGALACGSQPGGEANQHQQLR
FSSTGASGSSSSSWQTQSPYGQYQGTTPSMSTQMSRIHHESGHGNLAGGEQTASSSSFQM
PYNISTSFSQQWRSTSGQWDCTGSPQSGEHWPVGGMTTGVLSGPGGGSRPAHAHLSVLVP
PNRSGISLDSPRGGMHPPGAGDSGGFSMESGPQMPFPKLGSGREARSATQLRQYPSEAGL
LITPKTDGAGAILCDPLQALLDGSSEQRLPHSGECTSSPTGEASSNLPEWEKNWMENFPD
SYNEGRSLDPNNARGFQQPEKFYLVAPPQPIAMRDPGTEQQQPGTASGSSPVPPAAFSSA
AEHVKTRLRWTPELHEKFVDAVAQLGGPERATPKAVLRVMGVEGITIYHVKSHLQKYRLI
PEVSSEDSRNDRRRNDSSLSPMDIHSSLQMTQALQMQMEVQKRLHEQLEIQRELQLRIEA
QGQSLKMMLEAQAKASGVFGVRSDRAGKELIPPATDAIADRDFPTQSSPAPESAAAGART
SSPGYEQSTKRARVEVPNLVIVPQEPYRSEAYRHQGNNKMGLLHDRSSSAWNSSSSLQGS
LPSRLGRSSSTDQVEGALAMSAVSDGGGGTPQRHTASQGCMPQQPGNMTSKSLQASV
>PpaPSR_Pp3c26_3820V3.1
MWNSDILQSPEEQAALPPEGVSLPPLPTAEPETDLFGGGGEWLTWNQLMPEEDVIASCWT
QLIDVEQDDGRTLNQSIKYVPLQTPLQVEPLASHPPTSEYPTSSSGAVSCGSPKPSSSAA
AKEASASKSRLRWTPELHEKFVIAVAHLGGADRATPKAVLRLMGVQGITIYHVKSHLQKY
RLAKYMPEISEEAKAERRKHDCLLTSLDLGSGHQIAQALQMQMEVQKKLHEQLEIQRELQ
LRIEAQGLSLQKMLEQQAKLNHPDLPSGEPSAPANVVVPTPSSLAPSNSSNTTTLLEEQP
TGSGLVTHTPSYTDTTMERSKQKQIEAGSTSPTSLDGHAAKRARTDASPQVNLSGQEHNF
LAPSTKPSCTWESFSHQSPQFQRLTRTGQDPQYLQTPTSGGSPQQAHADSIPQHPGRAAQ
KPVQA
>PpaPSR_Pp3c2_310V3.1
MYQNKKFSTTSLMPQRSASQAPEPRGSSATGASTNSASSPGGGSGGGGSAAKQRLRWTPE
LHDRFVDAVTQLGGPDRATPKGVLRVMGVQGLTIYHVKSHLQKYRLAKYIPESSSDGGKS
EKKNPADVLPTLDATSGIQITEALRMQMEVQKRLHEQLEVQRHLQLRIEAQGKYLQKIIE
EQQRIGSITNLQGTTETGAPAAEEANQRQVVSVVDAKPKLPLAPVTTSETPTSNSAATAP
VSSSGLGLPQAAILLSQQLSENQYAYGTFMNPPLPEGSPVQGPSKRIRIEDGAGQPQTGQ

LATQQGPGIPQPGQTQGCVSGAFHPTQSDFQSGATFSPRQNGGSYTQDPLNQQSFPQLPP
QAVFGNQAFLPANNTSTPVNAQVQGTGAQATSDRCTSGSTRQHPEVSGELDTGAFLNNDR
KASLLPSDGDRGDRSSAPLPQAGMYEQWDHVNRGGQLFHENG
>PpaPSR_Pp3c3_10940V3.1
MPSQMPPLLPSLAAHQQGYDSSGSPRASPGVSQAFPPAGAQFASQPFLPSSTAALGGIQY
QVCGANSTQQRSYYSVDTGGGSPSSWQSPSPYAVHPPSTTPEYNPPTMSRHTSSHVAPSS
QHNIHHGLPYGSVQDRSSPSFQIPSFHNLSATFSQQWSSSNSQWVGAESPQSDVHWPSDV
MSMGVSSPSSGGSRPAALAQLSALVPTSDSCLNFHSPRRGVVTAPPGDGGGFRMVGANGV
IESDLQMGYQDTASGRGGGSAASLGQYPASGLGIPPNTDAISLCDPLHVLLEGASDQNIP
RDGEGSSQTAEASVNISDWTDPEWMENFSDAYNENGPLDTNNRRVFQQPKRFHPPPRLIP
TKDLRTEQQHSGAASEGSPGASRNSAPVGATASSSAAASAAEAASVKTRLRWTPELHDKF
VDAVAQLGGPERATPKAVLRVMGVNGITIYHVKSHLQKYRLIPEASSEDARNDRKRNDNS
LGPMDLTSSLQMTQALQMQMEVQKRLHEQLEIQRELQLRIEAQGQSLKMMLEAQAKASGG
FIPRPELFCNASLPAVASEVPKSQVVPAQPSQASETAPQQSTNGSSPVRETSVKRARVEV
PSMVIVPQEQFREAYNKQDGPKTGLFLHGCSAPPRDFAASCQSPPSLPPPPQLSLPSRLL
LSIKGHSTDQTDGPSSSTSAPPPPPPITDDDGGGGATPQWHDASPQGGCMLQQPGSMTST
TTSLQASV
>PpaPSR_Pp3c3_14670V3.1
MYQMKKYSSVGLVPHRQHQVSPYASAVSCEEMSPTDPKPRLRWTSELHERFVDAVTELGG
ADKATPKSVMRVMGVKGLTLYHLKSHLQKYRLGKQLHRDSRVHEANKDGSHESSDMQGTS
NGTSDGTLTPTSQNPQDNIQIPEAMRLQMEIQCRLHEQLEVQRELQLRIEAQGKYLQTIL
EKAKETLAGHTSTSPHVKAAHDELTELASKVISDPGTFEVTLPQHLAGLNPPQLSNHASD
SMQVVSNRVQDVTSPSLPGLQSGRTSDTSSPKSQLTTFQEDSGGVSGGGEQPTVPGGNSN
TGNQMQLSAGGNSSSNNDEVDDDHGPRLAISATEFGASAGPGVPPERPTPRRGAIPIRFT
DSQKNNNDNRFHSNVIAASFPLQQRMSSTVNAEGALDLNHGDHHHSHTNSSNSNGNSIQP
ERDGDLDLNAVGWER
>PpaPSR_Pp3c3_23200V3.3
MYQIKKYPNPGMNPHRQQQIMSFQDRGPSMYASHMSGDLSPTDPKPRLRWTPELHERFVD
AVNQLGGADKATPKSVMRVMGVKGLTLYHLKSHLQKFRLGKQLQRDSHEANKDATYGSSH
LRGTSSDSKFSPANHQNPQDEYVNVNEALQLQMAAQIRLQEQLEVQKQLQQRIEAQGKYL
QSILEKAKETLADHTSASPVLKEVHEELTTLASKVINYEAPKPAFESIKLPGLNPPELSH
KHGTADSQGDIPHAAHNLQQQQSRVQSGAVSQKSFLANLSTNQEDNGSERQTGTEGGSNC
SSHDAFTVENGVSGSAFERPPPRNAPSPADHRDNNTSSGGYSWPSAASCHGNIMTAPPNR
LVAPQRVKAEASSLDLNNHGSNGNGNATPFQGNKGGLDLNAFGWEQR
>PpaPSR_Pp3c3_34700V3.1
MGDALGHEAGETRGEVALPRTHIEVGYELQSVVTGFANHNLIGNQQEQDSIPPRVDEPLN
YISPTQGFTACHQQTPFGTTHNSHFLPPQYRWQPIEDAGQVQEHQVSPLEQWQPTRTTPT
TRAPRPWNSADLLEGHAVPRQSSAPTPIVPITPFRRRSSRVIRPMPTLRSRGSNRPPASL
EAQRNLFGAVPRCTDLPWPFLYVQPNLDVRPTIPFRLQPAATLDGIPPPPPSATCTPTKA
TPNESNSSQTRFASKWSDRGQMTGSNSPTDSEVPFSWPDLIDYGTPNPVDASAEFTMQSD
SVAPKQLSVSTSAASVPQLVASPQLNQISGAAVSQESPSQPSDDDSAIYNFLAAHAARTR
LRWTDALHDRFVAAVAECGGPDRATPKSVLLAMGCPGITIYHVKSHLQKFRLQSEASTAD
SMRRRPRECFRLDPVVQAQMERHAEVQKLLRQELESQRELQVRIEHQHLQLQRMLEEQLA
RPRRELGVVIEPEAVAAKSQLEEANTAIVPQEIFYSPDEDPYMGISLEDSLPSSDESHIG
EAASNLFIHSLDIEDEISPSLLETTPSGSLNCLV
>PpaPSR_Pp3c3_34900V3.1
MWKKWGLLDSSDGAGVPLLEGSEAAAEAGAVVAMNELEPVPASDVYGAGAEWLQWTQFVP
QDESLAKCWTEIIDVEQDDGPTLLQTTRTAPVSAPPPVVEKRSSTYTGAVSAACPGSPSL
SSGAAPSVSAPGKTRLRWTPELHEKFITAVAHLGGADRATPKAVMGLMGVQGITIYHVKS
HLQKYRLARYMPEITEEQKAERRRTESLLTPLEISSSYQITQALQMQMEVQKKLHEQLEV
QRELQLRIEAQGQSLQKMIEAQAKVGGMLLDKIPDSTAVALAPTPVSIGPAASPLPDTGP
EALPEGSCPETPNPTTSDVLCTNEINKREPIEATTIDNMPIDEPLIKRARIDEGCLQASQ
SAHPKPKVSTTIYSSSQAVQGQRPSHLMDEKSLSDGQITKGTPQRLPQQSHAPCVTQHPG
PVARTPVQA

>PpaPSR_Pp3c4_16540V3.1
MYQVQGKNYTNMELVPANGAQSASSNQDIPSAYGATFSSDGGVSSADPKPRLRWTPELHE
RFVDAVERLGGADKATPKSVMRVMAVKGLTLYHLKSHLQKFRLGKQLHRDSSGHEGAKGG
SADIQVTISACSDGPSTPKPQNQESFQISEAIRMQMEVQRRLQEQLEIQRQLQLRIEAQG
KYLQSILEKAKEALGSHIGASPGLETVHAELTELASKVNSEPMNMCFPPLTLPELPTQSA
DARIGNLPRQESRVSDSSSQKSHVTNGAANPEDSGGASASCENQASIGIQKFET
>PpaPSR_Pp3c4_20290V3.1
MYQMKKYSSLTVVPPQGQQHLMTSPHNQDHRSSSPYGVVLMSAGEVSPVDPKPRLRWTSE
LHERFVDAVTQLGGADKATPKSVMRVMGVKGLTLYHLKSHLQKYRLGKQQSQREASGHEL
PYKDASHGMQGTSTGASDGPPPTPTSQDPQNNAKINEALRLQVEAQRRLQEQLEVQKTLQ
LRIEAHGKYLQTILEKAKETLVSHMTSLAPDLQAAHAELTDLASYASLVPPHLDGVHPSG
PFSHAPDPTHQSGRIPNTSCQKSHSLPHLSAMQDGELRVSSAGIVDQTTAALANQGWSPT
APKHSA
>PpaPSR_Pp3c4_23130V3.4
MITALHQMLFLCLSIVWSSSNATRFVDFPSSLAQRHHHHHAEETLREIECEHRASDPGSF
FGKIRNFFWTFLRASNNRCCPLSTPMWNSDLLQPPEEEAALPQEERPLAAVPMAEPETDL
FGAGREWLIWNQLVPEEDVVASCWTQLIDVEQDDGKTLNHTIKYMPLQTPLQVEPVASRP
PTTEQRPKSSSGATSAESPKPSSSEAGTSGSASKSRLRWTPELHEKFVIAVAHLGGPDRA
TPKAVQRLMGVQGITIYHVKSHLQKYRLAKYMPEISEEAKAERRKHDSLLTSLDLGSSYQ
IAQALQLQMEVQKKLHEQLEIQRELQLRIEAQGQSLQKMLEQQAKLNHPDLPIGMPSSST
NTVVPTPSSLALSDSSNTTTRSEDQHAENSAVSTTLAHSQTTMDKIDQKQTEAGLTSNTL
LTEPAAKRACHETSSQVNLSGQANNIVVPRTKPSYAWESPSPQVQRSSRTMQESQQLQTS
TAAGSPQKVPQQAHAASISQHPGRVAQKPVQA
>PpaPSR_Pp3c4_6870V3.10
MCVVASMAPISSASASHVPASQSCVAGASEECVAPLARFVSRQEDGTSLSLNRAQSISEV
FGGSQVISPKASSGQSFFVAPQGFHEDYDLQYVVNVAMASHEHGSRQQVPILADSASELK
SPEWSHRNEHNNSQFREPNCAPPWQGAPSAIETSNVLAPGSPPSLALTHSQGETLSLPLQ
AETNPLWNCPYTQFAKPASESALRKPSPMMPVAVSDFVSSQEATAKVPAATTNTRLDSFQ
VLLAEASSSQQPDGSNLSQPVKSPILKSEWETSGWPHLIHRREMANPPDMLQEPSIAPQT
VASSPLLSTQHPQAEQSQRSTVATSGGSPGPSIEATPMFSAAEVAKARLRWTPALHEKFV
AAVAKLGGPDRATPKSVLRLMGCNDITIYHVKSHLQKYRLIPETSTAESKCERKRHNHCQ
GGFDVTSTTKMSQALQMQMETQRQLQLRIEEQGANLQRMIIEQVIAGHALGIPSDQITNG
ELLSNAVSQALHPSDSSNLFPGITPSKLSQQPTSWTHTASTSTGPPPKRTRVEVPALVLI
PKIAPKVKYTMPAETSCSHGSAPFANIDPLSQMGNTAIQAAAAPRGDADAMPASRHVNRP
QGCVPMQHDGAAAAPMQTSA

>SfaPSR_Sphfalx0001s0234.1
MLMNRQGMEGWQQPSQVSNSHAAAAASPDPAAPVVVTMPTPQLNPHRAFARPNPPGKTQG
GRGPAWAAMEGSSSSSSSSAAFTQPRHGYYQPPPPLPPQMENQRQVGMHGDSGMSSQYQY
PQSQGPYPAQPGPGTEPSSPHYGQQQQMSAHHSYQSPAHWSPGAQWPNHHQQQQQHPVPL
PPALQQSGGHWQAGNKQLLPSPTGLAGQSDTTAALHSCALDQANGPKTQDNRDGTDFRDQ
LQVLLDSPDGDGDSSQMGEQSVNDGFGAGPEWLKWTELVPQEETIANCWTEIIDVESGEE
RTLFQTTRFEPLAPTTLQPRSLPTDQQPSGSAGGQYPESPGPSSAAAAAAAAVKSRLRWT
PELHEKFVAAVSDLGGADRATPKAVLRIMGVQGITIYHVKSHLQKYRLAKYMPEISEETR
AERRRNDTYLQPMGISSSQQITQALQMQMEVQKRLHEQLEVQRELQLRIEAQGKSLQKMI
EQQGGMVSRCWSEPHDPSSLATVPATPVPETPVRSSIVPVTDSPHQASALESDIPVTDRA
PLPASTVDQPDLKHERLISEPKEDQPPPALITSNEEPSLKKARMDVNPQASSLVQVPVSV
TSDAQRGGGSDTQENVQQEVGLETSPEMVTQKLTSPALEAQEVTEGHQAHSFAQCTPQQG
HTACAPPHPGSIAHKPVHASG
>SfaPSR_Sphfalx0002s0165.1
MRLDSGCGAEAGGGRGRGEGGDCGDGEVVAGLQQSSASMRYRHGYCQPSHMDNQQVGLQG
DSDLPFQYHYPPQRPYLAQPSSEPETSSPHFRHQRMAPLPSYQSPEHWGLDVQRSNRHPQ
GHLHSFPHAHPQQQQGGGHSYVDRQQLLLPSPSAVDRPQQTLARSSSVSSHPGFRVGCEQ
LEKPGGGGSYALGHTGRESHQNFLNVDLEVASQSIGVVAHPNSAPDQPIGPKTEEDKDDA

DFGDQLKILLDSPDRGGTDSSQMGDRSMNDGFGTGAEWLTWTELVPEEETIANCWTEIID
VKSGEERTLFQTTRFEPLAPTPVKLGGSPTDQQPSGSAGKQYPESPGLSSGAAVAAAAAK
TRLRWTPELHEKFLTVVGQLGGADRATPKAVLRLMGVQGITIFHVKSHLQKYRLAKYSPD
ISDEARAEWRRSDTYLSPIGINSSHQITQALQLQMQVQKQLHEQLEIQRELQLRIEAQGK
SLQKMIEQQAKVGGMVLGYHSEPCDPSSSPLFPATPVPEILRSSSKIPITGSEGSPLASI
TEIVISMTDKTPLPESNIDQSNPSHGLLSASKAVSQEQPPMSLNASNEDPLSKRPRTDSK
PQVSSFLHVPVSVASDAQSGGGSAAQGGILQAAGLETSLEMVTQYATCQEVEAHTATEGQ
LAHSFGKCAPQQAHTACAQPNPGSITHKPVRASG
>SfaPSR_Sphfalx0006s0186.1
MYQTTKFSASSVMPERPKSQVVEPWAPSPAGGPTYSAGSPGASGGGGAAGSAAKQRLRWT
PELHERFVDAVTQLGGPDRATPKGVLRVMGVQGLTIYHVKSHLQKYRLAKYIPDSLSDGG
TSDKKKTSADLFPSLDATSGIQITEALRMQMEVQKRLHEQLEVQRNLQLRIEAQGKYLQQ
IIEENQRRTALSGTPVTTNVGEAGPSALEPHLGEATGQLEIKQLPVVVATTQAPGSLVPT
QAPAGLNSHTPQSEFVGVSNLSNPGGAPSQQLTDATHSEDGAAGVKVFQLEGTEVSGTSQ
AVVEFDLPAQGQEVVGSLQPNHSSLPPGTSFSPPPQAEDSHCPDAPSQEGYSKGKLQRFC
SAQTNDHANLV
>SfaPSR_Sphfalx0009s0225.1
MYHHHHHHHTKRFNGVTAAAATNFASSANQTSPAGGGASSGAAAAAAKQRLRWSPELHER
FVDAVSQLGGPDRATPKGVLRLMGVQGLTIYHVKSHLQKYRLAKYVPDSASDGGTSKVDQ
KSNKKKNPAAAAQPDIILPASDSTSGVQVDDALRMQMEVQKRLHEQLEVQRHLQLRIDAQ
SKYLKKIIEEQQRVGGSTNTTTTTTTHDDTKLHTVAAASVNERGLGRSHSLMAAAADDQE
PSSNEEVCIVCQDDDDDKKQQQHDQHQIPDLRLSVVVTPGEEAGIVTDDDAYKLACSKAS
SNEQAGASISPPGIRELHFEIMRSSESTTSQASSARTTTQLQQQQQQQQHQSTRVAQYN
MGTGSLYTASLNSTFTSSWTPSSQSPSKRTRFAGHGEIVATGQQPGSSQSVSSELEPVVQ
QHDGGIMVATCTTSSGGSDFQHQSIHLVQRVVTPGFPLSSPASNCDHSDFKLAMDFPLGG
GSSSSNLQQAPPANLAAALYTSQTSDLHSLTGDDLTLRPLRSSSSSLQGSHIVAMDQQWA
ERPSTTTAESDSASDAVRQMFSRWEENSGNAVAHSQGKT
>SfaPSR_Sphfalx0014s0203.1
MYQTTKLSASSSLMPQRPGGQAVEPWARPPAGAGGPSFSAGSPGVSGGTAAAAKQRLRWT
PELHERFVDAVTQLGGPDRATPKGVLKVMGVQGLTIYHVKSHLQKYRLAKYIPDSLSDGG
AGGISDKKNPADMFPSLEPTSGFQITEALRMQMEVQKRLHEQLEVQRNLQLRIEAQGKYL
QQIFEEQQKLNTINSTPMASDTDEAGAPGFKPPLAETVDKVEIKVPPDVTTQAPGPSGSL
IPTEASGLNSQTPQSEYAKS
>SfaPSR_Sphfalx0016s0215.1
MLMNRQGIEGWQPPQMTNSHAAGPSADVVTMPTPQLNPQLTYTQPNMTKAQGGGGGAAWP
AVEASSRSSSASLQPRHGYYQSPQMDNQQMGMQGDPAAVAMASQYQYLPPQGQYLAQPGP
GAQWPNLHAHGHPHPLPHHPHSQQQGHWQGGSQQLLPSAALAVHNPPPTLARSGVVAGSH
HPLRVGSGGQQQQQENPGGGGVSAAYSLEHTGRESNQAFQDVNLGVAAEGQSGLSAALLH
SAPDQAIGPKTEDRDDTDLGDQLQVLLDSADGDGTDSPQMGEQSMNGVATGAEWLTWTEL
VPDEETIANCWTEIIDVESGDERTLFQTTRFEPLAPTTLQPRGPHADYQAAGSARGQCPE
SPGPSSAAAVKTRLRWTPELHEKFVNAVSQLGGADRATPKAVLRLMGVQGITIYHVKSHL
QKYRLAKYIPEISEEARAERRQNDAYLSPMGINSTHQITQALQLQMEVQKRLHEQLEVQR
ELQLRIEAQGKSLQKMIEQQAKVGGLVLGYRSEPQHPSSSAIVSATPLAESSGLSSRISI
NEGPLPASTIESAIPVTDRAPLPASNVDQPNPKHVLISAQKAVSQEQPPPTLIASNDESL
SKRARTEFNSQLGSSVQVPMSVASDIQRGGGSATQQNARQAGEFEASPGMVTKLASPAVE
AWAAVEGEQAHSFLQLPPQQGHTACAQPHPSSVPHKPVHASG
>SfaPSR_Sphfalx0021s0179.1
MYHQTKKFAIVTHHGTARSASTCNDNIVSPGLESPGGSSPSAAAGGGGGAKQRLRWSPEL
HERFVDAVSQLGGPERATPKGVLRVMGVQGLTIYHVKSHLQKYRLAKYVPESASDGSKLE
KKNIPGASDSVVPASDATSSGVQVVDSDALHMQMEVQKRLHEQLEVQRHLQLRMEAQSKY
LKRIIEEQQQRAAGTQDVAGRHPVVTAASGHEQEERRVSLVKEPSSEAESVDVNERKLTR
LPDLRLSIVEQEVAMPSSWASDKQRATRSSPCNITREHQFAITASGAASSQASALLRPTQ
LQQAAQVQYSSASGYSHAASAFTSSWTLSQPSSKRTRFGSELMSSFSQSLSPELDLRQHG
RGMGSSSNEFQQSIQVQRTPIIPLNPASNRYGFNLAMGFQRSSYIHESAQVLYAPQPLLS

SPGDSLHLQSHGFQRPAATQGAAHQHVFMQPERGMSSFQVAKGEEEGSPLLPAPKSRASD
ALRQMFRRWEENSGGNAGGTSSQGNTS
>SfaPSR_Sphfalx0078s0059.1
MYQMKKYTSPGFMPHRTQTSSSNLQDRLALYASGNLSGDAGAGAGGDPKPRLRWTPELHE
RFVDAVMQLGGSDKATPKTVMRVMGVKGLTLYHLKSHLQKYRLGKQLHREGSVPEAKDGP
HGCSDGQGANVASGQSMVTPKSQNPQESIQITEAIRMQVDFQRRLQEQLEVQRHLQLRIE
AQGKYLQSILEKAKETLANHTAATPGLEAAHVELTELASKVITDPLGPSFPSLGLPNLSS
AHEDNSQGVSLQHQFSRVSDTSSQKSYLTNLTVKPEDSGGGSGSSEHQASTGMRKH
>SfaPSR_Sphfalx0078s0062.1
MYQIKKYTSSPPGGLMPHRPQTSPHLERPSLYAGSLAAHGVNAADPKPRLRWTPELHERF
VDAVAQLGGADKATPKSVMRVMGVKGLTLYHLKSHLQKFRLGKQLQRDTNIHEANKDGPR
GSSDMRLSSNVASDSMATLQPKHPQDGIEITEAIRVQMEVQHRLQQQLEVQRSLQLRIEA
QGKYLQSILEKAKETLAGHASSSPGLEAAHAELTELASKVSNDPLGQSFSWMNLPGINNP
ESLQHAVDTRGGASSLPRQQQLPRQQSRVSDSSSQKSYLTSLTANPEDSGGASGCGEQQA
ATGKKRACSSCTYSEDDDGEGQQMVQLREMNANFPPMQGSSNARAEDRLAKAAAALKGVN
EWKGPPSWSVASQDSLSLSAGIVGAEGAGDKFSPADGSMGRSKETISNSHAFMMKSSEGG
ASALERPTPRRGTIQSLAEERNAGGGSPHTSITGSSLQVTTKYCSNMGAPHRPAQTPAST
NMKVGGALDLNYGNGISMPRGSELDLNTRGWER
>SfaPSR_Sphfalx0088s0085.1
MYQIKKYTAANPAGSLMQQPGLHRPQQSTTSSPAHQQQLQERPSLYGRNSNGLLSADAGV
MNSSSSAADPKPRLRWTPELHERFVDAVTQLGGADKATPKSVMRVMGVKGLTLYHLKSHL
QKFRLGKQLHRDTNVHDANKDGPHESSDHMQVTSTTASDSVAALQTKNPQDNIQITEAIR
LQMEVQHRLQQQLEVQRNLQLRIEAQGKYLQSILEKAKETLAGHRSSSPGLEAAHAELTE
LASKVGNDPSGCSFSLVSLPGIHNNPELLLQHVVDPRGGASSQQQQLPRQQSRMSDSSSQ
KSYLTSLTANPEDSGGASGGGGEQQQQAATEGGGGKFSPTVAAGSMIMRRSNEATTGAS
SHAFVLESSGSRAPALERPTPRRGAIQPQPVVEDSNNAGGGGSPHTSSAASSQHQGTTTY
RSDMQGGASHKSSSSAGQTVTTHHHHVKVEGPLDLNHHGHHGIVMPRSGELDLNTYDCWE
R
>SfaPSR_Sphfalx0119s0044.1
MYQTTKYSPSSNLMPQRSASQAGEPWASASPAAGPTYSAGSPGTSGGGGSGSAAKQRLRW
TPELHERFVDAVTQLGGPDRATPKGVLRVMGVQGLTIYHVKSHLQKYRLAKYIPDSLSDG
GTSDKKKNPADLFPSLDATSGIQITEALRMQMEVQKRLHEQLEVQRNLQLRIEAQGKYLQ
QIIEEQQRLSALTGTHSIGTDSGEAGAAPALEPPPPIAEVSDKVEIRPLPDRTITGAPAS
SVSVIPTRASELSSQPGPQSEHIKSESVNPTTVTDLAMPSSSPIQPVTNATHIADAAVLE
SNAPQQTKQQEVSPGTAETVLPFQSPVQKQGDVGGELLLT
>SfaPSR_Sphfalx0140s0042.1
MYQVKKYAASPPGAGGLMSSQTTTPSPHLAAAERASVLYGSDGMNSADPKPRLRWTPELH
ERFVDAVTQLGGADKATPKSVLRVMGVKGLTLYHLKSHLQKFRLGKQLHKDTTNVHDADK
DGSHHGSSDMQRTSTPASDSVAPLQPKTPQDSIQITEAIRLQMEVQHRLQQQLEVQRNLQ
LRIEAQGKYLQSILEKAKETLAGHTSSSPGLEAAHAELTELASKVSNSDPLGGGHSFSYT
SLPGITTTHHEFLQQFGADYNHHHHHPRQARSGGTSSPPPQYQQQKQQQFPHRQSRVSNS
SSQKSSYLTNLTANPEDDSAGGSASGGVEQQQQHPTATMEGIGDKSSPGGRRSIVRQSNE
GNSSHVFMNLESSGEASAVIVELPTTARRGGGAIQSLEEESSAGGDGSPHTTSSAASSLQ
GITTTTTTYDHGDMRPNTTTVHQRSHQSSLARTHVKVEDSAGSLDLNHGTPGLAMQRASDL
DLNTYSWER
>SfaPSR_Sphfalx0140s0043.1
MYQMKKYTSPGLIPHRSQTSSSNQQDRSSLYSGTLSGDGGVTSADPKPRLRWTPELHERF
VDAVMQLGGSDKATPKTVMRVMGVKGLTLYHLKSHLQKYRLGKQFHREGSVPEAKDVLHG
SSDGQVVNNAASQTMTTPKSQNSQESFQITEAIRMQMEAQRRLQEQLEIQRNLQLRIEAQ
GKYLQSILEKAKETLANHTGVAPGLEAAHAELTELASKVNTEPLDPSFSSLALPNLSAHE
DHAQGGSLPRQSSRISDTSSQKSYLTNLTVKPEDSGGASGSCEHQASTGIVTTSSN
>SfaPSR_Sphfalx0170s0003.1
MYQTSKFSAASSLMPQLGGGSQPAVEPWAQSSAAAPSYSAGSPGASSGGGGTGSGSAAKQ
RLRWTPELHERFVDAVTQLGGPDRATPKGVLRVMGVQGLTIYHVKSHLQKYRLAKYIPDS

FSDGGTSEKKKNPADLFPSLDATSGIQITEALRMQMEVQKRLHEQLEVQRNLQLRIEAQG
KYLQQIMEEQQRLSAPALTSTPVAATNDARGELAVIAPALELPEIGDVGGKLERKSSLPD
AATTPEVPCDPAGSLVPTTQASRLSSQPPQTEYVVEQEYVGS
>SfaPSR_Sphfalx0173s0028.1
MYQIKKYTNPPSLMPHGRAQHHTSLHQQLEQRPSSLYAGNLISSASDASGMNSSSSSADP
KPRLRWTPQLHERFVDAVTQLGGADKATPKSVMRVMGIKGLTLYHLKSHLQKFRLGKKLH
SDLTNVHDTKKDGLSHGSSDMQVTNTSSASDSVDPHQLNNPQDGIQITESILLQMEVQHH
LQQQLEVQRSLQLRIEAQGKYLQSILEKAKETLAGHTSSSPGLEAAHAKLAELASKVTND
PVLMGHGSFSLMSLPPGMINNPEFLQHGVDPQGGGGGGGGGALSLPHLWQQQQQLIPRQQ
SRVSDSSSQKSYLTSLTANKPEDSGGASVGGEHQTATEGGGNKSSSQAGGISKSMQHGRS
NKAGSSSSYANYMMQQGSLAGAGGALSSSAFALESPTSRRATALQSMQAEESLNGAGDGS
PHTSSSAASSLQSGKNMYHHGEMGAASPSHIRSSAATPTLATPPYVKVEGSLDLNPGNGI
RMLGGSELDLNSYGWER
>SfaPSR_Sphfalx0173s0030.1
MYQMKKYTSPGLIPHRTQIPSPNYQDRSSLYGGSLSGDGGGLTSPDPKPRLRWTPELHER
FVDAVMQLGGSDKATPKSVMRVMGVKGLTLYHLKSHLQKYRLGKQFHREGSMPETKDAPH
VSSGGQGANTAAACQSMVTPKSQNPQENFQITEAVHMHMEVQRRLQEQLEVQRHLQLRIE
AQGKYLQSILEKAKETLANHAGVVPELEAAHAKLTDLASTVITEPGGPSFPSLGVSNLSA
HERNAQGVSLPRQLSLVSDTSSQKSYLTNPSAKPEDSGGASGSCEHQVSTGTRKQ
>SfaPSR_Sphfalx0191s0003.1
MLTEGWQQPPQVPNSSHAATPAVVTMPTPQPPYPQPNMAKVQGGGRTGLPWAVEGGSSSS
SSASMQQRRGYYEPPPPHVGLPGISAMPSQYQYASQGPDTESSSPHVLQQQQFATLPSYQ
SLPHWGPGVQWPNQHQHPDGHLHPLPHPYPQQGADQWQASSQQLPPSASDVHQPPQQPLV
RGSSGSHHPGAQLEKPSGGAGSYALQHTGRESHQAFQDVNLGPAAQSGPVTLSPSPTSLA
IAPKTEDRDDADLGDHLQVLLDSPDGDGTDSSQMGDQSMNEGFGTGAEWLKWTELDPEEE
TIANCWTELIDVESGEEQTLFQTTRFEPLAPSTLKLKDQPTDQQPSGPAGGQHPESPGPS
SATATAAAAVKTRLRWTPELHEKFVTAVSQLGGADRATPKAVLRVMGVQGITIYHVKSHL
QKYRLAKYMPEISEEARAERRRNDTFLSPMGINSSHQITQALQMQMEVQKRLHEQLEIQR
ELQLRIEAQGKSLQKMIEQQAKVGGVVLGYSSEPHDPSPSAIVPATEGPETPGPLSKISI
SGSKGSLPASIMESAISATNGAPLPVHIIDQPDSNQSLLSAPKAVSEEQPPPVLLALNVE
PSSKRARTDIEPQVSDSRQVPVSVASDAQRYGASAAEDNVQQAAVLEMPPDMVKQHLTCP
AAEAWAAADGQHTCSFVQCSPQQAHTACAQPQPGSITQKPVRASG
>SfaPSR_Sphfalx0326s0015.1
MCRLLPSTESLAGTYTSLIRLIIFRRAATTTTAAAAAAGSAEATMYQTKKFAPISSSFRT
ATNSAQISPAGAAAAGSSSAGSAAKQRLRWSPELHERFVEAVAQLGGPERATPKGVLRLM
GVQGLTIYHVKSHLQKYRLAKYDPAAVAAAAAVDSASADDDAAAAAARKNLKNHLLLQPA
AYSDHVVPAAARPAASDTTSSDGTAATAARGVTVDDASLVRNMQLEVQKRLHEQLEVQRH
LQLRIAAQAKYLRKIVEEQQQQGSSTRPALRLAVAQGEEPSTEANTRDDVAANVVEEEEE
EEEAEAEVEEQQLPDLRLSLAAEEVQEELEEQEQGSAAVVPGASCSRDASSIVEQAVAAG
DAASSRSPPSIKELHFEITRSAALAGFPNDQASSAATPPPPPPLQTHDDDQQWRPRQSAQ
GTGASASSTVFTSCSSSWTYSEPASSTHDESTDHAAPSLKRMRRFEDLYLQPPNMQQISF
QGQTVKKEPSTQQQQQQQQQQQEGSSTLVWPAAGLDQQGDAAVRISLNSLQSAAAGTDDH
QAKTSHDLQADSEADHAVRQMFSRWEENSAAAAHKSSTSVTAAAIGNARRDSAGAH
>SfaPSR_Sphfalx0466s0002.1
MGVQGLTIYHVKSHLQKYRLAKYIPDSLSDGGTSDKKKTSADLFPSLDATSGIQITEALC
MQMEVQKRLHEQLEVQRNLQLRIEAQGKYLQQIIEERQRRSALSGTPVTTNSFFEEMDMT
AAASRESSAAKGEIVVPFAHDPITSTSDETEASHVHATKEEEEEAIEVHTMPRSIKDVIR
DVSLPDPQDLEEEKSPNKHDEFLAHGKQGESAAADDNELLVGHYYTTQNGDIPDPSNSDD
RSSQGRDSVSNFATIHTRSDGDSSKWCERSFVSDVVEVEEQVNFITTIRPGSPKPEVYLM
M

>SmoPSR_149357
MYQMKKYPSPQLIPHRGGAMPAQSEPLYIASGGDSVVSSIEPKPRLRWTPELHERFVEAV
TQLGGADKATPKSVMRVMGVKGLTLYHLKSHLQKYRLGMQMHKENNGDGKKEGGAKAQTT

GSQNSMNSNLSDGYEINRALSMQMEVQRKLHEQLEVQKHLQLRIEAQSKYLQNILEKARD
AFVGHIPTSAELEAAHAELTELAKGTLEGFEFTVPPLPELPSPSSETKAANQTPAAPTGK
KSTTSNLTLNQQQQQRDGQELDSDQAAEIDITT
>SmoPSR_270428
MVPAAQSSRISQISQQQQQPCSIISPVPKPRMRWTPELHERFVRAVEELGGAENATPKCI
LRVMNTYSSVDGVNILHVKSHLQKYRLVKDLPPSPVAKQQQSKQCSLELPSLNVETGLQI
TETLRLQLEVQKQLHEQLEIQRDLQKKIEDHGRYLERMYNKTEEATRSCHKNSDEPSPGP
QEQEQRKHRQTCDDQIHQASGTTTSTADSFQPPLKKLKPSQEQEELQPVGDEDVDFSELW
KFT
>SmoPSR_35907
ISASEPKPRLRWTPELHERFVEAVTQLGGAEKATPKSVMRIMGVKGLTLYHLKSHLQKFR
LGKQLNKDTNVANRNACPHHFASSQITEALRLQMEVQKKLHEQLEVQRHLQLRIEAQGKY
LQALLEKARETFS
>SmoPSR_405399
MLYEALKQSTTKMSQHQRPSLPSPSPHHHQALSDPQRIASGSSPGATSSGGGGGASSNAV
SNAAAAAAKQRLRWTPDLHERFVEAVGQLGGADRATPKGVLRVMGVQGLTIYHVKSHLQK
YRLAKYIPDPMGDGKSDKRRHPDLPSLGGSVQINEALRMQMEVQKRLQEQLEVQRHLQLR
IEAQGKYLQKIIDEQKKMSGGLDNQPGASPGSTQVFSSDITVQGSDLKYELTESVPGLMD
PPGGGGGGALPAITSALSALGSMAANNSLSRENSPASVPNPLGASYTLLNNNNNNIRTDD
AVVVSPRHQESVEEYLHSKSLQQQQQSYHHSSYLQCNTAGRLP
>SmoPSR_417629
MYQVKNFSPGTNFMSQQQQQQPTLYTGLSVEQAKISASEPKPRLRWTPELHERFVEAVTQ
LGGAEKATPKSVMRIMGVKGLTLYHLKSHLQFLMLCFHQKFRLGKQLNKDTNVANRNASI
VSYNTPNAQDLIAQQGHLSSSSSDSQITEALRLQMEVQKKLHEQLEVQRHLQLRIEAQGK
YLQALLEKARETFSVGGQDLNASVKLELCKAASSAETTLEQQLLFAREQRLQSHNLHQPQ
AHHIMGNKMEQGLDLNCKEVNGLDLNEFGWAAGPTES
>SmoPSR_423505
MDDAEFSAALEMMLCSPQEQGGGGNGLMKNEVEWAQEGWTSLLNPSDGGGDGEESSSKKG
VVVPQFVQVRDNSGSATPSNSSSSGGSGISPNLKQRLRWTPELHQRFVEAVALLGGPETA
TPKSVLSVMAVPEITIYHVKSHLQKYRLNKQIPEDPEGAPKPEKKKLTLNKLAETTAVTE
NLRLQMEVQRRLHETIEIQRQLQLQIEARLQLMHDGELKLKHTQTKSIQSEDKGEPEATC
SSKRDEVVAEAAANPRLAVPKTKRPLDHEEDHTEETSEVTCNNATVSPKRRLKLSQPGAS
ESSVVSH
>SmoPSR_438638
MGVTELQASVKALVRSGGEFNQDKACSTFYYVPSSGEASAHLSASGLGNDQKPRLRWTPE
LHDQFVKAVAQLGGPEKATPKSVLKLMGVQGLTLYHLKSHLQKYRLGMQIPRPETSGDGR
SNSEDSSKQQESLPLTQIIAVHAEVEKKLREQMEIQQQLQARIDEQCQHLYKLMESASPQ
KKSIMADLEAARKLQLDGIMELSKMYSGLQSAVEQHGKLDQATSNNTAGIKRRPNDDDPQ
RQLEEEQERVTKSLVQWKKEPVWDDDEAQHASGSTAEISTGLNLSSGRGLDLNG
>SmoPSR_442062
MMLQPKNSPSSVVAAAPVAPQVPSSPAAAGDLVCFSTPPPAAAVPSTPAAALANNVASTP
AASSGNVASVKQRLRWTPELHDRFMEAVNQLGGSDKATPKGVLGLMGVQGLTIYHIKSHL
QARILNLLLPKFRLAKYLPDTLGDGELEKGRDLEADSRGRQLSEALRMQMEVQKRLHEQL
EVQRHLQLRIEAQGKYLQRILEEQQKMNKLLRGDDGLPLSPIKLDADPDPEPDPAIDLTD
DSAEDEEHKPTTADVILVDEEKLAGGSLAAAAVDSGEEPHAKRKKVDDLQHSQHKDNGAE
GDAKENGR
>SmoPSR_57477
KQRLRWTSELHDRFVEAVTQLGGPDRATPKGVLRIMGVHGLTIYHVKSHLQKYRLAKFIP
DSSGDGTLFDSYLSSKCLCRGIQLTEALRMQMEVQKRLHEQLEVQRQLQLRIEAQSTYLA
KIIEEQQKMRGML

>AfiPSR_Azfi_s0093.g043264
MKKQKTDHPHTLSPPPFLPNEPERQNQNQNQNQNQNQNQQNQNQNQNQSSPGLSLSHKPPIRLSFYSPSTRQH
STTTLTLPLQASTVSVNNEPRSFQATQSSCQIEFDPIQKFLVCLNDPSDCAELQADASAEWEHLGGVNMSEQE

DPLASCWSDLLAIDGCSVSGLQETCYQLTKTWTTLNTQSQGPIQQLQDCSPIAGSQGTPPGPQEGNATGKARL
RWTLELHQLFVESVNKLGGAEIATPKGVLKQMNVEGLTICHVKSHLQKYRMAKYIPESPGYKVDKKPITNGGP
PSSDPKMVAGISEALRLQMEMQKRLHEQLELQRNLQLRIEEQGKHLQKIFEEQQKATGDIINASPSSRATHTR
SSEPVILECEPFLLPQCNNNLDELTHLVKVHNTDE
>AfiPSR_Azfi_s0031.g024472
MYQAKKFSTTNLIPHRGAPGGSDQQLNYVHGSSMGMNNDGGDNAARVSVASTKQRLRWTSELHDRFVEAVTQL
GGPDRATPKGVLRTMGVEGLTIYHVKSHLQKYRLAKYIPDSGKDGEKSEKSASTDMLNNLDSAAGLQITEALR
MQMEVQKRLHEQLEVQRQLQLRIEAQAKYLQKIIEEQQKLSETLKPTSSSSSKEESDEQPELLPEASISRETS
QQRGQAIDGCSVSPNVLNSSLLSTRRNCEDYPKNNILESDDKELGFTSSDGEPPLKKKKVTDGLAIKTEIEPG
LTEIGGP
>AfiPSR_Azfi_s0078.g038309
MYQAKKFSTTSLVPHRGPPGVADQHYPPGSSMMGMSNDGGDIAARVAAASTKQRLRWTSELHDRFVEAVTQLG
GPDRATPKGVLRAMAVEGLTIYHVKSHLQKYRLAKYIPESGKDGETSEKSGSTEMLNNLEGAPGIQITEALRM
QMEVQKRLHEQLEVQRQLQLRIEAQAKYLQKIIEEQQRLSETLKPTGSSQDPNNETPESNANHLQIFNATHGQ
GQGIESCSPPLTSLPSGLLLPTTNGAFEAYPQSRRVGKNVAESVEGDLGLARSDGGIDHHPPSKKMRVNIDGG
AGTLQRDDKTGLRKVGPETNLKPSEDRGP
>AfiPSR_Azfi_s0002.g001571
MEATNPSSKRSAEQGLERVERKRAIGYLEVEQDQDQDQDQEEEGGGDEEEQYHQSYHSVTDETSQCGDGLVGS
AHVYVHGHAIPPRVEPQCDDGLEAWDNDDANPFVDCWTSLLGIENREHSSHQQQQQQATNWNLDQINEAPAV
SSVMQGNPYHPSSTDSMPFAYASETPDNRALNKPRLRWTPELHERFVESVNHLGGAEKATPKGVLKIMNVEGL
TIYHVKSHLQKYRTAKFIPTGDSSEGKLDKRQSVDEPPDHDAKTSIQMMEALQMQMEMQKKLHEQLEAQRELQ
LRIEAQGESLRKLFEEKKQTGEPSDYKAS
>AfiPSR_Azfi_s0005.g009061
MKKGVHKPENKAGVEESARRPTEGSEGVTCARNPKISPRTDIPIPVPAIEPKLLSTWSDMSRSVSQSPYQEGA
LLSCGNNLVYPAIDPSLVSMSPYNPQSLESGFPEHEPPQFYHRKPQDRQQDQHNHQMPVCGNTLLPYHSQDGS
HLPYGEYHQDPEDKKKPYDYNPPEISQWKQQQQKGQSHHFSQTLTQSQVLGHGHRDEFLQKQPQFVNGEGLPH
HTSLRQGQKALEVLQQQDYSFQPPLDLEGFFDLGGDGSASSSLNPSILPYDIHSHCVLKARSCAPEAIQTLSD
SDKIASLAEALMSDFDEDPKCVGLTPTTNNSISGKSMNSSNTIHKQRLRWTPELHKRFVEAVDELGGAEKATP
KCVLNLMGVLGLELRHVKSHLQKYRITKDIPELQDGSTETSSNCLHTVSSTAATQMTEALQLQMEMQKQLHKQ
LEIQRSLQLRIEEQGKNLQRMLEQQQRVMDPSFSGGSDKDCSIQIRCETDDDNVHSSEQMSRRSDQEEESHPK
RLKLDAIAKQGTC
>AfiPSR_Azfi_s0481.g073115
MFLLLSSLDPIFGVEIPSFSTLLISGFSTATRSQIVNIPNRTSKIEKCRVTEAVFSHQRQDFSGKMQDFSGEM
QGNAICRDGSVVLSTDPKPRLRWTAELHERFVDAVSQLGGADKATPKAVMKVMNVKGLTLYHLKSHLQKYRLG
KQPHRETTPDAAMPDRNSHITVLLRFSLIVFPSNVQITEAIRLQMEVQRKLHEQLEVQRLLQVRIEAQGRYLQ
SILEKAQQTLAGQTVTSDGLEVARAELSDLATKVSNECLNPSFSLVISDVSNSVSERRVPENDQDIAECSPKS
CLTHVTTNERSEHGCSRDIFQNLTKRSRLFLGNPEMLVSRSGNDEEDLKPNEGILNADICSLGERVLSSSLCG
WDREIARHGEVYRNGCNKTFDLVEEFSIVKEDGEKCFRSIGRPEPRRAGLSAEQVLTLSSSYKNSVIATAGFG
NKLSTCNPKLGKGLDLNMDSDGNKVGGIQEFDLNGYINGS
>AfiPSR_Azfi_s0121.g046895
MTHIDSYGTYSDARRSFTFSHLHGTNDFDVGGVILSTDPKPRLRWTTELHERFVEAVHHLGGAGKATPKAVMK
VMNIKGLTLYHLKSHLQKYRLGKHSKEANTKIEHFSGANENIKPSITISTGSENQATPENTQIAEALHMQMEV
QRKLQEQLEVQRLLQARIEAQGRYLHSILRKAEENLATQTLGLVGTDQARLMDITNKEVCMDDFSKCGAMDEC
ESHIKTECSFDSSITNVISKERYETEENGGLEKKYRGDLVGDQESVGMMMLKYKEDIQVQDEEDDNVGMGGLS
LMEQVSAHKRCFIDRLLVSTTRGFERGLPYVQVIDDNELVDKRTLLSDRLLVSDEASETKPLSKQAMHSPICM
PNQLGISKPMGPQNMLSFHSCIDLRLGLDLNFQCDNIANQN
>AfiPSR_Azfi_s0319.g064433
MYNPPPHFGIPNARAFALDGSSSQFLHHHHQQQTEVSLCDTNNNLHGEELSVMVSSDPKPRLRWTPELHDRF
VNAVAHLGGPEKATPKSLMKVMNVKGLTLYHLKSHLQVYFCNSMNYVPPEVQAWKTTHAGYEYDGDSSMDDIP
SLKDKKESTDITDALRQQMEVQKQLHEQLEVQRRLKMRMEAQGRYLQLVLEKAKDLLNLQPCEVQETTKFDFD
SNEISHYSNTPLHSMPKLHRMTNEKKQPLAQEFLSVLYPEPNQPCPTSSIFPQDLSRIGKRSQFENAENTDFS
LLSDGILVNEPHFMHGNTLIDEDARHVTLCANSKTGLDWEYNAYENLSHSMW
>AfiPSR_Azfi_s0020.g015442

MASSTSSSSSKQRLRWTPELHQRFQDAVAELGGPDRATPKGILKVMGVPGLTIYHVKSHLQKFRLAKYMHDAN
QVGKFDKRKSLEMVNMEVADQAEPSGMHLTETLQVHMEVQRRLQDQLEVQRQLQLRIEAQGRYLQSIIEEQER
ITKTMDNIDPGKGNIKLSMVPCNPLTLECSPGPESIIEEGACDNSSPESSHAFNSSENTTLGSSTNTRISNIT
MDGSSNSCDDQLPLVYHTPSHIMQMSGSQDHHPTKSSHLGSYNNLKRGDVYVDYGQKPSSFHNTLSNDHHDDL
WHDDELYHLGQQRF
>AfiPSR_Azfi_s0003.g007457
MSNPSSSFRSKTNPSTPFIASSDHVSSIPSSTQPYPSQTMSNQSSHSKQNMHEQYIPQQIKQHQYQHPHTHPH
HLHQYPQVNNEGIIQPSHFSTHSQHPHQMPMDLRELLEFAQTSLQEPYPSPTNASLPGIAQRDSNFDWCDLDR
LPNNDGPGHVASWAQMFISTTNKASPKPTNAKAIKMAISSEVIHSSTLATGNSINLMSAKHKQRFRLRWTPEL
HKCFREAVKELGGAEKATPKCVLKLMNIPGLQLSHIKSHLQKYRMTQDIPGSQRDEHGRKRRSSSSMEAITML
DGSMALQMTEALQLQMEMQKQLHEQLEVQRNLQMRIEEQGKHLHNMLQQQKQQQQQQQQQQQQKEKEAESAL
NIL
>AfiPSR_Azfi_s0003.g007690
MAVASLQFNRHDFSLPGFNSTKNKFVEISVEDFAHSLQYQFEPDKRYVIIKNGFILIDPEDFDHGPFESNKSS
HQKQLSLSDDNLPGDLNAVASNDPKPRLRWTPELHEHFVSAVTQLGGADKATPKSLMKVMNVKGLTLYHLKSH
LQKYRLGKQPTRDTNVVDGCKENRGENCTIARSIDITDALRQQMEVQKTLHAELEVRRLLQMRTETQGRYLQS
VLEKAKELPSQTSKAMVTTDKFEHNSYQNALPMISHPRANFEGEDKGALAYDLCSIYPEPYQDQSCPSFLWSQ
DLSWVRKKSKLDNVENQDLPMVRDGVFVNESLGINEEDGRRIVCANSKVEFGWEQYNSYESLPNSIW
>AfiPSR_Azfi_s0027.g023545
MYQMEQYGAMDRFPINHIQPFFKINDKTEGGRGRGRGRGGDTDYEDDHGISSSMSDGVILSTDPKPRLRWTSQ
LHQRFVDAITQLGGADKATPKSVMRLMNVKGLTLYHLKSHLQKYRLGKQSNKDMVEADMPKFGDKSTSESHRS
VTGRLDSIPIQDSPENAQITEAIGLQMEVERKLHEQLEVQRLLQVRIEAQGRYLQSILEKAQLTLARQNDSAT
ELEPECPESSDQTTKVSAMDCSGSNYSRSLLPAMGDGTFEDRGIGHPPERSECSPQSCLTHLASNERSDTNGS
NDEACLNYKRPRLFNVEEYEDAEVGQNMDEEGLQLNQSNGSRSYGRLLDVNAKDREIGLHWGFDNYRVVDPYM
LKEEEGKGGFRSFESIDKPVPHFHREHSVKPTEIGAYRDRISNHTAIYRLKKDLDLNAIGDGSTFQHREFI
>AfiPSR_Azfi_s0064.g035511
MYHHHHRRRLEAHGIAAMAASEINDDQRHGVVTLGTSLCEGLLSGTNLPGDVSSLLMSSSADPKPRLRWTPEL
HQRFVDAVLQLGGADKATPKSVMKVMNVKGLTLYHLKSHLQAGQTATEREHWGKDISDALRQQMEVQKKLHEQ
LEVQRRLQTRIEVHGNYLQSILEKAKQLLAPQGAGLRFNDISCIDELPWQSCHVDKIYESTGSENYQSRWERD
KPMAASRDIHLELIGRSVAYSESMALSKEDGQIGFCEQDDGLLHRIDAFNRSRLVTNGLSS
>AfiPSR_Azfi_s0035.g025591
MMKELGFTHEIMQESLVSGHGSLTLATDPKPRLRWTADLHRRFVNAVAHLGGPEIACGSPSSMTAVQNTISNQ
NDLGNAPIAEALRLQIEVQQRLHDQLEVQRLLQMRIEAQGRYLQSILEKAQQTLAGQVVASDGLDTAQAELSA
LATKFPNKCITNLELATKVSTKCLTNSDLPLGALVDRRILCPEVTECSPDSCLTHSTSERSETIGSKDYYQSV
DTKRPRLFLRDSEAKVDTRNSQEDMKQNKHIINVRSESTGSKDFYRNIDTKTARLFFGDSEAKNDEDDIKQKN
GGNMERDRWSSLDERLSSSTSSCVWTHESECEPLERIERFMGKGNLAFRSVCYVERPEPRKAGLSIENITNLS
AGQEENIAMQFPNRVSHCNSKLGGKTLDLNTNNDGDQTWDEA

>ScuPSR_Sacu_v1.1_s0064.g015782
MYQAKKFSTTNLVPHRGAAPLDQHYHHGSSVAPSLDDGGDVASRASAATKQRLRWTAELHERFVNAVTQLGGP
ERATPKGVLRAMAVEGLTIYHVKSHLQKYRLAKYIPDSGKDSEKPEKSSTDMLNNLHATSDLQLTEALRMQME
VQKRLHEQLEVQRQLQLRIEAQSKYLQKIIEEQQKMSDTLKPSQDSDGSQVPERIAEEETLKSNAKELQIICV
RDQNIDGHSPLPSSRHSTRECDDNPSHPQDDKNVEVGSPLPTSDQTHDEPSSKKPRLVDGGRKTENQSGLKK
>ScuPSR_Sacu_v1.1_s0009.g004589
MIFFWPSFFLCTAHPPVRTPSSLCINLPFLPFPSADPALRSVSFLATHSLSLPAPSFFTHPTSICGSSSVFKP
KLFSRPCEIRNRLRDNFTAYGMYQAKKFSTTSLMPHRGTSDPHYSSSGSTMVPTNDAGDVAARVAAASTKQRL
RWTSDLHDRFVEAVTQLGGPERATPKGVLRNMGIEGLTIYHVKSHLQKYRLAKYIPESGKEGDKSAKSDILNN
LDGAPGLQLTEALRMQMEVQKRLHEQLEVQRQLQLRIEAQAKYLQKIIEEQQKLSETLKPTNSRSQEPEGVKV
PETPESVADQLQIYSGRPGQSMDGSSPSPSPLMSSTQLGKNVSDNNSTEEEVGLANSASAAAFGQPPSKKMKL
VEDAAVAGAAQREEKPGFQPEIRAST
>ScuPSR_Sacu_v1.1_s0011.g005338
MYHQAHYAICQTDHDLPSVHDSICLSGTNLPGDISLMSSADPKPRLRWTPELHERFVDAVTQLGGADKATPKS
VMKVMNVKGLTLYHLKSHLQKYRLGKQPPRENTAEYQKGQSENSSKEDASAAAAKDEKEGKDISDALRQQMEV

QKKLHEQMEAQKRLQTRVEVQEKYLQSILDKAKQLLAVPETTTAASAPTPNLLLDSTPAVDFDDLASSNGSVN
ASSSCLSDHSPRNVPDVAHGAIAWNMKVVEDLSGRMSYVFSEGTGMLESCPPYATGMDVKLRHTAGEEELLAG
REKAAASVGEMAWSPCPRNAAATVYGSFADQTQWRSDGVGEEASSQGVIGQSVNVVFDESMLLLGSKGAVDDN
RDSEQDGAMQRLDAYNQWWCSNSNSRARGNCREASVKTNNGGSNTPSSMLLPRQSSSREFQKFSGLPYEWATS
S
>ScuPSR_Sacu_v1.1_s0004.g002208
MEPFMDQWFQVPSRDLERPVRLGRREVKEVTSSIADEANANRTACSDEYLCTLRHRGCWDSVVDLALHPYILA
FYLLPHSQQLPPRRAIHQLNFHSLIGSSRRWLRSNSATDRDLQIDMELSSRDCGGEAAVILSTDPKPRLRWTA
ELHDRFVHAVSELGGAEKATPKSVMKMMNVKGLTLYHLKSHLQKYRLGKQSTTHREASHGVDSSAGDAESSLT
AVAHDHQAPTNQNLINAADNLQINEAIRLQMEVQQRLHEQLEVQRVLQARIEAQGKYLQTILDKAQQTLCADS
NMKNHDCSNDVNNTNNNLNPRVGFYYDGNLSLNLHSSFSTSVSELKKQLRDVDKAAKQLEQQQLSDNAVIAMA
VNNYNYDRNSNSNNSNSEVQLKVKDSEMVALMAMRKSTSSRKYVDSNDTNPLTPNSELGMQQQQCRVLEDVSA
ECSPHQSRANYGRRQQSCATEPILLHDNSYFGSATTSTRSATTDYNVNSELIRSTAAAADEYDDSQMGRRKRP
RLSTVGLQLLQLQGSSADDDHDPYCSRSNIIDVDDESELTTSYRGGQQLVAHALDPANLHLQQSSNNYDSDRG
YGSSCCCCHCGCDCSSRRKMKESCNRAITTHYEYSCRRGCRPSDGSAAAAADRHGAAGHGSSKSFKGFDLNAS
DISKQPASRAVQGASLSSKQLNPQAELFKGHTSLSSKQLNPASRAVQGAYISQFQAAESGKQSCSRGICTIFK
SLAPGRAMHA
>ScuPSR_Sacu_v1.1_s0007.g003614
MYSSSYSDATTLAAAEDTVLSHKQKQLMPSGTNLPGDANGVTSNDPKPRLRWTPELHDRFVDAVTQLGGSEKA
TPKSVMKIMNVKGLTLYHLKSHLQKYRLSKQPHRDVQKDASSEDFALVGSIDGATQVQNPSLPTVATQNQKES
LDDLKSHMDLQKQLQEQLEVQKRLQMRIEAQSRYLQSILERAAAALSHQATAPVELTNVKAELADLVSDVTNE
SMVSSFSTFPITSPPPLPHLETSTLSRHAQFVTPCITVTQNSASSRSNLHECMDSRSTPRFWSTEDARFKFCN
GDVHADESEGSNLRLEELPGAMRMDSENHVTWKGFASC
>ScuPSR_Sacu_v1.1_s0013.g005903
MMASFSNRMQTFAAQMQGSITSASDTVVLSTDPKPRLRWTSHLHQKFVDAVTQLGGADKATPKSVMRLMNVKG
LTLYHLKSHLQKYRLGKQANKDIGTADQLTKIDCTTASKADGSLPCSSKSDSVLTAAMESPDDVQINDTIELQ
MEVQRKLHQQLEVQRLLQVRIEAQGRYLQSILEKAQLTLAGQKDIGIKAACAQLSDLASKVSTECLRSGSPCP
GEDWRIQPPHERAECSPQSCLTHLTAIDRSDATGNGESYPCNKRPRFSIEVDLDDPQVSHCTEEKLHLQRPPF
SDANLSNMLRVWNAKDREVILHSSHHRDLDDRGGLERCIIKQEEGQELAKTTIFCTERIENSVGNAIKQDAPN
RQNLHRTSSNNKMRRGLDLNAVVDGLHREIDLNHI
>ScuPSR_Sacu_v1.1_s0124.g021606
MHFLRTLPSTSCSDGAAHALAYNSNTGGCDAPQSLRHNQTPIMGGGDANDDDDSSSSYDDYNCNKNNNNTSCV
QAATVPQLLQQQDDIHNNNNNNNSNNNIIQLSTDPKPRLRWTSDLHKRFADAVAQLGGPDKATPKSVMRVMNV
KGLTLYHLKSHLQKYRLGKQPHKEVNTELSKLGNSGSILSSDVNHKSEPSVNMSTQTLEEQSKQIFEAIRMQM
EVQNRLHEQLEVQKQLQVRIEAQGKYVQEILEKAQETLTRAGGLHLDKDSRAELSHLATRMASLDSHPMLLST
SAAATTTCTTAAAAAPPPPPPSLVAPQSTLPPNTNSNVHNFGGNIHTVSNGGNLILVSNSRDTHLHNTNNKNP
DRECSSDSSLSDPPSDNTTASTSPCKKRLRMFLHPALVPATTVSAQCERGSAPHYHLDLNA
>ScuPSR_Sacu_v1.1_s0183.g024956
MSHVEPTVICSNDLFPHLQAFGSSRLHGTNVSDRGGAVILSTDPKPRLRWTVELHERFVEAVNQLGGPSMLVP
NTREVHAEATPKSVMKVMNIRGLTLYHLKSHLQKYRLGKHMKEAGTKIELCFGENVPERFLCLVKTLNDMSPL
FEHLETLNLPVQIRQTKFHLSMCPVCNNFSKIFNIQIAEALHMQMEVQRKLQEQLEVQRLLQVRIEAQGRYLQ
SILKKAEESLVNQTEKNVGFGTMAQDHLSDLGNKVRMDFSVTDFPRRSAKGEHRSCAKTESSFESSMSNVISK
DVSEPEDDGDLQYLDKDKYGPFFKTVCMQRDRGEAKLHDDGVLMGGVSRRLGDKEATLQEGAYSRACLGRLLA
AAEGQFLSCKQGIENRHLVEQNSAYKGTFLDQFPTYDGHYERLSPSKHTISSTPTIMNMDNKSNTLETKGSWD
GLSSCSIHADLRLGLDWNSQTAKSTNLNRQIDLNGCS
>ScuPSR_Sacu_v1.1_s0069.g016534
MGGNNLDMHQAAVHAQVQQHAAGPQGATIMSEIAVLCSPDPKPRLRWTSELHDRFVEAVMQLGGPDKATPKTV
MKVMNVKGLTLYHLKSHLQKYRLGRKSQRDQAGGNASGSAPTSAVNVDMNPAIQQQISGGVVKCEQVINGGGG
MMEMNGMCMTKQLGVSLPVDHMRDSPQKRFLEHNHMERSLKQRKDAQDKYLQSILMPENSKAADSYLAVHAQG
APTIMVDELSLLCSADQHTNMSNNVNTSIAMAHAHAQQQQQVDMDMNPMHAHAHHHHHHPHHTGHVHHMVDLD
DPADVVDAHYDVFSAAHLFGSGGGGGDHLTKSGHGGAATYLFDTSPTQPPPSDHHHSVATAASNNGSCFNVPG
GEDQGSVVHSSYAAFILNDSSHDSFLAPYNNSSAASNSHLHLNVSSS
>ScuPSR_Sacu_v1.1_s0107.g020383

MNRLGVPYDMPSDLSPYIYMAPLCPNSNSSMSLSPSYAMVQSDAASSDSVFYPHQLYHSLEDDSQQHHHFHEP
QMYRDGGDCQYMKQQESPAGSDNDYQRMQLQYQQSSLLFRQAQAINQHIQQQRQICECECPQNNHFELDNTIQ
PAHVLDEFLRLEDSSFYSNLNSPLSSGSCLQYISDAQSCPPELHVSNVQPPVLALTDEAKIASLTEALMSDID
LDPQHEQTSTQTTILSGESFNVSNATNQKQRLRWTPELHKRFLEAIAELGGAEKATPKCVLNLMDVPGLELRH
VKSHLQKYRMTKDIPTVREDNSKPAKLREANATHDMLQNLHTGSRATQMTEALQLQMQMQKQLHNQLEIQRNL
QLRIEEQGKNLQRMFEQQEKVMLPALRVSNSNACVPNRSEANELLVAGKMNQKLAESGHNVFVEEYALSKRLK
LDGDSKNSAC
>PabPSR_MA_10266929g0010
MFQNKKFSTDGLVPQRFHGSEQRGAGPLGVTSSSSGSLVTSGGSGSTKQWLHWTPDLHDQ
FVEAISQLGGADRVTPKGVRKMMGVQGLTIYHVKIHLQKYRIAKYIPESLEGGKSGKKKG
NKIVSNLEAISGTQIVEALQMQMEVQKRLHEQLEVQHYYSIYIASAI
>PabPSR_MA_10433112g0010
MSSGGGPPPESLDASGLPVLDGGGSTSVSAGCRPFQVSLMDSNERSMRASLGDGGGGGGGVGGACDDAPLPQL
PQVLTILCAAGSFLGYDVWLDAWSYVAQLDGFGGSIQIQNTAGSQARSIPDHQYGAPLDQPLTTWPITPLTNS
VSVRYSSLILSASASIMFQNKKFSTAGLVPQIFHGSKQRGAGPLGVTSSSSGSPVTSGGSGSTKQWMCSTPDL
HDQFVEAISHLGGADGVTPKGVLKVMGVQGLTIYHVKSHLQKYRLAKYIPESLEGGKSDKEKGNKIVSNLEVT
SGTQIAEALQMQMEVQKRMHEQLEVQRQLQLRIEAQGRYLQKIIEEQQKISTAKPETSSSPETSPPRYGEPCE
ITAKAFPDSESPGKTGRHRQDGNAVMPPADIAQNDQSAGIAYPRDEPQLQTSSYA
YRVSDGMPCSDNLEFGSPSKRSRLNNDTGQPKQENLSSQPHDQQQTLPMSYSSQTSHLKVEELPGSIYQNRVA
GLVGQIESTRPSQSTADLHAQQLPAKLPINNLDHISHAQDFYRPLCAKNASDSIRFISPGSQIQVPASNLPAP
SQDQFSSALDGHDGCHNVSSSQVSLFEPWP
>PabPSR_MA_120020g0010
MYQMEYPSNPGIVLNHGRAPSQPERQHQQPALTGSSHYPCDPGVVVSADPKPRLRWTPEL
HERFVDAVSQLGGPDKATPKSVLREMGVKGLTLYHLKSHLQKYRLGKQPHREVSVDRSND
GGSSDGGHCLSRMARDSSITPNHKEAMQIADALRAQIEVQRRLHDQLEVQRRLQLRIEAQ
GKYLQAILEKAQQTLASEASTSAGIEATRAKLADLASKLPTNSVRPNYSAMKFASHDMEQ
SLNESESPIEKKRCRKLFHESNGGNSPTKLRELGLNASSCKTEEEEDEEEEQLIKPTTVE
RPAATRALVDERMAMLVQSNGLSLIQANHEPGLYHPPSMLGNCRKVAEGLDLNRKGEGSV
PQQGRELDLNAYGWGR
>PabPSR_MA_181986g0010
MEKMYQVKNTVSSNGSVPSHSERSLLLQGSNGPGEMGLVLSTDPKPRLKWTAELHERFVD
AVSQLGGPDKATPKSVMRVMGVKGLTLYHLKSHLQKYRLGKHLHMVVNLDNNKNGCSAAA
SKTSEMQGASSAASSMIGANKTDALQINEAVRMQFEVQRRLHEQIEVQQHLQLRIEAQGK
YLQSVLEKARETLAGHNLGSVGLEAARAEISELASKVSSECLNNAFSSLPEMPSLYKQQQ
HMVQAVHSLNQQSQVTDYSMESCLTSNESSEKNLDRNMQNAGKKRLRTFQNRNVPFWQTE
PREDASLRELSATQLSWEENLKEDEKFLSSSANPKDSDRNSIPTKGNLRERTTSFRVQTE
ENGYGMPDAREKQMEVNGSCVGQVIKPEVSRSFSLQEVCQKEFHRDEDESPSVSFNKLSK
LTEELDLNANDDTSIASNCREFDLNGYGWTYEH
>PabPSR_MA_21538g0020
MNSFVPNFNSQPDNRNPTAKLTHQTEKMYQVKKPAMESHSERPAMLQEISISTETGLVLS
TDPKPRLRWTSELHERFVEAVTKLGGPDKATPKAVRTRMGVEGLTLYHLKSHLQKYRLGK
QPHKEVNVDTSRDGLSHGGPVAGYGSTACRVSEVQVACNSVPSIIPANMTDTLQITEALR
MQMEVQRGLHEQLEVQRHLQLRIEAQGKYLQSILEKARETLAGYNVGSIGLEATRAELSE
LASKVKSECFNSTFSALTFSTMPEIPGLLPESQIARQQSQAADCSAYSCLTSNESSEKSP
TENLQAGGRKRPRSAYDNKLQFWQDDIREDMSLQELTGEENLSSPSSRTKDSSIVFHERA
TSDIVQAEENGNRLSDQKGKECEMNGVLVDPPITRTLAFSSALSLQENSQMYEATKVYTN
QDKTRSKGTLQITEALRMQMEVQRRLHEQLEVQRHLQLRIEAQGKYLQSILEKARETLAG
YNVGSIGLEATRAELSELASKVKSECFNSTFSALTFSTMPEIPGLLPESQIARQQSQAAD
CSAYSCLTSNESSEKSPTENLQAGGRKRPRSAYDNKLQFWQDDIREDMSLQELTGEENLS
SPSSRTKDSSIVFHERATSDIVQAEENGNRLSDQKGKECEMNGVLVDPPITRTLAFSSAL
SLQENSQMYEATKVYTNQDKTRSKGYNKFSKMAQDLDLNSNCDTTVSSKERGLDLNGYVW
SR
>PabPSR_MA_8183372g0010
MGLHRAQLPLSQRDHQRGLSTGNIGQNELGVVVSTDPKPRLKWTPELHERFVDAVSQLGG

PDKATPKSVMRVMGVKGLTLYHLKSHLQKYRLGKQSCKENNSDTNKDGGNSDCSKDPNIK
SRDNNTRTNQKEAMHITEALNAQVEVQKRLHEQLEVQRCLQLRIEAQGKYLQGILEKAQQ
ALIYQTSTPARLGAEPAELVEMVSNVATDCNETASFSRSSASSSLMDMPTQDEAQYERDR
SRVYFCDNGSSIWHNEVSAHDKLQEPGFNDSNGCLYKTDSEHELQQTREFNVCSMTNSMW
ESNSHNRLLSSDSMKLIEAGQDHPQNENSLRIMDETIMKPTVERLTSRISLPEERLPMLV
QPPNAHLFHACLSGHEQGLFKNYDPCPPSMLVKRRKAVEGLDLNTRGDQGIPPQQGIQLD
LNTYGWGR
>AtrPSR_scaffold00010.492
MERSANEGVIVPLPLTHATKPMLKWTPELHQLFLDTVARLGGLDEVSPKVMVRAMGIPGL
TVFHLKSHLQRERYKRAKAQGLVHSRSLRNAMSNGDPWIQSMDAMGGIRRMRGAIFPPDE
NIRMEEEAIQKPISMKREMLQRIEVQKHLKLRMEAQTRYLQTVLERAKTALSSYNLDSSG
VEAFRAELSELALSVVIHDYMDTAISKNKASEFASPQMENQRMTGMGASGYHAPKPQVNY
GSGNSTLSSPKSPLTSLTMDVSMQGHNYELRFNKSCVELIPVMRGGPMVATNRASRSING
NNAVSLILQMGADRKEEGQEPPLGRKSVRMEECKRVKETDIGIRETDMEPPHVDQVSSVS
FMKSVEPSDKLGNHENGGLKLVMGFDLNACCDPLIIAIDK
>AtrPSR_scaffold00022.209
MNSQKIECPEENFPASNNYSFFSPQINIKQGNSQSPDAFSFSNGSLAPPPVTHQLPCSFS
SSNYFGLLPFSRYTNDRLMGFSHYDSLSPTPTKNSLGGSLPQFSVDSRKETNNNIGSFDN
SQYVMGFPLLGEHCVKTTSYCNLNQQKDSKRSEFYPSLPIDGSDVDVSLPRRHLLAIEGI
QNCLGNLQSSNCSTTELAFQNQQERCFSSCHVGTSVASTHVTCGVGASNKTRMRWTQELH
EKFVESVNRLGGAGKATPKAILQQMGSEGLTIYHVKSHLQKYRSAKYLPDAAEGKLERKT
TNDIRHDLKTGIQITEALRMQMDVQRRLHEQLEIQRKLQLQIEEHGKYLQKMVDQQQKRS
ENLFGSHNINVLYSEKQPINLAGVIESSFDDELLDKTPSRFSLGDGQSHE
>AtrPSR_scaffold00025.171
MDSEANGSFRQIDGQDSSDSSRLILSADPKPRLRWTAELHQRFVDAVTQLGGPEKATPKS
VMRIMDVKGLTLYHLKSHLQKYRLGKQTQREAVINSNGRDVSLEEVTTVSDSSICLGSEN
RPTDTCSPTMVTDTLQISEALRLQIEVQKSLHEQLEVQRHLQMRIEAQGKYLQSILEKAR
ETLARHHMGSAGIEETRRELSELSAKVSDGCRNPATFSLIGHQGLTGSQAKCEVLERLKK
KPMAPVLDLNSNLEPSNEDFGSGGCRKKEFDLNGFELGL
>AtrPSR_scaffold00025.314
MALNIEPEIFDEPLPPQFRNESSSQQGSTNELWNPSPYIETTNPGSSFSHSESSRHMLSQ
SSSFCTSLHVSSSSVPENNRHLANLPFLPDPLKGKLPASKASSLNSFLSISEDLKTESKE
HDTSENLIQDLFNLSGNASDTGLCSENYPNDDMIVTEQFDWQIISDHLDLAITDIGENPG
LDDIYGAPQISSVSTSGLECSPKHHQSLHTEATQSYSAPSPSGTSTGNKPRLRWTPELHE
CFVEAVNRLDGAEKATPKGILKLMNVEGLTIYHVKSHLQKYRIAKYLPEVKEDKKNSEFE
EKQQPSTDGESRIDIKMGMKVTEALRLQMEVQKQLHEQLEIQRALQLRIEEHARQLQKML
EEQTKAGYNLMGGHSSTAPSTSAREAGSPLSTSHVEPMATTANSGGTDSETESKASESRK
RARVEAGSEEICNE
>AtrPSR_scaffold00026.77
MHEEIAFVNMLHEGEFSESSSVPQNAHRYLGLPSYSIRSNMASRGGPSTRQRLRWTPDLH
DRFVEAITQLGGPERATPKGVLKIMGVQGLTIYHVKSHLQKYRLAKFIPESIGGEKIDRN
DKEVDADQTASSRLDIDEALQMQIEVQKRLNEQAEVQRHLQLRMEAQARYIQKILEEQQQ
NEKPSSPTTEGKSLWEKDQVPLFPPLSSEELSLPCGSPSKRT
>AtrPSR_scaffold00048.151
MFNSGSGEQSGANGAGNLSERDAMLSRDPKPRLRWTPDLHECFVDAVTKLGGPEKATPKS
VLKVMGFKGLTLYHLKSHLQKYRLGKQPRRESSTEGGKDGSLGSQKPPILATDSALRQNN
QGEIQIAQAIRYQIEVQRKLQEQLEVQKKLQMRIEAQGKYLQAILEKAQQSLRCKEGVEE
ESKVGSDYQKTAAGSLDLNVNQGSLDLQIYKRYS
>AtrPSR_scaffold00061.72
MYHPKNISSTTLVPTKPSSRDPQIDNAKMAANSGGNPVNNSNMASRQRLRWTNELHERFV
DAVAQLGGPDRATPKGVLRVMGVQGLTIYHVKSHLQKYRLAKYLPESSSDGTKPDRKESA
DMLSNLDAASGMQITEALRMQMEVQKRLHEQLEVQRQLQLRIEAQGKYLQKIIEEQQRLS
GVLAGTPAIVPESGDHMPEPDAKTDPATPAVTSEAPLKDEPTKTKCLVAKSFSQEESFSS
RHEPPTPDSGSPGSSPCAERPVKKQRGLEPSGLGGLGHPILESSSGSVFRQADMCSAYSS

LEGVSGRFDSLGSVSLCEREGFGDASGAEL
>AtrPSR_scaffold00061.73
MYPGLNHHHEQVLSNEETHGSVEAPGLAIDACLVLTTDPKPRLRWTAELHERFVDAVIQL
GGPEKATPKTIMRTMNVKGLTLYHLKSHLQKYRLGKQSSKEFSNENRDASSLSEDQATSS
PPSSRVMTRDMNEGYQVTEALRVQMEVQRRLHEQMEVQRHLQLRIDAQSKYLQSMLERAC
KYMTDQAMTSGSNLSVEPQNNHLSFSLLSQSTPDDISETAGVQNNLEDPRAQIGDCSVDS
CLTSQSEYSNGGKKRSRPPCFEANSTNWGEDIVGLNMVGQVKMVRNEGIDSCASNVGLHS
PRLEHSGLRRPSVFLDAGPVHAKGGIYYL
>AtrPSR_scaffold00089.17
MYSAFHSLEKGLGREDLQGALEGTNLPGDACLVLTTDPKPRLRWTAELHERFVDAVTQLG
GPDKATPKTIMRTMGVKGLTLYHLKSHLQKYRLGKQSCKEFTDNSKEESQGTTSSSSSKL
ASQDMNEGYQVTEALRVHMEVQRRLHEQLEVQKHLQLRIEAQGKYLQSILEKACKALADQ
TVASAGLEAARQELSALVIKVSNGCLSAPSEFLNLPILPEMASVCVDDKKLNRQAQMADC
SADSCATSNESSAGAPLQSGGKKRLRPMYCEGDSLVWEGEARQDPQWMTPL
>AtrPSR_scaffold00094.1
MFHPKKPPNMCSQERQMCVQGDSGLVLTTDPKPRLRWTVELHDRFVDAVTQLGGPDKATP
KSIMRVMGVKGLTLYHLKSHLQKFRLGKQPHKEFNDHSSKESERAAVLELQRSGASSSGI
MGRDGNDNMQVREAVRMQMEVQRRFTEQLEVQKHLQLRIEAQGKYMQNILEKAFQTIAGE
TMTSGSLKAGGSSHQGLLDLGASCMNYPCLQDLHIYGGDQNPHHHQLELQSGGQIDSAFM
SSQEDTQEEFDGAFETKPLQVMEGDCSINKKFEMRAPSNHMSMPRPITPSARNHSYG
>AtrPSR_scaffold00095.37
MYYPKKFSTMGLVPHKPQSQGADQLANIGVSGGSPINKSPTSGGAGKQRLRWTSDLHDRF
VDAIAQLGGPDRATPKGVLRVMGVPGLTIYHVKSHLQKYRLAKYLPDSPADGTKDEKKES
SDILPNMESAPGAQISEALRMQMEVQKRLHEQLEGLELEPSLTDSAECTWSRTWRLGWGK
VIYGVQWFKTRIQLGIRQDPSSSNV
>AtrPSR_scaffold00119.73
MYYQGKNCFSSSRATMPPERPLLLHGANIQGDPGLVLSTDAKPRLKWTPELHERFVEAVA
QLGGADKATPKNVMRVMGIPGLTLYHLKSHLQKYRLSKNLRSQSNGSDKNGPAAAAERMS
HTNGPHMGMANMAVSGTNKGLQINEALQMHIEVQRRLHEQLEVQRHLQLRIEAQGKYLQS
VLEKAQETLAKQNPGSSGLEATRAQISELVSQVSAECLNSAFSGLTEAPSLNNQQAQKSH
LADCSMDSCLTSCEGPQKDQETQNISIGLGYHSNSLLWQKAEREEFRVQRPNHSTGESLK
DTKHYSPSPERKMHLLTSSFGDTINNVRAPGDKGACSTNSDARRKERGFEGACGEPPRKR
SVASRTQALDIEQTEVFDRLSNHTAVLDLNAHDENDASSECKEFDLNGFSWS
>AtrPSR_scaffold00131.37
MYHKVFAYQASPFGCNQKEMNEEYCKCGKQCGSLQAPLVSFSTSKVASYGQSSAPSSLCN
TREQPLTPLEDLSVAAGETTCAYSFSLSPNLRNMCSQSSMFCPSLHSSHKEFQIGNIPFL
PYPQISDHPVSSSQSLKFMLLGHGDSSLYQYKECNNLENSMNDDDNVSPQDASGCNLVGQ
SCRGSVMTSEQADWQVVADPTGLVITNNCEDSMLDEFFVATKGSSSNTIGFQGNHMDGQF
VFPIGEQLNTGPLTSRNGTVNKTRMRWTQELHEQFVKAVTFLGGPDVATPKSILRIMNVA
GLNIYHVKSHLQKYRLARYQPGNGEDKRSPGSETKKKSSADTDKENVYVKMGAQDTEYLR
LQLEVQKKLHKQLEEQRELQLWIEANARSLQKMVEEQHRALIHATLSLSSSSPLASTSEL
KDNSEPCRPPCFGLTNKEGLSSMETDDGSTLELEIHSSASKRARME
>AtrPSR_scaffold00142.28
MEVRPAFSIRSSNTMQHNSSLGMPQGPSSSLPCLSAPLEDRQPKFPSTQQAGMGKELRPS
TVGPLTGPVASNNGVVGPIFSTGSAFSNDLHFSSSISQRQGHSSSSPFISQASSNFISQE
SSNFISQASSNSTSLSLLHSSQSGDIFQPVASDFTKESNDVSWSSDPLQGFLDFPDNVGE
SSQHNGMDGDDQTRRTDWGAWTDQLIGDDETLPAGWSDLLLDTNAADPQPKTPKLSSDFK
VQQGPLPQLAPALSGEFCSVSSPSSGNGGTNKARMRWTPELHECFVEAVNQLGGSERATP
KGVLKLMKVEGLTIYHVKSHLQKYRTARYRPDISEGNAEKKMTRLDDISTLDLKTGIEIT
EALRLQMEVQKRLHEQLEIQRNLQLRIEEQGRYLQMMFEKQSKGGVDKLKAQIASESDPS
VPSDSTLSLSKTDFSAKESVERNNATITLEEPHVNTQSDSGSQYPPAKRLKSGDPSSSIS
E
>NnuPRS_000838
MGMQKMQNQQIGLILSTDAKPRLKWTPELHQRFVVAVTELGGADKATPKALMRVMDVPGL

TLNTGLGRASSQNPTVTAGKKVRSLDLMEADYGDKEVRGGYFSGELIGRTPTPINESLKI
AQALQMQMEVQKKLHEQIEVQRHLQLRIEAQGKYLQSVLKKAQETLAGYNSSSLGLEAAK
AELSQLASMVDNRCPSSSFSGLTEAGGFSLQEVGKQSMRGMDCSMDSSLTSSDSSGRKDE
NSQKLESGHPHDCNRHSNAARLMEMYPNEGHLCKSDPSDRLHGQKRSWSTISDDISVDQP
TAKRSPTHKEKLGGQCTKLGLTDKLDLNSQYQSEMESGCQEFDLNSRVEEPIYVSKLFGA
>NnuPRS_004596
MYHAKKFSTMTLVPHKPQPQGADQLPNVGVLGGSTINNSATSGGTGKQRLRWTSDLHDRF
VDAITQLGGPDRATPKGVLRVMGVPGLTIYHVKSHLQKYRLAKYLPESPADGSKDEKKDS
GDSLSSMDSAPGVQINEALRMQMEVQKRLHEQLEVQRQLQLRIEAQGRYLQKIIEEQQKL
GGALKASEPLPLADEKQKQSTSVTAAAVEGPMSPLKKQRVDDGSSDPASSVLVPPPPSTE
KKSDFIGQWDGDLYGGNGVGFEFDVVNEFKEKEDGGAEQRATSKVGTGYGTGGSIS
>NnuPRS_004697
MRTMGVKGLTLDHLKSHLQKYRLGKQSCKEFTNSSKDESQSVVSSSSSASSRMMAQDGYQ
VTEALRVQMEVQRRLHEQLEVQ
>NnuPRS_005315
MHQPKSTSSSSLVRSNALLPRQHLNSTDSAMGSSSRGSIIETSNSSKNSPILSTRQRLRW
THELHERFVDAVAKLGGPDKATPKGVLRAMGVLGLTTFHVKSHLQKYRLATHHPELHGKM
GREARALVQLTEGSSKTDTLKVPTMALHWCTSSQIQKMDTLKLQAEVQKRLHEQLEASLF
KFVTGIFKVKFSLFPFVQRQLEVGKEDNSKPPKITEEQKSSSDVLAEASGAVTSVPLLES
EKTEACTPSPTASDSAPTSNSPNQELETPAQVDASKEHPQAKNASASKSSSPGESSGERV
VKKQRVSEGPECA
>NnuPRS_005316
MFQPKAVPSSNLASSNLLVHAQNMDNSVSAVAHCSGGSSLNNNTNSSTIASRQRLRWTNE
LHERFVDAVTQLGGPDRATPKGVLRIMGVQGLTIYHVKSHLQKYRLAKYLPESTSDVKKS
EKKEPSELLSTISSSSGMQITEALKLQMEVQRQLQLRIEAQGKYLKKIIEEQQRLSGFLG
ESNLMIPPPISGSNFPESDKTDPSTPAPTSECPLQNKAAQECAPAKSLSCNESISSYHEP
LTPDSVCHVTSPSESPKGERPLKQKRVGMDAVSAKPEMILTHQILESSLSSSFQQPCSGF
PIREQLDPSSGRSHCNGDQFENVSLRDL
>NnuPRS_005317
MFPGLIQPHEAVVPQEEIQGSNHRGDPCLVLTSDPKPRLRWTADLHERFVDAVTQLGGAD
KATPKSIMRTMGVKGLTLFHLKSHLQKYRLGKQSGKDLTEPSKDASYLLESPHNGTSSPG
MSASDLNEGYEVKEALRAQMEVQRKLHLQVEAEKHLQICKDAERRYVESMLERACKLLAG
QTISCLVPDAEGHELPGSAPKASRTSSSHEPTRFYSLRPAEAIRVQVPEEEPPNLQPPRA
DCSTESCLTTNGSPAGLSLESSPAGCKKRVRSLDTTDALIWSEASTRTQDIDATRIDPHV
ITGYGRLGEI
>NnuPRS_006931
MERTCGSYALETGMVLSRDPKPRLRWTPDLHDRFVDAVTKLGGPDKATPKSVLRVMGLKG
LTLYHLKSHLQKYRLGKQSKKGTNIEQAKSEELSVDSVGDSGYNHMHFSATSSSTSKENS
ERGIPIAEALRYQIEVQRRLQEQLEVQKKLQIRIEAQGKYLQAILEKAQKSLSTDTKCPR
SLEATRAQLTDFNLALSGLMENMNQVREDESKGEMIGKSLLNDNQENNHSSGFQLYRELE
EAKKKEIKIKVEESSLLFDLNVKGGYDFLGPKGSELDPNMNIQRI
>NnuPRS_007976
MNNRKVEYQGLNNECVSDYSFESFDRSSGDFGSRQTWNMGACFHSPIPALEGGSLQPNPR
PSALGSSTISSHYGLPTSAFYATERYMGFPQYQGARSWFSEFLKNYDQAVSSCEPSEEPF
SVDPERRDDLDLQPRDTLQSVMELPFCSNQNSRFSEISNTSSYRNFPGIEPHPLLRHKLV
GEDATLNSRHPSILFEVNQNPGVGYKPVTSPLAQTSFNVQLEKQSPIPPAVVSVASGSIV
SAGAAVSNKTRIRWTPDLHERFVESVNRLGGAEKATPKGILKLMASEGLTIFHVKSHLQK
YRIAKYMPESSEGRSERACINSMAQIDAKTGMQITEALRLQLDVQKRLHEQLEIQRNLQM
RIEEQGRQLKKMFEQQQKANKNWFETQNLDIMFPDDQSNSHEDAQVSTVGEGSEDANFTS
KTDNQNTSHPR
>NnuPRS_008138
MYHHHQGKTSFPSSRMSINVRGDTGLVLSTDAKPRLKWTPELHERFVEAVNQLGGVDKAT
PKTVMKLMGISGLTLYHLKSHLQKYRLSKNLHGQANTGINKNAVAGDRISEASGAFISNT
SLCSQTNKEYELKRISNIRSLQISEAIQMQIEVQRRLHEQLEVQRHLQLRIEAQGKYLQS

VLEKAQETLGRQNLGSAGLEAAKVQLSELVSKVSTECLNSTLSELKELQDLHPQKTQTTQ
PTDCSMDSCLTSCEGSQKDQEIHNIGVGLRSYYSNTSLGQKEIVDDSRLEQPEHAWGEDQ
NENKMFCSSKGRDTERIMFPMKRNSSDLTMSIKVHGEKWNASSFFTEATTKEGDEDDSFH
YQTGARRPVTNVENEKMSKGFRLPHLTPKLDLNSQEENDSTSTCKQFDLNGFSWT
>NnuPRS_006931
MERTCGSYALETGMVLSRDPKPRLRWTPDLHDRFVDAVTKLGGPDKATPKSVLRVMGLKG
LTLYHLKSHLQKYRLGKQSKKGTNIEQAKSEELSVDSVGDSGYNHMHFSATSSSTSKENS
ERGIPIAEALRYQIEVQRRLQEQLEVQKKLQIRIEAQGKYLQAILEKAQKSLSTDTKCPR
SLEATRAQLTDFNLALSGLMENMNQVREDESKGEMIGKSLLNDNQENNHSSGFQLYRELE
EAKKKEIKIKVEESSLLFDLNVKGGYDFLGPKGSELDPNMNIQRI
>NnuPRS_007976
MNNRKVEYQGLNNECVSDYSFESFDRSSGDFGSRQTWNMGACFHSPIPALEGGSLQPNPR
PSALGSSTISSHYGLPTSAFYATERYMGFPQYQGARSWFSEFLKNYDQAVSSCEPSEEPF
SVDPERRDDLDLQPRDTLQSVMELPFCSNQNSRFSEISNTSSYRNFPGIEPHPLLRHKLV
GEDATLNSRHPSILFEVNQNPGVGYKPVTSPLAQTSFNVQLEKQSPIPPAVVSVASGSIV
SAGAAVSNKTRIRWTPDLHERFVESVNRLGGAEKATPKGILKLMASEGLTIFHVKSHLQK
YRIAKYMPESSEGRSERACINSMAQIDAKTGMQITEALRLQLDVQKRLHEQLEIQRNLQM
RIEEQGRQLKKMFEQQQKANKNWFETQNLDIMFPDDQSNSHEDAQVSTVGEGSEDANFTS
KTDNQNTSHPR
>NnuPRS_008138
MYHHHQGKTSFPSSRMSINVRGDTGLVLSTDAKPRLKWTPELHERFVEAVNQLGGVDKAT
PKTVMKLMGISGLTLYHLKSHLQKYRLSKNLHGQANTGINKNAVAGDRISEASGAFISNT
SLCSQTNKEYELKRISNIRSLQISEAIQMQIEVQRRLHEQLEVQRHLQLRIEAQGKYLQS
VLEKAQETLGRQNLGSAGLEAAKVQLSELVSKVSTECLNSTLSELKELQDLHPQKTQTTQ
PTDCSMDSCLTSCEGSQKDQEIHNIGVGLRSYYSNTSLGQKEIVDDSRLEQPEHAWGEDQ
NENKMFCSSKGRDTERIMFPMKRNSSDLTMSIKVHGEKWNASSFFTEATTKEGDEDDSFH
YQTGARRPVTNVENEKMSKGFRLPHLTPKLDLNSQEENDSTSTCKQFDLNGFSWT
>NnuPRS_008492
MGTQKMQNQHMSLVLSTYAKPRLKWTRELHQRFIEAVAKLGRADSTGNPKSSVEGDGYPW
PYFVPPKEPFTDIVEGDYVGTEVRGGNVSGEMIGRIPTHMNDNLQIAQALQMQMEVHKKL
LEQIEVQRHLQLRIEAQGKYLQSVLEKAQETLAGCNSSCVGTRSCKS
>NnuPRS_009935
MFHSKKPTMNAHDRPMCVQGDSGLVLTTDPKPRLRWTVELHERFVDAVTQLGGPDKATPK
TIMRVMGVKGLTLYHLKSHLQKFRLGKQPHKEFNDHAVKDGDRALELQRNAASSSGIMGR
SMNENVRLTEALRMQMEVERRLHEQLEVQKHLQLRIEAQGKYLQNILEKAFQTLAGDQNM
ATASYKTVGNQGGVLDVSAMKDFGSPVNTFPSLQDLHIYGGEQLDLQSQMDRTSLDGFMP
AAANDNICLGKKRPSPYSSSSGKNPLIWADDLRLQELGGTAAACLGPQEDAFKGDHQLQI
SPSAIDNGPDMDSISDVYETKPIVSGDSMGEKKFEASAKLERPSPRRATLPVERINPMIK
SSALTQGRNMPYG
>NnuPRS_010973
MSSSLPVLPTPLEEKYPKLPDSQEVSLGREIMTNPVVPQVPPLVSNSEVVGHIFSSASGF
STDLQFSSVAHEGHSKNAPFISQSLSNGVALPLTHSSHSGLYQSGALNHYPKENSNISWT
ADQLQGFLDFPENVPTQSSQTESGNGGVMPSEDRTKRNDWQEWADQLITDDDPLASNWNE
LLLDTNVADTEKKAAYQVPKPPPNFTVHQPQIHQQLPDPSGQLCSVASPSSSXNGAPTKT
RMRWTPELHERFVEAVNQLGGSERATPKGVLKLMKVEGLTIYHVKSHLQKYRTARYRPDS
SEGSAEKKKTAIEEISSLDLKTGIDITEALRLQMEVQKQLHEQLEIQRNLQLRIEEQGRY
LQMMFEKQYKANNDEFKGSSSNPQEPSTALSDTMQQQSSTQNRVDALKVCAETGNDSIDT
SAIAEASYKKLGVKQKAPETHSCEDLEPDGIDDITIPSAIPSTKRAIADDTTIPSAKPTS
DRQANTADIPAPKDTVSMSTEDLSNG
>NnuPRS_012506
MSSSLPVLPTPLEEKYPKLPDSQQVSLEREIMANSVPPHATQLASNSGVVGHMFSSASGF
TTDLHFSSAPHERHFTNAPFISQSSSDGISLPLTRSSHSGVFQSRPMSHYPRENNNISWS
ADQLQSFLDFPENVPTQNNQVESSNSGIMPSDDNTKRSDWQEWADQLIRDDDPSTPNWNE
LLADTNVADTELETAYQAPKPPQNFLVQQPQVHQQIPVLSGDLSSVASPSSSVTGVPNKP

RMRWTPELHECFVEAVNQLGGSERATPKGVLKLMKVEGLTIYHVKSHLQKYRTARYRPDS
SEGSSEKKMSAIEEMTSLDLKTGIEITEALRLQMEVQKRLHEQLEIQRNLQLRIEEQGRY
LQMMFEKQCKSHNDRLKTLSSNLEESSAPLLDVMQHSAKNEVEPTKSCAETGNDIMDASA
IPEESSRKMSRKQKEPESHSEDLNLEGVSGSDSPPIKRVKADDRSVASAKLASD
>NnuPRS_019826
MYHHHHHHGKTSFPSSTMSIPPERHLFLQNGNGRGDTGLILSTDAKPRLKWTPELHERFI
EAVNQLGGADKATPKTVLKLMGIPGLTLYHLKSHLQKYRLSKNLHGQANMGTNKNVAGDR
ISEASGALMSNTSPSSQTNKNLQISEAIQIQIEVQRRLHEQLEVQRHLQLRIEAQGKYLQ
SVLEKAQETLGRQNLGSVEFDAAKMQFPELVSKVSAECLNSTFPELKELQGLCLCPQQTQ
KMQPTDCSVDSCLTSCEGSQKDQEIHNIEVGLRPYQSSASLGLKEIVDDSRIEQPEPTWG
EDLNENKMFSSSIGRDAERIMFPMQRSCSSLSTTSRVQGENRNASSSFAEARTKEGNEDD
SFLNQNSARRPVANLENDMSKGFRLPHLTAKLDLNTQENDSTTHKQFDLNGFSWT
>NnuPRS_021120
MDANCASACPPLFIRSESLNSPSRPPAPMFQLQKISPDPEPSSPVSYESYPQYPTSVFSR
SSTFCTDLYISSSTSSETHRHLGNLPFLPQPQTFDQPIATVQSSKSPLLLSGDSSNQTEE
KRSEDLMKDFLNFPGDASDGSFHGANFASDSLTFTEQLDLQILSAELDIAMTDNSENPRL
DEIYEAPTPDSRLKCKPFHQPIHPIQPVVPPVAGQIHSSPSTPGAAAAHKPRMRWTPELH
ERFVEAVNKLDGAEKATPKGVLKLMNVESLTIYHVKSHLQKYRLAKYMPETKEDKKASSS
EDKKVASTSKESDARLRRGIHITEALRLQMEVQKQLHEQLEVQRDLQLRIEEHARYLQKI
LEEQQKAGSTLIPACELPSLSKDPEGQPVPPSACALSQQAESKTESPSLPAKHKATDCAE
HEQLACQKRLCRETSPEIAEAKPEVTSDEHVVENPVP
>NnuPRS_021405
MERRCGSYSYETGVAVGVVLSRDPKSRLRWTPDLHDRFVDAVTKLGGPDKATPKSVLRVM
GLKGLTLYHLKSHLQKYRLGKQAKKEANLESRSKSSVSDGKADGHNHMDSSVISSSAPTQ
NNQEERKIAETLRYQIEVQRRLQEQIEVQKKLQKRIEAQGKYLQAILEKAQKSLSTDRKC
PRRLEATGSQLTDLSPSGFMTKLREDERKGEMMRKNLLNDNHDSNIQRSSFQLYQEVGEA
KNKENMIEGDEGSSLFDLNVKGGYEFFGPKGSGS
>NnuPRS_022348
MYSGIHSLDGSVGHEDLQGSLEGTNLPGDACLVLTTDPKPRLRWTAELHERFVDAVTQLG
GPDISHLKLIIIFPNSFAEATPKTIMRTMGVKGLTLYHLKSHLQKYRLGKQSCKEFTESS
KDASSVAESQGAVSSSSSASSRMMAQDLNDGYQVTEALRVQMEVQRRLHEQLEVQRHLHL
RIEAQGKYLQSILEKACKALNEQTIASVGLEAARQELSELAIKVSNDCLGMVPLDTIKMP
SLSEIAAALEEKSASNGPTRIGDCSVDSCLTSTGSPVSPMCMGSQAAMMKKRSRPMFGGD
SLAWESDMRQDVEWMLPGIR
>AcoPSR_Aqcoe1G011200
MYQARKFSTVGLVPHKAQSAEQHTSIGSLGESAKNDSGTSGGNGKQRLRWTTELHDRFVD
AITQLGGPDRATPKGVLRVMGVPGLTIYHVKSHLQKYRLAKYLPESPADGSKDEKKDSGD
SLSSMDSAPGIQINEALRMQMEVQKRLHEQLEVQRQLQLRIEAQGKYLQKIIEEQQKLGG
VLKASETMQVSKELQQPSTPPSNPRVDPEPPLKKHRGEDKTLEPAPSAGPPRTDKKPDFV
SPWDQDYGINVVGGFEFGVASDFKIGKDGEPQRTSQVAGPAYGPGGSSS
>AcoPSR_Aqcoe1G017700
MNTQKIGCYPLEFSGCSTDNSHPWPWNMGVCCQSSSSASQAGLQLSNLGSANSFTVASDH
LGSEASVFNTAEHYYQLGTPMFSQLPKSNTYISSHSPFEDYLPVESERGVDFDIQSRDTL
QSIVKYSLPNNQNPRPSENSYRTCRSLLGSEHSSPWQNKLVIDNAALNRKNPSNPSEGSH
NTRVAYNPFASELDPPSFQLIPDKQSAQTSPGVISYTSASPVSSRAAVSNKTRIRWTQDL
HERFVECVNCLGGADKATPKGILKLMDSDGLTIFHVKSHLQKYRMAKYIPESAEGKSDKK
VPTSDTIKLDPKAGLHIAEALRLQLDVQMRLHEQLEIQRNLQVRIEEQGKQLKKMFDQQA
KNSDLMCPDVQSTNQEDVSLEVSADKLASQNT
>AcoPSR_Aqcoe1G159000
MFHCKKPTMNSHERSMSIQGDSGLVLTTDPKPRLRWTVELHERFVDAVTQLGGPDKATPK
TIMRVMGVKGLTLYHLKSHLQKFRLGKQPHKEFNDHSIKEALDLQRNASSSSGLMGRSMN
DRNVHISEALRMQMEVQRRLHEQLEVQKHLQLRIEAQGKYMQNILEKACQTLASDNMASG
SYKSVNNQEVSDMSNMKDLSSSMSFPSLQDLHIYGGEQLDMQQQMDRSSLDGFSPINDNI
CLGRRRPNSYGSTSKNPIIWSDDIRLQELGMSVPCLGSQDDSFKNDQIQAASLTIDSGNE

MDNIGDGYETKPILPGDSIGDRRYDASSKLERPSPQRGSLSMNRVNSMIKGGGRNYS
>AcoPSR_Aqcoe1G374700
MSVKGLTLFHLKSHLQKYRLSKHSEKTKTAHSASYVMDSPGTDGSFESFSSSELNEGFDV
SQALRKEIEGQRKIELQLEAEKSIKDRVSSEQRYLEYVLERACQQHYQILGAADFAANIP
KQPEIPPTKMPSAIGTWNPLGFYTFSSQIGPAEAVRQQVLPGEEFPTLYTNTVNYSPNSY
LTATPVIFSSGTTSAEGMNTEFDHRIATRGMHMAADDSFFWGAAAKGIPEFCAHIMKQGG
GGGGGGGFKGYSGCPNY
>AcoPSR_Aqcoe1G374900
MQIMSVKGLTLFHLKSHLQKYRLSKHSGTTKTAPSALYGMDSTGTDGSFESFSSSELNEG
FDVSQALRKEIEGQRKIELQLEVEKSIKDRVSSEKRYLDYMLERASQRHYQTLRVANCVT
NIPKQPEIPSTEMSSTIETWNPLGFYSFSPQIGAAEAVRPQVPASEEFPTLYTNTVNYSP
NSYLTATPVILPSESTSAGGMSTDFDPRIAKGGMNMAADDSSLWGVAANGIPEFGAHIMN
QSGGGGGFNGYSGCPPY
>AcoPSR_Aqcoe1G443600
MYHHNHHNGKNSFPSTRMSIPQERSHMFLQGNGNACGGDAGLVLSTDPKPRLKWTPELHE
RFIEAVNQLGGADKATPKTVMKLMGIPGLTLYHLKSHLQKYRLSKNLHGQANPGNNKTGF
MGMVAAGERMCEGNGALTTGSQPNKNLQISEALEMQIEVQRRLNEQLEVQRHLQLRIEAQ
GKYLQSVLEKAQETLGKQKVGGLDDAKVHISQLVSKISTECLNPGPPDCSLDSCLTSSCE
GSQKDQEMPCITLTPYHHHNDNTNNAVVGLTEEHHLQANEALWCQNIKETKLFPSCSLKK
DTEKTMFPAQKSSSDFSLRIPSQREKENNGGSISDMRPKAGDSIQLGNENKSKAFGLPYL
TSKLDLNVHDDNDTASQLDLNGFSWS
>AcoPSR_Aqcoe2G092800
MVSNRSGSDSSGKQRLRWTQELRDRFEEAVNQLGGADRATPKGILKTMDFPGLNIYHVKS
HLQKYRISKFIPETYDRGRLDRRKVSELLPNFSTTSAAQINELLQTQIEVPRRTIDQTEI
QRNLRFKIEAQARYFENIAAEEHRNLSGKDTKVYKSTSSLPSLCEESELTARDHECNLEV
SDKEEDHFKGNIKPSSIIQRAPEQYSLPNIIEGRMRNDGVQPHKRLRVENYTIFPTSRFE
LPSCSLEHNQAKMVDKTFFVKDLPSSDSTYDTTFPWPVASAKSTSTSSYVFR
>AcoPSR_Aqcoe2G381900
MEGRSSVTIQRPSNQAGNVGVSGCMSSSLPVLPLPLEEKYTKFSYSQQICFERELMSDPV
APFDTPLASTGENVGHKISSTSGFCTDLHSSSSSTHASHLRNSPFISQVSNNGVSSPIAF
SSHSGIIRSTPSSSCNKRNNNASCGNILDQKSQIENSNGATASEEHTSQSDWQQWADQLI
TDDENLAADWDELLLDTNAVDSEPKTASQAPKQSSVISPHQERTHQQPSVQSRELAMVTS
PLSPVCGSSIKSRMRWTPELHECFVQAVNQLGGSERATPKGVLKLMNVEGLTILHVKSHL
QKYRTARYGPDLTDGTSEKKMSAAEETPSLSPKMSIEISEALRLQMEVQKRLHEQLEIQR
NLQLRIEEQGKVLQMMFEKQCKKVGDNFNLSTSTSTSTLNEPSTQNLPLHDAKSEPEATE
SDHDTKHGPINDTITVENSHNNGKEQKMLEIEVIEVCALDMGEGSDKSPKKRARTDDTTL
SSGKSASD
>AcoPSR_Aqcoe2G398700
MYSSIHPLECHEDLQGSLDACVVLTSDPKPRLRWTAELHERFVDAVTQLGGPDKATPKTI
MRTMGVKGLTLYHLKSHLQKYRLGKQSCKDFADSSKDASCIAESQGTESPSSSSSSRMMA
QDLNDGYQVTEALRVQMEVQRRLHEQLEVQRHLHLRIEAQGKYLQSILEKACKALNEQPV
ASVGLEEARQELSELAIKVTNDCLGAVPLDKLPSLAEIAAGCLDEKFLSNVQARIGDCSV
DSCLTSSGSPLSPMGVGMQSATIKKRPRPMFASGESLAWESDMRQDIEWIPTSI
>AcoPSR_Aqcoe3G208000
MEFGNSNSYETAGGVILSRDPKPRLRWTAELHDRFVAAVTNLGGPDKATPKSVLRVMGLK
GLTLYHLKSHLQKYRLGKQTQRDTKLDQSLDSAESSEGHSYMQSSGTTTTTPKVNNEGGF
QIAETLRYQIEVQRRLQEQLEVQKKLQMRIEAQGKYLQTILEKAQKSLSSDINCPENLEA
TRDQITDFNLALSGLMENMTQVNEEERKGDAIGKNSTVQHKKPHSSAFQLYQGEEEEKER
KEVKVKVEGGSLLLDLNMNGNYEFMGGNGNGNEYDTNIHTHRRQGWF
>AcoPSR_Aqcoe3G226300
MEQAMMLNNKQNQQKEQGNQEPPSVILTSDAKPRLRWTNDLHRRFVDAVMQLGGPKHAKP
KALWRIMGVKGLTLYHLKSHLQKYRLSTHLKKDWAEPTKTATSTSYQVDSPRTNGSFGSF
LSSELNEEFDVSEALRKEIEGQRKIQLELEVEKRIQDRISSEQRYLEHMIERACQQHNQT
LRAAGGASNIPKQPEIPPTEVSAMRTWNPLGFYTFSSQTGPVEAVRPQVPVGEGFPTVHT

NTVKHSPNSNLTATSLIFPNETSLAGGMNTELDPRIPTGGMNLAGDDYFFWGVPDNGIPK
FDAPIMNQGGGGGFNGYSGCPTS
>AcoPSR_Aqcoe3G342000
MSLQKMQNQEMDLALSSDAKPRLKWTPVLHQRFIEAVTQLGGAERATPKSLMRVMGIPGL
TLYHLKSHLQKYRLGKSQLSESCDDNEQEDNRENEIEGGHMNVKVNEKIHAQSDEGLHVT
QALKMQIDVQNKLHEQIEVQRHLQLRIEAQGMYLQSILKKAQETLSGYNSSSTELEAAKT
EVSELVSMVETGCLDSSFSEFTKTGCFVLDNEGKKLVRTTDCSLDSSLTSFESYGEIRGS
NFHWRSIQVIEATMQQ
>AcoPSR_Aqcoe4G246400
SNQNPKPSENSYHTCTSLLGSEQSSPWDNKLVTDNTALNQKYSSNPSDNTRVASNPFASE
LDPPSFRLIPDNKQSSQTSSDVISYTSASPVSSRAAVSNKTRIRWTQDLRERFVECVNCL
GGADKATPKGILKLMDSEGLTIFHVKSHLQKYRMAKYIPESAEGKSDKKVPTSDTIKLDP
KTGLQIEEALRQQLKLQMYLHEQLKIQRNLQVRIEEQGKQLKKMFDQQAKNSDLMCPDVQ
STHHEVARLEASSADIHL
>AcoPSR_Aqcoe5G018400
MEVRSALSIRGSGTKQLVDSGISVAMSSSLPVHPIALEKQYSQFPNSQQVSEERGLLTNS
LVSHSTLLATNSGSVGHIFSSASGSSSDLHFSTVSSHENDSRNTSFNSQFLNNGTTFSFP
HSSQSEIFGPTDLGNYTKENNNVSWGEDSLQVLLDFPENVPVENRQLENSGTDIMKQSDW
QEWDQLITDVEPLGSDWSEILANTNANQSEQKPATYEPIASSNSSLHQPQIQQQRPVDSG
DSNPAAVPTSSANGTSTKPRMRWTPELHESFVEAVNKLGGSERATPKGVLKLMKVEGLTI
YHVKSHLQKYRTARYRPEPSEGNSEKRATTIDDISSLDLKTGIEITEALRMQMEVQRRLH
EQLEIQRNLQLRIEEQGKYLQKMFEEQCKSTADRQKASSSTKDEASAPLSDLKQQSPSKD
ELKAKVNDLPEVGTEVLESRATLVECSQKDRGNQKASSHEDSGDVGADVSESTRSHSPPA
KRSKVDDRATSSANSALN
>AcoPSR_Aqcoe7G341000
MESNSGPNTLSKNPSLNSRQRLRWTNDLHERFVDAVAQLGGPDRATPKGVLRMMGVQGLT
IYHIKSHLQKYRLAKYLPDAASDGKGEDKKESGDLSSMDNSSGMQITEALKLQMEVQKRL
HEQLEVQRQLQLRIEAQGNYLKKILEEQQRLSGTLAEPHGLNSPAPANNNSPALDNKSDP
STPAPTSETPLDKGSKDSAPAKFPALDDTFASFHEPLTPDSSCQGGFSVESPKGGRSAKK
QKRNMSTSHAQPQMAFQHQILESSLSTGFQQQAASALPIKEHFDHSGDSYSDDDRL
>SlyPSR_Solyc06g066180.2
MGSNLYENDGVVMTRDPKPRLRWTAELHQQFVDAVTKLGGPDKATPKLVLRLMGLKGLTL
YHLKSHLQKYRLGQLAKKQNAAEANKENSGDSSGQFGLHSSGPSTSSLSMNFMQGEVPTT
EAVMSQIQVQKILQEQLQVQQKLQMRIEAQGSNNLSSPCFRFTSVKFNGLWREPGLQHRA
TNV
>SlyPSR_Solyc04g015290.2
MFPRLIQSQEDEFIHGCDVNVGIHHHNRVNGDPCLVLTSDPKPRLRWTADLHERFIDAVT
QLGGPSKATPKAIMRTMGVKGLTLFHLKSHLQKYRLGKQSQKDLDEASKDGLTATYSLES
PCSGGTPQQLPASDLNEGFEVKEALRAQMEVQSKLHLQVEAEKHLQIRQDAEQRYITMLE
KACKMLADQFIGDVVTENHQETYQGLNTKTQLSPLCNPHGLCPSESADFVGVHGPEDVSP
RIHPQRTDCSTESCLTSHESPAGLPVEGSSPGGKKRGLSGDSTQASYVWGEADMRSSGVH
VLPVNCFGISGSNVQNVSN
>SlyPSR_Solyc05g007890.2
MALHQNAQQNKDMNLVLSSDAKPRLKWTPDLHQRFVDAVSQLGGPDKATPKSLMRVMNIH
GLTLYHLKSHLQKYRLGKSSVTDQSFDENKQEVKLDLCEIVPNDDDTKGNEISRGLKQRT
YFDTELPQSDHKLSLGVCDGSQNHMNESFQIARALQMQMDVQRKLHHQIEVQRHLQLRIE
AQGKYLQSVLKKAQETLAGYGTSSGVELAKAELSQLVSMVNMGCCPSSSLTEIDCSISKD
IENKTSKEGILCSIESSLTSSESSARKEGQNNSMNNKNTTCIGLPLNQESKGKKRGRHNI
CGDEQSSAKRYLETIDLNRKCPNEYDENVTKVIDLNEYS
>SlyPSR_Solyc05g055940.2
MNTRRALGIQRSSETQHSNMGVCGAMSTSLSILPSFEEKHLKVTDSLHVLTEKEQTTNLI
SSRAIPSASKSGTVGHLLSSTSGPHKHFHFSSTPSRESRKHNYPFISSEASAATCQSSLS
NIHSTSLDNYPMENSNNSWGEDAYHDCINFSTNVPVQNDQVESLAVVMTSDDQVKRSDWQ
DWADQLINDDDVLDSSWSNILVDINPPDTKFLRASEVHAGQSDTLPPPLPAAAASSGQNY

PVGSPSSTAAQTKPRMRWTPELHEVFVEAISKLGGSEKATPKGVLKLMNVEGLTIYHVKS
HLQKYRTARYKPESSEGTPEKKTTSVTEMPSLDLITTMGITEALRMQMEVQKQLHEQLEI
QRKLQLRIEEQGKYLEMMFEKTKDIGKDLKVSSSSRTDEHPSPSNKMKHTSPND
>SlyPSR_Solyc06g008200.2
MEARPALSVQRTGAHQLSNFGASGAHSSLPVLQTSLEEKYPKLPGSALSSFPSNSGAVGH
LFSSPSEFSTNLDFSSVLPYDEHLAPATFISQPTINEASIPLANSGVLQGAASSQYVNVN
SESWCTELLPDYLDYSVNAPVQNTQLDCRNSDDCQIPPEDPSKQSDWPEWADLVMSDDDA
LTSSWMVDTSIADAEPKMQYQEQNQLSNFPVHQVQPFQQIPTASVETSAVVPASVETSAV
VPASVETSAVVPASSTGSGSSSKQRMRWTPELHEAFVEAVNKLGGSERATPKGVLKLMKV
EGLTIYHVKSHLQKYRTARYKPEASEGSSEKKESSIGDLSALDLKTGIEITEALRLQMEV
QKQLHEQLEIQRNLQLRIEEQGRCLQMMFEKQCKSMPGIDLAKGSSSMAEDASAQLNDAV
QSSSNKNDPGASLVGHHEAGSHTTQQLVREKQKEHEREDITSSSNSPPTKRAKVDE
>SlyPSR_Solyc06g066340.2
MRPIRGIPLYNQNPLPLPLPLSHHHNNYPIFAHQTFENTAPTTTPTLPIHSSGYCISSFS
NNSNKTIPNSSPTTIPFHHSNHQGGLMRSRFLSRFPKRTMRAPRMRWTSSLHARFVHAVE
LLGGHERATPKSVLELMDVKDLTLAHVKSHLQMYRTVKTTDRAAVPASSGQSEVFDNGSS
GETSEDLMPDMENSKKPDLSDQQGKNSMHLQEIDYHGLWSNSSSRESWQLHGKHGDYPGN
IPSLEKTVKNQHIDLEAKCLSYDRLSGEVSSSSITETSPKKPNLEFTLGRP
>SlyPSR_Solyc08g076010.2
MTITHLLVSSIWEDVARDCTNILFSSFFAEYSTLSPTCHFSIRSSRAHISQNSRHRLPLV
FSDLWKSFEDFEEFGSGGFMDRMYNGGGDMGYGYENGVVMTRDPKPRLRWTADLHDRFVD
AVTKLGGPDKATPKSVLRLMGLKGLTLYHLKSHLQKYRLGQQTKKQNAAEQNRENIGESF
RQFSLHSSGPSITSSSMDGMQGEAPISEALRRQIDVQKRLHEQLEVQQKLQMRIEAQGKY
LQAILDKAQKSLSTDMNSASAVDETRAQLTDFNIALSNLMDYVHGHNEDETSAGKRIQDD
TNKDLQRSTYLTEGEQKKIMNIKLEESSVSFDLNSRSSYDFIGMNSAALEAKQFSNGRLE
I
>SlyPSR_Solyc09g072830.2
MEARPAVSVQSVVASQLSNCGASGALSSSLSIFPTALGEKYPKFPDLQQASMGKELKQHP
ATVVSSLPSNSGAVGLMFSSSSGFSADLQFSSVSPQEKHSGTAPFISQSTYSETSIQLPH
SGVLQSTASSQYLNENHEPWCIDPLPNFLDYSMDNPVQNSQVASSNKQSDWQEWADLVLN
DEDALPSDWNDIIADTSIGDSELKMQFQEEKQPLNFPMQQVQASQQIPPVPVETSAIAPV
SSPASSAATKQRMRWTPELHEAFVEAVNKLGGSERATPKGVLKMMKVEGLTIYHVKSHLQ
KYRTARYKPEASEAGSSEKKQSSLDDLASLDLKTGIEITEALRLQMEVQKRLHEQLEIQR
NLQLRIEEQGRYLQMMFEKQCKSMPGADLAKGSSSSTADDAFAQLSQVDTAKEVDHEKQK
EREIEVLGDPETNITSTSDSPPLKRSKLDE
>SlyPSR_Solyc09g091880.2
MYSTLPIRNLVIEENGDFFNHHMNHNLQASLSVDGTNLPGDSCLVLSTDPKPRLRWTTEL
HERFVDAVTQLGGPEKATPKTIMRTMGVKGLTLYHLKSHLQKYRMGKQSAKEATENSKDV
SCPAESQETGSSTSGSSRVIVQDINEGLQVTEALRVQMEVQQRLHEQLEVQRHLQLRIEA
QGKYLQSILEKACKAFNSQSLELNGLEMNREELSELAFINGPSLPDIGPSFENKNACNIP
AMLGDCLLDDCLPSNGISSLKKRPRGFTNVNGLPMESNTREVEWMMSNI
>SlyPSR_Solyc10g078720.1
MSIQNYELASDCNLEFPQMGFCFQPESSVENGCQQQQQPNFWPSTDSSSSRTIISRIGSS
PSAFFATERYLGLTQYDYQDNNNSCTQLSKNLDPQTTSYTQQCGNGFSADSSARVDTDFP
KISMPSFIRSQFSSSQPFGPEGLYGNPFSNLSEKERILLLKSKLFREIDSSNRQPASIPF
QGNQDYGVSNNTCGFNLVHIRQQSGSQPANSFNNSGCSGGSLSSKARIRWTQDLHDRFVE
CVNRLGGADKATPKAILKLMDSEGLTIFHVKSHLQKYRNAKFIPESTEGRSGKTDCPNNV
TQIDSKTGMQIKEALHMQLEVQRRLHEQLEIQRKLQLRIEEQGEQLKKIFEQQQQTTRSL
LETRNSSISSPADQFTPHEDEVFAAESFNNTHFQSNISYNDM
>SlyPSR_Solyc10g080460.1
MSIQRFIITQGNEYFSSDYSIEVSKNSAQNLVDQEIQYPSYMGTCLPIASSSSSSIINCI
GSPSSAFFATERCLGLTQYDNQYDTSELIKNCDVQMSSFDPQQCKNGILKDPLVQAEPDF
RHEISMPSFIRTEFSTSPFSDVSEAEKESLLHLKNELLGEFDTSYRRHPSLPFHGNQDYC
LSHDLCCSQLANTRQQPASPSLTFHNSASSGVFHKPSKTRIRWNEDLHDRFLECVNRLGG

ADKATPKQILNLMDSDCLTLDHVKSHLQKYRNAKHPESVGKSEKRNSSDAMTDIESKTGR
EIKEALKMQLEVQRCLHEQLETQRTLQMRIEEQAKKLKMILDQQRKTNMTLLGTRNSNIS
SPGVEILVVQDSDN
>SlyPSR_Solyc10g083340.1
MYHHHHQAPNMHPSTRMSFPERHLFLQGGNANGDSGLVLSTDAKPRLKWTPDLHERFIEA
VTQLGGADKATPKSVLKLMGIQGLTLYHLKSHLQKYRLSKNHHGQANISGVNKAAASMEK
ICESTGSPKSNPSIGHQPNNNIPISEAIQMQIDVQRRLHEQLELRIEAQGKYLQAVLEKA
QETLGTQNLGTIGLEAAKVQLSDLVSKVSNQCLNSAFSEIKELSGFHTPQTQATQRLADC
SMDSCLTSSEGPLRDLQEMHNNQLGLRNLNFRPCTEEIENQTRLQQTALRWRDDLKENRL
FPKIDEDTEKEFAKETNWSNLSMNVGIQGGKRNVNSSYVDERLNGIDADIKLFHQTATDR
SDSTKPEKQVSPQEYKLPYFAPKLDLNTDDQTDAASNCKQLDLNGFSWN
>SlyPSR_Solyc10g085620.1
MSVPERHLFLQGGNGNGDSGLVLSTDAKPRLKWTPDLHERFIEAVNQLGGADKATPKSVL
KLMGIQGLTLYHLKSHLQKYRLSKNLHGQANASGANKAVAAAGVERISENSATCMSNPSM
VPQPNKNIQISEAIQMQIEVQRRLHEQLEVQRHLQLRIEAQGKYLQSVLEKAQETLGRQN
METVGLEAVKVQLSEFVSKASNQCLNSPFTDIKELSGFHSQQTQATQPTDRSIDSCLTSR
DGSLRDNTMHDNQIGLRPFGFTPSIECKDIENDTRLQQTELRWCDNLKENRRLFSPMNEG
REKTFTRETNCNNLSMSIGLQDEKLNGSMNHSDGNFNGTERDVKLFHQVTNRSESVPQRH
KSSQEYKLSYFEPKLDLNMHDETDAASSCKQFDLNGFSWS
>SlyPSR_Solyc11g022470.1
MDPANEGNNLNNNPSLASKQRLRWTHELHERFVDAVAQLGGPDRATPKGVLRVMGVQGLT
IYHVKSHLQKYRLAKYLPDSSSDGKQSDKKESGDMLSSLDGSSTGVQINEALKLQMEVQK
RLHEQLEVQRQLQLRIEAQGKYLKKIIEEQQRLSGVLSEVPDSGVTPSAAGDNGLDSDNR
TDPGTPAPTSESPHIDTSVQEHGRSKSLSIDQSFSSQHEPLTPDSGCRETSPINSSEGER
SSKKQRVGTFTKADMLLPHQILESSLSSRYEQPNPVFVAREQFNLSSGLSLGNEGPNVRG
SNI
>SlyPSR_Solyc11g067280.1
MDPSCGGNNSSSLASKQRLRWTNELHERFVDAVAQLGGPDRATPKGVLRVMGVQGLTIYH
VKSHLQKYRLAKYLPDSSSDGKNSDKKEPRDMLSSLDGSSGVEITTALKLQMEVQKRLHE
QLEVQKQLQLRIEAQGKYLKKIIEEQQRLSGVLSDVPGSGVTALPTGDNGPESDSRTDPG
TPAPSSEAPHVDKPVNAHTSTKSLSMDESFSSHHSPLSPDSDCQETSLMESPNGESSSKK
QRVGNVALPHQILESSVSPPYQQPHSVFMMRDQFNHTSGLSLDIEDHKVSGSNI
>SlyPSR_Solyc12g017370.1
MFHAKKPSTMNSHDRPMCVQGDSGLVLTTDPKPRLRWTVELHERFVDAVTQLGGPDKATP
KTIMRVMGVKGLTLYHLKSHLQKFRLGKQPHKEFNDHSVKDGDRATSLELQRNSASSSGM
IGRNMNEMQMEVQRRLHEQLEVQRHLQLRIEAQGKYMQTILEKACQTLGGEENMSLPTRT
FKGIGNHQGGLIPDISAAFKEFGTPPLTFSSFQDLNICGEHIDLHAQSSMGERSSSFDGF
MNLSNTSTDHNLSLGKKRASPYNTSNGKSPFMWSDDFRLHELGSNNEDDHQIIQMERSCN
PEIDSVSDMYESKPLLQDDKKFDTKPERPSPRRAAQVSSLSAQGGRNSVFG
>SlyPSR_Solyc12g098370.1
MKMYSSLGIDGNGGVNEYHHHHHHLPSLQSSLSGEMTNLAGDACLVLTADHRPRLRWTAE
LHERFVDAVSQLGGPDKATPKTIMKAMGVKGLTLYHLKSHLQKYRLGKQSKEAAESYKDE
SCVAESQDTSPSASGSSKVVAQDINDCGYQVTEAFRVQMEVQRRLQEQLEVQRHVQLRIE
AQGKYLQTILEKACKVLNYTSVESPDLDTAREQLSELAIKGATNNCDGIVPVSSLPEVVT
SFENKNASDMPASIGECLSSTTTTASPTSISALKKRPRALANGDVLPVENNMTQVQWMMT
SS

>MguPSR_Migut.A00576.1
MERMYGGGGGGGGLMMTRDSKPRLRWTSDLHDRFVDAVTKLGGPDKATPKSVLRLMGLKGL
TLYHLKSHLQKYRLGQQQQQAKKQNVLDHNRENSESSYGHHNMRIASTSANSSSMNSEQG
DIPIAEALRCQIEVQKTLQQQLEVQKKLQMRIEAQGKYLQSILEKAQHSLSTDINQSENL
ESTKAQLTDFNLALSNFMQTINGDEINGNKEARETRDGHIKLETEGPSIEFDLNSRSSYD
FIGINNRAVLEANQFQNR

>MguPSR_Migut.B00920.1
MLGLKNNPVSEDFIIAGAAHIYLKLRFKIHIIFTKATPKSVLRLMGLKGLPLYHLKSHLQ
KYRLGQQQEAKKQNVLDHNTENSGDSYGHHNMRIASTSANSSSMNSEQGDIPIAEALRCQ
IEVQKTLQQQLEVQKKLQMRIEVEGKYMQSILEKTQHSLSPDINQSENLELTKSQLTDFN
LALSNFMQTKRETI
>MguPSR_Migut.D00466.1
MEARPALSIQRSGGRQISSFGASAALSPSFPARPAALQESSPKFSDSQYISVERELMHRQ
PAPLSSNNGVVGHIFSSSSGFSTNLHFSTIEQQERHPRQPPFISQSTSSGKSVMLAHPVD
SRVLQSTASSQFNKENNDSWCTETLPDFLDFPMNTTILNNELDVSTNGGIPVPSEDLGKP
SDWQDWADQLITDNDALATDWNGILADATVADPAPKPSYQISEQSPNISMNQSQISLHLP
GTPGEICTSGGQSSSANAAPAKQRMRWTPELHEAFVEAVNKLGGSERATPKGVLKQMKVD
GLTIYHVKSHLQKYRTARFKPETPEESSEKKLASIEDLSSLDLKTGIEITEALRLQMEVQ
KRLHEQLEIQRNLQLRIEEQGRYLQMMFEKQCKSGVDLIKGGASSTSENALKELADAVQN
SCAKDNLLVEAENNETKGETAKTSGETSQLVGEKQKAVEAVVLENPEPSAGEKSESPPTK
RAKVDE
>MguPSR_Migut.E00086.1
MIKTKEMRLVLSSDAKPRLKWTTELHHRFIEAVEKLGGPDKATPKSLMRIMSIHGLTLYH
LKSHLQKYRMGKNQHSQTYQQSKQEECEEKQRNVFTSKICDGAKEQINESVEITQALQMQ
MEAQRKLHEQIEVQRHLQLRIEAQGKYLQSVLRKAQETLSEYSSCSIEVEHAKAQLSQLM
SMVDSGCTSSSFSILTQSDGSVLEKDERDKLLVHDSLESSLTSSESSWKKNKETRRKHRN
GKITKEKGNSIALPLMEMHSSKRVQLLEKIDLNSDTMNEFDQGRKVIDLNINGVEFFNGN
F
>MguPSR_Migut.F01810.1
MAPALFRKKSSEAHSNDVQDSPRPNFVQELTTSLISSCPNTLSSNEKLPWNRDEIENFLE
IETDSNSREVTCTGIVASGNRTNRNDWREWAHQITTVDDSLNSSLTDLLIDVNVPDLDAK
MLEFHGPVSTVQSCGAVVSSSNCSSPSTKARMRWTPELHELFMDAVNKLGGAERATPKGV
LKLMNVEGLTIYHVKSHLQKYRTARYKPESSEGALERKSKNIAEMTSLDLKTTMGITEAL
RVQMEVQKQLHEQLEIQRNLQVRIEEQAKHLQMMFDKQRKMEEDKSKATSSNLDTIEYQV
PDGNVKAESCGQDRASSPEIIATGNCILADENSNEKEISSKREMPEVEESGGGECSPPPP
LKRAKVDEINGETGR
>MguPSR_Migut.F02036.1
MHSPLEEHFPELLESPRVTLANEPACNMHKPVGQFCLPSSEMMNSISVSPNDGFDNSDGQ
WNADSVEYFLDNPFNSIPVENGQRDNTADLAMSENGVEKTDWPDWAAQLIAVDDDETLNS
NWNDLIDIDVPDPDPKLLDLPPDVSTFQPQIHQLHHHPMPVGDAYGGLNSPNAPTAKSRM
RWTPELHEVFVDAVNKLGGSEKATPKGVLNLMGVETLTIYHVKSHLQKYRTARYKPESSE
GTSEKKSSTPTDMTSVDLKTTMGITEALRVQMEVQKQLHEQLEIQRNLQLRIEEQGKHLQ
MMFEQQRKMEEDKTTRASSSNALLSVEKQPSLCTDKSEPSDNNRATTKEVATDVSISADE
NPSKKEMLSETKRANDPFDSVTKRARVDETGTVEVTRDT
>MguPSR_Migut.G00679.1
MRTMGVKGLTLYHLKSHLQKYRLGKQANKELTENSKDASCIAESQDTGSSTSASSKTMAQ
DINDGYQVTEALRAQMEVQRRLHEQLEVQRHLEIRIEAQGKYLQSILEKACKALDEQTVV
PDGGLEAAREELSELAIKVANDCNNNNGLLSYNNMNNKNVPIMPSRLGDCSTDSVLISNG
APFSQSGFDPAFKKRARPLFSNGDSVMPLDYNSNMRAVEWMMSNIV
>MguPSR_Migut.H01640.1
MSVKGLTLFHLKSHLQKYRLGKQSGKEFGEASKDGSYHLDSPRASTPPQNLLSSDMNMGY
EVKEALRAQMEVQSKLHLQVEAEKHVQIRQNAESRYMAMLERACEMFANQILGAADTSND
GDGYEGKGTESCLTSHESCVGFLPEGSSKRRVINTGSANASFVWGESDAYTPDLHLVQVS
NSHGIAGCGV
>MguPSR_Migut.H01641.1
MDSGGNNSNLASRQRLRWTHELHERFVDAVAQLGGPDRATPKGVLRVMGVQGLTIYHVKS
HLQKYRLAKYLPDSSSDGTKAEKKESGDILSSLDGSSSLQITDALKLQMEVQKRLHEQLE
VQRQLQLRIEAQGKYLKKIIEEQQRLAGGLSEAPGSGVPAPATDEICPESDNKTDPATPA
ATSEPPFLEIPIKERASIDESFSSHHEPLTPDSDCQVGPPLEIENERPTKRPRGSGDVAF
TKSEIVQTHSILESSLSPPYQQQNSIFLTGEQFDGSSVLSIGSENQLERISGSNL

>MguPSR_Migut.H01655.1
MFHQKKPPNMNTNNSNNNSSHMFVDGDHSGLVLTTDPKPRLRWTVELHERFVDAVAQLGG
PDKATPKTIMRVMGVKGLTLYHLKSHLQKFRLGKQPHKEFNDQYSMKDVERASSLELQRN
NTSSSGIAARNMNEMQIEVNRRLHEQLEVQRHLQIRIEAQGKYMQTILEKACQTLSSGES
NAAAAAASASSSGMYGFSAAKNFLPPPPPENNFPYLQDLNIYGGGDHHQIISNNKSGKSP
LIWADDFRLRELGQLSAVDDDDGGGDERHQIQFTSLSIDHTASDININ
>MguPSR_Migut.J01031.1
MYHHHHHHHHHQGKNIQASTRMSIPSERHLFLQGPTSSNNNGDQSGLVLSTDAKPRLKWT
PDLHQRFIEAVNQLGGADKATPKSVLKLMGIQGLTLYHLKSHLQKYRLSKSLNGQANTVS
NKSATVDRVPGEPNGVHITNQNTAPPLTTTNKNMHIGEALQMQIEVQRRLHEQLEVQRHL
QLRIEAQGKYLQSVLEKAQETLGGQNMGTVGLEAAKVQLSDLVSKVSTQCFSSAFSDMKE
LSDLCMSQQKQANKPTDCSIDSCLTSSFEGSIRDQEVVLYNNLMGLNDKHALGFRTNTEE
RDRKDDGARWREETKESGDDVDRGFINSGNNLSMSIGIEGGDQWNDSGKYYGTEDMFNKG
DDGSDGKLLMKTERSEKTKMPLFSTRLDLNTGSENNAASSYKHHQLDLNGFSWS
>MguPSR_Migut.J01825.1
MKGIQQQQSYGFPGDFPSEFPDNNNFSGNYNNNNINNTQLENQSWHPNSSSTIISRIGST
PASSAFYATELLMGLSNSNFQSGSNIFFGNNSQTRITHEPDFLSKNNQRYRNLAESEQLI
HIENKLLSDIDDSNTISPSLPFDANLDLEVPQNIYGSFSTTTTTTTNNSFQSHASVSSSK
TRIRWTQDLHDRFVECVNRLGGPEKATPKAILKLMETEGLTIFHVKSHLQKYRNAKYVPE
SDAPGKSEKKTSSNNAAQIDIKTGTQLKEALQLQLDVQRRLHEQLEIQRNLQLRIEEQGK
QLKMMFDKQQKTKRNPTTEARNTDKKPGPYNNNLSSTTLEEDTEVLVFDGSDDEILFPYK
IS
>MguPSR_Migut.K00047.1
MMKRTHQNYGFSTDYNFTSDFHYNSQLQSSPMGLVNQSSPPAFYATEEVYTGLSRYNNTV
CSDVQIPNTNHYIMSDNSLSESEQLLYLKNKLLGDLDDENHNLGASQDLYASHFDNINQL
GRQPGCFSAVSSSNNCGSVSSCKTRIRWTQDLHDRFVESVNRLGGPEKATPKAILKLMCT
EGLTILQVKSHLQKYRQVKFVPESVEGRSEKKNSTNNVAQIDMETGMQIKEALLLQLDVQ
RRLHEQLEIQKKLQLRIEQEAKRLKVMIDQQQKTTSDFTHEREEDDEEEEDSEVFVLEGS
DEHVI
>MguPSR_Migut.L01279.1
MALKNAHGKETTLVLSSDAKPRLKWTRELHQIFIDSVNHLGGAEKATPKSLMRIMGIDGL
TLYHLKSHLQKYRLGKSQLPHTCHQNKPEDIEENQLTDKTCDGAHEQIKESLQVAHSLQM
QMEVQKKLHEQIEVQRHLQLRIEAQGKYLQSVLRKAQETISEYSSCSIEVEHAKAQLSQL
ASMVDSTCPSSSFSVLTESGGSLLKNEGNKPLGHNGYSLESSLTSSDSSAMKEETQTTQQ
LAKNYTTINKGKRNSAMLSLMEMNPAKNSESENQARNVKRSKSTTTDEINHREKFDCDKQ
VQRFNLNDFDSGPKVIDLNRNGLDLFGASFSI
>MguPSR_Migut.L01782.1
MYHRQGKNIHPSSKMSIQPHERPLFLQGANGNGDSGLMLSTDAKPRLKWTADLHERFIEA
VNQLGGAEKATPKSVLKLMGIPGLTLYHLKSHLQKYRLSKTHYGQATSGNNNKASTADRV
PEPHTSNPNIATQTTKNIHLGEAIQMQINVQQRLHEQLEVQRHLQLRIEAQGKYLESVLE
KAQETIGSQNLGAFSGMKELSDTSLQQRQKAQAADCSMDSCLNSFEGSVMDQELHNNLGL
KPFNFRASMESKVNDKDTSLQKTELMWRENLKESGKYLSRANNNVENAETNLNSLSMNID
LLGDQWNGDSNYREGTFKEAKADAKFFDHNTNRSDLSRPEKQKASTELKLPFVSTNLDLN
TNDENNIASNCTKLDLNGFSWS
>MguPSR_Migut.L02017.1
MGSNRCNNSSKERLKWTKELHDLFEKAVHQIGGPDRATPRGILRAMAIPELTILHVKSHL
QKYRMSVFTREEDHVGGKFEKTSASEMLPNFSAISSAQLNEALLIQMEAQRRLSDQLEVQ
KSLKTKIEAQGRYLDKIMDEYKNRAPGTKPVKPYSPILSMPSLSEESDQSNIGKEFESDS
EVMSKDEFRAQKRIKIQDDVILPHVHKHSGFFNPAGSHHQSLSMFCVDGLDNNIYTEQEI
SFPWSVSAFQSQLLPRIYNPSNQ
>MguPSR_Migut.M00636.1
VAMERMYGGGGGLMMTRDSKPRLRWTSDLHDRFVDAVTKLGGPDSELKSVLRLMGLKGLT
LYHLKSHLQKYRLGQQQEAKKQNVLDHNTENSGEDIPIAEALRCQIEVHKTLQQQLEVQK
KLQMRIEAEGKYLQSILEKAQHSLSPDINQSENLELTKSQLTDFNLALSNLMQTINGDEI

NGNKVARETRDHMKLETEGPLIEFDLNSRSSYDFIGINNRAVLGANQFENR
>MguPSR_Migut.M01478.1
MFPRLIHPRDAILSQENIQVPNSSGYGQRGDPCLVLTSDPKPRLRWTADLHERFIDAVTQ
LGGASKATPKAILRTMGVKGLTLFHLKSHLQKYRLGKQSGKEFEASKDGGYLLDSPGASN
SSQNLSASDINEGYEVKEALRAQMEVQSKLHLQVEAEKHLQIRQDAERKYMAMLERACKM
LADQILGGNDDGDCYQGGKATQAIMPNGPQNPLLSYASQSANELGVPPPLEMPPNLRQQL
ADCSTESCLTSHGSPVGLPAEGSSPGGKRAILNVGSTNASFVWSESDVYNPVCI
>MguPSR_Migut.M01479.1
MDPISGANNMSSNSNLASKQRLRWTHELHERFVDAVAQLGGPDRATPKGVLRVMGVQGLT
IYHVKSHLQKYRLAKYLPDSSSDGKKAENKESGDILSGLDGSSGMQITEALKLQMEVQKR
LHEQLEVQRQLQLRIEAQGKYLKKIIEEQQRLSGVLPEAPGTEDNKICPESDNKTDPATP
APTSESPFSDKPPKEHTLVKSLSLDESFSSHHHEPLTPDSDCHVTSPVEAPTSDDVAFAK
SEMVNLGHSILESSLSPCYNQQQQHSVFLTRQQFDCSMEVPAGGENTMEKVSGGNV
>MguPSR_Migut.M01502.1
MFQTTKKATSMNNPHEPRGMCVQGDSGLVLTTDPKPRLRWTVELHERFVDAVTQLGGPDK
ATPKTIMRVMGVKGLTLYHLKSHLQKFRLGKQPHKDFHDHSMKDDSSLELQRNNASSSGM
LGRSMNEMQMEVNRRLHEQLEVQRHLQLRIEAHGKYMQTILEKACQTLAGENNMAAAAAA
ASGSYNKAAVGPDMSCGLKDFGPAMNNSFPSLQDLNLYGPTDQQLLDHHHHNHMERSSSI
GNINDVFNIQTNNENFLANKKRPNPYGKSPLIWADDLRLQEHLGSNDDNDCQIHMGPPFS
LDRGGSTSDHNIDDSVSDHNIYETKPILSGDRIHRSNNIIGEKKFDKLSDRSSPPPHHRR
AAERMNHHSAAISAGGIMPQAGRSSPFG
>MguPSR_Migut.N00196.1
MYGGGGGLMMTRDSKPRLRWTSDLHDRFVDAVTKLGGPDSELKSVLRLMGLKGLTLYHLK
SHLQKYRLGQQQEAKKHNVLDHNTENSGEDIPIAEALRCQIEVQKTLQQQLEVQKKLQMR
IEAEGKYLQSILEKAQHSLSPDINQSENLELTKSQLTDFNLALSNLMQTINGDEINGNKV
ARETRDHMKLETEGPLIEFDLNSRSSYDFIGINNRALLFYLICRQWEMNRVMN
>MguPSR_Migut.N01645.1
MNINEQIGVQCNNLGSFRISNSFFEFQSYYSTSSPPQDSCVHPQLPAIFEEQDEEMRRRH
QNVGPAILSDDHDYLYSYRPLKDPADNNNSPHREKIESGDTLQSIVQSCYCGNNLTVEKG
VIQECKHRNNNATISNSCSMLSFNNNINAENCQIEKEQPQINFDNGTTIKFGSIKVVVSG
KEAITNKKKKSRIKWTKELDEKFIDCVKRLGGARKATPKEILKLMKCNGLTIFHIKSHLQ
KYRFARLTPEATEAAARRLKNSQKKLQMRIEEQAKQLQNMIKCQLNTTSE

>VviPSR_GSVIVT01001374001
MNTQKIDVQKQKTGSVNCYSGKGSTFSRQPWKMELCFQPDQFPASEGGSSKQMINLGNTA
NMSTTIAGHFGSPAASAFYATEVYMGFPECDSYPVDSETLSYPSSNLDPASSQSRDTQNI
PSCLEKSFGTPYRNSPVCDILFIKSEVEDEHPYRILRENQNQRIPYQLVEPSPRFQLRRQ
SANPSHSTYSVASGNSSSPAAAASNKSRIRWTHDLHKRFVESVNRLGGAAKATPKGILRL
MGSEGLTIFQIKSHLQKYRIARHLPGSTEEKSEKGTCADFITKFDPETGLRVAEALQLQL
EVQTRLHEQLEIQRNLQMQIEEQGKQLKKMLDSNRIQIRP
>VviPSR_GSVIVT01001376001
MAHPKLEAFFHCREKTPPAPVRVHIHSELFTFFTKIDVVQNQNTGSFNCYSGKGIFSRQP
WKMGFCFQPDQLPASEGGSTQQIINLGSTPTTIAGHFGCPAASAFYATEFYMGFPECDSY
PADSVTSSYPSSKFDPAGSQSKDTQNLPSCENQNSTRTPYRNSQVCDILFIKSDVEDAQP
YRILRENQNQRIEPSSRFQLRRQPANPSHNTTSFASNKTRIRWTQDLHKRFVESVNCLGG
AEKATPKGILKLMGSEGLTIFHVKSHLQKYRIARHQPGSTEENSEKRTCADVITKFDPET
GLRIAEGLRLQLEVQRHLHEQLEIQRNLQLQIEEQGKQLKKMLDSNRDQIRP
>VviPSR_GSVIVT01007064001
MFPGLIQVQPHESIVPQEDLRGSNHRGDPCLVLTSDPKPRLRWTADLHERFVDAVTQLGG
ANKATPKAIMRTMGVKGLTLFHLKSHLQKYRLGKQSGKDMGEAPKDGISASYLSESPGTS
NSSPNLPTSDINEGYEVKEALRVQMEVQSKLHLQVEVKANSGARSQENKASCMPSEERGE

>VviPSR_GSVIVT01007065001
MDPINGGNSLNNNPSLASKQRLRWTHELHERFVDAVAQLGGPDRATPKGVLRVMGVQGLT

IYHVKSHLQKYRLAKYLPDSSSDGKKADKKESGDMLSSLDGSSGMQITEALKLQMEVQKR
LHEQLEVQRQLQLRIEAQGKYLKKIIEEQQRLSGVITETDPATPAPTSEGPLLDKAAKET
APAKSLSIDESFSSHHEPLTPDSGCHMVLTHQILESSLSSSFHQPHSVFLNRDQFDPQAG
ISISNEDQLEKVPSSNH
>VviPSR_GSVIVT01011163001
MPGGLGSVSANMGSSRSDGSGKERLRWTQELHDRFEEAVNQLGGADRATPKGILKAMAVP
GLTIYHVKSHLQKYRISKFVPESSSRAKFERRSISEMLPNFSTTSGAQLKEALQMHMEVE
RRLSDQLEVQKSLKLKIEAQGRFFERIAEEQRNWVSIMKPTNLFSPTSLPSLCEESESNA
KESDSDSDSNKTDMQYEGFQAKKPRLMDDHIYPPRDIISYCVLKGKKNLSILMNAMHSSA
TSL
>VviPSR_GSVIVT01015900001
MHSHDRPLSAQGDSGLVLTTDPKPRLRWTTELHERFVDAVAQLGGPDKATPKTIMRVMGV
KGLTLYHLKSHLQKFRLGKQPHKDFNDQAVKDGEKASALGNQRNATPTPVLMGRNINDRN
MHFNEALRMQMEVRRRLNEQLEVQRHLQMRIDAQGKYMQTILEKACQTLTGKNGDCQSYH
GVGNQGYTEVGSMKDFSSSVNFPCLEDLHIYGERPNPYDSSKAPIIWSNDMQLLQEVGTA
AACIPSQEDAFKSYLKQHHEEESSGSGEA
>VviPSR_GSVIVT01019971001
MSLVLSTDAKPRLKWTPELHHRFVEAVAHLGGPDKATPKTLMRVMGVPGLTLYHLKSHLQ
KYRLGKSQQAETFSDNNQEDYCENQNREIHFDRETGDGTQNPINESLQIAQALQVQLEVQ
RKLHEHIEVQRHLQLRIEAQGKYLQSVLKKAQETLAGYNSSSVGVELAKAELTQLVSIFD
TGCPSSSFSELTETGGSGLKDKERKPMRGTGCSLESSLTSSESSGRKEEKQPKNENGNTN
KCTSMAPTLMEIHPGDSQAWKSGSSNQASGRKRNGSTISDGNCVEQPSAIDLNCKE
>VviPSR_GSVIVT01021072001
MYSAIHSLPLDGGVAHADFQGSLDGTNLPGDACLVLTTDPKPRLRWTAELHERFVDAVTQ
LGGPDKATPKTIMRTMGVKGLTLYHLKSHLQKYRLGKQSCKELTDNSSCIAESQDTGSSS
TSSSRMIPQDLNDGYQVTEALRVQMEVQRRLHEQLEVQRHLQLRIEAQGKYLQSILEKAC
KALKDQAAATAGLEAAREELSELQIKVSNDCEGMNPLETIKMPSA
>VviPSR_GSVIVT01022645001
MERGYENGVVMTRDPRPRLRWTPDLHDRFVDAVTKLGGPHKATPKSVLRLMGLKGLTLYH
LKSHLQKYRLGQQSRKQSITENSDYRTHASGTSAKSSSRNNEQGGILIAEAVRCQVEVQK
QLLEQIEVQKKLQMRIEAQGKYLQAVLDKAQQSLSINVNCPGSLEAMRAQLTNFNMALSS
LTENTNEEDMKVNIIEKSIPSNRANGSVFTAYREAEELVNKDVKFREGEGFRHFDLNTKG
NYEFAGANGADFEAKMIAYRR
>VviPSR_GSVIVT01023401001
MTNPISPQAHQLASNNGTVGYMLSSGSQNSLFIPQSSTNGVSLPPISSSGIEIQSTPSVT
FSRERKENSWCTDSLKDFLDFPENVPIQNNNQVEGSGGGMSYEDCAKTTDWPDWADQFLN
DDDSLEPNWNGLLIDVDVPDPESKVLKPSSSVLTHQSEICQHHPAQSGEISAVPNSLSPA
PSSKPRMRWTPEMHEAFVEAVKQLGGSERATPKGILKLMNVEGLTIYHVKSHLQKYRTAR
YKPKLSEGTSDKNLTSIGEITSLDLKMSMGITEALRLQMEVQKQLHEQLEIQRNLQLRIE
EQAKHLQMMFEKQGKMEDKKLKVSSSIPDEPSSPISNVMQPSPVNHTSKVSEQPHVASGF
DAEESSQNVEQKQKAPETSGCEFINQDNGMSSTPPTKRARAENFLSS
>VviPSR_GSVIVT01025867001
MYHPKKFSTASLVPHKAQGSEPLATVGALGGSSVKNSNPTGGSGKQRLRWTSDLHDRFVD
AITQLGGPDRATPKGVLRVMGVPGLTIYHVKSHLQKYRLAKYLPESPADGSKDEKKGSGD
SGSSMDSAPGVQINEALRLQMEVQKRLHEQLEVQRQLQMRIEAQGKYLQKIIEEQQKLGG
ALKASEAVPLVDDKQNPSQSKPLPDASIGSSSPRKKQKVDDGMTHDCNPPLDPPKPDHGC
NPPLNPPKPDPKHGFIDRWDRDMYGNDGVFGFNLEIEFKEQEDGGSEQRAAPLELEPMCG
SK
>VviPSR_GSVIVT01028403001
MHASPLPSNSGAVGHIFSSSSGYSTDLHFSSVSPHERHSRRQLQSTASSHYIEENNNASW
CTDSLSGFLDFPVNTPVQSSQIESRSASGVIASEDLSKRHDWQEWADQLITDDDALNSNW
NEFLVDTNVADVEPKMAYQVPKPSSNFSANQPQVHPQLSAPSGEVHNVVTPSSSVNTAPT
KPRMRWTPELHEAFVEAVNQLGGSERATPKGVLKLMKVEGLTIYHVKSHLQKYRTARYRP
ESSEGSSEKRLTSIEEMSSLDLKTGIEITEALRLQMEVQKRLHEQLEIQRNLQLRIEEQG

RYLQMMFEKQCKSGIDKLKTSSSALENPSSLSSDTIPNSPAKSEMEASHDEHDKTGTDLV
NDSKTSSGNPQKLSREQNAIETEALLGSEQIAIEPEAPRNVEQDAACESNSQPSKRAKVD
ESSISSVKSASS
>VviPSR_GSVIVT01029458001
MNHHSVLSAKQTESTKGFTQSYCAAVSPIHNLLNVELEVQPQKLCSKSGPYSSVSSDTDA
QYPKCTFSRSSVFCTSLYLSSSSSTETHRPLGNLPFLPHPSMSYQSISAVHSTKTPFLSG
DSSGLYDEGNSEDMMKGFLNLSSDASDESFHVMNCASDNITFSEQLELQFLSDELDIAIA
DNGENPRLDEIYEMPQDSSTPAMALGLTVNQNHQSVAPSTDASSGQPSPGAAAAHKPRMR
WTPELHERFLEAVNKLEGAEKATPKGVLKLMNIEGLTIYHVKSHLQKYRLAKYMPERKED
KKASGSEEKKAASSNNESDGRRKGNIQITEALRLQMEVQKQLHEQLEVQRTLQLRIEEHA
RYLHKILEEQQKAGSALISPPSLSSPTSPHPDSERQPSSPSATTTLPQPAECKADSSSPP
PSKHKAATETTDSEQQACSKRSRLESNPEPVSDEAVVENPSQ
>VviPSR_GSVIVT01033381001
MYHHHHHQGKNIHPSSRTPITPERNLFLQGGNGPGDSGLVLSTDAKPRLKWTPDLHERFI
EAVNQLGGADKATPKTVMKLMGIPGLTLYHLKSHLQKYRLSKNLHGQANSATSKTVVGER
MPEANGALMSSPNIGNQTNKSLHLSETLQMIEAQRRLHEQLEVQRHLQLRIEAQGKYLQA
VLEKAQETLGRQNLGAVGLEAAKVQLSELVSKVSTQCLHSAFSELKELQKQEIHNCGMGL
RPYTNGNGSNSYSEGRFKGRAEADNFMSHGYRLPCFGAKLDLNAHDENDVTLSCKQFDLN
GFSWN
>VviPSR_GSVIVT01033515001
MGFSQYDHQASNPLLYSQSSTSCDSKLDNFSVDSNSTPPADPNFQFRNTFQSVMRASSEN
PNTIQCPSIPFDGNQDIRVCSYLYGSPLAQQAQCARSSSSGGVSVAPANPVSPILHSKAR
IRWTPDLHERFVECVNRLGGAEKATPKAILKLMDSEGLTIFHVKSHLQKYRIAKYMPESA
EGKSEKRASTNDLPHLDNKTGMQFKEALQMQLDVQRRLHEQLEIQRNLQLRIEEQGRQLK
MMFEQQQQTNRSFMEADEDLDIMSLEDPSTSLDQVENLSAQGSGNTRFPSKIS
>VviPSR_GSVIVT01033528001
MRQKGHISTCSYTPHDFPQKVNIWIVSIAGKSEKGASSSDVPHLDNEDGMQIREALQLQL
DLQRRLHEQYSEVYGYLCKSSPLAQQAQCATSSSEGVSIASADPVSPVLHSKPRIRWTPD
LHEHFVECVNRLGGAEKATPKAILKLMDSEGLTIFHVKRHLQKYRIAKHKPGFAGGKPLF
DVDS
>VviPSR_GSVIVT01034259001
MDNQVPEAAAAQKQRIRWTPELHELFLDAVSKLGGPDKATPKGILRLMNVEGLNICHVKS
HLQKYRLAKAVQMKQDKKASSSEERKVATKTDERETPIERAMHVTEALRVQVEVQKTLHE
QLKLQKVIQLNLEQNGEYLRRILEDQHKAGVALPSLMGSHSNPQPITLSSSDGASSPKQY
DFEVDCFPSLLSKHKASHTTESEQPRCHKKPRRSSSVEIISIDSPAESVEG
>VviPSR_GSVIVT01036717001
MFHHKKPSTMNSHDRPMCVQGDSGLVLTTDPKPRLRWTVELHERFVDAVTQLGGPDKATP
KTIMRVMGVKGLTLYHLKSHLQKFRLGKQPHKEFNDHSIKDASALELQRNIASSSGVMSR
NTNEMQMEVQRRLHEQLEVQRHLQLRIEAQGKYMQTILEKACQTLAGENMALGNYKGIGN
QGVPDMGAMKDFGSSLNFPSLQDLNIYGGDQLDLQQSMDRSLDGFIQNTENMCLGKKRPS
PYSGNDPFKGDQIQIAPPSMDRGADMDSISDIYETKPILSGDPMGEKKFDGSGSAKLGRP
SPRRAPLPTDRMNPMINAGAMPQGRSSYG

>TcaPSR_Thecc1EG002071t1
MFHSKKPTFNTYDRTCVQGDSGLVLTTDPKPRLRWTVELHERFVEAVTQLGGPDKATPKT
ILKIMGVKGLTLYHLKSHLQKFRLGRQPQKDFNDQSVSDSRKVSEFYFSKTATGIRACCL
SDQSVQQTGGVRLQMEGGGILNEELEVQKHLQMRIDAQGKYMQTILEKAAAQTLSEENTN
SQCYKYFGIQKYGNMGSMNNLDFVSHVNVPSLEELHSIPASETAHMEKQWKKLNIENLEN
PIVFWSDDLLPENIVIATADYNSDGDSFKDDHL
>TcaPSR_Thecc1EG003939t1
MEARPALSIQRSGARPLSNLGVSGGLSSSLPALTTPLEETYQKLSDTQQVSADRELMTRP
LVHATCVPTNSGVVGHIFSSSSGFSSDLHYSSASPHEKHSRNAPFISQSPTDATALPLPQ
SSNSALPQSTISSHFNKESSGSWCTDPGFLDFPVNTPIQSSQVESNSCSGIMISEDFSKR
NDWQEWADQLITDDEALASDWNELLVDNNVTDLEPKMAYQVAKPCTTMPAQKPQAQQQLP

SPSVESRSVVNPSSSANNAPAKPRMRWTPELHEAFVEAVNQLGGSERATPKGVLKLMKVE
GLTIYHVKSHLQKYRTARYRPESSEGSSEKKLTPIEDLSSLDLKTGMGITEALRLQMEVQ
KRLHEQLEIQRNLQLQIEEQGRYLQMMFEKQKSGLDKLKVSSSNHLENPPAPPSDATKES
PAKSDLEASQMDHVNSGTETVNANSMLESSSQEIVAKQKAPETGDLEKAETCVSESSSQP
SKRPRIEE
>TcaPSR_Thecc1EG011693t1
MGLQNLQNQNMSLVLSTDAKPRLKWTPELHQRFVEAVNQLGGADKATPKSLMRVMGIAGL
TLYHLKSHLQAITLHYKYRLGKSQQTEICLSNKQDDYREIQSSNGDFHSDISDETHKQMN
DLQIAQALQMQMEVQRKLHEQIEVQRHLQLRIEAQGKYLQSVLKKAQETLAGYSSSSVGV
ELAKAELSQLVSMVNTGCTSSSFSELTEGGGSSLKIERKPMRGTICSMESSLTSSESSGR
KDEEPPKNENISTPKSNASVELSLMDIHPEKKPLIGGSSNQAKGKKRSGSNISDGTCVEQ
PLAKRLELPEEETGHWLRKSGLLGSFDLNSQCQSDIELGPKAIDLNCRE
>TcaPSR_Thecc1EG016647t1
MERLSYGGGAGGGAVGGENFGYENGVVMTRDPKPRLRWTADLHDRFVDAVTKLGGPDKAT
PKSVLRLMGLKGLTLYHLKSHLQKYRLGQQARKQNAVDQNKDNGGSSYVQFSNHSPGTIT
NSPSADNDQRQIPVADALNTHLEVRRTLQEQLEVQKKLQMRVEAQGRYLQAILEKAHKSL
SFDINCEGNVEETRAELTNFNLALSSLMENVNGGADRKPNVVQMNEVPKKATSCSAFQNY
AVGERERNKDVKLKVEGESINFDLNTKDSFEFVAVNGNELQSHMFSYKR
>TcaPSR_Thecc1EG020332t1
MYPAIRSLPLDGSVGDYQGSLDGTNLPGDACLVLTTDPKPRLRWTAELHERFVDAVTQLG
GPDKATPKTIMRTMGVKGLTLYHLKSHLQKYRLGKQSCKESTDNSKDASCVAESQDTGSS
TTSTSRMVAQDLNDGYQVTEALRVQMEVQRRLHEQLEVQRRLQLRIEAQGKYLQSILEKA
CKALNDQAAASAGLEAAREELSELAIKVSNDCQEMIPLDNIKLPSLSELAAALENKTASS
MPVSPMGVGSQAAIMKKRPRPLFGNADPLPLDGNIRQEIEWVMPNIS
>TcaPSR_Thecc1EG021688t1
MRRPSRSDGLAKERLRWTQELHDRFEDAVNQLGGPDRATPKGILKAMGVDGLTIYHVKSH
LQKYRILKFVPETNTKCKFERRDISEILPNFGTTSGAQLNEALQMHKEAERKQGDHQLEA
QRNLKIKIEAQVIFLERLAGQHGNRATPTKATKPFSPTSLPSLCEESESTAKDGFETDPE
ADRNEIESGERVQAMYAFESWDQYDHQDMVLNREERVSYLANDISFPWNVPVCSSSPLVP
SFL
>TcaPSR_Thecc1EG024878t1
MDGASSQPKQKRLVLHSVTTPTTPLEQLASAFCATDQSKSHIDFQLAERDSDEPQAPGYE
AIPTINQAKDTLLSIVNPHLWRNQCQRFTEKSNGKSYWSQQGSLLSLDHGKLLAEATRNS
KVISLCSSRPEQQYPNISRISGTRASSVSSGGRPSNIKTRIKWTVELHKKFVNCVNLLGG
AEKATPRAILKLMESNGLTVLQVKSHLQKYRYAKYITGSTQAKPDQGVDDDDLPMLYLKS
GMQVKEILQMQLEVQRHLCEQLEIQQNLQQAIQEQGKQIQMMLEQLKKTNKA
>TcaPSR_Thecc1EG024886t1
MNSRKIDCQEHLEQNLGFSSVCNFEYVNHDGFQQPWNMGIRIQAPAMEEGSQQENPGAAK
TSNTIMSGFLSPASAFYATERCMGFSEYGCQGDRSSYTSQYNKSCNSHLPSFHASGDNFS
IESVAQDETNYELRNTFESLVKSQIYCNQYQKSSEKSYKIPCCNSQGSQVSPHDQSNFLG
NNAVTVGSHYSVPFRGNQDQRAYCNSYSSPLAQLSIFQQGKQSSNCSSGTFSVSSGNSVS
TGAALASKTRIRWTQDLHDKFVECVKRLGGAEKATPKAILKLMDTEGLTIFHVKSHLQKY
RIAKYMPDSAEGKSDKRSSTSDVTQLDVKTGLHLTEALQLQLDVQRRLHEQLEIQRNLQL
RIEEQGRQLKMMIDQQQKTNESLLKKQDLDITPFDHDPSFSLEDVEVSIAENSGDAHFPS
KIS
>TcaPSR_Thecc1EG026171t1
MSSSFPALPTPFKEKYPKLPDSFQVSSERKVMKNSISPQESSLAPSNRTLGNSFSSPSIA
NNDMCASALAHDRHSQSPAFISQRSRDLASLPSIDSSYSDQSTALFNHPQEKKDVSWCID
RLQDFLDLPENVPDPNGLLESSTGVMASEDHSKRTDWQEWADQLISVDDPLDTDWREFLD
DTNASDPKVKVLNSSGDISKQQPQFHQNQPAPHGEFSSDAYPLSPAPPTRPRMRWTPELH
EAFVDAVNILGGSERATPKGILKLMKVEGLTIYHVKSHLQKYRTARYKPESSEGTLENKM
ASIGEMKSLDLKAGMGITEALRLQMEVQKQLHEQLEIQRNLQLRIEEQGRYLQMMFEKQK
RMEDERTGAPSFNLDDASASLPGLTCPSCANDKSEALEQVHTKTGIDTRNASTTEDKSSQ
DVSRKQKALETNTADHIETNDNESGSPLSKRARTEK

```
>TcaPSR_Thecc1EG026448t1
MYSLFHSLPSNNLQGRASFTPPFYFSFFLYVSNSKASSFTVNIDNPKKRLQQQGEEQSSF
WRQKMYHHHHQHQGKNIHPSSRMPIPPERHLFLQGGNGPGDSGLVLSTDAKPRLKWTPDL
HERFIEAVNQLGGADKATPKTVMKLMGIPGLTLYHLKSHLQKYRLSKNLHGQANNGSNKI
GAVAMAGDRMSEANGTHVNNLSIGPQANNGLQIGEALQMQIEVQRRLHEQLEVQRHLQLR
IEAQGKYLQAVLEKAQETLGRQNLGSVGLEAAKVQLSELVSKVSNQCLNSAFSDLKDLQG
LCPQQTQATPPTDCSMDSCLTSCEGSQKEQEIHNNGMCLRPYNTSGALLEQREIAEDPLL
PQTELKSFEDIKENKMFLSSLGKDAERRMFFADRSSSDLSMSVGLQGEKGNGGNSSSFSE
AKFKGRNEDDSFLDRGNKRADEVNRLPYFATKLDLNVHEENDAASSCKQFDLNGLSWN
>TcaPSR_Thecc1EG027630t1
MFNLEAALSWMKVGMSSKTMNHPSYISVTQSEPSKGIGESHHTAASPIHNFLSIGSEGQS
SLAGECSSPHPFPFIRTESFKNNLKSGPSSPISPSSHAKSAFSRSSVFCTSLYLSSSSTS
ETQRQLGNLPFLPHPPTCGQSISAVDSSKSPVVFSEDLHNPYNEDHSEIIMKDFLNFPGD
DCDGNFHGLHCESNNFTLTEQLELQFLSDELDIAIADHGENPRLDEIYETPQKLNVAFTC
NQNSASVVPSTDACSSIRLSGPAAVHKPRMRWTPELHECFVEAVSKLDGPEKATPKGVLK
LMNVEGLTIYHVKSHLQKYRLAKYMPEKKEEKKTSSSEEKKAALSGNESDGKKKGGTHIT
EALRMQMEVQKQLHEQLELQRSLQLRIEEHARYLQKILEEQQKAGSALLPSLSMSTPTDP
SQNSELQPSSSSAIASPTQPSESKTELSSSLPSKHKAPEVNDCEPESSPKKLRTENKPES
AADEAVVENPAQ
>TcaPSR_Thecc1EG030479t1
MDMYHPRLQAHLHHPHHQEEMLQNLNHKGALAEPCLVLTSDPKPRLRWTADLHDRFVDAV
TQLGGPNKATPKAIMRTMNVKGLTLFHLKSHLQKYRLGKQSGKDMGEGPKDGMSASYLLE
SPGTNNSTPSLPSSDMNEGYEVKEALRAQMEVQSKLHLQVEAEKHLQIRQDAERRYMAML
ERACKMLADHFIGGGADTETENLGFSSKVPRNYCVDPLGFYSSQSAEVVNACGQDEEMPS
GLHSQRADCSTESCLTSHESPGGLTMEGSAVEGKKKMLNLDSTTGSLIWGDAKVNPHGLT
GYGM
>TcaPSR_Thecc1EG030480t1
MQKSWLWPGLWGWWVCLCVCSASLDFFFPFDFGLQTLSEGRNLTGWVAPRSASSSDDQQM
YQPKTVPGSSLVRNNSIVHGQHLDCGASQMDPISGGNSLTNNPNLASKQRLRWTHELHER
FVDAVAQLGGPDRATPKGVLRVMGVQGLTIYHVKSHLQKYRLAKYLPDSSSDGKKADKKE
TGDMLSNLDGSSGMQITEALKLQMEVQKRLHEQLEVQRQLQLRIEAQGKYLKKIIEEQQR
LSGVLAEAPGSGASVPALGDNGLESDKKTDPATPAPTSESPLQDKAAKERAPAKSHSIDE
SFSSHHEPLTPDSGCHVGSPAGSPKGERLMKKQRVSMAAAFAKPEVVLPHQILESSISSS
FQQSHSVFMTREQFDPSSGISMGNEDQLEKASGTEL
>TcaPSR_Thecc1EG033672t1
MFQPKKPSTMNSHDRAMCVQGDSGLVLTTDPKPRLRWTVELHERFVDAVTQLGGPDKATP
KTIMRVMGVKGLTLYHLKSHLQKFRLGKQPHKEFNDHSIKDDLQRNAASSSGMIARSMNE
MQMEVQRRIHEQLEVQRHLQLRIEAQGKYMQSILEKACQTLAGENMAAGGYKGMGNQGVP
DMGAMKDFGPLNFPPFQDLNIYGGDQLDLQQNMDRPSLDAFMPNNDNICLGKKRASPYSG
SGKSPLIWSDELRLQDLGTAASCLGPQDDPFKSEQIQIAPPSIDRSTDLDPISDIYEAKP
VLSGDGMGDKKYEASPKLERPSPRRAPLQADRMNPMINSGSVAQGRNSPYG
>TcaPSR_Thecc1EG039819t1
MTIFSLSGLVEVLFRASFLLLLLVLCHHLIESEKHNYHEQSVKGAMQTSRSGPSSVYKLL
SGEPIGLSSANGNSSFTILQPCYEQSEAPASAPQASVSQPRDLIPNPVPHNSYIICHKST
RLCTGFHSSTSESSNTEKHVGNSRCSSPLPMFDHLIPATNTPDSSLLFGEGKTTTCDKED
FEKFMVFPHVPAENSTVIPEHSAYPKGTLTLKEQLQLQYLTRELEIEMNGNGSENENPGL
AETHKVPRVATVPVIQLEHNRNCLSSVVNLDGCIHSTHQHPEAVAANKQRIRWMPKLHEL
FLNAVDKLGGPESATPKNILKLMNVQGLHICHVKSHLQKYRLAKNVSELKHDKRSSRFEE
KETLTETHGDGNIAKERDTQVLETLRMQVEVQKLLHEQLKVRSCAALCW
>CpapPSR_10.14
MKHRSIMSVAPSESRNGVRHSFCTTTSPIHNFLGVETERETSPFIQSESLGSPNQVRGYM
IQPQKHFLKSGPSSPVSPSPNGQHPRSTFSRSSLFCTSLYLSSSSKSETQRQLGNLPFLP
NPPTYNHSTSAVESAKSPAIFSEDLGNPYDDDQSEALMRDFLNIPGDTSDGSFHGMNCGN
DSLALTEQMELQFLSEELDIAITDHGETPRIDEIYEIPQASPKSALGFTCSQSFASVVPA
```

ADAVSTQPSSGSATVHKPRMRWTPELHERFVEAVNKLDGPEKATPKGVLKLMNVEGLTIY
HVKSHLQKYRLAKYMPEKKEEKKPSSSEEKKAASSSTESDGRKKGGIQITEALRMQVEVQ
KQLHEQLEIQRALQLRIEEHARYLQKILEEQQKAGGTLIPSQGFSFMSNSHQNSELQSST
SAAAALPSQPSGSSSPSPSISNHKDDPDKNDSKAEKSPKRPRLEHETKKASDEAEVGRQE
ASPLHVVSTL
>CpapPSR_128.40
MNLVLSTDAKPRLKWTPELHHRFVEAINQLGEATPKSLMRVMGIPGLTLYHLKSHLQKYR
LGKGHQAETCIDNNQEDYGEVQSGSHGHFNTDMNNGNHNQINENFQIAQALQVQMEVQRK
LHEQIEVQRHLQLRIEAQGKYLQSVLKKAQETLAGYSSSSVGIELAKAELSRLVSMVNTG
CPSSSFSELTEVGDSIMKDVNQKGTRGTICSMESSLTSSESSGGNDNGGPSKVNTCLELP
LMAIHPEKKPWSGDSSNQAVGRKRSCSTISDGICVEQPLSKMLEIHQEKRNDQLGNSGFL
GTLDLNSQYHLNDSEQGPKAIDLN
>CpapPSR_2085.1
MTNPMSSQAPPLVPDSQTVGHLFSSSRGLPHSMTLASVSSRERNYGKFPLISQSSSGGDS
FPQLLRSGSSLPTNRASRTEMQSRALINRPEEGKDISWSIDSLQDFLDLPENIPVQTVQM
ESSKGAVASEDHSKRTDWQEWADQLISADDTLEQNWSELLCEFNAQEPKPKTVSSGEFSA
IANPLSNAPSNKQRMRWTPELHESFVEAVNQLGGSERATPKGVLKLMNVEGSSEKKSIPL
DDVKSLDLKTSMSITEALRLQMEVQKQLHEQLEFPLRWYSKQNSTGLLQGFEGFHGKIKV
RLA
>CpapPSR_27.185
MDPINGGNNLNNNANLASKQRLRWTHELHERFVDAVAQLGGPDRATPKGVLRVMGVQGLT
IYHVKSHLQKYRLAKYLPDSSSDGKKTDKKETGDMLSNLDGSSGMQITEALKLQMEVQKR
LHEQLEACFISAIFHRQLLKIKQYEFAGIYCSFGGHYSGRQVQRQLQLRIEAQGKYLKKI
IEEQQRLSGVLAEAPGSGVSAPVSGDNCLDSDTKNDPGTPGPTSESPVQEKAGKDRAPDK
SLSVEESFSSHHEPLTPDSGCHGGSPADSPEGERSSKRQRVGMGASYSKQELVLTHQILE
SSLSSSYQQPHTVFLTREGFDPLSALPIRNEDQLEKVSGSDL
>CpapPSR_27.186
MYGRVIGSHESGIVGHEEMVGGPVGHTHAHRPPSGDPCLVLTSDPKPRLRWTADLHDRFV
DAVTQLGGPNICLTCMHGNLPSIPPWEKEVSFPVCVNRMNFSLKKCGESLLYESEFYYML
MRMCNNCRSNTKGNHADNECEGTNTLPLEESPPGFCSLLKFLLSQKYRLGKQSGKDMGEA
PKDGMSASYLLESPGISNSSPNLPSSDVHEGYEVKEALRVQMEVQSKLHLQVEAEKHLQI
RQDAERRYLTMLERACKMLADQFLGGALSDSENQKCVGVAGKMARNCSLDPLGFYSSHST
EVGNANGQEEEVPAGIHQQRTDCSTESCLTSHESPRGLTMEGPLVGGNKRIMNLDTTTGS
LIWSGTKMRSQEVNVAAQSPQGTAGCGIHQN
>CpapPSR_32.94
MFHSKKPSTMNSHDRPMCVQGDSGLVLTTDPKPRLRWTVELHERFVDAVTQLGGPDKATP
KTIMRVMGVKGLTLYHLKSHLQKFRLGKQPHKEFNDHSIKDGERASALDLQRSAASSSGI
IGRSMNEMQMEVQRRLHEQLEVQRHLQLRIEAQGKYMQSILEKACQTLAGENNIAAAAAS
YKGLGNHCGADMGTMKNFVPPLNFSSFQDLNIYGGDQLDHHLQQNMDRSSSLDHGFMSND
DTLCLGKKRSNPYSTCSGKSPLIWADDPRLQEPSCLGPQDHDLFKHSAHDQMQIAPPNSI
DRGADHLESISDIYETKPVLSGDHAMADKKFESTKLERPSPQRPSDRMSPMMNTGGMPQG
RNSPFG
>CpapPSR_33.181
MYHAKKISTANLVPHKPQGAEQFPSTGTLGGNVVNNSTSAGGGGKQRLRWTSDLHDRFVD
AITQLGGPDRATPKGVLRVMGVPGLTIYHVKSHLQKYRLAKYLPESPADGLKDEKKDAGD
SITTPDSSPGVQINDTLRMQMEVQKRLHEQLEVQRQLQMRIEAQGKYLQKIIEEQQRIGS
KLKATEAVDKEKASESEASADASEGLLSPRKKQKRNDDQAASKS
>CpapPSR_785.2
MYHHHQHPGKNMHLSSRMPIPPERHLFLQGGNGPGDSGLVLSTDAKPRLKWTPDLHERFI
EAVNQLGGADKATPKTVMKLMGIPGLTLYHLKSHLQKYRLSKNLHGQTNSGTNKLGAVVM
AGEKMPEANGTHMNNLSIGPQTNKSLHINETLQMQIEVQRRLHEQLEVQRHLQLRIEAQG
KYLQAVLEKAQETLGRQNLGAVGLEAAKVQLSELVSKVSTQCLNSAFSELKELQGLCPQQ
TQLTQPADCSIDSCLTSCEGSQKDQEIHNNGLGLRHYHGNSLLEQKERVQEPMLHQTELK
WSRDVKENKMFLSSIGKDTERRNSFVERHSSDLSMSVGLQVEKGRGSNSFSEGRLEEEMT

TIFLETKPMGGQTQLDLNARDENDAASSCKHFDLNGFSWS
>CpapPSR_85.82
MNTQKITCQERIRLTQGLISDCKVDFVNHSSQFSGTQQRSDMGIQIQAPAIEGGISQLHY
QFRSPNNILSPFESPTYDFYATEGCMGFPLYKSETTAVSMDSIEQFNTSSELTNTLQSLV
KSQYCGRKFSRSPSLDQNRFLVDEATDFGRNKSRVALKGNQDHRIFCNSNYNSPLVRQSF
YAKQEKQQSLRLSPVKSVKNGSELVSKTRIRWTQDLHEKFVECVNRLGGTEKATPKEILK
LMDSNGLTIFHVKSHLQKYRIAKYMSDSAEGKSEKRSSTNYIPYQLDTKTGMQIKEALEL
QLDIQRRLHEQLEDLRTTLSGLIFSSKPVTKHPLGNSKREVSCPQFIALSPYGYEFLSAR
GRRIRESALCNTKRRTKKKNGGYEEEEDGGDGRCSNFDAGTITYFARSSR
>CpapPSR_34762
MGSSRSRGTSKERLHWTQDLHDRFEEAVNQIGGPDRATPKGILNAMGIEGLTIYHVKSHL
QKYRISKFVLESSSKGKFERSNISEILPNFGSTCAAQLKEALKMHKEAQRRLSDQNEIHR
NLKLKIEAQGRFLGRIAAEYQNRTTSRRTRKSLSLQSTPSLCEESESNAKETERDLEAER
IEMESADHREIILARKRPRVVEDNNNVFTPPPTSEVTNSDPYNQNLFLAEEEISKQSASD
VIFPWNIITCPSPLIASYF

>AthPHR1_AT4G28610
MEARPVHRSGSRDLTRTSSIPSTQKPSPVEDSFMRSDNNSQLMSRPLGQTYHLLSSSNGG
AVGHICSSSSSGFATNLHYSTMVSHEKQQHYTGSSSNNAVQTPSNNDSAWCHDSLPGGFL
DFHETNPAIQNNCQIEDGGIAAAFDDIQKRSDWHEWADHLITDDDPLMSTNWNDLLLETN
SNSDSKDQKTLQIPQPQIVQQQPSPSVELRPVSTTSSNSNNGTGKARMRWTPELHEAFVE
AVNSLGGSERATPKGVLKIMKVEGLTIYHVKSHLQKYRTARYRPEPSETGSPERKLTPLE
HITSLDLKGGIGITEALRLQMEVQKQLHEQLEIQRNLQLRIEEQGKYLQMMFEKQNSGLT
KGTASTSDSAAKSEQEDKKTADSKEVPEEETRKCEELESPQPKRPKIDN
>AthPHL1_AT5G29000
MTLANDFGYSTAMSSSYSALHTSVEDRYHKLPNSFWVSSGQELMNNPVPCQSVSGGNSGG
YLFPSSSGYCNVSAVLPHGRNLQNQPPVSTVPRDRLAMQDCPLIAQSSLINHHPQEFIDP
LHEFFDFSDHVPVQNLQAESSGVRVDSSVELHKKSEWQDWADQLISVDDGSEPNWSELLG
DSSSHNPNSEIPTPFLDVPRLDITANQQQQMVSSEDQLSGRNSSSSVATSKQRMRWTPEL
HEAFVEAVNQLGGSERATPKAVLKLLNNPGLTIYHVKSHLQKYRTARYKPETSEVTGEPQ
EKKMTSIEDIKSLDMKTSVEITQALRLQMEVQKRLHEQLEIQRSLQLQIEKQGRYLQMMF
EKQQKIQDNKSSSSEASPKQCNGSFAEVEVGLETLTGDQNESASASRKRVRED
>AthPHL2_AT3G24120
MYSAIRSLPLDGGHVGGDYHGPLDGTNLPGDACLVLTTDPKPRLRWTTELHERFVDAVTQ
LGGPDKATPKTIMRTMGVKGLTLYHLKSHLQKFRLGRQAGKESTENSKDASCVGESQDTG
SSSTSSMRMAQQEQNEGYQVTEALRAQMEVQRRLHDQLEYGQVQRRLQLRIEAQGKYLQS
ILEKACKAFDEQAATFAGLEAAREELSELAIKVSNSSQGTSVPYFDATKMMMMPSLSELA
VAIDNKNNITTNCSVESSLTSITHGSSISAASMKKRQRGDNLGVGYESGWIMPSSTIG
>AthPHL3_AT4G13640
MYSAIRSSLPLDGSLGDYSDGTNLPIDACLVLTTDPKPRLRWTSELHERFVDAVTQLGGP
DKATPKTIMRTMGVKGLTLYHLKSHLQKFRLGRQSCKESIDNSKDVSCVAESQDTGSSST
SSLRLAAQEQNESYQVTEALRAQMEVQRRLHEQLEYTQVQRRLQLRIEAQGKYLQSILEK
ACKAIEEQAVAFAGLEAAREELSELAIKASITNGCQGTTSTFDTTKMMIPSLSELAVAIE
HKNNCSAESSLTSSTVGSPVSAALMKKRQRGVFGNGDSVVVGHDAGWVMPSSSIG
>AthPHL4_AT2G20400
MIPNDDDDANSMKNYPLNDDDANSMKNYPLNDDDANSMENYPLRSIPTELSHTCSLIPPS
LPNPSEAAADMSFNSELNQIMARPCDMLPANGGAVGHNPFLEPGFNCPETTDWIPSPLPH
IYFPSGSPNLIMEDGVIDEIHKQSDLPLWYDDLITTDEDPLMSSILGDLLLDTNFNSASK
VQQPSMQSQIQQPQAVLQQPSSCVELRPLDRTVSSNSNNNSNSNNAAAAAKGRMRWTPEL
HEVFVDAVNQLGGSNEATPKGVLKHMKVEGLTIFHVKSHLQKYRTAKYIPVPSEGSPEAR
LTPLEQITSDDTKRGIDITETLRIQMEHQKKLHEQLESLRTMQLRIEEQGKALLMMIEKQ
NMGFGGPEQGEKTSAKTPENGSEESESPRPKRPRNEE
>AthPHL5_AT5G06800
MMDNINFEFSNASQGSRLQLQQQPPQPFNLQDLNMIQYNQPSSPWTTETFSGLTPYDCTA

NQSFPVQCSSSKPYPSSFHPYHHQSSDSPSLDQSVSMIPMQPLPDQYMKPLYQRSCSNDF
AATNASSASYSLSFEASHDPQELCRRTYSNSNVTHLNFTSSQHQPKQSHPRFSSPPSFSI
HGGSMAPNCVNKTRIRWTQDLHEKFVECVNRLGGADKATPKAILKRMDSDGLTIFHVKSH
LQKYRIAKYMPESQEGKFEKRACAKELSQLDTRTGVQIKEALQLQLDVQRHLHEQLEIQR
NLQLRIEEQGKQLKMMMEQQQKNKESLLKKLPDAEASLSLLDPHIHSPPSPFLVHDAEAL
MLTSYEDTQLQSTKS
>AthPHL6_AT5G06800
MMDNINFEFSNASQGSRLQLQQQPPQPFNLQDLNMIQYNQPSSPWTTETFSGLTPYDCTA
NQSFPVQCSSSKPYPSSFHPYHHQSSDSPSLDQSVSMIPMQPLPDQYMKPLYQRSCSNDF
AATNASSASYSLSFEASHDPQELCRRTYSNSNVTHLNFTSSQHQPKQSHPRFSSPPSFSI
HGGSMAPNCVNKTRIRWTQDLHEKFVECVNRLGGADKATPKAILKRMDSDGLTIFHVKSH
LQKYRIAKYMPESQEGKFEKRACAKELSQLDTRTGVQIKEALQLQLDVQRHLHEQLEIQR
NLQLRIEEQGKQLKMMMEQQQKNKESLLKKLPDAEASLSLLDPHIHSPPSPFLVHDAEAL
MLTSYEDTQLQSTKS
>AthPHL7_AT2G01060
MEADNGGPNSSHASKQRLRWTHELHERFVDAVAQLGGPDRATPKGVLRVMGVQGLTIYHV
KSHLQKYRLAKYLPDSSSEGKKTDKKESGDMLSGLDGSSGMQITEALKLQMEVQKRLHEQ
LEVQRQLQLRIEAQGKYLKKIIEEQQRLSGVLGEPSAPVTGDSDPATPAPTSESPLQDKS
GKDCGPDKSLSVDESLSSYREPLTPDSGCNIGSPDESTGEERLSKKPRLVRGAAGYTPDI
VVGHPILESGLNTSYHQSDHVLAFDQPSTSLLGAEEQLDKVSGDNL
>AthPHL8_AT1G69580
MCLLMEINNNANNTNTTIDNHKAKMSLVLSTDAKPRLKWTCDLHHKFIEAVNQLGGPNKA
TPKGLMKVMEIPGLTLYHLKSHLQKYRLGKSMKFDDNKLEVSSASENQEVESKNDSRDLR
GCSVTEENSNPAKDRGLQITEALQMQMEVQKKLHEQIEVQRHLQVKIEAQGKYLQSVLMK
AQQTLAGYSSSNLGMDFARTELSRLASMVNRGCPSTSFSELTQVEEEEEGFLWYKKPENR
GISQLRCSVESSLTSSETSETKLDTDNNLNKSIELPLMEINSEVMKGKKRSINDVVCVEQ
PLMKRAFGVDDDEHLKLSLNTYKKDMEACTNIGLGFN
>AthPHL9_AT3G04030
MYYQNQHQGKNILSSSRMHITSERHPFLRGNSPGDSGLILSTDAKPRLKWTPDLHERFIE
AVNQLGGADKATPKTIMKVMGIPGLTLYHLKSHLQKYRLSKNLNGQANNSFNKIGIMTMM
EEKTPDADEIQSENLSIGPQPNKNSPIGEALQMQIEVQRRLHEQLEVQRHLQLRIEAQGK
YLQSVLEKAQETLGRQNLGAAGIEAAKVQLSELVSKVSAEYPNSSFLEPKELQNLCSQQM
QTNYPPDCSLESCLTSSEGTQKNSKMLENNRLGLRTYIGDSTSEQKEIMEEPLFQRMELT
WTEGLRGNPYLSTMVSEAEQRISYSERSPGRLSIGVGLHGHKSQHQQGNNEDHKLETRNR
KGMDSTTELDLNTHVENYCTTRTKQFDLNGFSWN
>AthPHL10_AT5G18240
MYYHNQHQGKSILSSSRMPISSERHPFLRGNGTGDSGLILSTDAKPRLKWTPDLHERFVE
AVNQLGGGDKATPKTIMKVMGIPGLTLYHLKSHLQKYRLSKNLNGQANSSLNKTSVMTMV
EENPPEVDESHSESLSIGPQPSMNLPISDALQMQIEVQRRLHEQLEVQRHLQLRIEAQGK
YLQSILEKAQETLGRQNLGAAGIEATKAQLSELVSKVSADYPDSSFLEPKELQNLHHQQM
QKTYPPNSSLDSCLTSSEGTQKAPKMLDNRLGLRTYIGDSTSEQKEIMEEPFFHRMELTW
AEEESLRENHNRPYLSTMVNNAEPRISSSRRSPGRLSIGVGLHEHRGRSSNNSEYTEERF
NENNEDCKLETHTRTALDLNTHDENYGTTRPKQFDLNGFSWN
>AthPHL11_AT5G45580
MMTRDPKPRLRWTADLHDRFVDAVAKLGGADKATPKSVLKLMGLKGLTLYHLKSHLQKYR
LGQQQGKKQNRTEQNKENAGSSYVHFDNCSQGGISNDSRFDNHQRQSGNVPFAEAMRHQV
DAQQRFQEQLEVQKKLQMRMEAQGKYLLTLLEKAQKSLPCGNAGETDKGQFSDFNLALSG
LVGSDRKNEKAGLVTDISHLNGGDSSQEFRLCGEQEKIETGDACVKPESGFVHFDLNSKS
GYDLLNCGKYGIEVKPNVIGDRLQ
>AthPHL12_AT3G12730
MMQSREEIRDDSSSGLVLTTDPKPRLRWTTELHERFVDAVTHLGGPEKATPKTIMRVMGV
KGLTLYHLKSHLQKFRLGKQPHKEHSQNHSICIRDTNRASMLDLRRNAVFTTSPLIIGRN
MNEMQMEVQRRIEEEVVIERQVNQRIAAQGKYMESMLEKACETQEASLTKDYSTLFFDRT
NICNNTSSIPIPWFEDHFPSSSSMDSTLILPDINSNFSLQDSRSSITKGRTVCLG

>AthPHL13_AT3G04450
MTLASDFGFPSAISSSFTILEERYHNNFPNTLCVSSGQESMNNNPVPCQVFPLVSGGSSG
GNLFSSSSGFCNGVYVSSSSQARPSVSTVPRDRITVAHVSGEGQRQECPVETHSLQLINQ
PQEQKIMTWSSDQIRGFFDFPVPDPQAASSRTMVSSKEVLSKCEWPDWADQLISDDSLEP
NWSELLGDPNVLNLYSKIETQSSDIARQEIVFRNQHQVDPSMEPFNAKSPPASSMTSKQR
MRWTPELHEAFVEAINQLGGSERATPKAVLKLINSPGLTVYHVKSHLQKYRTARYKPELS
KDTEEPLVKNLKTIEDIKSLDLKTSIEITEALRLQMKVQKQLHEQLEIQRSLQLQIEEQG
RYLQMMIEKQQKMQENKKDSTSSSSMPEADPSAPSPNLSQPFLHKATNSEPSITQKLQNG
SSTMDQSESTSGTSNRKRVRED
>AthPHL14_AT1G79430
MFHAKKPSSMNGSYENRAMCVQGDSGLVLTTDPKPRLRWTVELHERFVDAVAQLGGPDKA
TPKTIMRVMGVKGLTLYHLKSHLQKFRLGKQPHKEYGDHSTKEGSRASAMDIQRNVASSS
GMMSRNMNEMQMEVQRRLHEQLEVQRHLQLRIEAQGKYMQSILERACQTLAGENMAAATA
AAAVGGGYKGNLGSSSLSAAVGPPPHPLSFPPFQDLNIYGNTTDQVLDHHNFHHQNIENH
FTGNNAADTNIYLGKKRPNPNFGNDVRKGLLMWSDQDHDLSANQSIDDEHRIQIQMATHV
STDLDSLSEIYERKSGLSGDEGNNGGKLLERPSPRRSPLSPMMNPNGGLIQGRNSPFG
>BraPSR_LOC103827611
MEADDGGNNSSHASKQRLRWTHELHERFVDAVAQLGGPDRATPKGVLRVMGVQGLTIYHV
KSHLQKYRLAKYLPDSSSEGKKTDKKESGDVLSGLDGSPGTQITEALKLQMEVQKRLHEQ
LEVQRQLQLRIEAQGKYLKKIIEEQQRLSGALGESSGPVTGESDPATPAPTSEFPLQGKS
GKECEPDKSLSVEESHSSYREALTPDSGCNIGSQDESAGEERSSKKPRLMRGGAAGYTSE
MVVAHPILESGMNTSYHQADHALAFDHPSTSLLGDEDGLNKVSEDVL
>BraPSR_LOC103827790
MERVNLGGLGYENGGVMMTRDPKPRLRWTADLHDRFVDAVAKLGGADKATPKSVLKLMGL
KGLTLYHLKSHLQKYRLGQQQAKKQNRTEQNKENAGSSYVHFDNCSQGGISNESRFDNHL
RQSGNVPFADTLRHQVDAQQRFQEQLEVQKKLQMRMEAQGKYLLTLLEKAQKSMPCGGNG
AETDKGQFSDFNLALSGLVGNDHKSNKAGLITNISHVNGDLTDNFRLCGERDTGETEDAC
VKPESGFVHFDLNSKDGYDLLNSGKYGIEMKPNVIADRHQ
>BraPSR_LOC103828177
METRPFRSMSSLALTSSVPNPSVNADISFSSEHNQSMASPYHLLSANGGAVGHNIYSNDL
HNHPSMASHIGSSSDTPFISEILDWDHPAPIPDLFDFPLDIPIQTNQMDDIHRPSELAEL
DEELITDDENPLMSALLNDLFLDTSSTSAASKVQEPTMQSQIQQPQVVLHQTSPIVRTVS
SNSFTSNNTAATKGRVRWTPELHEAFVEAVNQLGGMENAKPMAVLKHMKVQGLTIYHVKS
HLQKYRTARHVPEPSEAGWREKKLTPVEHVTSLDTKRFGNITVHNGMYISEALRIQMEVQ
KQLHEQLEIQRKMQLQIEKQGKALVMMIEKQNMEFGKPEQEEEAEADSRRSKRPRKG
>BraPSR_LOC103828237
MYYQNQHQGKSILSSSRMHLPSERHHQFLRGGNSLGGSGLILSTDAKPRLKWTPDLHERF
IEAVNQLGGADKATPKTIMKVMGIPGLTLYHLKSHLQKYRLSKNLNGQANSGLNKIGMMT
MMEEKSPDADEIQSETLSIGPQPNKNSPISEALHMQIEVQRRLHEQLEVQRHLQLKIEAQ
GKYLQSVLEKAQETLGKQNLGEAGLEAAKVHLSEFVSKVSAEYPNTSFLEQKEFRNLCTQ
QMQPPDCSLESCLTSSEGAQKNPKTLENNRLGLRTYLGDSTSEQKEPMFKRMELTWTEGL
RGNPYLSTMVSDAEQRVSYSDRSPDRLSIGVGMHGHRGHQQGNNKFKDERFNDKGEDHKL
ETQGTTTELDLNTQVDNYCTTRPKQLDLNGFSWN
>BraPSR_LOC103830485
MFHAKKPSSMNGSYENRAMCVQGDSGLVLTTDPKPRLRWTVELHERFVDAVAQLGGPDKA
TPKTIMRVMGVKGLTLYHLKSHLQKFRLGKQPHKEYGDHSTKEGSRASAMDIQRNVASSS
GMISRNMNEMQMEVQRRLHEQLEVQRHLQLRIEAQGKYMQSILERACQTLAGENMAAAVA
AGGGYKTNLGSTSLSAAVGPQPLPLSFPPFQDLNIYGNTTEQVLDQHNFHHQNIENHYTA
NNAADTNIYLGKKRPNPSYGNDVRKELLMWSDQDHDLSGNQAIDDEHRIQIQMATHVSTD
LDSLSEIYERKSGLPGDEGNNGGKFLERPSPRRSPLSPMMNPNSGLIQGRNSPFG
>BraPSR_LOC103830924
MCLLMASNTNNGNNQKTKMSLVLSTDANPRLKWTCELHHRFVEAVNQLGGPNKATPKSLM
KAMEIPGLTLYHLKSHLQKYRLGKSLKFDDNRLEVSSALETQEAESGNYSRDFRGSVNEE
NNNPANEGLKITEALQLQTEVQKKLHEQIEVVQRNLQVKIEAQGKYLQSVLMKAQQTLAG

YTSSTLGMDFSRTKLSRFASLVNPSSSFSELTQVEEYEEEAEDARESFLYRKKTENRGIK
LLRCSVESFLESSESSETKRNNDNDERISVELPLMEIKSEVMTNKKKRNHNDVVCMECQL
LKKIDFEVDDDEQELKLSLNSYKKNYGDVPEPKKRLGFN
>BraPSR_LOC103831359
MCLLMESNTTNGNNQKTKMSLVLSTDAKPRLKWTCELHHRFIEAVNQLGGPNKATPKGLM
KAMEIPGLTLYHLKSHLQKYRLGKSLKFDDNNLEADSSASETQEAESKNVSTDFRGSVNV
ENNNPANEGVQITEALQLQMEVQKKLHEQIEVQRHLQVKIEAQGKYLQSVLLKAQHTLAG
YTSSTLGMDFARTELSRLASMVNQSSSFSELTQVEEYKEEGFLWCKKQENRGTTHPRRSV
ESSLTSSESSETNLNKNNDERMSVELPLMEIKSEKLTEKKKRSLNDVVCMERQPPKKRNL
GAYDDDDEHLRLSLNSYKKDMGTCPNIGLGFN
>BraPSR_LOC103832766
MEADNGGTNSSHASKQRLRWTHELHERFVDSVAQLGGPDRATPKGVLRVMGVQGLTIYHV
KSHLQKYRLAKYLPDSSSEGKKTDKKESGDVLSGLDGSSGMQITEALKLQMEVQKRLHEQ
LEVQRQLQLRIEAQGKYLKKIIEEQQRLSGALGEPSGPVTGESDPATPAPTSESPLQDKS
GKDCGPDKSLSVDESLSPYREPLTPDSGCNAGSQDESAGEERSSKKPRLVRGGAAGYTPE
MVVAHPILESGLNASYHQPDHALAFDHPSTSLLGTRDKVSGDDL
>BraPSR_LOC103854324
MEARPVQRSGSRELSNLTRTSSIPSTPNPSSAAAAEVAFMRSENNNNNTQFMSKPLGGQN
YHLLASSNGGAVGHICSSSSSGFSTNLHYSSMEKHYAGSSSNAAAAAAASRDDSSWCSDS
LHGGFLDFPENHHPASQIEDGGGIGAAFDDIQKRNDWHEWADHLITDEDPLLSTSWNDLL
LETSSNSDSKDQKTLQVPPQQPQIVQQQPSPSVSVELRPVSTTSSNSNNGNGKARMRWT
PELHEAFVEAVNSLGGSERATPKGVLKKMKVEGLTIYHVKSHLQKYRTARYRPEPSETGS
SEKKLTPLEHITSLDLKGGMCITEALRLQMEVQKQLHEQLEIQRNLQLRIEEQGKYLQMM
FEKQNSGLGKGTASTSDSPSKSEQEDKKVTDSKELTPEGTGKCKEPESPQPKRPKTDN
>BraPSR_LOC103834747
MEARPVQRSGSRELSNLARTSSIPSTPNPSSAPEAAFMRPDHKPLGQQTYHLLSSSNGGS
VGHICSSSSSGFSTNLHYSSMEKHYAAASSNDDSSWCNGGFLDFPEDHQAVHNNSQIEDG
CIGIGAGFDDIQKPNDWQWADHLITDEDPLLSTNWNDLLIDTSSNSDTKDQKSLQPIPPQ
SQTVQQQPSPSVSVELRPVSTTSSNSNNKARMRWTPELHEAFVEAVNSLGGSDRATPKGV
LKKMKVEGLTIYHVKSHLQKYRTARYRPEPSESGSPEKKLTPLEHITALDLKGGIGITEA
LRLQMEVQKQLHEQLEIQRNLQLRIEEQGKYLQMMFEKQNSGLGKGKGTASTSDSPSKSE
QEDNKKTADSEELAPEETRKCQEPESPQPKRVKTDN
>BraPSR_LOC103836250
MDNDKNHKTEEPKPRLRWSYELHHRFIDAVNQLGGPNKATPKGLMRVLEIPELTLYHLKS
HLQKYRLGISERFIGNKQDAGRSQECQSQEDLGDQLDIIVPEEKHDEPNKNLQIKEAVEI
QMEVQKKLHEQIEMQQQLQVRIEAQGKYLQSVLLKAQETLSGYKSSNLYAVASMANRNCL
SSSISALTQADEDNEVEEEYDFLCTKKPENRGNESTRSSVDCSLASSESSEAKLDHRSQT
IMRRSDELQFMEIKPEEVMDRKKRRWDDDVLCVEQSIRKKAFGGLDGEDLGLNLNSFKVM
ETSYKSNYKSNK
>BraPSR_LOC103837308
MTLANDFGYSTAMSSSFSPLHATVEDRYRKFPNSYWGSSGQELMNNPVPYQVVSSSGGYM
SGYCNVSAVSAHGRTSQTQPPVSTMPSDSLAMQSSLINNHHPQEFSDPLDEFFDFSHHVS
APNPQTDGSGVKLVPSVELHEKSEWQTWADQLMSVDNGSEPNWSELLGEPSPHNPNSQVQ
SCHDLQQTPTPPRQEIIANQQHQAVSSEEQLGGRNSSASGATSKQRMRWTPELHEAFVEA
VNQLGGSERATPKAVLKLLKNPGLTIYHVKSHLQKYRTARYKPETSEATGEPEEKKITSI
EDIKSLDMKTSVEITQALRLQMEVQKRLHEQLEIQRSLQLQIEKQGRQLQMMFEKQQKLQ
ENKSSSSELSPKQCNGTSAEVEFGVETQTRNQTEPALASRKRTRED
>BraPSR_LOC103838005
METRNLAHTCSSIPPPDIPFNSQQHNHLLSEAAGHIYSNDLPNAAAASMVSHEQQHIGST
DTPFIPEILDWDHHLLDLPLDFSPIQMEEEDGGILPSDENPIMSPYWNDLILDTSSTSAS
KVHESTMQPQVPLQQPPSPCVELPPLVRTVSSNSNDNTSAAAKGRMRWTPELHEAFVEAV
NHLGGMNNAKPKAVLKHMKVQGLTIYHVKSHLQKYRTARYVPEPSQGSQETKLTPLEHVT
SLDTKRGIDITEALRIQMEVQKQLHEQLEVQRKMQLRIEEQGKVLLMMFEKQNMDFCKPD
QEDKTSEKTPESCSEEADSPRPKRPRNNE

>BraPSR_LOC103845853
MYYHNQHQGKSLLSSSRMPIPSERHPFLRGNGPGDSGLILSTDAKPRLKWTPDLHERFVE
AVNQLGGGEKATPKTIMKVMGIPGLTLYHLKSHLQKYRLSKNLNGQANSSLNKTSVMTMV
EENTPEADESHSESLSIGPQPSMNLPISDALQMQIEVQRRLHEQLEVQRHLQLRIEAQGK
YLQAILEKAQETLGRQNLGPAGIEATKAQLSELVSKVSSEYPDASFLELQNLHHQQMQTA
YPPQNSSLESCLTSSEGNQKAPKMLENRLGLRTYLGDSSSEQKEIMEEPFFHRMELTWAE
EEEEEGIRENNRPYLSAMEPRNSSSRRSPGRLSIGVGLQEHRGGYTEERYRENGEDCKVE
TRTSTALDLNTHDETYGTTRPKQFDLNGFSWS
>BraPSR_LOC103847021
MKFGRRHASDNDPSHEMCSEQATHCCKCYDISEIFSTETRQTTSLSKQSSECVETDISTE
PAGSSLASEFTSSSPIKTRIKWTTDLHDKFVECVNHLGGPMITYYFQLIIKLLEATPKQI
LKMMRTDELTIYKVKSHLQKYRTHKHVQNSMQGTSLKEEEIHLEGSGIKEFMRLQDKVGQ
HLQEQLEIQRELYLVVEEQNKKLHEMITFQKQNNS
>BraPSR_LOC103848918
MTLTGDFGFPPTAMSSFFPLIPTTLDERYHHHKFPNSLWVSSSGQEPMNNPVPCQVFPLA
SGGTSSGYCNGTYVSAQERSSSQTRQSVETQSLPVTNQPQEQRDMSWPEDQLQGFFDFPP
QGESSSRAKSEWPDWANQMISVDDGLEPNWSELLGDPNVLNQDSKIPTPSCDIARQEIVV
STQHQVDSSSAKSPQASSMTSKQRMRWTPELHEAFVDAINQLGGSERATPKAVLKLINSP
GLTIYHVKSHLQKYRTARYKPEISIDTEKPPLKTLKTIEDIKSLDLKTSIEITEALRLQM
EVQKKLHEQLEVQRTLQLQIEEQGRYLQMMIEKQQQKMQEKKIGSSSGTSSMPEADTSSA
PSPNLSQASVTERLQSGGSSTLDQSGYPSGATKKRVRED
>BraPSR_LOC103849708
MMQSREERRSESPSGLVLTTDPKPRLRWTAELHERFVDAVTHLGGPDKATPKTIMRVMGV
KGLTLYHLKSHLQKFRLGKQPHKDHSHGHSTNIRDPNRVSMLDLQRNVVFTTPHITGHNM
NEMQTEVHRRIEEEVELERQVNQRIEAQGKYMESMLEKACETQEASLTKDYSTLFFDRTT
ICNNSSPLTIQWFEDQFPSSSSMDSAMHLPDINSNFSLQDSRSSITKNHTVCLG
>BraPSR_LOC103849734
MNSHRLVVSTAQEECNKGIQKACSSSLSPAFNFLNVQPETTSKSPFIRSQSPDWPKNSTF
SRSSTFCTNLYSSSSSANETQKHLGNSLPFLPDPSASASGVESARSPSVFSEDLGNPFDG
DNNSSLVKDFFNLSADACSNGGGYHDLDCSNDTLSDQLELQFLSDELELAITDRAETPRL
DEIYEKPMASPNPVTVLSPSQRCDAGAMCIDPVSSLPSPRSSAAANHKPRMRWTPELHES
FLKSVNKLEGPEKATPKAVLKLMNVEGLTIYHVKSHLQKYRLAKYMPEKKEEKKNVNSEE
KKLAMSNSEADEKKKGAIQLTEALRMQMEVQKQLHEQLEVQRVLQLRIEEHAKYLEKMLE
EQRKAGRLFSSSSSSQTLLSPSDDETRPDSQNMSKTEASLPQPSSSAKNIASETEDDQCE
SPQKRRRLENNTEPQDSER
>BraPSR_LOC103850633
MDNINFETSNASQGSRLHLHSQSQPPQLFNLQDVNMNHYNQSSPWTSETFSGYTPYDSTV
NQSLSVQCSSSKPCHALFHPYHHQSSDHPSVDQSQSMVPMQLLQDQYLKPLYQKSCANDF
AATNASSSSYSLSFDASQDPQELCRRTYSTSNVTQLDFSSSHHQAKQTHPRFSSHSFSAH
GGSMAPNCGTVGNKTRIRWTQDLHEKFVECVNRLGGADKATPKAILKLMDSEGLTIFHVK
SHLQKYRIAKYIPDYQEGKFEKGSCSKELSQLDTKTGVQIKEALQLQLDVQRHLHEQLEI
QRNLQLRIEEQGKQLKIMIEQQQKTKECLLLKSPNAEASLSHSASDHSPPPYSIQDAEAL
LLTTYGDTQFQSKIS
>BraPSR_LOC103852594
MESNTNNGNNHKAKVSLVLSTDAKPRLKWTCELHQRFIEAVNQLGGPNKATPKGLMKAME
IPGLTLYHLKSHLQKYRLGKSLKFDDNKLEVSSASETQEAESKNDSGNFRGNVNQENNDP
ANEGLKITEALQLQMEVQKKLHEQIEVQRHLQVKIEAQGKYLQSVLMKAQQTLAGYTSAS
LGMDFARSELSRLASMMNPSSSFSEQTQVEDYEEEGFLWCKKPENREKRQPRSSVESSLT
SSESSETKLNNNNEERKSMELPLMEIKSEAMTAKKTKRSLNDVLCVEHQPLKKRDFGVDD
DDEQHLRLSLNSYKQDMGTCPNMD
>BraPSR_LOC103853188
MFHAKKPSRMNGSYENRAMCVQGDSGLVLTTDPKPRLRWTVELHERFVDAVAQLGGPDKA
TPKTIMRVMGVKGLTLSHLKSHLQKFRLGKQPHKEYGDHSTKEGSRASAMDIQRNVASSS
GMMSRNMNEMQMEVQRRLHEQLEVQRHLQLRIEAQGKYMQSILERACQTLAGENMAAASG

GGFKGNLGSSSLSAAMGPHPLSFPPFQDLNIYGNTTDQVLDHHNFHNQNIENHYTANNAA
DTNIYLGKKRPNPSFGNDIRKELLMWSNQDHEPIDDEHRIQIQMATHVSTDLDSLSEIYD
RKPGLSGDEGNDCGKLLERSSPRRSPLSPMMNPNAGLVQGRNSPFE
>BraPSR_LOC103854619
MNNPVPCQAVSGGNSGGYMFPSPSGFCNVSPLSTHGRASQTQPPVATTPSDRLAMQDCFL
EAQSMNHPLPEFSDPLDEFFDFSDHVPALNPQAESSDVMVGSTVEVHEKSEWQSWADQLM
SVDNGSEPNWSELLGDPSPHNLTPPTPSLDVPRQEVVANQQQHQVVSFEEQLNSSASGGA
TTKQRMRWTPELHEAFVEAVNQLGGSERATPKAVLKLLNNPILTIYHVKSHLQKYRTARY
KPETSEATGEPQEKKMTSIEDIKSLDMKTSVEITQALRLQMEVQKRLHEQLETQRALQLQ
IEKQGRYLQMMFEKQQKLEENKSSSSSKQCNGASAEVEFESGVVTQAGDQTESAVSVSRK
RAREDE
>BraPSR_LOC103855697
MDNNFESSNASQGSRLQLHPQPPQPFNLQDVDTIHYSQTSPWTTETFSGFTPYDCIANQS
FSLQCSSSKPYPPSLHSYDHQSSDPPSLDQSQSMVPMQPSPDQYLKPLYKRSCVNDFAAT
NASSASYSLCFGASQDPQEICRGNYSNSNVTQLSFSLSHHQSKQTHSRFSSPSFSTYGGS
MVRNYGTVTGNKTRIRWTQDLHDKFLECVNRLGGANKATPKAILKLMDSDGLTIFHVKSH
LQKYRIAKYIPDPREGKFEKRSCSKELSQLDTKTGVQIKEALQLQLDVQRHLHEQLEIQR
NLQVRIEEQGKQLELMIEQQQKTKESLLKSPNAEVSLPLSAFDHSPPPFSLQDAEAMMLP
SYEDTHFQSKIS
>BraPSR_LOC103859491
MNRHRLVSTAQDECNKGLGQACSSSLSPVHNFLNLQPENRNSPFIRSHSPDSPWPKNSPQ
GTFSRSSTFCTNLYISSSSTSESQKHLGNTLPFLPDPSTNSQPPSAVESARSPSIFSEDM
SNPFDGDNTLVKDFFNLSGDACSDGAFHDLDCSNDSYCLSDQMELQFLSDELELAITDRS
ETPRLDEIYETPLASSSNPVTGLSQTQRSLSGGMSIEVVLSHPSPGSAAASNHKPRMRWT
PELHESFLKSVDKLEGPEKATPKAVLKLMNVEGLTIYHVKSHLQKYRLAKHIPEKKEEKR
NVNTEEKKLALSNNEAAERKKGAMQLTEALRMQMEVQKQLHEQLEVQRVLQLRIEEHAKY
LEKMLEEQRKTGMLMSSSSSQTSDCQNMSKTEVSSLSQPKNIASETEDDKCESPQKRRKV
ENNTESEDPEER
>BraPSR_LOC103831963
MYSAIRSLPLDGGDYHGPLDGTNLPGDACLVLTTDPKPRLRWTAELHERFVDAVTQLGGP
DKATPKTIMRTMGVKGLTLYHLKSHLQKFRLGRQACKDSTDNSKDASCVGESQDTGSSSS
SSLRMAAQEQNEGYQVTEALRAQMEVQRRLHEQLEHGQVQLRLQLRIEAQGKYLQSILEK
ACKAFDDQAAAFVGLEAAREELSELAIKVSNSSQGTAVPFFDTTKMMMMPSLSELAVAVD
TKNNITTNCSVESSLTSNTNGSSVSAASMKKRLRGDDVGLGYEAGWNVPSSSTIG
>BraPSR_LOC103860014
MYSAIRSLPLDGGEYHGPLDGTNLPGDACLVLTTDPKPRLRWTAELHERFVEAVTELGGP
EKATPKTLMRTMGVKGLTLYHLKSHLQKFRQGRQACKESTDNSNKDASCVGESQDTGSSS
PSSLKLAAQEQNESYQVTEALRAQMEVQRRLHEQLEHGQVQRRLQVRIEAQGKYLQSILE
KACKAFEEQAAMFTGLETAREELSELAIKVSNSSQGATVPYFDATKMMMMMPSLSELEVA
AIDHKSNITTTNCSVESSLTSNTNGSSVSAASMKKRHRGGDNVGYEGSWTVPSSTIG
>BraPSR_LOC103870231
MMQSREERRAESPSGLVLTTDPKPRLRWTAELHERFVDAVTHLGGPDKATPKTIMRVMAV
KGLTLYHLKSHLQKFRLGKQPHKEHSHGHSTNIRDTNRASMLDLQRNVVFSTPHIIGRNM
NEMQMEVKRRIEEEVEVERQVNQRIEAQGKYMESMLEKACETQEASLTKDYSTLFFNNTS
PLPIPWFEDHFPSSSSMDSTLNLSDISLNFSLQDSRSSITKNRTVCLG
>BraPSR_LOC103871004
MNSHSLLSIDGGNTSCSSSSLSPLHNFHPTETTTRSFQFKPSHFSNGSFSRCSSFCNLSS
SSSSDTQKHLGSTLPFLPNPLTENERSPAAASFTDGDLLSIPYEEEVDPSFLSLHGDGGF
QDVDNFSLSEEQMELQFLSDELQLAITDRAQTPRLDEIYQVTGSSPGQNCVPAAMPVLSQ
QPSPGGTEAVNQKPRMRWSPELHDCFLEAVKKLDGPEKATPKAVMKMMNVEGLTIYQVKS
HLQKYRIAKHMPERKEEKKSGGNPEDKKPASNTNGEANGRKKGAIQITEALRMQMEVQKQ
LHQQLEVQRSLQLRIEEHAKYLEKILDEQRKATPTSKQESQLSSASSAKDDECQASPKRP
RIEN

>GmaPSR_Glyma.01G009600.1
MEGGGREGYNGVIMTMTRDPKPRLRWTADLHDRFVDAVTKLGGPDKATPKSVLRLMGLKG
LTLYHLKSHLQKYRLGQQAQKQNEEVHKENSRCSYVNFSNRSLAPNTSYRGDDEGGEIPI
AEALRCQIEVQKRLEEQLKVQKKLQMRIEAQGKYLQSVLEKAQRSLSLDGPGSLEASRAE
LTEFNSALSNFMENMNKDSKQNIIEVNNFYSKSHGSAFYNQEVGREQNRDQKPKVEGGSI
QFDLNIKGSNDLVSAGGAEMDANMVSSYRV
>GmaPSR_Glyma.01G049100.1
MDPQSMHFVLSTDSKPRLKWTPELHRRFIEATNQLGGADKATPKSLMRVMGIPGLTLYHL
KSHLQKYRLGKSQELETCSDNKQEDYIETKSSSDGHCSREISIGAQNQLTENMQIAQALQ
MQMEVQRKLHEQIEVQKHLQLRIEAQGKYLQSVLKKAQEALAGYNSSPVGIELTKAELSQ
LVTIINNACPSSPISELTETRGLSLSCGERKRDRGTMCSLESSLTSSESSGRKEEKQPME
EIVEFKSSINVSLQLPLMEILTEDKASNGGSSNEASGRKRSATAAESDDGSCVVEQPCGK
RCGNKLRKAKLSEMLDLNSQCQSDMDSTSSKTLDLNCSLNFWEP
>GmaPSR_Glyma.02G070900.1
MYYQQQQQAKNMHALRMHSPTERHMMMQGGNGSGDSGLVLSTDAKPRLKWTPDLHERFIE
AVNQLGGADKATPKTVLKLMGIPGLTLYHLKSHLQKYRISKNMHGQTNTSNNKIASTMEA
AARISEASGVQMKHLSIGLQTNKNSEINDALQMQIEVQRRLHEQLEQVQRHLQLRIEAQG
KYLQAVLEKAQETLGRQNIGAEGVEATKVQLSELASRVSPQSLDSRFSELKELQVLWPQQ
TQEGQATDCSMGSFLTYSEESQRDRETHNMNLNLRAYNGPPFSVSKGSVEESMHLKPDHT
LCDEVKENLMFLSSSSDSKVVKGDFLSERPSSLLSMNEGVQEEENFGRTTVPKEEGWKGR
KSTETGRAPVKLNHEKISQDYRLANFEVKLDLNSHDDNDASSHCQQFDLNGFSWNC
>GmaPSR_Glyma.02G108500.1
MDPQNMQNQTMHFVLSTDSKPRLKWTPELHRRFIEATNQLGGEDKATPKSLMRVMGIPGL
TLYHLKSHLQKFRLGKSQQLETCSDNKQEDYIETKSSSDGHCSREISLGAQNQITENMQI
AQALQMQMEVQRKLYEQIEVQKHLQLRIEAQGKYLQSVLKKAQEALAGYNSSPVGIELTK
AELSQLVTIINDACPSSPISELTETRGLSLSCGERKRDRGTMCSLESSLTSSESSGRKEE
KQPMEEIVEFKSSNNVSFELPLMEIHTEDKASNGGLSSNEGSGRKRSLAATESDDGSCVV
EQPCGKRCGNKLLRKVKLSEMLDLNSQCQSDMDSTGSKTLDLNCSLNFWEP
>GmaPSR_Glyma.02G177800.1
MFGSVVDSFLSSDEDTFCETYREYSELPCHDVSLYEHFRHEEKRLEGDDSAPVIEAEVVC
ATSGNSASSMVPTRKNRIKWTKDLHEQFVAAVNSLGGPQKAKPKAVLQMMNSKSLTIFHV
KSHLQKYRTTMYMQNSSKEGYKESKGIDMVTELQQKIYMQIEESRLLQLEIGRGIQEQLE
AQRNLQMLVEEQKKQVNSVCGKNQIKQIGGSKS
>GmaPSR_Glyma.02G178100.1
MNENRIDCVGRTQQSYGLNGDWNSEFGNCSSQYFDVRQASNMGTCNQPLAMASGGGVEQE
PHIGQNKSSSSIISRFKSPASAFYATEMCMGGFPQYDSQIGNPSLMSHSSKFNDMEFPLY
QSLRQSLFMPSLANQPPPKFDLSNPLQEMLKFHLNSDQCVRSLETFNKIPCGDFPGSNFL
PIEQHKLFIDDAAPISRSPSIPSKGNQGQTVSCGSFNLPSAQLSFSSQQEMLSPTGSMPT
NSGNSSSNGSVVSSKTRIRWTQELHEKFVECVNRLGGAEKATPKAILRLMDSDGLTIFQV
KSHLQKYRIAKFMPQPTQGKSDKRTNAENVHLDVKTGFQIREALQLQLDVQRRLHEQLEI
QRKLQLRIEEQGKQLKMMFDQQQKTTDSHLITENSDRPISSKDVLVSIYEGSENSLFSSK
IS
>GmaPSR_Glyma.03G003500.1
MVPHKSQGGVEQLANAGVLGGSAVKIAAAPAGGSGKQRLRWTSDLHDRFVDAITQLGGPD
RATPKGVLRVMGVPGLTIYHVKSHLQKYRLAKYLPESPADGKDSKVEKRNSGDSISGADS
SPGMPINDALRMQMEVQKRLHEQLEVQKQLQMRIEAQGKYLQKIIEEQQKLGSNLTTSEA
LPLSHDEQNHPQSEASGSSEALASTVSPLKKQRIDDGSKDGFTASQVRNAAQKNDCNVGQ
LDPNLYDDAGFEFDLETKKDEDNESGQ
>GmaPSR_Glyma.03G143500.1
MYQYSSQLYGTEWESYMGISNLDKVVGSEQLSFEPTKSPFNIVTTPLQITPPGFCASAIK
SLVSFQAQQQQQHAHHQHQIEVPAAWCFEFPKTTTTTDTHMVMNNICQASGDNFTTKQDR
PSSQFTSSFLSSSGADCRSSSEKYCKIASYSEEKHSSIQPDSIQYYDLYSQEDDKLLRDD
AATDEGPLEISFQRNQLDSCTKREKQAPHRTCGVACVASRRGKRRIKWTKDLHEPFMMIV
NSLGGPEKAKPKAILDMMKSDLLSISHVKSHLQKCRSTIHMHKALQERSEKGQRTNGVSE

LQVKIHMQIEESRQLQLEIRRNICQQLKMQRNLHTLIQEQSQQLKVMLDYQKERTKLEKT
PYTELEATVPK
>GmaPSR_Glyma.03G143600.1
MNENKIDWGGLIQQSHVLSGDFNSEFGNRYCQYFDIRQAWNMGPLAMVGGEAIDNELQNI
GHAKSSGTIMSRFESPASAFYAAENCMEFAEYDCQVGIHSLSSQLCKINDLEFPLYQSPR
ENLFLDSANQSETNFDLSNTLQSIVKSQLNSANQCSRSPEKSNKISCGNFHSTKFLPVEQ
QKLFIDGLISGSSFPNKGNQDHMLRFSSQIERLSPTLSAGSVSTIGNSASNVAAVVSSKT
RIRWTKDLHEKFVECVNRLGGAEQATPKAILKMMNTDGLTIFHVKSHLQKYRIAKFIPEP
SHGSFPILNLKNCPVSGKSDKRTHTKDVHHLDVKTGIQIREALKLQLDAQRCLHEQLEIQ
RKLQLRIEEQGRQLKKMFDQQQKTSNDVSNTQNSTIEETSISHKDGENASEGANNNSFFP
SKTSPSQ
>GmaPSR_Glyma.03G166400.1
MEAHSAFSIERSNANDMGMSEVFPSSLPVLPSPLEETFPKLSDSRPAFMEKELKTKPFTH
SSHLTSSGAVGHMFSSSPGYSTDLHHSSFSSYEKQPRNTHFISQPLGNMASLPLSYSSNS
EPIPSTTSTPYSNGNSVSWHTDSLPSFLDFTANTSIDNSQVENRACNIMATEEYSKRNDW
QEWADQLISDVDPLTSNWNDLLADNIQDLEPKVTKSSSQLPIEHQSQSHQQLCASSGENR
VGVAPTSSANSAPAKPRMRWTPELHEAFVEAVNQLGGSERATPKGVLKLMKVEGLTIYHV
KSHLQKYRTARYRPESSEGAAEKNLSRIEEMSSLDLKTGIEITEALRLQMEVQKRLHEQL
EIQRNLQLRIEEQGRYLQMMFEKQCKPGIETFKASSSVIESQSGVSSDAIKDSPAKTESE
TIKVDHCKSGADLANGSTTVEESSLEVAEKQDAPEIQASDNPEQHASEDSGNASKRPRTE
E
>GmaPSR_Glyma.03G250000.1
MTLVICQCLPLTNGQTKLIHLGASRAMSTSSRVLPTPLENKYMKPPDSFQLSPVRDLTAN
SASSNSIRSAGKMFSSPSKCPDDFPFSSVSQHDRQYQDPPFVSQTIGDSVSSEIHSMTFI
SHPQENEDLSWGPDPCQDILGFPENVSVQHDQVENNGCYINDDNVKRSDFGEWVDQLMSI
DDSLHPSWSQLLADDNVAEPKPKASQVPPQQHIPSGEVVGNSASTASQTKARMRWTPELH
EAFVEAVNHLGGSEKATPKGVLNQMKVEGLTIYHVKSHLQKYRTARYKPEPSEGTSEKKV
TPMEEMKSLDLKTSKGITEALRLQMELQKRLHEQLEIQRKLQIQIEDQGKRLQMMFEKQR
EMGDSKVNVSLDEPSAAAPSETVETTNEERHKFESIPKAIPEEKESSTTKQIAGEAEEMI
KEDEVAPPTKRVKSS
>GmaPSR_Glyma.07G229800.1
MDLQNVQNHSMMRLVLSTDAKPRLKWTPELHQRFTEAINQLGGAERATPKSLMRVMGIPG
LTLYHLKSHLQKYRLGKSQPLETCSDNKQQGYCEIQNSDGHFSKEISIGTQNQMTESLKI
AEALQMQMEVQRKLNEQIEVQKHLQRRIEAQGKYLQSVLTKAHEALARHSSSTTGMELAK
AELYQLESIINNACPDSPLSELTETRGLSLNCGERKQDRGTMCSLESSLTSSESSEQHTM
DEAENPQKFNGVSVELPLMSIHPEEDKAFKGDTSDGRKRSAATDSDHCVDQPCGNKLRKS
EVSEMLDLNCQYQRDIDSSVKEIDLNFSSSF
>GmaPSR_Glyma.08G163500.1
MERMFPPKKPSTMNSHDRPMCVQGDSGLVLTTDPKPRLRWTVELHERFVDAVTQLGGPDK
ATPKTIMRVMGVKGLTLYHLKSHLQKFRLGKQPHKDFNDHSIKDGMRASALELQRNTASS
SAMIGRNMNEMQIEVQRRLHEQLEVQKHLQLRIEAQGKYMQSILEKAYQTLAGENMASAA
TNLKSAIVPHHQGIPDMGVVMKEFGSPLGFSSFQDLENIYGGNQIDLQQNMEKPSLDHGF
MPINESLCLGKKRSNNPYSGSGKNPLIWSDDLRLQDLGGPASSCLGPQDDPFKGDQIQIA
PPGSLDRGASTDIDPMSEIYDSKPVLQSEEKKFDASSMKLERPSPRRAPLQPERMSPMIS
TGTMAQGRGSPFG
>GmaPSR_Glyma.09G017300.1
MYPRLIHPHDGIVTQDELQGGAASNLSHAHKGDPCLVLTADPKPRLRWTQDLHERFVDAV
TQLGGASKATPKAIMRTMNVKGLTLFHLKSHLQKYRLGKQSGKDVGEGCKDGSYLLESPG
ADNTSPKLPTPDTNEGYEIKEALRAQMEVQSKLHLQVEAEKHLQIRQDAERRYMAMLERA
CKMLADQFISATVIDTDSQKFQGIGSKAPRGTLVDPLGFYSLPSTEVAGVNVPEEEILPS
LPPQRADCSTESCLTSHESSGGLALEGSPGEGKRRMLGMDSMAAPLIWSEAKMRTQAINV
AQGNHPQGITRYGM
>GmaPSR_Glyma.09G017400.1
MYHSKNFPSASLIGVNSLVHGQHIDCGGSTMDPGSGGNSLGNNSNLASKQRLRWTHELHE

RFVDAVAQLGGPDRATPKGVLRVMGVQGLTIYHVKSHLQKYRLAKYLPDSSSDEGKKADK
KETGDMLSNLDGSSGMQITEALKLQMEVQKRLHEQLEVQRQLQLRIEAQGKYLKKIIEEQ
QRLSGVLSETPGSGVAAVAPGDACQEPDNKTDPSTPDPEKAAKDRAPAKSLSIESFSSHL
EPMTPDSGCHVGSPAESPKGERSAKKQRVIMDGVYSKPEMVLPHQILESSMSLYQQPNTV
FLGQDQFDPSLGISTRSGEELDKVGGGNL
>GmaPSR_Glyma.09G113000.1
MNENRIDSCVGRIQQSYGLNGDWNSEFGNCSSQYFDMRQASNMGTSNQPLAMASGGGVEQ
EPNIGQTKSSSSIISRFESPASAFYATEMCMGGFPQYDSQIGNNLSLMSQSSKFSNMEFP
LYQSPRQSLFMASLANQPPPNFDLSNPLQEILKSHLNSDQCVRSPEDSNKIPYGDFPGSN
FLPIEQHKFFIDDAISTGISPSIPSKGNQDQSVSCGPFNLPSAQLSFSSQQEMLSPTGSM
STTSGNSNSNGPVVSSKTRIRWTQELHEKFVECVNRLGGAEKATPKAILRLMDSDGLTIF
HVKSHLQKYRIAKFMPQPTQGKSDKRTNVENVHLDVKTGLQIKEALQLQLDVQRRLHEQL
EIQRKLQLRIEEQGKQLKMMFDQQQKTSNGHLITENNDRPISSKDVLVSTSDGSENSLFS
TNIS
>GmaPSR_Glyma.09G113100.1
MYHFSQVSNGEHSSVANVGGLETDSFNFQQHVFDAQFSCLKFDSQNLLCQSSGDNSAPIL
FGSVVDSFLSSDEDTFCDTYSEYSELPCHEVSLYEHFRHENNILESNYSAPVNEVEVVCA
TSGMVPTRKNRIKWTKDLHEQFVVAVNSLGGPQKAKPKAVLQMMNSKLLTIFHVKSHLQK
YRTTMYMQNTTKEGYKESQGRDMVTELQQKIYMQLEESRLLQLEIERGIQEQLKAQRNLQ
MLVEEQKEQVNSVTGQNQTKQTGGSKSLEEK
>GmaPSR_Glyma.09G211400.1
MEGGGREGYNGIVMTMTRDPKPRLRWTADLHDRFVDAVKKLGGPDKATPKSVLRLMGLKG
LTLYHLKSHLQKYRLGQQARKQNEDMHKENSRCSYVNFSNRSSAPNTSYRGDDEGGEIPI
AEAMRCQIEVQKRLEEQLEVQKKLQMRIEAQGKYLQAMLEKAQRSLSLDGPGSLEASRAQ
LTEFNSVLSNFMENMKKDSKENIIEVSDFYSKNHDSAFHYQEVGRDQKPKVEGGSIQFDL
NIKGSNDLVCAGGAEMDANMISYRV
>GmaPSR_Glyma.10G039700.1
MEAHPTFSIESSKQLNNMGMSGALSSSLSILPIPPEELFPKLPESQLDFVEQELMIRPFT
HSSYLNSGGVIGHIFSSSPGYSTDLHHSTLSSAEKHSTNAHLISQSSTNITQFPLSYSSN
IGPPASATPSHYSKENSVSWHTDSLPGFLDFPENGSIDNNPVESSACPIMASEEYSKQND
WQEWAERLISDDGTLTSNWNDLLADNIQDLEPKGPFQVSKPLSQIPGHQSQGHQQLPASY
GENCAGAALSSSANSAPAKSRMRWTPELHEAFVEAVNQLGGSEKATPKGVLKLMKVEGLT
IYHVKSHLQKYRTARYRPESSEGVMDKKTSSVEEMSSLDLRTGIEITEALRLQMEVQKRL
HEQLEIQRNLQLRIEEQGRCLQMMFEKQCKPGTETFKAPSFTTIETPFGMSSNATKDSLS
KNEMEASLVLDHCRSGPDQVNGSTRVEEGSLEKCGKPDSPKTQHAIASEDSAQAPKRQRT
E
>GmaPSR_Glyma.10G196600.1
MQKRREESDKIPVKRDSASSLPQPQSLVLFTSPFSPFPFLLPLSFAKWVIISFSHNTIFF
TFSFFFASLRWGFKFHHRNRMFSRLIHPHEGQEDMQGGSNHAHLGDPCLVLTSDPKPRLR
WTADLHERFVDAVTQLGGASKATPKAIMRTMNVKGLTLYHLKSHLQKYRLGKQSGKDSDE
GLKDGMSASYLQESPGTDNSSPKLPASDANEGHEVKEALRAQMEVQSKLHLLVEAEKHLQ
IRQDAERRYMGMLERACKMLADQFIGDVIIDRDGQKFQGLENKTSRSPLVDHGGFFPAAC
TEVGGMHVSEVPPILQPQGAECSSESCLKSLESLGGLTLEGSPGGSKKRMLNLDSMVAPL
IWSEANTRTQGIHLAQVNPPGMTRYGM
>GmaPSR_Glyma.11G183400.1
MYSTHQQHQGKNIHSSSSSSRMPIPSERHMFLQTGNGSGDSGLVLSTDAKPRLKWTPDLH
ARFIEAVQQLGGADKATPKTVMKLMGIPGLTLYHLKSHLQKYRLSKSLHGQSNNATHKIT
INSGSATDERLRENNETHVMNNLNLAPQSINKDLHISEALQMQIEVQRRLNEQLQVQRLL
QLRIEAQGKYLQAVLEKAQETLGRQNLGVVGLEAAKLQLSELVSKVSSQCLNSAFSELKE
IQGFSPHHQKQTQTNNNQPINANDCSMDSCLTSCEGSSQKDQQEIQNRGMNLIPFNVHTF
MEGPNLNNLPNTDLKWCDPVKKNNTFLTRLSMHAERSPSNLSMSIGLLEGETTENRSTIV
RTESIKPAVAEKVSQDYGLPSNYFAASKLDQTTEDNKDTKTSCKQLDLNGFSWN
>GmaPSR_Glyma.12G089100.1
MYTHQQHQGKNIHSSSRMPIPSERQMFLQTGNGSGDSGLVLSTDAKPRLKWTPDLHARFI

EAVQQLGGADKATPKTVMKLIGIPGLTLYHLKSHLQKYRLSKSLHGQSNNMTHKITINSG
AATDERLRENNGTHMNSLNLAPQSNNKDLYISEALHMQIEEQRRLNEQLEVQRLLQLRIE
AQGKYLQAVLEKAQETLGRQNLGAVGLEATKLQLSELVSKVSSQCLNSAFSDRLKEIQGF
SPHQQTQTNQPNTNDCSMDSCLTSCEGSQKEQEIQNGGMSLRPFNVHTFMERKEVIEGPN
LNNLPNTDLNWCDPVKKNTFLTPLSMHADKRSPSNLSMSIGLEGETENGSTIRTESVKPV
ADKVSQDYGLPSNYFAASKLDLTTEDNKDTKTSCKQLDLNGFSWN
>GmaPSR_Glyma.12G184700.1
MYHHHQHQGKNIHSSSRMPIPSERHMFLQTGNGSGDSGLVLSTDAKPRLKWTPDLHARFI
EAVNQLGGADKATPKTVMKLMGIPGLTLYHLKSHLQKYRLSKNLHGQSNNVTYKITTSAS
TGERLSETNGTHMNKLSLGPQANKDLHISEALQMQIEVQRRLNEQLEVQRHLQLRIEAQG
KYLQSVLEKAQETLGRQNLGVVGIEAAKVQLSELVSKVSSQCLNSAFTEPKDLQGFFPQQ
TQTNPPNDCSMDSCLTSSDRSQKEQEIQNGLRHFNSHVFMEHKEATEAPNNLRNPELKWC
EDGKKNTFLAPLSKNEERRNYAAESSPNNLSMSIGLERETENGINLYPERLITESQSDGE
FQHRNRIKPETLKPVDEKVSQDYRLPASYFAAARLDLNTHGDNEAATTCKQLDLNRFSWS

>GmaPSR_Glyma.13G126200.1
MEAHPTFSIERSKQLNNMGMSGALSSSLSILPIPPEEMFPKLPQSQLDFVEQELMTRPFT
HSSYLNSGGVVGHIFSSSPGYSTDLHHSSLSPDEKHSTNAHLISQSSTNITQFPLSYSSN
TGPPTSATPSHYSKESSVSWHTDSLPSFLDFPENGSIDNNRVESSACPIMASEEYSKQND
WQEWAERLISDDDTLTTNWNDLLADNIQDLEPKVPFQVSKPLSQIPGHQSQGHQQLPASY
GENCIGAALSSSANFAPAKSRMRWTPELHEAFVEAVNQLGGSEKATPKGVLKLMKVEGLT
IYHVKSHLQKYRTARYRPESSEGVMEKKTSSVEEMASLDLRTGIEITEALRLQMEVQKRL
HEQLEIQRNLQLRIEEQGRYLQMMFEKQCKPGNETFKAPSSIIETPSGGSSNATKDSLAK
NEMEASQVNHGRSGPDQVKGSTTFEEGSLEKCGKPDSPKTEHAIASEDSAQAPKRQRTE

>GmaPSR_Glyma.13G316600.1
MYHHHQHQGKNIHSSSRMPIPSERHMFLQAGNGSGDSGLVLSTDAKPRLKWTPDLHARFI
EAVNQLGGADKATPKTVMKLMGIPGLTLYHLKSHLQKYRLSKNLHGQSNNVTHKITTSAT
TGERLSETNGTHMNKLSLGPQANKDLHISEALQMQIEVQRRLNEQLEVQRHLQLRIEAQG
KYLQSVLEKAQETLGRQNLGIVGLEAAKVQLSELVSKVSSQCFNSAFTELKDLQGFCPQQ
PQTNPPNDCSMDSCITSCDRSQKEQEIQNGLRHFSSHMFMEQKEAKEAPNNLRNPEIKWY
DDGKKNTFLAPLSKNEERRNYAAECSPSNLSMSIGLERETENGSSMYPERLITESPSDGR
IKPQTMKPVDEKVSQDYRLPTSYFAAARLDLNTHGDNEVL
>GmaPSR_Glyma.15G123000.1
MYSRLIHPHDGIVTQDDLQGAASNLSHAHKGDPCLVLTADPKPRLRWTQDLHERFVDAVT
QLGGASKATPKAIMRTMNVKGLTLFHLKSHLQKYRLGKQSGKDVGEGCKDGSYLLESPGA
DNSSPKLPTSDTNEGYEIKEALRAQMEVQSKLHLQVEAEKHLQIRQDAERRYMAMLERAC
KMLADQFIGATVIDTDSQKFQGIGSKAPRGTLVDPLGFYSMPSTEVAGVNVPEEEIPLSL
PPQRADCSTESCLTSHESSGGLALEGSPGEGKRRMLGMDSMAAPLIWSEAKMRTQAINVA
QGNLPQGITRYGM
>GmaPSR_Glyma.15G123100.1
MYHSKNVPSASLIGGNSLSHGQHIDCGGSTMDPGSGGNGLSNNSNLTSKQRLRWTHELHE
RFVDAVAQLGGPDRATPKGVLRVMGVQGLTIYHVKSHLQKYRLAKYLPDSSSDEGKKADK
KETGDMLSNLDGSSGMQITEALKLQMEVQKRLHEQLEVQRQLQLRIEAQGKYLKKIIEEQ
QRLSGVLSEAPGSGAVAVVPGDACQEPDNKTDPSTPDPEKAAKDRAPAKSLSIESFSSHP
EPMTPDSGCHVGSPAESPKGERSAKKQRVTMDGVYSKPEMVLPHQILESSMSSYQQPNTV
FLGQEQFDPSLDISTKSDEELVKIGGGNL
>GmaPSR_Glyma.15G215000.1
MERMFPPKKPSTMNSHDRSMCVQGDSGLVLTTDPKPRLRWTVELHERFVDAVAQLGGPDK
ATPKTIMRVMGVKGLTLYHLKSHLQKFRLGKQPHKDFNDHSIKDGMRASALELQRNIGSS
SAMIGRNMNEMQMEVQRRLHEQLEVQKNLQLRIEAQGKYMQSILEKAYHTLAGENMATNM
KGMGAPLGTTEMGVMKEFGSLNYPSFQDLNIYASGDQQLDGFMPSNNNNETLFVGKKRPN
CPYSGSGKSPLIWSDDLRLQDLGTASSCISPQDDPFKGDHKVQISPPSMDSDPISEIYDT
KPMILHGESVSDQNKFDASMKVERPSPKRSPLQPDRTSPMINSSSVAQGRSSPFG

>GmaPSR_Glyma.15G263700.1
MERMFPPKKPSTMNSHDRPMCVQGDSGLVLTTDPKPRLRWTVELHERFVDAVTQLGGPDK
ATPKTIMRVMGVKGLTLYHLKSHLQKFRLGKQPHKEFNDHSIKDGMRASALELQRNTASS
SAMIGRNMNDNSHMVDAIRMQIEVQRRLHEQLEVQKHLQLRIEAQGKYMQSILEKAYQTL
AGENMASAATNLKGAIVPHHQGIPDMGVVMKEFGSPLGFSSFQDLENIYGGDQIDLQQNM
EKPPLDHGFMPINETLCLGKKRSNNPYSGSGKSPLIWSDDLRLQDLGGPASSCLGPQDDP
FKGDQIQIAPPGSLDRGASTDIDPMSEIYDSKPVLQSEEKKFDASSMKLERPSPRRAPLQ
AERMSPMISTGTMAQGTYNTEFRKIEQ
>GmaPSR_Glyma.16G152200.1
MYYQQQQQAKNMHALRMHSPTERHMMLQGGNGAGDPGLVLSTDAKPRLKWTPDLHERFIE
AVNQLGGADKATPKTVLKLMGIPGLTLYHLKSHLQKYRISKNMHGQTNTSNNKIASTMEA
ATGISEASGVQMKHLSIGLQTNKNSEINDALQMQIEVQRRLHEQLEQVQRHLQLRIEAQG
KYLQAVLEKAQETLGRQNLGAEGVEAAKVQLSELASRVSPQSLDSKFSELKELQVLWPQQ
TQEGQATDCSMGSFLNYSEESQRDRETHSMNLNLRACNGPPFSVSKGCAEESMHLKPDYR
LCDEVKENMMFLSSSSNSKVVKGDFLSERPSSLLSMNVGVQEEENFGRTTVPKEEGWKRR
ESTETGRVPVKLNYEKISQDYRLANFDVKLDLNSHDDNDASSHCQQFDLNGFSWNC
>GmaPSR_Glyma.18G201800.1
MGSKQRLRGSATAKERLRWTQELHDRFVEAVNRLGGPERATPKGILKEMKAMGVSELNIY
HVKSHLQKYRISKLIPESPTRGKLEKRSMSDILPNFSSITALQLKEVLQMQTGMQNRLRD
KTEVQRSLKLKIEAQGKYFERIGQSNHSKTIIGKACKPFASTIASLPSLFEESESLKTQP
EEEHQSAKKQKISGEGVFPTSFDHESSTPPECYNETWDFSWSQLAAACQSPLVPSFL
>GmaPSR_Glyma.19G122700.1
MYHTKKFSPASMVPHKSQGGAEQLANAGVLGGSAVKTAAPSGGSGKQRLRWTSDLHDRFV
DAITQLGGPDRATPKGVLRVMGVPGLTIYHVKSHLQKYRLAKYLPESPADGKDPKDEKRM
SGDSISGADSSSGMPINDALRMQMEVQKRLHEQLEVQKQLQMRIEAQGKYLQKIIEEQQK
LGSTLTTSETLPLSHDKQNHPQSEASGSSDALASTVSPLKKQRIDDGSKEGFTASQVRKN
DNVGQLDPNLYDDAGFGFDLETKKDEDNESGQ
>GmaPSR_Glyma.19G146500.1
MYQYSSQMYGTDWESYMEISNLAKVVGSEQLSFEPTKSPFNIVTTPLQTPPGFCASATKS
LVSFQAQQQHTHQHQSEVPAWCFEFPKTTTMTTTTADSHMLNIRQASGDNFTTTITIKQD
PPSSQFTSSLCRPVAESFFSSSGADCRSSSEKYFKIASYSEKHSCIQPDSIQYYDLYSQE
DDTLLRDDAATDEGPLEISFQRNQREKQAPHRLCGVACVASSNSASRRGKRRIKWTKDLH
EPFMMIVNSLGGPEKAKPKAILDMMKSDLLSISHVKSHLQKCRSTIHMHKALQERSKEGQ
RTNGESELQVKIHMQIEESRQLQLEVRRNICQQLEMQRNLHTLIQQQNQQLKVMLDYQKE
RTKLEKTLYTEIEATVPK
>GmaPSR_Glyma.19G146600.1
MNENKIDWGGLIQQSHGLSGDFNSEFGNRYCQYFDIRQAWNMGPLSMFGGEATDHELPNI
GHVKSSGTIMSRFESPASAFYATENCMGFAEYDCQVGVHSLSSQLCKINDLEFPLYQSFS
RENLFLDSANQSETNFDLSNTLQSIVKSQLNSGNQCRRSPEKSNQISSGNFPSSKFLPIE
QQKLFVDGLIRGSSFLKKGNQDHMVGHGSFNLSVPQLRFSSQIEKLYPTLSAGSVSTIGN
SASNGAIVSSKTRIRWTKDLHEKFVECVNRLGGAEQATPKAILKMMNTDGLTIFHVKSHL
QKYRIAKFIPEPSHGKSDKRTHTKDVHHLDVKTGLQIREALKLQLDAQRRLHEQLEIQRK
LQLRIEEQGRELKKMFDQQQKTNNNLPNTQNSATDETTINH
>GmaPSR_Glyma.19G167500.1
MEARSAFSIDRSNAKQLNNMGMSEAFPSSLPALPSPLEETYPKLSDSKPVFMEKELKTKP
YTHSSHLTSSGAVGHMFSSSPGYSTDLHHSSFSSHEKQPRNTHFISQSLSNMASLPLSYS
SNSEPIPSTTSTPYSNGNSVSWHTDSLPSFLDFPANTSIGNSQVESSDCNIMATEEYSKR
NDWQEWADQLISDVDPLTSNWNDLLADNIQDLEPKVAKSSSQLPIGHQSQSHQQLPASSG
ENRVGVAPTSSTNSAPAKPRMRWTPELHEAFVEAVNQLGGSERATPKGVLKLMKVDGLTI
YHVKSHLQKYRTARYRPESSEGAAEKKLSPIEEMSSLDLKTGIEITEALRLQMEVQKRLH
EQLEIQRNLQLRIEEQGRYLQMMFEKQCKPGIETFKASSSAIESQSGVSSDAIKDSPAKT
ESETIKVDHCKSGADQANGITTVEESALEVGEKQDAPESQASENPEQHASEDSAKASKRP
RTEE
>GmaPSR_Glyma.19G247600.1

MYTPCAGQTKLIHLGASRAMSTSSRALPTPLENKYMKPPDSFQLSPVRDLTANSASSNSI
RSAGKMLSSPSECPDDIPFSFVSQTSGDNVSSEIHSTALISHPQDNEDLSWGPDPFQDIL
GFPENVSVQHDQVQNNGCYINDDNVKRSDFGEWVDQLMSIDDSLHPNWSQLLGDDNVAEP
KPKASHVPQQQHIASVEVVGNSASTAPQTKPRMRWTPELHEAFVEAVNQLGGSDKATPKG
VLNLMKVEGLTIYHVKSHLQKYRTARYKPEPSEGNSEKKVTPMEEMKSLDLKTSKGITEA
LRLQMELQKRLHEQLEIQRKLQIQIEDQGKRLQMMFEKQGEMGDNKVNGSSDTNEEGDKF
ESIPKAMPEEKDSSTRKQIAGEAEEVINEDEAAPPTKRVKSS
>GmaPSR_Glyma.20G035300.1
MDLQNVQNQSMMRLVLSTDAKPRLKWTPELHQRFTEAINQLGGAEKATPKSLMRVMGIPG
LTLYHLKSHLQKYRLGKSQPLETCSDNKQEGYSEIQNSDGHCSKEISIGTQNQMTESLKI
AEALQMQMEVQRKLYEQIEVQKHLQLRIEAQGKYLQSVLTKAHEALARHSSSTTGVELAK
FELSLLVSIINNACPSSPISELTETRGLSLNCGERKQDRGTMCSLESSLTSSESSEQQHI
MDEAENPQKFDGVSVELPLISIHPAEVKAFKGDTSVDGRKRSAETDSDHCVDQPCGNKKL
RKSEVSQMLDLNSQYQRDIDSSVKEIDLNFSSSF
>GmaPSR_Glyma.20G193600.1
MIGGKQRAANAKKEEKKGKAAESDKIPVKSDSASSLKPKSCAFHIPIFPFPFLLPLSPNG
SPITQSSPFFFPLSFFFASRWGFKFHHRMFSRLIHPHEGQEDMQGGSNHAHLGDPCLVLT
SDPKPRLRWTADLHERFVDAVTQLGGASKATPKAIMRTMNVKGLTLYHLKSHLQKYRLGK
QSGKDSDEGCKDGMSASYLQESPGTDNSSPKLPDANEGHEVKEALRAQMEVQSKLHLLVE
AEKHLQIRQDAERRYMGMLERACKMLADQFIGDVTIDMDGQKFQGLESKTSRSSLVDHVG
FYPQACTEVGGMHASVVSPILQPQGADCFTESCLTSLESLGGLTLEGSPGGSKKRMLNLD
SMVAPLIWSEANTRTQGIHLAKVNPSGMTRYGM
>MtrPSR_Medtr1g053800.1
MYINHWDYEHQSYEPCSIANPLEDISAIIHESNANETYMNFQAQQHHEFDIPIWSNEFSM
PTTTDSPFLYCQGCKDNYSENITKQDQPTTCFKLGSESLMSSSEDSISSCEKCSEFPSCS
DKRVLESDFSPDHKSHEISFQKTQWESCTKQDKQSSPCGFDFATSTNSDFKITAKGKRRL
RWTKELNESFIMIVNQLGGPEKAKPKAILKMMGVDVLTISHVKSHLQKYRSTLHTHKCLK
GISEEVQITDGINELQVKIQMQIEESRQLQLEVERSNQRQFEIQRNLQLVIEQQKKQLKL
MLDQQKKITKQEKMIDLKRK
>MtrPSR_Medtr1g053835.1
MNENKIDYLGSVQQNHILNGDFNSEFGNCSSQYFDMRNASNIGNFSQPLVMEQSSYIVQS
QNQNQGQGKSSSSSSSSTIMRSFESPTSAFYATEICMGFPQYDYQVGNESSNPLLISQFS
NKVNDLEFPLYQRENHYLDSTNQSSHNFELSNLNTLQPILRSPEKSNRIECGNFPRENYL
SVEQHKFFIDDAASVSMSPLIHSNGNQDHKVSCGSYDFPGSQLNFSYQQDKLSPTMSTGN
VSTNSGNPACNGSSVSSKTRIRWTQDLHEKFVECVNRLGGAEKATPKAILRLMDSDGLTI
FHVKSHLQKYRIAKYMPEPAQGKSEKRTHVENVNLDAKSGLQIREALQLQLDVQRRLHEQ
LEIQRKLQLRIEEQGKQLKMMFDQQQKTNSTCQLNTQNLDNTPNNDTPISPKDIEVTIFE
GSHSQYS
>MtrPSR_Medtr1g080330.1
MSSSIPSSSMQAASINSNIRSVGHMFSTPSEQPDNVHFSSASEIHSMTFPQESDVMSWGT
DPFEDILQFHDNVPTQNDHVEYNGSEVLGGNAKTTDFKEWVDQLMSVDDDSIQPNWNELL
GDNNMAEPKSQDAQMSPSLLMQETQVSQQQYIPSLPSKEVNDLPNSSVSTTSQSKPRMRW
TPELHEAFVEAVNQLGGSEKATPKGVLNLMKVEGLTIYHVKSHLQKYRTARYKPESSEGI
PEKKLTSIDEMPSIDLKTPKGITEALRLQMELQKRLHEQLEIQRNLQIQIENQGKHLQMM
FEQQMKSDEPSAPLSSAAVPSPVENLENTNEGHEKIGINGSASENMPEGSSQNTSTEQKG
DDAKATGELELGEDQLTAPPTKRVKTDK
>MtrPSR_Medtr1g090670.1
MFSRLIRPHDHEGSVVVVQEDSNHHLHTHLAASDPCLVLTSDPKPRLRWTTDLHQRFVDA
VTQLGGPTKATPKAIMRTMNVKGLTLYHLKSHLQKYRLGKQAGKDFDEGCKDGSYLLESP
GTENSSPKLPASDANEGHEVKEALRAQMEVQSKLHLLVEAEKHLQIRQDAERRYMAMLER
ACKMLADQFIGDTTIDTDIQKFQELPSTELGGNGMHISEVPYILQPQGANCSTESCLTSL
ESLRGLTLEGSPSGTKRMLGLDSMVSPLIWSETNMRTQGIHLAQVNPQGIARYGM
>MtrPSR_Medtr2g027800.2
MFPRLIHPHDGIVVGQDDLQGGVSNHKGDPCLVLTADPKPRLRWTQDLHERFVDAVTQLG

GPSKATPKAIMRTMNVKGLTLFHLKSHLQKYRLGMTGSYLLESPGTENPSPKLPTSDTNE
GYEIKEALRAQMEVQSKLHLQVEAEKHLQIRQDAERRYMAMLERACKMLADQFIGATVID
TDSQKFQGIENKAPRGPLVDHLGFYSLPSTEAAGVNVPEEEVPQTIPPQRADCSTESCLT
SHESSGGLTLEGSQVGGKRRMLGMDSMAAPLIWSEAKMKTQAINLGQGNHPLGISRYGM
>MtrPSR_Medtr2g027860.1
MYHPTNVPDSSLVGSNPLVHGQHIDSGRSAMDPGSGGNSLANNSNLNSKQRLRWTHELHE
RFVDAVAQLGGPDRATPKGVLRVMGVQGLTIYHVKSHLQKYRLAKYLPDCSSDEGKKTDK
KETGDMLSNLDGSSGMQITEALKLQMEVQKRLHEQLEVQRQLQLRIEAQGKYLKKIIEEQ
QRLSGVLSEAPGSGVSAPTPGDMFQQELDNKTEPATPDPEKAAKEHAPAKSLSAESFSSH
HEPMTPDSGCQVGSPADSPNGERSTKKQRVSVEGAYLKSDMVLPHQILESSMPSYQQPNA
IFLTQDHFDPSLGLSTRSGEELDKLGGRNM
>MtrPSR_Medtr2g086450.1
MYHHHHQGKNIHSSSRMSIPSERHMFLQTGNGSSDSGLVLSTDAKPRLKWTPDLHARFIE
AVNQLGGADKATPKTVMKLMGIPGLTLYHLKSHLQKYRLSKNLHGQSSSNVTHKINTHAT
SVSDERLSETNGTHMNKLTLGPQTNNNKDLHISEALQMQIEVQRRLNEQLEVQRHLQLRI
EAQGKYLQSVLEKAQETLGRQNLGIVGLEAAKVQLSELVSKVSSQCLNSTFSEMKELQGF
CPQPNDGSMDSSCLTSSDRSQKEQEIIQNGGFGLRHFNNNNNNHVFMERKEQQATELAGS
VQNLRNNEVLKWCVEEVKKNSNFLTPLGNNNELERNHGNLSMNIGVENHLDIGEFQQRNT
ARLDLNSRGDNNEGATTCKQLDLNRFSWN
>MtrPSR_Medtr4g081710.1
MDQQNQSMRLVLSTDAKPRLKWTHELHQRFTDAINQLGGAEKATPKSLMRVMGIPGLTLY
HLKSHLQKYRLGKSQLVETCSDNKQDYIEIQNSDGQCSREISVGNQNQTTESLKIAEALE
VQMEVQKKLYEQIEVQKHLQFRIEAQGKYLQSVLMKAQEALAGYSSSSSTTGVEHAKAEL
SQLLSIINNACPSSPLSELTETRGFSLNFGERKQNRGTMCSLESSLTSSESSERKEEKQT
INEAENTPNYNSISVELPLMAIESEGRTFRTNANDGGSGRKRSATIDLDGRCVDQPDGKI
CGKKPRKSEFSQMLDLNSKYERDIDSSSLEIDLNCSSSF
>MtrPSR_Medtr5g027440.1
MMESGGRGGIEGYDGMMMTMTRDPKPRLRWTTDLHDRFVDAVTKLGGPDKATPKSVLRLM
GLKGLTLYHLKSHLQKYRLGQHARKQNEEQFKENNRCSYVNFSNHSSGTNTNYGGDNEGG
EIQIGEALRQQIEVQKRLEEQLEVQNKLQMRIEAQGKYLQAVLEKAQTSLPQDGPGNLDA
SKAQLAEFNSALTNFMENMNKDSKENILDMNEFHNKNHAQAFNYQEVIGTEEHKELKPQV
EGGAVQLDLNIKGGNEHLVSADGAEMESNMVSYRVYHF
>MtrPSR_Medtr5g041350.1
MQNQSMHFVLSTDAKPRLKWTPELHQRFIDAINQLGGADKATPKSIMRVMEIPGLTLYHL
KSHLQKYRLGKSQQLETCSDNKKQVYTETMSWDEQCSREIGQGDHNQITENMEISHALEM
QMEVERKLNEQIEVQKHLQLRIDAQGKYLQSVLMKAQEALSGYNSSPIGIKLTKDELSQL
VTMINNACPSSPISDLTESRGLSLNYEERKHENGTSLCSLESSLTSSESSERKEEKHSLE
DIRDFKNSSAISLELPLMAMHSEEKSLSCPMSVSVLRSDDSSNEANGRKRNEETKFDGSY
VENDSLRKRCGNKLKKAKLSEKFDLNRQCQNDMESTSSKMLLDLNCSLNFCEP
>MtrPSR_Medtr6g032990.1
MYYHHHHQQQQARNMHALRMHMQGGGNGSSDSGLVLSTDAKPRLKWTPDLHERFIEAVN
QLGGADKATPKTVLKLMGIPGLTLYHLKSHLQKYRISRSMNGQTNTGSSKIAPTSEVVTS
RMSESSGIHMKDLNIGLQTNKNSDINEALNMQIEVQRRLHEQLEVQRHLQLRIEAQGKYL
QSVLEKAKETLGRQNLGAMGLDAAKVQLSELASRVSTENLDSKFSELKEMNVLWAQQTQE
GETIDYSMGSFLTNSEDSQRDQEIHNKSMNFRAYNGTLCEEVKENMMFLSSSNDKVLKGS
DEVPERTSSLLSMNIGVHEEENFWRRNISKEDLEGEEWKRRKSIDTSGVQLKLNSDKISQ
DYRLANFDMKLDLNSHDDNGASSHSQKFDLNGFSWNC
>MtrPSR_Medtr6g444980.1
MERMFPPKKPSTMNSHDRPMCVQGDSGLVLTTDPKPRLRWTVELHERFVDAVTQLGGPDK
ATPKTIMRVMGVKGLTLYHLKSHLQKFRLGKQPHKEFNDHSIKDGMRASALELQRNTASS
SAMIGRNMNEMQIEVQRRLHEQLEVQKHLQLRIEAQGKYMQRPHGMAEMGLLKEFGSPLS
FSSFQDLDLFGGGGGGGDQLDLQQNMDKQTLDHHGFMQINENLCLGKKRPNPNTNPYSGN
GKNPLMWSDDLRLQDLGTASSCLDDPFKGDQIQIAPPSLDRGSDIETIDIYDTKPLLQGE
ILGEKKFDASMNKLERPSPRRGPSLHAERMSPMISTGTMAQGRGSPFG

>MtrPSR_Medtr7g068600.1
MGSTRSDGSATHKERLRWTQQLHDLFVEAVNRLGGADRATPKGILKGMKAMGVSELKILH
VKSHLQKYRISKLIPESTTRGKIEKRSISDILPNFCSISALQLKEVLQMQAEVQNRMSDK
VEVPKSLKLKIEAQGKYLARIGQSNQIRTITRKACKPFVGKATPLPSLSEESESFKTQSD
EEHRTAKKKKVTNENVFPTGFELGSSTTSEFSNQTWNLSWSQLAEATYQSPLVPSFLL
>MtrPSR_Medtr7g088070.1
MYHAKKFPEETMMLHKSQGGGGGGGEQFANGGGLNGSAVKNVQPAGGGGKQRLRWTSDLH
DRFVDAITQLGGPDRATPKGVLRVMGVPGLTIYHVKSHLQKYRLAKYLPESPGDGKDSKD
EKRNSGDSISGADSSPGLQINDALRMQMEVQKRLHEQLEVQKQLQMRIEAQGKYLQKIIE
EQQKLGSTLAASETLPLSHDKQNQPLSEPSGSSDALADTFSPHKKQRIDEGSKDGTAPQV
TIKTAQKNDCNVGPLDPNLYEDDAGYGFDLETENDDGNE
>MtrPSR_Medtr7g093030.1
MSQIWNIGACNQQQLSIVEEGESTKSCNTNTIMSRFESPASAFYATENCMGFAEFDHQVD
NNQSLSSQSYKVNDLEFPLCQSLRENNHLLDASNQHDPNFELSNTLQALVKSQLNGNQHL
RFPENLNKFSCGNFPFEQQKLFFDGLASVSNSSSFCNKGNHDYMVARGSYHLSVEQLNFS
SQHEKLSPTISAGSLSTSLGSTSSSGNVVSSKTRIRWTKDLHEKFVECVNRLGGAEKATP
KAILKMMDSEGLTIFHVKSHLQKYRTAKFMPESAQGKSDKRIHIDDVQHVGVKTGFQIKE
ALQLQLDAQRRLHEQLEIQRTLQLRLEEQGRQLKKMFDQQQKTCSNLFNTPNTINDDTKN
IGKDVEVSISERAENSLLQSKSSERII
>MtrPSR_Medtr7g098250.1
MEARPAFSIERSSSQQLNNIGMSGALPSSLSVHPTPLEETYPRFSDSQPTYVEKDLKTKT
FNHSSHISSSGAVGHMFSSSPGYSTDLHHSSLSPHEKHSRSAHFISQSLSNMASVPLPYS
SNNGPVPSTTSTHYSNGNSASWHADPLPSFLDFSANASIDNNQVESGACNIMATEEFSKR
NDWQEWADQLISDDDTLTSNWNDLLADNIQDLEPKAVESISKSSSQFPAGHQSQDHQQLP
ALSGENHVGVAPSSSANSATTKPRMRWTPELHEAFVEAVNQLGGSERATPKGVLKLMKVE
GLTIYHVKSHLQKYRTARYRPESSEGAGEKKLSPIEDISSLDLKTGIEITEALRLQMEVQ
KRLHEQLEIQRNLQLRIEEQGRYLQMMFEKQCKSGVEPFKASSSAIENPSGVSSDTMKDS
PTKNELEASKMDHCKSGPDQANGSTTVEESSLEAVEKLDTSKSQQASKDLEQNENEDSPQ
APKRQRTDE
>MtrPSR_Medtr7g115530.1
MPSSSSPVLPSLMESNYMKPPDSFNCSPVRSLTANPASLQATSSYPIRSVRPVFSTNAEC
PGGVPFPPVSQHDRQYQDSPFTSQALGDNVSSEIHSATFTSHLQENDDISWGPDPLPDIL
GFPDIFSVQHDQVENSACYMNEDNVKKTDFGEWVEQLMTSDDSVNPNWSQLLGDDNVVEP
TQKAMHVSQKQHTSSGEVNNLCNPASASASASTASQTKPRMRWSPELHEAFVEAVNQLGG
SEKATPKGVLNLMNVEGLTIYHVKSHLQKYRTARYKPESPEETSEKKMSSIEEMKSLDLK
TSKGITEALRLQMELQKRLHEQLEIQRKLQIQIENQGKHLEMMFEKQKQIGDNKGPSPSN
APSAAVLDTTLPSSPVDNLKTSKDECDKSGCNADIPKDIPGESSHDVSRKQMADEAEVTN
EHERVDDQFSDVPPTKRAKIQ
>Potri.001G133400.1
MERTTFGGGGNYPYENGMVMMTRDPKPRLRWTADLHDRFVDAVTKLGGPDKATPKSVLRL
MGLKGLTLYHLKSHLQKYRLGHQARRQNISEQSRENRGASYVNFSHGSSATSTSSPRMDK
EQGEIPVAEALDSQIEVQKTLQEQLEVQQKLQMRIEAQGKYLQSILEKAQKSLSQNLNDD
GNGNLEATRAQLTGFNLAISSLIENLNAEDRKPCITDLKGVNIRTNGSAIHIDREGQTQE
TKDVKHHLQGDSIHFDLNTKGTYDFVAANGSELELKMLSYRR
>Potri.001G314800.1
MYSAIHSLPLDGHGDFQAALDGTNLPGDACLVLTTDPKPRLRWTAELHERFVDAVAQLGG
PDKATPKTIMRTMGVKGLTLYHLKSHLQKYRLGKQSCKESTDNSKDVGIAASVAESQDTG
SSTSASSRMIAQDLNDGYQVTEALRVQMEVQRRLHEQLEVQRRLQLRIEAQGKYLQSILE
KACKALNDQAVATAGLEAAREELSELAIKVSNERAGIAPLDTMKMPSISELAAALENKHA
SNVPARVGDCSVESCLTSTGSPVSPMGVGAQVASTKKRSRPVFGNGDSLPFDGNIQQEVE
WTMNNIV
>Potri.003G100100.1
MEKTTFGGGGGSYPYENGVVMMTRDPRPRLRWTADLHDRFVDAVTKLGGPDKATPKSVLR
LMGLKGLTLYHLKSHLQKYRLGQQARRQNNTEQSKESRGASYVNFSKGSSGTSTSSPRID

EEQGEISVAEALNCQIEVQKTLQEKLEVQKKLQMRIEAQGKYLQAILEKAQKSLSQNLND
DSNGKLKATRAHLTGFNSAVYSLMENLNAEDRKPSITDLKGINMKENGPAMHIQREGQTQ
ETKDVKHHLQGDSIHFDLNTKGNYDFVSANGSELELKMLSYRR
>Potri.002G257800.1
MTFPVSHYCEPIRRVLSEAMEGRPAFSIQRAGAKQLGNLGVSGTLSSSLPVVPTPLEETY
SKLPGCQQVSMERELMTRPLVHASHLPSNNGVVGHLFSSSASFSTDLQYSSVTPREKHSR
NTPFISQSSANAGALLMSQSSPSALLQSTTTSHYVNENSASWCPESPPGFLDFPTNTTVQ
NNQIESNSCAGVMASEEFGKRNDWQEWADQLITDDDALTTNWNELLADTSIVDMEPKMAY
QVSKPSSNTPVQHSQGHLQLPSLSAEIRPVLTPTSSANSAPTKPRMRWTPELHEAFVEAV
NNLGGSERATPKGVLKLMKVDSLTIYHVKSHLQKYRTARYRPESSEGSSEKRLTSIDEIS
SLDLKTGIEITEALRLQMEVQKRLHEQLEIQRNLQLRIEEQGRHLQMMFEKQCKSGIDVD
KLKAASSALENPSTLSSDAIQDSPAKNDLETAQVDCGKTPTDTIYANPALEGGSQDLNRK
HKVSPTETPENSEPYNTDSSLQPAKRPRTDA
>Potri.006G000800.1
MYQLESVPSSSSVHKNSLVNDQYLDCDDMTMDPINGGNNLNNNPNLASKQRLRWTHELHE
RFVDAVAQLGGPDRATPKGVLRVMGVQGLTIYHVKSHLQKYRLAKYLPDSSSDGKKADKK
ETGDMISNLDGSSGMQITEALKLQMEVQKRLHEQLEACFPCTRHPINCAIMCGDFYAHVS
LVQRQLQLRIEAQGKYLKKIIEEQQRLSGVLEDVPGSGVAAPVSGDNCPESDKTDPATPA
PTSESPLQDKAAKERAPAKSLSIDESFSSQPEPLTPDSRCNAGSPAESPRGERSMKKQRV
SIGVTYGKQEMVLTHQILESSLNSYPRPHSAFLGREQFDPSSGLSMGIEDQMEKVSGSDV

>Potri.006G101000.1
MGSSRSDVSNKERLRWTQELHDRFEEAVNQLGGPDRATPKGILRAMGISGLTIYHVKSHL
QKYRISKFIPETNRGKYERRNISEMLPNFSATSGAQLNEALLMQMEVQKRLSDQLEVQKS
LKIKIEAQGRFLERIVEENRNRSASINPIPKHSKSFSPVSQPSLCDESESNAREFETDSE
GEKAEIQPEEYLQALKRLRTENHALPSRYQLQPLNPDPYNQNMVLQRDAKFSYPSHDANF
PWNILATCPSPLVPSFF
>Potri.006G191000.1
MNTRNIDCEEGVQQNHGVMIGDFVNLSSQYFGNQQIRNMAPRLQPAVMEAGCQQQNISPE
RSSSSILSRFESPASSFYATERCMRFPQYDCQVGSSFCSQYSKSYDSHQSSDPNYSINLG
EQADHNFGLNSTLESVVKPHYSYYNSFDKSDKGLSSSSGNKLPSQQHNKFLDIHGTVSLG
NNFSVPFQGNQDRQVGCNPYSSPFAGLSFNSLEGKQSPRFSLGGGPTSSGKDLSSKTRIR
WTQDLHEKFVECVNRLGGAEKATPKAILNLMDSDGLTIFHVKSHLQKYRIAKYMPEPSEG
KAEKRNSINDVSQLDIKTGFQIREALQLQLDVQRRLHEQLEIQRNLQLRIEEQGKQLKMM
FDQQQKTTNSLLNKQNLDITSPDEPAFSLEDIDVSILEGSDNNTQFPSKIS
>Potri.007G003200.1
MNQHAVVSVTKSETSKGVTQPFCTTLFPIQNSSSSKSDCQTSLTGESSSPRPSPLIRTES
LGSPSKMQLSTAQHQMCCLKFGPDSPLSPTSHVQSSKSTFQRSSVFCTSLYLSSSSISET
NRQLGNLPFLPHPPTYSHSVSATDSTKSPLLFSEDLSNQCDEEHSDAFMKDFLNLSGNAS
EGSFHGMNYTGGNLELTEQLELQFLSDELEIAITDHGENPGLDEIYGTHETSSKPATGFA
CNQDSPSVDALSSHPSPGSSTAHKPRMRWTPELHERFVEAVNKLDGAEKATPKGVLKLMN
VKGLTIYHVKSHLQKYRLAKYLPEKKEEKKASCSEEKKVASINIDGDVKKKGTIQITEAL
RMQMEVQKQLHEQLEVQRTLQLRIEEHARYLQKIIEQQNAGSALLSPKSLSASTNPPKDS
ELPPPSPSAVAESKTDLSSPLPSSKHKAADSDNFEKQTSEKRIRLEEKSESASEDAVVED
PPV
>Potri.008G081800.1
MFHTKKPSTMNSHDRPMCVQDSGLVLTTDPKPRLRWTVELHERFVDAVAQLGGPDKATPK
TIMRVMGVKGLTLYHLKSHLQKFRLGKQLHKEFNDHSIKDASALDLQRSAASSSGMISRS
MNDNSHMIYAIRMQMEVQRRLHEQLEVQRHLQLRTEAQGKYIQSLLEKACQTLAGDQDLA
SGSYKGIGNQGVPDMGAMKDFGPLNFPPFQDLNIYGSGQLDLLHNMDRPSLDGFMSNNHD
DICLGKKRTNPYAGSGKSPLIWSDDLRLQDLGSGLSCLGPQDDPLKGDQIQIAPPLMDSG
TDLDSLSGLYGTKPVHQGDALDEKKLEASAKTERPSPRRAPLAADRMSPMINTGVMPQGR
NSPFG
>Potri.008G087600.1

MGLQHQSMNLVLSTDAKPRLKWTQELHQRFVEAVNQLGGADKATPKSLMRVMGIPGLTLY
HLKSHLQKYRLGKSQQSLFSIESEQEDDKEIQSSDDHFKESAVTRSSRGICSDGNHHPIN
ESFQIAQALQMQMEVQRKLHEQIEVQRHLQLRIEAQGKYLQTVLKKAQETLAGYNSSSMG
IELAKAELCRLVSMVNSGCPSSSISELTETGGSILKDIERTQMRNTVCSMESSLTSSESS
GRKEDMQKENEIHDTNKSNTAFVELPLMDIHPQENLLDNDSSNQGKKRSGSIISDGVSVE
QPLARRLKNGDQLRLGTFDLNS
>Potri.010G167900.1
MGLRHQNMNLVLSTDGKPRLKWTQELHQRFVEAVNQLGGADRATPKSLMRVMEIPGLTLY
HLKSHLQKYRLGKSQQSLISIENNQEVLFVADAKEIQSSDDHFQESAFIQSSGGICSDGN
QHPINGSFQIAQALQMQMEVKRKLHEQIEVQRHLQLRIEAQGKYLQSVLKKAQETLAGYN
SYSMGVELAFNIYDL
>Potri.010G174100.1
MFHTKKPSTMNSHDRPMCVQGDSGLVLTTDPKPRLRWTVELHERFVDAVTQLGGPDKATP
KTIMRVMGVKGLTLYHLKSHLQKFRLGKQPHKDFNDHSIKDASALDLQRSAASSSGMMSR
SMNEMQMEVQRRLHEQLEVQRHLQLRTEAQGKYIQSLLEKACQTLAGDQNLASGSYKGMG
NQGIPGMGAMKEFGTLNFPAFQDLNIYGGDQLDLQHNMDRPSLDGFMPNNDNICLGKKRP
SPYDGSGKSPLIWPDDLRLQDLGSGPACLEPQDDPFKGDQIQMAPPSMDRGTDLDSISDM
YEIKPALQGDALDEKKFEASAKLKRPSPRRSPLAAERMSPMINTGAMPQGRNSPFG
>Potri.011G023600.1
MPSALMQTHDREPPCFQRESNHLLITTDPKPRLRWTLELHERFVDAVTLLGGPDKATPKA
IMRIMGVKGLTLYHLKSHLQKFRLGKQPQNYLNEQAIRDATGHLKNLQDAATARIFGDGL
NKNIHRNEVLGTQIQAQRTLDEQLKVKHHLQKRIDAQRKYMQTILENAYRTVSAENRLFD
DQRVVSEMGNMKEIVSASNFPPIQDLQTYGDHSHDGFLPTDDSMSSCTIPMISYDNMQLQ
HITLAPCLASEEELHQMSYN
>Potri.013G048000.1
MMRNSASQQASSLSPGNRSVGPLFSSSSRFSNDMHVSSVSPQGRQSHNSPFISQSLRDRG
NFTPTHDSHSEVQSTEFIAYSDENKDLSWPVDPLQDLLDFAGNVHVQNGQVESSAGVFAS
EDHAKRTDWQEWADQLISVDDELEPNWSEILNDVNKTDSRQKELKPSPNISVKQPPIHQH
QTAHSGEVCAVANPLSAAPTTKPRMRWTPELHEAFVEAVNQLGGSERATPKGVLKHMNVE
GLTIYHVKSHLQKYRSARYKPESSDEKKTSPIEEMKSLDLKTSMGITEALRLQMEVQKRL
HEQLEIQRNLQLRIEEQGRHLQEMFEKQRKIEDDKSKAPSSSQDDPSPLQAKLEQSSANY
KLEASELDLVKTSNESALLEESSQSISRKQKAPEERNDQVLDQIDEESSPAPIKRPRRDE
TAELSTGAASN
>Potri.013G060200.1
MYHHHQHQGKSIHSSSRMAIPPERHLFLQGGNGPGDSGLVLSTDAKPRLKWTPDLHERFI
EAVNQLGGADKATPKTVMKLMGIPGLTLYHLKSHLQKYRLSKNLHGQANIGSSKIGTVAV
VGDRMPEANATHININNLSIGSQPNKILKSRSLHFSEALQMQIEVQRRLHEQLEVQRHLQ
LRIEAQGKYLQAVLEKAQETLGRQNLGTVGLEAAKVQLSELVSKVSTQCLNSTFSELNDL
QGLCPQQTPPTQPNDCSMDSCLTSCEGSQKEQEIHNIGMGLRPCNSNALLEPKEIAEEHA
LQQTELKWGEYLRDNKMFLTSIGHETERRTFSAERSCSDLSIGVGLQGEKGNINSSFAEG
RFKGMSEDDSFQDQTNKRAESVKFEDEKMSPGYRLSYFTTKLDLNSHDEIDAASSCKQLD
LNGFSWN
>Potri.014G000700.1
MQVSTVQHQKYHPKSGPDSPVSLAYHVQLSKSTFQRSSVFCTSLYLSSSSISETNRQLGN
FPFLPHPPTYSQSVSATDSTKSPQLVSEDLSSPFDEERSDGFMIDFLNLSGDASEGGFHG
MNCTSDNLELTEQLELQFLSDELDIAITDHGENPRLDEIYGTPETSSKPVTGFACYQNFP
SIAPPVDALSSQPSLGSSTAHKPRMRWTTELHERFLDAVNKLDGAEKATPKGVLKLMNVE
GLTIYHVKSHLQKYRLAKYFPEKKEEKKASCSEEKKAVSIIIDDDGKKKGTIQITEALRM
QMEVQKQLHEQLEVQRTLQLRIEEHARYLQKIIEEQQKAGSALLSPKSLSSVTDPPKDSE
LPPPSPSAGAESKTDSSSALPSSKHKATDSENFEKQASEKRIRLEEKLESESEEAEVEDP
PAQ
>Potri.016G001100.1
MDLDCEAMTMDPINGGNSLNNNPNLASKQRLRWTHELHERFVDAVAQLGGPDRATPKGVL
RVMGVQGLTIYHVKSHLQKYRLAKYLPDSSSDGKKVDKKETGDVLSNSDGSSGMQITEAL

KLQMEVQKRLHEQLEVQRQLQLRIEAQGKYLKKIIEEQQRLSGVLEDVPGSGVSAPVSGD
NCPVSDNKTDPATPAPTSESPLQDKVAKECAPTKSLSIDESFSSQHEPLTPDSRCNTGSP
AESPRGERSLKKQMVSMGVAFGKPEMVLTHQILESSLNSYPQPHSAFLTREQFDPSSGLS
MGNEDQSEVLGSDL
>Potri.016G001200.1
MLESRGVGIGGMIMEEMHQNQSGIGIGLGMQGGAQVVLTSDPKPRLRWTADLHQRFVDAV
SQLGGPNKATPKAILRTMNVKGLTLFHLKSHLQKYRLGKQSGKDMSDTFKDGLSGSYLLE
NPCTGNSSLNMTASDVNEGYEVKEALRAQMEVQSKLHLQVEAEKHLHIRLDAERRYLAML
ERACKMLADQFIGAAVIDTDSQKGLGTRTTRIASLDPLGFYSLQTSEVAEVHGPEDVLPG
LHHQGADCSTESCLTSNESPGGLNLEGSPAGGKKGMLSLESATSLIWGETRMGNAEVNAT
QVNSYGASLYGIWN
>Potri.016G047900.1
MASRLQPAAVEAGCQNQNISLERSPDSIMSRFESPASAFYATERYMRFPQYDCQVGNYFC
SQYSNSYDSHVSSHQSSGADYINSGEQADHNFGLKSTLESVVKPQFSCHKSFDKSDKGLS
SSSGNKLPSEHHNKFLDNPGVSLENHFLVPFQGNQNRQVDYNPYNSPFSELGRFNSREEK
RSPRFSLGGFPISSGKDLSTTLSSKTRIRWTQDLHKKFVECVNRLGGAEKATPKAILKLM
DSDGLTIFHVKSHLQKYRSARYMPDSSEGKAEKRTSIDDVSQLDVKTGFQIREALEVQLD
VQRRLHEQLEIQKILQLRIEEQGKQLKMMFDQQQKKTNSLLNNQNLNITSPDESTFSLED
IDVSIVEGSNNNTHFPSKIS
>Potri.016G117000.1
MGRSRSDRSNKDRLRWTQELHDRFEEAVNQLGGPDRATPKGILRAMGIPGLTIYHVKSHL
QKYRISKFIPETNRGKFERRNISEMLPNFSATSGAQLNEALLMQMEVHRRLSDQLVVQKS
LKLKIEAQGRFLERIVEENQNGNPKHTKSFSPVSLPSLCDSESNAKEFETDSEGEKVEIQ
SEEDFQALKRLRTEHHVLPSRYQLEPLNPDPYNQNMVLQRDAKFSHPSHDVNFPWDILAT
CSSPLEPSFF
>Potri.017G054800.1
MYSAIHSLPLDGHGDFQASLDGINLPGDACLVLTTDPKPRLRWTAELHERFVDAVTQLGG
PDKATPKTIMRTMGVKGLTLYHLKSHLQKYRLGKQSCKESTDNSKDVGIAPSVAESQDTG
SSTSASSRMIAQDLNDGYQVTEALRVQMEVQRRLHEQLEVQHHLQLRIEAQGKYLQSILE
KACKALNDQAVATAGLEAAREELSELAIKVSNECAGIAPLDTMKMPSLSELAAALGNRNA
SNVPARIGDCSVESCLTSTSSPVSPMGVGSQVASTKKRSRPVLGNGDSLPFEGNFRQEVE
WTMSNIVDE
>Potri.018G151400.1
MKFYSTVKSFNRRKGAMQTHPSSLNTESVGLYCLNKNLQYSAVLPDCVQPKPPNSLPQAS
LFLTRTLNASPILNDAVSYGSIGQPRKGTMFSASLQPSPLTIAESDLQLPTSPFLQRPLP
VDHITPPSDSLFLGGQDVITEPENENLMGVLDECSGGSFTGLHCTKESLTSTEKLVLQYL
SKELEIPVDDVSQNSTLNEAQRVSSIPVTELNHNANYRSSAAQMDDSINRLPEAATSQKQ
RIRWTTELHDLFVDAVKSLGGPDVATPKSILGIMNVKGLSIYHVKSHLQKYRLAKKFPET
NHDKSTSTVVENKAASSNSNNDALVIESNRDVQVTEALRTQIEIQKLLHEQLKAQKELQI
RIEQNEKFLRELMEQKAISIYEPSSFAVPASEPKLLPHSPSADVSSPGQAAVNSDCYLFQ
PSNHKDSDAVESEKAKCPKRDRGQKEHPILH
>Potri.019G020900.2
MIRNPASQQASPLSSGNGSVGHLVSSSSRFTNDTSVSAVSPQGKQSHNSPFISQSLRDGG
NFPPTHYSHSEGQSTAFINHSDDNKGLSWPIDPLQEFINFVENVPVQNGQVESTAGVIAS
EDHAKRTDWQEWADQLISVDDELEPNWSEILNDVNMKDSKQKMLSPNNSVQQPLIHQHQT
AHSGEVCAVTNPLLAAPPTKSRMRWTPELHEAFVEAVNQLGGSERATPKGVLKQMNVEGL
TIYHVKSHLQKYRTARYKPESSEGTSEKKLSPVEEMKSLDLKTSMEISEALRLQMEVQKQ
LHEQLEIQRNLQLRIEEQGRYLQEMFEKQKKMEGDRSKAPPPSQNDPSLLQSKLEQSPAN
DKLETSDLDCVKTRFDICNASALLEESSQSINRKQKAPEDRNCQVVDKNEEKTSLAPVKR
PRTDEATALSAEPASN
>Potri.019G032700.2
MYQHHQHQGKNIHSSSRNSIPPERHLFLQVGNGPGDSGLVLSTDAKPRLKWTPDLHERFI
EAVNQLGGADKATPKTVMKLMGIPGLTLYHLKSHLQKYRLSKNLHGQANSGSNKSGTVAV
VGDRMPEVNATHINNLSIGSQTNNEALQVQIEVQRRLHEQLEVQRHLQLRIEAQGKYLQS

VLEKAQETLGRQNLGTVGLEAAKVQLSELVSKVSSKCLNSAFSELKDLQGLCPPLTQPTH
PNDCSMDSCLTSIEGSQKEQEIHNTGMGLRPYNGNALLEPKVIAGEHALQQTELKWGEDQ
RDNKMFLSSMRNDTDRRTFSAERSCSNLSIGVGLQGERGNVSSSFAEARFKGRSEDDSFQ
DKTNRRIDAIKLENEKLSPGYRLSYYATKLDLNSHGEIDAASGCRQLDLNGFSWN

>MesPSR_Manes.01G035100.1
MGFQNLQNQNMNLVLSTDPKPRLKWTPELHHRFVEALRQLGGEDKATPKSLMRVMGVPRL
TLYHLKSHLQYRLAKRHPTELQSEFQSQSQSQSQGSIENKKVLLDADYKEIQISNPALRG
GISEGNQSPLNESFQIAQVLQMQMEVQRKIHEQIEVQRHLQVRIEAQGKYLQSVLKKAQE
TLSGYSSSSLGIELAKAELSRLVSMVNNECRSPSISELTEIEGSSLKGTERKKMRGTVCS
MESSLTSSESSGRKEDKQLTNEFGNTHKSNPASIELPLMDIHPREKPWNSGTSDQVKKRT
CCTISDGICVEQPPVKRSKIGDRLRIFDLNSHCQNEFESGSKTIDLNCKGIEQVNGQL
>MesPSR_Manes.03G077700.1
MRVMGVKGLTLYHLKSHLQKFRLGKQPHKEFNDHSIKDAQALDLQRSGGSSSVMMSRSMN
DNSHMVDATRMQMEVQRRLHEQLEVQRHLQLRMEAHGKYMQNMLEKAYQTLASENMASGI
YKGIGNQGVPDMGGGMKDFSPLNFSLFQDLNIYGGDQLDLQHNMERQSSLDGYMQNNDNI
CLGKKRPSPYSGNGKGHLFWPDDLRFQELGSAPTCLGPQDDHLFKGDHLIQIASQSVDRG
SVLDSISDIYESKPPVFQGDSVSDKHTRLERPSPRRAPFPADRMSPMINTAGAMLREGNS
TFG
>MesPSR_Manes.03G148100.1
MYSSIHSLPLDGSVGHGDFQGSLDGTNLPGDACLVLTTDPKPRLRWTAELHERFVDAVTQ
LGGPDKATPKTIMRTMGVKGLTLYHLKSHLQKYRLGRQSCKESNDNSKDVGIAASVAESQ
DTGSSTSTSSRLIAQDLNDGYQVTEALRVQMEVQRRLHEQLEVQRRLQLRIEAQGKYLQS
ILEKACKALNDQAAASAGLEAAREELSELAIKVSNECQGIVPMDNIKMPSLSELAAALEN
KSTANLPARIGDCSVESCLTSTGSPVSPMGVGSQAAGAGAASIKKRPRSAFGNGDSVPLE
GNIRQEVEWMMSNIG
>MesPSR_Manes.05G101800.1
MGLHNLQNQNMNLVLSTDAKPRLKWTPELHQRFVESVRQLGGADKATPKNLMRSMGIPGL
TLYHLKSHLQKYRLGKSQPSQSQSQSQSQSQSQSQSQSQASIENKKEDYKEIQSSNCDPR
AGIADGNQNPINESFQIAQALQMQMEVQRKLHEQIEVQRHLQLRIEAQGKYLQSVLKKAQ
EALSGYNSSLGIELAKAELSRLVSMVNTGCQSSSISELTEIGGSNLKDTERKLMRGTVCS
VESSLTSSESSGRKEDMKQKNQLGNTNNSNPVIIELPLMDIHPQENPWDNHKSEQVKKRS
CSTISDGICVEQPLVKRSKSGDKLRIFDLNSHYQNDFESGSKTLDLNCKGIEQVNGQV
>MesPSR_Manes.06G035600.1
MYQPKCIPSSSLVHKSSLVHGQYLDCGANSMDPISGGNSLNSNPTLASKQRLRWTHELHE
RFVDAVAQLGGPDRATPKGVLRVMGVQGLTIYHVKSHLQKYRLAKYLPDSSSDGKKADKK
ETGDMLSNVDGSSGMQITEALKLQMEVQKRLHEQLEVQRQLQLRIEAQGKYLKKIIEEQQ
RLSGVLGEVPVQGASAPISGDNCPESDHKTDPATPAPTSESPVQDQVGKECTSTKSLSID
ESFSSRHEPLTPDSRCNVGSQSESPKGERSLKKQRVCVGAAYSKSEMVLTHQILESSLNS
YPQPHSVFMAREQFDPSSGLSIGNEDEMEKVPGNDL
>MesPSR_Manes.06G178000.1
MNRNDVGSATKNECKGITQPYYTTISPIHDFFSIESEGQSSTTTECSSCPSPLIRTESLS
SPKNMQASTVQPVKYSLKSGPDSPPPVFHVQHSKSTFQRCSVFCTSLYLSSSSSSETNRP
LGNLPFLPHPPTHSQSVSAVGSTKSPALFSWDISGFQVEKETSDAFVKDFLNLHGNASEG
SFHGITSASDNLALTEQLELQFLSDELDIAITDHGENPRVDEIYETSESYTAMGLTCNQN
FASVAPSVDDNSSLPSPGPATMHKPRMRWTPELHECFVEAVNKLGGAEIEATPKGVLKLM
NVEGLTIYHVKSHLQKYRIAKYLPERKEEKKVSCSEEKNAAPSCIEVDGRRKGTIQITEA
LRMQMEVQKQLHEQLEVQRVLQLRIEEHARYLQKILEEQQKAGSALLSPKSLSSQDSELV
QPSSLEGAPHARSSESKTDASSLSKKRASDRSDFEQQACVKKMHLEEKPESAVEEVVVEN
PVQ
>MesPSR_Manes.07G000800.1
MRVFFQSDHLPIPLGCSTDGNPGASDIPSSAFCTMKGCMSFAQGDDYDFNSLSGSQATET
LGLQSTVKSESNKNLTIKETMCQDLHPHSSGGDLFLPILQKSANELNQPNQSHINFKNHQ
NCENGYQLFSSQFTKPINYNFKPAAKQPKHPCYGIVSANSKSVTSGPAGTCKTRIRWTKD

LHKRFVECVDILGGPVKATPKLILKLMGVEGLTIFHVKSHLQKYRISRYIPKSTEGKTNR
NLTNTIALDPKAGMQIAETLRMQIDVQKRLQEQLEIQRNLQSRIEEQGRQLRKMLDLQLV
TNKTLTETKNSSDLAFQEDPSNDPIEGLGDFH
>MesPSR_Manes.08G014900.1
MYQTKSIPSSTLVHSCCSLVHGQHLECGASKMEPINGGNSLNNPSLASKQRLRWTHELHE
RFVDAVAQLGGPDRATPKGVLRVMGVQGLTIYHVKSHLQKYRLAKYLPDSSSDGKKADKK
ETGDMLSNLDGSSGMQITEALRLQMEVQKRLHEQLEVQRQLQLRIEAQGKYLKKIIEEQQ
RLSGGLGEVPGAGVSAPVSGDNFPESNSKTDPATPAPTSESPVQNKAAKECAPAKSHSID
ESLSSRHEPLTPDSRCNVGSPAESPKGERSAKKLRACLDTTYAKPEMIFPHQILESSLNS
YPQPHSIFRTTGHFNPSSGLSVGSEDDIEKVSGNDH
>MesPSR_Manes.08G045400.1
MNTSKIDCQDRIHHNHGLIGDFAVHFSQCFGNQQSWNKEIHMQQPVMEAGLQQQQYLRPD
RSSASIMSRFESPASAFYATERFMGFPQYDCQVEAPPLCFSYSKSYDSQQSSRENYAIDS
GEQAEHNLEMRSNLQPIVKSHFSDDQFYKSYKSSCSSSSENKLYLLERNKVLNNGTASVG
NHGSIPFQGDQDHRVGCNPCTSPFAQLGFNSSQGIQSPRFSSAGACVSSGNPVANGAVLS
SKTRIRWTQDLHEKFVVCVNRLGGAEKATPKAILKLMDTDGLTIFHVKSHLQKYRIAKYM
PDSLEGKPERRNSISNVSQIDTKTSGMQITEALQLQLDVQRCLHEQLEIQKNLQLRIEEQ
SRQLRMMLDQQQQRTSNSLLRNQNLDNTTSPDEPELNLDDIEISITEDFNNTQFPSKIS
>MesPSR_Manes.08G140300.1
MSSSLPVLPTPLEDKYQKLPDSFQVASERQLIRNPVPLQASQLGPDSGTTAHLFSSSLRI
PKELNTSSVSPQNSPFIRSSKDRGTLLVAPTYSSHSEAQSTTLISNSEENKDISWSVDPL
YDLLDFPGNVTTQNGQVESNIGVIASEDLSKRTDWQEWADQLISVDSDLEPNWSELLNDA
NATDTEQQVLKSSPEFSVQPQIHLYNGEPCAVANPISTAPSTKPRMRWTPELHEAFVEAV
NKLGGSERATPKGVLKLMNVEGLTIYHVKSHLQKYRTARYKPESSEGTSEKKLSPIEEMK
SLDLKTSMGITEALRLQMEVQKRLHEQLEIQRNLQLRIEEQGRHLQMMFEKQRKLEDGKS
KVSSSSLDDPTLLQSDAVLPSGNDKSEISEPDHAKTVSDRNDCGVALEKNSPSVSRKHKA
LENGTGEDLLDESSPASTKRARANETAISSTSCI
>MesPSR_Manes.08G149600.1
MYHHHHQHRGKSVNSSSRMSIPPERYLFLQGGNGPGDSGLVLSTDAKPRLKWTPDLHERF
IEAVNQLGGADKATPKTVMKIMGIPGLTLYHLKSHLQKYRLSKNLHGQAISGSSKIGATT
VESDRMSEANVIHINNLSIGSQTNKSLHIGEALQMQIEVQRRLHEQLEVQRHLQLRIEAQ
GKYLQAVLEKAQETLGKQNLGTMGLEAAKVQLSELVSKVSTQCLNSAFSELKELQSLCPQ
QIQTTPPTDCSMDSCLTSCEGSQKEQEIHNAGMGLRPYNGNTLLEAKEMVQDRLLRPTEL
KWDVHLKDNKMFLSPVGNNAEKSSNLLMRVGLQEENGNTSSSFTEGRFKGRSNDNNFADQ
TNKRTDSVKLHNGNISPGYGLPYFATKLDLNSHEEIDAASSCKQLDLNGFSWN
>MesPSR_Manes.09G033400.1
MEGESLCQNLEAKLSSSVIAHHLDQLASVFYAADEDFIDFQREDECEIVAMNSSQVPEPG
DIVSESSLQTILMKPYFCSNEHHRFYEKFRRCLSHDEHKKLLGENPNLIGTQVQVPSDEQ
NNSNLILSSPRDMEVASFTSSSSCASSVLSLPSKKRIKWSQALHKKFVHCVNSLGGAEKA
TPKAILKMMESKELTIFHVKSHLQQARVKSFDQIIFCALQKYRSEKYMSEYKQDKPLLQG
KAESITSDISILCMKNNMKINEALKLQLDVEKHLRQQLEIQRQLQQQIEENARQLKMMMQ
QQQKINKSNL
>MesPSR_Manes.09G033500.1
MNTSKIDCQERVQQNNGLNGDFALHFSQCFGNQQSWNMGIRTQQPVMEAGLQQQNHRPND
KSSTTIMRSFESPASAFYATERYMGFPQYDCQVGVPPLSFPYSKPFDSQQSSRENYAIDP
GEQAESNLDLRRNLQPIVKPHFSVDHYHKSYKGPCSNSFGNKLHLFERNKLSNNGAASMG
HHFSIPFQADQDHRVGGNPCASPFSQLGFSSRQEIPSPRFSSPGACISSGNPEAAGAVLS
SKTRIRWTQDLHEKFVMCVNRLGGAEKATPKAILKLMDTDGLTIFHVKSHLQKYRIAKYM
PDSSEGKSEKRTTINDVPQIDTKTSGLQISEALQLQLDVQRRLHEQLEIQKNLQLRIEEQ
GRKLQRMFDQQQRTSNSLLRNQNLDRKSSPDDPAFSFEDIEVSIIEGSSDTQFPSKIS
>MesPSR_Manes.09G105800.1
MGSSRTDGSGKERLRWTQELHDRFERAVNQLGGPDRATPKGILKAMSIPGLTIYHVKSHL
QKYRISKFIPESNNKGKFERRNISELLPNFSATSGAQLNEALKMQMEVQRRLSDQLEVQK
SLKNKIEAQGRFLERIVQEHRNRTVNIQKHKQSFSPTSLPFLCEYSESNAKDFESDDEGD

RSDQIHYNEDDEFQPLKRFRTENDVLPSRYKLEELNSDPYNNQTSELNLYPWSNMSCSSP
LVPSFF
>MesPSR_Manes.09G142300.1
MYHHHQHQGKSIHSSSRMPIPPERHLLLQGGNGPGDSGLVLSTDAKPRLKWTPDLHERFI
EAVNQLGGADKATPKTVMKLMGIPGLTLYHLKSHLQKYRLSKNLHGQAVSRSSKIGANAV
AVDRMSEANVTHLNNLSIGTQTNKNLHISEALQMQIEVQRRLQEQLEVQRHLQLRIEAQG
KYLQAVLEKAQETLGRQNLGTVGLEAAKVQLSELVSKVSAQCLNSAFSELKELQGLCPQQ
TQTTPPTDCSIDSCLTSCEGSQKEQETHNTGMGLRPYNGNAFLESKEMAEDHMLHPTELK
WVENLRDSKMFLSPMGNNTESRIFSTERSSSNLAMRVGLQGESGKASTSFTEERYKGRND
DDNFPDQTNKRTDSVKLQHDNISTGYRLPYFATKLDLNSHDEIDAASNCKQLDLNGFSWN

>MesPSR_Manes.09G147400.1
MSSSFPVLPTPLEDKYPKLPDSFQIASERELMRDPVNQQSSPLGPNSGTIKHLLSSSSTF
PNELNDSLVSPRGGQSQNSPFIYLSSRNRGTLPVARSHSSHPEELSIALINHSEEHKDMS
WSMDPLHDLLDFPENVTVENGQVESNIGVITSEDLSKRTDWHEWADQLISVDDNLEPNWS
DLLNDSNATDVKQKVLKLSSEIPVHQPQIHQHQHLNNGEPYTVANPTCTAPPTKPRMRWT
PELHETFVEAVNKLGGSERATPKGVLKLMNVEGLTIYHVKSHLQKYRTARYKPEASEGTS
ENKLSSIDEMKSLDLKTSIGITEALRLQMEVQKQLHEQLEIQRNLQLRIEQQGRYLQMMF
EKQRKMEDEKSNPSSSSMDDDPLVPQSNAVQPSDNNKIQVSEPDHAKLVGDRSESRVALE
ESSPSVSRKHKSLANTTDKSLDPKDDESSPASAKRPRADETAL
>MesPSR_Manes.12G035600.1
MEARPALSIQRSGAKQISNHGLSAALSSSLPVLPTSIEETCPKLSNSQQVSLEGGLMTKP
IAHASQLPSNSGVVGHLFSSSAGFLSDLQYSSVSPQEKQSGNAPFISQSSINVTALTLPQ
SSHSGLLQSTKSSQYAKETNASWCPESMPGFLDLPVNNPVQNNQVESNSCSGVIVSEEFS
KRNDWQEWADQLITDDDALTSNWSELLIDTSVVEVEQKMAYQVSKPSSNVLSQQPQVHQQ
LPAPSGEIHPVLTPTSSTNNVPSKPRMRWTPELHEAFVEAVNQLGGSERATPKGVLKLMK
VDGLTIYHVKSHLQKYRTARYRPDSSEGSSEKKFTPREEISSLDLKTGIEITEALRLQME
VQKRLHEQLEIQRNLQLRIEEQGRYLQMMFEKQCKSGSDKLTASSSAMENTSAISSDAIH
DSPAKNEMEASQLDHDKTNCNPLDAKSMVEEGSHDPSEKQKAPEVDGSENPQPDNCESSS
PSAKRQRTDA
>MesPSR_Manes.15G053300.1
MYSAIHSLPLDGSGGHGEFQGGSLDGTNLPGDASLVLTTDPKPRLRWTAELHERFVDAVT
QLGGPDKATPKTIMRTMGVKGLTLYHLKSHLQKYRLGRQSCKESNDNSKDVGIAASVAES
QDTGSSTSTSSRMIAQDINEGYQVTEALRVQMEVQRRLHEQLEVQRRLQLRIEAQGKYLQ
SILEKACKALNDEAAASAVLEAAREELSELAIKVSGKCQEIVPVANVKMPPFSELAAALE
NKSTANLPARIGDCSVESCLTSTGSPVSPMGVGSQAAASIKKRPRSGFGNEEALPLEDNM
REVEWMMSNVG
>MesPSR_Manes.17G009200.1
MERGVLGGNYPYENGVVMSRDPKPRLRWTADLHHRFVDAVTKLGGPDKATPKSVLRLMGL
KGLTLYHLKSHLQKYRLGQQQAKKQTAKEQTKESGSSYVNFSNHSSAVSTSSSAIEIQQG
PGELPVTEPLKSQMEAHKRLEEQLEVQKKLQMRIEAQGKYLQEILEKAQKSLSLDMKCNG
NNLDLALSNLIQSMNEEGRKDNITDLKDIYNKENPWAFYISKQGEREEINDLEHKVEAHP
IHLDLNTKGTYDSLSVNGSTIGTQHAFI
>MesPSR_Manes.18G144000.1
MNTHTVISLNRRENSRVMHTHYSGVSPLYSAVNFDSLQPQCIPPKPVNSITTQLQQKTNA
NPVPNHSPSSTSHSPGLKSTMFCTDLHLSSMKSSETNRPLGISPFLPHPHPSPPIVNLVI
PSIESSLFMGGAMTTESENDNLNSLMDLFDIPSKLPIGRFQGSQPATQGLVFTEQMGLEY
ISKELGIDDDRNDPMLDGIFEMPQVSSSISFNGLNCKSNGDSSLAHQAGGACYNHGRLHE
AAASSKQRIRWTTELHDLFLDAVKALGGPEIATPKKVLGIMNVKGLNIYHVKSHLQKYRL
AKDFPEQKHDKKTENKAASSNNDNDARIKSDMQVTEALRMQIEVQKLLHQQLKTQKELQL
RVEKNGELLRKLMEEQPSLLNSLSKSDQQLSPSSSDIVSSTRPPVDGSSSQPSTHEPSKS
ESTCHKRLRGESSSRAASIDRSDV
>MesPSR_Manes.S107600.1
MERSLIKKPVVHASQSPSNSGVVGHLFSSSAGFSSDLQYSSVSPQEKHSRSIPFISQSSR

NVTDLTLPQSSCPGLLQFTTSSQYAKETNASWCPESLPDFLDYPVNNPVQNNQIEGNSCG
AIVSEEFSKRSDWQEWADQLITDDNALTSNWSELLIDTSVAEMKPKMAYQVSKSSSVPSH
LPQVHQQLLAPSGEIQPVLTPTSSANSAPSKPRMRWTPELHEAFVEAINQLGGSERATPK
GVLKLMKVDGLTIYHVKSHLQKYRTARYRPDSSEGSSEKKLTPMDEISPLDLKTGIEITE
ALRLQMEVQKRLHEQLEIQRNLQLRIEEQGRYLQMMFEKQCKSGADKLKVSLSTMENPTA
LSTDAICESPAENETEGSQSDHGKTNIDAVDAKPLLEEGSHDTSEKRKTPQVEDSENPEP
DSSKRLRADA
>RcoPSR_28582.m000316
MNTSKIDFQGRIQQNHGMIGDLALHSFQSFGNQQTWNMGIRAQSPVMESAHLQQQNLRPD
KSSSSIMRSFESPASAFYATERYMGFPQYDCQVNAVLSCPYSKSYDSQIPSQQSSGEIYV
IDAVNQQPDHNLELRNNLQSITKSHLSDDHYYKSYKGVCSNSLGNKLHQLEQNKLSRNGA
VSVGNQFSIPFYGDQDHNNNHNRFGSNPFVQLGVSSRQEMQSPRFSSGVVSVSSASSGNSM
ATGAVLSSKTRIRWTQDLHEKFVECVNRLGGADKATPKAILKLMDSDGLTIFHVKSHLQK
YRIAKYMPDSSEGKAEKRTSINDVSQMDPKTGLQITEALQLQLDVQRRLHEQLEIQKNLQ
LRIEEQGRQLKRMFDQQQRTNNNLFRNQNLDSISPDEQAFSLEDIEISFAEGSSNNSHFP
SKIS
>RcoPSR_28582.m000317
MGGSQWQFLGPNMLPSISADDLDQLASAVYATDCYLEFRKHDDEYGIASPSSQVPEPNST
IVQMPPNKSSGDGISIDSNLQTILKPYFCSNQLHKSCEIFQKGLSCIESRGNDFLSHDNI
RFLAANATLIGTHIQFPSIGQNNPELISSSQQEKSSLRGLEVASFTCSSSSASSGPNVSS
KKRMRWNQELHEKFINCVNNLGGAEKATPRTILKMMESKGLTIFQVKSHLQKYRAEKYMS
ERKQGKTETASSDIPQLCMKNTMQIKETLKLQLNFQKHLNEQLEIQRHVQQKIEENGKQL
KMMLQEQQKINKRHIT
>RcoPSR_28582.m000318
MEGGSKCQKLGPHMLSMSDLDQLASAVYATEKHDGEYGIASHLSSQAPEANTLPPIESSG
VEIVTESNLQTILKLLFCSNQHHKSCEMSQKSLFSIESRGNDHFLSHDNIKLLTANPNLI
GTHLQFPSIPQNNPELIPPSEREKSSLRDLEVASFTCSSSSAYSRPRHSRKNRMRWSREL
HEKFINCVDNLGGAEKATPKTILKMMESKGLTIFHVKSHLQKYRAEKYMSERKQGETERT
SSDVPLLYMENIMQIKETLQLQLDFQKQLNEQLEGPDPHMFLAQAFRVHGEIEEGAVIYD
KATGKSRGYGFITYKHMESTQIALRAPSKLIDGRLAVCNLACEGLTGASTTPDLAQRKLY
IGGLSPEITSEMLLNFFGRHGEIEEGSVAYDKDTNESRGFGFVTYKTVEAAKKAIDDPQK
VLGGRSIIVKLADTHKGKTVQTQMPASVAPVPLLMTAGYTQPGKAHPGPAPVGYSYPQTM
PSYPASSYPSPPTAPAPYPLQSQYQYPPVAMKKEPFGPPAPMGMGGYPYYLPKQ
>RcoPSR_29637.m000753
MEARPTLSIQRSGVKQLSNLGVSGALSSSYPVLPTSREETYPKLSDAQQVAMEKGLMTRP
LVHTNHLPSNNGVVGHLFSSSAGFSTDLQYSSVSPQEKHSRNTPFISQSPTNVAALTLPQ
SPHSGLLQCTASSQYDKENSASWCPESLPGFLDFPVNSHVQNNQIESNSCTGAITSEEFS
KRNDWQEWADQLITDDDALTSNWNDLLVDNSAPEMEPKMAYQVSKPPSDISAHQPQVHQQ
LPAPSIDIRPVLTPTSSVNTAPSKPRMRWTPELHEAFVDAVNQLGGSERATPKGVLKLMK
VEGLTIYHVKSHLQKYRTARYRPDSLEGSSEQKLTPLEEISSLDLKTGIEITEALRLQME
VQKRLHEQLEIQRNLQLRIEEQGRYLQMMFEKQCKSGTDVLKASSSAVENPSSALSSDAV
HDSSGKNEMEASKVDHGNAITNTDDTKSKLEERSQDPNEKKKALQIEASGNPEPDNGESN
SQSAKRPRLDV
>RcoPSR_29680.m001711
MSSPFPVLPTPLEGQYPKLPDSFQVSSERELMRNPILQQTSPLCSNSGTVGHLFSSSMRS
SIEAQASSLAPQGGQSQNSPFISQSLRDKGSLPVLTTHSSNSEVQSTALINHSEENKDMS
WTIDPLHDLLDFPENVAVQNGQVESTIGVITSEDFSKRTDWQEWADQLISVDDDLEPNWS
ELLNDANNADRKQKVVKSSSQISVQPTVHQPQPVHNGEPYSAANPMSAIPAAKHRMRWTP
ELHEAFVEAVNKLGGSERATPKGVLKLMNVEGLTIYHVKSHLQKYRTARYKPESAEGTSE
KKLSPIDEMKSLDLKASMGITEALRLQMEVQKRLHEQLEIQRNLQLRIEEQGRHLQMMFE
QQRKMEDDRSKASSSSLDDPSLPQSNIVQSPGNNKLEVSELDHARTEISSGGGALEGSSQ
NGSRKQKAPENRTGEDLDPEDDESGPASAKRPRADEIAA
>RcoPSR_29805.m001543
MFQRTIQPHQQQGGIMDQQQEEMQIQEHNQGIREPCLVLTSDPKPRLRWTADLHNRFVDA

ISQLGGPNKATPKAIMRTMNVKGLTLFHLKSHLQKYRLGKQSGKDMGEASKDGLSGSYLL
ESPGAGSSSPNIVTSDMNEGYEVKEALRVQMEVQSKLYLQVEAEKHLQIRQDAEKRYLAM
LERACKMLADQFLGGTVIDSDIQKDSGSKKKRSASVDPLGFHSLQTEAEAEARGLEEVPS
SLHQQGADCSTESCLTSNESPGGLNLEGSPAGGKKQMPSLDSTLNWEEAKMRTSEVNMVQ
VNSHGMSGYGIWS
>RcoPSR_29805.m001544
MYQLKSVPSSSLVHKSSLVHGQHLDCGASRMDAINGENSLNNNPSLASKQRLRWTHELHE
RFVDAVAQLGGPDRATPKGVLRVMGVQGLTIYHVKSHLQKYRLAKYLPDSSSDGKKADKK
ETGDMLSNLDGSSGMQITEALKLQMEVQKRLHEQLEVQRQLQLRIEAQGKYLKKIIEEQQ
RLSGVLGEVPGAVAAAPVSGDNCPESDNKTDPATPAPTSESPIQDKAAKERAPAKSLSID
ESFSSRHEPLTPDSRCNVGSPAESPKGERSMKKQRVCMGTSYGKSEMVLTHQILESSLNS
YPQPHSLFLSREQFDPSSGLSTGNDDHIEKVPGSDP
>RcoPSR_29813.m001501
MGLQDLQNQNMNLVLSTDAKPRLKWTPELHQRFVEAVNQLGGADKATPKSLMRVMGIPGL
TLYHLKSHLQKYRLGKSQLLHSESPSQSQSQASIENKQEDYKEIQSTNCELKAGIAEEIQ
NPTNESFQIAQALQMQMEVQRKLHEQIEVQRHLQLRIEAQGKYLRSVLKKAQETLSGYNP
SSAMGIEIAKAELSRLVSMVNTGCSSSSISELTEIGNSILNDTTDKNQMIRRGTVCSAES
SLTSSESSGRKEDDMQQKNEIGATNKSITTSLELPLMDAHRQDKSWNNNNNNHNNNTYQP
TNQAKKRSSTTISSDGICVQQPIAKRSRNGEQLRIFDLNSQYQIDFEPGSTRAIDLNCQG
F
>RcoPSR_29827.m002675
MERAAAAFGGTYGYENGVVMTRDPKPRLRWTPDLHHRFVDAVTKLGGPDKATPKSVLRLM
GLKGLTLYHLKSHLQKYRLGQQQARKQNTKEQYKENSGASYVNFSNHSSSSGLHATSSSN
HNQQGEIPIAEALKSQIEVHTRFKEQLEVQKKLQVRIEAQGKYLQDLLEKAQKSFSSLDM
KGSCNLDLALIKNTNAEERKENLKDIFAEPNYSVFHVSREAEMQKINDFEHKIEGSIHLD
LNTKGSYDFAAVNGSQLELR
>RcoPSR_29904.m002952
MYSAIHSLPLDGHGDFQGSLDGTNLPGDACLVLTTDPKPRLRWTAELHERFVDAVTQLGG
PDKATPKTIMRTMGVKGLTLYHLKSHLQKYRLGRQSCKESNENSKDASVAESQDTGSSTS
TSSRMIAQDVNDGYQVTEALRVQMEVQRRLHEQLEVQRRLQLRIEAQGKYLQSILEKACK
ALNDQAAVSAGLEAAREELSELAIKVSNECQGIVPADNMKMPSLSELAVALESKSTSNLP
ARIGDCSVESCLTSTGSPVSPMGVGSHTASIKKRPRPIFGNGDSLPLEGSMRQEVEWMMG
NIG
>RcoPSR_29950.m001149
MYHHHQHQGKSVHSSSRMSIPPERHLFLQGGNGPGDSGLVLSTDAKPRLKWTSDLHEHFI
EAVNQLGGADKATPKTVMKLMGIPGLTLYHLKSHLQKYRLSKNLHGQANSGSNKIGTGAV
VGDRISETNVTHINNLSMGTQTNKGLHIGEALQMQIEVQRRLHEQLEVQRHLQLRIEAQG
KYLQSVLEKAQETLGRQNLGSIGLEAAKVQLSELVSKVSTQCLNSAFSELKELQGLCHQQ
TQTAPPTDCSMDSCLTSCEGSQKEQEIHNTGMGLRPYNGNALLESKDITEGHVLHQTELK
WSEDLKDNKMFLSPLGNNAARRNFAAERSTSDLSMTVGLQGENGNASSFSEGRYKDRNDG
DSFPDQTNKSLDSVKLPKGDVSQGYRLPYFATKLDLNSHEEIDAASSCKQLDLNGFSWN
>RcoPSR_30017.m000317
MNSHDRPMCVQGDSGLVLTTDPKPRLRWTVELHERFVDAVTQLGGPDKATPKTIMRVMGV
KGLTLYHLKSHLQKFRLGKQPHKEFSDPSIKDGPALDLQRSAASTSAMMGRSMNEMQMEV
QRRLHEQLEVQRHLQLRIEAHGKYMQNMLEKAYQTLAGENMASGSYKGIGNQGVPDLGAM
KDFGPLNFPQFQDLNIYGGDQLDLQQNMDRPSLDGFMPNNDHICLGKKRSSPYSGSGKSP
LIWSDDLRLQDLGSAPTCLGPPDDLFKSDQIQIAPPSMDRGTDLDSISDIYETKPMLQGD
AMGEKKFDASTKLERPSPRRAPLPTDRMSPMINSGAMPQDRNSPFG
>RcoPSR_30169.m006598
MAELLFNLGMVSCWKRLFAANLIQLFLRCYFFGFPVCIIFGSTSPVKLWKVLGLLASENE
SLSSKGITQPYFTTLSPVHDFFNSESEGQSCLTSEFSSSRPSPFMLTDSLSPNTMQSIVQ
PPKYFLKSGPDMPLSPASHIQHSKSTFQRSSVFCTSLYLSSSSSSETNRQLGNLPFLPHP
SAHAHSLSAIDSTKSPLLFTDDISNPYDEEHSDCLMKDFVNFPGDASRSSFHGMTCASDN
LVLADQLELQFLSDELDIAITDHGENPRVDEIYETPEASSNPAIGSTCNLNVASVKPSAD

APSSHPSPGTAAVHKPRMRWTPELHESFVEAIIKLGGAEKATPKGVLKLMNVEGLTIYHV
KSHLQKYRIAKYLPDKKEEKKASCSEEKKAASSSTESDNQKKGMTQITEALRMQMEVQKQ
LHEQLEVQRALQLRIEEHARYLQKILEEQQKAGGTSLSPKDLSSLTNPPEASVLPPSQEV
VTLHPQSTESKTVSSSSKKKPTVDSEIEQPQRDKKIRVEKKPESAKEEAAVESPVQ
>SpoPSR_Spipo0G0051900
MVDDQSVAPSSHARPGLKWTGQLHERFLEVVSRLGGASKATPRSVERLMGASGLTLYQLK
IQLQARFKYRLAVKRSSEASSHGTEEEGGGGGGSFPGNQRARQAPMELQRKSEERLEVQR
HLQLRIEAQGRYLQSVLERAREMLEMDAADSQPSPAADADCLSSDGSCLTENDGPSG
>SpoPSR_Spipo11G0061900
MGQIQSHAMVGSASRARQNDWPDWVSSDSLGGSWNDYVLRTDAPLAESKALYPAASHLLV
QMTAPPQPQFYQQMPSNSGGVSAANGSLSSGSSGPPTKPRIRWTPELHERFVEAVSQLGG
SEKATPKGVLKLMRVDGLTIYHVKSHLQKYRTARFRPDSSEGAPETNLAPLEGLSSVDQG
TGLGITEALRLQMEVQKKLHEQLEVQRKLQLQIEEQGKYLQIIFETQRKMENGGHNPRST
LDSTASATAATNETPGRVGAAAADPEPAGESSAGAGAEQRRVNPESPHELDSVPSEAWPS
PPRGKRPRQGDVHP
>SpoPSR_Spipo12G0022900
MYQPNPASRLNLIPSSLIVHDQRGEMSGNAVIPGNGGRNLNNSTLASRQRLRWTHELHEC
FVDAVSQLGGPDRATPKGILRVMGVKGLTIYHVKSHLQKYRLAKYLPESSPDGIKAEQKE
PGSLLSTLENSSGIQITEALKLQMEVQKRLHEQLEDKFRVIS
>SpoPSR_Spipo12G0037300
MASSVNAGATPKGILKLMESEGLTICHVKSHLQKYRAAKNIPDPQECASSSLRVAGGSKG
EGGSFGRLDRLDPKNRMHMAEALRLQLEVQRRLHEQLEIQRTIQKKFEEQGRQLQKMFEQ
QERAARSLFEKQGPEVQAHDEGRGPEITHFPPKIS
>SpoPSR_Spipo1G0009300
MRWTPELHERFVEAVNQLGGSERATPKGVLKLMKVDGLTIYHVKSHLQKYRTARYRPEAL
EGAAEKKVSPIDKISSLDRKSGIDITEALRLQMEVQKQLHEQLEIQRNLQLRIEEQGRYL
QMMFEKQCKTGKEMFNLAPTSDIPSVQSSKSGNEAPEKDPARACDQVGADLERAEESPRK
AGAKEKMSEAEPAADATLSGAGDPESPPLKRARASNEVNPPAPPASD
>SpoPSR_Spipo22G0018300
MGGEKELKMYSKKFATMALMPHRSQGDEQLSNIGALRGSPVNSTASSGVGGKQRLRWTSD
LHDRFVDAITQLGGPDRATPKGVLKVMGVPGLTIYHVKSHLQKYRLAKYLPESPADGLKD
EKKDCSDGLGNLDPASGLQITEALKMQMEVQKRLHEQLEVQRQLQMRIEAQGRYLQKIIE
EQQKLGSYVGVNEPGR
>SpoPSR_Spipo2G0037400
MVSNRRLAPASAAKPRLKWTRQLHQQFVGAVSHLGGAAKATPKSVMRLMGVPGLSLCHLK
SHLQALSISFFLSRNILFSVKATTNLAAVIERRNEVRGSFSSKSIRFSVNGSNVPSVSLT
LPQVQTEVPNKLQEQIEVQRRLQLRMEAEEKYIRSLLKKAQEKLAVCSRSYPEMAAAVDR
GIAAGSFFGGGPAPSPPADCSSSSCLTTLERPEEVDEWLLTNEPPTMGTRDGGGGGIYSG
HSCRRRSGAQEGESGTVEIDLNR
>SpoPSR_Spipo5G0044100
MFQAKKACMNSHERPMCVQGDSGLVLTTDPKPRLRWTVELHDRFVDAVNQLGGPDKATPK
TIMRVMGVKGLTLYHLKSHLQKFRLGKQPHKELNDHHSTKDATGRAVSLDFQGNAASSSG
MMGRATNEMQMEVQRRLHEQLEVQKHLQMRIEAQGKYMQTILEKACQTLARENTATGGGE
MCFLKEMGSPLGFPSLHDLHLYAGGVHHLVDETCRGDLTHMDSTALNSGKPIFSPDRRGE
AKPEMSSPSAASPVVRTTAISARNLSYG
>SpoPSR_Spipo5G0019100
MWRNATEPRNGLNLPNPPLSSKQRLRWTDELHGRFVDAITQLGGPDRATPKGILRVMGVK
GLTIYHIKSHLQKYRLASYVPECSSDGRKAGNKEHGEEICHLDDSSIPITEALKLQMEVQ
RQLQLRIEAQGKYLQRIIEEQRRLSSVLTEGPGSTGNGKCVITSPASQRLDLGPEDKSGQ
RSQAGSFPGLVNSPLT
>SpoPSR_Spipo6G0056100
MYGHHEGKHGILTSRAAFHQDRNLLLQGANAPGDAGLVLSTDAKPRLKWTQDLHDRFIEA
VNQLGGADKATPKTIMKLMGIPGLTLYHLKSHLQKYRLSKNLHTQANNGTSKNASAGRQA
EVSGALTGTAVPSPTNKTLQIGEALQMQIEVQKRLHEQLENSMDLADHPKYASKEVVSVE

DTDWIFSFIGIKPEG
>ZmarPSR_Zosma101g00780.1
MNQSKVELVSSSSNNNNGTKIQEHGLEKSGSSVVSFNGRRNAITINKPLSSRQRLRWTHE
LHERFVDAIAQLGGPERSTPKGVLRVMDVQGLTIYHVKSHLQKYRLSKYLSVPESSPDGP
KAENRDLSDFDYPSMQITESLKLQMEVEKRLHEQLEVQKQLQLRIEAQGRYLKRIIDQQQ
RRNMVIEENPSNIDDARFQPFQQCENSEDLMIQSGSHPGTPEHEKQIKRQKDDGYVKQES
NSN
>ZmarPSR_Zosma13g00280.1
MYHHRGKNISSRLSFGHDRHLFLQDSNAQCDNGLVLSTDAKPRLKWTPELHDRFVDAVNQ
LGGPDKATPKTVMRLMGIPGLTLYHLKSHLQKYRLGKNLHAQSNGVTKTGASILASSERT
SPEISGTLILNKNQINNKSLQISEALQMQIEVQQRLHEQLEVQRHLQLRIEAQGKYLHSV
LEKAQETLGYQNLGSAGLEAIKETFLVSNDDRLNTAFQPSMVDLQKLKTQTPAASECQLD
SCLTSCDHKDVSHMSNVKMSLKICDKQQDNNDDIINQRKTTKEEDIRIMQHQHQPWLPEK
MKNSKGFKLHIMPEPQLDLNCRSQNDDVQTSKHFDLNGLFELELLEK
>ZmarPSR_Zosma175g00180.1
MFHSKKPTMNSHERPMCAQGDSGLVLTTDPKPRLRWTVELHDRFVDAVSQLGGPDKATPK
TIMRVMGVKGLTLYHLKSHLQKFRLGKQPHKEFNEHSMKDAFDMQRNASSSSSGMMGRTP
AENYHVAEALRMQMEVQRRLHEHLEVQKHLQMRIEAQGKYMQSILEKACNLAGENMTMGS
YNSSTVSNQQGVGDMGLLNSMGSPMNTFPCMQDLNIYGNGVGDQQQVDQMHQNLDGLMNT
NNDHHAGSSLVKK
>ZmarPSR_Zosma262g00120.1
MSGNSGVSYNGGKNITNSPLSSRQRLRWTNDLHERFVDAVSQLGGPDRATPKGVLRLMGA
QGLTIYHVKSHLQKYRLAKYVPDEPSSDGVNDEKKDRDELLSNSNNSSGSQITEGMKLQM
EVQKRLHEQLEVQRQLQLRIEAQSKYLKEILKEQQRFSDQCKNSDEKNPSSTSSCAPKSD
KLLPLLQDKEIEPINSLIKNIKQNNFSYRPDRSPEHEKPNKRQRESTEICHENPELVLAN
HILESSSGPDFQQNHLPLAEYNISL
>ZmarPSR_Zosma265g00260.1
MYRTKNTSTLTLFGVSSPTTATKQRLRWTSELHELFVNAVSQLGGPDRATPKGILTAMAV
PGLTLYHIKSHLQKYRLSICLPVYQATDNSKIERKDNIDNITNMDSASLDKTLELQIEVQ
KRFDEQLEIQRQLQLRIGAQVRYLQEITEDLPLVFEDKQKKQSENDNYPVKKVKKPRIGD
QAALFLSANKLDSYVTD
>ZmarPSR_Zosma289g00260.1
MQKKIRVVQTSMDGGSNIHPPPDFKLRRLRWTKQLHDQFIDAVTHLGGAEKATPKSVMLM
MDVPGLNLYHLKSHLQKYRLGQSREQRLLPDCRKQVANYIKPAEISSLDFRLQLNESMTK
MQTEIEVQRHLQLRIEAQAKYLQTVLKKAEESLAAGTDSVLIDSCLTSLERKEEDNDDDD
GEVEEDNLCKLKIGASCSSIDQSPHPLFIETQLGKHKFQYELDLNK
>ZmarPSR_Zosma296g00010.1
MEDMEIGVSAGSCRSFYSTTGDDNDNSGVMLTRDPKPRLRWTPALHDRFVDAVTILGGPD
KATPKSVLRVMGTKGLTLYHLKSHLQKYRLGKQARKDTPELKGGGEKSSCDDSNSDVLTP
DIGIMDKHGLPLTEALRYQIHVQAKLREQLEVQKKLQTRIEVQGKYLQTILEKAHDSFSF
GTMMTVAKVGEKEETTHRRDHGGATVFREMDGEGRADESRLVSLDLNVNNETFSGFRPCD
LQLGLQPHT
>ZmarPSR_Zosma305g00050.1
MFSDLIHKQKEVSPFPLEEIHGPSVVLTADPKPRLRWTADLHERFVDAVAQLGGPEKATP
KTILRTMGVKGLTLFHLKSHLQKYRLGKQSCKELGDQSKDSSYPLDATNSSGSLSPLVQT
PDANDGIEVKEALRAQMEMQRRLYEQVEVQKHVQIRMDAHQNYIKSLLEKACKIASEQIT
ASASGPETAGNDDLADLVSMGTSSVSSSGFHQTTAASEGMTFYQKLPNLRHAN
>ZmarPSR_Zosma320g00060.1
MWNMGLPAHYERCYSQHQPRSSNAPPPPLDVSGHGLSAKRSIFYVADQYDTECPRFDYRK
AEATVDPFLFTSLEKKNSSSQTPQLDLLPARNKILQPKVASKSAHSAFQPNKMRIRWTPE
LHDRFLDSVGLLGGAEKATPKGILKLMDTSGLTIYHVKSHLQKYRTARYAPEIMEGRRRC
MERTQQFDLKTGVQISEALRQQLEVQRQLHEQLEDQRNMQIRIEEQAKQLSKMFEKQAAG
TNK
>ZmarPSR_Zosma50g00270.1

MDADSSGAAKMKTKHTDCLDKTTTTFSRFSTFCSSSFSPSLVSLDGYKTLGKLPFLPNPP
KFDDNVQITAPIQPSPDPPVLSNAEMGHIDDFMTDLLDLSGGEASDATIDGYNFTCSEQV
DLEMLSEQLGINMTDNGENPQLDEIYELPQTSPLPQTAIAGYKPRMRWSVELHDRFVDAV
NKLDGAEKATPKGVLKLMNMEGLTIHHVKSHLQKYRHARCLPEAKQDTNCSEEKKSDQEV
RINRNVQLEESLRMQIEVQKQLHEQLELQRTLQLRIEEHARYLQKIFDEQEKTAASLCIG
PSQSSTSSIDQQPCNEIRHTSPERSVNKRARMDI
>ZmarPSR_Zosma58g00470.1
MCCGQSPTLETSLFDQMESFNMAKRKQTNSLQETSSQTSTSCSSLFSYDKPFGKLPFLPN
SLLEKTHHMDVSQSPSSMLLTSDGVFSQYGDRDQSGADDLSTGIFNVSDDLDGNKTQISE
DFSEQFNLQILSEQLGIQIADTDEYPQLNDIYQMPQTSSMPMMTGSSENSQLNEVYQMPQ
ISSVPMMAVSCEYSQPDQKLVTSPVMLEKSNPSTPTNAPSNKQRIRWTVELHDRFVEAVN
KLDGLSERATPKSVLKLMNMEGLTIHHVKSHLQKYRLAKYLPEAREDKKTTRFREEENVS
SRNQTEICINGNVQIAEALRMQIEMQKQLHEQLEKQRALQLRTEENARYLQKLLDGQEQA
NPVCPSLTDIFR
>ZmarPSR_Zosma6g00200.1
MYHAKTISTATLVPHKTKVTNQHPNSEYLGVSPASNPVTLGGSSKQRLRWTSDLHERFLD
AISQLGGSDRATPKGVLRVMGVSGLTIYHIKSHLQKYRLVKYLPESPINGSTSDKKESTD
NLMNMASEPGIKINEALKMQMEVQKRLHEQLEVQRKLQLRIEAQGKYLQKIIEEQEKLGN
ALTTSEPIPQPPNTPASSIKKRRIIGDKQAEGSPNAALEQENMTNAPRALR
>ZmarPSR_Zosma70g00660.1
MEMNGDVAIPYNGGTNQTTHPLTSRQRLRWTHELHERFVDAVAQLGGPDRATPKGVLRVM
GSQGLTIYHVKSHLQKYRLAKYLPDSSSDGLKAEKKDSGDLLSNLDNSSSGSQITEAMKL
QMEVQRRLHEQLEIQRQLQLRIEAQGKYLKKIMEEQQRLSGVLSEGSRLEISKPDPIDQC
QTSEKNKPCTPTPTSESHLHVKSTNGAGVVDGVLKCLSGADEDSLSSCHEPLTPDSSYQI
VSPSRSPENEKVIKRQRSSIEFGQQVN
>ZmarPSR_Zosma9g01070.1
MEASNKSNRCSYFSHSRTKSSSLPTNPNSQLVSIERELQSNTAEPSFAAHLLSNSSTMSS
LYSPDINFSVHKQNMSSSSFVLDHQSPSNNIVSPIPLSYSLPSGESLPSQQLIEGEHWND
DAIQELLNFNSGVSVDHCSIQNGQVLASEDLTPHNRQKSQNQWSNWTASDACLTSNWNEI
LNDVTAMDSIHTPEFTNTLESHQLMPVPSNSGELRAFPSHVSSSNCATPTKQRMRWTPEK
HERFVKSVNQLGGCEKATPKAVLKLMNFEDMTIYHVKSHLQKYRTARYRPDSSEGTSERN
NTLDEISSLDLKAGSIGITEALRLQMEVQKQLHEQLEIQRNLQVKIEENGRRLQMMFEKQ
QKASGEKVHLSDTTEGQSSDDNDNVNLKITSNDDKDRQVGEREKIINPNTNNGGSISQNC
QPAKRSKNSHIPDSLASK
>ZmaPSR_AC155434.2
MYHQQQLHTHNQHLSSSRPGLPPEKQFLLHGAGGGGGGGDAGLVLSTDAKPRLKWTPELH
ERFVEAVHQLGGPDKATPKTIMRLMGIPGLTLYHLKSHLQKYRLSKNLQAQAHTASAKNA
LVGCRTGADNALCQGSASPPPPPPPPHLNLEPPQINRSMHISEALQMQIEVQRRLHEQLE
VQRHLQLRIEAQGKYLQSVLEKAQEALAKQSGGADETTTTTQQQQQLLLPDLISRATATR
RGNVQQEHLHQQHHLGGDGSVDSCLTACEGSRCQRERDQDLLSIGLSSAPPPTPTPSRGY
NDRGGGGSASCEEFLQFLDEPSRRGAGGGGSSDEQQELDLSISGGRSNPRPRGSQSQRID
LNGSSWN
>ZmaPSR_AC233960.1
MNMFDGMERAGYGGGAGPMGGVVLSRDPKPRLRWTPDLHERFVEAVTKLGGADKATPKSV
LRLMGMKGLTLYHLKSHLQKYRLGKQTKKDTGLDAGRGAFAAQGINFSTPVPPPSIPSTA
SDNTGETPLADALKYQIEVQRKLHEQLEVQKKLQMRIEAQGKYLQTILEKAQSNLSYHAT
GAANLEATRSQLTDFNLALSGFMDNVTQACEQNNGELVKAMSEDGLRANNLDFQPYHGVH
DGDDVKCATDEGLLLLDLNIRGGYGHRISTDLKMNQHMR
>ZmaPSR_AC234155.1
MEQHPFLRGGNAQGDSGLVLSTDAKPRLKWTPELHQRFVDAVNQLGGEEKATPKTVMRLM
GIPGLTLYHLKSHLQKYRLSKNLQAQANASTSKNAIGCTPVADRIPGTTAATMSSTNVLP
QAEKTIQIGEALQMQIQVQRQLNEQLEVQRHLQLRIEAQGKYLQAVLEQAQETLGKQNLG
PANLEDAKIKISELVSQVSNECFSNAITDVKESSSVHRLEPIQIEFVESSTNSYLSVAEG
FIKEHKLQHHGVLKAYDDSSLFCRKRSHEHETQFALNRSLSEHRMAHLQNEEGYHKAEFG

YESDTEMAHEYTAPQKNGRCSTTSSASGSKVDAEKLYLEEQKCARQAVEYPRESKLVDFE
HPCSVNKLDLNTHNVDDTNQAYRHFDLNGFSWG
>ZmaPSR_GRMZM2G006477
MRNFNLMQSQKSRVLGAMSSSLPILPNPLKGSFSRPHNPQHIPMLRQLPDDSMPLCIDTH
QSASLHPRAGVIGVPYSGYTASPLDSVSNLDSQTMAAPFISQSSNFEALQSLSNNTPETH
TKAAWFTSSMDVSPLNTDNIAASDVNQIQSIRPAMTSDESATQNDWWADIMNDDWKDILD
ATATDSHSKAMIQISNSATSLPAVNQSASSHSREICPVASPPNSSNASVAKQRMRWTPEL
HECFVDAVNQLGGSEKATPKGVLKLMKVDGLTIYHVKSHLQKYRTARYKPDLSEGTSEKR
TATEELVLDLKTSMDLTEALRLQMEVQKRLHEQLEIQRKLQLRIEEQGKYLQMMFEKQSQ
SSTEKVQDPSSRDTTAKPSSNQSQSTNKDCGATMDPNGTGGIVRTAELGERSSELGVKQK
LVEIEESGTEVATGDRSKISQEKRRKLQDS
>ZmaPSR_GRMZM2G009060
MSSCQMYHHQQQLQSHSHFLSSRQTFPPERHMLLQGGSIPAEPGLVLSTDAKPRLKWTPE
LHERFVEAVNQLGGPDKATPKTIMRLMGVPGLTLYHLKSHLQKYRLSKNIHAQANGVNAK
NVIGCTMAMDKPLEGNGSPASHLNLGTQTNKSVHIGEALQMQIEVQRRLHEQLEVQRHLQ
LRIEAQGKYLQSVLEKAQETLSKQNAGSVGVETAKMQLSELVSKVSTECLQHAFTGFEEI
DGSQILQGHTIQLGDGSVDSCLTACDGSQKDQDILSISLSAHRGKEIGGMTFDIQEKERG
REDLFLDKLSMTPPGQLDRRERGSFSMTRKAAKLDLNINDTTDGPQNCKKIDLNGFNWT
>ZmaPSR_GRMZM2G010920
MNFLESIAVNGHRDRQHGGVMSRQSMVAVEQLTGPASARPAHTCAAQPHAHQLLSARYDH
RGACASQVELPDRISPSNASTFCTSMYSSSSSSSAAADSSKSCRRAGALPFLPHPPKREQ
QQQQQSSSSPSLLVCTDLGNGGQDEAEAEHPAELKDFLNLSGEASWRGGFHGESSGVAFR
FGEQVEFQFLSEQLGIAITDHDESPRLDDIYDRPPQTSSCPVLSCSDQEEGLQRAGSPVK
AQPSSSRAASCNNKPRLRWTLELHELFVKAVNKLGGPEKATPKGVLRLMKVEGLTIYHVK
SHLQKYRFAKYLPETKEDMKSSSEDKISKSEMPGSNAGRKKILRSLQVAEALRMQMEVQK
QLHEQLEVQRQLQVRIEEHAKYLHKILEQQKARNSLSATTSSIETELSESTKEEKPETQA
DTSSAQLPNKKNSDTDME
>ZmaPSR_GRMZM2G035370
MLLIKLVYLLLRLISWFAVPVCFPARLFSPDTWSGDRWMMDAKKIKLHDYHLYGSQQLFP
AAGGLSFHPAAGLMSSLPQPPHAAWLHEDHHTTTPRSVLATHGSLQGSRCVGSDAAAFFA
AEELMMGMPRFDDCPLGGTEMTAFAKRPTTEDEQLHYRRPGPVDPLPLRDSAAVRTYYVR
PQQRDGATEAPPSLELPFQQGRRQQQQLFGNASTGRLLGGEPKAHSFTAHVATSTLLPAM
EAPAGMQQSPTENPLSRSCSTIGAPATHVGSGNVAAAAAPGHGAPSKTRIRWTQDLHERF
VDSVNQLGGADKATPKGILKLMNSDGLTIYHIKSHLQKYRIAKYMPASSTSEGNLIYRKQ
EKRAVGNDVQNLDPSTGMKITEALRVQLDVQRRLHEQLEIQRNLQLRIEVQGKKLQKMFE
EQMKASRTVMEPRQEEEDDDDDAFNDVHVQLLAAAARSDAGFQLNIS
>ZmaPSR_GRMZM2G052544
MFPSKKATSSAAAAAAAAVSSNDSRQQAMCVQADSGSGLVLTTDPKPRLRWTVELHERFVD
AVTQLGGPDKATPKTIMRVMGVKGLTLYHLKSHLQKFRLGKQPHKEFNEHSVKDAAAAME
MQRNAASSSGMMGRSMNDRSVHMNEAIRMQMEVQRRLHEQLEVQRHLQMRIEAQGKYMQS
ILEKAYQTIATGDLAACSPVAAGYKSLLGNPQAMLDVCSLKDMAPSMGFPSLQDLHMYGG
GGGHCLDLQQQMERPMEAFFASCDIGSLAKKRPVSPYADDDGGKSPMLWGEDDEGKGIVD
HFQMAPPMMDAAGIDVMDSIADVYGDAKHMTMSSDSTGSKGGGFDVRLERPSLRRPHMGG
SPSVLGGGQTRNLSYG
>ZmaPSR_GRMZM2G064197
MFPPGLIHHRPDATAPGDGPPRSGPGGPSLVLTADPKPRLRWTADLHERFVDAVAQLGGP
EKATPKTILRTMGVKGLTLFHLKSHLQKYRLGKQSDKEGSEQSKDASYLLDAQSGMSVSP
RVAAQDMKESQEVKEALRAQMEVQRRLHEQVEQVQKRVQIRMEALEKYIDSILESACKMV
TEQFASSGFSISNPDLPEISPGGVMCGSTDTLGSSVLNQLSVSSIDSHSPGGKPSPSGME
GPTLLQKSPELKRRSS
>ZmaPSR_GRMZM2G081671
MFPSSKKQASTGAANPNDRPMCGQGGDSGGLVLTTDPKPRLRWTPELHDRFVDAVAQLGG
PDKATPKTIMRVMGVKGLTLYHLKSHLQKFRLGKQHKELGDHTAMEMQRSVASSSGMIAR
SMNDRSVNVNEALRIQMEVQRRLHGELEVQKHLQMRVEAQGKYMQSIVEKAYQALGSSDC

ATWPAGYRTLGSQGVLDIGTSSTSFSSVQDLQCFYGGSSHMDQLLHQMERPMDGFLTLGE
SCFIGSADNKKDLNNHCSSSGKSSMMWASEEQQAKSGNDQLQMGSSTRMEGAGIDVMDPV
TGLYEGAVSGDSMDSKGFEGSNSRLEMKPPAQQAPVGSQRIRI
>ZmaPSR_GRMZM2G113742
MFPSLIHHASHGRPDVAVAEDAPRSGCGLGLPGHGGGPSVVLTADPKPRLRWTADLHDRF
VDAVAQLGGPDKATPKAIMRTMGVKGLTLFHLKSHLQKYRLGRQSGKELTEQSKDASYLM
EAQSGTTLSPRGSTPDVKESQEVKEALRAQMEVQRRLHEQVEVQKHMQIRMEANQKYIDT
ILDKAFKIVSEQLSGFSISDQDLPILTSARAMLSPADHLSSSVFPQLSVSSVSLHNPGPG
GGKGLPHVADSHVFSQRPPEQFKRKSR
>ZmaPSR_GRMZM2G117854
MMYHAKNFSVPFAPQRAQDNEHASNIGGIGGPNISNPANPVGSGKQRLRWTSDLHNRFVD
AIAQLGGPDRATPKGVLTVMGVPGITIYHVKSHLQKYRLAKYIPDSPAEGSKDEKKDSSD
SLSNTDSAPGLQINEALKMQMEVQKRLHEQLEVQRQLQLRIEAQGRYLQMIIEEQQKLGG
SIKASEDQKLSDSPPSLDDYPESMQPSPKKPRIDALSPDSERDTTQPEFESHLIGPWDHG
IAFPVEEFKAGPAMSKS
>ZmaPSR_GRMZM2G124495
MNAKKIKLHDYHHCYGSPMCDPQLFPRAAAATTAGLSLHPGPGLVGSLPQRHGGGGGGWV
HEEHTATTPRAAQGQGGCVVGSDAAAFFAAEELMMGTARFDSPLGGTTTALQELTAFAKG
PPFGRPRPTTGGERERERLYPVDPLPLRDCAAVRTYYVRPQQRDGATEAPPSLELPFQRR
QQQQVHGLFGDPSTGRLLGGGEPQAHSFPAHTLKQVPASTFVPAMEAPPGMQSLMDNPLS
RSCSIIGAAATHAGSGNAAAPGQGAPSKTRIRWTQDLHERFVDCVNKLGGADKATPKGIL
KLMNSDGLTIYHIKSHLQKYRIAKYMPVSSTSEGKEKRAAAANDVQNLDPGTGMKITEAL
RVQLDVQRRLHEQLEIQRNLQLRIEAQGKKLQKMFEEQMKTSRTVMGPPQGADVAFIGAG
EQEEEVEVEDAFDDVQLLAAVSSASVGYRDGGFQSKIS
>ZmaPSR_GRMZM2G125704
MQGSYGYDGAASRDPKPRLRWTPDLHQRFVDAVTKLGGPDRATPKSVLRLMGMKDLTLYQ
LKSHLQKYRLGIQGKKSTGLEPASGGVLRSQGFGSTTAHPPPGVPDQGKNTREIALSDAL
RYQIQVQRKLQEQTEVQKKLQMRIEAQGKYLKTILEKAQTNISFHTNASNGIESTRSQLM
DFNLDGFMNNATQVCKEHREQLVKAMSDENDKDSLGLQLYHLGSQEAKEVKCTPKTEDSL
LLDLNIKGGYDLSSRGMQACELELKINQQIV
>ZmaPSR_GRMZM2G162409
MERLSTNQLYSSGVPVTVPTSLPCIPVSLDESFPRLPDAQSVLMERELRSTPLPPHQTTV
APIRGLFHSNTGSVGPLCSPPSVRFSSHSNPEQYPNHNSYNSQVPSTASSSTLNYGSQYG
GFEPSITDFPRDIEPTWCPDPVESILGYSGDVPAGNNLTGTTSIGASDDLTKQTEWWTEF
MNEDWKDMVDNPTSTETQPVGQPVQSSNSVHQSATQQTVSSQSVEPLAVVAPSPTAGSNT
GKARMRWTPELHERFVDAVNILGGSEKATPKGVLKLMKADNLTIYHVKSHLQKYRTARYR
PELSEGSSEKKVASKEDIPSIDLKGSFDLTEALRLQLELQKRLHEQLEIQRSLQLRIEEQ
GKCLQMMLEQQCIPGAEKAQDASTAADELRSPSEIPESSTVKEVPENSQNGSTKQTK
>ZmaPSR_GRMZM2G168002
MASSLGVGGQQQGHGHGHGEARARLRWTRQLHDRFVLAVAQLGGADKATPKSVLRAMAVP
GLTLYHLKRHLQKYRLVAVSRGVASPLGDSGDGTDERSSSSSENQPADECDDGTVAEPHG
DSSRSVARMQRKLQEQIEVQRHLQLRIEAQGRYLQSVLRRAQEVLADHGLGSAAGAEAEL
ASAVDTGSLSSSCCCSPTRRRSADSCVTSSSSEAESQAPVGGATWLHTCAGTRDCSVEQQ
PVQGEKSTFLQRHGAAEDGTSSEIDPNGNR
>ZmaPSR_GRMZM2G173943
MYSPKPESSFGPNPNSGTHQQQMELTGANMGPGNGANNNTNMAGRQRLRWTNELHERFVE
AVTQLGGPDRATPKGVLRIMGVQGLTIYHVKSHLQKYRLAKYIPDASTDGNKTDNKDPGD
LLAGLEGSSGLQISEALKLQMEVQKRLHEQLEVQRQLQLRIEAQGKYLQKIIEEQQRLTG
VKSETPAGGASVTVSSDQFPDSERTEPSTPAPASESPTQVGASNRDTGDRTEATKSTCHG
DSLSRNEPLTPDSNCQNGSPVASPNHERAAKRQRGSGTEFLDSEAEFSLPRHIFESSSGS
EFQQYSMSYSGQ
>ZmaPSR_GRMZM2G379167
MGITCSHTSTMNRLHSGRVPPCLSSAITPVPAFSEASFPSFTEQRSSFAERKLPVSSVAS
IGTSMAAYDTMSTHMLASNFTNESPNRQLCSGSYVTEPSDPDTPLLATHSSLMADSTPLF

VDFSEVSEHICLNQGQLLGLFDYPASVDFYKSKNMTIFGQQIQDTITVDPNTHLTPQNEL
FSGSSMQLQKTVGSAESVLKTVDARSATPQSYLYCHTESERSVPEPFNCDKLGADSLPFS
NTAPKPRMRWTPELHECFVDAVNKLGGSEKATPKAVQKVMKVDGLTIYHKHRIVQHRSAG
VPGRRGSHTEVDDDSIPQSKGEGGVEGGLVSQIGLQKQLHEQLEIQRRLQLQVEEHNKYL
ETVIAKQNESLKKLRALRGFRDRVRRVLQDSEAPGVRAHCAQ
>ZmaPSR_GRMZM2G701218
MYHQHQGLSELFTTRTSFPMEQHLFLRGGNAQGDSGLVLSTDAKPRLKWTPELHQRFVDA
VNQLGGAEKATPKTVMRLMGIPGLTLYHLKSHLQKYRLSKNLQAQANVSTSKNAIGCTSI
ADRIPGTSAATMSSTNVVPQAEKTIQIGEALQMQIEVQRQLNEQLEVQRHLQLRIEAQGK
YLQAVLEQAQETLGKQNLGPANLEDAKIKISQLVSQVSTECFSNAITDVKGSSSVHRLEP
RQIQFVESSTNSYLSVAEGFIKEQRLQHQQHHGVLKAHDGSSLFCRKRPHEHETQFALNR
SLSERRMAHLQNDEQYSKAEFGYESDTEIVHEYTAPQNGGGSTTSSASGSKVDAEKLYLE
EQKCARQVAEYPRESKLIDFENSCSGKKLDLNTHNVDDTDQAYRHFDLNDFSWS
>ZmaPSR_GRMZM5G869984
MRESEELDIEIAKPAGSLVEEMDVGRSVCSPAVEIEGFAERVGSSTIAPSPCIPSHDMNL
QSLPVATGAPYPASMAPSSSAFSTVTTHGCFPYSTPSTAHPSLSGILPCNNNMISYSVLP
EEPSGGIFSGQSPEGCADPGDIDYRVESQQIPGPGRTVDESDNRDEWFVPTDITWNWHQM
AESAPVPSYPNSKSWQHDEQHMNFGTVSLSPYKVLLAFYCAKPKAILDIMNVEGLTRDQV
KSHFQKYKLQVKQHPSEVPGTSVEMTMRSEAIPSDVQKFVTTRKNKAFANYHLCHHRHFY
KKSLIFLCNQPGVHLTRHIQDYALQVQVEFQKKLHDMVESTMLETSRSSEQGRALQRDGD
DDAVSEPEPIGGRYVPVLGHGDREDLVRGGDRGGHEDLAPMLVYDDGRDDHDGLAPVLVH
GGRGDHDDLPPVHHRGHEHDENPTAAPPGRDDHGGGDEGSERRQEHRPAPDARSAP

>SbiPSR_Sobic.001G384300.1
MQSQKSRVLGAMSSSLPILPNPLKGSFPKPCNPQHIPMSRQLPDDSMPLRNDIHQSASLH
PRAGVIGAPYSGYSASPLDSVSNHDSQSMVAPYISQSSSFESFQSLSDTTPGTHTEAAWF
TSSMDVSPLYTDNIAAPDDNQIQSIRPAMTSDETAKQNDWWADIMNDDWKVILDATATDS
HSKAMIQTSNSATSLPAVNQSASSHSMEICPVASPPNSSNASVAKQRMRWTPELHECFVD
AVNQLGGSEKATPKGVLKLMKVDGLTIYHVKSHLQKYRTARYKPDLSEGASEKRTATEEL
VLDLKTSMDLTEALRLQMEVQKRLHEQLEIQRKLQLRIEEQGKYLQMMFEKQSQSSTDKV
QDPSSRDTVAKPSSTLSQSSNKDSGATMDPNGTGDSTKTAKLGERSSGSGVNQKLVEIES
DTEGGTDDGCKISQEKRHKLQDS
>SbiPSR_Sobic.001G386700.1
MYHHQQQLQSHSHFLSSRQTFPPERHMILQGGSIPAESGLVLSTDAKPRLKWTPELHERF
VEAVNQLGGPDKATPKTIMRLMGVPGLTLYHLKSHLQKYRLSKNIHAQANGGNAKNVVGC
AMAMEKPPEGNGSPASHLNLGTQTNKSVHIGEALQMQIEVQRRLHEQLEVQRHLQLRIEA
QGKYLQSVLEKAQETLSKQNAGSVGVETAKMQLSELVSKVSTECLQHSFTGFEEIEGSQI
LQGHTIQLGDGSVDSCLTACDGSQKDQDILSISLSAHRGKEIGGMAFDMQAKERREDLFL
DKLSMMPPSHLDRHERDSFSMTRKAAKLDLNINDTTDGPQNCKKIDLNGFNWT
>SbiPSR_Sobic.002G121600.1
MERLSTNQLYKSGVPVTIPTSLPSIPASLDESFPRLPDTQSVLMERELRSTPLPTHQTTV
APIRGHFHSNTGSVGPLCSPPSVRFCSLSNPDQYSNHNPYNSQPPSTASSSTLNYGSQYG
GFEPSITDFPRDVEPTWCPDPVESILGYSGDVPAGNNVTGTTSIGASDDLTKQTEWWTEF
MNEDWKDMVDIPTSTETQQVGQPVQSSISVHQSATQQTVSSQSVEPLAVVAPSPTAGSNT
AKARMRWTPELHERFVDAVNQLGGSEKATPKGVLKLMKADNLTIYHVKSHLQKYRTARYR
PELSEGSSEKKAASKEDIPSIDLKGSFDLTEALRLQLELQKRLHEQLEIQRSLQLRIEEQ
GKCLQKMLEQQCIPGTEKAQDASTTADELKLPSEIPESSTVKEVRENCQNGSTKQTGLNQ
VVFVLLWEQMNAIGNFFAKPGKVHVTRS
>SbiPSR_Sobic.002G161700.1
MGVKGLTLFHLKSHLQKYRLGKQSGKEGSEQSKDASYLLDAQSGMSVSPRVPAQEMKESQ
EVKEALRAQMEVQRRLHEQVEVQKRVQIRMEALQKYIDSILGSACKMVTEQFASSGFSIS
DPDLPEISPGGIMCGPTDTLSSSVFNQLSVSSIDSHSPGGKPSPSGMEGPPLLLQKSPEL
KRRSS
>SbiPSR_Sobic.002G161800.1

MYSPKPESSFAPNPNSGTHQQQMELPGANMGPSNGANNNTNMAARQRLRWTNELHERFVE
AVTQLGGPDRATPKGVLRIMGVQGLTIYHVKSHLQKYRLAKYIPDASTDGNKADNKDPGD
LLAGLEGSSGLPISEALKLQMEVQKRLHEQLEVQRQLQLRIEAQGKYLQKIIEEQQRLTG
VKSETPAGGASVTVSSDQFPDSERTEPSTPAPTSESPTQVGASNRDPGDRTEGTKSTCHE
PLTPDSNCQNGSPVASPNHERAAKRQRGSGTEFLDSESEFSLPRHIFESSSGSEFQQYSM
SYSGQ
>SbiPSR_Sobic.002G422000.1
MYHQQQLHNHNQHLSSRPGLPPEKQFLLQGGGDAGLVLSTDAKPRLKWTPELHERFVEAV
HQLGGPDKATPKTIMRLMGIPGLTLYHLKSHLQKYRLSKNLQAQANAVNAKNALSCRTGT
DNPCEGSGSPPPHLNLEPQINRSMHISEALQMQIEVQRRLHEQLEVQRHLQLRIEAQGKY
LQSVLEKAQEALAKHSGVHLDGGETSTQQLSELISRATATRRAHVQQDHQHQHQHQHQRH
LGGDGSVDSCLTACEGSQCQRERERDQDLLSIGLSSATPPTPSRRGYNNDRCGAICEEFL
FLDEPGRRERGGGGGSSDEHGQQELDLSINGRSNPPKPRDSQRIDLNGSSWN
>SbiPSR_Sobic.004G036500.1
MTKETNSLNLKAVKQFTSPGRMAHTRTAPQPSAHQLLSAPHDLLPSSSCAASQSSKILPF
DLQKCSPDSTNGTPVSRVSQAGLLSKDLVSPSSSSTFCTSMYSSSSTNSKSCRQADALPF
LPHPPKSEHQQTSGGQSSSSSSLLFGTDLNNGGQSEAEPAADLKDFLNLSGDASGGSFHG
ESNGMAFREQMEFQFLSEQLGIAITDNEESPRLDDIYDRPPQTSSCPVLSCSDQDDLQRT
GSPVKVQLSSSRAASCNKPRLRWTLELHELFVKSVNKLGGPEKATPKGVLKLVKVEGLTI
YHVKSHLQKYRFAKHLPETKEDMKFSSEDKISKSEIPGNNAGRKKSLQLAEALRMQMEVQ
KQLHEQLEVQRQLQVRIEEHAKYLQKILEQQKASNSLPAMTSSIERELSESKEEKKPKTQ
ADTFSAPLPN
>SbiPSR_Sobic.004G058700.1
MFPSKKATSSAAAAAAAAVSSNDRQAMCVQQGDSGSGLVLTTDPKPRLRWTVELHERFVD
AVTQLGGPDKATPKTIMRVMGVKGLTLYHLKSHLQKFRLGKQPHKEFNEHSVKDAAAAME
MQRNAASSSGMMGRSMNDRSVHMNEAIRMQMEVQRRLHEQLEVQRHLQMRIEAQGKYMQS
ILEKAYQTIAAGDVAACPAAGYKSLLGNHHQAMLDVCSLKDMGPSMGFPSLQDLHMYGGG
GHLDLQQQMERPMEAFFASCDIGSLAKKRPISPYADGKSPMMWCDDEDGKGIVDQLQMAP
SMMDAAGIDVMDSIADVYGDAKPMMSGDSTGSKGGFDVKLERPSPRRPHMGGSPSVIGGG
QQTRNLSYG
>SbiPSR_Sobic.004G270600.1
MSMFEGMERAGYGGAGPMGGVVLSRDPKPRLRWTPDLHERFVEAVTKLGGPDKATPKSVL
RLMGMKGLTLYHLKSHLQKYRLGKQSKKDTGLEASRGAFAAQGINFSTPVPPSIPSTASN
NTGETPLADALKYQIEVQRKLHEQLEVQKKLQMRIEAQGKYLQTILEKAQSNLSYDATGG
ANLEATRSQLTDFNLALSGFMDNVTQVCEQKNGELAKAMSEDGLRASNNLGFQLYHHGVQ
DRDDVKCATDEGLLLLDLNIRGGYDHRASADLKMNQHMR
>SbiPSR_Sobic.004G342200.1
MGTACSHTSTMNRLHSGRVPPCLSSAITLVPAFSEVSFPSFPEQRLSFVERELPISSVTS
FDTSMAAPDTMSTHLASSFTNESPNGQLCSGSYVTEPCDPDTPLLATHSSLMGNSRPLVM
DFPEISEQICSNQEQLLGLFDYPASVDCSKSKNVTTFGQQVQATITVDPNTHLTSQNEWF
SSGSPMQLQKNVGSAESVLKTVDARSATPQNYLFCHTQRSVPDPFNCDRLGADSLPSSNT
APKPRLRWTPELHERFVDAVNKLGGSEKATPKAVQKVMKVEGLTIYHKYRTVQHRSDGVS
GRSGKADEDSIPQSKGKGNVEGVMAQIGLQKQLHEQLEIQRKLQLQVEEHSKYLETVIAK
QKESLKKLGALRGFRDQVRQILKDGEAPEEWTHSAQQR
>SbiPSR_Sobic.006G202300.1
MLRGSYGCGGAASRDPKPRLRWTPDLHQRFVEAVTRLGGPDKATPKSVLRLMGIKGLTLY
HLKSHLQKYRLGIQGKKSTGLELATGGVFSQGLSSTTAHPPGVPDEGKSTREIALSNALR
YQIQVQRKLQEQIEVQKKLQMRIEAQGKYLKTILEKAQTNISFDTDASNGIESTRSQLMG
LNQALSGFMDNATQVCKENRKQLVKALSDDNDKDNLGFQLYHVGSQEAKEVKCTPKTEDS
LLLDLNIEGGYDLSSRGMQACELELKINQQIG
>SbiPSR_Sobic.006G256200.1
MMNAKKIKLHDYHRYGSPLCDPQLFPATAAAGLPFHPAPGLVGSLPQPHGGGWVHEEHTT
TTPRSVLATQGLQGGSCVGSDAAAFFAAEELMMGMPRFDCPLGGTPPPLPDLTAFAKRPP
FGRPTTEAEQLYHRRPVDPLPIRDGSVRTYYVQPQQRDGATEAPPSLELPFQRRQQQQQ

QQQERVHGLFGNASAGRLLGGTGGEPKAHSFPAHVAPSTLLPVMEAPAVMQQSPMENLLS
RSCSIIGAAATHVGSGNAVAAAAAAPGQGAPSKTRIRWTQDLHERFVDCVNQLGGADIFS
CCCVTLAEATPKGILKLMNSDGLTIYHIKSHLQKYRIAKYMPASTSEGKQEKRAAGNDVQ
NLDPTGMKITEALRFQLDVQMRLHEQLEIQRNLQLRIEEQGKKLQKMLEEQMKVSRTVME
PQQGAAAAAAAAFLGVGERDEEEVEDAFDDVQEQLLAAMASSDAGFQSKIS
>SbiPSR_Sobic.007G137300.1
MSSLGGQQQQHGHGEAARARLRWTRQLHDRFVLAVAQLGGADKATPKSVLRAMAVPGLTL
YHLKSHLQKYRLAVSRGVASPLGDNGDGTIERSSSSESQPDEYDDDGTIAELHGDSSRTM
ARMQREVQRKLQEQIEVQRHLQLRIEAQGRYLQSVLRRAQEVLADDHSLGSPAGAEAAKG
ELSELAASAVETAGCLSSSCSCCCCSPSPSPTRHRSADSSCVTSSSSSEAESHQAPAAGA
KWRLHTCGAGTTRDCSDVDQPVLQAEESTFLQRHDAEDAEDGTSSEIDLNR
>SbiPSR_Sobic.009G178000.1
MYHQHQGPSELFTTRTSFPMEQHLFLRGGNAQGDSGLVLSTDAKPRLKWTPELHQRFVDA
VNQLGGAEKATPKTVMRLMGIPGLTLYHLKSHLQKYRLSKNLQAQANVSTSKNAIGCTSV
ADRIPGTSAATMSSTNVVPQAEKTIQIGEALQMQIEVQRQLNEQLEVQRHLQLRIEAQGK
YLQAVLEQAQETLGKQNLGPANLEDAKIKISELVSQVSTECFSNAITDVKGSSSVHRLEP
RQIQFVESSTNSYLSVAEGFIKEHRLQHHGVLKAYDDSSLFCRKRSHEHETQFALNRSLS
ERRMAHLQNEEGYSKAEFGYESDTEMAHEYAEPQKNGGGSTTSSASGSKVDAEKLYLEEQ
NCARQAVEYPRESKLVDFEHPCSGKKLDLNTHNVDDTDQAYRHFDLNGFSWS
>SbiPSR_Sobic.009G180500.1
MMYHAKNFSVPFAQQRAQNNEHASNIGGIGGPNVSNPANPVGSGKQRLRWTSDLHNRFVD
AIAQLGGPDRATPKGVLTVMGVPGITIYHVKSHLQKYRLAKYIPESPAEGSKDEKKDSSD
SLSNTDSAPGLQINEALKMQMEVQKRLHEQLEVQRQLQLRIEAQGRYLQMIIEEQQKLGG
SIKASEDQKLSDSPPSLDDYPESMQPSPKKPRIDALSPDSERDRTQPEFESHLMGPWDQE
ICAKNICGVAFPVEEFKADPGMSKS
>SbiPSR_Sobic.010G220000.1
MFPSSSKKQAASTGAASSNDRPTTMCGQAGGDSGGLVLTTDPKPRLRWTPELHDRFVDAV
AQLGGPDKATPKTIMRVMGVKGLTLYHLKSHLQKFRLGKQHKEFGDHTAMEMQRSVASSS
GVIARSMNDRSVNVNEALRIQMEVQRRLHGELEVQKHLQMRVEAQGKYMQSIVEKAYQAL
GSSDCTTWPAGYRSLGSSQAVLDIGTSSTSFSSVQDLQGFYGGSSHMDQLLHQMERPMDG
FLTLGESCFIGSADNKKGPNHCSSSGKSSMTMWASEEQQQQQAKSGNDQLQMGSSSTRME
GAGTDVMDPVTGLYEGAISGDSMDSKGFEGSSSRLKMKPPAQQAPVGSQRIRI
>SbiPSR_Sobic.010G254300.1
MESVNHLQAMNSQSLLAMEQIAASDKTAHVSALSVHKLFDAKLDHHSLMDGTSASTSQSS
NIKTELIRSSSLSRSLSVNLQKRSPESDPESPQSHVSHPKFSEPMFSNSSTFCTSLFSSS
STKTEPCHQMGTLPFLPHPPKCEQQVSAGQSSSSSLLLSGDTGNGLDEAEQSDDLKDFLD
LSGDASDGSFQENNALAYDEQMEFQFLSEQLGIAITDNEKSPHLDDIYGTPPQLSSLPIS
SCSTQSIQDLGSPVKVQLSSSQSSSSSATTNKSRLRWTLELHERFLEAVKKLEGPEKATP
KGVLKLMKVEGLTIYHVKSHLQKYRLAKYLPGPKEDKKASSEDKKAQTGKSGSDSSKNKN
LQVAEALRMQIEVQKQLHEQLEVQRQLQLRIEEHARYLQKILEEQKAGSLSLKAPTKAQA
TESPESTLDEVSTTPQPSRNRNPVVDTECKSPVDTECKSPARPSKNRIPVVDTECKSPAR
IKRTKVQVDLENETLCS

>SitPSR_Seita.1G069400.1
MFSSKKATSAAAAAAAVSSNDRQMCVQGDSGLVLTTDPKPRLRWTVELHERFVDAVTQLG
GPDKATPKTIMRVMGVKGLTLYHLKSHLQKFRLGKQPHKEFNDHSVKDAAAMEMQRNAAS
SSGMMGRSMNDRSVHMNEAIRMQMEVQRRLHEQLEVQKHLQMRIEAQGKYMQSILEKAYQ
TIASGDVAACPAGGYKSLGNPAIVDVCSFKDIGPSMGFASLQDLHMYGGGHLDLQHQMER
PMEAFFANCDMGSLGKKRLSPYAAGKSPMMWGDDEQGKGGIDHLQMAPPMMDAGGIDVMD
SIADVYGDVKPMMSSDSTGSKGCFEGKLERPSPRRPHMGNERMGSPSVIGGQTRNLSYG
>SitPSR_Seita.1G092500.1
MSTQSIISVKQFAGPHRMAHTCTAPQPSAHNLLSAKSDNCGSAHDPQSSWAAVQTSSIKS
EMVGSLSLTKILPFNLEKCSPGSNPDSAVSHVSQAELSDPVSSSSSTFCTSMFSSFQTNS
ESCRQKGALPFLPHPPKCEQKLQQQISAGQSSSSSSLLFGADLRSGGHDDAGDLKDFLNL

SGDVSEGSFHGESSAMAFSEQMEFQFLSEQLGIAITNNEESPRLDDIYDRPLQTSSCPVP
SYSGQEDLPSAVSPVKVQLSSSRPEACNKTRLRWTLELHERFVEAVNKLGGPEKATPKGA
LKLMKVEGLTIYHVKSHLQKYRFAKYLPETKEDKKSSSEGKKSQSAIPGNDAGKKSLQVA
EALRVQIEVQKQLHEQLEVQRQLQLRIEEHARYLQKILEEQQKARNSLSTTRNSAQEELP
ESTEKEETGMKVETSSEPLSRSKISDTDV
>SitPSR_Seita.1G286400.1
MFEGMERAGYGGAAAMGGVVLSRDPKPRLRWTPDLHERFVEAVTKLGGPDKATPKSVLRL
MGMKGLTLYHLKSHLQKYRLGKQSKKDTGLEASRGAFAAQGINFSAPVPPSIPSTAGNNT
GETPLADALKYQIEVQRKLHEQLEVQKKLQMRIEAQGKYLQTILEKAQNNLSYDASGAAN
LEATRSQLTDFNLALSGFMDNVSQVCEQNNGDLAKAISEDNLRASNLGFQLYHGVHDGDD
VKCTPDEGLLLLDLNIRGGYDHRSAADLKMNQHMR
>SitPSR_Seita.2G118400.1
MERLSTNQLYSSGVPVTVPTSLPSIPASLEESFPRLPDSQNVLIERELRSTPVPPHQNTV
APIRGQFHTSTGSVGPLCSPPAVRFSSVSNPEQYSNPSPYNSQAPSTASSSTLNYGSQYG
GFEPSLTDFPRDVGPTWCPDPVDSILGYSGDVPGGNNLTGSTSLGASDDLTKQTEWWTEL
MNDDWKDIVDNPASAETQQVGPPAQSSISVHQSAAQQTVSSQSGEPMAVVAPSPTGGSNT
AKARMRWTPELHERFVDAVNQLGGSEKATPKGVLKLMKADNLTIYHVKSHLQKYRTARYR
PELSEGSSEKKAASKEDIPSIDLKGSFDLTEALRLQLELQKRLHEQLEIQRSLQLRIEEQ
GKCLQMMLEQQCIPGAEKATDALTSAEGSKISSEVPESSTAKEVPETSQNGLTKQTESGD
TR
>SitPSR_Seita.2G168800.1
MALLTNQIFKSHEEGGEREREQGGREESLPVESGRSSREARTSKKGERESRGTFPWIGSF
AFAAEEGGEEGPVSPLSAAAESSRSQEGAEEEEMFPPGLIHHRPDGPAPGDGAPRSGPGG
GPGGPSLVLTADPKPRLRWTADLHERFVDAVAQLGGPEKATPKTILRTMGVKGLTLFHLK
SHLQKYRLGKQSGKEGSEQSKDASYLLDAQSGMSVSPRVAAQDVKESQEVKEALRAQMEV
QRRLHEQVEVQKRVQIRMEALQKYIDSILESACKMVTEQFASSGFSISDPDLPEISPGGV
MCGPTDTLSSSVFNQLSVSSIDSHSPGGKPSPSGIEGPPMLLQKSPELKRRSS
>SitPSR_Seita.2G168900.1
MYSPKPEASFGSTHTNSVANQQQMELAGNNMGPSNGANNNNNLAARQRLRWTNELHERFV
EAVTQLGGPDRATPKGVLRIMGVPGLTIYHVKSHLQKYRLAKYIPDASTDGNKADNKDPG
DLLAGLEGSSGLPISEALKLQMEVQKRLHEQLEVQRQLQLRIEAQGKYLQKIIEEQQRLT
GVKSETPAAGASVTVSSDQFPDSERTEPSTPAPTSESPTQVGASNRDTGERAEATKSTCH
GDSLSRHEPLTPDSNCQNGSPPASPNHERAAKRQRGSGNEFVDVETDFSHPRHIFESSLG
PEFEQYSMSYSGH
>SitPSR_Seita.2G436900.1
MYHQQQLHGHNQHLSSRPGLPPEKQFLLQGGDAGLVLSTDAKPRLKWTPELHERFVEAVQ
QLGGPDKATPKTIMRLMGIPGLTLYHLKSHLQKYRLSKNLQAQANATNAKNVLGCRTGTD
KPCERNGSPASHLNTEPQINRSMHISEALQMQIEVQRRLHEQLEVQRHLQLRIEAQGKYL
QSVLEKAQEALAKQNVDLDAGVLGAAAAAETQQLSELISRASATKCAQHEHFHHQHLGGV
GDGSVDSCLTACEGSQRDHDMLSIGLSPAPTPRGGGYPFEAARSSGNERGGASTSCCEEF
LFLEESGTGRRGSTDEQQELDLNINDRNTRRPRNCEKIDLNGSSWN
>SitPSR_Seita.3G029000.1
MMNTKKIKLHDHHFGSPLYDPQMFPATAAAGLSFHPGLVGSLPQQHGGAGGWLQEEYSPT
PRTVLATQGSCVGSDPAAFFAAEHLLGMARFDCTLGSTALPAMTATKTAAPFVRSPEAEQ
LYRPLDPLLLRDGSVRTYYVRPQQRDATEAPPPLKLPLQQQQERGHGLYGNGSTGRLLGG
GEPKAPSFSPHATANTLIQAMESPGMQSPIENTLSRSCSIGAPASHTGNVVAAAGHGAPS
KTRIRWTQDLHERFVECVNQLGGADKATPKGILKLMNSDGLTIYHIKSHLQKYRIAKYMP
ASTSSEGKQEKRAAGNDAQNLDPSTGSQITEALRVQVDVQRRLHEQLEIQRNLQMRIEAQ
GKKLQKMFEEQLKASRTVMEPREELQLQDAGGIGAAAFPGVSEQEGEDAFDDVQLLSVAS
SGYNDARFPSKIS
>SitPSR_Seita.3G195600.1
MTLEIRSTGLTLTGVNGKEHARYTVRLDPVRLPLSGPARAPGRPDPRGPVREAPAPPETA
GSGKQRLRWTSDLHNRFVDAIAQLGGPDRATPKGVLTVMGVPGITIYHVKSHLQVQRQLQ
LRIEAQGRYLQMIIEEQQKLGGSIKASEDQKLLHSPPSLDEYPESTQPSPKKPRMDALSP

DSERDTIQPEFESHLIGPWDQEICGKNICGVAFPVEEFKADPGMSKS
>SitPSR_Seita.3G199300.1
MYHQHQGPSELFTTRTSFPMEQHLFLRGGNAQGDSGLVLSTDAKPRLKWTPELHQRFVDA
VNQLGGAEKATPKTVMRLMGIPGLTLYHLKSHLQKYRLSKNLQAQANVSTSKNAIGCTNI
ADRMPGTSAPTMSSTNAIPQAEKTIQIGEALQMQIEVQRQLNEQLEVQRHLQLRIEAQGK
YLQAVLEQAQETLGKQNLGPANLEDAKIKISELVSQVSNECFSNAITDIKESSSVHRLEP
KQIQFVESSTNNYLTAAEGFIKEHRLHHHGVLKAHDDSSLFCRKGSHEHETPLALNRSLS
ERRMAHLQIENGYSKAELGYENDTEMAPEYIGPPKNGGGSTSSSASGSKGDAEKPYLEEP
NCTRQAVEYPRESKLLDFGHSCPGKKLDLNTHNVDDTDQAYRHFDLNGFSWS
>SitPSR_Seita.4G185800.1
MFPSSKKQPNTGAGSSNDRSMCVQGDSGGLVLTTDPKPRLRWTAELHDRFVDAVAQLGGP
DKATPKTIMRVMGVKGLTLYHLKSHLQKFRLGKQHKEFGDHTAMEMQRNVASSSGVMGRT
MNDRSVNVNEALRIQMEVQRRLHGELEVQKHLQMRVEAQGKYMQSILEKAHQALGTSDYA
TWPAGYRSLGNQAVLDIGSSTGFSSLQDLHFYGGSSHMDHLLHQMERPMDSFLTLGENFI
GSSSADKKGPNHCSSSGKSSMIWAGEEDQQVKSGTDQLQMGSSTTMEGGISVMDPITSLY
EGALSGESMGSKGFEGSSSKLEMKSPPHNKHL
>SitPSR_Seita.4G270600.1
MSTQSVATGEQIIAPNETVHACTSTQTSVLQLFDSKSDHRLLIDDTLSSTSQSSSIKTEL
IRSSSLSRSLSVNLQKRSPETDPESPLSHISHPKFSDPILSNSSTFCTSLFSSSSKNTDP
CRQMGTLPFLPHPPKCEQQVSAGQSSSSSLLFAGDTGNALDEAEHSDDLKDFLNLSGDAS
DGSFHGETNALAFDEQMEFQFLSEQLGIAITDNEESPHLDDIYGTPPQLSSLPVSSCSNQ
SIQNLGSPVKVQLSSSRSSSVSATTNKSRLRWTLELHERFVEAVNKLEGPEKATPKGVLK
LMKVEGLTIYHVKSHLQKYRLAKYLPETKEDEKASSEDKKAQSGSSSSDSSKTKNLQVAE
ALRMQMEVQKQLHEQLEVQRQLQLRIEEHARYLQKILEEQQKAGNLSLKAPTKAQAVSPE
STASKERSETEAGTSSPRPSKNRNLDAHSECKSPAVSKRTEFQVDPESEVPCS
>SitPSR_Seita.6G120200.1
MFPGLIHHAGRGGRPDAAGAEDSPRGGGLGLHGHGGGPSVVLTADPKPRLRWTADLHDRF
VDAVSQLGGPDKATPKAIMRTMGVKGLTLFHLKSHLQKYRLGRQSGKELTEQSKDASYLM
EAQSGTNSSPRGSTPDVKESQELKEALRAQMEVQRRLHEQVEVQKHMQIRMEANQKYIDA
ILDKAFKIVSEQLSGISISDRDLPDLASAGVMFSPADPLSPSVFHQLSVSAVSLLSPGGG
KALPHVAIDISQKPPELKRKSR
>SitPSR_Seita.6G121500.1
MYQPNPSIGPTQSNPAPHDQMGPGDGAMVPHNGGNSNPNMAARQRLRWTNELHDRFVEAV
TQLGGPDRATPKGVLRIMGVPGLTIYHVKSHLQKYRLAKYIPDPSTDDNKAEKKDPGDLL
AALEGSSTMQISEALKLQMEVQKRLHEQLEVQRQLQLRIEAQGKYLQKIIEEQQRISGAG
ASRDTSSEQLPDSERTNPSTPVPTSESPLQAVPFSKDNGSRVEPMESASHDDLPHGEPVT
PDSNCRPGSPTLSPKHERAAKRQRGSSDGTPIADGDFALPHHIFESSTDSEFQQCSMPYS
SH
>SitPSR_Seita.6G158900.1
MSSSGGGQQQQGEGRPRLRWTSQLHGRFELAVAQLGGADKATPKSVLRAMAVPELTLYHL
KSHLQKYRLAVSRGLITTSPGDNGEGANDRSSSSENEYDEDAVAELHGAFTADDGAGAKE
GLCDSSRSMARMHREVQRKLQEQIEVQRHLQLRIEAQGRYLQSVLHRAYEVLSDDHNLGS
PAATELSELASAVESGCLSSSSSLSPSPPRRRAAGSCVTSSSSWEAESHAAAGSKRPCTC
AVEQPAQGKRTFLQQSHDHGAEEADADAEAEDGSSSEIDLNR
>SitPSR_Seita.7G223500.1
MMEGRYGGGGGGAMSRDPKPRLRWTPDLHQRFVDAVDKLGGPDKATPKSVLRLMGMKGLT
LYHLKSHLQKYRLGKQGKKSTGLELANGGGFAAPGLSFPTPTPIPGVPAEGKNTGEMPLA
DALRYQIQVQRKLQEQLEVQKKLQMRIEAQGKYLKAILEKAERNISSDVNAPSDNIESTR
SQLMDFNLALSGFMDNATRVCEENNEQLVKALSDDKHKDNNLGFQLYQVGCQGAKEVKWT
PKTEDLLQLDLNIKGGYDLSSRGMQACEVDLKINQQMI
>SitPSR_Seita.9G416300.1
MRNFNLRQSQNSRVLGAMSSSLPILPNSLKESFPRPCTPQHIPMSRQLPDDSMPLHNGTP
QSDTLQPRTGVIGASYSGYSANPLDSVSNHEAQSMVSPFICQPSNVDVFQSLSDNTPGAH
AEATWFPSSMDVLPVYTDNIAASDNQIQSGSSAMTSDEVAKQNDWWAEIMNDDWKDILDA

TATDSQSKAMMQPSNSIASLLAVNQSASSHSGEICPVASPPNSSNASAAKQRMRWTPELH
ECFVDAVNQLGGSEKATPKGVLKLMKVDGLTIYHVKSHLQKYRTARYKPDLSEGTSEKRT
TTEELSLDLKTSMDLTEALRLQMEVQKRLHEQLEIQRKLQLRIEEQGKYLQMMFEKQCKS
TTEVVQDPSSGDTSANPSSDPSHSANKDSSAAMDRNRIGDRPGTAELGERSTKLGVKQKI
SEIDSDPEAAADGGPKISLEKRRKLQDS
>SitPSR_Seita.9G419000.1
MYHHQQQLQSHSHFLSSRQTSPPERHLLLQGGSIPAEPGLVLSTDAKPRLKWTPELHERF
VDAVNQLGGPDKATPKTIMRLMGVPGLTLYHLKSHLQKYRLSKNVHAQANGGNAKNMVGC
TMAMEKPPEGNSSPASHINLGTQTNKSVHIGEALQMQIEVQRRLHEQLEVQRHLQLRIEA
QGKYLQSVLEKAQETLAKQNAGSVGLETAKMQLSELVSKVSTECLQHAFTGFEEMEGSQM
AQGHTMQLGDGSVDSCLTACDGSQKDQDILSISLSAHRGKEIAGMAFDIQAKERGSEDLF
LDKLSRTPPSHQERRERDSFIMAAKLDLNINDTNDGPQNCKKFDLNGFNWT
>PhePSR_PH01000008G1310
MSLSDHPHQHGEARPRLKWTRQLHERFVLAVSELGGADRATPKSVLRAMGVPGLTLYHLK
SHLQKYRLVVSRDLAGNGGGLNDRSSSSESPPNEYDDANTTELHTFVYAGDADADNGDDD
TKEALCDSSRSMAQMQREVQRKLQEQIEVQRHLQLRIEAQGKYLQSVLRRAQQVLADHNL
ASSPEATKAELSELVSEVETECLSSPSPPRHRSTDSCVTSSSSSEAESKAAGSKRLYTSM
HCLDCTVEQPVQGKRTFPFLQRHEAEQEAEAEAEDGSSSVIDLNR
>PhePSR_PH01000095G1450
MCVQGDSGLVLTTDPKPRLRWTVELHERFVDAVTQLGGPDKATPKTIMRVMGVKGLTLYH
LKSHLQKFRLGKQPHKEFNDHSIKDAAAMEMQRNAASSSSMMGRSMNDRNVHMNESIRMQ
MEVQRRLHEQLEVQKHLQMRIEAQGKYMQSILEKAYQAACPAGYKSVGNQAILDVCSIKD
IGPSMGFHSLQDLHMYGGGHLDLQQQMERPMESFFACNESSIGSMGKKRPSPYAAAGKSP
MMWGDDGDQDKSDQLQMAPPMMDGSIDVMGSIADVYEAKPIMSGDSTGSKGFKGPGSKLE
RPSPRRPQIGGERMGSPSVIGAQTRNLSYG
>PhePSR_PH01000123G0060
MFPSKKATSTAVSSNDRPMCVQGDSGLVLTTDPKPRLRWTVELHERFVDAVTQLGGPDKA
TPKTIMRVMGVKGLTLYHLKSHLQKFRLGKQPHKEFNDHSVKDAAAMEMQRNAASSSGIM
GRSMNDRNVHVNDALRMQMEVQRRLHEQLEVQKHLQMRIEAQGRYMQSILKKAYQTLASG
DGAACPAGYKSLGNQAVLDVCSIEDIGPSMGFPSLQDLHMYGGGHLDLQQQMERPMESFF
ACNESSIGSMGKKRPSPYAAAGKSPMMWGDDGDQAKSDHLQMAPPMMDGGIDVMDSMADV
YEAKPLMSGDSTGSKGFDGPGSKLERPSLRRPQIGGETRNFSYG
>PhePSR_PH01000152G0240
MEGSYGGGAGAVALSRDPKPRLRWTPDLHERFVDAVTKLGGPDKATPKSVLRMMRMKGLT
LYHLKSHLQVQKKLQMRIEAQGRYLKAILEKAQKNISFDANGAANLESTRSQLMDFNLAL
SGFMDNATRVYEENSEQLVKAITDDNLKANNLGFQLNHVGNQEAKDVKCTPKTEDSFLLD
LNIRGGYDLSST
>PhePSR_PH01000164G0530
MSSHSVVAVKQVAAPDKTVHACTSTQPSVHKLLDTKLDHQGLMDASQSSNIKTELIQSSS
LPKSLPFNLQKRSPESDPEKPIFSNSSTFCTSLFSSSSTNSVPCRQMGTLPFLPHPPKCE
QQVSAGQSSSSSLLLNGDIGNDHDESEQSDDLKDFLNLSGDASHGSFHGEDNASAFAEQM
EFQFLSEQLGIAITDNEESPRLDDIYGAPPQLSSLPVSSCANQGLQNIGSPVKVQLSSSQ
SSSGSATTNKARLRWTMELNERFVEAVNKLEGPEKATPKGVLKLMKVEGLTIYHFYHSKL
AYSMYYVDKKASSEDKKAQSSSSGNDSGKTKNLQVAEALRMQMEVQKQLHDQLEVQRQLQ
LRIEEHARYLQKILEEQQKTSNSLSLKNPTEAQAESPESASNED
>PhePSR_PH01000176G0870
MSSQSVVAVKQIAAPDKTVLGCTSTQPSVHKLFDAKLDHQGLMDDNLSSTTQSSNIKNEL
IRLSSLPKSLPFNVQKRSPESDPESPPSQISHPKFSDPIFSNSSTFCTSLFSSSSTNSEP
CRQMGTLPFLPHPPKCEQQVSAGQSSSSSLLLNGDIGNAHDEAEHSDDFEDFLNLSGDAS
DGSFHGENNALAFAEQMEFQLLSEQLGVAVTDNEESPRLDDIYGTPLQLSSLPVSSCSNQ
GLQNIGSPVKVQLSSSRSSSGSATTNKARLRWTLELHERFMDAVNKLEGPEKATPKGVLK
LMKVEGLTIYHVKSHLQKYRLAKYLPETKEDKKASSEDKKAHSSSSENDSGKKKNLQVAE
ALRMQMEVQKQLHEQLEVQRQLQLRIEEHARYLQRILEEQQKARSSVSLKNPTDAQAGLP
ESTSKEKTDSEAATTSPQPSKNRAPDADADAECSSPVDKKRAKVHADLDFLVRASAPSLP

VEVVAPLGPGVPAVVLARRRRLPQRRRAPAEPAALDQDRVHLFLGAARETYGFRWGRTDL
GPRGFERGFRGPEAVDVVDARWSHGGMGKIGKRGWGGCVRMISGRKW
>PhePSR_PH01000225G0780
MEGSYGGRAEAVALSRDPKPRLRWTPDLHERFVDAVTKLGGPDKATPKSVLRLMGMKGLT
LYHLKSHLQKYRMGRQTMKGSGPELANSGGFTAQGISFSTAAPPCVPAGENNTGEIPLAD
ALRYQIQVQQKLHEQLEVQKKLQMRIEAQGRYLKAILVKAQKNISFDANGAANLDSTRSQ
LMDFNLALSGFMDNATRGYEENNEQMVKAISDDNLKSNSLGFQFNHVGSQEAKDVKCTPK
TEDSLLLDLNIRGGYDLSSRGMPGCELDL
>PhePSR_PH01000261G1090
MAKSDAEPKTQDGSTLSGAFTAANSLSWALNYRWRFKSHDVLLQKVESNVKQKMYHQHQG
PSELFAARTTFPMERHLFLRGGNTQGDSGLVLSTDAKPRLKWTPELHQRFVDAVNQLGGA
EKATPKTVMRLMGIPGLTLYHLKSHLQKYRLSKNLQAQANVGTTKNTIGCEIVADRIPGI
SAPAMSNTNVIPQEEKTIQIGEALQMQIEVQRQLNEQLEVQRHLQLRIEAQGKYLQAVLE
QAQETLGKQNSGPANPEDAKIKISEPASQVSNECFSNTMSETKESTSMPRLETRQIQFVE
SSTNNCLTSAEGFIKEHRLQNHGMLKAYVDSSLFCRKQSHDHESQFSLNRSLSERRMGLG
HLHNVKEYHKAEFGSESDTEIPHEYIRPQKNSGGSTTSSASGNKERDAEKLYLEEPTCKR
QAIEYPRESKLLDFEHPSSGKKLDLNTHNIDDSDQGYRHFDLNGFSWS
>PhePSR_PH01000468G0380
MVHGRLQMAWLRSTAAFTYRGRLHYYHPLFSSHMVWDMKVFRKLFIVFEQIFMCRMLGQS
IKITYALVMFVNVIKANSCIRLTCFNRWRYLPLFGPMYQWILSKDFLVFMAVFIKAWGVG
TNDGSRTNIGNPNQCRPTANVALSFHNMYHHQQQLQSHSQFLSSRQTFPPERHLLLQGGT
IPAESGLVLSTDAKPRLKWTPELHDRFVEAVNQLGGPDKATPKTIMRHMAVPGLTLYHLK
SHLQKYRLSKNLHAQANVGNARNAVDCAITAEKPSEGNGSPVSHLNLGTQTNKSMHIGEA
LQMQIEVQRRLHEQLEVQRHLQLRIEAQGKYLQSVLEKAQETLAKQNAGSVGLETAKMQL
SELVSKVSTECLHHAFTGFMEIEGSPMLQGHTMQLGDGSVDSCLTACEGSQKDQDILSIS
LSAKKGKEIGGMAFDPQVKERGSEYLFLDKLSRRPPNHQERRDRRDGFSMSCQATKLDLN
INDTNDGPQHCKKFDLNGLSWI
>PhePSR_PH01000542G0010
MVQPSNLTASQPAISQSASSHSGEICPVASPPSNASAAKQRMRWTPEHHECFVDAVNQLG
GSEKATPKGVLKLMKVDGLTIYHVKSHLQVQQKKMSSTDELSLDLKTSMDLTEALRLQME
VQKRLHEQLEIQRKLQLRIEEQGKYLQMMFEKQCKSSTEKVQDPSSVDTAAPSSDPSHSA
SKDSDAALDPNRTGDNPGIAELGPNSTPVGGNQKTSDYGLQTYKQYYMIRLRITDI
>PhePSR_PH01000542G0130
MSMVQPSNLTASQPAISQSASSHSGEICPVASPPSNASAAKQRMRWTPEHHECFVDAVNQ
LGGSEKATPKGVLKLMKVDGLTIYHVKSHLQKYRTARYKPDLSEGSTEKMSSTDELSLDL
KTSMDLTEALRLQMEVQKRLHEQLEIQRKLQLRIEEQGKYLQMMFEKQCKSSTEKVQDPS
SADTAAPSSDPSHSASKDTDAALDPHRTGDNPGTAELGPNSTPVGGNQKTSETESSDSLA
TANDGSKLPQEKRRRVHSS
>PhePSR_PH01000548G0730
MSTESVIAVKQFTSESMAHTCTASQPSVHELFNAKYDHCGSTNDTPSSCTTQSSDIKTEL
IRSSSMPRILPFELQKCSTDSNPESSLSRISKADLADPILSKFSTFCTSLYSSSSTNSKS
CRQTSASPFLPHPPKCEQQQNSAGQSSCPSSLLFGADLSNGGHGDAEYSDDLKDYLNLSG
DGSFHGERNAMTFNEHMEFQFLSEQLVIAVTNNEESPRLDDIYEYDRPPQHSSLPVSSFS
DQEGLQNAGSPVKIQLSSSRSSSGAAACSKTRLRWTLELHERFVEAVNKLEGPEKATPKG
VLKLMKVEGLTVYHVKSHLQKYRHAKYLPETKEDKKSSSEYKKAQSATNVNGPGKKKSFQ
MAEALRMQMEVQKQLHEQLEVQRQLQLRIEEQGRYLQKILEEQQKARNSLSSMRSSTGES
PESTLKGKSKIKADTSSAPSSKHKISDIDMECYSQVDNQKAKLQVDL
>PhePSR_PH01000810G0130
MAQFDCPLGEALPVMQTAKTPFRSSENELYRPLDPLLLRDHSVRTYYVRPQQRNAAQAPP
SLKLPLPQQQQQQQERVHGLFSNTSTTRLLGGEPETHSFSPHVLMMDASCVLGYLKTLKV
ASPILPVMEATSLQSQMENQLSRSCIGAPVVTPTGNVSAPAGPEATPKGILRLMNSDGLT
IYHIKSHLQKYRSAKYMPASSEGKQREKRATGNDIQNLDPKTGMQITEALRAQLDVQRHL
HEQLEIQRNLQLRIEEQGKKLQKMFEDQLKVSRNVMESSPELDMVFPSEQDDAFDDVQLL
SVASGSYDDAGFPSKMS

>PhePSR_PH01001079G0490
MEKITTNQHCSSGVPVTVPSPLPAIPAALEESFPRLPDAHNQTTFAPLHGQFHSSTGSVG
PLSSPPAIRFSSVSNPEQYTSANPYNSQAPSTGCSSTLNYGSQYGGLEPSLTNFPRDVGP
TWCPDPVESMLGYSDDVPGGNNLTESSSIAATDELAKQTEWWTDFMNYDWQDVVDNTAGT
EPQSQVGPPAQSSIAVHQSVTQQTVSSQSGEPSAVAAPSPTAGSNTSKTRMRWTPELHER
FVDAVNLLGGSEKATPKGVLKLMKADNLTIYHVKSHLQKYRTARYRPELSEGSSEKKVAS
EEDIPSIDLKGSFDLTEALRLQLELQKRLHEQLEGGRVGADAFVIL
>PhePSR_PH01001153G0050
MEHRRRQESFFPVAGHTTSLRAPVASLIARISSSSSSRPKLIRRPISFSDSDIAESSLDQ
SARVTQNFDEGPKSQDLLGLVPWTPRRSDWTSWKDCPRDPLDLLGGPGGLLGRTTVMLQR
WLQSSHGYSLLDYLTQPTFLWPGSSEQVANLPKGIKENSMLIPFLGFVQSPEGIKELKNG
HLHFSSMQNFAQFFFWMPAVKRRKFSSSALSTDIPCPTHRREAQKCSGETGIGSQQLKDI
IQESNKRRGRQMYHQQQQQLQSHSQHLSSRSSLPPEKHFLLQGGGGDSGLALSTDAKPRL
KWTNEQHERFVEAVNQLGGPDKATPKTIMRLMGIPGLTLYQLKSHLQVQRHLQLRIEAQG
KYLQSMLEKAQEALEKQNVAGTTVDLEAAKTQLSELVSKASAKRLQNERSIIYQQRGDGS
IDCCLTSCEGSQRGDHGILSIGLSAANQSGTPFETARSGKDRGTISTSCEEYYLFPGKPS
RGASSDEHQERREIRRDDGFSNMTLQTTAELDLNINDGNCRPRNREKIDLNGSGRN
>PhePSR_PH01001295G0130
MERAGYGVSAGVVLSRDPKPRLRWTPDLHERFVEAVTKLGGPDKATPKSVLRLMGMKGLT
LYHLKSHLQKYRLGKQNKKDTGLETSRGAFAAQGITFSSPVPPNIPSAGNNSMGNALMTR
KRHRTNLQHRLVCRSTLAEENAVLNLYTKRCLLILYWTTNDLEDEVQKKLQMRIEAQGKY
LQAILERAQKNLSYDASGTVNLEATRSQLTDFNLALSGFMDNVTQACEQNNGQLAKAISE
DNLRNSNLGFQLYHGVQEADDVKCTPDEGLLLLDLNIKGGYDHLSSSGMRGGEADLKISQ
HRR
>PhePSR_PH01001459G0190
MPSSLPILPKPLKENFPRPHNPQHMPMPRQLQNDSMPLHQSSPQSATLHPRAGVVRSSYS
ATLGYSASPHDFVSNHEGQSIVAPFFSQSSNVEVFQSLSNNTPGGHAEATWFPASVDGLP
DYTDNMTAPDNQIPSGSSNMTSDEIAKQNEWWAEIMDDDWKDILDATATDSQSKSMVQPS
NLAASQPAVNQSASSHSGEICPVASPPNSNNASAAKQRMRWTPELHECFVDAVNQLGGSE
KATPKGVLKLMKVDGLTIYHVKSHLQKYRTARYKPDLSEGATEKRSTTDELSLDLKTCIC
QIPPLLVLDSSRCILSPLCVHFYVGSEIIVLRLDYMRKSMDLTEALRLQMEVQKRLHEQL
EIQRKLQLRIEEQGKYLQMMFEKQCKSSTDKVQDPSSGDTATPSSDPIHSASKDSDAALD
PSRTGDNPGIAELGQNSTHVGGNQKTSETESSDSLATANDGLFALEAQKLDIEISEPYWV
LQFPELLVYVVLNMNRWCLVFSPAWAPNQNSSSLTGLASQLLGEFVRVERARHGDLVVLG
VVVALDVLLL
>PhePSR_PH01001536G0490
MTYHAKKFSVPFAPQRAQSSEHASNIGAFSGSNISNPSNPVGSGKQRLRWTSDLHNRFVD
AIAQLGGPDRATPKGVLTVMGVPGITIYHVKSHLQKYRLAKYIPESPAEGSKDEKKDSSD
SLSITDSAPGLQINEALKMQMEVQKRLHEQLEVQRQLQLRIEAQGKYLQMIIEEQQKLGG
SFKTCEEKKLSESPRSLDDYPDSAQPSPKKPRMDDLSPDLVRDVAQPEFESHLIGPWDHE
LCGKNICGAAFRVDEFKADPGMSKS
>PhePSR_PH01001779G0420
MGVEASQTQKMAGPQQPHALYSLAHSCCLALYPSPLYPYMLLHKRKGKTKREGERERGGG
IYLSSGVQGKEPTEPPPLRVAAASHAVHATDPAAIPRTVVGSLEMSSSDHPHQHGEARPR
LRWTRQLHERFVLAVSELGGADGATPKSVLRAMGVPGLTLYHLKSHLQKYRLAMSRDLTA
SLAGNGGLNDRSSPLESPPNEYDDDSTTELHTFVCADDGDDDTKEALCDSSRSMAQMQRE
VQRKLQEQIEVQRHLQLRIEAQGRYLQSVLRRAQQVLADHSPASSPEAAKAELSELVSAV
ETECLSSSPSPPRRRHRSTDSCVTSSSSEAESMAAGSKRLYTSMHCRDCTVEQLVQGKMS
FPFLQRHEAEQEEAAEAEDGSSAEIDLNR
>PhePSR_PH01001931G0070
MQFDDVTVATKRSLQQQIPIHEATPKTIMRTMGVKGLTLFHLKSHLQKYRLGKQSGKEMT
EQSKDASYLMEAQSGTNLSPRVPTPDVKESQEVKEALRAQMEVQRRLHEQVEVQKHVQIR
MEAYQKYIDTLLEKACKIVSEQLSGFSISDHDLPDLASAGVICDPADPLSPSVFHQLSVS
SISLQSPGGKASPSALDGQLFFQKPPELKPKSC

>PhePSR_PH01002141G0040
MERYFSKRNRSSDNDAGTSRPPLRPRQDASNAAATPASLPNPTIPREINLDELPFDPADR
KRISQYSRNPKKQDEIRRIYLTRGPYRPQCGNYEQRVIGDAPRRFNPEWFEEYGGWLEYR
QEIALRDRLGMMHLQLMLSVAMMNQRSQKNKGNFRELLDTLANQNDAIRNVVLRNAPENC
QWVCPDIQKEIANYFAKITLNSILEEIGGDVFSLLIDEASDVSDKEQMAVVLRYLEKRAF
IVERLVGVVHVKETSAICLNSALQKLFTDIGLSIKQVRGQCYDGASNMREFDDRFNEVNS
ALLGHMASFNPKDSFAAFDLESLKKLAEFYPDDFDSNKLKDLGHQLRIYINNVRADKRFS
NLDGIADLAKLMVETKKHLTFHLVYRLLKLVLILPVATASVERCFSTMNVVKNMLRNKMG
DSGLVLTTDPKPRLRWTVELHDRFVDAVTQLGGPDKATPKTIMRIMGVRGLTLYHLKSHL
QKFRLGKQHKEFNDHSVKDASSSAMMGRSMNDVHMNEALRMQMEVQRRLHEQLEVQKHLQ
MRIEAQGKYMQSILEKAHQALASGDCGTWPGGYKSLGNQAVIDIGSSISFPSLQDLHLYE
GRHLELQQQMERPMDSFLAFNESCTGSVGKKSPNHYSSTGKSPMVWAGEEQAKISADQLQ
MASSMMMET
>PhePSR_PH01002430G0200
MYHHQQQLQSNSQLLSSRQTFPPERHLLLQGGTIPAESGLVLSTDAKPRLKWTPELHDRF
VEAVNQLGGPDKATPKTIMRFMGVPGLTLYHLKSHLQKYRLSKNLHAQANVGNARNVVAC
TIATDKPSEGIGPPVSHLNLGTQTNKSMPIGEALQMQIEVQRRLHEQLEVQRHLQLRIEA
QGKYLQSVLEKAQETLAKQNAGSVGLETAKMQLSELVSKVSTECLHNAFTGFEEIEGSPM
LQGHTMQLGDGSVDSCLTACESSQKDQDILSISLSAKKGKEIRGMAFDPQVKERGHEDLF
LEKLSRRPPNHQERRERRDGFSMSCQTAKLDLNINDTNDGPQHCKKFDLNGFSWA
>PhePSR_PH01002891G0030
MELGVNDMGPNNGANNNPNLAARQRLRWTNELHERFVEAVTQLGGPDRATPKGVLRIMGE
PGLTIYHVKSHLQKYRLAKYIPDSSTDANRTDNNDPGDLVAGLEGSSRLQISEALKLQRE
VQNRLHQQLEVQRQLQLRIEAQGKYLKKIIEEQQRLSGVKSETPAAGASVTVSSDRFPDS
ERTDPSTPAPISESPTQGVPSNRDNGGRAEATKSPCHDDSLSLREPLTPDSDCQPGSPTV
SPKHERAAKRQRGSGTEFSEADFALPHHILEPSSESELQQCSMSYSGH
>PhePSR_PH01003413G0110
MFHSSKKPGTSTVSSNDRPMCVQGDSGLVLTTDPKPRLRWTVELHDRFVDAITQLGGPDK
ATPKTIMRVMGVKGLTLYHLKSHLQKFRLGKQHKEFNDHSVNDAMEMQQNAASSSSMMGR
SMNERDRSVHMNEALRMQMEVQRRLHEQLEVQKHLQMRTEAQAKYMQSILEKAYQALASD
DCATWPGGYKSLGNQAVLDIGSSMSSPPQDLQLYGGSHLELQQQMERPMDSFLAFNESCI
GSVGKKSPSHYSCIGKSPIVWAGEEQVKISADQLQIAPSMMMEAGDIDVMDPIADVYEAK
QIMSGDSMGSKGFENPSSKLEMRSSPQRAPVGSERINSPMVGAQARNLSYG
>PhePSR_PH01005938G0020
MERITTNQHYSSGVPVTVPSPLPAIPASLDESIPRLPDSHNQTTFAPLHGQFHSSTGSVG
PLSSPPAIRFSSVSNPEQYASANPYNSQAPSTGSSSTLNYGAQYGGLEPSLTNFPRDVGP
TWCLDPVESMLGYTDDVPVGNNLTESSSIAATDELAKQTEWWTDFVDWKDVVDNTAGTET
QSQVGPPAQSSVSVHQSATQQIVLSQSGEPSAVAAPSPTAGSSTSKTRMRWTPELHERFV
DAINILGGSEKATPKGVLKLMKADNLTIYHVKSHLQKYRTARYRPELSEGSSEKVASKED
IPSIDLKGSFDLTEALRLQLELQKRLHEQLEIQRSLQLRIEEQGKCLQMMIEQQCIPGTD
KVLDASTSAEGSKLSSDPPESSTVKDAPENSQNGITKQAESGDS

>BdiPSR_Bradi1g17400.1
MYHQQQHQLHTHSQHLSSRPSLSPEKKFLRQGQGRGDSGLILSTDAKPRLKWTSELHERF
VEAVNQLGGPDKATPKTIMRVMGIPGLTLYHLKSHLQKFRLGKNLQTQAAVVNVKNVLGF
VTATDKACEGHGSPADHLNRETGTSKSMHISETLQMQIEVQRRLHEQIEVQRHLQLRIEA
QGKYLHSVLEKAQEALAKNQHAAGAGAGHEAGKPPARQRLQRNDGSADGSCLTAASDEGI
LSIGLSSAAAAGARRAAVPPLETPARSSSGRDRGPEEEGEEEEEECFLFLGKPEHHEERI
KLGLSIGGGERIDLNGSSWKD
>BdiPSR_Bradi1g28920.4
MERITTNPFYSPGIPVTVPAPLPSIPASLDESFPRHPDVQNVLMERELRRTPLPPHQSTV
APISGQFHPSAGSVGPLCSPQDVRFSSVLIPEQYTSANPYNPQTQSTGSSSALIYGSQHG
GFEPTFTDFPRDVEPAWCPDPVESMLGYSDDVSGGNSLNGMSPIAATDELAKQTEWWTDF
MNDDWKDIVDNPSGAKSQPQGGPPVQSSTSVHQSATEQIVTTQSVEPCAVAAPSPSASSN

TSKTRMRWTPELHERFVDAVNLLGGSEKATPKGVLKLMKADNLTIYHVKSHLQKYRTARY
RPELSEGSSERLDASKEELPSIDLKGNFDLTEALRLQLELQKRLHEQLEVQRSLQLRIEE
QGKCLQIMIEQQCVPGTDKVRDASTSAEGSKLFSDPPESSTLKDIPNNSQNGTTKQAESG
DNE
>BdiPSR_Bradi1g31837.1
MFPAAAVCSKKPGAVVSSSPNDRPCVVQGQQGGDSGLVLTTDPKPRLRWTVELHDRFVDA
VAQLGGPDKATPKTIMRVMGVKGLTLYHLKSHLQKFRLGKQHKEFGDHSSVKEAMEMQRN
AASSSGMMGRSMNDRSAHMNEALRMQVEVQRRLHEQLEVQKHLQLRVEAQGKYMQSILEK
AYQTLASGGDCATWPAAGYRSLGGASMDVGSSMSFQDLTLYGSGSSHLDLQQQMEIRPTM
APMDSFLAFNESCIGRRSPADAGGSCYGRAAGKSPMMMMWAGDDQQAKSCGTDGDDQLLQ
MAPASTMMMEAGGVDAMDPIMSMSGDSLGSNKGFDRGPELQA
>BdiPSR_Bradi1g34470.1
MSSQAVVAVKPTTAPDKTVHSYACGSTQSSVHKLPDAKLDHLRFLDDNLSSTSQSSNIKT
ELIRSSSLPFRLQKRSPLSDPESPLSHVSQPNFSDPMASNSSTFCTSLFSSSSTNSAPCQ
QKGALPFLPPPPKCEQQVIPGQSCSSSLHLTGDIGNAYDEAEHSDDMKDFLNLSGDASDA
SYHGEDNAMAFTEQMEFQFLSEQLGIAITDNEESPRLDDIYDTPPPQLSSLPVSSCSNQS
LHNLGSPVKLPLCSARSSSGSTTTNKSRLRWTLELHESFVEAVNKLEGPEKATPKGVLKL
MKVEGLTIYHVKSHLQKYRHARYLPDMKEDKKASLDCKKVQSAQSGSNGSYLDKNKNLAE
ALRMQMEVQKQLHEQLEVQRQLQLRIEEHAKYLHRILEEQQKASNGGSSSLKISTEPTES
TSINGTAPEEATTSSPHPSKNIAAPEAGTECDSPVRNKRAKVHGDLESESPCS
>BdiPSR_Bradi1g36236.2
MRGCQMYEPKPLSSTGSAHNNPIPHNQQIVPTANNAASNIGGNSSNINFATRQRLRWTDE
LHGRFLDALTQLGGPDRATPKGILRTMGVQGLTICHVKSHLQKYRLSKYIPDPTADGAKS
DKKELGNLLAGIESSPGMELSEALKLQMEVQKRLRDQLEVQRQLQLRIEAQGKYLQKIME
EQQRLTGVLCESGTLNALVPVQELRQDFNKTDPLTPVPTSEPLIRDKASTVSDDHEGTDG
LLKDLSSHDECHSSGREPLTPDSSCGAASPLDIPRDRLDHQNSESVLHHNILESRSGSDF
QQASSVFPSSTRRSDSPAALYVSEDDFKNSPDV
>BdiPSR_Bradi1g63530.1
MRKCDLRHSHNNRVSGAMSSSLPILPNSLKENFPRSHNPQLIPMPRQLMNDSVPLHHSAP
QSATLHPRAGVMRSSYSASLGFSASPADSVPNHESQSMSAPFISQPLDIELFQTLSDNIP
GGHTEATWFPGSVDGLTDYGDNVGAPGNQIRNGCPAVTSDVVAKQNEWWEEIMDDDWRGI
LDATTTDSQSKAMIQTSNSAVSQPAVNQSASSHSGEMCNVASPPNGNNVSAAKQRMRWTP
ELHECFVDAVNKLGGSEKATPKGVLKLMKVDSLTIYHVKSHLQKYRTARYKPDLSEGTTE
KRTSTEELTLDLKSSMDLTEALRLQMEVQKRLHEQLETQRKLQLRIEEQGKYLQMMFEKQ
SKSSTENVQDLSGNTAAPSSDLSHSVNRNRDSEAAEDPNRTGDNPGSVKSGEYSTHTSGN
QRTAERDSSDPLANTNDGLKAPQEKRRRVHDS
>BdiPSR_Bradi1g63690.1
MYHHQQQLQRHSQLLSSRQTFPSERHLLLQGGIVPGESGLVLSTDAKPRLKWTPELHDRF
VEAVNQLGGPDKATPKTIMRLMGVPGLTLYHLKSHLQKYRLSKNLHAQANVGNSRNVVGC
TMATEKHSEGNGSPVSHHLGAQTNKSMHIGEALQMQIEVQRRLHEQLEVQRHLQLRIEAQ
GKYLQSVLEKAHETLAKQNTGSASLENAKMQLSELVSKVSTECLHNAFTGFEEIQGSQML
QTMQLGDGSVDSCLTACESQRDQDILSISLSAKKGKEIGAMAFDLHMKEGHGNLFLEKLS
RRPPNHQEGHERTDGFSISCQTTKLDLNINETNDGPQNCKKFDLNGFSWT
>BdiPSR_Bradi2g21407.2
MMYHAKKFSVPFAPQRAQNSEHVSNIGAFGGSNISNPANPVGSGKQRLRWTSDLHNRFVD
AIAQLGGPDRATPKGVLTVMGVPGITIYHVKSHLQKYRLAKYIPESPAEGSKDEKKDSSD
SLSNTDSAPGLQINEALKMQMEVQKRLHEQLEVQRQLQLRIEAQGKYLQMIIEEQQKLGD
SLEGSEERKLSHSPPSLDDYPDSMHPSPKKPRMEDLSPDLARGVTQPRFESHLIGPWDQE
LCGKNISDPAFQVDEFKANPGLS
>BdiPSR_Bradi2g21880.1
MYHQHQGPSELFSTRTTFPMERHLFLHGGSTQDSGLVLSTDAKPRLKWTPELHQRFVDAV
NQLGGAEKATPKTVMRLMGIPGLTLYHLKSHLQKYRLSKNLQAQVNVGTTKNGCAVVADS
MPATSTPAMTNTNVIPQAEKTIQIGEALQMQIEVQRQLNEQLEVQRHLQLRIEAQGKYLQ
SVLEQAQESLGKQNLGPANLEDAKIKISELVSQVSNECFSNAVTDIKESSRMHRLEPRQI

QFVESSTNNCLTAAEGYINEHRLHSHGVLKAYDDSSILYRKQSHGHEYQFPLNRSLSERR
MGHLHNVKEYHKAELGSESDTEIQQEYITPQKNVGGSTTSSASGTKEGDIKKLYLEEPSC
KRRAMDYPSFERPNSGKKLDLNTHNTDDSDQGFRHFDLNGFSWS
>BdiPSR_Bradi3g03538.2
MSTQSVIAVKQPPVQHLHSMESQWNLNDAHTCSAKQPPVHRMFSVISDDCASTSDARSSC
TTQFSTIKTELIRSSSMTKILPFELQKCSTTDFNPGRSFSHVSQTDLSDPILSSSSTFCT
SLYSSLSTNSKCRETGALPFLPHPRKLEQQHSSAGQSPNSSLLLAADPSNSGHGDAEHSD
DLKDFLNLSGDASGSGFHEGISNCMDFNEQMEFQFLSEQLGIAITDNEEIPRLDDIYGRP
QQPLALPLSSSSDQEDGRDAGSPVKVQLSSSSSSGAAGCNKTRMRWTLELHERFVEALKK
LGGPEKATPKGVLKLMKVEGLTIYHVKSHLQKYRLAKYIPEKKEEKKPSSEDKKAQSTAD
GIDPAKKKSLQMAEALRMQIEVQKQLHEQLEVQRELQLRIEEHARYLQLILEQQKVRKCP
SSMKSSMEGESSGSKPKEKTEMRAETPSAPSPKRKFPDIDIEHNQQTDN
>BdiPSR_Bradi3g05500.2
MFSSKKGGGSSGGGGAAVQVQGSNNSSVAAAAGMCVQGDSGLVLTTDPKPRLRWTVELHE
RFVDAVAQLGGPDKATPKTIMRVMGVKGLTLYHLKSHLQKFRLGKQPHKDFNDHAVKDAA
AAMEMHRNAASSSGMMGRNMNDRNVHMNEAIRMQMEVQRRLHEQLEVQKHLQMRIEAQGK
YMQSILEKAYQSLGSGEPAAGYKSLGGVLDVCSIKDIGAASMGFPSLQDLHLYGDLQQNQ
PIESFFACSDGGIGPPLGKMKRSSAGHYTAGGKSPMMWGSDDDGDQDKDDQLLQMAPPMM
MGEDMVDSIAAGVHEAAKPMMSHAMSGGGESKLERPSPRRPHLGGGQRMGSPSVIYG
>BdiPSR_Bradi3g22290.2
MYQPNQISSIGHNHGSPAAHEQMELGGTSMSMVPCNGGNDNPNMASRQRLRWTNELHDRF
VEAVTQLGGPDRATPKGVLKIMGVPGLTIYHVKSHLQKYRLAKYIPDPSASDDNKAEERD
PGDLLAALEGSSGMPISEALKLQMEVQKRLHEQLEVQRQLQLRIEAQGKYLQKIIEEQQR
ITAAGPSRATSSDQMPDSERTNPSTPGLTSESRVHGSTKNNKNQIELTKSSPNDDPLACR
QPLTPDCSRPSSPTLSPEHERPVKRQRGSDPVDDTSFDDDQFVLPRTIFESSTATEFQDC
SSPYSGH
>BdiPSR_Bradi3g22297.1
MFPGLIHHHHQHAADDAPRGHGGGSAPSLVLTADPKPRLRWTADLHDRFVDAIAQLGGPD
KATPKTILRTMGVKGLTLFHLKSHLQKYRLGKQSGKEITEQSKDGSYLMEAQSGINLSPR
IPIPDVEESQEVKEALREQMEVQRRLHEQVKVQECVKIRREAHQTYIDSLLEKACMLVSE
QLSGFSISDYDLPDLASAGFQIP
>BdiPSR_Bradi3g36710.2
MSSPSAQQHGGEAATARARLRWTRPLHERFVLAVSELGGADRATPKSVLRAMGVQGLTLY
HLKSHLQKYRLAVSRDLAGNAGGGSLNVDRSSSSESQSNEYNDDDDTTAELRDSSRSMAQ
MQREVQRKLHEQIEVQRTLQLRMEAQGRYLQSVLRRAQQVLTDNSLAKAELSELVSAVDT
ECLSSSSSQPRQQHRSAYSSCVSSSSSEAESKSAGSRRRLHAGRHQGDCCTVEQPAQGKR
TFPFLQQMQDAEQQAAAEAEPEEEEEAEDGSSSEFDLNM
>BdiPSR_Bradi3g52500.2
MMFEGMERAGYGVGAGGVVLSRDPKPRLRWTPDLHERFVEAVTKLGGPDKATPKSVLRLM
GMKGLTLYHLKSHLQKYRMGKQSKKDTGFETSRAAFATHGISFSSATPPVVPSAGNNNMG
ETPLADALRYQIEVQRKLHEQLEVQKKLQMRIEAQGKYLQTILEKAQKNLTYDSSAATNL
EATRSQLTDFNLALSGFMDDATQVCEQNNGELAKVISEDNLRAGNLGFQLYHGVQDAEDV
KCTADEDLLLLDLNIKGGYDHRLSSHGMRRGDADLTVGQHRRWTGVKIWYGHGG
>BdiPSR_Bradi3g55120.1
MYTAILDTSTMSQFRDGEVPWYLSSDITVLPALPEVMVPSFPQYSASYIERGLCRSSITP
LNTFLPASSGLQSTYMSASNFTSDLCINNGLPNEKLSSGSHIIAISGFDTSLSSTHSSYK
DDPSSLRMLYPKVSEEIYWVQEPVPGVFDYPASLNVSDQRNLVVSQEMQDIITLDHDTHL
AKQKEWFSSESSGKFLENSGCGGSVLKAVDTRSTAPPNYTYFHMQNNVSSHFNVDELCSD
NFPSSDTAPTKSRMRWTTELHELFVGAIIKLGGSEKATPKAVQKIMKVEGLTIYHFSFIR
FHLCTNLEGYLCLHNFKHISKYPKGGVMPWVCARDLEDMDTSEGLRTQIGLQKQLHEQLE
IQRKLQLQVEEHSKYLEMAIAKQGESLKQLGALPVFENSRTQVLDHIKACEDQTVDFSGR
EALRN
>BdiPSR_Bradi4g27620.1
MELGGNNMGSNDGANSKASLAARQRLRWTDELHEQFVEAVTQLGGPDRATPKGVLRIMGT

PGLTIYHVKSHLQKYRLAKYIPDSSTDGNKSDNKDPGDSLAGLDGSSGLQISEALKLQME
VQKRLHEQLEVQRQLQLRIEAQGKYLKKIIEEQQRYGGIKSETPGAGGTATVSSDQFPDS
ERTDPSTPAPTSESSQGVPFKRDNGGQTEATKSPCHDEQLTTDSNCHPGSPTVSPKHERA
AKRQRGNGTEFSEADLSLPQHIFESSSGPEFQQCSVPYSGH
>BdiPSR_Bradi5g20520.1
MFEGSYGGGCAAGAEAALSRDPKQRLRWTPELHRRFVDAVAKLGGPDKATPKSVLRLMGI
KGLTLFHLKSHLQKYRMGRQTKKATDLELASSGGFAAGDISFSIGTPRLVPAGDDNREIS
PTDTLRYQIQVQRKLHEQLEVQKKLHARIEAQGRYLKAILEKAKKNISVDINGSPNIEST
RSQFMDFNLDLLGLMDNGTQMYEENSEQLMKAISDNNLKDNNLDFQLYDVGSQEAKNVRC
TPRTEDLLLLDLNIKGGHDLSSTGMQ
>BdiPSR_Bradi5g25320.1
MMDARKMKLHGRHQYCSGMMPAPAPRLMAPAASAYGSAGGFHHGFSSPAALHHQQQIQQQ
QQHGGGWLQDPDQYAAVAPSCVVGSDTAMFYAAEKLLGMPQLDCCPPLRMLPPHMPLELD
HPAVMTYYVRPQQRRGDGADLPLTPQQQQGEGVQLNHGLYGNGSAIKPHSFVPAAMDLQA
PSGSLQMGQMTDSSHGHMPRSCVGAPASHTSNGSLAAPAPAPSKTRIRWTQELHERFVDC
VSKLGGADRATPKGILKLMNSDGLTIYHIKSHLQKYRTVKCVPSSSSSSEGKQQEKRAAG
SDDVPNLDPKTGMHITEALRVQLDVQRRLHEQLEIQRKLQVRIEEQGKRLQEMFEEQLKA
SGNAAAAAPGSPEPGCAASDDVVIFPVSDDEDEDDDVQLLSVASSSYDEDLLAL

>OsaPSR_LOC_Os02g04640.1
MSTQSVIAVKQFSGPDKIAQAYTVPQPSAHVLSNANYDYDLCGSTNSTSLSCAIQSSNIK
TESISSSSLPKILPFSTDSNGESSLSRMSQAEFSDPILSSSSTFCTSLYTSSPMNSGSCR
KTGYLPFLPQPPKCEQQQNSAGQSSSSLMLLDADLRNSGHADDEHTDDLKDFLNLSSDCS
FHGKCSAMAYNEQMEFQFLSEQLGIAISNNEESPRLDDIYDRPPQLMSLPVSSCSDQEDL
QDARSPAKVQLSSSRSSSGTASCNKPRLRWTPELHERFVDAVNKLEGPEKATPKGVLKLM
KVEGLTIYHIKSHLQKYRLAKYLPETKEDKKQEEKKTKSVANGNDHAKKKSAQMAEALRM
QMEVQKQLHEQLEVQRQLQLRIEEHARYLQKILEEQQKARESISSMTSTTEGESPEFAPM
EKTEDKAETSSAPLSKCRITDTDAECHSKVDNKKTKPQADLEMVHDE
>OsaPSR_LOC_Os02g07770.1
MCVQGDSGLVLTTDPKPRLRWTVELHERFVDAVTQLGGPDKATPKTIMRVMGVKGLTLYH
LKSHLQKFRLGKQPHKEFSEHSVKEAAAMEMQRNAASSSGIMGRSMNHDRNVNDAIRMQM
EVQRRLHEQLEVQKHLQMRIEAQGKYMQSILEKAYQTLAAGDVAAAVACGPAGYKSLGNH
QAAVLDVCSMGFPSLQDLHMYGGAGGGHLDLQQQQPPASTMESFFACGDGGGSLGKTAAK
TRHYGGAGKSPMMWGVDDDDDDDDPAGKCGGGGHHQLQMAPPPMMDGGIDVMDSLAADVY
ETKPIMSGDSTGSKGGGYDVAAAASKLERPSPRRPPQLGSPSVMAGAQTRNLSYG
>OsaPSR_LOC_Os02g47190.1
MFEGMERAGYGVGVGGAGAVGAGVVLSRDPKPRLRWTPDLHERFVEAVTKLGGPDKATPK
SVLRLMGMKGLTLYHLKSHLQKYRLGKQNKKDTGLEASRGAFAAHGISFASAAPPTIPSA
ENNNAGETPLADALRYQIEVQRKLHEQLEVQKKLQMRIEAQGKYLQTILEKAQNNLSYDA
TGTANLEATRTQLTDFNLALSGFMNNVSQVCEQNNGELAKAISEDNLRTTNLGFQLYHGI
QDSDDVKCSQDEGLLLLDLNIKGGGYDHLSSNAMRGGESGLKISQHRR
>OsaPSR_LOC_Os03g20900.1
MYHHQQQLQSHNQLLPSRQSFPSERHLLMQGGSVSGESGLVLSTDAKPRLKWTPELHERF
VEAVNQLGGPEKATPKTIMRLMGVPGLTLYHLKSHLQKYRLSKNLHAQANAGNVKNALVC
TTATEKPSEANGSPVSHLNLGTQTNKSVHIGEALQMQIEVQRRLHEQLEVQRHLQLRIEA
QGKYLQSVLEKAQETLAKQNAGSVGLETAKMELSELVSKVSTECLQHAFSGFEIESSQML
QGHTMHLGDGSVDSCLTACDGSQKDQDILSISLSAQKGKEIGCMSFDMHVKERGSEDLFL
DKLNRRPSNHPERCERRGGFSMSCQTANLDLNMNDTYDGPKHCKKFDLNGFSWA
>OsaPSR_LOC_Os03g21240.1
MSSSLPILPKSLKDIPRSHNTQNILMPGQLPNDSMPLHQSATQSSISHPRASVVRSSYSA
MLGYAANPIDSVSSHEGHFMAAPFISQSSNAEMLQYLCNNNTHGGHTVPTFFPAPACGAP
DYMDTITVPDNHTQSGSSTVTSDAAKQNEWWADIMNDDWKDILDATATDSQSKSMAQPSN
SAASQPAFNQSTSSHSGDICPVTSPPPNNSNASASKQRMRWTPELHESFVHAVNKLGGSE
KATPKGVLKLMKVDGLTIYHVKSHLQKYRTARYKPDLSEGKTQEGKTTDELSLDLKASMD

LTEALRLQMEVQKRLHEQLEIQRKLQLRIEEQGKYLQKMFEKQCKSSTQSVQDPSSGDTA
TPSEPSNSVDKDSEAALDPNRIGDNHPKNSTNVGANLKTAATESPDSPVIATDGSELPQE
KRRRVHES
>OsaPSR_LOC_Os04g56990.1
MLQDIMNTKKIKLHDCHFGSPLCDPSPAPHLLSSAAAAGLSFHPGLVSSAAQHQQHGAGG
WLHEEYYAPRSSPPSSLLAQTCVGSNATAFYAAENLPQFDFPALGTAAAAAAKAPFRSSE
SELYRPVDPLLLRADHSVRTYYVRPQKRDSGERTPLPPPSQQQHQDRIHGLFAGAPTTRL
LSGEPKIHSFPPQVAAKPILPAMDAPSLQNQMENQLTRNCIGAATPVTPTGNLAGSGAPS
KTRIRWTQDLHERFVDCVNQLGGADKATPKGILKLMNSDGLTIYHIKSHLQKYRIAKYMP
ASSEGKQLEKRATGNDMQNLDPKTGMQITEALRVQLDVQRRLHEQLEIQRNLQLRIEEQG
KRLQKMFEDQLKASRSVMEPQELDDVVAFAAGDGDDDAFDDVDVQLLAVAGSGYDDAGFQ
SKIS
>OsaPSR_LOC_Os05g40960.1
MYHQHQGRSDLFTTRTSFPMERHLFLHGGNTQGDSGLVLSTDAKPRLKWTPELHQRFVDA
VNQLGGAEKATPKTVMRLMGIPGLTLYHLKSHLQKYRLSKNLQGQANVGTTKNALGCTGV
ADRIPGTSALAMASASAIPQAEKTIQIGEALQMQIEVQRQLNEQLEVQRHLQLRIEAQGK
YLQAVLEQAQETLGKQNLGPASLEDAKIKISELVSQVSNECLSNAVTEIRESSSIHRLEP
RQIQFVESSANNCLTAAEGFKEHRLQNHGVLKAYDDSTLFCRKQSQDQESQYSLNRSLSE
RRMGHLYSGKQYHKSEGSDSDTEVLHEYITPQKNGGGSTTSSTSGSKEINVEKLYLDEPS
CKRQTVDYQRESKLLDFDQQSSGKNLDLNTHNIDDNDQGYRHFDLNGFSWS
>OsaPSR_LOC_Os05g41240.1
MMYHAKKFSVPFGPQSTQSNEHMSNIGAFGGSNMGSPANPAGSGKQRLRWTSDLHNRFVD
AIAQLGGPDRATPKGVLTVMGVPGITIYHVKSHLQKYRLAKYIPESPAEGSKDEKKDSSD
SLSNTDSAPGMQINEALKMQMEVQKRLHEQLEVQRQLQLRIEAQGKYLQMIIEEQQKLGG
SLKACEEQKLPHSPPSLDDYPDSMQPSPKKPKMDNLSPDSVRDVTQSDFESHLIGPWDQE
AAFRVDEFKADPGLNKS
>OsaPSR_LOC_Os06g40710.1
MYEPKPFSSIVLAHNDPVSHNQQIERINNNVVSNSGGNSSNSNFAARQRLRWTDDLHDRF
VDAVTQLGGPDRATPKGILRIMGVQGLTIYHVKSHLQKYRLAKYIPDPTADGAKSDKKDL
GDLLADIESSSGMEIGEALKLQMEVQKRLHEQLEVQRQLQLRIEAQGRYLQKIIEEQQRL
SGVLGESGKLGALGPAPGEPYQDSNKTDPSTPVPTSESPIRDKAGSGLFKTISSHDDCRE
PLTPDSSCRAGSPLESPPRASKRIRVSSDIDHRGNNEFPPPLKVPEPSSGSDFRQESSVL
LSSSAVHFDSLESLDADENVFTNGSGSDD
>OsaPSR_LOC_Os06g45410.1
MFPSSSPSPSPSSSKKQQQLSGGVGVGGAAAAASSNDRPPLCVQGDSGLVLTTDPKPRLR
WTVELHDRFVDAVTQLGGPDKATPKTIMRVMGVKGLTLYHLKSHLQKFRLGKQHKDFNDH
SVKDAMDMQRNAASSSGIMGRSMNDRSVHVNEALRMKMEVQRRFHEQLEVQKHLQMRVEA
QGKYMQTILEKAYQAISSSGDCATWHAGYKSLGSQAVLDIGSSMSFPSLQDDLQLYGGSH
LDHLHQQHEQMEIRPSIDTFLAFNYSSSTGKSPMVWPGADDGGGEPAKISGDHQLQMAAP
ATTTMMMEAITMSGGDSMGSKGFEGQMSSKLDMRSPPPQQTVLPVGSERMSSPIVGAKAR
NISYG
>OsaPSR_LOC_Os06g49040.1
MSSQSVVAVKQITAPDKIVETCPSTKHSAHKLFDVKPDFQGLIDDNLSSSSQSSSIKIEL
IRSSSLPNILPFQKRSSEPEPESPLSHVSHPNVSEPVYSNSSTFCTSLFSSSSMETEPCR
QLGTLPFLPHPPKCEQQVSAGHSSSSSLLVPGGDGDIGNAHDEPEQSDDLKDFLNLSGGD
ASDGSFHGENNAMAFAEQMEFQFLSEQLGIAITDNEESPRLDDIYGTPPQLSSLPVSSCS
NQSVQKAGSPVKVQLSSPRSSSGSATTNKARLRWTLELHERFVEAVNKLDGPEKATPKGV
LKLMKVEGLTIYHVKSHLQKYRLAKYLPETKEDKKASSEDKKSQSGSSGNDSVKKKNLQV
AEALRMQMEVQKQLHEQLEVQRQLQLRIEEHARYLQRILEEQHKVSISSNSLSLKPPAES
QPESPKPTSEKKEAESEAGAATSAQPSSEDKSPDAECKSSPPVGSKRARVHVGDEHQCS
>OsaPSR_LOC_Os07g25710.1
MERISTNQLYNSGIPVTVPSPLPAIPATLDENIPRIPDGQNVPRERELRSTPMPPHQNQS
TVAPLHGHFQSSTGSVGPLRSSQAIRFSSVSSNEQYTNANPYNSQPPSSGSSSTLNYGSQ
YGGFEPSLTDFPRDAGPTWCPDPVDGLLGYTDDVPAGNNLTENSSIAAGDELAKQSEWWN

DFMNYDWKDIDNTACTETQPQVGPAAQSSVAVHQSAAQQSVSSQSGEPSAVAIPSPSGAS
NTSNSKTRMRWTPELHERFVDAVNLLGGSEKATPKGVLKLMKADNLTIYHVKSHLQKYRT
ARYRPELSEGSSEKKAASKEDIPSIDLKGGNFDLTEALRLQLELQKRLHEQLEIQRSLQL
RIEEQGKCLQMMLEQQCIPGTDKAVDASTSAEGTKPSSDLPESSAVKDVPENSQNGIAKQ
TESGDR
>OsaPSR_LOC_Os07g48596.1
MTGTEQMMYHQQQVQSDSQHLSSRPGLPPEKQFLLQGGADSSSGLVLSTDAKPRLKWTSE
LHERFVEAVNQLGGPDKATPKTIMRLMGIPGLTLYHLKSHLQKYRLSKNLQSQANASRAQ
GVLGCSTTEIDKPCEGNGSPASHLDLETQTNSSMHINEALQMQIEVQRRLHEQLEVQRHL
QLRIEAQGKYLQSVLEKAQEALGTIAVAETAATANASSSKRLQNEHTQLHHHQQQQVGD
GSVDSCLTACDCEGSHHSRSHGHRGEQDILSIGLPPFEPAAARSGKEHHYLLFPNEPSRR
RSCSDERRREMSTLQASELDLSINGRSSSHSHRRENIDLNGAGWS
>OsaPSR_LOC_Os08g25799.1
MFPGLIHHHRLLDADVGGGGGGSSAGLVLTADPKPRLRWTADLHDRFVDAVAQLGGPDKA
TPKTIMRTMGVKGLTLFHLKSHLQKYRLGKQSGKEMAEQSKDASYILGAQSGTNLSPTVP
TPDLKESQELKEALRAQMEVQRKLHEQVEVQRHVQIRMEAYQNYIDTLLEKACNIVSEQL
NGFSISDHDLTSAGVMLSSSDTLSPSIFHQLSVSSISLHSPGGKSSPFAADADLFFQKAP
EKRKSY
>OsaPSR_LOC_Os08g25820.1
MYQPNPISSSGQTHGNPTAHEQMELGNNAIVPSNGGNNNPNMAARQRLRWTNELHDRFVE
AVTQLGGPDRATPKGVLRIMGVPGLTIYHVKSHLQKYRLAKYIPDPSADDNKDEDKDPGN
LLSALEGSSGMQISEALKLQMEVQKRLHEQLEVQRQLQLRIEAQGKYLQKIIEEQQRVIG
AGASRATSSEQLPDSEKTNPPTPVPISESPVQGAPHSKNSQSQVEPTKSPSHDDALPCGE
PLTPDSSCRPGSPTLSPKHERAAKRQRGSDAGDVTAFADGEFVLPPGIFESSTGSEFQEC
SMPYSGH
>OsaPSR_LOC_Os08g33750.1
MSSQQQHAGEAPASAAAAARARLRWTGQLHERFVLAVAELGGADRATPKSVLRAMAVPGL
TLYHLKSHLQKYRQAVSRGGNGGGGGSGSLNDRSSSSERQPADHDGDSAADEPRTIAYDG
DSDGDAKEGLRDSSRSMVQMQREVQRKLQEQIEVKRHLQLRMEAQGRYLQSVLRRAQQVL
ADHSLASSPEAATAELSELASAVDIECMSSSSPPRHHRQSAATDSCVTTTSSSEAESKAA
GSKRLHTSDCTVEQPVQGKRAFNFLQRHNQADQEEEQEEEYAGAEDGSSSEIDLNR
>OsaPSR_LOC_Os09g12750.1
MFPPGLIHHRPDGGEAGRAAGGGPSLVLTADPKPRLRWTADLHERFVDAVAQLGGPEKAT
PKTILRTMGVKGLTLFHLKSHLQKYRLGKQSGKEASEQSKDASYLLDAQGGMSVSPRVST
QDVKENQEVKEALRAQMEMQRRLHEQVEVQKHVQIRMEAYQKYIDTLLEKACKIVSEQLA
SSGFSISDNDLPELSGGVMCGSADTLSSSIFHQLSVSPINLHSPEGKPTPSGIEGQMILQ
KSPELKRKSC
>OsaPSR_LOC_Os09g12770.1
MELGGNNMGPDNGANNNSNLAARQRLRWTNELHERFVEAVTQLGGPDRATPKGVLRIMGV
QGLTIYHVKSHLQKYRLAKYIPDSSADGNKAENKDPGDLLAGLEGSSGLQISEALKLQME
VQKRLHEQLEVQRQLQLRIEAQGKYLKKIIEEQQRLGGVKSETPAAGASVTLPSDQFPDS
ERTDPSTPAPTSESPTQGVPSNRDNGGQNEATKSPQRDDSLSRHEPLTPDSNCQPGSPTA
SPKHERAAKRQRGNGAEFSETDFALPHSIFESSSGSEFQQCSMSYSGH
>AcomPSR_Aco001094.1
MGSTEMPDQTGNSRLALPADAKPRVRWTRELHEQFLRAVSHLGGADIHSLISEATPKSVM
RMMGVPGLTLYHLKSHLQALFAAVKKKVQKHLQLRIEAQGKYLQSVLRKAHETLAAYSSS
SITELSELVSAAETLCPSSLLSQTIFNPNRAQQGDCSTDSCVTSEEFERENKMGCKIKSS
LSKGGDELTGFKRSCSRINGEYGDERPINAKRSLCETENGGFMLQEQIDLNS
>AcomPSR_Aco001685.1
MYSHFANQPRQKMYNNHHQGHNNNLLIPSRGAFPADKHLFLQGGSVQGESGLVLSTDAKP
RLKWTPELHERFIEAVNHLGGADKATPKTIMRLMGISGLTLYHLKSHLQKYRLSKNLQTQ
SSNGSTKNIVSCTVATDRTLEGNGAPISDLNIASQTKKTMQISEALQMQIEVQRRLHEQL
EVQRHLQLRIEAQGKYLQSVLEKAHETLAKQNLGSAPGLEAAKAQLSELVSKVSTECLHN
AFTFEDVHRHAAQFNDGSVDSCLTSCEGSQKEQELHTKCKRTAAEDINSQALWFRDSNKR

KIASASAARDQDMNDYHVQRDSGMLSIGLAAHDAKEMRGNVFEARRNDPCLTTQLDLNAN
YSSDAAPNCKKFDLNGISWT
>AcomPSR_Aco002890.1
MSNHNSNQGLVLSADAKPRLKWTQQLHDRFVDAVARLGGADKATPKSVMQVMGVPGLTLY
HLKSHLQKYRLAKSRDTNILHDYSEGEIQCKILERSTPDADENKAQNQLDETMLQMQMQV
QRKLQEQIEVQRHLQLRIEAQGKYLQNVLRKAQETLSAYSSNSIGTDVVKAELSELASAV
GNECLNSPPPQPCLVFNISQQSDCSIDSSLTSSEKPDKATEKSRRIGSSSSSCNGSEELQ
SRGINYDSGEAVGTKRSRSSICGEATLDEHTTWKRIFHQKECSFEEPEELDLNR
>AcomPSR_Aco007475.1
MRVMGVKGLTLYHLKSHLQKFRLGKQPHKEFNDHSVKDAANMSMQRNAPSSSSIMGRSMN
ENVHITEAIRMQMEVQRRLHEQLEVQKHLQMRIEAQGKYMQTILEKAYETLQGDSNIGLG
GYTKALGNNTQGVIDMVSMKEMASPMCFPSLQDLHLYGGDHLEIPQQIERPLDMFFPTNE
SIAMGSKKRPYPYGSNGKSPMIWADDLRLQELGSQEEASKCDQLQIAPSMMDSSIDMGSM
ANVYESKPMLDITGNKKFEGTSKLDRPSP
>AcomPSR_Aco008334.1
MFPTKKPTMNSSHERSMGVQGDSGLVLTTDPKPRLRWTVELHDRFVDAVAQLGGPDKATP
KTIMRVMGVKGLTLYHLKSHLQKFRLGKQPHKEFAEHSIKDAATQELQRNAASSSSIMGR
NTNRNLHTNNDALRMQMEVQRRLHEQVEVQKHLQMRIEAQGKYMQTMLEKALQTLAVEST
GSGDYKEIGNQGVIDMSSMKEIGSNYSASFPNSLYDLHSHGANYGLDMQYHTDRPLVDQF
FSTVDERICLGKKRPVNPHCRYRQNPMVGAWADNLGLQELGSFAACMAGSQEEASKNIDN
HLQVAPSTVIDVRIGMDSTRGAYEAKPILSSDGDVGGEKKFEYPTKLERPSPRRASLLIE
RLNQMIRGS
>AcomPSR_Aco010145.1
MSFCRSRGLSEAVHEGTDQPSVSNYPKESTELAWHPEPLQRVLDYPDNPNFGNNLSHGVN
VMASDEINKQSEWSDLMDFMNGNWGELQGDTNTTESKPEVALPADQAPTSFSVHHPQTHP
SVPSNSNDLSAVASPSSSATAAPIKARMRWTPELHERFVEAVNQLGGSESSTQAPCFDFT
EATPKGVLKLMKVEGLTIYHVKSHLQKYRTARYKPESSEGTSEQRVIQVEETKSLDLKTV
TEINEALRVQMEVQKQLHEQLEIQRKLQLQIEEQGRYLQMMFENQNKSIHKLKESASLEI
PIDRMEISEKDHSETENHLHDKGAAEEVSKNKMPKNDDGKI
>AcomPSR_Aco011622.1
MNSHSIVTVKQSNSPERTAHYCQAAPSKISKIFKGQQDHQSLSGNNSSSSGEFSYLLNPK
LSPESDSESPISHVSLPHYSEPIFSRSSMFCTSLFSSSSKSSDSCRRLSNLPFLPHPLKC
EQQTSAAQDSDSPFHFTSDINNTPNSEDEHSDELMKDFLNLSGDASDGSFHGENCDSNSL
AITEQMELQILSEQLGIAITDNEESPRLDDIYEKPQISPQPVSSCQNQNPQPSGSPIKVQ
LHSTPSASRATTNNNKPRLRWTLDLHEHFVEAVNKLDGPDKATPKGVLKLMNVEGLTIYH
VKSHLQKYRLAKYLPETKEDKKSLPVEDKKVTPVTNDSGGKKKNMEMTEALRMQIEVQKQ
LHEQLEIQRALQLRIEEHARYLQKILEEQQKASNSFASSMGVSAEAQVESPDKTSQDQDE
SKTDSVSSPTALKHRIPESDGDCKPPEDHKRARLQVEQ
>AcomPSR_Aco012613.1
MYHQHQGHSDLFSPRAAFPQDRHLFLQGGNVTGESGLVLSTDAKPRLKWTPELHQRFIEA
VNQLGGAEKATPKTVMRLMGIPGLTLYHLKSHLQKYRLSKNLQAQANIATTKNVIGCVAA
ADRTPGQSGLIVSNTSATCEAEKTMQISEALQMQIEVQRQLHEQLEVQRHLQLRIEAQGK
YLQAVLEKAQETLGKQNLTSSGLEAAKIKLAEQVPEVSNEFLNNGFPNSEEITNISILGA
QAIQLADCSRRDLVYQTVKFERDTNILPVHLKGQTENGGNKNNSQSKQKERENEKFYLEQ
QLNYKRPAVQHMAGNGSNDFGLACFGAQLDLNAHDNNDSDSSNKEFDLNGLSWS
>AcomPSR_Aco014587.1
MKKSATSQLYDIGGSGAMSSSLPALPTPSEVSFLKLPDSQQVSMERELSSSNPLPPHRAP
SHYNTAVVGPLYSSASGPLYSAPSPNVSSFISQPSNAGNFMGSANLSHTGIYHPSTSNYQ
GEPLESNWRPDSVQGILDYSDNVIASGNQIQSSPIMASDELIKQNEWWNDFINDDWKDIY
DANASEPQPKMLSLSKSLAQQTKSGSNQVVCPPSNLSMQQPAIPQSAPSNSGELCTVASP
SPTASAPAAKARMRWTPELHESFVEAVNQLGGSEKATPKGVLKLMKVDSLTIYHVKSHLQ
KYRTARYRPESSEGASEKKGTSQEELSSLDLKTSFDLTEALRLQMEVQKRLHEQLEIQRN
LQLRIEEQGRCLQMMFEKQYKSGVGKPPSTSADPLTLSSDQSHSDTNNRIQENAQAETND
IPSSSNLEAKSPRLVGEKKKKKAEAESFDQGENTTDGPKLKRARGQEIEPSSANSEP

>AcomPSR_Aco015674.1
MNTKRVAFSDRNYGSLNDSSHGTLNPKHCYAYYTPQQCNPNSFSSLLPTTGSNCVGSTPA
AFYAAEQLLGIPQFEYQIGDVPPPCRLPRRSSESGLSRRPDAYFCNNSVKNYDHGLHATN
ELLSVVKFPFQESRNSRALGNFDGFLCRSRPEDELQPILHRQEASRNTTVGCNASSSLAE
KPNQLVTNSAGSSSNVPSNLTVQNKTRIRWTQDLHERFVECVNRLGGAEKATPKGILNLM
KYEGLTIYHIKSHLQKYRIAKNIPEQTEGRNDRRTVDDIQQLDPKTGVDITEALRIQLDV
QRQLHEQLEVQRELQLRIEAQGRKLQQMFEQQMKTSRDIVENQNYDMLFPDDAQVTLDDA
QISVLGGGSSENSDFMSKID
>AcomPSR_Aco017021.1
MFEGMERSYPAAAAAATMYEAVEGSGGVVLSRDPKPRLRWTPDLHDRFVDAVTKLGGPEK
ATPKSVLRLMGMKGLTLYHLKSHLQKYRLGKQTKKETGGQANTGSNSSVNSYSCVTPSNV
SIGNNIGELPLAEALRYQIEVQRKLQEQLEVQKKLQMRIEAQGKYLQAILEKAHRSLPLD
VNGTRDFEAASAQLTDFNLALSGLMDNVNQVCEEKSPEPGKAMTQDSIERSNNNASTLLR
RNREEEVRDMKVNLNAVWGEGGGRCERSSERTKEAKKWIAEWIRGEFATTFPPSRVSLSP
SPSGRVRETLAVSLADPQFAESKLVRNPSLLISIEESGYVFSMSSGRAKNFRRRSEDSGV
NGDEKPGPTPTTTIASSSSSSTANRSQTLGSAAKPKSSKPSGPKRLSFADEEEDEDSEGG
GGVAVRRPPARAPSSAAPVHKLTAAKDRSKSSVPLPPIPSNVQPQTGEYTKERLRELQKN
ARPLGSMPKPQPTTAPSFSEPKSQKPEPASSEPVIVLKGLLKPTSVTATAGGTQRGVAKD
EDEEEEEEEEEEEERADGDEERADGERQRKLPVIPDRATIDAIRAKRQQMQQPRRLGPDY
ISLDGGGMLSSRPSAGGSSDEEDSDFQGRIALFGDKVGGKTKKGVFESIEEPVSISSKFQ
EVGINREEDEDDEEERKWEEEQFRKGLGRRLDETSSHRGATGVSTVVPVQPQPSIYSGIA
PSMGNVPIGASVLVTRSAEVMSIAQQAEVATRALQDNISKLKESHKITLNSLVRTDTHLS
EALSEMSDLEKSLEAADKKYVFMQQLREFISVMCDFLNDKAFYIEELEEQMQKLHEKRAL
AISDRRAFDFADAESEVEAAVSAAMSVLSKGSGPTYVSAATAAAQKAMAAARECPNLPVE
LDEFGRDINLRKRMDFTRRAKERKIKRAKAESKRMSSLGKGNKSEQIEGELSTDESETEM
CAYGFSRSELLDTAEQIFADASEEYSSLRIVKERFEGWKNKYSSTYRDAYVSLSVPAVFS
PYVRLELLKWDPLYKTTDFFDMEWHKLLFDYGKPGGDHDFELDDADVNLIPGLVEKIALP
ILHHEIAHCWDILSTRLTKNAVFATNMVISYIPGSSKALHELLAVVRNRLNEAISSLSVP
AWNTTVTKAVPGAAQFAAYRFGLSLRLLRNICLWKNILALPILEKLALEELLGGKLLPHL
KSIISDIHDAITRTERIVASLSGVWAGPEVKSEPSQKLRPLVDFVAELGSKLERRHASGA
SEEETRGLARRLKNMLVALNEYDKARAILKTFQLKEAL
>AcomPSR_Aco017080.1
MFSGLIHREASISSGEDAAAAAAGAGAAHGGPSLVLTADPKPRLRWTADLHERFVDAVAQ
LGGPENVVYFAEATPKTIMRTMGVKGLTLFHLKSHLQKYRLGKQSGKEMTEQSKDAAYLS
EAQSSANLSPRVATPDVNEYELGQEVKEALRAQMEVQRKLHEQVEVQKHVQIRMEAYKKY
IDSLLDKAYKIASEQIAMSSLNNSTEHDLHDLATRGVMCAPSDPLSPSIFHQLSVGSISL
HSPGGRTAPTSAIDGQFFFQKPPELKRKSY
>AcomPSR_Aco017082.1
MTSRDPCRHVAIFNSRLTQIRSSHSEATWIGSNLLILASILGSKHLLFGVRRSKFVLWIL
SVLFWGDFALFFDLEMYQPKPIAGMGPTHGNSIALNKQAELARNGVGSPNDASNSASAAS
RQRLRWTNELHERFVDAVAQLGGPDRATPKGVLRIMGVPGLTIYHVKSHLQKYRLAKYIP
EPSSDGSKPEKKDPGDLLSGIESSSGMQITEALKLQMEVQRQLQLRIEAQGKYLKKIIEE
QQRLSGVLATSEADNPSDPDKTEPSTPAPTSETPTAGTAPSGHFFSAAHEPPLTPDYPSK
DNSNDERASKRPRTGETADLVLAHQILESN
>AcomPSR_Aco019267.1
MEDLAVLKSLRVFAMYHPKKFSTLHMAPHKSQGSEQTGNVSVMDGSSLNNSTNSGGGGKQ
RLRWTSDLHDRFVNAITQLGGPDISILESWRLVEAANLAVTSSSKNDVTMPCNFWTDIGV
SLMHEISFVVVWKVSNLVLLRGATPKGVLRIMGVPGITIYHVKSHLQKYRLAKFLPESPAD
GSKDETKDSGDTLSGNDSAPEIQINEALKMQMEVQKRLNEQLEVQKQLQLRIEAQGRYLQ
KIIEEQQKLDGAVKSSEGNEGNKNASASSSNLEDCLGHTPFPSKKPRISNPPAEPTLENT
NPELKPDFIRQLEHEIFESSVGDFVLGEAIEFKGEGGSSEQLGKGLVDVTSAQYLEVSNL
RVDCGSRNQEIVVEQQLHHSCGTI
>AcomPSR_Aco023131.1
MGPDSGGHHPNNASMAARQRLRWTHDLHERFVEAVTQLGGPDRATPKGVLRLMGVPGLTI

YHVKSHLQKYRLAKYIPESSADGMKTEKKDPGDLLSGLESSSGMQITEALKLQMEVQKRL
HEQLEVQRQLQLRIEAQGKYLQKIIEEQQRLSGVLAEGPGLSLPSPPASGGDNLPDPDKT
DPSTPAPTSESPFQDKALLQDGLLKSQFSHDDDSFSSFREPLTPDSTCHAAGSPSASPDR
NPTHERASKRLRGAMAAGHGKNEVVLAHHILESSSGLDFQQHCSVFPAARGGSGGRFSSS
STS
>PdaPSR_PDK_30s1141531g006
MNSHERPMCVQGDSGLVLTTDPKPRLRWTVELHDRFVDAVTQLGGPDKATPKTIMRVMGV
QGLTLYHLKSHLQKFRLGKQPHKEINDHSIKDASALDMQRNAASSSGIMGRTMNENVHIT
EALSMQMEVQRRIQEQLEVQKHLQMRIEAQRKYMQTILEKACQTLAGEGMASVSYKGHGN
QGVIHVGAMKDMGSPMSFPSLQDLHLYGGDDQLDMQQHMDRPLDGFFQANEGVCLGKKRF
NPYSSNGKSPLICADDLRLQELGSAAACMGAQEEPSKNDQLQIASSVIGGGIDMDSITDV
YETKPVLSTDGTGEKKHEGSPKLERPSPRRAPLPVERINPMIRGGTLPQTRNISYG
>PdaPSR_PDK_30s6550950g002
MQIFETPILDILWKHAGFYGSIEIVFFSQFNCQELLDDKLSCASPSPHVQRELINSSSPE
KGLPFCLKKLCPESEPGSSLSNESHTQHFEHAFSGSSTFCASLYSSSSTSSESYRQLSNL
PFLPHPQKCKLENSVVQSLNSPLLHVGDTSNVHNEDEHSDDLMKDFLNLSGDASDSSFHG
ENYDNNSIALGEQMELQMLSEQLGIAITDNGESPRLDDIYETLAPSVPLSSNCNQTSQPL
TPPAKVQLHSKSSTSTTAAANKPRLRWTLDLHERFVEAVNKLDGAEKATPKGVLKLMNVE
GLTIYHVKSHLQKYRLAKYLPEAQEDKKASSSEDKKAPTISHESDLGNKRSTQVIEALRM
QIEVQKQLHEQLEVQRALQLRIEEHAGYLQKILEEQQKASNTFVFSMEMPDTEMQLESPH
HSSPHQAESKVDSICPPSSSNSKHKGTDSDTDSKPLEDHKRARLEVE
>PdaPSR_PDK_30s6550973g003
MYHHHLEGHSNLLASRTTFPPKKHLFLQGGSVPGESGLVLSTDAKPRLKWTPELHERFVE
AVHQLGGADKATPKTIMRLMGIPGLTLYHLKSHLQKYRLSKNLQSQANTGTTKNDKQVRD
RKYSRMLLINENLQLIVQRHLQLRIEAQGQYLQSVLEKAQETLGKQNLGSTGLEAAKVQL
SELVSKVSHECLNTAFPGLEEISGLHPLQAHAAQFTDCSVDSCLTSCEGSQKEQETNYVG
VGLSTYPGNSPLCLQQFRADTELERAQPAWHADFNAQKTFSPFIVRDSEGTVFPAQRGSR
TLSINMKAQREKVDSSTDSEARRKERDSKDTFLEASRKRPAVLQERGKEPNEFGLSSMTT
QLDLNAHEENDGLPNSKQFDLNGFSWS
>PdaPSR_PDK_30s6550989g009
MYHAKKFSTISMLPHKAQGTEQLANAGVMGGSNVSNPINSGGSGKQRLRWTSDLHDRFVD
AITQLGGPDRATPKGVLRVMGVPGITIYHVKSHLQKYRLAKYLPESPADGSKDEKKDSGD
SLSSMDSASGIQINEALKIQMEVQKRLHEQLEVQRQLQLRIEAQGKYLQKIIEEQQKLGG
ALKVTEESQKSSESPSASQDCLGGPSPLKKPRIADQSPDATHETPLPESKPDFIGQWNQE
FYESSVTHFAFDSRSEFKEDGNNCGQPEKHFVGANADRS
>PdaPSR_PDK_30s661191g015
MTNFNYESSGKSDAPSSIYFPTDCQAQNNDNVGILQSKSTSLEHACTANNSDLSWREVGG
SKHGEMLYSSNSRIGTTAVHSSMQSGGAATGKQRIRWTHELHEQFVEAVNHLGGAMKATP
KGILALMKSQGLTIFHIKSHLQKYRTAKCMPNFLGGKNGKRINNDMLPLSNFRGYAGLQI
SETLRLQMDVQKSLHDHLEFQKKLQMRIEEHANYLERMFDQQKRSCSFLENHNSANAVDQ
SIIGCSSNGEYENSKEAILAKREP
>PdaPSR_PDK_30s674801g004
MFSGVLHRPEASISQEEAHGPRLVLTADPKPRLRWTDLHERFVDAVTQLGGPEKATPKTI
MRTMGVKGLTLFHLKSHLQKYRLGKQSGKEMTDQSKDASYPLENPSSSGLSPRLPTPDVN
EGQEVKEALRAQMEVQRRLHEQVEVQKHVQIRMEAYHKYIDSLLEKACKIASEQIASSGF
NGTGHDLPDLATGVMCSPSDPLSPSVFHQLSLGAISLHSPGGKTSPSSAIEGQSFYPKAP
ELKRKLC
>PdaPSR_PDK_30s679311g001
MEGDTFFSSEALFAGQNRALGRDYETPFTANNTSDKNQSPPLSETGATPKGVLRVMGVQG
LTIYHVKSHLQKYRLAKYIPESSSDGTKSEKKEPGDLLSTIDNSSKSIDLSESYNELSKV
QRQLQLRIQAQGKYLKKIVEEQQRLSCVLTGASGLGNTATTSGNPNLDSDKTDPPTPAST
SDIPIQDKGIFSFTDENARINVLKKRV
>PdaPSR_PDK_30s681141g002
MERELRSTPLIPLRTPFVSNSAVVGPLYSSASGFSSDLHFSSMAPNERHHNGASFVSQSP

NVGIPFPSTPSHSGSFCPATSNHPRESADVSWCPEPIQGMLDYSDNITAGNNQIQSSCDV
ASDDLAKQNEWWTDLMNDDWKDILNETSASECQTKSVQPATQASPSISVHQLQIHQSVPS
HSGELCAVSNSSPAAIAAAAKPRMRWTPELHECFVDAVNQLGGSEKATPKGVLKLMKVES
LTIYHVKSHLQKYRTARYRPDSSEGASQKKVTPQEEVSSLDLKTVSNILMKTPKIREDPE
IGKSVTGCLNISMMQISFRSIDLTEALRLQMEVQKRLHEQLEIQRNLQLRIEEQGRYLQM
MFEKQSGISNLQASSSLEEPLTVSSDQTHSAAKIELPGKGRDESGIDPNGAKMSEGSRQV
GDKQKTLEAESCNEMEANIVSGSHSPSCKRARGPDGESLTPMSALD
>PdaPSR_PDK_30s695651g003
MNTRRINFHDRNDDSLNSSSFELLNPSCGNFTSQQAWNTGFPFQTTSLDCSPRQYAANSS
DSSLSSEKGSPQPDFMAPNRSITAPICVGSTPSDFYAAERLMGFPPLEYQFGTPPLLSKS
PGIHLEASHNRPSESYMCVNTVKQSDFGCQTGDALESITKFPLQESGTSRFSDNTNRFPC
RDHQEIELRSLLQSQQSDGYSTLDSRHLYSPQGIHDPCVGYNLSNASAAKPSFQPPTEKQ
LVKTSVGRPTTPTGVSSGTSVPNKTRIRWTQDLHERFVECINRLGGAEKATPKGILKLMN
LDGLTIYHVKSHLQKYRIAKYMPESSEGPSQIELNEAGNLMFPYFCSGMQITEALQLQLD
VQRRLHEQLEIQRNLQLRIEAQGRKLRQMFEQQLKANANLTESQNLENLFPDEQPISLDD
DVQILNVEDGSQNTNFPSKIS
>PdaPSR_PDK_30s714041g001
MGPGSGGTDPNNPNLASRQRLRWTHELHERFVDAVTQLGGPDRATPKGVLRIMGVQGLTI
YHVKSHLQKYRLAKYVPESSSDGSKGEKKDPGDLLSGLENSSGMQITEALKLQMEVQKRL
HEQLEVQRQLQLRIEAQGKYLKKIIEEQQRLNGVLAETPGVSIPAPISGDHCQDSDKTDP
STPAPTSESPRQDKAASGDHGGTGGLFKSLSRDDSFSSRREPLTPDSSCHAGSPESPKPE
RRIKRQRGAGDPGNGKMELVFAHHILESTAPSDHLFSSLQPQSTPSNCQLWCRI
>PdaPSR_PDK_30s714681g011
MYQPKPISNLVSAHSNPLVCDQPIELAGNNMGPCSGGTNQNNPNLASRQRLRWTHELHER
FVDAVTQLGGPERATPKGVLRTMGVTGLTIYHVKSHLQKYRLAKYIPESSSDGTKAAKKD
SGDLLSGLENSSGMQITEALKLQMEVQKRLHEQLEVQRQLQLRIEAQGKYLKKIIEEQQR
LGGVMAETPSASIPAPISGNQCLDSDKTDPSTPAPTSESPLQVKAASGDHGGSGGLFKSL
SHDDSFSSHHEPLTPDSSCRAGSSFESPKHERPIKRQREASDPGNGKTELVLVHHILESS
SGSDFQQPCSIFPARGGHFDSTGTSICDDDRLKNGSGSDV
>PdaPSR_PDK_30s714681g012
MQKEGWFATPTAFTSKEPLLATKLVAWSDGPLPFGPPLFPRKEMFSGVIHRSEASVPQEE
AHGLRLVLTADPKPRLRSVYYTFEGIAAYAEATPKTIMRTMGVKGLTLFHLKSHLQKYRL
GKQSGKEMTDQSKDASYPSETLSSSGLSPGVPAPDVNEGQEVKEALRAQMEVQRSLHEQL
EVQKHVQIRMEAYHKYIDSLLEKACKIASEQIAASGFNATGHDLPDLATGIICSPSDPLS
PSVFHQLSVGAISLHSPGGKTSPSSAIEGYLFYHQKAPELKRRPC
>PdaPSR_PDK_30s728141g003
MEKRRAETLLLFFIFSIAVVAGDWNILSYMAKKRDHHQVGISLKNYCESWRMNAELNNIR
CFDVVPGECVGYIGKYMTSTQYKVDVQRAAEEATLFLTNSFLLGGDGKDAWVFDIDDTLL
STVPYFKKHQFGGATTNRTSMEQWMEERSAPAVEHMLNLYHQIRARGLKVFLISSRREHL
RDATIDNLVKVGYLGWAELILRCKRHYKLDIDLGLGQEHRGSIALVDCAQLLKRVPLQLS
HLSLEENIFKMFCASDAKVTMERELRSDPMAPLHTPYVKNNGVAEPFFSTASGFSLDVHC
TSVPPYERQPVSASYISQPSSSALSLSSTYSSYSGTCQPSMSNYPKESAEITWYSDSLQG
VLDYSDNVNTVNDQIQSNCVMTSDNLIKQNEWPDLTDLMNGDWGEFLDNRDATEPQPKAS
DLHVVYPAAQASTDSSAYHSQIYESVHSHSGEQCAVGSLSSSANAAPAKQRMRWTPELHE
CFVEAVNKLGGSEKATPKGVLKLMNVEGLTIYHVKSHLQKYRTARYRPESSEGTSAKKIT
PLEEMPSLDLKTGIEITKALRLQMEVQKRLHEQLEPLKKSEAGHHYPFLQTIHIQRKLQL
QIEEQGKYLQMMFEKQGKAGSDKLKAPSNEEDPSSQYSDLSHNVTKDGISEKDRAETENG
PSGASILEECSRQVGDKQKIVEPDADMRSNTIRSHSSPHKRARSHDADVSSATSAFY
>PdaPSR_PDK_30s738571g007
MERGYGSAYEAAAGGGGGGVVLSRDPKPRLRWTPDLHDRFVDAVAKLGGPEKFWAVDHGH
MPKESQESYKLTFRWKALRDNLLTTKCFSKLRLEVINSIEASFGNEMPLAEALRYQIEVQ
RKLHEQLEVQKKLQMRIEAQGKYLQAILEKAQKSLSFDMNNSGSLEATRAQLTDFNLALS
GLMDNVNQVCEEKNEDLGKFISQDNLKRTQNSGFQLYQKQQEDREDIKLAPNGENHVFPH
HYNLITFSTPVQLCLRESESWYGSRLLILAIVGCSRNSQVSTTMGARVHDQSCPSTSTGS

ENLQLASTWPIRLQPKQNSGKRFRARMTELASPFPSECLLRAGRTSPLNPLSRFDLPFLP
GFPSPAQSD
>PdaPSR_PDK_30s771701g004
MYHHQHHQGPSNLLASRVSFPPERHLFLQGGSVPGESGLVLSTDAKPRLKWTPELHERFI
EAVNQLGGPDKATPKTVMRLMGIPGLTLYHLKSHLQKYRLSKSLQAQPNTGTTKNGCPLA
ADRTAEGNGSLMSNTTVASQTNKTMQISEALQMQIEVQRRLHEQLEVQRHLQLRIEAQGK
YLQSVLEKAQETLGKQNLGSAGLEAAKVQLSELVSKVSNECLNTTFPGLEEISGFHTLQA
HAAQFADCSVNSYLTSCEGSQKEQEKYNVSMGTYLGNSHPCLQQFRADTGLEQAQLAWHG
DLNEQKTFSPSILQDSEVTFFPVQRDSRTLTMNIKAEREKVDSSTVSDARRKEKDSKDAF
LERNSKRPAALQERGKESNGFGLQSVTTQLDLNAHEENDGPPNCKQFDLNGFSWS
>PdaPSR_PDK_30s785961g002
MYHQHQGKNDLFSSRAAFPPERHLYLQGGNAPGNSGLILSTDAKPRLKWTPELHQRFIEA
VNQLGGADKATPKTVMRLMGIPGLTLYHLKSHLQKYRLSKNLQAQVNTGSTKNGSVIAAD
RTSEKVCANDMFLDLRTMQISKALQMNIEAQRQLHEQLEVQRHLQLRIEAQGKYLQSVLE
KAQETLGKPSVGSVGLEAAKVQLSELASKVSKECFGTAFPNLKESSSLHTLQGQTTQFSD
YSGDSCLTSCEEFQKDRETHNVCVGLGTCPSNSPLCTQQIGKDPTLEQNQFMWCGNVNEH
RLLHSSTVKKNDMSIFPVEGESNIVTKNIKVQGENGCSISISEARRKGIDGEDPYLELPN
CKRAAVKQERGNQLNEFALPCLTSQLDLNSHDDNDPPPGCRQFDLNGFSWS
>PdaPSR_PDK_30s803301g004
METQRVKNFNVKHFRSTGNSEVMSSSLPVLPAPFEEKFPKLPDSQQLSVERELRSSPIIP
HHTPFVSNSGVVGPLYSSASGFSSDLHFSSVSPNVRHPNGSSFVSRSSNVGISFPSTPSH
SGAFQSSTVNHPRESTKVTWCPEPIQSMIDYSDNITDGNNEIQNSCDVVSDDFAKQNEWW
TDLMNEDWKDILNETSATVSQPKAMQSAAQASPSISAHQPQIHQVVPSQSGEICTVSNSS
SAANATTTKPRMRWTPELHECFVDAVNQLGGSEKATPKGVLKLMKVEGLTIYHVKSHLQK
YRTARYRPDSSEGASEKMVTPQQEVSSLDLKTSINLTEALQLQMEVQKQLHEQLEQYGGK
VPPPRGLPLYMYSMWIIALKNARKVFCPKPLLDVVQRNLQLRIEEQGRYLQMMFEKQCKS
GIDKLQAPSTKEEPSTVSSDQTYSADKELPGKVRGESGNDPAGTEIAEGSRQVGDKQKMP
EAESCNERESNTVSGSHSPPCKRA
>PdaPSR_PDK_30s806121g001
MNSHERPMSCVQGDSGLVLTTDPKPRLRWTVELHDRFVDAVTQLGGPDKATPKTIMRVMG
VKGLTLYHLKSHLQKFRLGKQPHKELNDHSVKDASALEMQRNGASSSGIMSRTMNENVHI
TEAIRMQMEVQRRLHEQLEVQKHLQMRIEAQGKYMQTILEKACQTLAGEGMASGSYKALG
NQGVVDMAAMKDMGSQMSYPSLQDLQLYGGDQLDMQQQMDRPLDGFFPTNEGICLGKKRT
HPYSSNGKSPLIWVDDFRLQELGSAAACMGTQEEPSKSDQLQLAPSVIDSGMDMDSMTDV
YETKPVLSTDSMGEKKYEGSSKLERPSPQRATLPVERINPMIRGGALPQTRNISYG
>PdaPSR_PDK_30s850271g001
MWSSLMRAKYGDLVPGEGGARQGLELRAAWEGLFYARRVLGVEWVFLEGDSSMVIDWIQG
ADRFVILVLDLIRYHSITVVVRYGTARCTEYRYGTEATPKAIMKTMGVKGLTLYHLKSHL
QVQRHLQIRIEAQGKYLHSMLERACNALIDPNLGLIGLASIRCDLPELPNIETDDCLGHP
HVSLKLPSLSEIAAACMEEKSLNRAMAQIAECSVDSCLTSTESPGRAPVLGSQAAALRKG
SCPLLGTDESHAWDGDFLCSGCNEAMRVQMELQRRLHEQLEVQRHLQIRIEAQGKYLHSM
LERACNALIDPNLGLIGLASIRCDLPELPNIETDDCLGHPHVSLKLPSLSEIAAACMEEK
SNIETDDCLGHPHVSLKLPSLSEIAAACMEEKSLNRAMAQIAECSVDSCLTSTESPGRAP
VLGSQAAALRKGSCPLLGTDESHAWDGDVHEDGQWVSSI
>PdaPSR_PDK_30s916311g004
MYHAKKFSTVSMVPHKAQGTEQLVNAGVMGGSNVSNPTNSGGSGKQRLRWTSDLHDRFVD
AITQLGGPDRATPKGVLRVMGVPGITIYHVKSHLQKYRLAKYLPESPADGSKDEKKDSAD
TLSSMDSASGIQINEALKMQMEVQKRLHEQLEVQRQLQLRIEAQGRYLEKIIDEQQKLNG
ALKAPGEEQNPSPSPSASQDCLGDPSSPLKKQRIADQSPDPTHEMPPPESKPHFIGQ
>PdaPSR_PDK_30s951241g002
MTYHASRNVGSSMFGSEWNMSLPVANIPTNWHGPSGLNEENVPTTMPDIHEIGSYQQPCS
PLLVSLIHQEGLNQCHSLPNLTSSSSTSTKSTDICLQSSIHYTDRYASSSSALEQYPYSG
NMFPLLQIHLSSMQTSAPTSLQGLGLTYQPNEDAQIPGDIDDILPSAVEQEHRMESDQPK
QIILSNGNQDRYEIIGVNQEAQVSEVNVGNKSRMQWTQELHESFVQAVNDLGGADRATPK

CIVIKMGIPGLTKDHVKSHLQKYRLSRYRSEGKEDKRPSCPEGKKRNATEDSGNPNLSSN
ESETLHMQWKLQKTLHEQLKWQYYCANVVMALNIIRKLQLQTKENGRQLQKFMEDQYKVR
EAFFQATQSMSMTKGVATEGTLAPFSSDKLPVSLDEEHSMKDGMLSDLAKLDVSIDVELA
SLSYKRARIDAESSKATISGNVFQL
>MacPSR_GSMUA_Achr10P23260_001
MSSHSLITVKQNNSPEGTKHLCHSSPLSVQFNNEQDCQKLSRSSYVETVLHNSSSFSKDL
PLHTKKSSSGPEPGSSCSYVSHPQYPEHMFSRSSTFCTSLYSSSSTSSESCQKLQNLPFL
PHPPKCKHQNSAVQSSNSPLQFSGDISATSGEDEHTDDLMKDFLNLSGEVSDGSIHGENC
GSSGLALNEQIELQILSEQLGIAITDNGESPHLDDIYETPQVSSLPLSANHNQTDQPSKP
HTKVQLHSPPSINPASAANKTRLRWTLELHERFVEAVNKLDGAEKATPKGVLKLMNVEGL
TIYHVKSHLQKYRLAKYLPEAKEDKKASSREDKKSSSLVSNDGDLANKRSIQVTEALRMQ
IEVQKQLHEQLEVQRALQLRIEENARYLQKILEEQQKANNSSSSTQRFSSEPPLELRSPF
TEKADARVDSSPLNSVKQRGNESNDRESDSESVEDSKRMRLDVEHIRPS
>MacPSR_GSMUA_Achr11P02560_001
MSSSLPVLPTPSEEKFPKLTDLQQVSVEREIRNIALASHHTPFISDGGIVGSLYSSPSGF
SSELNGSSFSLHEGHPLFATQSPRVGVSLHPTNPSCPGTIQPITTNFPRQSTEVAWCPDA
VDNILDFTDNNIGVGNQMPSNSAMVSDDLSKQNEWWTDIIDEDWKEILNETTAIESQPKV
VYSTTQSTPNISVHEPPIHHSVPSNSGETCAVISSSSAATNAAKPRMRWTPELHECFVDA
VNQLGGSEKATPKGVLKLMKVESLTIYHVKSHLQKYRTARHRPDSSEEIFNKKITLKEEI
PSLDLKTSFDLTEALQLQMEVQKQLHEQLEIQRNLQLRIEEQGRYLQMMLEKQCKQSIPV
ATTVEGPSTTASDLMHSTDKVDVPENSDDSANTNEGSKQVGNDEKMPDAELSDKK
>MacPSR_GSMUA_Achr11P06340_001
MENIYQRRPPLAASLFEAPGGEGAGVMLSRDPKPRLRWTPDLHDRFVDAVTKLGGPDKAT
PKSVLSLMGMKGLTLYHLKSHLQKYRLGKQTRRETEQEAKKKGSNSSKINCSSTTSNYVS
RTDGAGEMPLGEALCYQIEVQRKLQEQLEVQKKLQTRIEAQGKYLQAILEKAQKSLCFDN
NRSSGSLEATRAQLTDFDLPQSGLMENVGRVCEEKHSELREVRPQENIKKRNNSGFQLYQ
EGRDEAEDSSLLLLDLNVKGSSGEMVGGSRGNDLDLRIQTQGL
>MacPSR_GSMUA_Achr11P14430_001
MFPSKKATVSSHDRGMCVQGDSGLVLTTDPKPRLRWTVELHDRFVDAVTQLGGPDKATPK
TIMRVMGVKGLTLYHLKSHLQKFRLGKQPHKELNDHSFKNAAALELQRNAASSSAMMGRH
MNENVHAEAIRMQMEVQKHLQLRIEAHGKYMQSILERACQTLEAESLVSGSYKGHGDQGV
ADMGAIKEMGSPMSFPSLQDLHLCGGDQLDLQSQVDGPLDGFYPISDSILGKKRMIWADD
LRLQELGSTSACVGPKEEPCCKSEQLQIVPSVIDAAISSDPMANVYEGKPVLSMERPGEK
QYEGSLKLDRPSPRRASLPMERINAMMAGGAVPQASNRSYG
>MacPSR_GSMUA_Achr1P16440_001
MFSGLIHRPEASIPPEDAHGPSLVLTADPKPRLRWTADLHERFVDAVAQLGGPEKATPKA
IMRTMGVKGLTLFHLKSHLQKYRLGKQSGKEMTEQSKDASYLLENPGSSSLSPRVPTPDV
NEGQEVKEALRAQMEVQRRLYEQVEVQKHVQIRMGAYQKYIDSLLARACKIASEQIALNS
FGTAEHELPDMTPRVVCPPSSPLSPSILHQLSVSSINLQSPGCKTSPSSSAIEGQLFCQK
PPELKNKPC
>MacPSR_GSMUA_Achr1P19430_001
MFSGLIHGSEASIPPEEARGPSLVLTADPKPRLRWTADLHERFVDAVAQLGGPEKATPKA
IMRTMGVKGLTLFHLKSHLQKYRQGKQSGKELTEQSKDASYVLENPSSSALSPRVPAPDV
NEGQEVKEALRVQMEVQRRLHEQLEVQKHVQIRMDAYQRYVDSLIAMAYKIASDQIASSS
FGMTEHELTEGT
>MacPSR_GSMUA_Achr2P01180_001
MYHHQHHHQGHNNILSCRTAFPAEKHLLLQGGSIPEDSGLVLSTDAKPRLKWTAELHERF
IEAVNQLGGADKATPKSVMRLMGIPGLTLYHLKSHLQKYRLSKNLQAQANGAESVIGCKL
AAERTSEGNGSRASNTNIIPQSNKTYPINEALQMQIEVQRRLQEQLEVQRHLQLRIEAQG
KYLQSVLEKAQETLGKQHLGTPGLEAAKVHLSQLVFKVSNECFSNALTGLEEIPAPNTLQ
VHPPQLADFSVQSCLTSSQGSQKDLDMANFRRSLRAYMHENARLEGSQSAWCYLNEHKTF
PSSMFGDSERTSFKVEDFVSPLPVRPRVEREGEAGSDAQQTERSDPNGKRTAEQQERGKQ
SDSFGLAGHTAQLDLNADEDDEGATNSKFDLNGFSWS
>MacPSR_GSMUA_Achr2P10180_001

METRSALPMGKLNMEQLSDARSSGVMSSTLPVRPNNSGEKFPKIPDSQLVLMDREIRSNP
LPSDYTPFVSDGGNVGPLFSSPSGLSSDLHFSSSLPHERHTNGTPFANQLLNAGVSLSST
YSSNTGIFQVPNNNLPKDPTAVTWCPETVQGIGNSKINDSLIVVSNDLSKQNDWWSDIMN
VDWKDLLNDTTIAESQSKVVCPAAQSSPDISMHQLQSHRSVPCHAGEVCSITSPMSATTS
TATKPRMRWTPELHELFIDAVNQLGGGEKATPKGVLNIMKVEGLTICHVKSHLQKYRTAR
YIPDSSEGMSEKRITQSEELPSLNLKTGIDFTEALRLQIEVQKRLHEQLEIQRSMQLRIE
EQAKRLQVMIEKQRKSTMEKQHASSTPLEPTPHSTAEIELPEGGNSSSSIQRTEDSRQVG
NKWKMPELETSNKKGTDAITGCPSTSKHIRVSNEDA
>MacPSR_GSMUA_Achr3P11970_001
MVLNSGGNNSDNHNLASRQRLRWTNELHDRFVLAVTQLGGPERATPKGVLRIMGVAGLTI
YHVKSHLQKYRLAKCVPDSSADDAKSEKKDPDGVSSGLESSSGTQITEALKLQMEVQKRL
HEQSEVQRQLQRRIEAQGKYLKTIIKEQQQLSGELAETPGGDISACSYVDNSSGSAKTGP
HRSVNGDHGGTGKLLKSLSHDNSLREPEA
>MacPSR_GSMUA_Achr3P11980_001
MFSGLIHRPEASIPTEEAHGPSLVLTADPKPRLRWTTDLHERFVDAVAQLGGPEKATPKA
IMRTMGVKGLTLFHLKSHLQKYRLGKQSGKEMTEQSKDASYLLENPSSSVLSPRVPTPDV
NEGQEVKEALRAQMEVERKLHEQVEVQKHVQIRMEAYQKYIDSLLAKAYKIASEQITSNS
FNTTEQELSDMATRVICSPSDPLSQSILHQLSVNSINLQSPGCKTSSSSAIEGQFFYQKP
HELKTKPC
>MacPSR_GSMUA_Achr3P23640_001
MNSHERPMCVQGDSGLVLTTDPKPRLRWTVELHDRFVDAVTQLGGPDKATPKTIMRVMGV
KGLTLYHLKSHLQKFRLGKQPHKEFNDQSAKNAALELQRNSASSSAIMGRNMNDRNMHMN
DAFRMQMEVQKHLQMRIEAHGKYMQNILERAYQTLAAESMVSGGYKGQASRGVPDMGAVK
EMGSPVSYPSLQDLHIYGGDQLEMQHQMERPLDGLFPTDDSIMSKKRSNPYHSNGKGPLI
WADDLRLQELGSTAACIVSQEEPPPSKSEQLQIASSVIDTGIDADSIANVYEAKPILSMD
SSREKKYDGRTSKLDRPSPRRDPLPMEMMNPMITGGAMPQARNLSYG
>MacPSR_GSMUA_Achr3P28410_001
MSPMNGVNNSNNPNLASRQRLRWTNELHERFVDAVTQLGGPDRATPKGVLRIMGVAGLTI
YHVKSHLQKYRLAKYVPESSADGTMSEKKDDRNLNSGLESSSGMQITEALKLQMEVQKRL
QEQLEASEHIIVQRQLQLRIEAQGKYLKKIIDEQQRLSGVLADLPGADIPAPTSGNLATP
TGPSRAFHVMIHSPPVASH
>MacPSR_GSMUA_Achr4P11370_001
MGPINGVNNSSNPSLASRQRLRWTNELHERFEDAVTQLGGPDRATPKGVLRIMGVPGLTI
YHVKSHLQKYRLAKYVPESSADGTRSEKKDDDNQISGPESSSGTQITEALKLQMEVQKRL
QEQLEAS
>MacPSR_GSMUA_Achr4P19300_001
MISVIFYVKIPSLIRGADRQVGVPPVHLLSFMIPLYVSLPLLFCSVLLFVLVGCGPSSSS
TEKQNSQLQTEKQLPKALSTAPGTALSNKTRIRWTQDLHERFVECANRLGGAEKATPKGI
LKLMNSAGLTIYHVKSHLQKYRIAKHMPERAEGKTLLLRAAANVAELDPKIGMQFTEALR
LQLDVQVRLHEQLEIQKNLQLRIEAQSRKLQQMLEEQAKSNTSHVQTENMDVSSAGDSPA
ESFDDAQLLHQVSIED
>MacPSR_GSMUA_Achr4P30200_001
MSSSLPVLPNTLEEKFPELPDPQHVPLEREIQRNPLPSQRPLFSSPSGFSPDLTFSSTLA
HERHTNNTPFVSRSLNAGVSLPSTYLSNMEIFQVPNNFPKDPTEITWCPDSVQGMLNCSD
GVIMGNNQIQNSSNKVSNDLNKQNEWWSDIMNVDWKDLFDDTTISESQPKVVYPPAQSSS
NISKQQPQTDQSVPCHSGEVCAVTGASSSATTAAAKPRMRWTPELHECFINAVNQLGGSE
KATPKGVLNIMKVEGLTIYHVKSHLQKYRTARYKPDSLEGMSEKTATQSEELPSLDLKTG
IDFTEALRLQMEVQKRLHEQLEIQRNLQLRIEEQGKYLQMMFEKQYKSTMDKTHCSSTVE
KPTTISSDQTHSTAKIDLPEAQNSSTDSKITEGFRQVSNKWKMSEVEPSNEKETDALTSS
LPPSKHSRINDEDS
>MacPSR_GSMUA_Achr5P07670_001
MYHQHQQGGHNNILSSRTAFPAERHLLLQGGRIPEESGLVLSTDAKPRLKWTPELHERFI
EAVDQLGGADKATPKSVMRLMGIPGLTLYHLKSHLQKYRLSKNLQAQASSGSAKIATGCK
LVAGRTAEGNGLLLGSTNIIPQSNKNIPINEALQMQIEVQRRLQEQLEVQRHLQLRIEAQ

GKYLQSVLEKAQETLGKQNLGSPGLEAAKVQLSELVSKVSNECFSNAFPGLEEIRNPNTL
QVNPSQLADCSAESCLTSSEGSQKDQDMTNFHRSLRAYGGGADSEAQQKERSEEHMFLEH
PNNKRVAGQQDRGKQSNSFGMPGHTAQLDLNADEDNEGDRDSKFDLNGFGWS
>MacPSR_GSMUA_Achr5P12960_001
MSDDLNKQNEWWSEIMNEDWEELLNDKTVAESQPKVVYPAAQSSQNMSVHQLQTHQSVPC
HSGEISAVTGPLSTATAAATKPRMRWTPELHECFVNAVNQLGGSEKATPKGVLKLMKVEG
LTIYHVKSHLQKYRTARYRPDLSEGMSEKKITQSQEIPSLDLKTGIDLTEALRLQMEVQK
QLHEQLEIQRNLQLRIEEQGKYLQMMFDKQCKTTMDKLHAPSTVEEPSNISSELTQSTAK
IEFPDAGSNPIDSNKTQITEQIGNKRKMPDVEPSNHKDVDAVTDSPSSAAKCARVDDEEA
>MacPSR_GSMUA_Achr6P01310_001
MMRHHRTRSAGAEASAYEAAGATLSRDPKHRLRWTPELHDRFVDAVAKLGGPDKATPKSV
LRLMGMKGLTLYHLKSHLQKYRLGRQTGRETKAETSSKGSNPADTSYSADSSVFGGPESV
GGTPLAEAIRYQLEVQRKMNEQLEVQKKLQTRIEAQGRYLQAILEKALNSLSLDMNASAS
VEATSSSQLTDCNLALSGSMDDATKNSTSLKRSNTSAFQLHREGRQEHEDSKLGNDAGTV
LLDLNAKGSYQLFGFFGSPYAHLLCYRTIDLEDKLDERRR
>MacPSR_GSMUA_Achr6P32220_001
MFQHQPHQGHNNLLSPRTTFPSERQLFLQRGNTTGEPGLVLSTDAKPRLKWNPELHERFV
EAVNQLGGADKATPKTIMRLMGIPGLTLYHLKSHLQKYRLGKNLLAQTNTGSTKNVIGCT
LAAEKTSEGNGSSMNIAAPTNKTMQINEALQMQIEVQRQLHEQLEVQRHLQLQIEAQGKY
LQSVLEKAQETLGKQNLGSAGLEVAKFQLSELVSKVSVDSCLTSSEGSQKEQDTHAAIMG
LRPQREIRDSGTVPEARKKDRDDEDMFLEHTSSKRAAVKQESRKQTDGFELPCLTTDLEL
KIHEGSEGASSCKKFDLNGFSWN
>MacPSR_GSMUA_Achr7P06180_001
MYQSKAVSSISPNSKNPLAHDQHLELSDNVVFSNSGRNDLGNPNLASRQRLRWTNELHDR
FVDAVTQLGGPDRATPKGVLRIMGVSGLTIYHIKSHLQKYRLANYIPDSSADGAKSENKD
SGDHTMRPENSSGMQMTEALRLQMEVQKRLHEQLEVQRQLQLRIEAQSKYLKKIIEEHQL
LSGEPAELPGVGIYAYSNDNCLNSDDKTDPPAPAPTSVSPVQDKAASKPLTPDSTCHVSS
PLESPKHERVKRQRFPGHETTDLMPAQHILESSSGSDFQQPCSVFLAGRGQFDASSASIF
NEQFGNDSGSEL
>MacPSR_GSMUA_Achr7P06370_001
MDLQTNDLHEFSHGSFRNATFEASNPSYQRNTSLGCFPAPRNPWSSVSLVSSSTSASFEE
EMDSSSHGLMPPTLPSCVSSCVGSSPAVFFASEQSMSFPQLYLYRSETLPSFSKSTKNRP
AAASFFFDRDDDSVKQYRLPSQPRDALDSDLKLPMLQNTRASSRDGFQASRIQPYPGRAD
WSASLQTKHLSPPQVVHDPSVSTEFFSSSSSCLLFGIGFYSEHLLVACLAVCSSTKVGWG
IPSCSMDKPNLRLQTEKQLPITSASVSSATAISNKTRIRWTQDLHERFVECVNRLGGAEK
ATPKGILKLMNSAGLTIYHVKSHLQKYRIAKHIPESTEGKFERRAAAVSVTELDPKIGMQ
ISEALRLQLDVQMRLHEQLEIQKNLQLRIEAQSRKLQQMFEEQVRTTKGPAELENLGDLF
SGSPAASLEDAQLWCAPDGPQSADFPLQKC
>MacPSR_GSMUA_Achr7P15740_001
MSSHSLTTAKQNNNPEGTKHFHHSSLLSVQLSNQHDCKNLLDDRLSSTSPYRAQTELLKS
SSFKPGSPRSYVSHSQCSDHMFSRSSTFCTSLYSSSSAISESSRKLSSSPFLPHPPKCEK
QNSAVQLSSSPLLFNGDTSARSCEDEHTDDLMKDFLNLSGDASDGSTHGEIYGNNGIALS
EQIELQLLSEQLGIAITDNGESPRLDDIYETPQVSSLPLSSNHNQAVQLSEPPAKVQLHS
SPSTATVSASAANKTRLRWTLELHERFVEAVNKLDGAEKATPKAVLNLMNVEGLTIYHVK
SHLQKYRLTKYIPEAKEDDKKASCPEDKKAPSVSDDNDLAKRRNIQVTEALQMQIEVQKQ
LHEQLEVQRALQLRIEENAKYLQKILEQQQKARNPTSSTQRSISEVPLEQHSPAPDQSEV
GIDCSPPNSLKHKENDSESDSKSVKDCKRIRLEVSCTLAFLSGENFVHIFAPPYLLSSPG
TFSDR
>MacPSR_GSMUA_Achr8P12580_001
MFSRETTKINLSPDTLHDTLDYPDNNFSLSNQILGNSTMVSNDPSKQKEWWTDIIDEDCK
ETAVESQPKVCDFIFVGTFIKLQKISIFSFFLLHQPPIHHSVPSNSGEVCAVTSAAKPRM
RWNPELHECFVNAVNQLGGSEKATPKGVLKLMKVEGLTIYHVKSHLQKYRTTWYRPDSSE
GPSNKKTTLKEELPSLDLKTSFDLTEALRLQVVVQKQLHEQLEIQRNLQLRIEEQGRFLQ
MILEKQCKSNIDKLQITSTVEGLSTTTSDIMHSIDKFELPENDRYIAGSSNGSAETKEGS

WQVSIRHKMPEAEPSAEKEADAVDGSHSSACKPAKGQVGETKTPP
>MacPSR_GSMUA_Achr8P21520_001
MYHHYHQSHSNIFSSRETFPADGQLLLQRGGSAPEESGLVLSTDAKPRLKWTSELHERFI
EAVNQLGGADKATPKTVMRLMGIPGLTLYHLKSHLQKYRLSKNLQAQANSGSTKSVIGCT
LAAERTIDVNGSFMVNTSTMPQSNKSFQINEALQMQIEVQRQLQEHLERYTRVQRNLQLQ
IEAQGKYLQSVLEKAQETLGKQNLGSPGVEDAKVQLCELVSKISSECFSNAFQGSEEIPG
SQSACKSLRSCNGSLPLCQQQVHENSRLENTQSAWCYLHEHKTFSSSSLGDSERTSFAAQ
KDFETLPTSIKTQNEGGTGSEAQGKESDDENISVEHPNSSRPADQQGRGKQSDRFGLLTH
TAQLDLNAHEDNEGGTSSRFDLNGFSWS
>MacPSR_GSMUA_Achr8P33560_001
MFPSKKPTMNSHDRPMCVQADSGLVLTTDPKPRLRWTVELHERFVDAVNQLGGPDKATPK
TIMRVMGVKGLTLYHLKSHLQKFRLGKQPHKEFNEHSLKDASALELQRNAVSSSRMMGRN
MNDTSAAHGTEAIRMQMEVQRRLHEQLDVQRHLQIRIEAQGKYMQSILEKAYQTLAADGM
TSGDHHHHHHHHHQGLADMAAMKDLDPSMCFPSPHDLQALYGGDQLEMHQQMNASKHPFV
WDDDLDVQKLGSAASCIGPQEEASKTHRLGTAPSAIDMDTVVDVFAAKPMMLSADSAGEK
KHEGSTKLEPPSPLRAPLPLDGINPTLIGGSFAQARNLSYG
>MacPSR_GSMUA_Achr9P01710_001
MYATKRGEEFHGSLEGANLPGDTCHLFSSDPKPRLRWTPELHDRFVDAVTQLGGPDKATP
KTIMKTMAVKGLTLYHLKSHLQKYRLGKQSNKEPSDNSKDAANFVESQGTGPLSSSPLSS
SKLLAKDLIDGCDEAMRVQMELERQLHEHLEAFLLAFPESFSSSSYLPFLNSFFFITLHQ
VQRHLQIRIEAHGKYLHSMFERACNIIDPNLASNGLALVRKTPSQFPTSEAADDRLSFLC
KALKLPSLSEIAMESVQRKPSNGVSATFAECSVDSCLTSTASPGTGRSKMLGPREAAVDI
RFSYVGN
>MacPSR_GSMUA_Achr9P05030_001
MYQHQPHQSHNNLISSRIAAPSERQLFLQRGSTTGESGLVLSTEAKPRLKWNPELHARFI
EAVNQLGGADKATPKTIMRLMGIPGLTLYHLKSHLQKYRLGKNLQAQTNIRSTKNIFGCS
LAAERTPEDNGSSRNIAAPTNKTMQINEAFTKQIEVQKQLHEQLEVQRHLQVRIEAQGRY
LQLVLEKAQERLEKQSLGSVGLEVAKVQLSELVTKVSGECFGNSFLGVEEIPVLHMSQLT
PAHFADCSVDSCLTSDRSQKEQDTHDTTIGIRACHKDLPSCMKQFGEDKLEQTQHAWCGD
LNEQKIFSSVPEETKKQSDGFGLSCLTTELELKIHEDDEGASGCKQFELNGFSWN
>MacPSR_GSMUA_Achr9P19070_001
MFSGLIHRPEASVPPPEEGHHCPNLVLTADPKPRLRWTADLHERFVDAVTQLGGPAKATP
KTIMRTMGVKGLTLFHLKSHLQKYRLGKQSGKEIAEQSKDASYITEKPNSSVSSPRMPTS
DVNEGQEVKEALRAQMEVERRLHEQVEVQKHVQIRMEAYHNYIDSLLAMAYKIVSEQIAL
SGLGSVGDLPNLANAFICSLSDPSSQSTFHESSVGSIKLHSAGGKHSPPSAIECQFNQQE
PLKHEPC
>MacPSR_GSMUA_AchrUn_randomP18240_001
MYHASKFSTVSMAPDNGQGTEQVGSIGISGGSSVNNPVNPGGSGKQRLRWTSDLHDRFVD
AITQLGGPDRATPKGVLRVMAVPGITIYHVKSHLQKYRLAKYLPDSPADGSKDDKKDSGH
THSDTDSAPSAQFNESLKMQMEVQKRLHEQLEVQRQLQLRIEAQGRYLQKIIEEQQKLSG
VVQKHKSSLSLPISDSNRNPSPLKKPRTAVVNQSSDAAQETPKQESKPDLCSDHCIISCG
FGAGTMLDEGTGSTQQNLPNEVEVGLRFVQFICTCHATDNFDGDATLVGFLSAISSVPYK
VFFCKYQFRQCFFFFFLG
>PeqPSR_PEQU_00348
MYHLDQSGNRVISYRTSNPSKRNLFLASENCPGDAGLVLSTDAKPRLKWTTELHERFVEA
VNQLGGPEKATPKTVMRVMEVPGLTLYHLKSHLQKYRLSKNLHVMSNGCGAMRNVIGHSV
APDKTSEGETIVNNMNVGSHSNTVVKINEALQMQIEVQKRLNEQLEVQQHLQVRMEAQGK
YLQSVLEKAEITLENQNLCPVGSEAAKVQISELASNMKEIPDPHNLKVNPIQFADCSINS
CLTTSEGSKMDRAMHCTNMILRAYHGNLSQSKAQTEKNSRLEQIQSTSYRELKGQERDPG
ILSMSITNQKDKDEGNNISKALHMRREKEDDYLEEPGRGRAPFQQKSSEKFSDHGIGTQL
DLNVQDANNHAAPDYREFDLNCSSWS
>PeqPSR_PEQU_06318
MIESFVLMGRGYQYGATEAAAAVLSRDPKPRLRWTPDLHDRFVDAVAKLGGPDKATPKSV
LRVMGMRGLTLYHLKSHLQKYRLGRQARKETGLDGIKDGKPSKASFNSSASSIMSIGNTG

GREIQLTEELRYQIEVQRKLHEQLEVQKKLQSRIEAQGKYLQAILEKAQKSLSFNNNNNN
NNMEAARAQLKDFNQALSGLMDNVNQFCVVKGAAELDKVAASFSLNSSPTNSAFRLYQEG
EETQNLRFSSHRGESLLLDLNLKGNQDYFTGNMGSTDLDPQMIMQS
>PeqPSR_PEQU_07132
MPNRLDHHGPVLSADGKPRLKWTRELHEQFVDAVSELGGAEKATPKLLMRLMAVPGLTLY
HLKSHLQKYRLAKNRDPSTLTNDKKTKDDELKNRNDKTQPQLNEAILQQIQIDVHRKLQE
QIEVQKHLQLRIEAQGKYLQSVLKIAQETLAGYSSTLGTEAARSELSELVSVVETECLSS
SLNHGGLISKGNQNADSSMDSCLTSELFELREKYAGQHKLEASSCSNPVNGRTAETRGEI
EEFDLNN
>PeqPSR_PEQU_09870
MDFPQFEPQVRSPSLFEPPVSSFCSSVSSASQFSFRPEKKRDVSYSWQTFESEVKLPLHE
YGTSAACDKRQKLFYRHHQDVLAQSVAQSTEEGLDSNLNKRQLASLGTADRSSVKYNLEN
FLVRNPNFQIQNVEKHSEASIVVPVTSFNPNLRSPLPSKTRIRWTPNLHEKFIECVNCLG
GAEKATPKGILKLMDLDGLTIYHIKSHLQKYRLAKYLPESSEGKFQRKANELQQFDPKTV
IPITEALQIQLQVQRHLHEQLEIQRNLQMRIEAQGRQLQKLFKEQMKSNKILFETQGLDD
LFPDKIHEKTPENDQVFGVGEDHQNNHFSSKIS
>PeqPSR_PEQU_10677
MQAARGSGEPGLVLSTDAKPRLKWTPELHSRFVEAVKQLGGAEKATPKTVMRLMGIPGVT
LYHLKSHLQKYRLSKNILAQSNVKIPTAALDKTSLEGNGSLVNNINFGPHKYKTMQISEA
LQIQMEQRIHEQLEVQKHLQRRMEVQGKYLQSVLEKAQEAFGKQNSGSAGLEASRTQLSE
LISKVSNECFTSTFPYLEKTYCSHNQEGHMAQINNFSMYSCLTSCQESQNGQEMHDTNMV
LTAYYHGNLSLGTQEMEQIQSGWSEDLNGSKIFSQTGVKDSTQNTLPVSRGFGILSICKS
EGNRNLEIHPRQNEEHNSYLDRADYERYVIQQRSLKNPSECEQPSATTQLDLNACDEYNV
SKNGNGNDLNGFSWS
>PeqPSR_PEQU_11035
MFSSKKPTMNSHPESRAPLCVPVDSGLVLTTDPKPRLRWTVELHERFVDAVAQLGGPDKA
TPKTIMRVMAVKGLTLYHLKSHLQKFRLGKQPHKEFYDHAKDALDMQRNAASSSGIMSRS
MNDRNVHITEALRMQKEVQRRLHEQLEVQKHLQMRIEAQGKYMQSILEKAYQTLAGEEKS
IPATASFKFHFLPRPPFIQQ
>PeqPSR_PEQU_15381
MKQICSSGVSAVISSSLPTQPSLFDERLHEPLSIVPQQTPLVSNSGVMEPLFSSASGFSS
DLHYFSVAPQGRYPNGAPFIAPSPSCGMPMPSVQSLASNYPLEATGISWVPESIQDIFSY
SDNMNIGSSQLQNSPINNQIKNSVIAASDDLAKQNEWWDLGSEDWKDLLNDPVGPETQLK
VFVLLSFLVFSVSVVHVHQQVPLHSGERSIMSSPSPSINGAPPKPRMRWTPELHERFVDA
VKQLGGSEKATPKGVLKLMKVEGLTIYHVKSHLQKYRTARYRPDSLEEKQVYSLDEKASL
DLKTGMELTEALRLQMEVQKRLHEQLEIQRNLQLRIEEQGRCLQLMFEQQYKVGLDKLKP
PASDQDEQTAEYCCTTPPSPKSSAFEKERDEKGNSSSTTIAKEVKLTAVADQQRVEDNGF
LRAKPPDDRRTEGDLKEHISE
>PeqPSR_PEQU_17183
MYHLDHNSSNLISLRESYPSERNLFLQAGNGPVDSGLVLSTDAKPRLKWTPELHERFVET
VNQLGGADKATPKTVMRLMGIPGLTLYHLKSHLQKYRLCKNLQAQTNGRSAKNGGAAEAP
DRTAEGSSSLVNNINVGLHNKTVQINEALQMQMEVQRRLHEQMEVQRHLQMRIEAQGKYL
QTVLEKARTTLEIQNTGSAGHEAAKEQLSELVSKVSNECFTSAFIGLGEIHGADNLRECR
VQHADCSIDSCLTSCEGSQKEQKMHGINLNLQGYHSNLSSGTNEKHENVRFESINSGWWC
GELNGHNMFSQPRVNVSENCSLPVSKTNCRQIDKYNSDHERPNYGGTVQQESSKNLNEFG
MPCLTPQLDLNTRDDNDAARNCNEFDLNGFSWN
>PeqPSR_PEQU_17379
MFPSKKPTMNSHEGPLSCVQGDSGLVLTTDPKPRLRWTVELHDRFVDAVTQLGGPDKATP
KTIMRVMGVKGLTLYHLKSHLQKFRLGKQPHKDFNDHSKDALDMQRIGASSSGTMGRTMT
DRNAQTAEALRMQLEVQRRLHEQLEVQKSLQMRIEAQGKYMQSILEKACQTLAACENLQP
SADAYKAAFSGHGSVQIDLMKEMTSPMAVSFHSFQDLNVFTNNNQLDHMNDQDDTFFRSN
NDDNNNMFMGKKKQHEFYSINQSGKSAMIWGDEEDYQFGSGQEENCNCKSEQMHEISASV
INGSLDVESMMEVYEAKPRVSGEECRKAPVLTKEMLSPVIRGSGLSRGRNLSYG
>PeqPSR_PEQU_22301

MGSTSVPKHTPYASNSEFVGPLFSSASGLSTNLHYSSISPQGRHPNGAPFISQSQSFGIP
SSLLQSSHSKSFQASTSNYPMETSGISLHTEFVHNALTYSDNLSGGINNQIQEGTINDQI
EDLVINDQIHSSTIAASDDLTKQNDWWELNSDDWKDLLNDPVVTEAEQKVSKASLTMSAD
QTHVCPQLPLQSGELCIVTSPLSSTSALPRPRMRWTPELHESFVDAVNQLGGSEKATPKG
VLKLMKVEGLTIYHVKSHLQKYRTARYQPEWADGTSEKKASSLDKLASLDLKTGMELTEA
LRLQIEVQKRLHEQLEVQRNLQLRIEEQGRYLQMMFEQQCKAGMDRFRPPSAPEEQATQS
SSTTTVSPTRNNASEKALDEQESEKPRLVGGKQKPPDPAAFLGSESPHGKRVKGDQNASS
E
>PeqPSR_PEQU_24486
MYQPSLSNISLANYPSLAHDQPVEMNSTAMSPHAAGNNPNNPSSASRQRLRWTNELHERF
VDAVAQLGGPDRATPKGVLRVMGVQGLTIYHVKSHLQKYRLAKYLPDSSAEGAKVEKKES
DDVLSNLESSTGMQITEALKLQMEVQKRLHEQLEVQRQLQLRIEAQGKYLKKIMEEQQRL
SGGLAPTSTNQTLEDKSDPSTPVPVTKSSLHGSLFKSFSFEDSHHGPLTPETGSPSGSPK
HERSVKMQGDRDSPKQELMLGHHILESSSSLFLGGGL
>PeqPSR_PEQU_26158
ATPKGVLRVMGVPGLTIYHVKSHLQKYRLAKYIPDSSSDGLKDESKEPGDLLSGLDSSSG
VQITEALKLQMEVQKRLHEQLEASQRQLQLRIEAQGKYLKKIMEEQQRLNGVLAETPGVN
LPPPPSSADFCLDSNPSTPAPICESLHKDKIPNGKHGNADRLIKSTSLDDSSRQEPLTPD
SSFHVDGSTSESVKHERPNKRLRCNGGGMHGAGNVDLVLGNHILESSSGSEFQQQCSMFN
AGSHFDSSGGSFGDDDQFGNGSGSD
>PeqPSR_PEQU_26160
MFPVLIHRQEASIPQEEIHSPSLVLTADPKPRLRWTADLHERFVDAVAQLGGPEKATPKT
IMRTMGVKGLTLFHLKSHLQKYRLGKQSGKEMTEQPKDVPGSSSLSPRMPTPDINENQEV
KEALRAQMEVQKRLHEQVEVQKHIKIRMEAYHNYIDSLLEKACKIASEQIASGGFNGQLG
PSQTSLNKSVFQQLPLTESSPHFSH
>PeqPSR_PEQU_30954
MNISAMAPHAGGNNQNNPGAASRQRLRWTNELHERFVDAVTQLGGPDRATPKGVQRVMGV
QGLTIYHVKSHLQKYRLAKCLPDSSAEVTRELKESDDVLSKLDNSSGIQISEALKLQMEV
QKRLHEQLEVQRQLQLRIEAQGKYLKKIIEEQQRLNGGGFMPISMDQSSEDKTDPSTPTP
IAKSSLHAKSNTNLLNSLSFEDSHHEPHTPETGSPLMSLKHERQVKSQ
>PeqPSR_PEQU_34364
MYIKGGEFVGSIEGAKLPVDANLVLTSDPKPRLRWTVELHDRFTDAVAQLGGPEKATPKT
IMRLMNIKGLTLYHLKSHLQKYRMGKQSLKELENTNDASGVLESRGSKTSTASTQVTQEF
NDGCNEAMRVQMELQRRLHEQLEVQKSLQLRIEAQEKYLQSFLDRACNALVDPNLTEIGF
DAIKQQFTELSNRNINGGIKYEPVKLPSLSENARVFAEDKPWAGLEQITEHSVDSCLTSA
GSPSAFCSKLSLKKRLHPMLCNGNLEVGVAVHEDDLWMCSI
>PeqPSR_PEQU_40820
APFIAPSPSCGMPMPSVQSLASNYPLEATGISWVPESIQDIFRYSDNMHIGSSQLQNSPI
NNQIKNSVIAASDDLAKQNEWWDLGSEDWKDLLNDPVGPETQLKVFVLLSFLVFSVSVVH
VHQQVPLHSGERSIMSSPSPSINGAPPKPRMRWTPELHERFVDAVKQLGGSEKATPKGVL
KLMKVEGLTIYHVKSHLQKYRTARYRPDSLEEKQVYPLVEKASLDLKTGMELTEALRLQM
EVQKRLHEQLEIQRNLQLRIEEQGRCLQLMFEQQYKVGLDKLKPPASDQDEQTAEYCCTT
PPSPKSSAFEKERDEKGNSSSTTIAKEVKLTAVADQQRVEDNGFLRAKPPDDRRTEGDLK
EHITE
>AofPSR_AsparagusV1_01.3515
MYQPKPVSSLGPNHGNSLARDSQLELSDNMMSPNSGSPNTNNPNMASRQRLRWTHELHERFVDAVTQLGGPDR
ATPKGVLRVMGVQGLTIYHVKSHLQKY
RLAKYIPESSSDGSKGEKKDPGDMISGIENSSGMQITEALKLQMEVQKRLQEQLEVQRQLQLRIEAQGKYLKK
IIEEQQRLSGTLTETPDRNTTPAPISD
PSTPAPISESPIQEKPTSATAIFKNESFTSHHEPLTPDSTARNGSSCESTMHERPIKRQRSNGKGDLVLGHHI
LESSSGSDFQQPSYDD
>AofPSR_AsparagusV1_01.3516
MFSGLIQRQESSIPAQDEIHGGPSLVLTADPKPRLRWTADLHERFVDAVAQLGGPEKATPKTIMRTMGVKGLT
LFHLKSHLQKYRLGKQSGKEMSEQSKD

ASYLLETPSSSALSPRLPTPDANEGQEVKEALRAQMEVQRRLHEQVEVQKHVQIRMDAYHKYIDSLLDKACKI
AHEQISSSGLNSPLMCSSSVPLSPSSL
HQITVGGITLQSPESKSSLSSIGEGQFFYQKSPELKRKLT
>AofPSR_AsparagusV1_01.3572
MNTVKKSNSPEGSLSSSHASSSSEFSYIQTGIINSSSLQTGLQFNHPNLIQEPEMRSPLSNVPHPQCSENPFS
RSSTFCTRLFLSSPSSSMTCRQLKNLP
FLPHPPKSEQLVSAVQCSSSPSLFSGDMCDVHTEGEHSDDLMKDFLNLSGDGSDGNFHQENHENSSTINEQMD
LQFLSEQLGIEMADNGESPCLDDLYDV
PQVSCVPPSTSESNQNNQQLSPPVKVQLHPTLSTSSAPAANKQRLRWTLELHERFVEAVNKLDGAEKATPKGV
LKLMNVEGLTIYHVKSHLQKYRLAKYL
PETKEGKRTSSSEEKKEPLVSNKNESFDRNKHVTEALRLQIEVQKQLHEQLEVQRKLQLRIEEHASFLQKILE
QQQKSSNSFISSKPAQSQEVQPESSDD
TSVEQAESKDGSISSRSGQKRKAIDLESDSVSSEKRQCVKLDEKLEGVSSTL
>AofPSR_AsparagusV1_01.763
MGREMGSNNMAHPYASLFSDDAVCGPMLSSAQGYSPDFHFPSIPPQGMPSFDPSFISHPSDDVLLPSMHSPCL
GTRQPSISFPKENSESWCPDSSQYIHS
GDAQIQSICNMTSDVPLSHEEWPEFSEIDDDLQALLNITDTTELQPKALCVADNSQKVPVQNKQVQQAIPSNS
GQLTVVSNPSSSCTAGATKSRMRWTPE
LHDSFVEAVNQLGGSEKATPKGVLKLMKVEGLTIYHVKSHLQKYRTVRYRPESTQDKSEDETTSAERIPSVSL
KKDFGLSEALRMQMEVQKQLHEQLEIQ
RNLQLKLEEQGRYLQKMLEQQCKADGETPKTSSTLNAVHTATSNVISSEKDDSEPRNVSISLESRESSQRVGD
KQNLLKIESHADQEFDPFDGSQSPPKK
RAKGDMSI
>AofPSR_AsparagusV1_02.1982
MMGMRGILSRLYLVYFSMSRKEFQFRARDIRRRMEWMKGELVPFEDVVDDQSITMYHAKKFSTMPLVPHRAQA
TEQPPDTGVMVGNTVNNATNSGGSGKQ
RLRWTSDLHDRFVDAIAQLGGPDRATPKGVLRVMGVPGITIYHVKSHLQKYRLAKYLPESPGDGSKDEKKDLG
ETSSGTESAGGIQINEALKMQMEVQKR
LHEQLEVQRQLQLRIEDQGKYLQKIIEEQQKLGGVLQASEDKQKPCNSPSTSQDLPSSPHKKARVGDLPRDPL
HNSHPSGSQESKPSLIPQWDQDLYGQP
SFEQQQMSVGTTSRGSELSS
>AofPSR_AsparagusV1_03.7
MSNETDDHGLALSATDAKPRLKWTRQLHQHFVEAVSELGGPERATPKSVMRIMRIPGITLYHLKSHLQKYRLA
KSRESKAGSNTEGIMAADYRPREDVVR
YVDESKTQPQFHNSTTMLNRQMEVQRRLLEQIEVQKHLQLRIEAQGKYLQSVLKKAQETLAAYTFSSSAQAEA
ADAGAECPSLLFSVPRQFLNAECSTES
CLTCSEREEELKNDGKRSNAETSECSNSWSPDESNESNPCYGGGVSEGTVERRRRVHQMERATVFGPEGIDLN
R
>AofPSR_AsparagusV1_04.938
MIAPKLCILAEATPKSVLRLMGMKGLTLYHLKSHLQKYRLGKQARKETGLLDANKNGSSSGVNYSSATNSDVS
RGNNGGEMPLAESLRYQIEVQRKLQEQ
LEVQKKLQMRIEAQGKYLQAILEKAQKSLSYDVNDTENIEESRAQLTDFNLNLSGLMENVSNIVSEDNKKTES
GKAKAHVSGFRLYQEDTEDTADVKFPS
GGGSQHLDLNMKGGGYDLFGGTLGSELDLRMQLQGR
>AofPSR_AsparagusV1_05.2141
MGINSEGNNSDNPGTSSRQRLRWSNDLHERFVDAVAQLGGPDRATPKGVLKVMGVPGLTIYHVKSHLQKYRLS
KYLPDSTSDGSNVEKGEQEGQLSTIDN
SSGMQITEALKLQMEVQKRLHEQLEVQRQLQLRIEAQGKYLKKIIEEQQKVNGDGNIPTPIPTNICHDSDKTY
PSTPAPTSEEKPTRDDHGLYNGHLHNE
SVASQHEPLTPDSSGRDDSPFRSPQNEYPNKKQRSENFVLSHQILESSSSSGFQQPGLMFPVVGSHFDSPFGN
>AofPSR_AsparagusV1_05.877
MFSGLLPAQGEIHGPRVVLTADPKPRLRWTADLHERFVDAVAQLGGPEKATPKTIMRTMGVKGLTLFHLKSHL
QKYRLGKQSGKEISEQSKDASYLLETP

TSSALSPSVGTPDANEGQEVKEALRAQMEVQRRLHEQVEVQKHVQIRMDAYHKYIDALLDKACKIAHEQIVSA
GLNNPFMCASSVPLSPSSLPQISVEAK
SSLSSITEGQFLYQKSPQLKRNLT
>AofPSR_AsparagusV1_06.154
MFSSKKPTTMNSHERPMCVQGDSGLVLTTDPKPRLRWTVELHERFVDAVTQLGGPDKATPKTIMRVMGVKGLT
LYHLKSHLQKFRLGKQPHKEFNDHAVK
DASALELQRNAASSSGFMGRAINDRNVHLSEALRMQMEVQRRLHEQLEVQKHLQIRIEAQGKYMQSILEKACQ
TLSSENIASIGYKCIENQGVVDMGVNK
DVESSMSFPLVQDLHIYSENQIDDFFPTNETLCLGKKRSNPYGSNGKSPMDWADDLHIQELEPSKSQIDMEGE
KKFEGSLKLERSSPMRPPLQMESMNPM
MSNGSLNQTRNLSYG
>AofPSR_AsparagusV1_06.309
MNMSCYNGRTLDDITDNLSLLYNANDSNFQSTCSANLADVIPSQSSMFPINMQEQYPSTGDIHNQEPTSKNFE
SILLNDDTSYKLDESFDSDVVMDALLS
DTVNQSSETHYMNVEINEANLEAEKFLSPFELSKIHTFSQKELPNIRRVEVAKPRMRWTPELHERFVQAINQL
GGPDRTTPRSILIKMGIDGITVDHVKS
HLQKYRLAKYQPEAKQDKRVSCLEGKKRNSNEKDNKVSKLSSEELGMLQTQWQLQKTLHDQLKITRELHLQTK
ENARRFQEFVENQSKLQEAFMQATQSL
SSTYTSVDDGLLNHNHTDESCFVESNGLRLAKKARFN
>AofPSR_AsparagusV1_07.828
MTPNVSMDSSNLFLSATKNLKCLSSNPLLSRGPNAESIPIALTYIALDLDQDPLSIMNQPQTSPFQFRNKTMF
PRARDPSAEFLRTASESGKMYNHHHHQ
GSNNLVSSRTAFPSERHLFLQGGTVQGESGLVLSTDAKPRLKWTPELHERFIEAVNQLGGADKATPKTVMRLM
GIPGLTLYHLKSHLQKYRLGKNLQSQA
NTGSPKNAVGCTQVADRIPEDNGLAGNMNMGVQPNKTIHINEALQMQIEVQRRLHEQLEVQRHLQLRIEAQGK
YLQSVLEKAQETLGKQNLGSAGLEAAK
VQLSELVSKVSNECFSSAFPGVEEISSLHTLQPHQAQLADCSIDSCLTSCEGSQKDQEIHNISIGLRAFHGNP
PLYTQQTGDDMRLDQSQSAWCEDLNDQ
KMFSSTIVKEPERTIFSVPRDPSILSMNITPQREKEGSGRXXXXL
>AofPSR_AsparagusV1_08.2515
MFPSKKPTLNSHERPMCVQGDSGLVLTTDPKPRLRWTVELHDRFVDAVTQLGGPDKATPKTIMRVMGVKGLTL
YHLKSHLQKFRLGKQPHKEFSDHSVKD
ASAFELQRNAASSSMMGRTMNDRNVHLSEALRMQMEVQRRLHEQLEVQKHLQMRIEAQGKYMQSILEKACQTL
SSDNMGASGSFKGIGNQDLGSPMSFPS
LQDLHIYSGDQLDGFFPNDQTLSLGKKRPNPYSSNEKSPIVWADDLRLQELGSAEPSKSDQVLQLDSGIDLES
IGDVYEAKPNFSSDVGEKKFDVSTKLE
RPSPRRAPPSMERMNPMMGSGGLNQTRNLSYG
>AofPSR_AsparagusV1_08.2758
MNTNKVRPHLCNLRSSSDAQFELVNSSCGNVSSKERWTPLKHTQKNIGDSIGSYMQSEAASSSLAFMLPSLSN
SGAKCVGSTLDAFYAAERLMGFPNFGE
GTLESLVKLPVQGGCNSMVFDDAQRFPDGSPQNCFVVCDPLNSLVNDPSLQSENDKGLSKISTVVAPNKTRIR
WSQDLHAKFVESVNILGGADKATPKGI
LKLMNSEGLTIYHVKSHLQKYRSAKFLPESSLGKADRRASTSEAEQIDPKTGLQITETLRMQLEVQRSLHEQL
EVQRKLQIRIEEQGKKLQKMFEEQLKA
NGSTFEQENSETLAPNEDTVSLEDDQILIPEEGSRNTNSPSKIS
>AofPSR_AsparagusV1_09.1226
MKPDAILFFILSLDGSGGNGNDWQQPGEATPKTILRLMNIRGLTLYHLKSHLQKYRLGKQSCKELIDNSSDAS
GAAETQSAKPSAPPSSILIAEELNDGC
NEAMRVQMELQRRLHEQLEVQRHIQVRIEAQDKYLQSILERACNILIDPDLVATEHESITHDIFELSTREIYG
CANFQCESLKMPSLAEMTVACLPQITE
CSVDSCLTSTESPAKVSALKKRPHPLLGTN
>AofPSR_AsparagusV1_09.1410
MAIVVSTVVNQFSKDFLYSLLCVGLHVPFGAYYFSNQLLALVNNRNASAGKRARTDGRALNFACWDVEACDEE
CVLFSRLIFGKGEVPFLKKSLAFRCSL

NLSTRLTIRLLDVRKIKVGINNVRLNLRTIHINEALQMQIEVQRRLHEQLEVQRHLQLRIEAQGKYLQSVLEK
AQETLGKQNLGSAGLEAAKVQLSELVS
KVSNECFSSAFPGVEEISSLHTLQPHQAQLADLCQLSSCG
>AofPSR_AsparagusV1_10.1900
MSSSLPVLPITSGENYPKLPDSQQVSMERELRSNPVNPHKTTFVPSNGAVGPLFSSAPGLTSDLQYSTISPNV
KQPINASFIPQSQSSCTFQSSLSNHSR
ESNEINWCQDTLQGMLDYSEEVTVENNQIQSSTIMPSDNLNQQNDWWSDLVNEDWKDILNETSGSEHQPKVVY
SAAPVSSNVSVPQSQSHQMVPSQSGEL
GAVNSPSTSAGGAPARARMRWTPELHELFVEAVNQLGGSEKATPKGVLKIMKVDGLTIYHVKSHLQKYRTARY
RPDSSEGTSEKKSNPLDEMSSLDLKTG
IEITEALRLQMEVQKRLHEQLEIQRKLQLRIEEQGRYLQMMFEKQCKSNSDKLTPSSSDPDNPNPQPSALTLA
DNGAKEKDQEISEGLGEADTVSGSQSQ
SPSRKRSRAEETTSASSA

**HRS sequences**

>PheHRS_PH01000456G0940
MGLDVAEIGLGTDLSLDLKMFASRSVGRVKDAPASAMDDCIRRLEEERGKIEVFRRELPL
CARLLADVIDVMKEEAEKKKVHREEEDKEDGDAGDKSKWMSTAQLWTGDSGRDDADSEKQ
DKERSSSEANSHGGGGFVPFKAVGSGAPAFAPPCFRKYDKANDVGMPDLSLLSPAPIKST
PAATTGAADESRRQVVGFAQAAARTAAMAPSSPNLNLQTQSQQQQQARKARRCWSPELHR
KFVAALQQLGGPQVATPKQIRELMKVDSLTNDEVKSHLQKYRLHNQRAPSSAVVGQPILF
VNPAGLWIPPEQSSSQSGSPQGPLHFSTSGIAVSSAATVSCEDEDGRSESYGWK
>PheHRS_PH01001535G0300
MGLDVGLNLKLFAAWSAGRMAAAAKEAPAVDACIKGLEEERRKIEVFRRELPLCVRLLAE
VIEVMKEEAGKRSGNGDADAKAEDGDKRKWMSTAQLWVDSRGSDADSEKEQESETTSPES
KLRGGEPVPLRTVGCGAPALPPAPYFRREDKVASTLGLPGLPLMSPAVKRPLSPVTAGDD
HRHVVTARFAAAMPPSGPRISLQTQTQTQQLQQQQEARKTRRCWSPELHRQFVAALRQLG
GPQVATPKQIREVMKVDGLTNDEVKSHLQKYRLHNRRSPGAAPASQQIVLLGDLWVTQEE
SSSQSGSPQGPLQFSGSGVAVSVATVGDSYEEDDRSEGYSRK
>PheHRS_PH01002901G0070
MGLDVAEIGVGTDLSLDLKMFAARSVGRVKDAPASAMDDCIRRLEDERGKIEVFRRELPL
CARLLADVIDVMKEEAEKKKGDRGEEEDGAGDKSKWMSTAQLWTGNSGRDDADSEKQDKE
RSSSEVKVNSHGGGAFMPFKAVGSGAPAFAPPCFRKDDKAYDVGMPDLSLLSPAPTKSTP
SAATGAADESRRQVVGFAQAAARAAAMAPSGPNHNLQTESQQQQARKARRCWSPELHRQF
VAALQQLGGPQVATPKQIRELMEVDGLTNDEVKSHLQKYRLHNRRAPSSAVVSQPILVGN
PAGLWIPPEQSCSQSGSPQGPLHFSTSGIAVSSAATVSCEEEDGRSESYGWK
>PheHRS_PH01003297G0100
MGLDLKLFAARSAGRMAAAAKDAPAVDACIRSLEEEHRKIEMFRRELPLCVRLLADVIEV
MKEEAGKRRGDGDAEAKAEDGDKRKWMSTAQLWIDSRVSDADSEKEQESESTSPESKLRG
GAPVPLRAVGCGAPALPPYFGREERVASTVGLPGLPLMSPAVKRPISPVTAGDDHRHAVT
ARFAAAMPPSGPVLSLQTQTQEQQQQQARKTRRCWSPELHRQFVATLRQLGGPQVATPKQ
IREVMKVDGLTNDEVKSHLQKYRLHNRRLPGAAPASQPIVLLGDLWVPQEQSSSQSGSPQ
GPLQFSGSGVAVSVDTVGDSYEEDDRSEGYSRK
>PheHRS_PH01005036G0050
MQEPSLHADRNRARRRCREYLLALEEERRKIQVFQRELPLCLGLVTQTIEGMRSQMDGVG
SEETVSDHGPVLEEFIPLKPSLSLSSSEEDSTHAAAPTDAGKKEEAETPERRSPPAENKK
ATPNWLQSVQLWSQEPQQSSPDKELPCKPVALNARKAGGAFHPFKKEKRTELPASSTTAA
VSSAVAGDSGDKATSDTERHKDKDSKDVEDKEKDKEEGQSQPHRKPRRCWAPELHRRFLQ
ALQQLGGSHVATPKQIRQLMKVDGLTNDEVKSHLQKYRLHTRRPSSTVQSSAAVAPPVPQ
FVVVGGIWVPPTQYAAAAAAAAAAAQPQVQLSAADALGSADPVYAPVATLPSGLQPHSQK
QSSRYSEGRRSGDTGDACSASPAVSSSSQTTSA
>PheHRS_PH01000930G0300
MGTSTSHSEAIPGLDLRLLVAKTIGNLLKEEGVASGSSEQEKAERFGECIRRLETEKAKM
EAFRKELPMSVHLVGDVIGWLEDEADQRRMPTRELFAPALAKRKVEDKAGAVMKPEVDAN
DKTDWMSSAQLWSCGNHSDTTDTSRAGSKGRGVKKPSHKVSSALPGSPTLATASEKAGNA
TPPVLAELSLSSPAIDRACPVSPNANANCNAIDQARAQGQPAAQRKARRFWSAELHRQFV
AVLQRLGGAQVATPKQIKELMNVDGLTNDEVKSHLQKYRLHTRRAAGRDQPAIGLWPQHE
QRCGASPDNTLHSSVSPEVHLQTSGSSQSRPGSPEARLQLSESSGRVVSVTAGDSYEADE
EDRRSENCSVEAQHRTPS
>PheHRS_PH01001881G0390
MDSSPSDLTLDYKPNGGGGTAAGAYPMIIPKQAPPIVDGHHLTTEQATQKLQEFLSRLEE
ERLKIDAFKRELPLCMQLLNHAMEAYRQQLEAYQMESQGGAAAARPLVLEQFIPLKNIGI
DAADKPGNPPSEKASWMVSAQLWNGPAADTAAKGPQTPKEHSEHPLDTSPKLSALDGGSG
QRNGGAFLPFSKDKAMAESAALPELALAPAEKDAIAGAGAEVDKKPYHDAGSNNVATPKQ
IRELMKVDGLTNDEVKSHLQEYATQAGAPAIYGAHPATQPHYTAAVAAQEYYQSHAAVHH
LQHHPAAAMVHRAVAPPPPPAYKAAAMVGSPESEGRGSGDGRERSESIEEEGEGEEREEE
EDDDEMAALAAKADAEENAAGAGVIKY
>PheHRS_PH01000185G0030
MGSSVQKETVGDVDLRLLAARTVTDSLRAALSRSSAGEKAARFEECIRSLEAEKAKMEVF

RRELPFSVHLIGDVIEWLKDEVEQHRRPAPEMFALASAPAAKRKAEEKVGGVKTEVDAND
KRSWMSSAQLWSCGSHNSTSNSNGGSVKKQAQKVPNAFMPLNGLPTFAKSPEKPEKAAMP
PVPDLSLSSPEIDTPCPVAAPSANSSAITDAGAQLQQPVQRKARRCWSPELHRRFLAALH
RLGGPQAATPKQIRELMKVDGLTNDEVKSHLQKYRLHTRRASDGDQQAAGLWPPPEQYTT
SQHSTSQSGSPQGPLRLTVSSRGAGDSCEGGEEEEEDGKSESYSWEMENGAKAPSSS
>PheHRS_PH01002274G0070
MDSSPSDLTLDYKPNGGGGTVAGAYPMIIGKQAPLVDHHLTTEQATQKLQESLSRLEEER
LKIDAFRRELPLCMQLLNHAMEAYRQQLEAYQMGSQGGAAAARRPLVLEEFIPLKNIGID
EADKPGNPPSEKASWMVSAQLWNGPAADTAANGPQTPKEHSEHPLDTSPEFSALDGSAGQ
RNGGAFLPFSKDKAMAESAALPKLALAPAEKDAIAGAGGEVDKMQYHDAGSNNGVASRRD
VQNGAKPTSTTPEGQAAPMPPQSHRKARRCWSPELHRRFVNALQILGGAQVATPKQIREL
MKVDGLTNDEVKSHLQKYRLHTRRPMTAPPTAAPAAPHLVVLGGIWVPPEYATQAAAPTI
YGAHPATQPHYTAAVAAQGYYQSPAAVHHPAAAMVHRAVAPPPPPAYKAATRVVSPESEG
RGSGGGRERWESIEEEREDDDDEEMAAAKADAEENARRCRSN
>PheHRS_PH01001128G0130
MGTSHSEAVPGLDLGLFAAKTVGNWLKEGVASASSEQEEAERFGECIRSLETEKAKMEAF
RKELPMSVHLVGDFIGWLKDEADQRRIPTRELFAPAPAKRKVEDKAGAVMKLEADANDKM
DWMSSAQLWSCGNHSDTTDTSKDGSKGRGVKKPSHKVPNALNGLPTLATASEKAGNATPP
VLAELSMSSPAIHGACPVSHNADANCSAIDQARAQGQPAVQRKARRCWSAELHRQFVAML
QRLGGAQVATPKQIKELMKVDGLTNDEVKSHLQKYRLHARRVSGPDQPMIGLWPKHEQRR
STSQDNTLHSSASPEVHLQTSGSSQSHPGSPEARLQLSESSRAVSVTAGDSYEEDRRSES
CSVETQHRTTS

>SbiHRS_Sobic.008G036900.1
MEMDPPPPPPLHADNRRRRRCREYLLALEEEERKIRVFQRELPLCLDLVTQTIEEMRSHM
DSVVVGSEEETVSDHGGGPVLEEFMPLKPTLSSSSSEDDDDDEHDSSRHHQHRAAGVGDD
DKKNDEASGDPETAAALASSRRPLLPQPETKKAMPDWLQSVQLWSNQQQPPPQQQHQDEL
LLPCRPVALNAYRKPGGAFQPFEKDKDKEKQRGELPLPASSSAAASSAVVGDSCCDRAGG
ATDTDTAENTNKLSIKGGKDKEAQSSSQAPGRKPRRCWAPELHRRFLQALQQLGGSHAAT
PKQIRELMKVDGLTNDEVKSHLQKYRLHTRRPNSTTAVAVQSGGTSVVAPPTAPQFVVVG
GIWVPPPEYAAAAAAAAAAAAAAAAASQPQQAPHLASTGDASGTPTATANKVYAPVATTLT
TPGLLQPRPERQSSSCSGGRRSGDACSGSPAVSSSSSQTTSA
>SbiHRS_Sobic.001G078900.1
MGLDVGEIGMGLDLGLDLRLFAARTAGGMAAAAAKGAPAGIESCIRSLEEERKKIEVFRR
ELPLCARLLADVIDELKEEAAKRGGGDADAKADDGDKRKWMSTAQLWVDSEAKSDESDKE
HQSEITSPEPKLLGGAPMPIRAVAAVPPLPPPFFRREDSSAGTCLSLVSPASKAPISPVA
TSDNASGRFCPTMPPSGSSGVTLHSQAQQQASRKARRCWSPELHRQFVAALHQLGGPQVA
TPKQIREVMQVDGLTNDEVKSHLQKYRLHNRRSPGVAPVSQPIMLVGGLWAPQEQSSSGS
PQGPLQFSGSGVAISTATVGGDSSGSSSDEDDKSEGYSRKCV
>SbiHRS_Sobic.002G016300.1
MELTFHDLPPFPSAARALLPSGTQDTTVDRFIATMGLDVVEIGMGADLSLDLRHFATKAV
RQSKDDAPAPDMDACIRRLEEERGKIEMFKRELPLCARLLADVIDVMKEEAGKKTTKRSD
RSLPAAEEDEDGAAGDKSKWMSTAQLWTGDSGREDAESEKQDKGRSLPEAKSRGGALLPF
KAAVGSGAPAFASLCLRTDDKAARVGMPDLSLLSPPATKSAAEESRRQVVGLAQAAARAA
AMAPAAPALGLQSQSQQQTAQQQQQQARKVRRCWSTELHRQFVAALNQLGGPQVATPKQI
RELMKVDGLTNDEVKSHLQKYRLHNRRAPGSGVVSQPIMLVGGLWIPQEQSSSQSGSPHG
PLHFSTASGIAVSSAATVSCEEEDGRSESYGWK
>SbiHRS_Sobic.003G046800.1
MASSPSDLTLDYKPNGNAAAYAVTIPKPQQEPLVDGHHHHHHLTTAEQATQKLREFLARL
EEEWLKIDAFKRELPLCMQLLNHAMESYRQQLEAYQMGSLQGAPARPLVLEEFMPLKNIG
IDAAAADKMGNPPSEKASWMESAQLWNGPAAATAADMAAKGPQTPKESSEHPLPIDTLCA
HDAAAAGQRNGGGAFLPFAKDKTASAAEGAALPELALAPADKDAGDAERKPYLDASSNNG
VLGARRDVVQNGVKPGTNAPEGQQAAATPPPQTHRKARRCWSPELHRRFVNALQILGGAQ
VATPKQIRELMKVDGLTNDEVKSHLQKYRLHTRRPMPAPPAPATAAPQLVVLGGIWVPPE

YATQAAGQAIYGAHPATQPHYTAAVAAQEYYPSPAAVHHLQHHPAAAAAAMVHRAAAAPQ
QAAYKAAAAAMAGSPPGSEGRGSAGGGSVGGGGGGRERSESIEEEGEGEEREEDDDDDDD
DMAAAKADGEEAAAGAGAGVKY
>SbiHRS_Sobic.008G147101.1
MMARKEDHQQQQQQLRALAARAVTDSLRAAASRCSAADRAARFRDCVRSLEAEKAKMEVF
RRELPISVHLVADVIQWLKDELAQAQHRRHPPPPPPSDLFAPATTEAAAAPAPPPQTQAQ
AQDGAAVKADDAEADANDKRSWMSSAQLWSCGSHDDSTANTNGVAAAAHSKVSTAFMPTL
ASPPTLARSPVDAACKLAGATPVPDLSLSPPPSSAAAAPSATSSAVTDADRAQRQHQQAQ
AQQQRKARRCWSPELHRRFVAALQRLGGAQVATPKQIRELMKVDGLTNDEVKSHLQKYRL
HTRRASSDVGGGDHLAAAAVGGLWSSSGAAEQQYATSQHSTSQSQSGSPQGPLQLTTVSS
RAMSATAGDSCDGDGDEAEGGGRSESYGWEMQQQQQQHGTKASSSS
>ZmaHRS_GRMZM2G173882
MGLDVVEIGMGADLSLDLRHFASKAVRQSKDYAPAPAPDMDACIRRLEEERGKIEMFKRE
LPLCARLLADVIDVMKEEAGKAVAEEDGAAGDKSKWMSTAQLWTGDCGREDAEPEKQDKE
RSSPEARSRAAGGAFLPLKAAVGSGAPAFAPLGLRMDDKAAARAGMPDLCLLSPPATKSA
AEESRRQVVGFAQAATRAAAATAPAAPSLSVGVQSPSQQARKARRCWSTELHRQFVAALN
QLGGPQVATPKQIRELMKVDGLTNDEVKSHLQKYRLHNRRAPGSGVVSQPIVLVGGLWIP
QEQSSSQSGSPHGPLHFSTSGIAVSSAATVSCEEEDGRSESYGWK
>ZmaHRS_GRMZM2G124540
MGLDVGGIGMGLDLGLDLGLFAARSAGGMAAAAKGAPAEIESCIRSLEEERRKIEVFRRE
LPLCVRLLADVIDELKDEAAKRGGDAEAKADDGDKRKWMSTAQLWLDSDAKSDESDKEQL
SEITSPEPKLLGGAPMPIRAVAAVPPLPPPFFRREDSSAGSGLSLVPPAAKPPIPPMSAS
DNASGRFCATMPPSGSGANLHSQAQQQARKARRCWSPELHRLFVAALHQLGGPQVATPKQ
IREVMKVDGLTNDEVKSHLQKYRLHNRRSPGVVAPVSQSVMLAGGLWAPPHQEQSSSQSG
SPQGPLQFSGSGVAATVGGDSSSSDEDDKSEGYSRKYV
>ZmaHRS_GRMZM2G159119
MEMDPPPPPPLHADRRRRFRDYLLALEEERRKIQVFQRELPLCLDLVTQTIEGMRSHMDS
VVVGSEETVSDHGGPVLEEFMPLKPTTLSSSSSQPHYQDEHDSAHYLEHRRGSAATANDV
VDADKDGEAVGDLETAAASSSRPLPHPETKKAMPDWLQSVQLWSNQQQPSASPPQHQDEL
LLPCRPVALNACRKPGGAFQPFEKEKKKKDKEEKQRAELELPLPAAASSAVVGDSCDRAG
ATDTDTDTAENNKASSNKGGNDKEAQLSSQSQAPSRKARRCWAPELHRRFLQALQQLGGS
HVATPKQIRELMNVDGLTNDEVKSHLQKYRLHTRRPNSAAAAVQSGGTSVVAPPAAPQFV
VVGGIWVPPPEYAAAVAVAAAAAAQPQQVHLAGDASGTTTTAADKVYAPVATTLTPAPRP
RPRPRPERQSSSCSGARRSGDACSGSPAVLSSSSHTASA
>ZmaHRS_GRMZM2G016370
MEMDPPPPPPLHADRRLRFRDYLLALEEERRKIQVFQRELPLCLDLVTQTIEGMRSHMDS
VVVGSEETVSDHGGPVLEEFMPLKPTTLSSSSSQPQSQDDHDSAHYLEHRRAAAATANDV
VDADKDDEALGDPETAAASSSRPLPHPETKKAMPDWLQSVQLWSNQQQPSVSPPQHQDEL
LLPCRPVALNACRKSGGAFQPFEKEKKKEDKEEKQRAELELPLPAAASSAVVGDSCDRAG
ATDTDTDTAENNKASSTKGGKDKEAQLSSQAPSRKARRCWAPELHRRFLQALQQLGGSHV
ATPKQIRELMNVDGLTNDEVKSHLQKYRLHTRRPNSAAAAVQSGGTSVVAPPAAPQFVVV
GGIWVPPPEYAAAAAAQPQQVHVHLAGDASGTTTTAADKVYAPVATTLTPAPRPRPRPGP
ERQSSSCSGARRNGDACSGSPAVSSSSSQTASA
>ZmaHRS_GRMZM2G171468
MGLDVGEIGMGLDLGLDLRLFAARSAGGMAKGAAPAGIQSCIRSLEVERRKIEVFRRELP
LCLRLLADVIDELKEEAAKRGEYDDGAATVDDGDKRKWMSTAQLWVDSDAKSDESDKEQQ
SEITSPPEPKLLGGGAPTPIRAAVSAVAVPQPLPPPLFRREDSSASSGLSLVSLSPATKA
AVPISPPVVAASGTGTASARFCGTTMPPCGSEVNNMHSQAQQQQQASRKARRCWSPELHR
RFVAALHELGGPQVATPKQIREVMQVDGLTNDEVKSHLQKYRLHNRRSSPGAAAAPVSQS
IMLVDGVWAAQEQSSGSQSGSRQGPLQFSRAGMAVGGGDDSSSSDDDEDDDKSEDGYSLK
CV
>ZmaHRS_GRMZM2G100176
MGLDVMEIGMGADLSLDLRHFASKAVRQSKDDTPAPDMDACIRRLEEERGKIEMFKRDLP
LCARLLADVIDVMKEEAGKKKTTTRRRSDRRLASAAADDEEEEADGATADKSKWMSTAQL

WTGDSGREDAESEKQDKGRCSPEARSRGALLPFKADVGSGAPAFAPLFLRTDDKAAAARV
GVPDLSSLLSPPATMPPADAGAEESRRQVVGFAQAAARAAAMAPSAPALGLQSQQQQQQ
QQARKARRCWSTELHRKFVAALDQLGGPQVATPKQIRELMKVDGLTNDEVKSHLQKYRLH
NRRAPGSGVVRQPIVLVGGLWIPQEQGSPQSGSPHGPLHHLSTSVAAVSSAATASCEEED
GRSESYGWK
>ZmaHRS_GRMZM2G348238
MASSPSDLTLDYKPNGNGAAYAVTTPKPPQETLVVDGHHHHHHLTAAEQTTQKLREFLAR
LEEERLKIDAFKRELPLCMQLLNHAMESYRQQLEAYQMGSLQGAPARPLVLEEFMPLKNI
GIDAAADKMGNPTSEKASWMESAQLWNGPAATADVAARGPQTPKESAECAHGAAGAGHGQ
RNGGGGGGAFLPFAKDKTASAAEGAALPELALAPADKDAADADRKPYLDAAAGSNNGGVL
GSRRDAVQSGVVNVKPAPNAPEGQQAAPPQTHRKARRCWSPELHRRFVNALQILGGAQVA
TPKQIRELMKVDGLTNDEVKSHLQKYRLHTRRPMPAPPAPATAAPQLVVLGGIWVPPEYA
SQAAGQAIYGAHPATQPHYTAAAAQEYYPSPAAVHHLQHHPAAAAMVHRAAAAPPPPQQA
AYSKAAAMAGSPPGSEGRGSAGGGSIGGGGGRERSESIEEEGEVEEREEDDEDDDDDMTA
NKADGEGAAAGAGVGAAMY
>ZmaHRS_GRMZM2G142748
MPPVKFNIPAPLAPTANATAAHPRVLPVLSASPCPRHRRHNPLGLPGLGVDADKDDEAVG
DLETAAASSSRRLPHPETKKAMPDWLQSVQLWSNQQQPSASPPQHQDELLLPCRPVALNA
CRKPGGAFQPFEKEKKKKDKEEKQCAELELELPVAASSAVVGDSCDRAGATDTDTDTAEN
NKASSTKGGKDKEAQLSSQAPSRKARRCWAPELHHQFLQALQQLGGSHDCLCLPRDNIKV
IILLVINGFLKKLAEKLGIALLPLMRSIVFLNGAMIFDLTIGLLLCLVLKECHVISIYLS
TCVSMVTKNKLIERRYSSLSSKTQLNSVEQAFDTKPSLVPHENGHGKSLDTKLPKLGSLP
SRVLLNGTDGDDHSDRLKQGFVTGDMVARSSNVKWDEKAAIIISTSFVYCDDVVMDKAED
EEP
>AcomHRS_Aco031310.1
MDSTERVRRCKEYVEALEAERRKIQVFQRELPLCLQLVTQTIETVKQQMESENVRSDDGP
VLEEFIPLKPSLSSTSEEESALHESKKRESSCDRKPDWLQSVQLWNQHPDLTFPVEPPRK
PIAVSAKKIGGAFLPFEREKQVPLPLSSATPQIVSCIASGSGEKEKEKEKEKEKDKEKEK
EKEKEKEKEKEKEEGQSSQSHRKARRCWSPELHRRFLDALRQLGGSQAATPKQIRELMKV
DGLTNDEVKSHLQLID
>AcomHRS_Aco009781.1
MASPSDLSLDYKPNNYPIIQKPVGEHPNQTQKIQEFLARLEEERLKIEAFKRELPLCLQL
LNNAVEAYRQQLEAYQTSQGPRPVLEEFIPVKQMSVDGLETPNGPSSEKASWMISAQLWS
NPAESKPQQAPKGGGDVSPKLGLLDAKQRNGGGGAFLPFSKEKAKGGGGVTPRGLPELAL
ASADKGEEEKKGGVAVMENGGFGGRRENGGKGVGTEQAKGGNVAEGAAAAAAVAPPPQTH
RKARRCWSPDLHRRFVNALQMLGGSQVATPKQIRELMKVDGLTNDEVKSHLQKYRLHTRR
PIPAPPASAAAAPQLVVLGGIWVPPEYATTAAGPAIYGAHPGPAHYCTAPVPQDFYPTAT
GVAPLPHHQIHPALHRSAATATVSAVPTAYRGHRGSPESDMRSGGEQSMESIEDEEEEEE
EEGGEQRAAEEEGDEGVAVEEAMKAGEGDGGDDVALKF

>MacHRS_GSMUA_Achr8P18270_001
MDLLGRARRYHDYIRALKEERKKIEVFQRELPLCLQLVTQAIESVRHRAGEEEGVYEGPV
LEEFIPLKPSPTSTLSEEEKGETRKSAAAIGWDRKPEWLRSVQLWNQEPDTDLKEPPKKP
IAVSLKKIGGAFQPFDREKHVAPPAVVAVAPPASSTAGGGDGKGGGGGCADVDGSRSGGR
DKEKEGQSQPHRKARRCWSPELHRRFLHALQRLGGSHVATPKQIRELMKVDGLTNDEVKS
HLQKYRLHTRRPTPTVQSSSSSTSSPPATQFVLVGGILVSPPDYAAASPGNGAYAPPDSI
YAPLASVPSDSKLRNQQLHQQSQRSIARPSDTEDRSGGEDDNSMRDDGDATNSASPTTSA
TSQTTTASPLF
>MacHRS_GSMUA_Achr6P34680_001
MGSAAAEFGLDFKLCAMRTVGGFLKEAAAAVQSSDGGVAKLENSVRSLEEERRKIEAFKR
ELPLCMLLLTDVIEGLKKELERCRGDRFVHVFQELMPIRSRCEEEGGAKLEVDCKDKKKW
MSSAQLWSDYSSDDNRNDDDQSVADERDGGPDRQEEEENLCLESKSGADTFVPFKGMAAL
VMSSKQVTPTAGLSDLSLRSPAAGSASFPVSVVAGNHPGSGSVSKCVGRAPTSTAGAHLS
LQLQQQRKARRCWSPELHRRFVFALQQLGGAQVATPKQIRELMKVDGLTNDEVKSHLQKY

RLHTRKMPNASPPASRPVVVLGGLWVPPENHTAPPQKSVSQSGSPQSPLQLADARRAISP
TAGDSCEEEGGKSECHNWR
>MacHRS_GSMUA_Achr9P26570_001
MGSAVADVVLDLNLCAVRAVGGSLKEAAALESGDGRVAKLEEYVRSLEEERRKIEAFKRE
LPLCMILLTDLIEGLKQELKQCRDGRPTHVFEEVIPVKRKREDEGGLKPEADCKDKMSWM
SSAQLWSVNSREDKSDGDRNVTEERSGSRDQREEKEKEDNLFLESTSRNGRGAFVPFKGI
SGPKIKSKEETKTTVMLPDLSLLSPAGNSASSPVSATVEDHPVGGSGSKGVGRAPVSAPT
ISGAHLSLQVQQQQQQQLPRKARRCWSSELHRRFVLALEQLGGPQVATPKQIRDLMKVD
GLTNDEVKSHLQKYRLHTRKMPNATSAISQPVMVVRGLCVPDENYTTLPRSASHSGSPES
PLQLANSNHEISANAGDSCEEVDEKSENYNLR
>MacHRS_GSMUA_Achr9P09630_001
MAGIGLELRLCASRTVGGFVKQAAAVVETTDRGVAKLEESVRSLEEERRKIDAFKRELPL
CMLLLTDVIEGLKKELERCRGQKLANAFEEFIPIRRKCEEEAGVKLEADYEDKKNWMSSA
QLWGVNSSENNDEDDKSITDERNGRPDCDAGKESLNLESKNRSAGGAFVQFKGISALAMK
PKEEVLQQAPRKARRCWSPELHRRFVLALHQLGGVRVATPKQIRELMKVDGLTNDEVKSH
MQKYRLHSRKLPNASASFSRPPMVLGGLWVPPENHTVSPQQSDSQSGSRQSSLRLAGCDG
TISAAAGDSCEEEGNLLSTMLHFPWNNKLCRFST
>MacHRS_GSMUA_Achr10P28540_001
MGSSAAADVSLDLKLFTARTVAGFLKEALVMEIGDGRVAKLEESVRSLEEEKRKIEAFKR
ELPLCMHLLTDVIEGLEKELEKCRGERCARVFGESIPIKEKFEEEGREKVEKDCEAKMNW
MSSAQLWSDNPNRNNCKDNKNEKKVTPEGGGELKRQQKKENLFLESKSLSGGGAFVPFKG
LSAVTANGEEGKPTVALPDLSLQAPAMDNGALTLSAVTEGHFVSGSGSKGAGRAPTSAPA
TAGDHPSSQAQQPPPRKVRRCWSPELHRRFLFALQKLGGAQVATPKQIRELMKVDGLTND
EVKSHLQKYRLHARRMPNSSDAVNRRVAVAGGVWVPEEQYSNSSQQSVSQSGSPQSPLQL
AGATPVVSVTAGDSCEEEDGKSESFSWK
>MacHRS_GSMUA_Achr6P00650_001
MGSAAADASLELRLFAARSVTRSLKEAPAMEKGDGRKAKLEEYVTRLEEEKRKIEAFQRE
LPHSMHLLTDVIDGLKKELGQCKMGESCARVLGEFMPKKSKFEEKGAVNPEADCKEKRNW
MSSAQLWSDNSNRSNYEENENEKEVLGERGEELSRPRKKESLHLESKSLSGGSAFVPFKA
PPALSASTKKQEKPSITLPDLLLQASAIDNGNLTPAAVAEGRLVGDPVSEGIGRAPALAT
ANDRTSSQTQQQQPRKVRRCWSPDLHRRFLVALRLLGGAHVATPRQIRGLMEEDGLTNDE
VKSHLQKYRLRMTSSSNAKNRPAAIGGGGWVHEEQHSASLQQSVPQSGSPESPLRLPVAS
TTGDSCEEEDRKSESHNCV
>MacHRS_GSMUA_Achr2P13380_001
MGSALAEMGLELELCAMRTTVCGFVKEASAIESVGGGRATRLEASIKSLEEEKRKIEAFR
RELPLCMRLLSEVIEELRREIDRCHGESFGCIVEEFIPIKSKVGDDGGIKVESDCKDKMN
WMSSVQLWSDNYIENNDDDKAIAKEKDGVVDRRQEEQSNLECKNRRSGAALSLFKGLPPL
AARSMTEDKPTASLPELSLQSAVIKSNPDVVTPVTVDHRGGSAKGARKSPELFRATLSVQ
SQQQPPRKERRCWSQELHRRFVLAIEQLGGTQVATPKQIRELMKVEGLTNDEVKSHLQKY
RLHARKMTTSWATASHSVPVIWVREEQYTSSSQQSISQAGSPTSPLQLTAADSSEEDGKS
ESYNWK
>MacHRS_GSMUA_Achr4P28210_001
MGSAVFEMGLELELCATRVVGDFVKEVSAIESGGSGRASRLEESIKFLEEEKRKIEAFRR
EVPICMRLLREVIERLRMEIERCRCESFGHVFEEFMPLKSKVEDDGGVKVETDCRDKMNW
MSSVQLWSDNYSENDDEEKIVSNEQDGAVDHRQGKGKSLECKSRSSGVEFLPFKALSPLA
ASSEEEEEKPASALPELSLQSPAINRTLDLARRCWSQELHRRFVLAIQQLGGAQVATPKQ
IRELMKVDGLTNDEVKSHLQKYRLHTERVHAQRLPNAVATADRAVVVLGGLWDREEQYTS
SSQQSFSQSGSPMSPFQLAGSR
>MacHRS_GSMUA_Achr4P23210_001
MAVAEEMGLEMKLCAMRTVGGFLKEASAIECGDGGRAARLEESIRSLEEEKRKIEAFKRE
LPLCMHLLGEVIEGLNKEIERFRCERFGRAFEEFVPVKSKVEEGGGIKEETEYKDKRNWM
SSAKLWNDNCSQNNNDSKKHDKIKFKEVERRIFTKDGDSHRRQGKENFFSERKSRNGGGA
FVPFRGLPPFADSCKREEEKPAAALPDLSLQSPAIRNHHVSGCASRAVGNAPEMTPATVG
AHVSLQSLQQQPPRKARRCWSQELHRRFMRAIQQLGGVQVATPKQIRDRMKVDGLTNDEV

KSHLQKYRLHTRRMSNASAIPQDQHTCSSQQSVSQSNSPQSPLQLAGSALALSVTAGESL
EDDGKSESYSWK
>MacHRS_GSMUA_Achr7P00660_001
MVGGFVKEASAIGSGDGGRAARLEESIKSLEEEKRKIEAFKRELPLCMLLLSEVIEGLNK
EIQRCRGDRFGCVFEEFIPMKSKVEEDGRVKAETDCKDKKNWMSSAQLWSDNHSDNDDDD
KAHANIISEEVKKKKRNLYGYHRNFFSECKNRNGGGAFVPFKRLSSFAASSKEEEKPPPA
LPDLSLQSPAIENTPGLVTCAAEDHRPPRKARRCWSQELHRRFVLAIQQLGGAQVATPKQ
IREQMKVDGLTNDEVKSHLQKYRLHARRVHNASATADRPVVGEGLWIPQKQYAGSSRRSA
SRSGSPQSPLHLAASAHALSITAGDSSEEDGKSESCSWK
>MacHRS_GSMUA_Achr4P32500_001
MIEPAAMGSEVGLALQLCAMRTVGGFVKDAATESAGRAARLVESIKILEEEKRKIEAFQR
ELPLCMHLLGEVIDGLKKETERCGGGCFGRVLQELLPVESRVEEDGEVKVDDKMNWMSSA
QLWSDNYGENDISHENHEVMICGKKKEELFNLFLECKSPGGDGAFLPFEALSTVSASSEE
KKPAAALPDRRGCGRGPKSVGRAPESSPAPATVCGRLGFQSQHQPSRKARRCWSQELHRR
FTVAIQQLGVATPKQIREVMKVEGLTNDEVKSHLQKYRLHAGKLPNASSAMASPPGAVGG
SSCVRLQQQCISPPPQQSVSPSGSLSTEPSSAERSPELYE
>MacHRS_GSMUA_Achr6P09870_001
MVDHSGREQRCHDCIRALEEERRKIEAFQRELPLCLQLVTRAIECVREQMGDDESVNDAP
VLEEFIPLKPSLASTSSEGSGEAKKAAMVGRLETKPDWLRSVQLWDQQPDTVLKVEPPKK
PMAVSLKKTGGAFQPFEREKHATTPPAATAAASNFTTSRGGGDSSRGGEMEKKEEQSPVH
RKARRCWSPELHRLFLHALERLGGSHVATPKQIREMMEVDGLTNDEVKSHLQKYRLHTRR
PNPADQSSSSRPQFVLVGGLVVPPASPPGGNGACAPPNGIYAPVASRPSGLRSQQRPRSG
IRPSNSGGRCSEDVNSMGDDDSASPTTSASSQTTTASPPF
>MacHRS_GSMUA_Achr3P21090_001
MDLLDRVQSCHDYIRALEEERKKIEVFQRELPLCLKLVTHAIESVRQQMSDGERVSNGPV
LEEFIPLKPSFKSPSSGAVAAAIGSDRKPDWLRSVQLWNQEPDTSLKEPPKKPIAVSAKR
IGGAFQPFEREKLVVPPPASRAVAASSTNSAGGRGEGGGCGDSDGSSSGGEKEEKEKEGQ
FQSHRKTRRCWSPDLHRRFLNALQQLGGSHAATPKQIRDLMKVDGLTNDEIKSHLQKYRL
HTRRPSPAVQSSSHPAPQFVLLVPPPDYAAAAAVAAAAQPGNGACAPTNGIYAPVASLPS
DPRFQQQLQTKKQYQRSCSGGEDSTGGDDDDATNTESLATSASSQTTTASPPCKLRSGTD
SVEQVLSRSRFRI
>MacHRS_GSMUA_Achr10P19700_001
MDLLGRAQRCQDCIRALEEERKKIKAFQRELPLCLHLVTRAIESVRHVMGDDEKVNHGPV
TEELIPPMSEQRSEAKKSAAVGSEMKLRSVQLRNQEPDTEPPKKPIVAISKKIGSAFQPF
QRQKYVVPPPASSAATAIPATTDGDGGRDYGSKEGNREKKEEQLQPHRKARRSWSPELHR
CFLHALHQLGGSDVATPKQIRELMKVDGLTIDEVKSHLQKYRLHGRRRSPAVQCSSNGSL
AVSPQVVLVGGIVVPSPDYNMADAAVAAAAQPANGARAPSNGMYAPVASHPSDSRYRQKQ
PQRSITLRWR
>MacHRS_GSMUA_Achr11P13410_001
MDFSATLPSCGDYVRALEEERKKIQVFQRELPLCLQIVTHAIDSARRQMNGHATMSEDGP
ILEEFIPLKPSSSSSEDRSVDGAAKKSGSAATRADEKPDWLRSVQLWNQEADAFGKVEPP
TKPIAVNARRIGGAFHPFEREKHVAAPIPASSSTTSGKGGVGGSGVGGEEKEKEGQSQPH
RKARRCWSPELHQRFLHALKQLGGSHVATPKQIRELMKVDGLTNDEVKSHLQKYRLHNRR
PSTAVHSSSSNTTNPPIPQFVVVSGIWVPPPDYTVAAPPADAPCTPAVGVFAPAASLPSD
SRVRPQQSKQSDRSPTGPQLEKGGRSEESNSSGNAAAFNSLSPAASSSSQTTTASHLEIT
CKSFMVGRQ
>MacHRS_GSMUA_Achr1P23220_001
MDDRSEKAFWYDELIDALKEERRKIQVFRRELPLCLHLIDQTIENYMQLMMMEDTVRTSN
ETVLKEFISLKPCSGASEENKDPDPVPEKDRVRKPVAVEARKVREDAHPGDKEDDLGSEG
GEGKGGDGEDKKKSASQGKTRRWWSEDLHKRFLDALQQLGGCHAAKPKHIRELMKVDGLT
NDEVKSHLQKYRLHARRRRSTAVENPSAGVPQLVVLGGIWIPPPGYGVAAATHPVVNEPR
NGTHPSLPFPSSNSVLEHQNQYNII
>MacHRS_GSMUA_Achr4P03680_001
MDRIEYARRFGELIVALDEERRKMEAFHRELPLCLHLINQTIESYKQLMVSAETLSHELA

VKEFIPLRPRSVSSEDDTIHHQEWIRSTQLVAQEQDPLPKTVHPQKPIAIKAKMTWGDAQ
YLDKEEEIAPSESLACQDTEVASGEEGNGDGGKKKKKKKERSQPERKVKRYWSEELHKRF
LHALEQLGGCHAATPKLIRKLMKVDGLTNDEVKSHLQKYRLHARRSCPVEEISTSSNIQF
VVVRGIWIPTPDCAILTSTDAAAHAAPVIGVSETGRYADVSPLPSPLMILDRSYQQTDKH
SKGRNPRE
>MacHRS_GSMUA_Achr10P27180_001
MGSPPELSLDYKLDNYTAVAKAAGAQPEQNQKIQEFLARLEEERHKIEVFKRELPLCMQL
LNNAIEIYKQQLQERVPLKYMNVDGSDKANWMVSAQLWSPPDDAAKQQAAPPPKEADVIP
KLALDTTKQRSGGAFLPFSREKSRGAHATSAEKVVEEEKKCAELENEIVNTGRHSAADGH
GNTADGQAAAAPPTHRKARRCWSPDLHRRFVNALQMLGGSQVATPKQIRELMKVDGLTND
EVKSHLQKYRLHTRRPIPAPHAAAGAGAPPQLVVLGGIWVPPEYATSAGPAMYGAHAAPV
PREFYSPVPHHHYHHLHYPPLGLGASTPAASRGRTADSPEPEVRSGGERSESIEEEEEEV
RKRDEEEEEEEAPASEEKALVPLMVKAEDDNGGDVVLKI
>MacHRS_GSMUA_Achr6P05330_001
MASPAELSLDYKPNSYTMIQKQIGEQPDQTQKIQDFLARLEEERHKIDAFKRELPLCMQL
LNNAIECYKQQLETYQTNQGPRPVLEEFIPLKHMNMDGSDKDPSAHSEKASWMVSAQLWS
PPVDAAKQQPVPPPKETEQAFDVSPKLSLDTKQRNGGAFLPFSKEKSKAARSASRALSEL
ALASPEKVGGSSSATEGQAAAPPTHRKARRCWSPDLHRRFVNALQILGGSQVATPKQIRE
LMKVDGLTNDEVKSHLQKYRLHTRRPAPAPQAAAAAAPQLVVLGGIWVPPEYATSAAAAA
AAGPAIYGAHAAPAHYCAATPVPQEFYPPPPPVAHHHHLHPPLHRGAAAYKGRSSGSPES
EVRSGGERSESIEEEEEGEEREEDEEEEEEEEGTPSMEEKALLPLPVKAEDGNGGGNVAL
KF

>PdaHSR_PDK_30s723941g001
MDFAERARKCQEYVEALEEERRKIEVFQRELPLCLQLVTQAIESIRMQIWSSEETVTDGP
VLEEFIPLKPTSSSTEEEKGGFMEMEQERSEKKPDWLRSVQLWNQETDHSLPKGDPPKRP
IAVNAKKIGGAFHPFERGKHAPAAPAASSSTVGGGGGGGVAATPKQIREMMKVDGLTNDE
VKSHLQDCRIW
>PdaHSR_PDK_30s1094441g001
MDLAERARRCHEYVEALEEERRKIQVFPRELPLCLQLVTEGSYLSLLFVRAFCGAFILGS
IRVQMGSSEETVTDGPVLEEFIPLKPTSSSIEEEKGGSMEMDQERSDNKPDWLRSVQLWS
QDPDHSLPKAEPPKKPIAVNAKKIGGAFRPFDREKHVPAPVPSAAAPASSTTVGGGSSGE
KEKDKEGQSQPHRKARRCWSPDLHRQFIHALQQLGGSHAATPKQIRELMKVDDLTNDEVK
SHLQKYRLHSRRPSPAVQSSGTSSPPTPQYVVVGEIWVRPPDYAAAAVATAAQPANGAGG
IYAPVASLPSDLRHQQQQVKQQSHRSPTGPLHSQGRCSGDDGAANSASPTTSSSSQTTTA
SPPF
>PdaHSR_PDK_30s1025391g002
MGSPSDLSLDYKPNSYPMIQKPLGEQPDQTQKIQEFLARLEEERLKIEAFKRELPLCMQL
LNNAVETYRQQLETYQTNQAPRPVLEEFIPLKHASIDGSEKAPNAPSEKASWMVSAQLWS
AANDATRNQPMAAPPLKETEHAFDVSPKLALDTKQRNGGAFLPFSKEKNKGMGSASRGLP
ELALASSEKAVEEKQSCARRCWSPDLHRRFVNALQILGGSQVATPKQIRELMKVDGLTND
EVKSHLQKYRLHTRRPIPAPQASVAAAPQLVVLGSIWVPPELGSPESEMRSGGERSESIE
EEEEGEEREEEEEEEEEEEEEDVEPATDGKAEEKALVPLAVKAEEGGGRGGEVALKI

>PeqHRS_PEQU_26724
MALSARKRDLRAYIEALEDERRKIQMFSRELPLCLELITQAIEIRRQEMADDEYSGEVPV
VEEFISLKPTSSSSPSDKSLKLAGDSKPDWLRSAQLWAPSTVAGVETSPEESSKLRKPVA
KIGGAFQLLEKEKPCPPPASEVLESSITMENIAAPRKQRRCWSPELHRKFLDALEQLGGS
QSGTPKQIRELMKVERLTNDEVKSHLQKYRLHTRRLPPTIQSSSRISPQIPPQFVLVGGI
LVPPPEYAVAASSAEQSSAAFNGLYTRNPQIKQPKQQKQLNNLPFRPSQSYEDDNNNPAD
HESETKSDSCNMSSASQTTTLPLL
>PeqHRS_PEQU_27343
DGDKMGLLAPELGLDLKLFTKRTIFSCLKDEIQAGSRAAKLEDFARRLEEERRKIEAFKR
ELPLCMLLLADVMVLKEFIPLETRSDEYKEEEKAMNLEKESKEKMEWMSSAQLWSDNSSH

ESIDGKEDSQEVFHQRTDQFVFVSSHGRKDCDALFLESNTGRAFLPFKAMAAPRKEEEKL
SISLPDLSLMPLPIKVPCSAPITTVAAEEDNRSSGLNSKAVTIASSSPRNEHLGLQPQQQ
YQQNPQRKARRCWSPELHRRFIIALQQLGGAQVATPKQIRELMKVDGLTNDEVKSHLQKY
RLHTRRIPNSSATQVNRPVVMVGGLWVSQDQQNASQSGSPQGPLQLTCAAQALSMTGGDS
CTDDGKSESYSWK
>SpoHRS_Spipo0G0029600
MLPEGAPPLHAIEDLEEELARCGGGGAAHPKLEEFIPINRKFEEAGRSKDEEDRREKANW
MSSAQLWSDTSSTSNSGDSKNRSRQRSPEEEDDRFKGLPAASYKNGRAAAAVPRLPVLSF
LPPPGMLLPSPQQTPRKARRCWSPELHRRFVDALHRLGGLKGG
>SpoHRS_Spipo2G0120600
MGPAVRPELALDLGPPSSGGFVPQREAAPGEGRAAKLEEYVRRLEEERRKIEVFRRELPL
CMLILNDTIEGVKEELAGCRDGGGGGRPALEEFMPVNAKLEEAEGGVRDEKDFRDKVNWM
SSAQLWSDNYSANTSSTTSTTVASSSNSQKKGSERAAEDRSGGGGGGAFLPFKEMAGKEK
AAAVPLPPPDLSLLSPATKASSSGEHQRSGGSISLTRVSSRGTGMASPAGNEQPPPSHQA
APRKPRRCWSPELHRRFVNALQQLGGAQGSATPKQIRELMKVDGLTNDEVKSHLQKYRLH
TRRLPSSSSPEKHPMMVLGGLWAPPENFPLLKQATGSPQGPLHLANAGEDSGEEEDGRSE
SYSWKGHLQRPAGEETD
>SpoHRS_Spipo27G0017800
MERESSQKAWMGSPSELTLDFRPHGHSAALRAIADQSDQTQNLEDFLSRLEKERLKIEAF
KRELPLCMQLLNNAMEAYRQQLDMHHTNRSPRPVLEEFIPLKNPSVAAEAPEKSAAADDV
LSEKTSWMTTAQLWGTAEGAAAVQRTVTPPEATDDKLRTGGAFLPFSKERDGRVRGGLPE
LALAPMEKEDEKRPYSENGGKGGAAADHAKAAAADGGHATAVQTQRKPRRCWTPDLHRRF
VNALQILGGSQGEFPTTLHVATPKQIRELMNVDGLTNDEVKSHLQKYRLHTRRPISAPQA
SAAAAPPQLVVLGGIWVPPEYTASSTTAKAGQLFYGGHPAAAHYCAAAVQQEFYSPPLQP
PHYHHGTASAARAALAAPHEMKRESPESEMRRSPGSTHSTEEDEEVEGDDDDDDVEEGRP
SEAAGGKEVDDKEAAPLREDGKLSGVRLKF
>SpoHRS_Spipo18G0031200
MDLASIFSKAEGKGRSYEDYLEALEEERKKIQVFHRELPLSLHLVTYAIESCRRQVAAEL
FPPRVDFRSEHEDEELISDGAPALEEFIPMKLNSSTPEADDEESGDTAVRKRDWLKSVQL
WTPHPSEEDNRGPWIPVSEKESKSTATLLTFLSQKRPLQEPAVLSVETQRIPDATPPPQT
ASSTSETRGARGSSSSGGDEKRPQTHRKTRRCWSQELHRRFLDALNQLGGTEVATPKRIR
EIMQEDGLTNDEVKSHLQGNSGNAQQQQVVFVGGIWVPPSEYTTTIAATATVRRPMAEQA
AAAP
>SpoHRS_Spipo8G0062400
MVGSMVVEDVIADTTTVEERAWGYKDYVEALEEERRKIQVFQRELPLSLRLVTQTIDSCR
QRMAGAGCRSEVDEEETSCEGGPILEEFIPIKRRSSPGADQGEEDGHRQEQISNASHHEE
DSFSTSKRKNQDFSEKKPDWLRSVQLWSPLPDPHHDEDDETSRQTIVAADRNKVTSAFHP
FLPQKRSPPPATGTVEVATAPGLVKPAPAVSCTSDGRGGKSGGSGGEDKDAHPNRKARRC
WSPELHRRFLDALQQLGGSHVATPKQIRDLMKVEGLTNDEVKSHLQKYRLHSRRPSPAMS
HSNAANAPPHQVVFLGGLWVPPPEYAVAAAAAVAHPPPLTEGMVCPPIYAPVASLPMLNQ
HQQQQQQQAKQPHRSPSGPLPNGSRGTREDQRGVDDGATRSNSPATSSSSHTTSASHFN
RHNVAQSC
>SpoHRS_Spipo9G0028800
MGCLPASELSLELRPPSTPRDTMAPFLSAPAAAGGGGGGPGSGTELEEYLRGLEEERRKI
EVFRRELPLCVIILNDVIEDLKEEMTQRRSKKKQEDDGRMNWMSSTQLWCNNSSTTTAST
GGRENKSSAALHGSVRGPSPPSTADTTPPQLSLLPLAAFSAAEKQAPPQPAAVKPRRCWS
PELHQRFVDALEQLGGAKVATPKKVKELMKVEGLTNDEIKSHLQKFRLHAQRSAAPLSPA
NQAAVAVAAAGGSSWKDRQRGNWS

>ZmarHRS_Zosma112g00420.1
MTGLVEEKGAGVRRWREYLDALEEERKKIHVFQRELPLCLQLVTQTIESYRQQIGCEGDV
ATSSDVPVLEEFVPINRCSSVEEEESSHENESKSMKRLSCGGGLEQQHGGEGGGKSSENK
TPHWMQSVHLSIRTPNSSDEETELMKPISLDSGKERGAFHPFHRDRKIAPHVADISLTVP
TVSSTADTDTRGRRGFEGKDQGNSHRKARRCWSPELHRRFLHALKQLGGSHVATPKQIRE

LMKVDGLSNDEVKSHLQKYRLHTKRPNSTTGSCQTPQFILMGGIWVPPSEYASAADTDAP
PVTVSDATPVSRIYAPVASVPLGSSQSNAKKTDVNSQSGEGGCRTDSSSPSTSRSSLIG
>ZmarHRS_Zosma178g00480.1
MSLQKQQEYVDALEKERDKIAIFRRDLPLSLDLLDRVIKNYRGEICKVAAAMPILVVDEE
EERCVDDSTPILKEFIPLKPTVLKKLNEDDDDDEGDRLEKMSRCSWLKSAQLWNNDKKTL
SSPNKNENKFLLKHHHRDQHHQQRKERRCWSPELHRRFVHTLQILGGPHVATPKQIRQIL
QVDDLTNDEVKSHLQKYRMHAKWSNRETKINGDPKSVKPVVMWEPRGLTYSNQLLGHRIR
STAPFQH
>ZmarHRS_Zosma208g00340.1
MGRSVTTSPHIHGDHLQTSSSQKIEDFLSRLEEERVKIDAFKRELPLCMQLLNNAMEAYK
EQLESYRGEKRTSVLLEEFIPIKKNYDEIQKVSSNSNNGDGVGTGTGISNMEKSSWMTSV
QLWTNDSTTSEATVTPKPATESNGTFLPFYKVERSSSSTSNKGNLPDLSLSSLDTVKDDV
VPTEGKSNTLSEVHVPTPAASQGGGTHNHRKARRCWSPDLHERFVNALNMLGGPQLATPK
QIRELMKVNGLTNDEVKSHLQKYRLHTRRPNPLMQAPPPGAPQLVVLGGIWVPPEYAAAA
GATFYGTHPSAHYCATATANQQYPPHHLHHSHNRQTFNNRSSVMGYNVSESDVNLSERSM
KTGKEDDNEKAVATVGAVKAVVEEREEKSLVSVVDGNDVILKF
>ZmarHRS_Zosma30g00060.1
MGSNELSLDYKPNVDITLSTTSEADQPDPTHKLKQFLARLEQERLKIDVFKKELPLCMQL
LNNAMEVYRQQLEEGGEKMAVTEEFLPAVVKKNCTGDDKSDNSEPLEKANWLASVQLWTQ
SNNEVPSSPKRWNTGGAFLPFSKEKTVDKASGCSDLALAPVEDNRPSVLETIEESRDDRC
DNNISTTPNMENSGGGGGGGGQTAAGMGSSGGQNQTHRKARRCWSPDLHRRFVNALQVLGG
SQVATPKQIRELMKVEGLTNDEVKSHLQKYRLHTRPLPAPAQHAVASGHPKLVVLGGIWV
PPEYATSATVYSTHPSSATQFCTTAATPLSRGFYSSPTAPQPPQASNAHSHCSVTRHRCG
LQGTVELDKKGNREETSESTEEEIKEEVEAERKSGTSCSNGAGGGEEDSNGSDMC
>ZmarHRS_Zosma9g01150.1
MTYYVHNLDGFLHDHHRIGSVTASGDVEVCWVSATDGEIEDGRDFSGHTVANYVMALEEE
KRKIEPFKRELSLCLQLVTDLINHLKRDLVNVTPVLEHFLPIKSTPVLDQNGATQNSEKE
TRDRNKMDWMSSAQLWSDNYSDEAISNNNHDETNNGFGRVFTATAFQEEETSKKKKKKKK
KKLNTIATSDLSLLTPGGFRFFSPAKNADSVLSDNNQQQQHARRKARRCWSSDLHRRFVS
ALHQIGGSQAATPKQIRELMKVDGLTNDEVKSHLQKYRLHNRRISSCPPLSIAIDETMVV
LGNGGMWVPSSISSPAHTDKHTTFL
>ZmarHRS_Zosma33g00510.1
MGLPTTELCMGKRTVTIGLVAEDGDANGMKIEDLVRCLEDEKQKIEMFKRELPLCLGFIK
HVIDVLNDDLMKRGAQSSNARPILEQFLPIKDDKEMMVAEKDCKDNKMNWMTSVQLWKND
GDDTNICSGQNKVHRQSDVSEKVDNQPLPDLSLLTPGIVKDDSNKCFFSATNPSTRKEHL
ASLFPSPNTPSSNNHNIQQQHAYQQNCSRKARRCWSLELHQKFVKALERLGGSHVATPKQ
IREMMRVDGLTNDEVKSHLQKYRLHNQRVLLHSSSSSGNNNETEHCDEKISSCNNPSFST
HFLSERKIIN
>ZmarHRS_Zosma1g03450.1
MPSMSFDLNLAPEIVEKASEWVERRKYGHSLLQLKEHIRCLEDELAKIQPFGSDLFLVVG
IVKHVITKLKEDLRRFEDVEMEGDRIINPNDNLSTTSTTTTTTAAAIPLTCVVMADDGG
ERMSYWRSQYPSRKTRRRWCRELHRRFVDALNELGGPRVATPKNIKKLMNVRGLTNDEVK
SHLQKHRLHFKKIFGHGLMDNWTSSSKF
>Sit_Seita.9G080000.1
MGLDVAEIGMGLDLGLDLKLFAARSAGGMAAAAAKGAPAGIEACIRSLEEERRKIEVFRR
ELPLCVRLLADVIEELKDEAAKRGEDLELEMKADDGDKKKWMSTAQLWVDSDAKSKSEKE
KRSEMTSPEPKLLGSPMPIRAVPAVAPPPPPCFRGDDNAASTVGLPGLSLLPPAAKTSIS
PAPAVDEHRQNATARFSAPMSPSGPALNLHAQTQQQQQARKARRCWSPELHRQFVAALH
QLGGPQVATPKQIREVMQVDGLTNDEVKSHLQKYRLHNRRSPGVAPVSQSIMLVGGLWVP
QEQTSSQSGSPQGPLQFSGSGMAVSAATVGGDGSSSDEDDKSDESYSRK
>Sit_Seita.1G146600.1
MDPPLPSPQYAHRRRCSEYLLALEEERRKIQVFKRELPLCLQLVTQTIEGMKSQMHGVGS
EGTVSDHGPVLEEFMPLKPSLSLSSDEHESADDAAATNDVGKKEKAAETHGRQSPPTEAN
KAMPDWLQSVQLWSQEPQQQPSSPRKELLCKPVALNTRKAGDAFQPFVKEKRAEMPASPT

TAAASSAVVGDSCDKVATDTSEKHSDKEMNKDAKDMGKYSKDKEGQSQAPNRKPRRCWAP
ELHRRFLQALQQLGGSHVATPTQIRELMKVDGLTNDEVKSHLQKYRLHTRRPNSTTVVQS
TSTSAAQPAPQFVVVGGIWVPPAEYATAAAAASAAAAAQSQVQLAGDASGTANTVYAPVA
TLPSGTRQGQRQSSRCSGGRRSGDASSDSPAVSSSSHTTSA
>Sit_Seita.2G013500.1
MGLDVLEIGMGSDLSLDLRYFASKAVRQARDAPASDVDACIRRLEEERGKIEMFRRELPL
CARLLAEVIDVMKAEAGKKARSDRKAAAEEDGAAGDKSKWMSTAQLWTGDSGREDDSESE
KQDKGRSSPEAKSGGGGAFVPFKAVGSGAPAFAPLCLRVDDKAADAGMPDLSLLSPPAIK
SAPAPAGAAEESRRQVVGFAQAAARAVAMAPSAPSLTLQPQPQQTAQQQQQQARKARRTW
SPDLHRQFVAALNQLGGPQVATPKQIRELMKVDGLTNDEVKSHLQKYRLHNRRAPGSAVV
SQPIVLVGGLWIPQEQSSSQSGSPQGPLHFSTSGLAVSSAATVSCEEEDGRSESYGWK
>Sit_Seita.5G124700.1
MASSASDLTLDYKPNGNSNGGGAYAVIPKQQQETLVDGHHLTTEQTTQKLREFLARLEEE
RLKIDAFKRELPLCMHLLNHAMEAYGQQLEAYQMGSLQGAPARPLVLEEFIPLKKIGIDA
AADKMGNQPSEKASWMESAQLWNGPGAALAAADTAAKGPQTPKESSEHQLPIDTLGALDT
AAGQRNGGAFLPFAKDKAAAEAAALPELALAPAEKDAAETDRKPYLDAAGANGGLGARRD
VQNGVKPASNATDGQAPPPPQTHRKARRCWSPELHRRFVNALQILGGAQVATPKQIRELM
KVDGLTNDEVKSHLQKYRLHTRRPMPTPPAPATAAPQLVVLGGIWVPPEYATQAAGQAIY
GAHPATQPHYTAAVAAAQEYYPPPAAVHHLQHHPAAAMVHHRAAAPPPPQAAAYKAAMAG
SPPESSEGLGSAGGGSVGGGGGRERSESIEDEDEGEEREDDGDDDDDVPAAKTDGEDSTG
AAAIKY
>Sit_Seita.3G370000.1
MMARKEDHHHHHLRALAARAVTDSLRAAASRATDADRAARFEDCVRNLEAEKAKMEVFRR
ELPISVHLVADVIEWLKEELAQHRRLAPSPAPAPELFAPAPTSPAPAAKRKAAAPEGGAV
KAEADANDKRSWMSSAQLWSCGGGHDGSSSSTATNNGAAAAAKPAHKVSDAFMPLSGLPT
LARSPDDAAEKPTAVPVPELTLSSPAIDAACPAAPSATSSAVTDGGGAAQRQHQQQQRKA
RRCWSPELHRRFVAALQRLGGAQVATPKQIRELMKVDGLTNDEVKSHLQKYRLHTRRASS
SDGGGGDHHAASAAAAGLWSSAAPEQQYTTSQHSTSQSGSPQGPLQLTVSSRAMSATAGD
SCDGDEAEGGRSESYSFGMQQQHGTKAASS
>OsaHRS_LOC_Os01g08160.1
MASSSSDLTLDDHHHLTAVAAASGQATQKLQEFLSRLEEERLKIDAFKRELPLCMQLLNH
AMEAYRQQLEAYQMGSQHSAAAAAAAARAPLVLEEFIPVKNIGIDVVAADKAAAAGGNSVS
SEKASWMVSAQLWNAPASASAADTAAKGPQTPKEHSEHHPLDTSPKLITALDGGGGGGGA
FLPFSKDNAMGDGSAAAAAALPELALAPAEKAADAITIAAGEVDKKPYAHDNGVVARSRE
AQNGGKPPSTPSDGQAVPPPPQPHRKARRCWSPELHRRFVNALQILGGAQVATPKQIREL
MKVDGLTNDEVKSHLQKYRLHTRRPMPSPAPPTAATPQLVVLGGIWVPPEYATQAAGPAI
YGAHPATQPHYTAAVAAQEYYHHHHHHLQHHPAAAALVHHRAVAPPPPLPPQQQLAPPYS
AKSSASARLGSPDSDGRGSGGGGGAAASGAGRDMSESIEEEGEGEEREDDDDDDEMAATN
NAHAVDGDDDNDEINTTTTTSAGAINY
>OsaHRS_LOC_Os02g22020.1
MEVDHADRDGARRRCREYLLALEEERRKIQVFQRELPLCFDLVTQTIEGMRSQMDAAGSE
ETVSDQGPPPVLEEFIPLKPSLSLSSSEEESTHADAAKSGKKEEAETSERHSSPPPPPPE
AKKVTPDWLQSVQLWSQEEPQQPSSPSPTPTKDLPCKPVALNARKAGGAFQPFEKEKRAE
LPASSTTAAASSTVVGDSGDKPTDDDTEKHMETDKDNDKDAKDKDKEGQSQPHRKPRRCW
APELHRRFLQALQQLGGSHVATPKQIRELMKVDGLTNDEVKSHLQKYRLHTRRPSSTGQS
SAAAGVPAPPAPQFVVVGSIWVPPPEYAAAAAAQQHVQLAAAGNNASGSANPVYAPVAML
PAGLQPHSHRKQHQQQQQGQRHSGSEGRRSGDAGDGSSSSPAVSSSSQTTSA
>OsaHRS_LOC_Os03g55590.1
MGLDVGEIGMGLDLSLDLKMFAARSAVRMAAAAAKEATGVEACIRSLEEERRKIEMFRRE
LPLCARLLADVIELMKEEAGKRRKDGDDAEAKAEDGDKTKWMSTAQLWVDSRGSDADSEN
DRRSGSTSPASRLLGGAEESSSRAVAPPPYFRREERVVLRPAMPLLPPASHRSPPPAAAA
AATAAGDDHRHVVASSFATAVPSPVPAALSLQAQAQQQQQQARKSRRCWSPELHRQFVAA
LQQLGGPQVATPKQIREVMKVDGLTNDEVKSHLQKYRLHNRKSPGTASASHSIVLVGDLW
ASQEVSCSQSGSPQGPLQLSGSGVAVSAATAGDSCCEDDDKSEGYVRK

>OsaHRS_LOC_Os07g02800.2
MGLDVGEIGVGAADLSLDLKMFAAKSFGRVRGKDTTTTAMGDCIRRLEEEMGKIEVFRRE
LPLCVRLLADVIDVMKEEVEKKGGDRKEDEEDAAGDKSNWMSTAQLWTGNSGGPDAAAAD
PEKQDKVRISSEAKSNGGAFVGSGAPAFARPKQSLMRKEDMAYDVRMPDLSLLSPPASAA
AADESRRQVVGFSQAAARAAAMAASGPALSLQPQPQPAAAQQQQQARKTRRCWSPDLHRK
FVAALQQLGGPQVATPKQIRELMKVDGLTNDEVKSHLQKYRLHNRPRVPSSTIVNQPIVL
MQGLCYIPQEQSSSQSGSPEGPLHFSGSGMAGGGSSAATVSCEEEDGRSESYGWK
>OsaHRS_LOC_Os12g39640.1
MGSSSASAAQHEEAAAAAAFVLGGVDMRMLAARTATGALARAGGGEAAAAAAAAARFEDC
IRSLEAEKAKMEVFRRELPISVHLIADVIEWLKDEVEKQRLLRRRQVEAPAAAPPPEMFA
PPATAKRKSAASAAAEGVKAEADANDKRSWMSSAQLWSCGSHTSTSTSNGGSVKKQQHKV
SNAFMPLATLPAFAKSLEKADAAVPDLSLSSRVAMADAPACPAAPSATSSAVTDVAVAQR
QQAVQRKARRCWSPELHRRFVAALQRLGGPQAATPKQIRELMKVDGLTNDEVKSHLQKYR
LHTRRASDGGDGGGDHQTVGGRLWPLPPEQYTTSQHSTSQSGSPQGPLQLTVSSSHAVSV
TAGDSCDGGEEEEEDGKSGSYSWEMQNGARASSSS

>AofHRS_10.37
MDYLKALEEERKKILVFERELPLSLQLVTQAIESCKKKKHEVVGRGEEIASEGPILEEFIPLRPSCSSSDEEN
NHKNSGNSGGSDKKPDCMRSVQLWNQQ
PEPKGDVVMKPIAVNAKRIGGAFHPFEKEKVVVKTPVVAVGPVSSTTENSSGGNVKTQWECR
>AofHRS_01.3080
MDYLKALEEERKKILVFERELPLSLQLVTQAIENCKKKNDEVVGRGEEIASEGPILEEFIPLRPSCSSSDEEN
NHKNSGNSGGSDKKPDWMRSVQLWNQQ
PEPKGDVVMRPIAVNAKRIGGAFHPFEKEKVVVKTPVVAAAPASSTTENTSGGNGGSGGSGGSGSGEEREREK
EKEGTSQRKARRCWSPELHRRFLNALQ
QLGGSHVATPKQIRELMKIDGLTNDEVKSHLQKYRLHTRRPTASAVQNSTNSSPQQPQIVLVGGIWMPPPQYN
ATQNAPLNTVFAPVASHPSQQPMHQSK
QKNNRSPTSPLNSKERCEGDDNSEDAEQTNSDSSATSSSSRTTTASAPLVRVC
>AofHRS_06.1057
MSLESCQDSPKFSRSILLGTYATPTLRLNLSGRDESKSSILSPCNVLTSASLDSVVAMDYIKALEKEREKILV
FERELPLSLLLVTQAIESCKKPEVGGS
EEISSEGPILEEFIPLRPSCSSSDEENNNNHGNGWIGDGFDKKPDWLRSVQLWNPQAEPPKGDIVRRPVAVNA
RKIGGAFHPFEKEKVGVKAPAAAPASS
TTENTNEGGRGNGGSMEEKEKEGHSQRKARRCWSPELHRRFLHALQQLGGCHVATPKQIRELMKVDGLTNDEV
KSHLQKYRLHIKRPTTPTVHNNSNSNA
QQPQIVLVGSLWMPPPNYTTEAESTPSNEIYTPVASLPFEFDLRKRDTSPERDDNVEDSERPKSHSSNTSSQT
TTASLP
>AofHRS_07.3042
MAPDLGLDLKLSATKIVTTLSKEASSIQDDDKKIAKLEEYVQNLEDERRKIEVFKRELPLCMFLLNDVIGGMK
EEMERCQNEKRHVLEEFMPIKSKLDEE
RGVKSEKDCKDKMNWMSSAQLWSDNYSDDNDHINNNDIKDSKKNIESGDESDHHQESETPSLESKSKSFGGAF
VPFKSVERKEEVKVTGGLPDLSLVSRP
VSAAPEDHRSGVISTPRPIARCQLSXXXXTSASSLSSRARERLDAAGRLSFIDDLSWPFNSSGVHKSQLPNKL
EN

>BdiHRS_Bradi3g10730.1
MQMAMEEDPASDTRRRFRDYLLALEEERRKIHVFQRELPLCLDLVTQTIEGMKSQMDAVV
GSEETVSDHGPVLEEFIPLKPSLSLCSSEEESTHAVAPVNSPKKDESERARPPTPETKKA
MPDWLQSVQLWSQEPQQSCPNKELPCKPVALNATKTRGAFHPFEKEKRAEPELPASSTTA

AASSAVVGDSCGDKATSDTEVAHEKIKDTNKDMEKDSKEGQSSQQQHNRKARRCWAPELH
RRFLQALQQLGGSHVATPKQIRELMKVDGLTNDEVKSHLQKYRLHTTRRPSSTVAQSNNA
AQAAAPQFVVVGSCIWVPPQEYAAAAAQQQANNNSPAVVYAPVATMPSSSSALHVPSKKK
QTTSRGSKHSEDQRSGDASSSSGSPAVSSSSQTTSA
>BdiHRS_Bradi1g07630.1
MGLDVGEIGAPLDLGLDLKLFVARTAGRLAAAKEAPSMDACIRGLEEERRKIEVFRRELP
LCVRLLAEVIEMMKEQAGKRSEVRDAEAKAEDNDKRKWMSTAQLWVDNRGSDSDSVVQKE
QKKETTLPKPMLLGGAGGAPAPLMAVGFGAMPPPAPPSSQYFSREDKVAASTEGLPALPM
MSPVLKRPFSPGVDDRRPAPSAKFATIMPPPALSLQSQEQQARKTRRCWSPELHRQFVAA
LRQLGGPQVATPKQIREVMKVDGLTNDEVKSHLQKYRLHNQRSSGSSSSSHSIVLVGDHW
PPQEQSSSQSRSPEAEGPLQFSSSGVAVSAATVSDSSEEDDRSDGHSRK
>BdiHRS_Bradi2g04810.1
MDSSPSDLTLDYTPNGNGNGGGGGGHPMTPKQAPLVVEQHHLTAAEQASATTTVQKLQEF
LSRLDDERLKIDAFKRELPLCMQLLNQAMEAYRQQLEACQMGSSHGGTAGPGRAPLVLEE
FIPLSKNIDAADQNQQAASENPKASWMVSAQLWNGPPPSSADAPQTPKERSEHPPLDAGT
RGNGGAFLPFTKEMITTTADHSPAALLPELALAPGAEKDALVGINNGLSASGDKKPYLHH
DAGGGGNNGLAVARRDDDVLQNGSQTTTAAAAPQTNRKARRCWSPELHRRFVNALQILGG
AQVATPKQIRELMKVDGLTNDEVKSHLQKYRLHTRRPMQAPAAPPAGAPQLVVLGGIWMP
PDYAAQAAQAAAGPAAIYGAHPATQTHYTAAVSSAAAQEYYHSAAAHHHHLQQQHHHPH
PAMAHGRAMAPPAAAYKVHHPVAASPESEGRGSGGGGGGGRERSESIEEEGEGEDDEDDE
DAIAADAEAEEIKY
>BdiHRS_Bradi1g58830.1
MGLEAGESAMGGDLSLDLQAFAARTVAGRIPAASREDVLRKLEEEKGKIEVFGRELPVCV
RLIAAVIDLLKEKVESNGGDHRDEAEGSGDKSSWMSSAQLWIGGGSSKAPEKEGRSSAPE
KRALGGGLGFGPFKAVGSGGPAFLPVSLRKEAPLRIPDLPFLSSGSLKINSPAPAAAASA
SAGLQVAGFGLDAARTAAIAAPQLTLQTQPQPQQVAQQTAQQQQQARKARRCWSPELHRK
FVNALNQLGGPHATPKQIRERMQVDGLTNDEVKSHLQKYRLHTSRMVSGGPLMHHRPVVL
SGGLWMLPSEESSSLSGSPPGPLSGMAVSSAVSGEDDDGRSQSHGWM
>BdiHRS_Bradi4g03160.1
MGPSSSSAAAARKQQDQAAVLLGGGAAAAEEDNHQRRRMRAVAARAVTDSLRADEGKDKD
KAARLEECARGLQAEKAKMEVFRRELPISVTLIADVIEWLKDEVEQHRRPPVLMAPAPPS
SSPSPAAKRKQAEADASDKRSWMSSAQLWTCGSHSNGGIRKQQAQKLSNAFMPLGGGPPR
TTMAKSPERPEAAAMAVVPADLSLSSPAATADAAPSSNSSAVTTDAGAQSAQQQRKARRC
WSPELHRRFVAALQRLGGPQVATPKQIREMMKVDGLTNDEVKSHLQKYRLHTRRASSSDG
DHQHQQQSAAVWPPSEQPQYTASQHSTSKSGGSPMQLTGSSRATAGDSCDGEEEEEDGRS
ASYGWGMQLNGTMASSSS

>SlyHRS_Solyc12g006800.1
MNMASSNSELSLECKPHQSYSMLLKSFGEKNVDQTQNLEEFLTRLEEERVKIDAFKRELP
LCMQLLTNAMEASRQQLHSHRENNHIGQMRPVLEEFIPLKNNNASVELGDEKLVANNTST
IVDNKANWMTSAQLWSPPSDHQETKQQVQISTTCNKENDHHNNIGFSIASKLALDNNNSS
CPSPTTLALASSNKQNNNETLLQVEDNNKNLEIQQDSQGIGGGTNSTSTSTQQQQPHRKA
RRCWSPDLHRRFVNALQMLGGSQVATPKQIRELMKVDGLTNDEVKSHLQKYRLHTRRPSP
SPQATTAPPHLVVLGGIWVPPEYAAAAAHGGAPPTATFYGPHSTSHAPSPHYIAAPQALA
QEFYNTPQPLQTLHHQQLYHPGSHMYKPSPKAHSHSSPESDARGTGQHGDRSSESIEDGK
SESYSSENGGERKVVVHKF
>SlyHRS_Solyc02g090400.2
MGSNSKEMNIDLNFVYVPKLISDVLTEVSAMDDISKKLKKLNQFLVPLEEELTKIEAFKR
ELPLCMLLLKHAIERLKAEALLYKEKDKSPVMMEEFIPLKKGNSDEIGRVKKSNDLSDKK
NWMSSAQLWSTPVQYESFNLQNLKRSGVEDKAKEKQYQLGACKLTSERGAFLPFQRQALK
VDKKCLAVKDLSLSMAVTVGEGENRENPIDVSVKRENGPSNSNGCDGFSHDKSLQRKQRR
CWSPELHRRFVDALHQLGGPQVATPKQIRDIMQVDGLTNDEVKSHLQKYRLHVRRVPASG
CSWSTVDENGESSKNNGTQSGSPEGPLHFTGSGSAKGVSVNEEEDNKSESYNWNGQLHKS

IEGSTIRSSSH
>SlyHRS_Solyc01g108300.2
MGSIPAELSLDCRPSFIPKSITDFLRQLSLIRNVPDKLSQIDDYITRLEDEMRKIDAFKR
ELPLSVLLVKDAIVALREESMQYRKSRTEPVLEEFIPLKKSSREDTKAEITKDKDSREKM
SWMSSVQLWNSESHCENTDEAINKQQSKSELKDTQRSPEEGNSSVTEDPFQSCKTMNAAK
AFSVPFKGYSGFSVTAVRKDNKDELPGLSLHTPGITKLREDTVTSGLNSKHSGSRGGLSS
VAGCQSNMRTGSLAQQQSSRKQRRCWSPELHRRFVDALQQLGGSQVATPKQIRELMQVDG
LTNDEVKSHLQKYRLHTRRIPNSQTQPANQSGVALGNLWMSQGQYGESSKQSSSQSGSPQ
GPLHLAGSCGGTSTTVGDSMDEEDDVKSENHSWKNHVHTSGKVDV
>SlyHRS_Solyc12g011240.1
MFPLSPSPVVEIGNFQSPIIPTMQLHYDELMSLQQSQFYYQPVNDFSFLENHGPMRLFGE
QNIINNNVVSYQDPFMLNPNVLNFNVAHQVQQELGQQPNNVGSTSRVKPPRMLWTEDLHR
KFVAVVDAYGGPWVVKPRHILKEMTHLGISHCQIKNRLQSYRAKLNPNDAGGRKCKASSK
KELIQDVQEQLKNVDGGKADDDVCIPQLPTEAEKTKAQISPEDAFIEELLNWMVVKQDI
>SlyHRS_Solyc12g011260.1
MFPLPPSPVVKIGNFQSPIIPTMQLRSDELMSLQQSPFNYQPANDYSFLENHGLMRLFGG
LNTINNNVVNYQDPFMFNPNVLNFNVAQQVQQELGQQPNNVGSTSRVKPPRMLWTEDLHR
EFVAVVDAYGGPWAVKPRHILKEMAHLGISHCQIKNRLQSYRAKLNPNDAGGRKYISKAS
SKKELIQDVQEQLKNVDGGKVDDDVCVPQLPTEAKRMEAHISPEDAFIEGLLNWI
>SlyHRS_Solyc12g011220.1
MIPLPPSPAVNIGNFQSPNIPTMQHCYDELMSLQQSQFNYQPTNDFSFLENHGLMRLFGG
QNTINNNVVGYQDPFMLNPNVINFNIAHQVQQELGQQLNNVGSASRIKPPRVLWTDELHR
KFLDVINAYGGPWVVKPRHIWKEMAHLGVSHCQIKNRLQRYRAKLNPNEAGGRKCKASSN
KELIQDVQEQLNNVDGGKADDNVYVPQLPTEAERMEAHISPDDAFIEELLNWM
>SlyHRS_Solyc05g009720.2
MMSNNNFSDKMERCQQYIDALEQERTKIQVFSRELPLCLELVTQAIETYKQQLSGTTTEY
NLNAQSTECSDDEHTSSDVPILEEFIPLKSTFSHEDEDEDENDEENQSHKSKSFNNNNNN
STSSKDAKNKKSDWLRSVQLWNQTSDPTPKEELTPKKVSVVEVKKNGSGGAFHPFKKEKN
TVAAVETTPALAGVVLAATGSSTAENSGGSKKEDKDGQRKQRRCWSPELHRRFLHALQQL
GGSHVATPKQIRELMKVDGLTNDEVKSHLQKYRLHTRRPSPSSIHNNNNQQPPQFVVVGG
IWVPPPEYASMAAGAPAASGEASGVANSNGIYAPIATHPKGPLHDHVSGGTLRQHNNKSS
RSERSSERDRSHSHSDCGGVHSNSPATSSSTHTTTASPAY
>SlyHRS_Solyc12g011250.1
MFPLPPSPVVEIGNFQSSIIPTMQLHYDELMSLQQSLFYYQPANDFSFLENHGPMRLFGG
QSTINNNVFSYQDPFMLNPNVLNFNVAHQVQQELGQQPNNVGSTSRVKPPRVLWTEDLHR
KFLDVIDANGGPWAVKPRHIWKEMAHFGISHCQIKNRLQRYRAKLNPNEAGGRKCKASSN
KELIQDVQEQLKNVDGGKADDNVYVPQLLTEADMIEAHISPDNAFIEELLNWT

>MguHRS_Migut.E00311.1
MSSINGHNHSDYSEKMQRCQDYVHGLEQERSKIQVFQRELPLCLELVTQAIEACKQQLHG
QLSECSEQTSSDVVPVLEEFIPIKRAFSSFHSDEDDEEEEEQESKKLKNDDDSSNKKSDW
LRSVQLWNNQTPDLSKEDSPRKVAVTEVKRNGGGGAFHPFKKEKSTTTTTVAVVAAPPQE
GSMPPASTSSTAETGGGNKKEDKESQSQRKSRRCWSPELHRRFLQSLQQLGGSHLATPKQ
IRELMKVDGLTNDEVKSHLQKYRLHTRRPNNQSMQNNNNNNQQAPQFVVVGGIWMQPEYA
TPMAAATTTSGEAASGVANSTGIYAPVASLPPPFHHASKQRQHISDNDRGCSHSDGGVHS
NSPATSSSTHTSTASPAIY
>MguHRS_Migut.L01092.1
MQKCNDYIHALEEERSKIQVFQRDLPLCLDLVTQAIETCKQQLTHEKSGCNSEQTSNDDV
VPVLEEFMPIKRAADSDSDQDQDQSEEIRNIGNFDINSKKSDWLKSVQLWNQTPDPPSKQ
DSPRKSDVVEVKKNGAFHPFKKAKSGGGGPITGAAQPASTSSTAETGGGNGQSQNRKARR
CWSPELHRRFLQAIQQLGGGHVGTPKQIRELMKVDGLTNDEVKSHLQKYRLHTRRPSTTI
HNNNTQAPQFVVVGGIWMQPEYAAMATTATSVEESGSGGGAAASKGIYAPVAAAAPPPPF
REAKHKPSSSHSSDGGGARSYSPATSSSTHTQ

>MguHRS_Migut.B00308.1
MASLFSSELTLDCRFSPNSCYKETIGGISAQVSRSEALSEKISELVDYVNSLQDEMNKIG
GFKRELPHCMRLLKEEIGKMKEEMAELEKCNAEPPVLEEFIPLKKICTDEKDDDKVEGNN
KEKDKDVSSKEKMNWMSSVQLWNSDANNQNPDTDFNNNKLSLKPENKKKRKEEEVNRGVM
DDLSQSGKIRTVGRAFVPFKGCANFPVMTVGKEDKNVLTVPKLSLCTPEIKNSMDAITSI
SFIPKSSSSKLGSLSSTNIRSSLKSGQQQNSRKQRRCWSPELHRKFVNALQSLGGSQAAT
PKQIRDHMQVDGLTNDEVKSHLQKFRLHTRKATLKSGGMWMSQEQCGESSKQNNCQSGSP
EGPLQLGGSSVGYSSEYEDDDKSESHTHLSVRDS
>MguHRS_Migut.H00466.1
MILHPHPLHAYLSGLCNNKTTHTHTHTQRMGENYLKQVNDSSCDGGSIFVAKTISDILTE
LSMIHDFSKKMSKLDFYIHSFQEEMRKIHVFQRELPLSMHLLKDAIERLKKESLKWEERE
RVPVMEEFIPLKSESSYDEGRAKPSNISQQEAKKWMNSLHLWITPVQYEEDNNNQHSSFK
SKNKEEESSESQSIFRNREGGAFSPFKKPSGVGVGVTIKEEKKAAAVVNPTVHGGGGGLT
LSIPVEADVAEHTTTAAQDSIVKDAPDQGIMIKKTKLKEPHQDEQQRKQRRCWSPELHRV
FIDALQQLGGAQVATPKQIREVMKVEGLTNDEVKSHLQKYRLHVRKVPDPSSAPAARMNM
MGIIPWFRQDYEHGESSNRAKTNCFSPQQHLRDDMAGGSTSAAQVGSATTTEEGESIEEA
DEDDQKSKSCSWKGKSAEHPF
>MguHRS_Migut.H01532.1
MMTSPSELSFDCKPQIYSMLPKPFGEQILSCDQQTQKIEEFLARLEEERLKIDAFKRELP
LCMQLLTNAMEASRQQLQSQRTNQEERPILEQFIPLKNTSNNSCNNNISDKEANLSADKA
NWMTSAQLWNQESGEEEEEEEETKPPPSPLRTILPPQEITDFKLGFTNEGKKVINGGGGA
FLPFSSSSSMNCGEGFPELALANSSRNEEMKNSSSSSEVVKNICRRENSVAVVGSEREQV
VVSGGAAASNGSQTNRKARRCWSPDLHRRFVNALHMLGGSQVATPKQIRELMKVDGLTND
EVKSHLQKYRLHTRRPSPSPQTSVAAGPQVVVLGGIWVPPEYAAAAAAHHHSGAQAAALY
GAHAAHHLSPSFCGSQSAQPGQQELYPAIAPPPPPHNHQLHRHAIHHHQQQQQQQQLHL
YNKPPPLTQRGRNISPETDVRGGGGAGDPSESIEDGKSESGSWKADSGGGENGGERRRLD
GEESNGSGVTLKF
>MguHRS_Migut.N01205.1
MSSLIPPELTLDCGGASPETRVPTTTGNFLVGISSIQNSPEKISKLEDYVSRLDDEMKKI
DAFKRELPLCMLLLNDGEFLFFLQFPHFTPHRNFFFSIAAIAKIKEELIQCRNSITEPII
EEFIPLKKSLPKKEDDKIDSVDKMNWMSSVQLWNSDNHPNTENKKKWGGVDLIKSGNNRT
EGRAFSSFKRCADLPALSLATPEIKNSKKEEICSSDFSPKSSSSRSGSSSATNTDQSNSK
SKLPNQTSRKQRRCWSPELHRRFISALEHLGGPQSATPKQIRDLMQVHGLTNDEVKSHLQ
KYRLLARKLAPKTNTLVGIGSLWMPQQQKESSKKSNSQFGSPVGPLLLASSSRVSSIVEE
DDEKSEL
>MguHRS_Migut.K00828.1
KAIVIYQESKQKIQIFFAHTKHTLMGDRNLKTISDILAEVSRTDDFSRKLRKLEFYVGAF
EDELRKIDGFKRELPNCMRLLIDAIQRLKEEELMLFKSEGGNGRAKASNDSRERKNWMSS
VQLWTTPVQYETNYLTRNQDSILHLTSRCDQEGEGNGSKFRHKEGSFMPFKKSYDVAMVE
EERGGNPLRQPTAVPVHGLTVKGHFGPGRPKLHQQELEKKHRRCWSPELHDRFVDALNQL
GGAQIATPKQIRGIMKVEGLTNDEVKSHLQKYRIHVRKVPTLSNPVAAHSASPQGHLCLL
PRPDEEGSATGGASLEEEEEGEKTEGCVSNGGAHKHVDRNV

>RcoHRS_28883.m000751
MDYAEKMQRCHEYVEALEEEKRKIQVFQRELPLCLELVTQAIEACKRELSGTTTEYMHGQ
SECSEQTTSTDGTANGTGTRSLVLEEFIPIKRINSSSHNDNDNDDDNENEKEDNDDDEEE
EDQDSHKPNKSIRDINNDQKKKSDWLRSVQLWNQSSPDSEPPKEDLPRKAAVTEVKRNGG
AFQPFHKEKGIAKTPPSVPASATSSSAETGTGGGTSGAGNNRKEDKDGQAQRKQRRCWSP
ELHRRFLHALQQLGGSHAATPKQIRELMKVDGLTNDEVKSHLQKYRLHTRRPSPTIHNNS
NPQAPQFVVVGGIWVPPPEYAAVAATTASMETVTTAAANGIYAPVAAPLGTIPKQQRAQS
QHLQSERRGSHSERSNSPATSSSTHTTTNSPVF
>RcoHRS_29676.m001636
MELSLDLSLVYVPKTISEYLKEVSKVKDSSLKLSKLDDYVQRLEDEMRKIDAFKRELPLC

MLLLNDAIVRLKEEAMQCKELEEVVSVKGNSCRNEERELENDMIDKKNWMSSVQLWNNNN
HTNNNNFDSENQESKSETKQRSEEDDDRSTCENPTQLCNHRSKGGGFMPFKSTSGFEKKE
EKEVVSQVTGLSLMTPVSELGSRNLMSKTNGTDQFKIQNKPQQQQQQPYKKQRRCWSPEL
HRRFIDALHQLGGSQVATPKQIRELMQVDGLTNDEVKSHLQKYRLHIRKLPASSAAQANA
LWMAQNGHKDDSSKQSNSKSSSPQGPLHGCGSAKGMSSTGGDSMEVEDDDRSVSNSWNGR
QHKQAGEVDV
>RcoHRS_29780.m001324
MGSLPPELSLDFRPSYVPKTISDILNEVSLIRNVSDKVSKLDGFVNGLQEEMKKIDAFKR
ELPLCMLLLNDAILFLKSELMQCAALDHQPILEEFMPLKKHSDRDDDDDGDQDGPIKKEK
DTKDKKNWMSSVQLWNTDDHPSTHNILETKQNLQLESKTTKKGNYQYANDDTFSACKSRS
AARAFMPFKTYPDLSRKEDKHSTQELPVPGLSLLTPGIKNLREESTSTGSRVSCSRAVSS
STPNPQSNLRNGPQSPQQQTARKQRRCWSPELHRRFVSALQQLGGSQAATPKQIRELMQV
DGLTNDEVKSHLQKYRLHTRRMPSATAAATNHSLVVLGGGLRMSQDQYGDDSKAAGSQSG
SPQGPLQLAGNTGATSTTGGDSMEDDEDAKSEGYSWKSQIHRSGKVHV
>RcoHRS_29807.m000474
MASPSELTLDCKPHSYSMLLKSFGDQNDHTQKLEEFLSRLEEEKLKIDAFKRELPLCMQL
LTDAVETSRQQLQAYRANQGPRRILEEFMPLKNSTSETLDKSSNMSDKANWMTTAQLWSQ
ESNEIKPQTTFNSPKETNIGFNVSPKLGLDTKQRNGGAFLPFSKERNLCPSPTLALASTD
KELMEDKKCLETENGLSCSKRDNSSKVSNGGIMIEHGKGNGNLASCDGQPTNTGTGNNST
NTSTTTQTHRKARRCWSPDLHRRFVNALQMLGGSQVATPKQIRELMKVDGLTNDEVKSHL
QKYRLHTRRPSPSPQAPGAPTPQLVVLGGIWVPPEYATAAHGGTPTLYGTHHHPVSHAPP
PHFCAAPPVPQDFYTAAAAAGATTPPLPPPHHQLHHHTLHHQLHMYNKATSQAHSSPESD
IRGTGDRSESIEDGKSESSSWKAESGENGGERKGLAAFREDGEESNGSEITLKF
>RcoHRS_29790.m000831
MASSSNARSQFDLALGLKLSYVPKSIPNLIKDLSRDESSDEVKILILNDYLLKFREELIF
AAVFNQELPHCRLLLLDAIELLKEEIRRIKKGMGLDVADKPGDINILATRKNMSFEGKYS
IDLNNPYTDNQHQWSNPTDHHHQHDKMQDGTTLITNQAASLSLAPSPREKQDGMDLKLRV
NDRKLMKYKLKPLTQPICRNNRRSWTPELHARFVVVLHMLGGPEVATPKQIKEAMKVEGL
TNDQVKSHLQKYRLNSRRAPADSIRDPSFPSVCLEKFQDPCSWTSEFYHNQIKHWTRDPQ
ELFSLDSNK

>MesHRS_Manes.06G071500.1
MDFAEKMQRCHEYVEALEEEKRKIQVFQRELPLCLELVTQAIEACRRELSGTTTEYMHGQ
SECSEQTLSEGTRPVLEEFIPIKRTHSSSDNDNDDDDENDNDDQEQHSHKRNKTVVNNSG
SSNNKDKNSSDHRKKSDWLRSVQLWNQSPDPSPEEDVPRKAVVTELKRNGGAFQPFQKEK
TTGKSIQTIAKTPSSVPASATSSTADTGTGAGIVGGSERKEEKNGQNERKQRRCWSPELH
RRFLHALQQLGGSHAATPKQIREIMKVDGLTNDEVKSHLQKYRLHTRRPSPAIHNNSNPQ
AAPQFVVVGGIWVPPPEYAAVATTTASGETSTLAAAAANGIYAPVAAPPPTSHQMLQKQE
HTQSEQMQSEERGSHSEDGVRSNSPATSSSTHTTTNSPLF
>MesHRS_Manes.14G098400.1
MDYAEKMQRCHEYVEALEVERSKILVFKRELPLCLELITKAIEACRRELSGTTTDYLHGQ
SESSEQTSSEGTKPVLEEFIPIKKTNSSSDNDNEEQYLHKRNKNSSTNKDKNSSDHKKKS
DWLRSAQLWNQSPDPTPKEDVSRKAVETEVNRNGGAFQPFQKEKTTGKNVQTTAKNPSSV
PASATSSTTETGAGGTGGGSTRKEEKDGQSERKQRRCWSPELHRRFLHSLQQLGGPHVAT
PKQIRELMKVDGLTNDEVKSHLQKYRLHTRRPSPPIHNNSNPQAPQFVVVGGIWVQPPEY
AAVAATTASGETSTIAAAAAANGIYTPVAAPLSTAPPKLQKQLLKHSEQMQLEERGSHSD
GGFRSNSPATSSSTHTTTNSPSF
>MesHRS_Manes.12G093900.1
MELSLDLGSVYVPKTITEYLREVSKVKDSSQKLSKLDDYVKKLEDEMRKIDAFKRELPLC
MLLLNDAIVRLKEEAMQCKKLEDRAEIEVDVSVKEDYSGADGGNDMRDKKNWMSSVQLWN
TGDVNSDSKQHDSKSETKQRSEEEDDRSTCENPVQLCNYRSKGGAFMAFKALPVFDGTGR
KEEKEVVSQVTGLSLMTPVSEFGSGNLMSKSNGNIQMKIQNKPLQQQQQPSYKKQRRCW
SPELHRRFVDALQQLGGSKVATPKQIRELMQVDGLTNDEVKSHLQKYRLHIRKLPASSAA
QANGLWMVQDQCKDPSKPTISQSNSPQGPFHGCGSLKDVSSTGGDSIEAEDDDVSESHS

>MesHRS_Manes.17G094500.1
MGSLPPELSLDFRPTYVPKTISDFLNEVSIIGDISEKVSKLDGFVKGLEEEMRKIDAFKR
ELPLCMLLLNDAILFLKAESTQCAASNNQPVLEEFIPLKTNCDDDDDEQDGPIKKEKDSK
DKKNWMSSVQLWNSNDHHSTDYIFDQKQNLKLESKTTKKGNQFANEDTFQACKGRSPART
FLPFKTYSGLSRKEDNDTSEELPVPGLSLLTPGIKNLRAESGSTRISCSRAVSSSAPNPQ
PNLRNGQPSQQQTARKQRRCWSPELHRRFVNALQQLGGSQTATPKQIRELMQVDGLTNDE
VKSHLQKYRLHTRRMPPATAASANQSVVVLGGLWMSQDQYGDSSKTTSSQSGSPQGPLQL
AGNTGGTSTTGGDSMEDDEDAKSEGYSWKSHIHRSGKDDV
>MesHRS_Manes.13G128300.1
MELSLDLSLVYVPKTISEYLMEISGIKDSRRKLSKLDDYIKKLEDEMRKIDAFKRELPLC
MLLLNDAIVRLKQEALQCKELEGEDRTGKQEFVSVKENSGGDGGGNMGNDLSDKKNWMSS
VQLWNTNNINSDSKQHDSKSETKQRSEEDDDQSTCENPVQLCNYKSKGGAFMPFKALSVF
EGTERKEEKEVVSRVTDLSLMTPVSQLGSCNLISKCNANIQTKMHNKPQQLQQHQQEHPS
YKKQRRCWSPELHRRFVDALQQLGGSQVATPKQIRELMQVDGLTNDEVKSHLQKYRLHIR
KLPASSAACQASGLWMAQDHCKDPSKPSISESNSPQGPFHACGYAKGISSTGGNSGEAED
DDKSESHSWTGRLHKAGEVDV
>MesHRS_Manes.15G143500.1
MGSVPPDLSLDFRPTYVPKTISDFLKEVSLIGDVSEKVSKLDGFVKGLEEEMNKIDAFKR
ELPLCMLLLNDAILFLKAESIQYAASNNPPILEEFIPLKKNCDDDDDDDEQDGRINKEKD
SRDKKNWMSSVQLWNTNDHPSTNYIFDKKQSLKLESKITKRGNQYEKEDTFQVCNSRSAA
RAFMPFKTYSGLSRNDDNDNNSNGELPVPGLSLLTPGIKNFREESSSTSSRISCSRAVSS
SAPNPHSNLQNGPQPHQQQTARKQRRCWSPELHRRFVNALQQLGGSQAATPKQIRELMQV
DGLTNDEVKSHLQKYRLHTRRMPPATAASANQSVVVLGGLWMSKDQYGDSMKATSSQSGS
PQGPLQFAGNTGGTSTTGGDSMEDDEDAKSEGYSWKSHIHRSVKDDV
>MesHRS_Manes.14G127500.1
MASPSELTLDCRPQSYSMLLKSFGDQNDHTLKIEEFLSCLEEERLKIDAFKRELPLCMQL
LTNAMETSRQQIQAYRANQGPRPVLEEFIPLKNHASETLDRSSNISDKANWMTTAQLWSQ
DSNETKPQTTLTTSPKETNIGFNVSPKLGLDTKQRNGGAFLPFSKERNLCPSPTLALASA
DQQEMENKKCLEIENGFSCSKRENSGKLGNNNGGAIVIEQTKGTGNSSSDGQATNTGTSN
GADNTTTSTTTGSQTHRKARRCWSPDLHRRFVNALQMLGGSQVATPKQIRELMKVDGLTN
DEVKSHLQKYRLHTRRPSPSPQAPGAPTPQLVVLGGIWVPPEYATAAAAAHSGAPTLYGT
HPASHAPPPHFCAAPPVHQDFYTAAAATPSPQQPPHHHLHSTLHHQLHMYKATSQGHSSP
ESDVRGTGDRSESIEDGKSESSSWKAESGENGGERKSWLH
>MesHRS_Manes.06G060000.1
MASPSELTLDCKPHSYSMLLKSFGDQNDHQTQKLEEFLSRLEEERLKIDAFKRELPLCMQ
LLTNAVETSRQQLQSYRANQGPRPVLEEFIPLKNSTSETLDNSSNISDKANWMTTAQLWS
QESNETKLQTTLNTFPKKTNIGFSVSPKISLDTKQRNGGAFLPFSKERNLCPSPILALAS
TEPEMEDQKCLETENGFSCPKRGNSEKIGNNNGGVVVIEQGKGTGNSSSDGQATNTGTGN
CTNNTTTSTTTSSQAHRKARRCWSPDLHRRFVNALQMLGGSQVATPKQIRELMKVDGLTN
DEVKSHLQKYRLHTRRPSPSPQATGAPARLVVLGGIWVPPEYATAAAAAHTGAPTLYGAH
PASHAPPPHFCAAPPMPQDFYTTAAAAAPSSPQPTHHQMHHHHHTLHQLHMYKATSQAHS
SPESDVRGGDRSESIEDGKSESSSWKSGENGGDRKGLASLREDGDDSNGSEISLKPLRY
>MesHRS_Manes.01G103800.1
MLSQLFLFCQEQKEMKGVQGMLHLWQKILHWSHLDNSHCTYIGAKTKCVFADEALRCLYA
FGLGLKLSYVPQSLPNAIKDLSIGESNEVKLLLLNDYLLKCQEEMTFTGVFKQELPQSRS
LLTDAIEALKEEMRKTRIAMGLESESSPEDVLATKKRGSDEQQGLDLNNPCSDQWSNTPT
HHHHNKEQICGSLSIDPSPSMEVNLTVGENQEPVSYKMKPLTQCIWKNNRRSWTPELHAR
FAVVLHLLGGPEVATPKQIRDQMQVEGLTTNQVKSHLQKYRMNWRSSPEGAADPQRRSKL
FCRNLARPPSAAADAINMEEIN

>PtrHRS_Potri.010G128900.1
MDFAEKMQRCHEYVEALEEERRKIQVFERELPLCLELVTQAIEACKRELSGTTEDHNMHG
QSECSEQTSSEGPVLEEFIPIKRTHSSDDEENDNNHDDDDHQEQQSQNDNKRNKSNSSIS

NNDHKKKSDWLRSVQLWNQSPDPPQKQDLPRKAAVTEVKRNGAGGAFQPFHREKSVGKSS
NQAISKAPPSVPASATSSIAGAVTGGTGGGGNKKEDKEKGNQRKQRRCWSPELHRRFLHS
LQQLGGSHAATPKQIRELMKVDGLTNDEVKSHLQKYRLHTRRPSPTIHTNSSQQAPQFVV
VGGIWVPPTEYAAVAATTTAGETSTISAANGIYAPIAAPPPAVPQNRQHKQSEHSQSEGR
GSHGERGGAHSNNSPATSSSTHTTTTSPVF
>PtrHRS_Potri.008G117500.1
MDFAEKIQRCGEYVEALEEERRKIQVFERELPLCLELVTQAIEACKRELSGTTTEYNMHG
QSECSEQTSSEGPVLEEFIPIKRTHSYDENDNENDDHQEQQSHDNSKRNKTSISSGNNDH
KKKSDWLRSVQLWNHSPDLPQKQDLPRKAAVTEVKRNGAGGAFQPFHREKSIGKTSNQAI
AKAPTSVPASTTSSTAVVATGGIGGGSNKKEDIDGGNQRKQRRCWSPELHRRFLHALRQL
GGSHAATPKQIRELMKVDGLTNDEVKSHLQKYRLHTRRPSPTIHNNSNQQAPQFVVVGGI
WVPPPEYAAVAATTASAETSPISAANGIYAPIAAAPPTVPQKPHHVQYEPLQSEGGGSHS
EGGAHSNNSPATSSSTHTTTT
>PtrHRS_Potri.007G039400.1
MELSLDLSLVCEPKTINECLKEVSMVKDGSQKLTKLYDYVERLEDERRKIDAFKRELPLC
MLLLNEAIARLKEEAMQCSELNDLIPLKGNSNEDGNDKKNWMSSVQLWNTNNHLDCKKQD
SKSEPKQKSEEDDDRSTCENPVQLGDHSNKGEAFVPFKALSGFEGSKRKEEKEVVSQVTG
LSPMTPVPDLSSCNWTRKSNNCSNQTKILSKSQQQAYRKQRRCWSPELHRRFVDALQQLG
GCQVATPKQIREHMQVDGLTNDEVKSHLQKYRLHLRKVPASPAAPASQDQCKDPSTGNIS
QSNSPKGSLHASGSAKATSNTGGDSMEAEDDDKSESHSWNGALHKPGEAHV
>PtrHRS_Potri.009G106600.1
MGSLSSELSLDFARPSTAKALPFLPKTIADFLKEVSVIGDSAVKVLKVDGFIKDLEEEKR
KIDAFKRELPLCMLLLNDAIQVLREEPMQRGASKNQQPVLEEFIPLKKNIDDHHENDGLI
EEKDSKDKKNWMSSVQLWNADDHHPSTDYLFDPKQNLKLESKTNKKGNQYGNEDAFQAGI
GRTAARTFMPFKACSGLSSKEEKLPVPSLSLSTPGIKSLKEESNSTGSRSSCSRSVSTIS
GPNSDSNLRNGPQSQQQTARKQRRCWSPELHRRFVNALQQLGGSQAATPKQIRELMQVDG
LTNDEVKSHLQKYRLHTRRVTPATAAAPANQSVVVLGGLWMTQDQYGDSSKATSSQSGSP
QGPLQLAVNTGGTSTTGGDSMEDDEDAKSEGYSWKSHSHRSGKDDV
>PtrHRS_Potri.004G144800.1
MGSVPPELSLDFVRPSSTTKTFTSAIKTFTFLPETITDFLKEVSMIGDAAVKGLKVDSFL
NDLEEEKRKIDAFKRELPLCMLLLNDAIQVVREELMQCGTSNNQQPVLEEFIPLKKKIDD
HGDDRESDGLIKEKDSKDKKNWMSSVQLWITDDHHPSTDYLFDAKQSFKLESKTNKKANQ
YVNEDAFQACKGRTAARTFMPFKAYPGSSRKEDDSNREELPVPALSLLTPGIKSIKEESN
STGSRSSCSRSVSTTAPNSESNLRNGPQSQQQSSRKHRRCWSPELHRQFVNALQQLGGAQ
VATPKQIRELMQVDGLTNDEVKSHLQKYRLHTRRVPPATASAPANQSAIVLGGLWMAQDQ
YGDSSKANSSKSGSPQGPLQLAVNTGGTSTTGGDSMEDDEDAKSEGYSWKSHIHRSGKDV

>PtrHRS_Potri.005G134600.1
MELSLDLSLVYVPKAISECLKEVSMVKDGSQKLPNPDDYVKRLEDERRKIDAFKRELPLC
MLLLNEAIIRLKEEAMQCKELNALVPLKGDSNEDGNDKKKWMSSVQLWNTNNNINLDCKN
QDTRSEPKQRGEEDDDRSTCENPIQLGNHGNKGGAFVPFKALSGFERSKKKEEKEVVSQV
TGLSLMTPVPKSCDFMSKSNNCGNQMKIQSKSQQQQQQRQHAYRKQRRCWSPELHRCFV
DALQQLGGYQVATPKQIRELMQVDGLTNDEVKSHLQKYRLHLRKVPASSATPANDLWKSQ
DQCEDPVMHNISESNSPKAPLHGSSSAKAASNSGGDSMEAEDDDKSESHSWNGVLHHPGE
VHV
>PtrHRS_Potri.018G074200.1
MASPSELSLDCKPHSYSMLLKSFGEQNDQTQKLEEFLSLLEEERLKIDVFKRELPLCMQL
LTNAVETSRQQLQAYRANQVPTPVLEEFIPLKTPTSEALEKTTNISDKANWMTTAQLWSQ
DSNESKPQTTLTSPKQTDIGFNVSSKLALDTKQRNGGAFLPFSKERNLCPSPTLALSSTD
KHMEFDHKKCSEAENGFSCPKRESSGNKIGNGGVVIEQAKGAVNNSSSDGQATNTATAST
TSTTQTHRKARRCWSPDLHRRFVNALHMLGGSQVATPKQIRELMKVDGLTNDEVKSHLQK
YRLHTRRPSPSPQAAGAPPPQLVVLGGIWVPPEYATAAAAAHSGGPTLYGAHQASHAPPP
HFCAAPPVSQDFYTAAATPPPPLHHHTLYHQLHLYKPTAQAHSSPESDVRGTRDRSESIE
DGKSESSSWKGESGENDGGERRGLAALREDCEESNGSEITLKF

>PtrHRS_Potri.006G155200.1
MASPSELSLDCKPHSYSMLLKSYGHQNDETDKLEEFLSRLEEERLKIDAFKRELPLCMQL
LTSAVETSRQKLQAYRGNQVPIPVLEEFIPLKTSTSETPEKTSNISDKANWMTTAQLWSQ
ESNETKPQTTLTSPILTDIGFNVSSKLILDTKQRNGGAFLPFSKERNLCPSPTLALAACT
DEHLELDHKRFSETENGFSCPKRESSGKKMVVIEQGKGAGNSSSSDGQATNTATIASAST
NTSTSQTHRKARRCWSPDLHRRFVNALHMLGGSQVATPKQIRELMKVDGLTNDEVKSHLQ
KYRLHTRRPSPSPHAAGAPAPHLVVLGSIWVPPEYATAAAAAHSGGAALYGAHPASHAPP
PHFCATPPASQDFYTAAAAPPPSLHHHTLHHQLQLFRPTSQVHSSPESDFRGSRDRSESI
EDGKSESSSWKGESGENDGGERKGRAGLREEGEESHGSKF

>TcaHRS_Thecc1EG011857t1
MINMDYAKKMRRCHEYVEALEEERRKIQVFQRELPLCLELVTQAIEACKKEISGGSTTTD
YMQGQSECSEQTSSDGPVLEEFIPIKRSSDCSEEDDEQESRKSKDHDTNTKEKNVAAADK
KKSDWLRSVQLWNNNQSPDPPLQEGAGKSGSAVEVKRNGGAFQPFHREKTVEKSVPSVGK
ANASATSTSTTESGSRGGGGGEANNGNTNSKKEEKEGQPQRKQRRCWSPELHRRFLHALQ
QLGGSHVATPKQIRELMKVDGLTNDEVKSHLQKYRLHTRRPSPTVHNNANAQAPQFVVVG
GIWVPPPEYAPMAATTASGETASVTPTNGIYTPVAAPLPKLPQPSGAIVQRPQRSQSEER
GSHSEGRVHSNSPSTSSSTHTTTDSPLF
>TcaHRS_Thecc1EG006998t1
MGSVPPELSLDFRPTFVPKTISNFLKEVSMVGNVSDKVSKVDAFVKGLEEEMRKIDAFKR
ELPLCMLLLNDAIVALKEESMQCVTRNVEPVLEEFIPLKNNKKETKHSEEDGASITTKKD
KDPNNNNYNINKDKKNWMSSVQLWNTDDDDYRSTDHKLDTKRNDEDPFQGCKNRGSARAF
MPFKPNLGLAVRKEEKEEIPVHGLTLLTPGIKNLKEESGSTGSRTSCSRAVSSSAPNAQS
NFRSGPQPLAHHLQQQQQQQTARKQRRCWSPELHRRFVNALQQLGGSQVATPKQIRELM
QVDGLTNDEVKSHLQKYRLHTRRLPPSTTTPANQSVVVLGSGLWISQDQYGESSKGSSSQ
SGSPQGPLQLAANTGGTSTTGGDSMEDDEDAKSESYSWKSHIHKPGKDDV
>TcaHRS_Thecc1EG000558t1
MELSLDLGLAYVPKSISEFLKEVSNIKNGFQRLSKISDYVKRLEDEMKKIDAFKRELPLC
MLLLKDGIERLKEEEMQCKEMSDGSVTEELMLPSKRNTEENRRANMEDDGGDMKNWMSSV
QLWNSNFNNVDHNKKANAVQELKLRSEDEDLSEKPIELCNNKSWGGAFVPFKGQDDKEIS
GLSLMTPSSELASGNPILKSDSSCRIGSGSSLYTEQNQIKFQTKSQHQQQQQQNSRKQRR
CWSPELHRRFVDALQQLGGSQVATPKQIRELMQVDGLTNDEVKSHLQKYRLHIRKLPPSS
AGQGNGLWSAQNQCSEHMKANISQSGSPQGPLLATGSAKDMSSTGGDSMEAEEDEKSDGH
SWRGGVHKQGEIDV
>TcaHRS_Thecc1EG037970t1
MASPSELTLDCKPHSYSMLLKSFGDQQIDQTQKLEEFLSRLEEERLKIDAFKRELPLCMQ
LLTNAVEASRQQLLACRANHGSRPVLEEFMPLKNSSSENSEKSQNISDKANWMTTAQLWS
QAGNETKPQSSITSPKETEIGFNVSPKLALDTKPRNGGAFLPFTKERNSCPGSALQALPD
LALASANKDMEDKRCSDTENGMSCQRRENSGKVSNGVVVIEQGRGTANTIDGQTANTNPS
ANTTQPHRKARRCWSPDLHRRFVNALQLLGGSQVATPKQIRELMKVEGLTNDEVKSHLQK
YRLHTRRPSPSPQATGAPTPPLVVLGGIWVPPEYATAAAAAYSGAPTLYGTHHPAAAHAP
PHFCAPPVPQEFYTAAATPAPPPPQLHHHAIHHQLHMYKANSQAHSSPESDVRGAGDRSE
SIEDGKSESSSWKGESGDNGGAGERKGLAALREEGEESNGSEITLKF
>TcaHRS_Thecc1EG019009t1
MASSSSLELNLSLKPSYVPKTIANLLKDLSHVDTASDKLAVLNDYINQHEEELNNIVAFK
HQLPQCMLLLMEALGTLKDEFMSIKNGMESETGRPLMEFLAIKRKHYEEEQTDVSAAEKY
DCKGKSLMRSSPQFWNSNCNNKKQKTFIEFIQAPNQNPIPFKESTFSSSLAEKGCISKEI
IRKSYSRFASVGTNNGDSRLNLRADNRTLNYHPRPLTQPIWKNNRRSWSPELHARFVEAL
NLLGGIEVATPKQIREVMQVEGLTNDQVKSHLQKYRLHCRRLPENYAKCLVKLQDPWHPQ
MYYGDPLSPEDSDENNSNPEGSLSRKS
>TcaHRS_Thecc1EG019003t1
MNPTYVDLSLSLKPSYVPKSISNLLEDLSKIDNESDKLSVLSDYICKHQEELSTVEALKR
ELPQCRLLLMDEVAIETLKEGFMNIKEKMEYQSRQPLVEYLPTMRKNRNEQEQREIWRTN

SAEEEYWNPIYAKKQKTLVLDSFQPPNLKVVQPCRDGNVEGLGLLSYKECSSNSLGSSST
GKGKEVAIDQSSRHQSLLYLKADDQTLNYHPKPLTQPIWKNDRRCWSSELHARFVEALNL
LGGIEVATPKQIRDLMQVEGLTIDQVKSHLQVEFDLIKLNNACMYSS

>GmaHRS_Glyma.01G086700.1
MQFAQKMQPLKMGFLQYIEALEEERRKIQVFPKELPLSLELVTQAIEACRQQLAGTVAEY
NLNGQSECSEQTSTDGPVFEEFIPLKKRASQDSVEEEDEDEEHFHKHKKTATDKKKSDWL
RSVQLWNPNPPPTKEDVVVPRKTDVVEVKRNGGAFQPFQREEKSGDAKASINNDASAIGK
APSSPPVPATSSTGPVRVENKKEEKGQAQRKQRRCWSQELHKRFLHALQQLGGADSATPK
QIRELMKVDGLTNDEVKSHLQKFRLHTRRSPIIHNNASSQAGPLFLVGNIFVQPPEYAAV
ATTSTASGEELTTVTTTTAPTGIYAPVAAHPPAVTHTLPIMKQKEHSHSEERPNHSVLSN
SPASSSSTHTITTSPPVPN
>GmaHRS_Glyma.02G098800.1
MQFAEKMQPLKMGFREYIEALEEERKKIQVFPKELPLSLELVTQAIEACRQQLCGTVAEY
NLNAQSECSEQTSTDGPVFEEFIIPIKKKASQASVEEDDDEEHSHKHTKTTTDKKKSDWL
RSVQLWNPNPPPTKEDVVPRKTNVVEVKRNGGGAFQPFQREEKSGVSKANNNNEASAIGK
APSSPPVPATSYTGPVRVDSKKEEKGDAQRKQRRCWSQELHKRFLHALQQLGGADSATPK
QIRELMKVDGLTNDEVKSHLQKFRLHTRRSPIIHNSASSQAGSLFLVGNIFVQPPEYATS
SASGGELTTATPAAPTGIYAPVAAHLPAGTHTSPIMKHSHSPSEEKNNHSVHSNSPASSS
STHTITTSPPA
>GmaHRS_Glyma.10G204200.1
MYFFTEKMHSLKMGFPDYIQTLEEERRKIQVFSKELPLSLELVTQAIEACKQQLSGTASE
YNLNGHSECSEQTTSTEGPVLEEFIPIKKMASSSPFCDEEDEDDEQHSHKQRVSKENNNS
DKRKSDWLRSVQLWNPDPPAEEDVNKKVPGLELKRSGNGGGAFQPFHKEERGAKTSESLS
KAPSSTSVAAASSSAAEPAAEKSLNEGHRKQRRCWSQDLHKRFLHALQQLGGADTATPKQ
IREIMNVDGLTNDEVKSHLQKYRLHTRRPSPMVHNSSNPQAAPFVLVGNIFVQSPEYAAV
ATSTASREVATVAAPAGIYAPVATHPIPVSHPPADSIKNPQFKKVQLFEHSISDERANHS
EGAVHSNSPTSSSSTHTSTSLGY
>GmaHRS_Glyma.20G186500.1
MYFFTEKMQSLKMGFPDYIQTLEEERRKIQVFSKELPLSLELVTQAIETCKQQLSGTTSE
YNLNGHSECSEQTTSTEGPVLEEFIPIKKRASSSSPCCDEDDEQHSHKQRVSKENNNSDK
RKSDWLRSVQLWNPDPPAEEDVSKIVCGVELKRSGSGGGAFQPLHKEEKSSKPSESLSKT
PSSTPVVATSSSAVEPAEEKSLNEGQRKLRRCWSQDLHKRFLHALQQLGGADSATPKQIR
ELMNVDGLTNDEVKSHLQKYRLHTRRPIPMVHNSSDPQAAPFVLVGNIFVQSPEYAAVAT
STASREVATVAAPARIYAPVATHPTPVSHPPDDSIKKPQFKKVKLFELSNSDDRANHSEG
AVHSNSPTSSSSTHTSTSPGY
>GmaHRS_Glyma.04G151000.1
MSSQVELSMDYKPYSYSTLLKSYADETETDQTHKLEEFLSRLEEERVKIDAFKRELPLCM
QLLTNAVEASRQQLQAFRSNQGTRPVREEFMPIKHPNSQESTEKTSNISDKANWMTSAQL
WSQASEGTKPQSTITSPKNGADMGFSVSPNPALDNKHRNGGAFLPFSKERNSCQGLRDLP
EVALASSEKEMEKKCELESEKCSKRENSGKGSGSCEGVVDQGKSASVASEAQTTNTTITT
TTNNTTGQTHRKARRCWSPDLHRRFVNALQMLGGSQVATPKQIRELMKVDGLTNDEVKSH
LQKYRLHTRRPSPSLQTGAPTPQLVVFGGIWVPPEYARAAAHSGGPTLCGPHPTSHVPPP
HYCAPTPMPQEFYNSAPSLSLPSPAHENILHHHHFHMYKTAPQTRSSPVSDVRSGGDRSE
TIEDGKSESGSWKAENGEKKGLAALRDQEGDESTGSEITLKF
>GmaHRS_Glyma.06G213400.1
MPSQAELSMMDYKPYSYSTLLKSFLDQTETDQTYKLEEFLSRLEEERVKIDAFKRELPLC
MQLLTNAVEASRQQLQAFRSNQGTRPVLEEFMPILKHPNSQESAEKTSNISDKANWMTSA
QLWSQASEGTKPQSTITSLPKEGADIGFSVSPKLALDNKHRNGGAFLPFSKERNSCQGLR
GLPELALASPEKEIEENKCELEAEKCSKRENPGKGGSCEGVNVDQGKSASVASEAQTANT
TTTTTNTSGQTHRKARRCWSPDLHRRFVNALQMLGGSQVATPKQIRELMKVDGLTNDEVK
SHLQKYRLHTRRPSPSLHTGGPTPQLVVLGGIWVPPEYATTTAAMTNSGDPTLYGPHPTS
QVAPPHYCAATQMPQEFYNSALPLSLLPPPHDYTLHHHHFHMYKTAAQTQNSPESDIRSS
GDRSESIVDGKSESGKWNGESGEKKGLAALRDQEGEESTGSEITLKF

>GmaHRS_Glyma.05G011500.1
MASPSELSLDCKPQSYSLLLKSFGDQTDHSYKLEEFLNRLEEERLKIDAFKRELPLCMQL
LTNAMEASRQQLQAFKVNHGAKPVLEEFIPMKHLASESSEKATNMSDKANWMTSAQLWSQ
ASSEGTKQQPPITTLKESDIGFSISPKLALDNKQRNGGGAFLPFSKERNSCQGSTLRPLP
ELVLASAEKEMEDKKRAEVEIKGVSCQSRKENSGSDGAVVDQGKGGSPVASSHAQTTTTT
TSAQTHRKARRCWSPDLHRRFVNALQMLGGSQVATPKQIRELMKVDGLTNDEVKSHLQKY
RLHTRRPSPSPQAGAAAPQLVVLGGIWVPPEYATAAHTATPTLYGAHPTSHAPPPHYCAA
TPMPQDFYTAAPPQPLLPPPPQHHNALHHLHMYKAASHGHGSPESDVPGGGERSESIEDG
KSESSSWKGESGENEGERKGIGEESNGSEITLKF
>GmaHRS_Glyma.17G119600.1
MESPSELSLDCKPQSYSLLLKSFGDQTDQTYKLEEFLSRLEEERLKIDAFKRELPLCMQL
LTNAMEASRQQLQAYKVNHGTKPVLEEFIPMKHLASDQSSEKATNMSDKANWMTSAQLWS
QASEGTKQQPTITTPKESDIGFSISPKLALDNKQRNGGGAFLPFSKERNSCQGSTLRPLP
ELALAYAEKEMEDKKLRPEVEIKGVSCQSKKENSGSDGTVVDQGKGGSPVASSHATTTTT
TSSAQTHRKARRCWSPDLHRRFVNALQMLGGSQVATPKQIRELMKVDGLTNDEVKSHLQK
YRLHTRRPSPSPQAVGAAPQLVVLGGIWVPPEYATAAHTATPTLYGAHPTSHAPPPHYCA
ATPMPQDFYTTAPPQPLLPPPPPHHNALHHLHMYKAAPHGQGSPESDVPSNGERSESIED
GKSESSSWKGENEGERKGISKGSNGSEITLKF
>GmaHRS_Glyma.01G183700.1
MVVCDFWELSLAQSQRKTPKSNIIMEPSLDLRLGFVPKPLSLFFGDVSGNRDKCDKVVTL
DGFVQRLEEELTKVEAFKRELPLCILLLNDAIARLKEEKVKCSGMQDPPLKTSSGGNENE
NSEKKNWMSSAQLWSTQKSKSRNEEDDRSVPANSINGNSCVPEKEGSQVPSFGLMARASE
LSHSNSKSVGGDTSSGSSLLRVEVQSQPQPPQHMQQNPRKQRRCWSPELHRRFVDALQQL
GGAQVATPKQIRELMQVEGLTNDEVKSHLQKYRLHVRRFPVFSIGQVDNGSWMTQDECGD
KSKGNMSQSGSPQGPLTPLLLGGAGSAKGLSSPGRNSVDAEDEQSSDCRNWKGGLHHQQL
ETDNHSL
>GmaHRS_Glyma.11G058600.1
MELSLDLSLGFVPKPLSLFFADVSANRDKVATLDGFVQRLEEELKKVEAFKRELPLCILL
LNDAIARLKEEKVKCSGMQDPPLKTSSGGNKNESSEKMNWMSSAQLWSTQKTKSRNEEDD
RSVPANPINGNSCVLEKEGSQVPRFGLMARASELSHSNSKSVGGDISSGSSLLRVEVQSQ
PQPPQHMQQNPRKQRRCWSPELHRRFVDALQQLGGAQVATPKQIRELMQVEGLTNDEVKS
HLQKYRLHVRRFPVSSTGQADNGSWMSQDESGDKSKGNNMSQSGSPQGPLTPLILGGGGG
GSAKGLSSPGQNSVDGEDEQSDCRNWKGGLHHHQLEADNQCL
>GmaHRS_Glyma.17G178500.1
MEVSLDLSLAFVPRRTVCEILGDIAKSKDGSRRMATIEDLVKRLEDEKKKIEAFKRELPL
CMILVNDAISKLKEEIKGGVRMKDEAVVEELMKLMKTNSEANGSLMIVGNESSDTKNWMN
SVQLWNVETKQRNEEGDLFVPSNPIEQKNDTNKSVSKTVMKDNKKMSQVPSLGLMSPAVL
ELNHRKTESGYGHGSSMIITSSVEIKGHHQSQQPQQNPRKQRRCWSPDLHRRFVDALQQL
GGPQVATPKQIRELMQVVGLTNDEVKSHLQKYRLHFKRPQGSSIGHANSGLCKMAQDKCG
DDKSGSPQGPLFLGGSGKGLSSSGRNSMDTEGDEESDCRNWKATTIVTFVIAISIVIITI
TIVAT
>GmaHRS_Glyma.02G135400.1
MGSVPAELSLDLRPTFIPKTITDFLRHLSDNNNPPATLHDFLARLEDELRKIHAFKRELP
LSMLLLNDAISVLKAESQKCRVARDSAPVLEEFIPLKKERGDQSEEEEEEEENDDDDDNE
CRDKRNWMSSVQLWNNNTTTATTNNNNPSDRKQLLHKLQTKKSEEGQSVVEDPFQTCSNI
NEGRRAFMPFSRYSSSSSSVPVTTVGLGAAASKEEKEESVMNRLSLLTPSSVKEGCGSRG
SRSSSNRAVSSSSPPMVQPSLRATSLQQTARKQRRCWSPELHRRFVNALQKLGGSQVATP
KQIRELMQVDGLTNDEVKSHLQKYRLHTRRVPAASSNQPVVVLGGLWMSQDQYNDSSKVS
SSGSGSPQSPLHLAAGSRGGTSPTEGDSMEDDEDARSESFSWKSHMHKAGKVDV
>GmaHRS_Glyma.07G209500.1
MGSVAAEELNLDLRSSFVPKTITDFLRHLSANNNHPATLRDFLSRLEDELRKIHAFKREL
PLSMLLLNDAISVLKVESQKCCRVARDSPPVLEEFIPLKKELGDQSEEEEENDDDKDDNE
CRDKRNWMSSVQLWNNNTTTTTTNNNNNASDRKQLLHKLQTKKSEEGQSVAEDPFQTCSN
RNGGRRAFMPFSRYSSSSSSVPVTTVGLGAASKEEKEESVRNRLSLLTPSVKEGCGSRGS

RSSSNRAVSSSPPTAQPGLRATSLQQTARKQRRCWSPELHRRFVNALQKLGGSQAATPKQ
IRELMQVDGLTNDEVKSHLQKYRLHTRRVPAASSNQPVVVLGGLWMSQDQYNDSSKVSSS
GSGSPQSPLHLAAGSRGGTSPTEGDSIEDDEDARSESYSWKSHMNKPGKVDV
>GmaHRS_Glyma.07G178500.1
MGLVIPEEELSLDLRPSFVPKTITDFLCHLSTTPNASVKVSLLDDFVHRLQLELAKIQAF
KRELPLCMFLLNDAISALKVESEKCMACKSEPVLEEFIPLKKECDQREESEKEKECRDKK
SWMSSFQLWNTDDKADINNNAYECDKKQNYGVEDKNNREERKSVAKDLFQYGGIRNGEKG
FVMPFSTYPASKEVKEDCVFNGLSLQTPGTAVKNTREGSGCRTSSCRVVSSAPSPLRQPQ
SGRKQRRCWSPELHSRFVKALEELGGSQATTPKQIRELMRVDGLTNDEVKSHLQKYRLHT
QRVPVAKAANSNRSAVALGGLWMHNESLKGRSSGSPQGPLQLATQSGEATSRTEGDNMID
DDVKSDL
>GmaHRS_Glyma.20G009800.1
MGLVVPEEELSLDLRPSFVPKTITDFLCHLSTTPNASHKVSLLDDFVHRLELELAKIQAF
KRELPLCMFLLNDAISALKVESEKCRACKSEPVFEEFIPLKKECDQRKEIEKEKECRDKK
NWMSSFQLWNNDDKADNNNNAYECDKKHNYRVENKNNGEKRKSVTKDLFQYGRIRNGEKG
FVIPFSSYPATKEEKEDCVANGLSLQTPGTAVKSTREGSGCRTSSCRVGSSAPSPLHQPQ
SSRKQRRCWSPELHSRFIKALEVLGGSQAATPKQIRELMRVDGLTNDEVKSHLQKYRLHT
QRVPVATAANFSRSAVDLGGLWMHNESLKGGSSGSPQGPLQLATQSGEATSQTEGDNMID
DDDVKSDL

>MtrHRS_Medtr5g017980.1
MEQQQQLNLDLTLASVPKTVSHFLNNVVQTKDMSQKLSMLDDLVHSLEEEMKKVLAFKRE
LPLSILLLNDAIARLNDEKEKVRLMKMDDLKENCDNKKNWMSSAQLWTNETKSKNEGDDD
RTVLHKGNGGEVFMGFTENLLKEVSQAKSFSLVSEVSHGNSKSGGGSSGSSLLRVEIQNQ
PQPPQPLQQSSRKQRRCWSSELHRRFVDALQQLGGAHAATPKQIREKMQVDGLTNDEVKS
HLQKYRLHVRRFPVSSIQEANKLALYMAHDQCEEDTSEGNFSESVSPQGPLTPLLLGGSA
QGLSSHGRNSMDAEDEQSDCRNWKSD
>MtrHRS_Medtr1g093080.1
MQRFKMGFSDYIQALEQERRKIQVFPKELPLSLELVTQAIEACRQQLSGTTTEYNLNGQS
ECSEQTTSTDGPVLEEFIPIKKRASSYSQEVFDDVEDYDQHFHHKQQKLSLDNKKKSDWL
RSVQLWNSDPSSEEDVTKKAVPVLELKRNGCGGGAFHPFHKEDRVNNTTSELLSKGQPSS
TGVAAVSSNAATVTSNNVENRKRDEKEEKRKQRRCWSQELHKRFLKALQQLGGADCATPK
QIREVMNVDGLTNDEVKSHLQKYRLHTRRPSSTNNESANSQTAAPFVLVGNIFVQQQEYG
GVASSTTTGEMTKVVAPSGIYAPVATHPQVATIKKPEFKKFEHSISEERGNNSEGAVHSN
SPTSSSSTHTTTNASTGY
>MtrHRS_Medtr5g054300.1
MVSMQFHHKMHSHKMEFRDYILALEEEKKKIQVFPRDLPLSLELVTQAIETCKQQLFGTQ
SECSEQTSTDEGLVFEEFIPIKKRALSPDCDENDEEDDDDEEQHSSHKMKSDWLRSVQLW
NPNPSSAKEDVPRKTNVVEVKRNGGAFQPFHKEEIAAEKDNALESDKAPTSSPQVPATSS
TEPVPESGSKKDDKGQRKQRRCWSQELHKRFLHALQQLGGSNSATPKQIRELMKVDGLTN
DEVKSHLQKFRLHTRRSPIIHNNSNSHTAPMFLVGNIYVQPQEYAAVATKTTVSGELTTV
TTPTGIYAPVATHPSSTTTVIKPKSKKFELSENSHSVERVVAHSNSPASSCSTHTPTTSR
C
>MtrHRS_Medtr4g113140.1
MASSSELGLDCKPHSYSMLLKSFGEQSDQSYKLEEFVSRLEEERLKIDAFKRELPLCMQL
LTNAMEASKQQLQAFRSNQGAKPILEEFIPVKQLTSSETLEKTTNNNVCDMANWMTSAQL
WSQTSELGTKQQQNSTKENNDNNNNIGFNISPKHRNGGAFLPFSKERNNSSCQGQGLPEL
ALASTQKEEDKKHVGEAEKGKTNSGNEVDNQGKGSPVASSQTQTTSNNSNQTHRKARRCW
SPDLHRRFVNALQMLGGSQVATPKQIRELMKVDGLTNDEVKSHLQKYRLHTRRPSPSAQN
GAPAAQLVVLGGIWVPPEYATAAHAGGTPTLYGGHPTSHHLTLPHYCTAPGQDQYYTTAP
PPQQLLPPPHHMHHVYKTTPHGQGSPETCGDGLDSIENGKTSESSSWKEGSSEDEGERKG
FVEESNGSEITLKF
>MtrHRS_Medtr4g086835.1
MGCVVPAELSLDLRPTFVPKTIANFLFHLSTIQTTSDKLSKLHDFLSRLEDELNKIDAFK

RELPLSMLLLNDAILVLKEELEKCTSKNSVPVLEEFIPLKKEIDQSEENKNNDRDKNNNN
ECSKDKKNWMSSVQLWNNNTTTSNNNVSDHHHHHKLNKLETTKKREEGQSVTVAEDLFQS
CSSNRNGGSAFMPFSAYSSVPVTTVTLSAPKEEKEEPVRNRLSFLTPEVKSLREGFGSRG
SRSSSNRAVSSSSPPTVQPSLRAAPLQPQQTSRKQRRCWSPELHRRFVNALQKLGGSQAA
TPKQIRELMQVDGLTNDEVKSHLQKYRLHTRRVPAASGTDQSVVVLGGLWMPQEHYNDSS
KGSSTASGSPQSPLHLATGSRGGTSPTEGDSMEDDDDEKSESYSWKSHIHRHGKVGV

>BraHRS_LOC103875351
MIKSLSNMKNYNEKREKCCEYIEALEEERRKINVFQRELPLCLELVTQAIEAYKKEISGT
TTDTLYGQPECSEQTTGECGPVLDLFLPIKQSSASTDEEEEEEEVDDVEHESHDTGIDFV
DKNMKSEWLKSVQLWNQPEAVPVSKSERSQQETQTLVEPIKENDNGAGSHQPPCYESNEK
NDYIISLATTSSGSGRQQTEAERDGSGGSGGSSSRRGQRKQRRCWSHELHIRFLNALKQL
GGPHVATPKQIRELMKVDGLTNDEVKSHLQKYRLHTRRPSQTIPNNRDSQTQHFVVVGGI
WVPQASHSKANAVASRETTTGIYGPMASPLPSEWPSHSNFGGEISEEISRCSDKGIVRCS
SPAMSSSTRTKTKDAKMS
>BraHRS_LOC103836112
MIKKLSNMDYKQKQERCGQYIEALEEERRKIHVFQRELPLCLDLVTQAIEACKRELPGTA
TENMYGQSECSEQTTGECTPVLEQFLTIKDSSTSNEEEELDDEHGNHDPDNDYEDKNMKS
DWLKSVQLWNQPDPLLPKEEGTQEKMVDTVVKKDESMKKEAMANGGERRKREAEKDGGRK
QRRCWSSQLHRRFLNALQHLGGPHVATPKQIKELMKIDGLTNDEVKSHLQKYRLHTRRPS
QTVSNNGNSQAQHLVVVEGIWVPKSDHSTGRITGGATTSGTTTRSTTGIYGAMAAPPLPQ
WPSPSNFRPSIIVEEKGSGSPSEEVVFRCSSPAMSSSTRNHYVKNI
>BraHRS_LOC103839360
MINSLSNMKNYNEKRVRCCEYIKALEEERRKINVFQRELPLCLELVTQAIETYKKEISEA
TMENLCGQSECSEQTTGECGGILDLFRPIKHSSTSLEGEEEEVDADDEHESNETGLDCDD
KNMKSEWLKSVQLWNQPDSVLSKKLERSQQETETVVEAINGNDNGTTSHQPPCYETSNGK
SGDDKSQASISGRQKMEAEKDISGGSGGIGRRKHRRCWSHELHRRFLNTLKQLGGPHVAT
PKQIRELMKVDGLTNDEVKSHLQKYRLHTRRPGQTIPNKRDSQTQHFVVVGGIWVPQASH
STANAATAGETTIGIYGPLQAEWPSQSNFGRTISEERSRCSNKGIIRCSSPAMSSSTRTK
TKDAKLS
>BraHRS_LOC103852516
MENDYAQKMQRCHEYVEALKEEQKKIQVFQRELPLCLELVTQAIEACRKELSSTSTTSEQ
CSEQTTSVSGGPVLEEFIPTKKIEENGEHESPTPEEIGNNVDKKKSDWLRSVQLWNPSPD
TNEVSNPERVVGKKAKVVEVKPNNYNNCGGFQPFQREKKRETDLQPAAVRAVALAPAVKV
VASAPATTTSSTTETCGVGREGEEHVLQQQSQSQTRRKQRRCWSPELHRRFLHALQQLGG
SHVATPKQIRDHMKVDGLTNDEVKSHLQKYRLHTRRPATTVTAQGNGNSQQPQFVVVGGI
WVPSPQDVPPPSDVANNGGGAYAPVTLQPPPQSLPPQSPKRSVERSSGQCNSQAASSSTN
TTTSSPVS
>BraHRS_LOC103831280
MMVQKMEVDHAHKMQRCHEYVEALKEEQKKIQVFQRELPLCLELVTQAIEACKKELSVTS
TTSEQYSEQTASVCGGPVLEEFIPIKKSNNEENREHESPREVDKSDVDSKKSDWLRSAHL
WNHHSQDPDMTVVVAKKARVVEVKPNSHNRGGFQPFQKEKKRVFSETDLHPAVKATTPAP
ATTTCSTTEVGGADKAGEEQIQKQQLQTQTQRKQRRCWSPELHRRFLHALQQLGGSHAAT
PKQIRDHMKVDGLTNDEVKSHLQKYRLHTRRPATAQGNGNSQQPQFVVVGGIWVPSPQDF
PPPSDVANKGDGVYAQATAQAPVAPQSPKRSVERSSGRCNSPAASSSTNTTTTSASPVS
>BraHRS_LOC103871943
MIKKIRNMDYNQEECGQYIEALEEERRKILVFQRELPLCLDLVTQAIERCTKEITETATD
NVYGQSECSEQTTGQCTPVLEQFLTIKDSSPFNEEEVEEEFEDEQGNHDPENDSEDKNMK
SDWLKSVQLWNQPDPLLPKEEQSQQMMETVVKRDESMREDAMANSGERRRRETERKQRRC
WSSQLHRRFLNALQHLGGPHVATPKQIRELMKVDGLTNDEVKSHLQKYRLHTRRPSQTVP
NNGNSQKQHFVVVGGLWVPQSDYSTGKTTGRATTSGTTTTTTTTTTTGIYGAMAAPPPPQ
WPSHFNFTPSIIVEEGSGSHSDEVVVRCSSPAMSSSTRNCFSKPTHIFKSNPK
>BraHRS_LOC103862509

MVQTDTDQRMSLDLNLYSLAKPLSQFLNDVSKIKDNHSKLFEIDEYVGKLEEERKKIDVF
KRELPLCMLLMNEAIERLKEEASSVKLEIDNKKNWMNSTQLWISKTNPQLPSTNGEEDRC
VTQTCSNNNNSNQGGAILSFNVPPRPPPLSLRTPILTDYSSRIEQSHPIQKKELRRRWSE
DLHRRFVDALQMIGGSQVATPKQIKDIMKVDGLTNDEIKSHLQKYRMHIHKHPAKILTAS
DQHGLSRSDSPQGPLVDRGLFSNNGHSSEEEEKSDGRSWKSESRRKRQGLLDLEL
>BraHRS_LOC103864545
MVQTDSDKMGLNLNLSMYSLAKPLSQFLDEVSRIKDYDSKLSEIDGYVGKLEEERRKIDV
FKRELPLCMLLLNEAIETLKEEASSSVMMASDGKLDVGEGAKVESDNKKNWMSSAQLWIS
NPNSQLQSTNEKEDRSVTHKPNQGWAFTPFNLPPPPPPLTLRTPPSEILMDYNHQFNKPM
QSLHSQKKEQRRRWSQDLHRKFVDALHSLGGSQVATPKQIRDMMKVDGLTNDEVKSHLQK
YRMHIRKHPLHPAAKTLSSSDQAVLLDKETQSLISLTRSDSPQSPLVDGCLFNNNGHSSE
DEEKSDGRSWKSESNKKRQALDLEL
>BraHRS_LOC103835011
MVQTDKDRRMSLNLNLSMYSIAKPLSQFLDEVSRIKDNHSKLSEIDVYVGKLEEERKKID
VFKRELPLCMLLLNEAIERLKEEASSVMMASNGKLDVGQGSKLENDYKKNWMSSAQLWIS
NSNSTNKEEDRCVTQTPFQTCNNLNQGGAFLPFKPPPAPLPLMTPMMDCNRTEQNHQFNK
PLQSHHHIPKKEQRRRWSQDLRRRFVDALRRIGGSQVATPKQIRDEMKVDGLTNDEVKSH
LQKYRMHIRKHPLSSSDQPRETQSLISLSRSGSPQSPLVGRGLFNNNGSHISEDDEEEEK
SDGRSWRGESDKKRQVVDFEL
>BraHRS_LOC103829805
MAKKGDNMDYTLKCVQKKKDYTLKKKRCEEYIEALVEEQKKIQVFKRDLPLCLSLVTQAI
ESYRKELSEFSRSEHIQPEGSERTTSECEGRDGCIALCEEFVPIKSCERVKDDEAENNID
KKKSEWLQLWNQSPDPQPIEDQTTTVVAAVKENCDAFQKGKLAYSQPLKAITPTPTASSS
TAETGGNKKEMEQQKQLQMHRKQRRCWSSELHRRFLHALQKLGGSHVATPKQIRDLMQVD
GLTNDEVKSHLQKYRLHTRRPATPTLTNGCENPQQQQLIVVEGIWVPSKDAVNNRVYAPV
AVQPPPRSSPSEPRSIQRCKPPTSSSSTHPLHLPLS
>BraHRS_LOC103875404
MASSSELSLDCKPQSYSMLLKSFGENFQSDQTTQKLEDLLSRLEQERLKIDAFKRELPLC
MQLLNNAVEVYKQQLEAYLANNNNQSVVTRPVLEEFIPLRNQPEKANNWMTTAQLWSQPE
TKPKSIDQTTDQSPKDELASSPKLGHVDAKQRNSGGAFHPFTKEQHLPELALSTEVKRFS
PTNEHTNDHGNDESIINSGKNNNNNINSNSNGVSSTTSQSNRKARRCWSPDLHRRFVQAL
QMLGGSQVATPKQIRELMKVDGLTNDEVKSHLQKYRLHTRRPSPSPQTSGGQGPHLVVLG
GIWVPPEYTTAHGGTPTLYHHQVHHHNGNAATQPPQHFCSSQEFYTTPPTPQPLHHQHFQ
TFNGSSAGGATSNDSAHHQLTDSAADEGKSLESGGGEGKGLAALRQEGEDQSNINGSEIT
LKF
>BraHRS_LOC103853942
MASSSELSLDCKPQRYSMLLKSFGDNFQSDQTTQKLEDLLSRLEQERLKIDAFKRELPLC
MQLLGNAVEVYRQQLEAYRENSNNSNQSVVARPVLEEFIPLRNTPEKENNKGNNWMTTAQ
LWSQPETKPKNIDPTTDQSPKDELASSPKLGHFDVKQRNGGGAFLPFSKEKTLPELALST
EVKRVSPANEHTNDHDCNGESMINGSNNNNNNNSSTTSQSNRKARRCWSPDLHRRFVQAL
QMLGGSQVATPKQIRELMKVDGLTNDEVKSHLQKYRLHTRRPSPSPQTSGGQGPHLVVIG
AISAPPEYTTAHGGTPTLYHHQVQNHQGNAAVQPPPHFCSSQEFYTARPPPQQLHHHHFQ
TCNGSSADGSASTDSAHQLTDSAADDGKSPESGGGERKGLAALREESGNQSNINGSEITL
KF
>BraHRS_LOC103868788
MGSLGDELSLGQRRHSSIPTTICRSKIMNVAVEQGCKLEEHVQKLEEEKRKIQGSELELP
LCLQMLNDAISYLKKETETDTQPLLKDFISLSKPIREEHDEELFKEKKFQLWRENDRISN
ARFSDTLEIERDEEKSSNSLYMLLSPGMTTPKVETGLGLGLTSSSMVKGRRKLVASSSVP
QPPPYLQQQSLRKQRRCWTPELHRLFVDALQQLGGPGVATPKQIREHMQEEGLTNDEVKS
HLQKYRLHIRKSDSNLEKQSVVVLGFNLWNSSQEVEESGEGGETSKRNNSQSDSPQGPLQ
LPCTTTTTCGDSSMEDAEDAKSERRY
>BraHRS_LOC103843187
MSIEDVLLDDLDAIEACKRELPEMATYNLYRRPECSEQTIGECRHVVLEKFLTIKDSSPS
NEEEDEEFDYEHGTHDPDNDSEDKNMKSDWLKSVQLWNQPEPILPKEERLQPKIIETIVE

IDESIGKYPIVNSGEERKREAEKDAGRKQRRCWSSQLHRRFFQHLGGAHVATPKQIRELM
NIGRLTNDEVKSHLQILLNTCNYFSRICKWLLTSSFVNPPTFINSL

>AthHRS1_AT1G13300
MIKKFSNMDYNQKRERCGQYIEALEEERRKIHVFQRELPLCLDLVTQAIEACKRELPEMT
TENMYGQPECSEQTTGECGPVLEQFLTIKDSSTSNEEEDEEFDDEHGNHDPDNDSEDKNT
KSDWLKSVQLWNQPDHPLLPKEERLQQETMTRDESMRKDPMVNGGEGRKREAEKDGGGGR
KQRRCWSSQLHRRFLNALQHLGGPHVATPKQIREFMKVDGLTNDEVKSHLQKYRLHTRRP
RQTVPNNGNSQTQHFVVVGGLWVPQSDYSTGKTTGGATTSSTTTTTGIYGTMAAPPPPQW
PSHSNYRPSIIVDEGSGSHSEGVVVRCSSPAMSSSTRNHYVKNN
>AthHOH1_AT3G25790
MIKNLSNMKNDNQKREKCCEYIEALEEERRKINVFQRELPLCVELVTQAIEAYKREISGT
STDNLYGQSECSEQTTGECGRILDLFIPIKHSSTSIEEEVDDKDDDDEEHQSHETDIDFD
DKNMKSEWLKSVQLWNQSDAVVSNNRQDRSQEKTETLVELIKINDEAAKKNNNIKSPVTT
SDGGSGGGGGRRGQRKNRRCWSQELHRRFLNALKQLGGPHVATPKQIRDIMKVDGLTNDE
VKSHLQKYRLHARRPSQTTPNNRNSQTQHFVVVGGIWVPQTNHSTANAVNAVASGETTGI
YGPMVSSLPSEWPRHSNFGRKISEDRSRCSNNGFFRCSSPAMSCSTRTKTKDAKIIS
>AthHOH3_AT1G25550
MMMFKSGDMDYTQKMKRCHEYVEALEEEQKKIQVFQRELPLCLELVTQAIESCRKELSES
SEHVGGQSECSERTTSECGGAVFEEFMPIKWSSASSDETDKDEEAEKTEMMTNENNDGDK
KKSDWLRSVQLWNQSPDPQPNNKKPMVIEVKRSAGAFQPFQKEKPKAADSQPLIKAITPT
STTTTSSTAETVGGGKEFEEQKQSHSNRKQRRCWSPELHRRFLHALQQLGGSHVATPKQI
RDLMKVDGLTNDEVKSHLQKYRLHTRRPATPVVRTGGENPQQRQFMVMEGIWVPSHDTTN
NRVYAPVATQPPQSSTSGERSNRGCKSPATSSTTTHTPHLLPLS
>AthHOH2_AT1G68670
MMVEMDYAKKMQKCHEYVEALEEEQKKIQVFQRELPLCLELVTQAIEACRKELSGTTTTT
SEQCSEQTTSVCGGPVFEEFIPIKKISSLCEEVQEEEEEDGEHESSPELVNNKKSDWLRS
VQLWNHSPDLNPKEERVAKKAKVVEVKPKSGAFQPFQKRVLETDLQPAVKVASSMPATTT
SSTTETCGGKSDLIKAGDEERRIEQQQSQSHTHRKQRRCWSPELHRRFLNALQQLGGSHV
ATPKQIRDHMKVDGLTNDEVKSHLQKYRLHTRRPAATSVAAQSTGNQQQPQFVVVGGIWV
PSSQDFPPPSDVANKGGVYAPVAVAQSPKRSLERSCNSPAASSSTNTNTSTPVS
>AthHOH4_AT2G03500
MASSSELSLDCKPQSYSMLLKSFGDNFQSDPTTHKLEDLLSRLEQERLKIDAFKRELPLC
MQLLNNAVEVYKQQLEAYRANSNNNNQSVGTRPVLEEFIPLRNQPEKTNNKGSNWMTTAQ
LWSQSETKPKNIDSTTDQSLPKDEINSSPKLGHFDAKQRNGSGAFLPFSKEQSLPELALS
TEVKRVSPTNEHTNGQDGNDESMINNDNNYNNNNNNNSNSNGVSSTTSQSNRKARRCWSP
DLHRRFVQALQMLGGSQVATPKQIRELMKVDGLTNDEVKSHLQKYRLHTRRPSPSPQTSG
GPGPHLVVLGGIWVPPEYTSAHGGTPTLYHHQVHHHHTNTAGPPPPHFCSSQEFYTTPPP
PQPLHHHHFQTFNGSSGGTASTDSTHHQVTDSPTVEGKSPESGGGERKGLAALREECEDH
SNINGSEITLKF
>AthHOH5_AT4G37180
MVQTETDQRMGLNLNLSIYSLPKPLSQFLDEVSRIKDNHSKLSEIDGYVGKLEEERNKID
VFKRELPLCMLLLNEEIVFLCVAIGALKDEARKGLSLMASNGKFDDVERAKPETDKKSWM
SSAQLWISNPNSQFRSTNEEEEDRCVSQNPFQTCNYPNQGGVFMPFNRPPPPPPPAPLSL
MTPTSEMMMDYSRIEQSHHHHQFNKPSSQSHHIQKKEQRRRWSQELHRKFVDALHRLGGP
QVATPKQIRDLMKVDGLTNDEVKSHLQKYRMHIRKHPLHPTKTLSSSDQPGVLERESQSL
ISLSRSDSPQSPLVARGLFSSNVGHSSEEDEEEEDEEEEKSDGRSSCRNDETKKKRQVLD
LEL
>AthHOH6_AT1G49560
MGSLGDELSLGSIFGRGVSMNVVAVEKVDEHVKKLEEEKRKLESCQLELPLSLQILNDAI
LYLKDKRCSEMETQPLLKDFISVNKPIQGERGIELLKREELMREKKFQQWKANDDHTSKI
KSKLEIKRNEEKSPMLLIPKVETGLGLGLSSSSIRRKGIVASCGFTSNSMPQPPTPAVPQ
QPAFLKQQALRKQRRCWNPELHRRFVDALQQLGGPGVATPKQIREHMQEEGLTNDEVKSH
LQKYRLHIRKPNSNAEKQSAVVLGFNLWNSSAQDEEETCEGGESLKRSNAQSDSPQGPLQ

LPSTTTTTGGDSSMEDVEDAKSESFQLERLRSP

>VviHRS_1011942001
MQRCHDYIEALEEERRKIQVFQRELPLCLELVSQAIESCRQQMSGTTQEYFHGQSECSEQ
TSSDGPVLEEFIPIKKTSDDEDEQQSHQPNDNKDKNNDKSGKKSDWLRSVQLWNQTPDPP
VKEDTPKKIPSMEVKKNGGAFHPFKRDKAVGTNPTSAPSAATSSTAETATGCSSGSRKEE
KEGQSQRKARRCWSPELHRRFLHALQQLGGSHVATPKQIRELMKVDGLTNDEVKSHLQKY
RLHTRRPNPAIQHNGNPQAPQFVVVGGIWVPPPEYTAVAATTSSGEATGVTTANGIYAPV
ASVPPSHPQGSTQRQQPMKPKKSQSEERGSHSEGGVQSNSPATSSSTHTTTTSPVF
>VviHRS_1031804001
MGSIPPELSLDLRPSYMPKTINDFLSGISTIGDVSERVTKLDEFLKRLEEEMRKIDAFKR
ELPLCMILLSDAILALKEELLRSKASNVQPVLEEFIPLKKDCDEEGGAKEEKECKDKKNW
MSSVQLWNSDDSASTDHIYDKKQDSKLEIKQRVDEENRPATEDPFQPCKSRTGGRAFLPF
KGYSGFPVATARKEDKDELPVHGLSLLTPGIKNPREESGSSGSKTSCSRGVSSSAPNLQP
NLRTGPQPPQQQTARKQRRCWSPELHRRFVNALQQLGGSQAATPKQIRELMQVDGLTNDE
VKSHLQKYRLHTRRMPTTSAAPPSNQPVVVLGSLWMPQDQYGDSSKASSSQSASPQGPLQ
LGGAAGGTSTTGGDSMEDDEDEKSESYGWKSQVHKSVKEDV
>VviHRS_1015462001
MASPSELSLDCKPHSYSLLLKSFGDQPDQTQKLEDFLARLEEERLKIEVFKRELPLCMQL
LTDAVEASRQQLQSYRANQGARPFLEEFIPLKHSTPKGSEKTSNMSDKANWMTSAQLWSP
AGDETKQQSTLTSSKEPDIGFTVSPKLGFDNKQRNGGAFLPFSKERNSCPSPNLRNCEAQ
TTAAAPTANTQTHRKARRCWSPDLHRRFVNALQMLGGSQVATPKQIRELMKVDGLTNDEV
KSHLQKYRLHTRRPSPSPQATGAAAPQLVVVGLGGIWVPPEYATAAHTGGTTLYGAHPPS
HTSPHYCGPPVPQDFYAAPAQPHHHSLHHQLHMYKASPQTQSSPDSDMRGAGDRSESIED
GKSESSSWKGESGENGGERKGLVTLRDDGEESNRAMEV
>VviHRS_1003551001
MAELRLDLNPGCELKTVSSFLREVSMMRDSSEKLSKLDDYVKGLEDEMRKIDGFKRELPL
CMLLLNDAILRLREEAMQCTESEGGHVTEEFIPLKGRSEGDEGAEMGKDSSDKRSWMSCA
QLWSCNRSSDNNNKKSVLEFQQRNEEDNNSGSENPIRPCSYGNRAGAFVPFKVSSGFPAK
GDKEVVPVAGLSLMTPMVEVDPNNSNSKCGNSGETGSGSVSTLLTDSMRLQCKSQQQPPQ
QGIRKQRRCWSPELHRRFVNALQQLGGSQAATPKQIRELMQVEGLTNDEVKSHLQKYRLH
VRRIPSTSAAPANGSWVPQVQVGDVSKKNISQSGSPQGPLHLAGSAKGVSGTGGDSMEYD
EDDKSEGHSWKDQLHKPGEDDV

>CpapHRS_1.213
MDYATKMQRCHEYVDALEEERRKIQVFQRELPLCLELVTQAIETCKKEISGTTADYMHVQ
SECSEQTMSEVGPPVLEEFIPIKRTSSPDEDEQESHKASTNTNTDKKKSDWLRSVQLWNQ
SPDPPPKKDVPKKTTTADQVKINGGAFQPFTKDQPTENCTSNKQPAGKSNNISVPAAATS
STDEKTGGKREEKEGQQTQRKQRRCWSPELHRRFLHALQQLGGSHTATPKQIRELMKVDG
LTNDEVKSHLQKYRLHTRRPTQNVQTNGNSQAPQFVVVGGIWVPPQEYATITAATAPASG
EGCNGIYAPVPVAALPTPAKERPQQTQSHRMQSEERGCSHSEGRVHSSSPSTSSSTHTTT
ASPVF
>CpapHRS_127.55
MASPSELSLDCKPHSYSMLLKSFGDQTTADHAQKLEEFLSRLEEERLKIDAFKRELPLCM
QLLTNAVEASRQQLQAYRANQGPRPVLEEFIPLKNPSSESTEKLNTISDKANWMTSAQLW
SQAGNETKPQPTTIISSSLSPKETDIGFNVSPKLALDTKQRNGGAFLPFSKERNSCPTPT
PVRAGLPELALVSREKDGMDQDSKFSDQMGESGMVRRESTGKAGNGINEQGKGPCNNNST
SEAQAGNNTATNAAQTHRKARRCWSPDLHRRFVNALQMLGGSQVATPKQIRELMKVEGLT
NDEVKSHLQKYRLHTRRPSPSPQATGAPAPQLVVLGGIWVPPEYATAHGGTPTLYGTHHH
PAAPHAPTHFCAPPPVPQEFYSSTSPAAAPPPALHHPTLHHQLLMYKAASAQTHSSPESD
IRGAADRSESIEDGKSESSSWKGDSGGENGNSGDQRKGLAALREDGEESNGSEITLKF
>CpapHRS_62.51
MGSIPPELSLDFRPSSFVPKTISDFLKEVSRIGSVSEKVSKLDGFVKGLEEEMRKIDAFK

RELPLCMLLLKDAIIALKEESMRYAAKNALPVLEEFIPLKKRDRDEEDEEDRAAARKKED
CRDKKNWMSSVQLWNSEDLPSSTVNHRSFDPKQNHKLETKKKDGQNQCANVDLLFQTARN
RASARTIMPFKGFSGFPGRKEDKEELPVQGLSLSTPGIKSLRDELNSGSRTSCSQAVSSS
ATISQSNFRSGPQPPHQQTSRKQRRCWSPELHRRFVNALQQLGGSQVATPKQIRELMQVD
GLTNDEVKSHLQKYRLHTRRVPPTNATSTTNSSAMMLGGLWVSQDQYVDSSKPSNSQSGS
PQGPLQLTYTGGTSTTGGDSMEDDGDAKSESYSWKGHIHKPGKDDV
>CpapHRS_6.397
MDLSLDLSLVYVPKTMSEFMRDISKTKNISAKVSKIDDFVKRLEDERRKIDAFRRELPLC
MLLLTDVIARLEEEKVQYAELNDRSAVEEILPLKRISEEDGAANRERDGCDKKNWMSSVQ
LWSNNVNPDPEQHSISRLKSTSEDDGSFAEEHPVQPSNHSSRGGAIGPFKGKSGFQEKGV
PSLSLMTPISELSSCTTVSASIMSCRVGFGSSLCADQNPVELQSKPQHQQQQNSRKQRRC
WSPELHRRFVEALQKLGGSKVATPKQIRELMQVDGLTNDEVKSHLQKYRLHIRKVPASSS
AAATSHWSAGNHSGDSSKPSISPSGSPQGPLLSHGADSNGADSMETEEDEKSDGRSWRGR
IHRPEEIHVYN

>AcoHRS_Aqcoe3G398400
MDFIKKMKGIRCEDYMEALEEERHKIQVFHRELPLCLELVTQAIDICKKQVGIKHECEDV
ELISSTETRKLYSPKEKKVDWLKSAQLWNHDFSDQKDKPSQNTIGIEPNRLKGAFRPFQR
DEGHRASTATSTSTTAETVGVSPVNRAEEKQRQVESHKKARRSWSNELHKKFLNALQQLG
GSYVATPKQIRELMKVDGLTNDEVKSHLQKYRLHTRRTSAAIHIKDNLHSTQLILVEGLL
VQPEESFQDRPTTGDAAQLVTNGVYAPVAARPQSPLLTSQRQHKQPSTVHPESKIRGSQA
QHCMEDEGTHSDHQSSSSSTQSTTVSVSLV
>AcoHRS_Aqcoe3G045700
MVMMMDYMETIKITNNKNKAYQDYMQALLEERRKIQVFHRELPLCLDLVNQAIENCKQYV
YSSERSDQCEEKQEETSSEGPVLEEFIPMKRSLSCEEEEEDKSNIDESNVNNNNNIGKKV
DWLRSVQLWNQNPEVPPPSQQQEYHQQQEQENSPRKEISADVKENGGAFHPFHKTKPIKT
VASETVTINTVPNSVQTASTTATPKQIRELMKVDGLTNDEVKSHLQKYRLHTRRPGPSMH
GNGNTPPTQFVVVGGIWVPPQDYTPSTTNANAPSADSEKIYVPVAATSQLTLSLRQQQQQ
QKHHKQSELSTGSSHSERKSSLAEHSTREKNARSSSPAVTSSSTETTTASPNF
>AcoHRS_Aqcoe6G095900
MGELSLDLKPTTTITTSSSSTITTNYMPESIHVFLGEVSLIGNVSEKLSKADEFIQRLEE
EMNKIDAFKRELPLCMVLLNDAIDLLKREVMQWKKSGSSSASVMMENFVPSKTNSSSEID
LRDKKNWMSSAQLWTSNGYSSKTDDEKPKSIVDEMVKKDDGSAITTETNLPRNGGGGAFV
PFKGFTGFLTSTRMEEKEREIHQLEASLSLLTPGVRKPRTEPTSCLMNSKTTGGSRGLPT
SSAEKTQTNLHLPLQSQPPRKQRRCWSADLHRRFLNALQQLGGSQAATPKQIRELMKVDG
LTNDEVKSHLQKYRLHTRKFPANASSPGPENQKLALFGGLWAPEDQCGASKASTSQSESP
QGPLQLMTSGGSTTGDDSMEDDEEDGKSESYNWKGHA
>AcoHRS_Aqcoe7G203800
MVSITSEVCQDSKPHSYSMLLKSFVDHPEQTQKLEEFLARLEEECLKIEAFKRELPLCMQ
LLTNAIETYRQQLQSYRGNQGTNKPVLEEFIPLKHSSSEGSEKTSSISEKSSWMVSAQLW
SQSNDGNKQQTITSSPKETDQSFSPKLSLDDKKRNGGGGAFLPFTKERNTCASPNTLRVL
SDLALASTDDKELNDNKKCLESDKVKGGNGGTISEQGKTGAGNQTEAQAASTNSQTHRKA
RRCWSPDLHRRFVNALQMLGGSQVATPKQIREFMKVDGLTNDEVKSHLQKYRLHTRRPSP
SHQAAGNQAPQLVVLGSIWVPPEYATAAHGGGPTIYSAHPASLATTHYCAPQEFYPAPSS
HQLHHTIHHPHPHHQPHVYKTSSKTHSSPESDTQGAGDRSESIEDGKSESSSWKGDSGEN
GGERRGLASLRDEGEKSDGNEITLSF

>NnuHRS_012707
MDNFAEKMQRCQTYMEALEEERRKIQVFQRELPLCLELVTQAIETCRQQLATSTEYFPGP
TECSEQTSSEGPVLEEFIPLKRSSSSEEEEELNKPNNKDRNDKKADWLKSVQLWNQAPDP
PLKQDLGPKLSVVETKKSGGAFHPFERDTKRVTKPTATSVAAAPSTAQATATTSSTAETI
TDDSRRDGKEGQSHRKPRRCWSPELHRRFLNALQQLGGSHAATPKQIRELMKVDGLTNDE
VKSHLQKYRLHTRRPSPAIHSNGNQQTPQFVVVGGLWVPPPDYAAVTATKREANGVAANG
IYAPVAPQPVPESCSQQQKKQKQSPQSSLQSEGRSSGQRDDDAHSNSPSASSSTHTTAAA

SPSL
>NnuHRS_001193
MNPQPTMTIMANYTENAHKCQTYMEALEEERRKIQVFRRELPLCLELVTQAIETCRQQLA
TTTEYFPGQSECSEETSSEAPVLEEFIPLKRCFSSEEDEESNKPKNEDINEKPGKKADWL
KSVQLWNQDPDPSLNQDLSRKPSVIESKKGGGAFHPFQRDDKLTAPSITVIPSSAQTTAA
ISSTADTINGDNRGEDKSQSNRKPRRCWSPELHRRFLHALQQLGGSHAATPKQIRELMKV
DGLTNDEVKSHLQKYRLHTRRPSPAVQNNGSPQPPQFVVVGGIWVPPHDYAAATETKGEL
NGVSGVTAHPTIYAPITAPSVPTVRSEQRQKLSPRSPLQAEGRSSCKKDDDTHSNSPSTS
SSSRSY
>NnuHRS_006595
MGSTSPELSLDFKPWQAPKTICCFLRELSAIEDIPERLSKLDEYIKKLEEEMKKIDAFKR
ELPLCMLLLDDAIITLKEHAMQCTTSDVRPVMEEFIPLKRNSDEDGAPKKENDCRDKKNW
MSTAQLWSSNGYSNKDNCYDRNKKSTVDRKEVGEDDHSVPENAFQPCKYRSRGGAFMPFK
GYSCFPATVVKEDKDVLPISALSLLTPGIRTHSIEASPGYLNSKSIISSRVINSSSTKHH
TQSNLQNTLQQQQSSRKQRRCWSPELHRRFVNALQQLGGSQVATPKQIRELMKVDGLTND
EVKSHLQKYRLHTRRLPAVSPPANQPVVVLGGLWMPQDQYGPSPKPSTSQSGSPQGPLQL
AGAAGGLSTTGGDSMEDDDDGRSESYSWKGHHQKSGEDDV
>NnuHRS_011780
MGSTSPELSLDFKPTYVPQTISGFLRELSTIGDISQRSSKLDDYVKRLEEEMKKIEAFKR
ELPLCMLLLNDAIVSLKEEGMQCTTSNVRPVMEEFIPLKRNSDEDVRAKKEKESRDKKNW
MSSAQLWSSSGYSNQDTSYDRNQKSVVDGKEVTEPEDHSVPENPFQSCKYRSRGGAFMPF
KGYTGFPTTPLKEDKEVLPVPALSLFTPGIKTPRHEPNPSCLNSKNSLATATTNFEEAKE
MLVPRVTSSFLATPKQIRELMKVDGLTNDEVKSHLQKYRLHTRRLPAASSPANQSVVVLG
GLWMPQDQFGASSKPSSASQSGSPQGPLQLAGTAGGISTTGGDSMEDDEDRRSESYSWKG
HLHKSGEDDV
>NnuHRS_006061
MGSPAELSLDCKPHSYSLLLKSFGEQTDQTQKLEEFLYRLEEERLKIDAFKRELPLCMQL
LTNAMEASRQQLQTYRTNQVSRPVLEEFIPLKHSASDQGLEKASTASEMASWMTTAQLWS
QSSDGSKQQPATAASAKEVGDQCFSVSPKLALDSKQRNGGAFLPFSKERNTCASPTLRGL
PELALASTDKEMEDKKCSESDNAVSVSCSRRENSSKSNAGGTATTEQGKGGRNSAEPQPA
TTTQTHRKARRCWSPDLHRRFVNALQMLGGSQVATPKQIRELMKVDGLTNDEVKSHLQKY
RLHTRRPSPSPQAAGTPAPQLVVLSGIWVPPEYATAAAAAAHPAAPTIYSAHPASHASTH
YCAPAVAQEFYPTPTHQLHHSLHHVYKASSQTHSSPESDGRPAGDGSESIEDGKSESSSW
NKGDSGDQNGGERKGLASLRDDGDESNGSEITLKF
>NnuHRS_003614
MGSPAELSLDYKPNSYSLLLKTFGDQTQNLEEILSRLEEERIKIDGLKRELPLCMQLLTN
AMEALRQQLQTYRTNQGPRPVLEEFIPLKHSGSEGSEKAASISEKVSWMTTAQLWSQSSD
GINKEQATVASAKEGDHCFSVSPKLALDNKQRNGGAFLPFSKEKSSCATPTLSVLPELAL
ASTNKEMEDKKRSESDNGVSVSCSTRDNSCSKAAGGGGTPTTEKGNAGGNAAEAQPTATT
QTHRKARRCWSPDLHRRFVNALQILGGSHVATPKQIRELMKVDGLTNDEVKSHLQKYRLH
TRRPSPSPQVAGTPAPQLVVLGGIWVPPEYATAAAAAAAHPAAASTIYGAHPASHPSTHY
CAPAVAQEFYPTPPHHQLHHSLHPQQHIYKASSQTHSSPESDGRPAGDGSESIEDGKSEN
SSWKGDSGENGGDRKGLGSMRDDGDDQSNGSEITLKF

>CatHRS1_Chlat1_8216_Chrsp76S07637
MLGSWAERLFYSASVRAAQGAGAGAGAGDGGGGGGGSAGGHDHAGGGPVSRTPPPPLTPAELEQYLTALTAER
SKVEVFKRELPCCTQLLDGAIAYIQTVVASQPQQAEAPSTQEADADQPSSSTPSVETTSPPVAPLPPPPLFSP
PWPHLPWPGAQPWAPNAQMWGASKDGMAIAKGQQVPFSAPVAIPAVAMKTEQTSPVHVSQSYSPPRMPSPVHA
SPPVVVPQPHRPAVVLASHGGAFTPFLPAREPDPNRSPPRSQSQEHASTSRPSAEACRTSHSKETRSDSDGDK
GAPRKARRCWNTELHRRFVDALEQLGGCHTATPKQIRELMLVPGLTNDEVKSHLQKYRLHLRRPQSSPQPRGP
PEPWQMPSGAMPPMMHPGMLPGMIPAMTAHMQDPQSYQAAVEAWHDPSGVTTLSLGRGQAPPQPTSSAPLMQE
TNPEYSGDASAQAAAPASAPGAPWPTPSGSHPMDGSTSGSSMSDAAGSG

>MviHRS1_Mesvir1_13691_Mv02124-RA.1

MVVSMSPRFTPWTGGANGINPAIQDAGCAAGAPSVEDLHVYWREILAERQKVEAFRRELPHTIAILDREIFRL
RLPLETAEASGHCGDEAVTGEVPAYRPAYLPAKDAATAGPKTGSQADPQPHYASAEAPTKKLMEAGVGVLEGL
MEKSSGAVTSGSSGGPNRYSSEEVKRQLRMSTEAMLAKKAPRDAATPSCSVSNGDGGKQSLERRETDAASGNP
RSSSRHKGSAQEGDAQGGVSPGSQRPPIKKRGDGGDVDARSREDSDGGDNTSGGGARRDTQRGRKERGDEEER
AAHENSSGGDLHARSRLSGRGLRKTGSASLGDGVANTSINNAPNRSPPGEGMVAPEPMYPPGAIPGVPLPPPA
VPPGSMADASAHAAYFQAAAAAAAAAAAAAAAGRPMPPLPGAPPPMPPMDMGHPHAWPSPYGPRPGDPMDQW
GAGFSKDQFAKMPYHLQQQMYQQSEVAARAAAAAAAAASMPIPGVGPPPPVPPPPGMPEMGPHGMMVRHGSGW
SDVSAGGAGDMPRLSRSKTWSPEKTLGIGGPGTSAATISRLPSDGAQRTIPLRLAGPLSPTGSGGESGDRGLK
RKASGGLSRENSGVTGDMFMNAADRMMHAAAAAAAAGCGPPPPGTMERMPSADLNMLMGRPGGMMPPLFSNPP
PPPYPGVPGGAPNPGRMMPPYSVVPPAPPPGMPPYGVPPPPVGLDPRHYNDAWQRAAAAAAGAPSGSPLGMDM
ASRMRGVHPDQQAHMDQWGMAMRGMAAPSANGGPPLGAPPPMGIPDGYSAMGMRGPLPPHRPQPQPHRDAAGV
AGVGGAPPPGGLRKEGEYGDAGGGGGIPGGGSGVGGGRSARKMRRCWGTELHNQFLAALEQLGGCQVATPKMI
KELMDVPGLTNDEVKSHLQKHRLNRKDKQAGDAGAAGEDAKEKSKGNDNDNDDDNDTDAGSDSKSEDEGDA

>MenHRS1_Mesen1_2781_ME000170S01887
MQKLEEYLKALQDEQRKLDAFKRELPCCMQLLTEAIERARQQLEGTQAKAAAAPFGARAPSLQLGRASPEPPA
TSAAHDVARPGDHSRGGIAEPTPGPASPLMRAQEFLPMPPSSAAWDRAQWRPELASAKEGAADQAKISEPTAE
RPSARPAWMVPSPAWIPPLDGQAAAANEHLKEERPEPYGPPARPGALPSVEHAGLYQSPGTKAVLVGSQRPTG
AFLPFTRDRDRQVAGTGIGASGGGGSSTTTRDNFSSPPSGSADHILHLPRTDLALRLSPSGGLHLGVGSGADS
GTGTGMAAGAGMGGGAGAASRGMSLEAGQGAGAGGAEPGGGGAGAYRGRMQVPSSPEERGRGGQLGGGGVRSR
SQDSDLHSGGGGDPRDVGAYQTGGGGGAGGHSGGSSGARKARRCWSPELHKRFVSALGRLGGPQVATPKQIRE
LMKVDGLTNDEVKSHLQKYRLHTRRPSPVPHATAAAAAAAAQPQPQLVVLGGMWVPPTYAAVTSGAQHQSPSG
YDLAAPTSGPHSLFGGGPQTAYPLQLPTCHKYGQVAAPHPGSRERAPGSEIDLERSPQQTRTAVHSPPARGLY
LTPSPRYSSRERDEARIRARPEGGAARDGGDGRAAERRPSPLEAALDYKAPNLDMTLGSRGEYTPPRLKQDEP
CEARGAPEGRGGHELAVRRASNGQQPAVRP

>CbrHRS1_Chabra1_322305_rna-CBR_g3557
MPSDARHSGGLSGSALDEVTSRAAGAAVTNQRERGAGVEAGKVCHEETGSRLEERLCALEAERRRIEGFMEML
PHCRDLLDKEIESMREKIEVKRLLKVHGKSSLLENHMDSSRRQPIEPEESVVPSNGGAVWDDSSTSGDAGIGV
NPVHLSPAQGERDCPELFKKRRTEQPMHDETQAETERCKARFPLSNCMEKPLGMLTLACDSQLVDGNDKFDNL
QASEKHDRRHGCRLEGKAAAEDGDNAKPSWLWRGVPREQHLVEQNASKEHNEKGHAEGYHHGVRQSWLHEPVE
QKDELGSGRCQHLQPHLALTSIKGGNENHSMQTAGNDISFLQPGRERPGGAFLPFVPNRDNKASTGVGTEKGA
STIQSASLPVPACIELVLGLAPGTVPEPRHRGMQMETFKGTDHGEHADMEMMQVDRTHRLLTAESAPNEDRLE
TPMTVAGNIQKKEEDMVVRDPSREGSKGRTVNGEGGGDANGNEGAESGGSESSDNSGGRESDATNTALQASPD
AMRTAPGTMLTTSFNGSSGGGHRKARRCWSPELHRRFVDALTILGGPQVATPKQIRELMKVEGLTNDEVKSHL
QKYRLHTRRPSPPQNISLPHPPQVFVLGSFYLPDYGSSPTLAQQPPIVATTPAGVYDTAAVAAAAAAAAAAAG
AGTAAGGFGTPVASLAAAGVSQQRSHFGLHASSLQTQAGLSTQPSPMLHQHQFFPCKPAHGLTATAIGQVSLC
RPSQRESPQHPSSQPEIEVESTPLSMGVGLGATRQAREQEPTAEQHLRPSCEDRTVELSGAERKYVRDVNGEH
LGAGAHAAPADGVDVRRVGSLEGGSRKRSLETANARGREGEASPMLLKLQLNEDHMNSAGYSNDCHQLGPRAA
RAMSGIGGHRSLYCTQPVLSQSG

>KniHRS1_kfl00369_0070
MSDDADRQELQEYLRGLEEERRKVEAFKRELPLCMQLLTDAIQSTQAQLKADDEPMDSDD
VSEPERPRDRSHHSHRPIPTIPPHPAHLPFPPPFRGSPAWLSEAISVRAAQQTGVEQRAL
EQRMEGLGPPRGERAPGRPLRDGGEERAGPQLFRTPGAFLPFSREQHSPHVARPSSPRAI
TYPLAVHLTPGGLAAGSGQDHDGSPASVLEERRSQTAPAGPSDTEGGHSGEGTAMATDGT
GRNVKITSLFPLFIYLPPGSFAWQHGGDVATGAMSRKQRRCWSPELHRRFVNALNQLGGS
QVATPKQIRELMKVDGLTNDEVKSHLQKYRLHTRRPSPIPTSGASPGPGQVVVLGSIWVP
PEYNAAASSAAALYDAATAAASAPSQAGHAPSLDEARGIPPGAFHSLDEARGIPPGAFPF
FQVASGLLAGRHRPSASSTEMDLGVSGSGEEGGQNQGGFLKAEVKEGSAFGAFRAFKPLG
GNQAERNVEDTEPVHRERRAAAAPDDASESMREEVPKVKPPGRDLDEDESSEGEVVHRGF
RNFLPPRPEASAQKPG

>MpoHRS1_Mapoly0119s0041
MGSPADLTLGCRPQSSGNEFKAVSTSGDQIERVRKLDEYLKALEDERHKIEAFKRELPHC
MQLLDYAIEVSKDRLVDSPRSSSGGGHAVDSYGEVETSTLGSFGKHVVEEFLKKSWEAPK
REIEEVEKVSKRSDEGRDADHRPSWMAEAQLWSQPSTRSDAWKEKESSTEDSTRRQRDTT
EQSTVTSPANLFNTHKRTGGAFLPFIKEKPVATVASRVKVLSGADLALSSVESRPASRIA
LGPVESEAGSLDVSTTRTRDVGMEVVQAKDNGRKLNGSGSGGNHSGGTGSTSSGGGGGGS
GQAQRKARRCWSPELHRRFVSALQQLGGSQIATPKQIRELMKVDGLTNDEVKSHLQKYRL
HTRRPSPAPQAPQQQAPQLVVLGGIWVPPEYTSGGMYHDPSSSVSQQGHFCQASLSQEIY
SQLNPSAQIQLHRSQSHSSPQGPLQSTSQLSSGARQTSAEMYREDSPGEDGKSESSSWKG
EDDSRGRSEKESDREVGSGGRDDDEDDTDVEDEGRESGMRLKLEMAFTRSHTVMKPEPQM
A
>PpaHRS1_Pp3c15_4480V3.1
MGSPVDLTLGCSTQAPSSNGYYNHQNEPTHSENYLESLREVEERVRSLEEERRKVEGFKR
ELPLCIQLLEDAIKTWKVKLVNGKHLPLQLRSGSMDSGRNQHVMPSSPGSRPTLEDFMPF
KRRWEQRDQPSESSDVEEEVRGSKRPAWLTEAPLWRQSSGCSEREGKEVQEHSEFGISER
EQSSITSSQMLLSAKQRPAGAFLPFIRDQPGASSFLSRPTPRTVASAADLSLSLGERTTT
PSSRNLRPSDSEVGSIDAGLQTPEVQIPKDIGVMSNGTSGAGSSSGGGNPTRKSRRCWSP
ELHRRFVSALHQLGGSQIATPKQIRELMKVDGLTNDEVKSHLQKYRLHTRRPSSSPQSGG
MGQSPQLVVLGGIWVPPQYAASGSQPSPGFYNPSAAHSPQPNQFCPTSLPQGYSCISNST
QGAQLEMHREPIFEPPRQAGTSQSQSSPQGPLQVTSQLSGATQGPSSAAYHDESAGEEDA
KSESASWKCDNPKGEELKKSNSNQGSGDGQSRAHVSEEDVNQAD
>PpaHRS3_Pp3c20_11430V3.1
MGSPANLALGPQVSTITSTIETALNCTVSADGLEADEIQKIAAHLRALQEERGKIEAFKR
ELPLCMQLLDEAIETAINKVMVDQCSPTASLQTSVLDAQDSDYERPKRTLVLENFMPLKR
RWEKQLKEKIFQNDAGNNSQQHEDKVVIGRPAWMKETQLWTKQSESIDYEKASPSRGSRR
QVEFKCSEQDRPLTSSRLLLNSKDRQGGAFVPFNREKQLPSPPFSSQSTGPASDAAGFST
SSAVRTPSRMGHTIESFDIGRVNGNVHVRDHSSDARNSIDHSVQTRTGANQPQRKARRCW
SPELHRRFVSALQQLGGSQVATPKQIRELMKVDGLTNDEVKSHLQKYRLHTRRPSPSPPS
PSHAPQLVVVGWVRPEFTAVEGAASQASTGDNPLPGISNECVPSQSTGQPQKQSPPNFQK
HLPAVSTSQTSLRGPSPVGVDSPHSSSDSYIDESDDEQDGNCTSSGENGRQYECNEKPLQ
SARESKGSTTCLADIDATINPQMKLRILRKTKAAA
>PpaHRS2_Pp3c9_8580V3.1
MGSPIDLTLGCNTQAPSSNGRYNDHRNSSACLEDCLESLRKAEDYVRTLEEERRKVEGFK
RELPLCIQLLEDAIRTSKEKLLRGKDLPQQLKANSMDSGKDQHGVSSGPGCRHTLEEFMP
LKRRWERDQPSVSSDGERGEDRSSKRPAWMTEALLSKRTPASPEKETRLTQEHSELSVSE
QEHSWVTSSQMLLGANQRPTGAFLPFIREQHGVNSFLSRPTPRPAGSGADLSLSLCEPTT
TQTGHTLRPSGSEVGSIDAGRQTVDVPVAEDMGVISNGTSGAGSSSVGGNPSRKARRCWS
PELHRLFVNALHQLGGSQIATPKQIRELMKVDGLTNDEVKSHLQKYRLHTRRPSSSPQPA
GAGQSPKLVVLGGIWVPPQYAASGSQPSSGVYDPSIAHPPQPTEFCPTSLPQDYFACINN
DTSSVSLQMHRQPIFEQPIFEQPKQAGTSQSQNSPQGPLHVTSQLSSATQGPSSEVYHDE
SPGEEEAKSEGTSWQREDLKSEEPTKPNVRFSLGAGSGDVQSHAHASEEEENQIE
>SfaHRS3_Sphfalx0001s0189.1
MASPPDLSLGCQTQAPSSASTITSSEVSNPLVNNQLGRIQALEDEQRKIEAFKRELPLCM
QLLEEAIAKSKEQLEEMQPAPGTPQQPLQLRCSTPEKSQSNVSSPNDNNKLVLKEFMPLK
KRRLERLQGGDQLLGGDEEGDRQVERTRGIDIGRPAWMAEAQLWTQHAAIAEARNEKDLS
PEDRCGQGECTRAAQANGSIETSRLLLSSKQRPTGAFLPFTRKKQLTPPAVVSQSNPVTS
NGANLSLSSVQQLPGLEGQAFLVAESEVGGIDAGLSTQEVNAEGCMKRPLLKVVSGAGST
CSEGGSQSQRKARRCWSPELHQRFVRALQQLGGSQVATPKQIRELMKVDGLTNDEVKSHL
QKYRLHTQRPSPPPQSASAHAPQLVVLGTIWGPDPDYVAAAAAAAQTAAAGVYDPSVSHH
SHQYSSYCKSTLPQDHFANFNNNRRSENKPQMQEPDCPQSQQAVSQSQTSPQAPLQVTNP
VSSGVGYQYDSGGEEGKSDSTSWKGRPLETGGGDREEESQIQIKPQAAAADQL
>SfaHRS2_Sphfalx0004s0229.1
MGSPADLTLGTSQAPSTNATTNTAAATTTTTVLSKSSPSSFVGDRLQRMRKLEEYLQSLE
DERRKIEAFKRELPLCMQLLDETIESSKEQLSAIQALRPTNLRGPLQLLQVVTQSSSQQL

VQDERDTGTTVSSPAASGGHQRGLEFIPLKRLSETQQLRDDEGDDDDDDDDDDRELGDQS
DSCLATRGIDIGRKAWMAEAQIWTQQSEPMEVHHEKEISSPEDVVGRQEHFGQSPEQEQS
SVTSSRLFFSSTQRPAGAFLPFIRGLQQVTAVAPLLSHSTNGTASAAAADRSLSSLDQMS
SPTNGHGFGGGAESEVGSIDVVSLKTADQNQTLDAQAYKEMQGPESNGASGAGSTSVGIV
SHSHRKSRRCWSPELHRRFVCALQQLGGSQVATPKQIRELMKVDGLTNDEVKSHLQKYRL
HTRRPSPSPQSVSQQPPPQLVVLGGIWVPPEYAAVAAAAAQPTSGVYNPSQQSVYCQPSH
PQDYFMHNNSTGSSGGQNVHHHPPTSFEPQKSTSVSQSSPQGPLQLTSQLSGGAQEPSLD
DTGGEDVKSDSTSWKGHQQEDEGKETANGANKQSSVSKEDNKAGKSRDKTPGGEIFADHN
SPVLELQISCGTSDIDMGVKDVA
>SfaHRS1_Sphfalx0010s0233.1
MESPVDLTLGSAHTASSSYTNATNTVNGLESSLSFVGDHLERIRKIEEYLQALEDERRKI
EAFKRELPLCMQLLEETIESSKEQLAENGVLPTTQLPVGAAAVSSPTRLRGLEFMPLKAQ
AWERQQQKDEGDDHDQEGDGSECPGVVDIEKMTWMTEAQLWTQQLEPSSELHCENDRMSM
EEDMGRHPNFDQSEQEQCSVTSSRFELFSSKQPPAGAFLPFVGEQQVVTAAPPPSGSTTG
TTGSGADLSLSSLDHVSSPNGNGFGGAESEVGSIDVTRLRTPDRILDAQGCNLGIRPQSN
GVSEAGSTRVFGSQSQRKSRRCWTAELHRRFVGALQQLGGSQVATPKQIRELMKVDGLTN
DEVKSHLQKYRLHTRRRSTSSQSVSLEAPQMVGMWVPLECAVAQAAAQPTSGVYDPSPPS
YCQPSHPQDYFTCNMSTGSSGGQLKQHPTTFEPQKSSLLSQSSPQGPLQLSSQLSGGAQE
LSNEDSGGGDVKSDSTSWKSCQLDVIEGKESGRSIQHTVKQDNNGQQIKR
>SfaHRS4_Sphfalx0040s0035.1
MGISSDLSLGCHTQALSCASTITSLKVSLPELEENQLGGHLHALEDERQKIEAFKRELPL
CMQLLEEAIAKLKEQLEEVQPAPGTPQPLQLRCSTPEKSSNGNNMLVLKEFMPLKKRQCQ
SLRKEEVGNDLQGDQPVEQPRGIDIGRPALMAEAQLWTQQVATSAAHIERGALPEEQCGP
LESGLAEQANSSIRSSRGLLSSKQRSTGAFLPFTRKQVNVPMLSQSDHVTSNGGNLSLSY
ANPLPGFQAQAFPVAEPELGSIDAGPSPQKLDAPGCMKSGPFSNGVLGAGSTCSAGGSQS
HRKARRCWSPDLHQRFVHVLQQLGGSQVATPKQIRELMNVDGLTNDEVKSHLQKYRLHTR
RPSPPPQTASAPTPQLVVLGSIWGSDPDYMAGSTAQTAVEVHDPSPRHPRQQYASYCKPS
VQQDQFAHFNNGRSENKQQLQESVCVKSQQAVSQSQTSPQGCLQITNQLSGSAGYQDDSG
GEDAKPDIAPVGKDRG
>SmoHRS1_402040
MNPEEEFGQAEGPDLTLGCAIAKRDSSDQQTRRRMEDYLQALQQERQKIEAFKRELPCCM
NLLDQAIQSTRGCLDESYKTEMEEENPPKLELFLENRSGFRLGKPKSEAEDHRDFEERPS
TPKQNWMQLACFPSQSPREDEEFETRKALRSSTASADSGAFAPFVRDKKWDQAELCLSSS
ICLSLEQKKTGFIKFQGNESNLPLVELSLQQMPSSGICSNSTLHQQRKARRCWSPELHRR
FVNALQQLGGSQVATPKQIRELMKVDGLTNDEVKSHLQKYRLHTRRPATVAPGTAAQKSP
QVVVLGSIWVPPEYAVFDHHQAAPSMQSTPLQQSKAQDFFANLHASPAEPMVFYEKQELL
KSSKEEGEVLSPRKASATTASEEEDDGDVTEAEEQR
>SmoHRS2_66160
VTALEGERRKIEAFKRELPLCMDIITEGNCSYRDEAIGHDSVTAKSDSVLEEFIPPKTLE
ESKRDEKACDPDLDKPSWMAQVQLWSEAKQIKTQVEIYSKHAVPGIFANSGGAFSPFVRA
APSTEEKPGLRSTSDAGLTLSSSESSPRSGVVEETRPTVEKRTVEATTPAARSTTSSQFE
SNSQRKARRCWSPELHRRFLNALQQLGGSHVATPKQIREIMKVDGLTNDEVKSHLQKYRL
HTRRPTVTTSQTSEPGGAQVVVVSGIW
>AzfiHRS1_Azfi_s0208.g057975
MRKIEAFRRVLPLSMQLLSSAIEKSKEELGDGNGYGRLTTEKGSRPGVGDGDHLQRSMEVISLSEEQSCNIDA
LPNDLCDLTLQYSRDFEPFSKDKHREVSSLVFPKARRLGAFLPFLPRISFNIESGKSPSSSMPTLNEDYMIRG
VEDHLDHLKPSEDNIHDVPIVLSLDYTSGSNGRYSSRRKRRRSWSPELHRRFINALQLLGGPKVATPKLIRQL
MNVDGLTNDEVKSHLQDTSFKLAHPTNGHSFIEPFLRG
>AzfiHRS3_Azfi_s0092.g042991
MKEQDLSLSNPLLTTPYEYNMPKLHSDFNPLNQCMERDDRLTYATHSILQGTISNGIIPSTKFWSQKEAYAHG
EEIVSTMSGRKEDDKEINNSITCSAKLSNGIISCNPERRRGAFVPFTMTRLRATNISPPDHDHGDLSSSTNIK
IGRDIKYVRREEDTLQIDGRTNTISPNNVINHDNDPSLTTNININIKIKIKSIQCNNVNQLNKKQRRRWSPELHRRF
VNALHQLVATPKQIRALLKVDNLTNDEVKSHLQKYRLHNATRQVKMGSSETTMLPHDVVVMGGICVPNHECLN

IRQQRNSLNFSSSSHLCLHYIDEDKKFRVRDSSHDLLCSEEENSEAEDGRVTNHFLNRVRYMDHQRSENSVDI
ALWLVLRWNVHHSVATQLVSVAHLLDECGKNVGSIFKLYNDLVDINDGVVAYK
>AzfiHRS4_Azfi_s0114.g046051
MGSPTTLVLECRRPSAAYTATTSTANSGCRSSNFTYDHHNIDDDGQGKQIHRIEEYLNALEEERKKIEVFKRE
LPLCMQLLQDAIQAYKERLCFYKPARPITNGKEPNHHDQSVIKEAIVVEECTPLKIIKPADGSYTDQTAEEKN
ICKPTISWMKVDCIDMQREVNSQEMCNSDLTIRKNYGLQCVQEQAFSTSSKPFFHSKNKSGGAFMPFNRDKLQ
AVVQPRARPGALNPSDLAFTPSDAKVYCNSPNNLSLSESPPPPSSSLLKLMKIKPKVENNHNELIEFKQTSMK
GTDGSNGTSIPRKARRCWSPELHHRFVNALHQLGGPQVATPKQIRELMKVEGLTNDEVKSHLQKYRLHTRRPS
PSPAVSAHTPQVVLVGGIWVPPEYALHNSTHNSIQNGIDQSPALYSASRPHAYQTPATSTSQEMCSQLESPEQ
INVQYCSSAPAHSQSSLRSATVRNNGQSSLVVYAHSDDVMGREESVGDDDDDDDENSSYKGLYMNEATRVRQN
VAHVDMDNEDRSLTESINANRSHCTSDDDSDTEIEDRRDCDETSLELS
>AzfiHRS5_Azfi_s0092.g042967
MSLYKDHHQVMMGSSHAELTLGLMPYMNNEGEGGARGSNTSKLLELEQYVESLEAERKKIEAFKRELPFCMTL
LTSAIEASKEKLANNFQNSPSNSKDQEVISNHTVTPINEDINIDTIFRPLNRSGTPKGLSSEEASPRINGTTT
QTTKFWSLEEAYAHGEEMVSAMGGGDITYCNSSSIMKPIPISCERRGGAFMPFLKTKTASTISAVEQQSPCEV
SSPAIGDKEDYTERTSSPNTSQEHYPAIMVTNTNSKTTPPTRANNATQHSRKQRRCWSPELHRRFVHALHQLG
GSQCATPKQIRELMNVDDLTNDEVKSHLQKYRLHTRRQVDGSFGKTTMSHDVPQLVVLGGIWVPQGYGNEGSK
QQKLASNLPSDSHMYIHSGEIKFRGDSSHELYSEEDNSEAEDGRVY
>AzfiHRS6_Azfi_s0015.g013719
MGSLAELTLGRPSSSSIITSYKHGVNDHHVESDIEEHNSYVNKIDRIHTYLKLLEDERRKIDAFKRELPICMQ
LLNSAIENSKGELLDYNHGPDHQTAGATRPDSEESGLVDHEEIFVPLTMKKRSTVQNQANPSGDQTNVLNQNE
FMSKSNQLKETRFWSPSEAYAHGQEMLASIMTKKPQQMVKRRATTAAAADNPPSFLTHEYSDHNNADNASGGP
DQFDQASLGEIDEPTPTTSSSLQHHDEIYTKEGNGSHHTNAPMPASRGGAGGVATPKLIRELMKVDGLTNDEV
KSHLQKYRLHTRRPNGCSGERTKAQTPQLVVLGGIWVTPPSQSSSEDPAFAATTTHSLDRQAGESDQEAMDEG
CSWIRGGPYISQRRKMGLMHLGLHHGEFSYVQNQ
>AzfiHRS7_Azfi_s0173.g055847
MHIMGLNMATDLSLGHHEVMNNEDIKNNTSDNKFTITSPLTFKDNYASNIILYDNHNEPDTTSIVRDELMSIN
AMYSDLRALEVYHKALHEERRKVRAFEHELPFCMQLLDDAIEASREHLEDRLTCKLHTPSLDDDKTPKFVHNG
GSMLPSSGEFYTKPNHLKRKWTSYVDVNSEYKQAARLTSSMSHPIAKWMPTTSPNDRIGYENPSSSNIQAQGN
IIKGSIGKFEELCVIKPEPLSAHPLIHQNYMKVVINGAPISNMEANNVSPSPVTCNMNGIAKLSNELHQVSSL
NVSKPTLAMSSAPAMQRKPRRCWSPELHQLFLKSLNHLGGCQVATPKQIRELMNVEGLTNDEVKSHLQKFRLH
MRRPNMNMGEMESSKQGGAGNGQHVVVFGCNPLMRGLSQADSFDAYHDRNDEDPRQIHRNLPLICDNSEVTSM
EDDETVENQQQEERRNMHSRHGVDHHRPPRLDINQENFDLRIACTAKYCQ
>AzfiHRS2_Azfi_s0208.g057980
MGSLSEVESQSMMMKRKKKIQEMERYIVCLEEEMRKIEVFKRELPLSMQLIFSAIQTSKKELCECNENERLIR
VKGRIPLDSTSPDQAKAPELGQRKTFTFWTPDQAYDHAEKILHSMTNPHKKHTLEAVCSSGERASSIDYSPTN
DPCDLTPPYLRDVLPFSRDKHRTEISLVFPKDRRQGAFSPFLPKDGMRTSSNVESGENPSSLTLNEDYISHQE
SLSHSVDDRLDHSMSSRDDEEPNMKLDEGNIQNASIVLSLNSGRNRRHSSKRKQRRCWSPELHRKFIKALQLL
GGPRVATPKLIRELMNVDGLTNDEVKSHLQKHRLNAQRPNRSSKLNETSPSLVLLGGVWVTHDPNGIQHDERE
HIYNNVLNPPLSEGSSEEISCAPMHTFIR
>AzfiHRS8_Azfi_s0008.g011579
MAMQGDLGLTLGFVREDQCKDLNDDSTNPTRPNALMCSNNHHEFGSSSSSFEHIRRSTPSPDANYVTINSGTN
TNASSTRPHGNHNQFLPTKLLNHNTHTNIGHMLSNDHHHDHNNNTNNNHIGISSDYHGTMMSNNTNYHDGAIV
NDDNLEWCLRALHTERLKIEAFRRELPMCMRLLDGAISATTAKLNATSNNAPSSINYMQLYDADNDANSTHNE
KTDYNHNNNNCSNISMNHDEIEDHHRHRHHHGHYVDGDGGGDGGGGGGDGDGSIMILKPRHPGINMSSSFLSP
LNMMKPSGSPLALPISSSSCQQHTSTTMTTVSAGGPTEGGVVGVGVGVNRKPRRCWSPELHQLFLHALHRLGG
PQCMHLSPLSLHFCPSLFNSPNGPPCGTTTTTTTTTTTTTTTTTATTTASGGGGGGGSSIATPKQIRELMNVQ
GLTNDEVKSHLQKFRLHTRRPIVSSSCSEAYQQPHHLVLFGGIWVPTSPVEQSSSPHQQSTSMGAHTCDTSLP
PQAE

>ScuHRS1_Sacu_v1.1_s0185.g025023
MGSPTSLVLECRHSSSTTTAANSGSCRNFSNTADDQACEIQRLEEYLNALEEERRKIEAFKRELPFCMELLHD
TIQAYKEKLAFFKPSSQTWSHIQQKSNSDDADSKPESRIIIEEFLPVNRAPTCTGRNAEEVQQRPNWMKEAGL
VDTHTGSNERTVKVLKSTSRFIREFSSIKDLISAFLCTFFFYLSQVCEPEPATKMNGSYSGQGQAFSNAWKPV

FHSKTSSGGAFTPFNRDKPTVIQPTARPGALNLADLAFTPKDSDVSPLSLDQNPGLLENMKQKENVIRQAAKE
AEMKVSVDKSNGSNNVTPARKARRCWSPELHRRFVNALNQLGGPQVATPKQIRELMKVEGLTNDEVKSHLQKY
RLHTRRPNASSAAMSTHPPQVVVLGSIWVPPPDQSPALYGSLQPRSRIYRSTTTPSEEMCSQLESPEQINVHF
SSAPAYSQSSLQGHVEHNGQLPPLIYSHSDDAGREESVGDDRISANSSSKAFKDLSSHHSLPAHHQDETVDKS
PDGSMYAQGSHCTSENGDDTEIEDERDCATSV
>ScuHRS2_Sacu_v1.1_s0042.g012676
MGSPTSLILGRNPSSCSIRTRNERGCLVNLHLGDGDYLVDEYVNALEEERQKEEVFKRELPLCLQLLHDAIQA
FKEKLAFRGIEDAIDDVNCTGSNVHEGSRKEEETVLVDEGFLQPWKWVQQQQQHGVFSERTAGKTKKNKKRSW
DEACGEKEGNYYRSEEEDVHCRRRQWNSRSVFPSNCCRIISSGGAFVPFNKHRQLPAAVSQPDVQKSSGVPLQ
QQEKNINTEGSNCNNVIAYARKARRSWSAQLHNRFVHVLDMLGGPHVATPKLIRELMKVEGLTNDEVKNHLQK
YRAHARIGCRSPTLTPRAPAATAAHNRALRAASTARYGLHHDLAHSITRYVPSSLLGSSPHHAANHN
>ScuHRS3_Sacu_v1.1_s0071.g016842
MAMDMSLMAAELSLGRSNSCEMAAAMAAEQSTIATTKKKKKAQSGATEEKRLMNGDGMEGLMGIELMNHNHNR
NQNQPIHQNSNYTHQFNSTHRHLHVSARHLPPDLHALEEYHKALEEERLKIRVFERELPFCMRLLDDAIEASR
EKLDDRLKCKLLTTAAEEDEEEDEEGEADRDGTAALARFRGQDPLTVISMSSAPQAKISVGAGTEPAEPNCSS
TPPRKRKLLLHHYHMTEPGHDVDESPGSDARGGAFGAVDLYHDVAANAASDHDHHYVKSPVNGAKHDHNHMIK
GPISVANHEPAGASASASHNSSYKWPLTPSPHDTVRDRRTSYATDHLLIQKGASTAGHGQGGRMSHNTTSTTT
AVMGDVSGATAATDITLPSIRTDEGDRNGVNLFNHMSRSAGVASAHAQLAAETISANDRLQAHHKDYSYNSLI
ATSPDLMTAMPDHHPAADAEESAAADPLLKKQIRINPLINNRFRPPASSSISMVNSEYSSSINANCSSNCCKP
IHNIQQDGSINANCKNMHQQRAMKPPERLVHGNHSGRISSDEAAVAAVDYLSATTTEINLRALTEGHLVNTRS
TPAGACQAAAVASSSCSATLSNSSGMGMMQAVQRKPRRCWSPELHQRFLNALNHLGGCQATPKQIRELMNVEG
LTNDEVKSHLQKFRLHMRRPHNMSLGSIQTDNPIVEKGNTNGHHLVVFGHIWVPPEQYQQPTNVHTLSHHSHP
INDTTRASPLLEHRCELDSSYNVVRQGNEIMNKSQLQMGTSNDVDGNAIKACIDNNKVTNVKLQLQFSDMCNN
MDCQELEEKALKSEGDRFVNMDSDGHGEDYKYDGEHKTRSNTDEEIKLRITCSNYGIRDQLINNAIDRV
>ScuHRS4_Sacu_v1.1_s0013.g005748
MGSSAGLSVMRTCHLYPFEGDLERMLKLEQYIKCLEDERKKIEAFKRELPLCMQLLNEGLLNSALQQETDIRS
RSTQPLLDMFLPSKKLVADSSLQVNYSKENDKTKGTISNSVFSNDEEFCRRTQETSENVYTDMEGMKPVIGNG
RAFLPFASLRKSILSSISAEHGVGEEYISSHVTRLSSVINNFDKESQIASTSSPAAGVSNSFPVRAAQRKPRR
CWSPDLHRKFVDALQQLGGSEVATPKQIRELMNIDGLTNDEVKSHLQKYRLHTRRPNQNFHNMPSHTPQMVLI
GGIWVPEYTEHAAGVSQRNSADQHNYHMSLTSLSKDHCVGRNNIDKGQVHPAITCSQGSSQEPPSQSSDGSKE
EIMEDGTEAFSWKAGYQEDNSEIYDSASLHGS
>ScuHRS5_Sacu_v1.1_s0010.g004815
MGSVAELCLRRSLDYGWRRRMFSQSEEQRIESIKTQLQSLQDERRKIEAFKRELPICMQLLTDAIETSNEELL
EWSSGSLGAENYKDGVSDDDDRHSPAVPDDSNNDGRPFPAASATAAKHEMLACRTTSNRGATGAATAAMIVKN
SKSKPQPFYLEQMPDVICRNSKSSTKRPAAVHVDPMQVQEEELEEEEGTMNRCMSRISGTTTTRRSAAAAAEQ
PSLHMKSTFWTAAEAFAKGEEMLSTILTTAASKRRKHSHDDAVFAIGSCSNPHNNNIHDIDVTVELRSSSWRP
RGGGAFSPFPSSSKQLCSSSSNGTGSTPDSNSPPQLYSVDKRTPVHMNIYIPTDALHPPNDSNYPPAPPSSSV
ISSSMHDDDDDDDQEINTDDGHRRPYVNGTTAAEDVAPTSQTSSDHEGLVDVVEHWVDDDDDELIAAATGCAYG
GGEARDRLPFTGRGIHANLHGLMSQAPTTTTTTVPLTSMSSFRRKQRRCWSPELHRKFLSALHQLGGSEAATP
KLIRELMRVDGLTNDEVKSHLQKYRLHMRRPGGAVDATVVSTSEVAERRRRVGPRQLQAVNAVLDGVSAPRAD
AAAAESQCNFVDDSRTTSSQLLLLPPDDIDCERIINIEQQYLSSTCPTPVEVAAAYNRSELEVEQQGEMWRHL
HGAEEQEVSDRWKRRMCVNGLEERSRHMHHALFNTPRRGGKTGQGRAAQVGGQGGAQDGTGLSDTDRAGQSEA
KRSEAKLSGARRAHGKPERRKAGQSRARRRTERRRAGQSRAKQHRPAKGGERGTEAKAG

>PabHRS1_MA_14087g0010
MRRHEEYIKGLEEERKKIEVFKRELPFCMQLLNDAIEACKEQLADCERASLHEDVQNAPN
TNGRPVLEEFIPIKKSYQATENKLEENHVAKKARFNNGDKPNWMISANLWNPDPETIDSR
KGGGISMEESLKEQDSQYQVHSFPSNHKLFSDSKHRMGGAFHPFSKERQVIAPRPVRSTV
ERTLPNLALSSAEREVDSSLVGNETVCLDATTTTTARISRSKETTEVHTPVHEKEAVING
ANTSPTTSTPNTSQTQTQRKSRRCWSPELHRRFVNALQQLGGSQVATPKQIRELMKVDGL
TNDEVKSHLQKYRLHTRRPSPSPPSSNPQAPHLVVLGGIWVPPEYAAHAAAAAQQGQGGL
YNPLPTTVPCPPQSHYCQPSLPQGHEYYCQMNPAASKLQLHHSVSHYCDQQQEQQQQTPS
QSQNSPQGPLQCNGHSSGARGTSADAGREESVGEDAKSDSSSWKDEDSGETTEVMNMGRG
LSLRKQPHEFLDEDDGDATDNRARVQMGKQTLVGR

```
>PabHRS2_MA_33594g0010
MHLLNDAIAASKEQLTDCQPSVQAPRPVQQAGFQFEEEQYNRPVLEELIPIKKRCDESGD
ESPENNQNDSKRTKFANECKKEKSSSPEEPLKQHDSQLRSDQEQLFPTNSSLFSDSKQRI
GGAFLPFSRERQSNPSVSARSNAVKHPPDLALSSAEQEVGSTFRCNETATMNNVTTMNNV
TTTSLSRSKEPTETTVPTPQKEVVVKPNGGVCAVPTPAASNATQSGGQSQRKARRCWSPE
LHRRFVNALQQLGGSQVATPKQIRELMKVDGLTNDEVKSHLQKYRLHTRRPSPSPPTPAS
QAPQLVVLGGIWVPPEYAAHAAVAAHQGPGLYGTLSNTTSSHSQSHYCQSPMPQEHYSQM
NMSSQLQLPQPAYCEQQQTPSQSLSSPQGPLQLTGQSSGARGTSVDGGREESVGEDGKSE
SSSWKGEEIAKNVEARNMGRGLSLRKQPLQSVDGDEEDSRGSETTLKF
>AtrHRS1_scaffold00122.65
MEASRQQLASCSELTRSSSKPVLEEFIPIKRSSSEGSETTNMTSDKANWMTSAQLWSDQS
FPSSQKPAFNGKQRNGGAFLPFSKDRESRKTLSDLALASTEKEEIESPKFTEHEASGANK
MKENYNQVQGGGGGGLEREENKGSSGGQTQRKARRCWSPDLHRRFVSALQQLGGSQVATP
KQIRELMKVDGLTNDEVKSHLQKYRLHTRRPSPTPQAVPTPAPQLVVLGGIWVPQPDFSN
PSPNVNPSNPSQTLYTTTPPTQPHHFCQPTMAQDHLYAPTHRTYCDQMALAHMNPSHPNG
SVIEQRGGSESIEEDGKSESGSAENEVRDAGPALSLSRMHGHRHGNRGHDGDDMDEDGDG
DDDDGDDGDEDDSTGSEITLKF
>AtrHRS2_scaffold00033.259
MGSPSELSLDYKPPCKGEERFSKPSGANERFFKAVLEVGGNGVDRVVRIEELVACLEEER
RKIDVFKRELPLCMLLLNDAIDDLKEQARQCRLRGPVLEEFIPIKKAQEERENEAKAERD
CKDKMNWMSSAQLWSDNSDNSEEKKEKLDEYEAPSEELQKTDFGEKAYTIGRYRNSGGAF
VPLNGNGNGNGLVRKEENDIVDLSLSNGGIKELGHVSPKGSCRGNAGSSQSALQRKARRC
WSPELHRRFVNALQQLGGSQVATPKQIRELMKVDGLTNDEVKSHLQKYRLHTRRPNVSVS
SANSNQQNPPLVLLSNLWVPHDKLYNQTSSTASGSSSQSFQLVKKTASSQSGSPQGPFQY
PGKARTSVNGGDSGGEEDGLSDEETN
>AtrHRS3_scaffold00093.40
MEALEEEKRKIQVFQRELPLCLHLVTQAINTCREQLELSNEEVPARSGYGETSGEDPPFE
EFFPLKRSTSETTQAPAGEKKRKPEDQNDKPDWLRSVQLWNSDADPSRNEDAGLNPKLGT
KKQRGAFQPFSREREGRAEAVSGSGNERVKVREKEREREKEGQPNRKSRRCWSPELHRRF
LHALQQLGGSHTATPKQIRELMKVDGLTNDEVKSHLQKYRLHTRTPAAVPASQPINPQAP
QFVVVGGIWGLPEYAVPTPSPVTCGAIYATPIRPLPPPCQEAHQEKCSPTGPLHSDAGQG
VQREGGASFRDEHSASASSSSARAVSSEGNSAVL
```
