## Supplementary Dataset 2 for "Stepwise origin and evolution of a transcriptional activator and repressor system integrating nutrient signaling in plants"

Supplementary Dataset 2. Analysis of novel motifs in PSR and HRS transcription factors.
In each case, the sequence logo is given above and the sequence and p-value are given in the table below

### PSR Motif 1

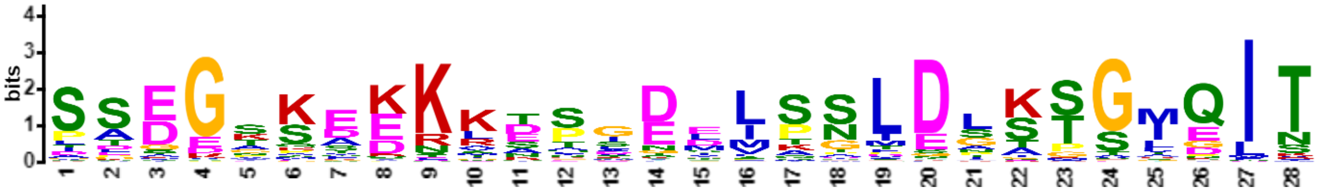

| Name | Start | *p*-value | Sites | | |
|---|---|---|---|---|---|
| F07065001 | 81 | 3.57e-26 | YRLAKYLPDS | SSDGKKADKKESGDMLSSLDGSSGMQIT | EALKLQMEVQ |
| M06G000800.1 | 111 | 5.64e-26 | YRLAKYLPDS | SSDGKKADKKETGDMISNLDGSSGMQIT | EALKLQMEVQ |
| O129805.m001544 | 111 | 7.07e-26 | YRLAKYLPDS | SSDGKKADKKETGDMLSNLDGSSGMQIT | EALKLQMEVQ |
| N08G014900.1 | 111 | 7.07e-26 | YRLAKYLPDS | SSDGKKADKKETGDMLSNLDGSSGMQIT | EALRLQMEVQ |
| G030480t1 | 170 | 7.07e-26 | YRLAKYLPDS | SSDGKKADKKETGDMLSNLDGSSGMQIT | EALKLQMEVQ |
| H27.185 | 81 | 3.98e-25 | YRLAKYLPDS | SSDGKKTDKKETGDMLSNLDGSSGMQIT | EALKLQMEVQ |
| K15G123100.1 | 112 | 9.04e-25 | RLAKYLPDSS | SDEGKKADKKETGDMLSNLDGSSGMQIT | EALKLQMEVQ |
| K09G017400.1 | 112 | 9.04e-25 | RLAKYLPDSS | SDEGKKADKKETGDMLSNLDGSSGMQIT | EALKLQMEVQ |
| PHL7_AT2G01060 | 77 | 1.11e-24 | YRLAKYLPDS | SSEGKKTDKKESGDMLSGLDGSSGMQIT | EALKLQMEVQ |
| G003939t1 | 323 | 1.11e-24 | KYRTARYRPE | SSEGSSEKKLTPIEDLSSLDLKTGMGIT | EALRLQMEVQ |
| M02G257800.1 | 344 | 2.43e-24 | KYRTARYRPE | SSEGSSEKRLTSIDEISSLDLKTGIEIT | EALRLQMEVQ |
| L2g027860.1 | 112 | 4.32e-24 | RLAKYLPDCS | SDEGKKTDKKETGDMLSNLDGSSGMQIT | EALKLQMEVQ |
| J3832766 | 77 | 1.10e-23 | YRLAKYLPDS | SSEGKKTDKKESGDVLSGLDGSSGMQIT | EALKLQMEVQ |
| F28403001 | 242 | 1.89e-23 | KYRTARYRPE | SSEGSSEKRLTSIEEMSSLDLKTGIEIT | EALRLQMEVQ |
| N06G035600.1 | 111 | 2.27e-23 | YRLAKYLPDS | SSDGKKADKKETGDMLSNVDGSSGMQIT | EALKLQMEVQ |
| EM101479.1 | 81 | 2.71e-23 | YRLAKYLPDS | SSDGKKAENKESGDILSGLDGSSGMQIT | EALKLQMEVQ |
| B010973 | 300 | 3.85e-23 | KYRTARYRPD | SSEGSAEKKKTAIEEISSLDLKTGIDIT | EALRLQMEVQ |
| NS107600.1 | 272 | 6.49e-23 | KYRTARYRPD | SSEGSSEKKLTPMDEISPLDLKTGIEIT | EALRLQMEVQ |
| AB10.1900 | 277 | 1.08e-22 | KYRTARYRPD | SSEGTSEKKSNPLDEMSSLDLKTGIEIT | EALRLQMEVQ |
| EH101641.1 | 75 | 1.52e-22 | YRLAKYLPDS | SSDGTKAEKKESGDILSSLDGSSSLQIT | DALKLQMEVQ |
| Y728141g003 | 530 | 2.12e-22 | KYRTARYRPE | SSEGTSAKKITPLEEMPSLDLKTGIEIT | KALRLQMEVQ |
| K19G167500.1 | 319 | 2.12e-22 | KYRTARYRPE | SSEGAAEKKLSPIEEMSSLDLKTGIEIT | EALRLQMEVQ |
| C5G018400 | 319 | 2.12e-22 | KYRTARYRPE | PSEGNSEKRATTIDDISSLDLKTGIEIT | EALRMQMEVQ |
| N12G035600.1 | 325 | 4.79e-22 | KYRTARYRPD | SSEGSSEKKFTPREEISSLDLKTGIEIT | EALRLQMEVQ |
| L7g098250.1 | 323 | 5.63e-22 | KYRTARYRPE | SSEGAGEKKLSPIEDISSLDLKTGIEIT | EALRLQMEVQ |
| Y714041g001 | 80 | 6.60e-22 | YRLAKYVPES | SSDGSKGEKKDPGDLLSGLENSSGMQIT | EALKLQMEVQ |
| B012506 | 300 | 7.74e-22 | KYRTARYRPD | SSEGSSEKKMSAIEEMTSLDLKTGIEIT | EALRLQMEVQ |
| AB01.3515 | 110 | 1.24e-21 | YRLAKYIPES | SSDGSKGEKKDPGDMISGIENSSGMQIT | EALKLQMEVQ |
| X23131.1 | 80 | 1.24e-21 | YRLAKYIPES | SADGMKTEKKDPGDLLSGLESSSGMQIT | EALKLQMEVQ |
| N08G140300.1 | 284 | 1.24e-21 | KYRTARYKPE | SSEGTSEKKLSPIEEMKSLDLKTSMGIT | EALRLQMEVQ |
| A061.72 | 108 | 1.98e-21 | YRLAKYLPES | SSDGTKPDRKESADMLSNLDAASGMQIT | EALRMQMEVQ |
| Y6550989g009 | 107 | 2.31e-21 | YRLAKYLPES | PADGSKDEKKDSGDSLSSMDSASGIQIN | EALKIQMEVQ |
| Mpo3115s0048 | 115 | 2.69e-21 | RLAKYIPESM | SDGGKSEKKKNPADIIPSLDATSGIQIT | EALRMQMEVQ |
| AA24486 | 109 | 3.13e-21 | YRLAKYLPDS | SAEGAKVEKKESDDVLSNLESSTGMQIT | EALKLQMEVQ |
| Y714681g011 | 110 | 3.13e-21 | YRLAKYIPES | SSDGTKAAKKDSGDLLSGLENSSGMQIT | EALKLQMEVQ |
| J3827611 | 77 | 3.13e-21 | YRLAKYLPDS | SSEGKKTDKKESGDVLSGLDGSPGTQIT | EALKLQMEVQ |
| AC2_310V3.1 | 114 | 5.72e-21 | YRLAKYIPES | SSDGGKSEKKNPADVLPTLDATSGIQIT | EALRMQMEVQ |
| C1G011200 | 107 | 8.91e-21 | YRLAKYLPES | PADGSKDEKKDSGDSLSSMDSAPGIQIN | EALRMQMEVQ |
| M16G001100.1 | 90 | 1.03e-20 | YRLAKYLPDS | SSDGKKVDKKETGDVLSNSDGSSGMQIT | EALKLQMEVQ |
| AD466s0002.1 | 29 | 1.38e-20 | RLAKYIPDSL | SDGGTSDKKKTSADLFPSLDATSGIQIT | EALCMQMEVQ |
| AD006s0186.1 | 117 | 1.38e-20 | RLAKYIPDSL | SDGGTSDKKKTSADLFPSLDATSGIQIT | EALRMQMEVQ |
| AD170s0003.1 | 122 | 1.85e-20 | RLAKYIPDSF | SDGGTSEKKKNPADLFPSLDATSGIQIT | EALRMQMEVQ |
| Y916311g004 | 107 | 2.83e-20 | YRLAKYLPES | PADGSKDEKKDSADTLSSMDSASGIQIN | EALKMQMEVQ |
| AD119s0044.1 | 118 | 3.76e-20 | RLAKYIPDSL | SDGGTSDKKKNPADLFPSLDATSGIQIT | EALRMQMEVQ |
| O29637.m000753 | 325 | 4.98e-20 | KYRTARYRPD | SLEGSSEQKLTPLEEISSLDLKTGIEIT | EALRLQMEVQ |

| | | | | | |
|---|---|---|---|---|---|
| ED00466.1 | 323 | 4.98e-20 | KYRTARFKPE | TPEESSEKKLASIEDLSSLDLKTGIEIT | EALRLQMEVQ |
| U0810G0130 | 200 | 8.67e-20 | KYRSAKYMPA | SSEGKQREKRATGNDIQNLDPKTGMQIT | EALRAQLDVQ |
| O29680.m001711 | 294 | 8.67e-20 | KYRTARYKPE | SAEGTSEKKLSPIDEMKSLDLKASMGIT | EALRLQMEVQ |
| W5g41240.1 | 107 | 1.31e-19 | YRLAKYIPES | PAEGSKDEKKDSSDSLSNTDSAPGMQIN | EALKMQMEVQ |
| K03G166400.1 | 316 | 1.31e-19 | KYRTARYRPE | SSEGAAEKNLSRIEEMSSLDLKTGIEIT | EALRLQMEVQ |
| W4g56990.1 | 302 | 1.71e-19 | KYRIAKYMPA | SSEGKQLEKRATGNDMQNLDPKTGMQIT | EALRVQLDVQ |
| D11g067280.1 | 78 | 1.71e-19 | YRLAKYLPDS | SSDGKNSDKKEPRDMLSSLDGSSGVEIT | TALKLQMEVQ |
| B004596 | 109 | 1.71e-19 | YRLAKYLPES | PADGSKDEKKDSGDSLSSMDSAPGVQIN | EALRMQMEVQ |
| M19G020900.2 | 261 | 1.96e-19 | KYRTARYKPE | SSEGTSEKKLSPVEEMKSLDLKTSMEIS | EALRLQMEVQ |
| AA26158 | 37 | 2.56e-19 | YRLAKYIPDS | SSDGLKDESKEPGDLLSGLDSSSGVQIT | EALKLQMEVQ |
| A142.28 | 333 | 3.34e-19 | KYRTARYRPD | ISEGNAEKKMTRLDDISTLDLKTGIEIT | EALRLQMEVQ |
| N09G147400.1 | 295 | 7.32e-19 | KYRTARYKPE | ASEGTSENKLSSIDEMKSLDLKTSIGIT | EALRLQMEVQ |
| D06g008200.2 | 324 | 1.39e-18 | KYRTARYKPE | ASEGSSEKKESSIGDLSALDLKTGIEIT | EALRLQMEVQ |
| V2g21407.2 | 107 | 1.79e-18 | YRLAKYIPES | PAEGSKDEKKDSSDSLSNTDSAPGLQIN | EALKMQMEVQ |
| S09G180500.1 | 107 | 1.79e-18 | YRLAKYIPES | PAEGSKDEKKDSSDSLSNTDSAPGLQIN | EALKMQMEVQ |
| R2G117854 | 107 | 1.79e-18 | YRLAKYIPDS | PAEGSKDEKKDSSDSLSNTDSAPGLQIN | EALKMQMEVQ |
| K03G250000.1 | 291 | 1.79e-18 | KYRTARYKPE | PSEGTSEKKVTPMEEMKSLDLKTSKGIT | EALRLQMELQ |
| AC14_26270V3.1 | 115 | 1.79e-18 | RLAKYIPESL | SDGGKSDKKKNQADLLPALDATSGIQIT | EALRMQMEVQ |
| K19G247600.1 | 271 | 3.34e-18 | KYRTARYKPE | PSEGNSEKKVTPMEEMKSLDLKTSKGIT | EALRLQMELQ |
| AC17_23560V3.1 | 115 | 3.34e-18 | RLAKYIPESL | SDGGKSDKKNNPTDLLPTLDATSGIQIT | EALRMQMEVQ |
| W9g12770.1 | 85 | 4.28e-18 | YRLAKYIPDS | SADGNKAENKDPGDLLAGLEGSSGLQIS | EALKLQMEVQ |
| V4g27620.1 | 85 | 4.28e-18 | YRLAKYIPDS | STDGNKSDNKDPGDSLAGLDGSSGLQIS | EALKLQMEVQ |
| K10G039700.1 | 320 | 4.28e-18 | KYRTARYRPE | SSEGVMDKKTSSVEEMSSLDLRTGIEIT | EALRLQMEVQ |
| D09g072830.2 | 312 | 4.28e-18 | YRTARYKPEA | SEAGSSEKKQSSLDDLASLDLKTGIEIT | EALRLQMEVQ |
| D05g055940.2 | 314 | 8.88e-18 | KYRTARYKPE | SSEGTPEKKTTSVTEMPSLDLITTMGIT | EALRMQMEVQ |
| X14587.1 | 311 | 1.00e-17 | KYRTARYRPE | SSEGASEKKGTSQEELSSLDLKTSFDLT | EALRLQMEVQ |
| K13G126200.1 | 260 | 1.00e-17 | KYRTARYRPE | SSEGVMEKKTSSVEEMASLDLRTGIEIT | EALRLQMEVQ |
| AA22301 | 271 | 1.62e-17 | KYRTARYQPE | WADGTSEKKASSLDKLASLDLKTGMELT | EALRLQIEVQ |
| J3854324 | 296 | 1.62e-17 | YRTARYRPEP | SETGSSEKKLTPLEHITSLDLKGGMCIT | EALRLQMEVQ |
| R2G173943 | 107 | 2.59e-17 | YRLAKYIPDA | STDGNKTDNKDPGDLLAGLEGSSGLQIS | EALKLQMEVQ |
| U1536G0490 | 107 | 2.91e-17 | YRLAKYIPES | PAEGSKDEKKDSSDSLSITDSAPGLQIN | EALKMQMEVQ |
| L1g080330.1 | 236 | 2.91e-17 | KYRTARYKPE | SSEGIPEKKLTSIDEMPSIDLKTPKGIT | EALRLQMELQ |
| L7g115530.1 | 279 | 3.67e-17 | KYRTARYKPE | SPEETSEKKMSSIEEMKSLDLKTSKGIT | EALRLQMELQ |
| F23401001 | 245 | 5.19e-17 | KYRTARYKPK | LSEGTSDKNLTSIGEITSLDLKMSMGIT | EALRLQMEVQ |
| A095.37 | 109 | 5.19e-17 | YRLAKYLPDS | PADGTKDEKKESSDILPNMESAPGAQIS | EALRMQMEVQ |
| G026171t1 | 291 | 8.19e-17 | KYRTARYKPE | SSEGTLENKMASIGEMKSLDLKAGMGIT | EALRLQMEVQ |
| P1G0009300 | 59 | 9.17e-17 | KYRTARYRPE | ALEGAAEKKVSPIDKISSLDRKSGIDIT | EALRLQMEVQ |
| T6G121500.1 | 105 | 1.29e-16 | YRLAKYIPDP | STDDNKAEKKDPGDLLAALEGSSTMQIS | EALKLQMEVQ |
| Z2P10180_001 | 305 | 2.01e-16 | KYRTARYIPD | SSEGMSEKRITQSEELPSLNLKTGIDFT | EALRLQIEVQ |
| T2G168900.1 | 108 | 2.24e-16 | YRLAKYIPDA | STDGNKADNKDPGDLLAGLEGSSGLPIS | EALKLQMEVQ |
| S02G161800.1 | 107 | 2.24e-16 | YRLAKYIPDA | STDGNKADNKDPGDLLAGLEGSSGLPIS | EALKLQMEVQ |
| Z5P12960_001 | 142 | 2.50e-16 | KYRTARYRPD | LSEGMSEKKITQSQEIPSLDLKTGIDLT | EALRLQMEVQ |
| P22G0018300 | 114 | 2.50e-16 | YRLAKYLPES | PADGLKDEKKDCSDGLGNLDPASGLQIT | EALKMQMEVQ |
| W8g25820.1 | 107 | 2.79e-16 | YRLAKYIPDP | SADDNKDEDKDPGNLLSALEGSSGMQIS | EALKLQMEVQ |
| J3834747 | 271 | 2.79e-16 | YRTARYRPEP | SESGSPEKKLTPLEHITALDLKGGIGIT | EALRLQMEVQ |
| PHR1_AT4G28610 | 287 | 2.79e-16 | YRTARYRPEP | SETGSPERKLTPLEHITSLDLKGGIGIT | EALRLQMEVQ |
| Z4P30200_001 | 277 | 6.65e-16 | KYRTARYKPD | SLEGMSEKTATQSEELPSLDLKTGIDFT | EALRLQMEVQ |
| F25867001 | 107 | 8.23e-16 | YRLAKYLPES | PADGSKDEKKGSGDSGSSMDSAPGVQIN | EALRLQMEVQ |
| H33.181 | 107 | 1.02e-15 | YRLAKYLPES | PADGLKDEKKDAGDSITTPDSSPGVQIN | DTLRMQMEVQ |
| Z8P12580_001 | 178 | 1.26e-15 | KYRTTWYRPD | SSEGPSNKKTTLKEELPSLDLKTSFDLT | EALRLQVVVQ |
| P12G0022900 | 110 | 1.26e-15 | YRLAKYLPES | SPDGIKAEQKEPGSLLSTLENSSGIQIT | EALKLQMEVQ |
| EF02036.1 | 238 | 1.26e-15 | KYRTARYKPE | SSEGTSEKKSSTPTDMTSVDLKTTMGIT | EALRVQMEVQ |
| Z3P28410_001 | 80 | 1.91e-15 | YRLAKYVPES | SADGTMSEKKDDRNLNSGLESSSGMQIT | EALKLQMEVQ |
| R2G162409 | 303 | 1.91e-15 | KYRTARYRPE | LSEGSSEKKVASKEDIPSIDLKGSFDLT | EALRLQLELQ |
| T2G118400.1 | 303 | 2.35e-15 | KYRTARYRPE | LSEGSSEKKAASKEDIPSIDLKGSFDLT | EALRLQLELQ |
| S02G121600.1 | 303 | 2.35e-15 | KYRTARYRPE | LSEGSSEKKAASKEDIPSIDLKGSFDLT | EALRLQLELQ |
| Z3P11970_001 | 80 | 3.20e-15 | YRLAKCVPDS | SADDAKSEKKDPDGVSSGLESSSGTQIT | EALKLQMEVQ |
| J3854619 | 247 | 4.35e-15 | TARYKPETSE | ATGEPQEKKMTSIEDIKSLDMKTSVEIT | QALRLQMEVQ |
| U1079G0490 | 289 | 4.82e-15 | KYRTARYRPE | LSEGSSEKKVASEEDIPSIDLKGSFDLT | EALRLQLELQ |
| EF01810.1 | 210 | 7.21e-15 | KYRTARYKPE | SSEGALERKSKNIAEMTSLDLKTTMGIT | EALRVQMEVQ |
| PHL1_AT5G29000 | 295 | 9.73e-15 | TARYKPETSE | VTGEPQEKKMTSIEDIKSLDMKTSVEIT | QALRLQMEVQ |
| H2085.1 | 228 | 1.07e-14 | ATPKGVLKLM | NVEGSSEKKSIPLDDVKSLDLKTSMSIT | EALRLQMEVQ |
| W6g40710.1 | 109 | 1.19e-14 | YRLAKYIPDP | TADGAKSDKKDLGDLLADIESSSGMEIG | EALKLQMEVQ |

| | | | | | |
|---|---|---|---|---|---|
| Y803301g004 | 310 | 1.59e-14 | KYRTARYRPD | SSEGASEKMVTPQQEVSSLDLKTSINLT | EALQLQMEVQ |
| X19267.1 | 178 | 1.59e-14 | YRLAKFLPES | PADGSKDETKDSGDTLSGNDSAPEIQIN | EALKMQMEVQ |
| PHL13_AT3G04450 | 302 | 1.59e-14 | TARYKPELSK | DTEEPLVKNLKTIEDIKSLDLKTSIEIT | EALRLQMKVQ |
| Z4P11370_001 | 80 | 1.76e-14 | YRLAKYVPES | SADGTRSEKKDDDNQISGPESSSGTQIT | EALKLQMEVQ |
| J3837308 | 288 | 1.76e-14 | TARYKPETSE | ATGEPEEKKITSIEDIKSLDMKTSVEIT | QALRLQMEVQ |
| Q262g00120.1 | 86 | 3.15e-14 | RLAKYVPDEP | SSDGVNDEKKDRDELLSNSNNSSGSQIT | EGMKLQMEVQ |
| J3838005 | 224 | 3.47e-14 | KYRTARYVPE | PSQGSQETKLTPLEHVTSLDTKRGIDIT | EALRIQMEVQ |
| AB02.1982 | 161 | 4.62e-14 | YRLAKYLPES | PGDGSKDEKKDLGETSSGTESAGGIQIN | EALKMQMEVQ |
| V3g22290.2 | 110 | 5.59e-14 | RLAKYIPDPS | ASDDNKAEERDPGDLLAALEGSSGMPIS | EALKLQMEVQ |
| L7g088070.1 | 114 | 6.15e-14 | LAKYLPESPG | DGKDSKDEKRNSGDSISGADSSPGLQIN | DALRMQMEVQ |
| Q70g00660.1 | 87 | 6.75e-14 | YRLAKYLPDS | SSDGLKAEKKDSGDLLSNLDNSSSGSQI | TEAMKLQMEV |
| PHL4_AT2G20400 | 292 | 7.42e-14 | KYRTAKYIPV | PSEGSPEARLTPLEQITSDDTKRGIDIT | ETLRIQMEHQ |
| AF031.g024472 | 115 | 8.95e-14 | YRLAKYIPDS | GKDGEKSEKSASTDMLNNLDSAAGLQIT | EALRMQMEVQ |
| Z7P06180_001 | 110 | 9.83e-14 | YRLANYIPDS | SADGAKSENKDSGDHTMRPENSSGMQMT | EALRLQMEVQ |
| J3848918 | 266 | 9.83e-14 | TARYKPEISI | DTEKPPLKTLKTIEDIKSLDLKTSIEIT | EALRLQMEVQ |
| AD014s0203.1 | 119 | 9.83e-14 | AKYIPDSLSD | GGAGGISDKKNPADMFPSLEPTSGFQIT | EALRMQMEVQ |
| Z11P02560_001 | 285 | 1.18e-13 | KYRTARHRPD | SSEEIFNKKITLKEEIPSLDLKTSFDLT | EALQLQMEVQ |
| AB05.2141 | 80 | 1.30e-13 | YRLSKYLPDS | TSDGSNVEKGEQEGQLSTIDNSSGMQIT | EALKLQMEVQ |
| V1g36236.2 | 114 | 1.30e-13 | YRLSKYIPDP | TADGAKSDKKELGNLLAGIESSPGMELS | EALKLQMEVQ |
| Y681141g002 | 262 | 2.26e-13 | KYRTARYRPD | SSEGASQKKVTPQEEVSSLDLKTVSNIL | MKTPKIREDP |
| V5g25320.1 | 289 | 2.26e-13 | VKCVPSSSSS | SEGKQQEKRAAGSDDVPNLDPKTGMHIT | EALRVQLDVQ |
| D11g022470.1 | 81 | 2.47e-13 | YRLAKYLPDS | SSDGKQSDKKESGDMLSSLDGSSTGVQI | NEALKLQMEV |
| R2G124495 | 330 | 2.96e-13 | YRIAKYMPVS | STSEGKEKRAAAANDVQNLDPGTGMKIT | EALRVQLDVQ |
| T3G029000.1 | 303 | 3.88e-13 | YRIAKYMPAS | TSSEGKQEKRAAGNDAQNLDPSTGSQIT | EALRVQVDVQ |
| K03G003500.1 | 100 | 5.08e-13 | LAKYLPESPA | DGKDSKVEKRNSGDSISGADSSPGMPIN | DALRMQMEVQ |
| H85.82 | 268 | 5.55e-13 | KYRIAKYMSD | SAEGKSEKRSSTNYIPYQLDTKTGMQIK | EALELQLDIQ |
| K19G122700.1 | 110 | 7.90e-13 | LAKYLPESPA | DGKDPKDEKRMSGDSISGADSSSGMPIN | DALRMQMEVQ |
| AC126_3820V3.1 | 189 | 7.90e-13 | KYRLAKYMPE | ISEEAKAERRKHDCLLTSLDLGSGHQIA | QALQMQMEVQ |
| X10145.1 | 217 | 8.63e-13 | KYRTARYKPE | SSEGTSEQRVIQVEETKSLDLKTVTEIN | EALRVQMEVQ |
| C2G381900 | 312 | 1.12e-12 | KYRTARYGPD | LTDGTSEKKMSAAEETPSLSPKMSIEIS | EALRLQMEVQ |
| C7G341000 | 80 | 1.22e-12 | KYRLAKYLPD | AASDGKGEDKKESGDLSSMDNSSGMQIT | EALKLQMEVQ |
| AA30954 | 84 | 1.33e-12 | KYRLAKCLPD | SSAEVTRELKESDDVLSKLDNSSGIQIS | EALKLQMEVQ |
| P11G0061900 | 159 | 2.89e-12 | KYRTARFRPD | SSEGAPETNLAPLEGLSSVDQGTGLGIT | EALRLQMEVQ |
| AF078.g038309 | 114 | 3.72e-12 | YRLAKYIPES | GKDGETSEKSGSTEMLNNLEGAPGIQIT | EALRMQMEVQ |
| J3828177 | 255 | 6.14e-12 | YRTARHVPEP | SEAGWREKKLTPVEHVTSLDTKRFGNIT | VHNGMYISEA |
| ZP18240_001 | 107 | 6.67e-12 | YRLAKYLPDS | PADGSKDDKKDSGHTHSDTDSAPSAQFN | ESLKMQMEVQ |
| AC4_23130V3.4 | 275 | 7.24e-12 | KYRLAKYMPE | ISEEAKAERRKHDSLLTSLDLGSSYQIA | QALQLQMEVQ |
| W7g25710.1 | 307 | 8.54e-12 | KYRTARYRPE | LSEGSSEKKAASKEDIPSIDLKGGNFDL | TEALRLQLEL |
| U2891G0030 | 85 | 1.77e-11 | YRLAKYIPDS | STDANRTDNNDPGDLVAGLEGSSRLQIS | EALKLQREVQ |
| R2G035370 | 354 | 2.08e-11 | YMPASSTSEG | NLIYRKQEKRAVGNDVQNLDPSTGMKIT | EALRVQLDVQ |
| V1g28920.4 | 304 | 2.25e-11 | KYRTARYRPE | LSEGSSERLDASKEELPSIDLKGNFDLT | EALRLQLELQ |
| AH10433112g0010 | 271 | 3.35e-11 | KYRLAKYIPE | SLEGGKSDKEKGNKIVSNLEVTSGTQIA | EALQMQMEVQ |
| AH10266929g0010 | 109 | 4.59e-11 | KYRIAKYIPE | SLEGGKSGKKKGNKIVSNLEAISGTQIV | EALQMQMEVQ |
| N109G033500.1 | 303 | 5.36e-11 | KYRIAKYMPD | SSEGKSEKRTTINDVPQIDTKTSGLQIS | EALQLQLDVQ |
| AC13_34900V3.1 | 194 | 7.30e-11 | KYRLARYMPE | ITEEQKAERRRTESLLTPLEISSSYQIT | QALQMQMEVQ |
| F11163001 | 83 | 8.51e-11 | KYRISKFVPE | SSSRAKFERRSISEMLPNFSTTSGAQLK | EALQMHMEVE |
| EJ01825.1 | 241 | 2.83e-10 | KYRNAKYVPE | SDAPGKSEKKTSSNNAAQIDIKTGTQLK | EALQLQLDVQ |
| Q6g00200.1 | 107 | 3.05e-10 | YRLVKYLPES | PINGSTSDKKESTDNLMNMASEPGIKIN | EALKMQMEVQ |
| N09G105800.1 | 72 | 3.28e-10 | KYRISKFIPE | SNNKGKFERRNISELLPNFSATSGAQLN | EALKMQMEVQ |
| AD002s0165.1 | 421 | 3.28e-10 | KYRLAKYSPD | ISDEARAEWRRSDTYLSPIGINSSHQIT | QALQLQMQVQ |
| K103G143600.1 | 313 | 6.77e-10 | GSFPILNLKN | CPVSGKSDKRTHTKDVHHLDVKTGIQIR | EALKLQLDAQ |
| AF002.g001571 | 241 | 7.27e-10 | YRTAKFIPTG | DSSEGKLDKRQSVDEPPDHDAKTSIQMM | EALQMQMEMQ |
| G021688t1 | 73 | 8.39e-10 | KYRILKFVPE | TNTKCKFERRDISEILPNFGTTSGAQLN | EALQMHKEAE |
| EL02017.1 | 73 | 8.39e-10 | YRMSVFTREE | DHVGGKFEKTSASEMLPNFSAISSAQLN | EALLIQMEAQ |
| N08G045400.1 | 303 | 1.11e-9 | KYRIAKYMPD | SLEGKPERRNSISNVSQIDTKTSGMQIT | EALQLQLDVQ |
| L7g068600.1 | 77 | 1.19e-9 | KYRISKLIPE | STTRGKIEKRSISDILPNFCSISALQLK | EVLQMQAEVQ |
| F33528001 | 26 | 1.28e-9 | DFPQKVNIWI | VSIAGKSEKGASSSDVPHLDNEDGMQIR | EALQLQLDLQ |
| AD191s0003.1 | 432 | 1.28e-9 | KYRLAKYMPE | ISEEARAERRRNDTFLSPMGINSSHQIT | QALQMQMEVQ |
| AD001s0234.1 | 415 | 1.58e-9 | KYRLAKYMPE | ISEETRAERRRNDTYLQPMGISSSQQIT | QALQMQMEVQ |
| AD016s0215.1 | 432 | 3.15e-9 | KYRLAKYIPE | ISEEARAERRQNDAYLSPMGINSTHQIT | QALQLQMEVQ |
| K18G201800.1 | 78 | 4.12e-9 | KYRISKLIPE | SPTRGKLEKRSMSDILPNFSSITALQLK | EVLQMQTGMQ |
| C2G092800 | 74 | 8.03e-9 | KYRISKFIPE | TYDRGRLDRRKVSELLPNFSTTSAAQIN | ELLQTQIEVP |
| Y695651g003 | 325 | 1.35e-8 | KYRIAKYMPE | SSEGPSQIELNEAGNLMFPYFCSGMQIT | EALQLQLDVQ |

| | | | | | |
|---|---|---|---|---|---|
| AE270428 | 94 | 2.12e-8 | KYRLVKDLPP | SPVAKQQQSKQCSLELPSLNVETGLQIT | ETLRLQLEVQ |
| D05g007890.2 | 86 | 3.50e-8 | LGKSSVTDQS | FDENKQEVKLDLCEIVPNDDDTKGNEIS | RGLKQRTYFD |
| F29458001 | 302 | 5.08e-8 | KYMPERKEDK | KASGSEEKKAASSNNESDGRRKGNIQIT | EALRLQMEVQ |
| AG064.g015782 | 111 | 6.89e-8 | KYRLAKYIPD | SGKDSEKPEKSSTDMLNNLHATSDLQLT | EALRMQMEVQ |
| AF003.g007457 | 274 | 1.79e-7 | MTQDIPGSQR | DEHGRKRRSSSSMEAITMLDGSMALQMT | EALQLQMEMQ |
| Mpo198s0044 | 117 | 1.20e-6 | VNVETIKDQG | SSEGQGPSSGTATDCVITQNPKESLQIT | EALRLQMEVQ |
| O29827.m002675 | 89 | 4.67e-6 | LGQQQARKQN | TKEQYKENSGASYVNFSNHSSSSGLHAT | SSSNHNQQGE |

### PSR Motif 2

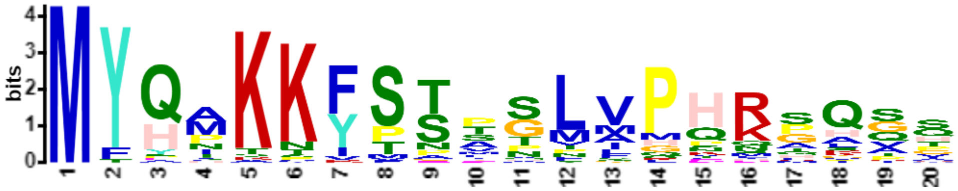

| Name | Start | *p*-value | Sites | | |
|---|---|---|---|---|---|
| Mpo198s0044 | 1 | 8.11e-21 | | MYQMKKYSNTSLVPHRGQTS | SQQERSMYGG |
| B004596 | 1 | 5.14e-20 | | MYHAKKFSTMTLVPHKPQPQ | GADQLPNVGV |
| AB02.1982 | 55 | 6.11e-20 | EDVVDDQSIT | MYHAKKFSTMPLVPHRAQAT | EQPPDTGVMV |
| F25867001 | 1 | 6.11e-20 | | MYHPKKFSTASLVPHKAQGS | EPLATVGALG |
| AG064.g015782 | 1 | 7.23e-20 | | MYQAKKFSTTNLVPHRGAAP | LDQHYHHGSS |
| AC3_14670V3.1 | 1 | 1.40e-19 | | MYQMKKYSSVGLVPHRQHQV | SPYASAVSCE |
| A095.37 | 1 | 1.65e-19 | | MYYPKKFSTMGLVPHKPQSQ | GADQLANIGV |
| AD1140s0043.1 | 1 | 1.65e-19 | | MYQMKKYTSPGLIPHRSQTS | SSNQQDRSSL |
| AF078.g038309 | 1 | 1.18e-18 | | MYQAKKFSTTSLVPHRGPPG | VADQHYPPGS |
| AF031.g024472 | 1 | 1.18e-18 | | MYQAKKFSTTNLIPHRGAPG | GSDQQLNYVH |
| AD1173s0030.1 | 1 | 1.57e-18 | | MYQMKKYTSPGLIPHRTQIP | SPNYQDRSSL |
| AD078s0059.1 | 1 | 2.39e-18 | | MYQMKKYTSPGFMPHRTQTS | SSNLQDRLAL |
| AC2_310V3.1 | 1 | 2.75e-18 | | MYQNKKFSTTSLMPQRSASQ | APEPRGSSAT |
| AC17_23560V3.1 | 1 | 2.75e-18 | | MYQNKKFSTTSLMPQRSASQ | ALELRGSSAA |
| AC14_26270V3.1 | 1 | 2.75e-18 | | MYQNKKFSTTSLMPQRSASQ | APEPRGSSAA |
| H33.181 | 1 | 3.15e-18 | | MYHAKKISTANLVPHKPQGA | EQFPSTGTLG |
| C1G011200 | 1 | 8.04e-18 | | MYQARKFSTVGLVPHKAQSA | EQHTSIGSLG |
| Y6550989g009 | 1 | 9.15e-18 | | MYHAKKFSTISMLPHKAQGT | EQLANAGVMG |
| K19G122700.1 | 1 | 2.53e-17 | | MYHTKKFSPASMVPHKSQGG | AEQLANAGVL |
| AC13_23440V3.1 | 3 | 4.66e-17 | MA | MYQMKKYSSLALAPHRHHQV | SSYAPTLSTG |
| X19267.1 | 15 | 6.68e-17 | AVLKSLRVFA | MYHPKKFSTLHMAPHKSQGS | EQTGNVSVMD |
| AH10266929g0010 | 1 | 1.70e-16 | | MFQNKKFSTDGLVPQRFHGS | EQRGAGPLGV |
| AC3_23200V3.3 | 1 | 2.99e-16 | | MYQIKKYPNPGMNPHRQQQI | MSFQDRGPSM |
| AE149357 | 1 | 4.65e-16 | | MYQMKKYPSPQLIPHRGGAM | PAQSEPLYIA |
| Mpo3115s0048 | 1 | 5.19e-16 | | MYAAKKFSTSSLVPQRPGQV | PDPRGSNAAG |
| AH10433112g0010 | 163 | 5.79e-16 | SLILSASASI | MFQNKKFSTAGLVPQIFHGS | KQRGAGPLGV |
| AC4_20290V3.1 | 1 | 5.79e-16 | | MYQMKKYSSLTVVPPQGQQH | LMTSPHNQDH |
| Q6g00200.1 | 1 | 3.81e-15 | | MYHAKTISTATLVPHKTKVT | NQHPNSEYLG |
| AC10_13030V3.1 | 1 | 8.80e-14 | | MYQMKKYVSPGLSSHMHQQI | MLLEDRGPSM |
| ZP18240_001 | 1 | 1.37e-13 | | MYHASKFSTVSMAPDNGQGT | EQVGSIGISG |
| L7g088070.1 | 1 | 3.01e-13 | | MYHAKKFPEETMMLHKSQGG | GGGGGEQFAN |
| A061.72 | 1 | 7.62e-13 | | MYHPKNISSTTLVPTKPSSR | DPQIDNAKMA |
| AD006s0186.1 | 1 | 1.59e-12 | | MYQTTKFSASSVMPERPKSQ | VVEPWAPSPA |
| AC18_20920V3.2 | 1 | 6.60e-12 | | MFSMKKYMNPGTFQQRHQSS | GSLYIPDKLQ |
| AC21_2850V3.1 | 1 | 9.68e-12 | | MFPMKKYMSPGTFQQRPRST | GSLYIPEKLQ |
| N06G035600.1 | 1 | 1.31e-11 | | MYQPKCIPSSSLVHKSSLVH | GQYLDCGANS |
| AD173s0028.1 | 1 | 1.52e-11 | | MYQIKKYTNPPSLMPHGRAQ | HHTSLHQQLE |
| AD088s0085.1 | 1 | 3.43e-11 | | MYQIKKYTAANPAGSLMQQP | GLHRPQQSTT |
| K09G017400.1 | 1 | 4.26e-11 | | MYHSKNFPSASLIGVNSLVH | GQHIDCGGST |
| AD078s0062.1 | 1 | 6.10e-11 | | MYQIKKYTSSPPGGLMPHRP | QTSPHLERPS |
| AE417629 | 1 | 8.70e-11 | | MYQVKNFSPGTNFMSQQQQQ | QPTLYTGLSV |
| W6g40710.1 | 1 | 1.87e-10 | | MYEPKPFSSIVLAHNDPVSH | NQQIERINNN |

| | | | | | |
|---|---|---|---|---|---|
| G030480t1 | 60 | 2.00e-10 | RSASSSDDQQ | MYQPKTVPGSSLVRNNSIVH | GQHLDCGASQ |
| O129805.m001544 | 1 | 4.82e-10 | | MYQLKSVPSSSLVHKSSLVH | GQHLDCGASR |
| AD119s0044.1 | 1 | 7.16e-10 | | MYQTTKYSPSSNLMPQRSAS | QAGEPWASAS |
| U1536G0490 | 1 | 4.52e-9 | | MTYHAKKFSVPFAPQRAQSS | EHASNIGAFS |

### PSR Motif 3

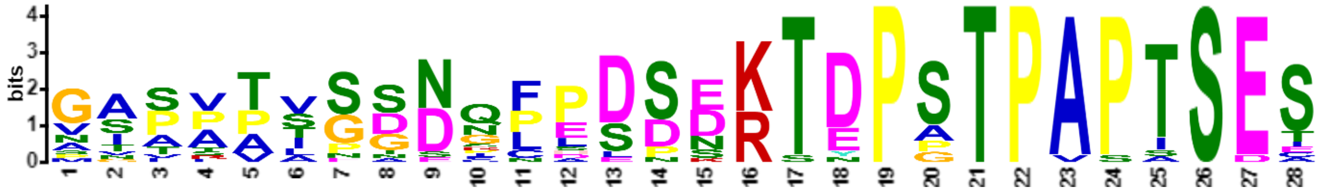

| Name | Start | *p*-value | Sites | | |
|---|---|---|---|---|---|
| W9g12770.1 | 167 | 2.93e-28 | GGVKSETPAA | GASVTLPSDQFPDSERTDPSTPAPTSES | PTQGVPSNRD |
| T2G168900.1 | 190 | 6.38e-28 | TGVKSETPAA | GASVTVSSDQFPDSERTEPSTPAPTSES | PTQVGASNRD |
| S02G161800.1 | 189 | 6.38e-28 | TGVKSETPAG | GASVTVSSDQFPDSERTEPSTPAPTSES | PTQVGASNRD |
| V4g27620.1 | 167 | 8.63e-28 | GGIKSETPGA | GGTATVSSDQFPDSERTDPSTPAPTSES | SQGVPFKRDN |
| U2891G0030 | 167 | 1.55e-27 | SGVKSETPAA | GASVTVSSDRFPDSERTDPSTPAPISES | PTQGVPSNRD |
| R2G173943 | 189 | 1.00e-25 | TGVKSETPAG | GASVTVSSDQFPDSERTEPSTPAPASES | PTQVGASNRD |
| Y714681g011 | 192 | 1.80e-25 | GGVMAETPSA | SIPAPISGNQCLDSDKTDPSTPAPTSES | PLQVKAASGD |
| X23131.1 | 164 | 2.26e-24 | VLAEGPGLSL | PSPPASGGDNLPDPDKTDPSTPAPTSES | PFQDKALLQD |
| B1005316 | 185 | 6.32e-24 | LSGFLGESNL | MIPPPISGSNFPESDKTDPSTPAPTSEC | PLQNKAAQEC |
| O129805.m001544 | 194 | 8.55e-24 | GVLGEVPGAV | AAAPVSGDNCPESDNKTDPATPAPTSES | PIQDKAAKER |
| N08G014900.1 | 194 | 8.19e-22 | GGLGEVPGAG | VSAPVSGDNFPESNSKTDPATPAPTSES | PVQNKAAKEC |
| D11g022470.1 | 165 | 1.26e-21 | GVLSEVPDSG | VTPSAAGDNGLDSDNRTDPGTPAPTSES | PHIDTSVQEH |
| G030480t1 | 253 | 1.63e-21 | GVLAEAPGSG | ASVPALGDNGLESDKKTDPATPAPTSES | PLQDKAAKER |
| AB05.2141 | 156 | 5.29e-21 | EEQQKVNGDG | NIPTPIPTNICHDSDKTYPSTPAPTSEE | KPTRDDHGLY |
| W8g25820.1 | 182 | 9.40e-21 | IEEQQRVIGA | GASRATSSEQLPDSEKTNPPTPVPISES | PVQGAPHSKN |
| X17082.1 | 246 | 2.67e-20 | KIIEEQQRLS | GVLATSEADNPSDPDKTEPSTPAPTSET | PTAGTAPSGH |
| C7G341000 | 162 | 1.71e-19 | SGTLAEPHGL | NSPAPANNNSPALDNKSDPSTPAPTSET | PLDKGSKDSA |
| D11g067280.1 | 161 | 1.99e-19 | GVLSDVPGSG | VTALPTGDNGPESDSRTDPGTPAPSSEA | PHVDKPVNAH |
| Y679311g001 | 156 | 7.53e-19 | SCVLTGASGL | GNTATTSGNPNLDSDKTDPPTPASTSDI | PIQDKGIFSF |

### PSR Motif 4

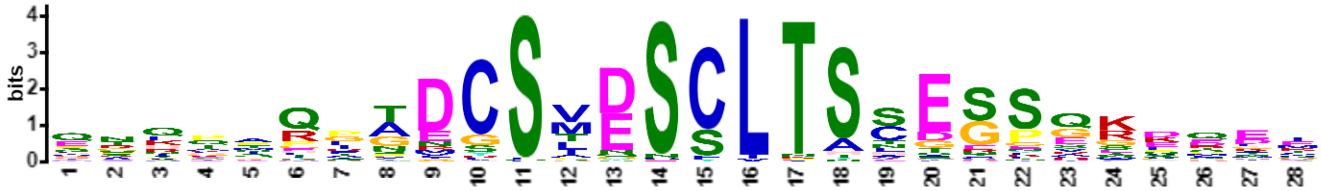

| Name | Start | *p*-value | Sites | | |
|---|---|---|---|---|---|
| B019826 | 238 | 5.28e-31 | ELQGLCLCPQ | QTQKMQPTDCSVDSCLTSCEGSQKDQEI | HNIEVGLRPY |
| B008138 | 235 | 1.09e-29 | LKELQDLHPQ | KTQTTQPTDCSMDSCLTSCEGSQKDQEI | HNIGVGLRSY |
| B108138 | 235 | 1.09e-29 | LKELQDLHPQ | KTQTTQPTDCSMDSCLTSCEGSQKDQEI | HNIGVGLRSY |
| O29950.m001149 | 240 | 3.00e-29 | LKELQGLCHQ | QTQTAPPTDCSMDSCLTSCEGSQKEQEI | HNTGMGLRPY |
| N08G149600.1 | 241 | 3.51e-28 | LKELQSLCPQ | QIQTTPPTDCSMDSCLTSCEGSQKEQEI | HNAGMGLRPY |
| H785.2 | 240 | 8.21e-28 | LKELQGLCPQ | QTQLTQPADCSIDSCLTSCEGSQKDQEI | HNNGLGLRHY |
| G026448t1 | 305 | 8.21e-28 | LKDLQGLCPQ | QTQATPPTDCSMDSCLTSCEGSQKEQEI | HNNGMCLRPY |
| AB07.828 | 332 | 1.42e-27 | VEEISSLHTL | QPHQAQLADCSIDSCLTSCEGSQKDQEI | HNISIGLRAF |
| M13G060200.1 | 247 | 1.87e-27 | LNDLQGLCPQ | QTPPTQPNDCSMDSCLTSCEGSQKEQEI | HNIGMGLRPC |
| Y6550973g003 | 209 | 6.88e-27 | LEEISGLHPL | QAHAAQFTDCSVDSCLTSCEGSQKEQET | NYVGVGLSTY |
| N09G142300.1 | 240 | 8.86e-27 | LKELQGLCPQ | QTQTTPPTDCSIDSCLTSCEGSQKEQET | HNTGMGLRPY |

| | | | | | |
|---|---|---|---|---|---|
| K12G089100.1 | 247 | 1.62e-24 | IQGFSPHQQT | QTNQPNTNDCSMDSCLTSCEGSQKEQEI | QNGGMSLRPF |
| U0468G0380 | 389 | 8.03e-24 | FMEIEGSPML | QGHTMQLGDGSVDSCLTACEGSQKDQDI | LSISLSAKKG |
| U2430G0200 | 242 | 9.74e-24 | FEEIEGSPML | QGHTMQLGDGSVDSCLTACESSQKDQDI | LSISLSAKKG |
| A119.73 | 235 | 4.39e-23 | LTEAPSLNNQ | QAQKSHLADCSMDSCLTSCEGPQKDQET | QNISIGLGYH |
| K12G184700.1 | 240 | 5.27e-23 | PKDLQGFFPQ | QTQTNPPNDCSMDSCLTSSDRSQKEQEI | QNGLRHFNSH |
| T9G419000.1 | 242 | 1.29e-22 | FEEMEGSQMA | QGHTMQLGDGSVDSCLTACDGSQKDQDI | LSISLSAHRG |
| M19G032700.2 | 235 | 1.54e-22 | LKDLQGLCPP | LTQPTHPNDCSMDSCLTSIEGSQKEQEI | HNTGMGLRPY |
| X01685.1 | 248 | 3.09e-22 | LHNAFTFEDV | HRHAAQFNDGSVDSCLTSCEGSQKEQEL | HTKCKRTAAE |
| S01G386700.1 | 242 | 3.09e-22 | FEEIEGSQIL | QGHTIQLGDGSVDSCLTACDGSQKDQDI | LSISLSAHRG |
| R2G009060 | 247 | 3.09e-22 | FEEIDGSQIL | QGHTIQLGDGSVDSCLTACDGSQKDQDI | LSISLSAHRG |
| Z5P07670_001 | 181 | 1.00e-21 | LEEIRNPNTL | QVNPSQLADCSAESCLTSSEGSQKDQDM | TNFHRSLRAY |
| K13G316600.1 | 240 | 1.18e-21 | LKDLQGFCPQ | QPQTNPPNDCSMDSCITSCDRSQKEQEI | QNGLRHFSSH |
| W3g20900.1 | 241 | 1.64e-21 | GFEIESSQML | QGHTMHLGDGSVDSCLTACDGSQKDQDI | LSISLSAQKG |
| Y771701g004 | 239 | 1.93e-21 | LEEISGFHTL | QAHAAQFADCSVNSYLTSCEGSQKEQEK | YNVSMGTYLG |
| L2g027800.2 | 224 | 1.72e-20 | GVNVPEEEVP | QTIPPQRADCSTESCLTSHESSGGLTLE | GSQVGGKRRM |
| AA17183 | 237 | 2.69e-20 | LGEIHGADNL | RECRVQHADCSIDSCLTSCEGSQKEQKM | HGINLNLQGY |
| G011693t1 | 217 | 7.47e-20 | TEGGGSSLKI | ERKPMRGTICSMESSLTSSESSGRKDEE | PPKNENISTP |
| K11G183400.1 | 256 | 4.59e-19 | PHHQKQTQTN | NNQPINANDCSMDSCLTSCEGSSQKDQQ | EIQNRGMNLI |
| J3828237 | 238 | 4.59e-19 | FLEQKEFRNL | CTQQMQPPDCSLESCLTSSEGAQKNPKT | LENNRLGLRT |
| N05G101800.1 | 230 | 9.00e-19 | EIGGSNLKDT | ERKLMRGTVCSVESSLTSSESSGRKEDM | KQKNQLGNTN |
| N01G035100.1 | 230 | 1.17e-18 | EIEGSSLKGT | ERKKMRGTVCSMESSLTSSESSGRKEDK | QLTNEFGNTH |
| M116G001200.1 | 239 | 1.17e-18 | AEVHGPEDVL | PGLHHQGADCSTESCLTSNESPGGLNLE | GSPAGGKKGM |
| L4g081710.1 | 211 | 1.34e-18 | ETRGFSLNFG | ERKQNRGTMCSLESSLTSSESSERKEEK | QTINEAENTP |
| Y785961g002 | 232 | 1.52e-18 | LKESSSLHTL | QGQTTQFSDYSGDSCLTSCEEFQKDRET | HNVCVGLGTC |
| EL01782.1 | 207 | 1.74e-18 | MKELSDTSLQ | QRQKAQAADCSMDSCLNSFEGSVMDQEL | HNNLGLKPFN |
| M08G087600.1 | 219 | 2.25e-18 | ETGGSILKDI | ERTQMRNTVCSMESSLTSSESSGRKEDM | QKENEIHDTN |
| F19971001 | 203 | 2.25e-18 | ETGGSGLKDK | ERKPMRGTGCSLESSLTSSESSGRKEEK | QPKNENGNTN |
| K09G017300.1 | 239 | 4.27e-18 | GVNVPEEEIL | PSLPPQRADCSTESCLTSHESSGGLALE | GSPGEGKRRM |
| T2G436900.1 | 234 | 4.85e-18 | TKCAQHEHFH | HQHLGGVGDGSVDSCLTACEGSQRDHDM | LSIGLSPAPT |
| G030479t1 | 240 | 4.85e-18 | NACGQDEEMP | SGLHSQRADCSTESCLTSHESPGGLTME | GSAVEGKKKM |
| PHL9_AT3G04030 | 239 | 7.07e-18 | PKELQNLCSQ | QMQTNYPPDCSLESCLTSSEGTQKNSKM | LENNRLGLRT |
| D04g015290.2 | 240 | 7.07e-18 | VGVHGPEDVS | PRIHPQRTDCSTESCLTSHESPAGLPVE | GSSPGGKKRG |
| H127.186 | 314 | 8.01e-18 | NANGQEEEVP | AGIHQQRTDCSTESCLTSHESPRGLTME | GPLVGGNKRI |
| N15G053300.1 | 245 | 9.07e-18 | LAAALENKST | ANLPARIGDCSVESCLTSTGSPVSPMGV | GSQAAASIKK |
| N03G148100.1 | 244 | 9.07e-18 | LAAALENKST | ANLPARIGDCSVESCLTSTGSPVSPMGV | GSQAAGAGAA |
| K02G108500.1 | 214 | 9.07e-18 | ETRGLSLSCG | ERKRDRGTMCSLESSLTSSESSGRKEEK | QPMEEIVEFK |
| K01G049100.1 | 209 | 9.07e-18 | ETRGLSLSCG | ERKRDRGTMCSLESSLTSSESSGRKEEK | QPMEEIVEFK |
| O29904.m002952 | 237 | 1.03e-17 | LAVALESKST | SNLPARIGDCSVESCLTSTGSPVSPMGV | GSHTASIKKR |
| AH181986g0010 | 249 | 1.16e-17 | QQHMVQAVHS | LNQQSQVTDYSMESCLTSNESSEKNLDR | NMQNAGKKRL |
| K15G123000.1 | 238 | 1.48e-17 | GVNVPEEEIP | LSLPPQRADCSTESCLTSHESSGGLALE | GSPGEGKRRM |
| D10g083340.1 | 231 | 1.48e-17 | KELSGFHTPQ | TQATQRLADCSMDSCLTSSEGPLRDLQE | MHNNQLGLRN |
| B022348 | 255 | 1.48e-17 | IAAALEEKSA | SNGPTRIGDCSVDSCLTSTGSPVSPMCM | GSQAAMMKKR |
| O29805.m001543 | 240 | 1.89e-17 | AEARGLEEVP | SSLHQQGADCSTESCLTSNESPGGLNLE | GSPAGGKKQM |
| Z2P01180_001 | 240 | 2.13e-17 | LEEIPAPNTL | QVHPPQLADFSVQSCLTSSQGSQKDLDM | ANFRRSLRAY |
| C2G398700 | 229 | 3.06e-17 | AAGCLDEKFL | SNVQARIGDCSVDSCLTSSGSPLSPMGV | GMQSATIKKR |
| AA00348 | 227 | 3.45e-17 | MKEIPDPHNL | KVNPIQFADCSINSCLTTSEGSKMDRAM | HCTNMILRAY |
| EM01478.1 | 234 | 3.45e-17 | LGVPPPLEMP | PNLRQQLADCSTESCLTSHGSPVGLPAE | GSSPGGKRAI |
| D10g085620.1 | 222 | 3.45e-17 | IKELSGFHSQ | QTQATQPTDRSIDSCLTSRDGSLRDNTM | HDNQIGLRPF |
| B000838 | 214 | 4.37e-17 | EAGGFSLQEV | GKQSMRGMDCSMDSSLTSSDSSGRKDEN | SQKLESGHPH |
| AH21538g0020 | 532 | 6.99e-17 | IPGLLPESQI | ARQQSQAADCSAYSCLTSNESSEKSPTE | NLQAGGRKRP |
| S02G422000.1 | 236 | 7.85e-17 | QQDHQHQHQH | QHQRHLGGDGSVDSCLTACEGSQCQRER | ERDQDLLSIG |
| Y850271g001 | 395 | 8.82e-17 | AAACMEEKSL | NRAMAQIAECSVDSCLTSTESPGRAPVL | GSQAAALRKG |
| M01G314800.1 | 241 | 8.82e-17 | LAAALENKHA | SNVPARVGDCSVESCLTSTGSPVSPMGV | GAQVASTKKR |
| K02G070900.1 | 240 | 1.11e-16 | LKELQVLWPQ | QTQEGQATDCSMGSFLTYSEESQRDRET | HNMNLNLRAY |
| M17G054800.1 | 241 | 1.25e-16 | LAAALGNRNA | SNVPARIGDCSVESCLTSTSSPVSPMGV | GSQVASTKKR |
| R155434.2 | 250 | 2.75e-16 | RRGNVQQEHL | HQQHHLGGDGSVDSCLTACEGSRCQRER | DQDLLSIGLS |
| AB09.1226 | 192 | 4.79e-16 | LKMPSLAEMT | VACLPQITECSVDSCLTSTESPAKVSAL | KKRPHPLLGT |
| K20G193600.1 | 319 | 5.96e-16 | VGGMHASVVS | PILQPQGADCFTESCLTSLESLGGLTLE | GSPGGSKKRM |
| U1153G0050 | 410 | 1.03e-15 | ASAKRLQNER | SIIYQQRGDGSIDCCLTSCEGSQRGDHG | ILSIGLSAAN |
| B0105317 | 233 | 1.03e-15 | IRVQVPEEEP | PNLQPPRADCSTESCLTTNGSPAGLSLE | SSPAGCKKRV |
| K20G035300.1 | 214 | 1.14e-15 | ETRGLSLNCG | ERKQDRGTMCSLESSLTSSESSEQQHIM | DEAENPQKFD |
| H128.40 | 201 | 1.14e-15 | EVGDSIMKDV | NQKGTRGTICSMESSLTSSESSGGNDNG | GPSKVNTCLE |
| L1g090670.1 | 222 | 2.40e-15 | GNGMHISEVP | YILQPQGANCSTESCLTSLESLRGLTLE | GSPSGTKRML |

| | | | | | |
|---|---|---|---|---|---|
| K07G229800.1 | 214 | 2.40e-15 | ETRGLSLNCG | ERKQDRGTMCSLESSLTSSESSEQHTMD | EAENPQKFNG |
| AA10677 | 207 | 3.28e-15 | LEKTYCSHNQ | EGHMAQINNFSMYSCLTSCQESQNGQEM | HDTNMVLTAY |
| K16G152200.1 | 240 | 6.08e-15 | LKELQVLWPQ | QTQEGQATDCSMGSFLNYSEESQRDRET | HSMNLNLRAC |
| A089.17 | 231 | 1.36e-14 | MASVCVDDKK | LNRQAQMADCSADSCATSNESSAGAPLQ | SGGKKRLRPM |
| V1g63690.1 | 239 | 2.46e-14 | TGFEEIQGSQ | MLQTMQLGDGSVDSCLTACESQRDQDIL | SISLSAKKGK |
| PHL10_AT5G18240 | 239 | 2.46e-14 | PKELQNLHHQ | QMQKTYPPNSSLDSCLTSSEGTQKAPKM | LDNRLGLRTY |
| C1G443600 | 220 | 2.46e-14 | ISQLVSKIST | ECLNPGPPDCSLDSCLTSSCEGSQKDQE | MPCITLTPYH |
| W7g48596.1 | 232 | 3.30e-14 | LQNEHTQLHH | HQQQQQVGDGSVDSCLTACDCEGSHHSR | SHGHRGEQDI |
| O29813.m001501 | 227 | 3.63e-14 | NSILNDTTDK | NQMIRRGTVCSAESSLTSSESSGRKEDD | MQQKNEIGAT |
| L5g041350.1 | 209 | 4.40e-14 | SRGLSLNYEE | RKHENGTSLCSLESSLTSSESSERKEEK | HSLEDIRDFK |
| EJ01031.1 | 249 | 5.34e-14 | KELSDLCMSQ | QKQANKPTDCSIDSCLTSSFEGSIRDQE | VVLYNNLMGL |
| Z9P01710_001 | 262 | 8.57e-14 | AMESVQRKPS | NGVSATFAECSVDSCLTSTASPGTGRSK | MLGPREAAVD |
| K10G196600.1 | 313 | 1.81e-13 | VGGMHVSEVP | PILQPQGAECSSESCLKSLESLGGLTLE | GSPGGSKKRM |
| L6g032990.1 | 237 | 1.98e-13 | LKEMNVLWAQ | QTQEGETIDYSMGSFLTNSEDSQRDQEI | HNKSMNFRAY |
| X02890.1 | 195 | 2.61e-13 | LNSPPPQPCL | VFNISQQSDCSIDSSLTSSEKPDKATEK | SRRIGSSSSS |
| Z9P05030_001 | 238 | 3.12e-13 | VEEIPVLHMS | QLTPAHFADCSVDSCLTSDRSQKEQDTH | DTTIGIRACH |
| C3G342000 | 213 | 5.85e-13 | KTGCFVLDNE | GKKLVRTTDCSLDSSLTSFESYGEIRGS | NFHWRSIQVI |
| X01094.1 | 147 | 6.40e-13 | PSSLLSQTIF | NPNRAQQGDCSTDSCVTSEEFERENKMG | CKIKSSLSKG |
| A061.73 | 227 | 1.08e-12 | SETAGVQNNL | EDPRAQIGDCSVDSCLTSQSEYSNGGKK | RSRPPCFEAN |
| PHL8_AT1G69580 | 238 | 1.41e-12 | EEGFLWYKKP | ENRGISQLRCSVESSLTSSETSETKLDT | DNNLNKSIEL |
| U0261G1090 | 292 | 1.53e-12 | TKESTSMPRL | ETRQIQFVESSTNNCLTSAEGFIKEHRL | QNHGMLKAYV |
| J3852594 | 224 | 1.67e-12 | EEGFLWCKKP | ENREKRQPRSSVESSLTSSESSETKLNN | NNEERKSMEL |
| P2G0037400 | 191 | 2.35e-12 | GIAAGSFFGG | GPAPSPPADCSSSSCLTTLERPEEVDEW | LLTNEPPTMG |
| J3845853 | 237 | 2.35e-12 | LELQNLHHQQ | MQTAYPPQNSSLESCLTSSEGNQKAPKM | LENRLGLRTY |
| J3831359 | 229 | 2.56e-12 | EEGFLWCKKQ | ENRGTTHPRRSVESSLTSSESSETNLNK | NNDERMSVEL |
| AF035.g025591 | 184 | 3.03e-12 | DLPLGALVDR | RILCPEVTECSPDSCLTHSTSERSETIG | SKDYYQSVDT |
| J3830924 | 235 | 4.24e-12 | RESFLYRKKT | ENRGIKLLRCSVESFLESSESSETKRNN | DNDERISVEL |
| D05g007890.2 | 243 | 4.61e-12 | IDCSISKDIE | NKTSKEGILCSIESSLTSSESSARKEGQ | NNSMNNKNTT |
| AB03.7 | 187 | 6.97e-12 | GAECPSLLFS | VPRQFLNAECSTESCLTCSEREEELKND | GKRSNAETSE |
| J3831963 | 240 | 1.34e-11 | PSLSELAVAV | DTKNNITTNCSVESSLTSNTNGSSVSAA | SMKKRLRGDD |
| EL01279.1 | 208 | 2.16e-11 | ESGGSLLKNE | GNKPLGHNGYSLESSLTSSDSSAMKEET | QTTQQLAKNY |
| AF027.g023545 | 263 | 2.53e-11 | MGDGTFEDRG | IGHPPERSECSPQSCLTHLASNERSDTN | GSNDEACLNY |
| Mpo198s0044 | 259 | 2.53e-11 | CGGVPHQLSS | PNPQSRLSDCSSQSYLTSLACSGNPDGN | EPSDQKQNTG |
| PHL2_AT3G24120 | 244 | 4.37e-11 | PSLSELAVAI | DNKNNITTNCSVESSLTSITHGSSISAA | SMKKRQRGDN |
| AA07132 | 188 | 4.72e-11 | LSSSLNHGGL | ISKGNQNADSSMDSCLTSELFELREKYA | GQHKLEASSC |
| V2g21880.1 | 236 | 4.72e-11 | IKESSRMHRL | EPRQIQFVESSTNNCLTAAEGYINEHRL | HSHGVLKAYD |
| J3860014 | 244 | 1.09e-10 | LSELEVAAID | HKSNITTTNCSVESSLTSNTNGSSVSAA | SMKKRHRGGD |
| EE00086.1 | 206 | 1.27e-10 | TQSDGSVLEK | DERDKLLVHDSLESSLTSSESSWKKNKE | TRRKHRNGKI |
| AA34364 | 222 | 1.47e-10 | NARVFAEDKP | WAGLEQITEHSVDSCLTSAGSPSAFCSK | LSLKKRLHPM |
| W5g40960.1 | 239 | 2.13e-10 | IRESSSIHRL | EPRQIQFVESSANNCLTAAEGFKEHRLQ | NHGVLKAYDD |
| Q13g00280.1 | 228 | 2.29e-10 | QPSMVDLQKL | KTQTPAASECQLDSCLTSCDHKDVSHMS | NVKMSLKICD |
| J3836250 | 209 | 2.47e-10 | EYDFLCTKKP | ENRGNESTRSSVDCSLASSESSEAKLDH | RSQTIMRRSD |
| AG013.g005903 | 225 | 8.32e-10 | GSPCPGEDWR | IQPPHERAECSPQSCLTHLTAIDRSDAT | GNGESYPCNK |
| U0008G1310 | 203 | 2.86e-9 | ELVSEVETEC | LSSPSPPRHRSTDSCVTSSSSSEAESKA | AGSKRLYTSM |
| Q289g00260.1 | 151 | 3.27e-9 | YLQTVLKKAE | ESLAAGTDSVLIDSCLTSLERKEEDNDD | DDGEVEEDNL |
| AF481.g073115 | 279 | 4.26e-9 | VSNSVSERRV | PENDQDIAECSPKSCLTHVTTNERSEHG | CSRDIFQNLT |
| D09g091880.2 | 237 | 6.32e-9 | IGPSFENKNA | CNIPAMLGDCLLDDCLPSNGISSLKKRP | RGFTNVNGLP |
| AF121.g046895 | 218 | 9.32e-9 | DDFSKCGAMD | ECESHIKTECSFDSSITNVISKERYETE | ENGGLEKKYR |
| PHL3_AT4G13640 | 236 | 1.99e-8 | KMMIPSLSEL | AVAIEHKNNCSAESSLTSSTVGSPVSAA | LMKKRQRGVF |
| AH8183372g0010 | 104 | 1.97e-6 | KQSCKENNSD | TNKDGGNSDCSKDPNIKSRDNNTRTNQK | EAMHITEALN |

PSR Motif 5

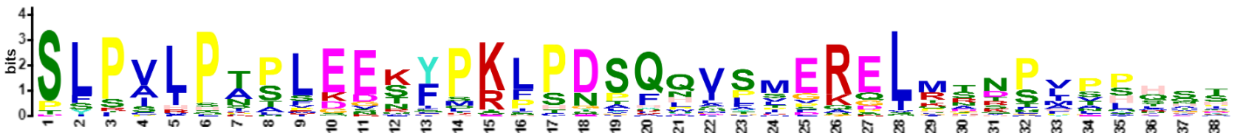

| Name | Start | *p*-value | Sites | | |
|---|---|---|---|---|---|
| B012506 | 4 | 1.33e-37 | MSS | SLPVLPTPLEEKYPKLPDSQQVSLEREIMANSVPPHAT | QLASNSGVVG |
| B010973 | 4 | 1.86e-33 | MSS | SLPVLPTPLEEKYPKLPDSQEVSLGREIMTNPVVPQVP | PLVSNSEVVG |
| AB10.1900 | 4 | 1.23e-32 | MSS | SLPVLPITSGENYPKLPDSQQVSMERELRSNPVNPHKT | TFVPSNGAVG |
| Y803301g004 | 26 | 3.97e-32 | STGNSEVMSS | SLPVLPAPFEEKFPKLPDSQQLSVERELRSSPIIPHHT | PFVSNSGVVG |
| N09G147400.1 | 4 | 2.06e-31 | MSS | SFPVLPTPLEDKYPKLPDSFQIASERELMRDPVNQQSS | PLGPNSGTIK |
| T2G118400.1 | 21 | 3.82e-31 | SSGVPVTVPT | SLPSIPASLEESFPRLPDSQNVLIERELRSTPVPPHQN | TVAPIRGQFH |
| M02G257800.1 | 48 | 1.84e-30 | NLGVSGTLSS | SLPVVPTPLEETYSKLPGCQQVSMERELMTRPLVHASH | LPSNNGVVGH |
| O29680.m001711 | 4 | 5.96e-30 | MSS | PFPVLPTPLEGQYPKLPDSFQVSSERELMRNPILQQTS | PLCSNSGTVG |
| N08G140300.1 | 4 | 9.46e-30 | MSS | SLPVLPTPLEDKYQKLPDSFQVASERQLIRNPVPLQAS | QLGPDSGTTA |
| R2G162409 | 21 | 5.17e-29 | SSGVPVTVPT | SLPCIPVSLDESFPRLPDAQSVLMERELRSTPLPPHQT | TVAPIRGLFH |
| S02G121600.1 | 21 | 1.56e-28 | KSGVPVTIPT | SLPSIPASLDESFPRLPDTQSVLMERELRSTPLPTHQT | TVAPIRGHFH |
| O29637.m000753 | 29 | 5.66e-28 | NLGVSGALSS | SYPVLPTSREETYPKLSDAQQVAMEKGLMTRPLVHTNH | LPSNNGVVGH |
| Z4P30200_001 | 4 | 1.62e-27 | MSS | SLPVLPNTLEEKFPELPDPQHVPLEREIQRNPLPSQRP | LFSSPSGFSP |
| K19G167500.1 | 29 | 2.00e-27 | NMGMSEAFPS | SLPALPSPLEETYPKLSDSKPVFMEKELKTKPYTHSSH | LTSSGAVGHM |
| V1g28920.4 | 21 | 2.46e-27 | SPGIPVTVPA | PLPSIPASLDESFPRHPDVQNVLMERELRRTPLPPHQS | TVAPISGQFH |
| G026171t1 | 4 | 3.72e-27 | MSS | SFPALPTPFKEKYPKLPDSFQVSSERKVMKNSISPQES | SLAPSNRTLG |
| Z11P02560_001 | 4 | 5.59e-27 | MSS | SLPVLPTPSEEKFPKLTDLQQVSVEREIRNIALASHHT | PFISDGGIVG |
| C2G381900 | 28 | 6.19e-27 | NVGVSGCMSS | SLPVLPLPLEEKYTKFSYSQQICFERELMSDPVAPFDT | PLASTGENVG |
| K03G166400.1 | 26 | 1.26e-26 | DMGMSEVFPS | SLPVLPSPLEETFPKLSDSRPAFMEKELKTKPFTHSSH | LTSSGAVGHM |
| G003939t1 | 29 | 2.53e-26 | NLGVSGGLSS | SLPALTTPLEETYQKLSDTQQVSADRELMTRPLVHATC | VPTNSGVVGH |
| W7g25710.1 | 21 | 2.85e-25 | NSGIPVTVPS | PLPAIPATLDENIPRIPDGQNVPRERELRSTPMPPHQN | QSTVAPLHGH |
| N12G035600.1 | 29 | 4.16e-25 | NHGLSAALSS | SLPVLPTSIEETCPKLSNSQQVSLEGGLMTKPIAHASQ | LPSNSGVVGH |
| V1g63530.1 | 21 | 1.68e-24 | NNRVSGAMSS | SLPILPNSLKENFPRSHNPQLIPMPRQLMNDSVPLHHS | APQSATLHPR |
| Z2P10180_001 | 29 | 4.18e-24 | DARSSGVMSS | TLPVRPNNSGEKFPKIPDSQLVLMDREIRSNPLPSDYT | PFVSDGGNVG |
| L7g098250.1 | 29 | 6.56e-24 | NIGMSGALPS | SLSVHPTPLEETYPRFSDSQPTYVEKDLKTKTFNHSSH | ISSSGAVGHM |
| U1459G0190 | 4 | 1.12e-23 | MPS | SLPILPKPLKENFPRPHNPQHMPMPRQLQNDSMPLHQS | SPQSATLHPR |
| X14587.1 | 21 | 1.46e-23 | DIGGSGAMSS | SLPALPTPSEVSFLKLPDSQQVSMERELSSSNPLPPHR | APSHYNTAVV |
| PHL1_AT5G29000 | 16 | 3.22e-23 | DFGYSTAMSS | SYSALHTSVEDRYHKLPNSFWVSSGQELMNNPVPCQSV | SGGNSGGYLF |
| S01G384300.1 | 15 | 4.97e-23 | KSRVLGAMSS | SLPILPNPLKGSFPKPCNPQHIPMSRQLPDDSMPLRND | IHQSASLHPR |
| T9G416300.1 | 21 | 8.34e-23 | NSRVLGAMSS | SLPILPNSLKESFPRPCTPQHIPMSRQLPDDSMPLHNG | TPQSDTLQPR |
| K03G250000.1 | 30 | 1.79e-22 | HLGASRAMST | SSRVLPTPLENKYMKPPDSFQLSPVRDLTANSASSNSI | RSAGKMFSSP |
| C5G018400 | 29 | 1.79e-22 | DSGISVAMSS | SLPVHPIALEKQYSQFPNSQQVSEERGLLTNSLVSHST | LLATNSGSVG |
| K19G247600.1 | 23 | 5.33e-22 | HLGASRAMST | SSRALPTPLENKYMKPPDSFQLSPVRDLTANSASSNSI | RSAGKMLSSP |
| D09g072830.2 | 29 | 6.30e-22 | NCGASGALSS | SLSIFPTALGEKYPKFPDLQQASMGKELKQHPATVVSS | LPSNSGAVGL |
| K10G039700.1 | 27 | 1.03e-21 | NMGMSGALSS | SLSILPIPPEELFPKLPESQLDFVEQELMIRPFTHSSY | LNSGGVIGHI |
| ED00466.1 | 29 | 6.60e-21 | SFGASAALSP | SFPARPAALQESSPKFSDSQYISVERELMHRQPAPLSS | NNGVVGHIFS |
| R2G006477 | 21 | 1.34e-20 | KSRVLGAMSS | SLPILPNPLKGSFSRPHNPQHIPMLRQLPDDSMPLCID | THQSASLHPR |
| L7g115530.1 | 5 | 3.42e-20 | MPSS | SSPVLPSLMESNYMKPPDSFNCSPVRSLTANPASLQAT | SSYPIRSVRP |
| J3837308 | 16 | 9.90e-19 | DFGYSTAMSS | SFSPLHATVEDRYRKFPNSYWGSSGQELMNNPVPYQVV | SSSGGYMSGY |
| A142.28 | 30 | 3.35e-18 | SLGMPQGPSS | SLPCLSAPLEDRQPKFPSTQQAGMGKELRPSTVGPLTG | PVASNNGVVG |
| U1079G0490 | 21 | 1.03e-17 | SSGVPVTVPS | PLPAIPAALEESFPRLPDAHNQTTFAPLHGQFHSSTGS | VGPLSSPPAI |
| U5938G0020 | 21 | 2.90e-17 | SSGVPVTVPS | PLPAIPASLDESIPRLPDSHNQTTFAPLHGQFHSSTGS | VGPLSSPPAI |
| D06g008200.2 | 28 | 3.18e-15 | SNFGASGAHS | SLPVLQTSLEEKYPKLPGSALSSFPSNSGAVGHLFSSP | SEFSTNLDFS |
| Q9g01070.1 | 11 | 1.17e-13 | MEASNKSNRC | SYFSHSRTKSSSLPTNPNSQLVSIERELQSNTAEPSFA | AHLLSNSSTM |
| D05g055940.2 | 28 | 3.89e-13 | SNMGVCGAMS | TSLSILPSFEEKHLKVTDSLHVLTEKEQTTNLISSRAI | PSASKSGTVG |
| V3g55120.1 | 26 | 8.51e-13 | DGEVPWYLSS | DITVLPALPEVMVPSFPQYSASYIERGLCRSSITPLNT | FLPASSGLQS |

PSR Motif 6

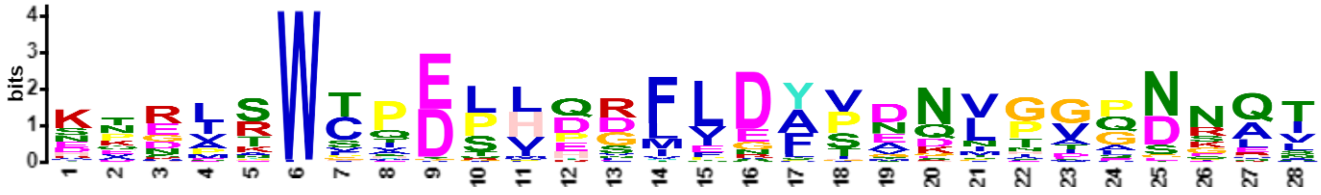

| Name | Start | *p*-value | Sites | | |
|---|---|---|---|---|---|
| NS107600.1 | 87 | 1.85e-24 | QFTTSSQYAK | ETNASWCPESLPDFLDYPVNNPVQNNQI | EGNSCGAIVS |
| G021688t1 | 12 | 1.17e-22 | RRPSRSDGLA | KERLRWTQELHDRFEDAVNQLGGPDRAT | PKGILKAMGV |
| Y681141g002 | 87 | 1.86e-22 | CPATSNHPRE | SADVSWCPEPIQGMLDYSDNITAGNNQI | QSSCDVASDD |
| L5g041350.1 | 15 | 1.12e-21 | SMHFVLSTDA | KPRLKWTPELHQRFIDAINQLGGADKAT | PKSIMRVMEI |
| J3836250 | 13 | 1.12e-21 | NDKNHKTEEP | KPRLRWSYELHHRFIDAVNQLGGPNKAT | PKGLMRVLEI |
| M06G101000.1 | 11 | 2.65e-21 | MGSSRSDVSN | KERLRWTQELHDRFEEAVNQLGGPDRAT | PKGILRAMGI |
| PHL11_AT5G45580 | 7 | 4.04e-21 | MMTRDP | KPRLRWTADLHDRFVDAVAKLGGADKAT | PKSVLKLMGL |
| AF020.g015442 | 11 | 4.65e-21 | MASSTSSSSS | KQRLRWTPELHQRFQDAVAELGGPDRAT | PKGILKVMGV |
| M16G117000.1 | 11 | 5.34e-21 | MGRSRSDRSN | KDRLRWTQELHDRFEEAVNQLGGPDRAT | PKGILRAMGI |
| O29637.m000753 | 138 | 1.06e-20 | QCTASSQYDK | ENSASWCPESLPGFLDFPVNSHVQNNQI | ESNSCTGAIT |
| N12G035600.1 | 138 | 1.06e-20 | QSTKSSQYAK | ETNASWCPESMPGFLDLPVNNPVQNNQV | ESNSCSGVIV |
| N09G105800.1 | 11 | 1.22e-20 | MGSSRTDGSG | KERLRWTQELHDRFERAVNQLGGPDRAT | PKGILKAMSI |
| F19971001 | 10 | 1.40e-20 | MSLVLSTDA | KPRLKWTPELHHRFVEAVAHLGGPDKAT | PKTLMRVMGV |
| EH101641.1 | 13 | 1.40e-20 | SGGNNSNLAS | RQRLRWTHELHERFVDAVAQLGGPDRAT | PKGVLRVMGV |
| O29680.m001711 | 116 | 1.83e-20 | STALINHSEE | NKDMSWTIDPLHDLLDFPENVAVQNGQV | ESTIGVITSE |
| F34259001 | 13 | 2.09e-20 | NQVPEAAAAQ | KQRIRWTPELHELFLDAVSKLGGPDKAT | PKGILRLMNV |
| B012506 | 114 | 2.39e-20 | SRPMSHYPRE | NNNISWSADQLQSFLDFPENVPTQNNQV | ESSNSGIMPS |
| Y803301g004 | 135 | 4.62e-20 | QSSTVNHPRE | STKVTWCPEPIQSMIDYSDNITDGNNEI | QNSCDVVSDD |
| M02G257800.1 | 157 | 4.62e-20 | QSTTTSHYVN | ENSASWCPESPPGFLDFPTNTTVQNNQI | ESNSCAGVMA |
| J3832766 | 15 | 4.62e-20 | NGGTNSSHAS | KQRLRWTHELHERFVDSVAQLGGPDRAT | PKGVLRVMGV |
| U1079G0490 | 117 | 5.26e-20 | LEPSLTNFPR | DVGPTWCPDPVESMLGYSDDVPGGNNLT | ESSSIAATDE |
| H34762 | 11 | 5.26e-20 | MGSSRSRGTS | KERLHWTQDLHDRFEEAVNQIGGPDRAT | PKGILNAMGI |
| AB10.1900 | 102 | 1.00e-19 | QSSLSNHSRE | SNEINWCQDTLQGMLDYSEEVTVENNQI | QSSTIMPSDN |
| W7g25710.1 | 134 | 1.29e-19 | FEPSLTDFPR | DAGPTWCPDPVDGLLGYTDDVPAGNNLT | ENSSIAAGDE |
| K01G049100.1 | 15 | 1.67e-19 | SMHFVLSTDS | KPRLKWTPELHRRFIEATNQLGGADKAT | PKSLMRVMGI |
| T2G118400.1 | 132 | 4.50e-19 | FEPSLTDFPR | DVGPTWCPDPVDSILGYSGDVPGGNNLT | GSTSLGASDD |
| EN00196.1 | 15 | 5.08e-19 | GGGLMMTRDS | KPRLRWTSDLHDRFVDAVTKLGGPDSEL | KSVLRLMGLK |
| G026171t1 | 112 | 6.48e-19 | STALFNHPQE | KKDVSWCIDRLQDFLDLPENVPDPNGLL | ESSTGVMASE |
| N09G147400.1 | 116 | 1.05e-18 | SIALINHSEE | HKDMSWSMDPLHDLLDFPENVTVENGQV | ESNIGVITSE |
| N08G140300.1 | 110 | 1.69e-18 | STTLISNSEE | NKDISWSVDPLYDLLDFPGNVTTQNGQV | ESNIGVIASE |
| M19G020900.2 | 84 | 1.69e-18 | STAFINHSDD | NKGLSWPIDPLQEFINFVENVPVQNGQV | ESTAGVIASE |
| B010973 | 114 | 2.13e-18 | SGALNHYPKE | NSNISWTADQLQGFLDFPENVPTQSSQT | ESGNGGVMPS |
| U5938G0020 | 117 | 3.40e-18 | LEPSLTNFPR | DVGPTWCLDPVESMLGYTDDVPVGNNLT | ESSSIAATDE |
| F23401001 | 65 | 5.38e-18 | STPSVTFSRE | RKENSWCTDSLKDFLDFPENVPIQNNNQ | VEGSGGGMSY |
| M13G048000.1 | 84 | 6.75e-18 | STEFIAYSDE | NKDLSWPVDPLQDLLDFAGNVHVQNGQV | ESSAGVFASE |
| ED00466.1 | 135 | 1.06e-17 | QSTASSQFNK | ENNDSWCTETLPDFLDFPMNTTILNNEL | DVSTNGGIPV |
| Z4P30200_001 | 102 | 1.18e-17 | FQVPNNFPKD | PTEITWCPDSVQGMLNCSDGVIMGNNQI | QNSSNKVSND |
| D09g072830.2 | 135 | 1.48e-17 | QSTASSQYLN | ENHEPWCIDPLPNFLDYSMDNPVQNSQV | ASSNKQSDWQ |
| C5G018400 | 140 | 2.30e-17 | PTDLGNYTKE | NNNVSWGEDSLQVLLDFPENVPVENRQL | ENSGTDIMKQ |
| U1459G0190 | 106 | 2.86e-17 | QSLSNNTPGG | HAEATWFPASVDGLPDYTDNMTAPDNQI | PSGSSNMTSD |
| X10145.1 | 29 | 5.46e-17 | QPSVSNYPKE | STELAWHPEPLQRVLDYPDNPNFGNNLS | HGVNVMASDE |
| N06G178000.1 | 264 | 1.03e-16 | LPSPGPATMH | KPRMRWTPELHECFVEAVNKLGGAEIEA | TPKGVLKLMN |
| Z11P02560_001 | 111 | 1.27e-16 | QPITTNFPRQ | STEVAWCPDAVDNILDFTDNNIGVGNQM | PSNSAMVSDD |
| V1g63530.1 | 123 | 2.13e-16 | QTLSDNIPGG | HTEATWFPGSVDGLTDYGDNVGAPGNQI | RNGCPAVTSD |
| V1g28920.4 | 132 | 2.13e-16 | FEPTFTDFPR | DVEPAWCPDPVESMLGYSDDVSGGNSLN | GMSPIAATDE |
| A142.28 | 158 | 2.90e-16 | QPVASDFTKE | SNDVSWSSDPLQGFLDFPDNVGESSQHN | GMDGDDQTRR |
| D06g008200.2 | 119 | 4.35e-16 | QGAASSQYVN | VNSESWCTELLPDYLDYSVNAPVQNTQL | DCRNSDDCQI |
| Q58g00470.1 | 222 | 6.49e-16 | PSTPTNAPSN | KQRIRWTVELHDRFVEAVNKLDGLSERA | TPKSVLKLMN |
| A010.492 | 20 | 2.55e-15 | IVPLPLTHAT | KPMLKWTPELHQLFLDTVARLGGLDEVS | PKVMVRAMGI |
| AA40820 | 28 | 2.81e-15 | QSLASNYPLE | ATGISWVPESIQDIFRYSDNMHIGSSQL | QNSPINNQIK |
| H128.40 | 10 | 3.40e-15 | MNLVLSTDA | KPRLKWTPELHHRFVEAINQLGEATPKS | LMRVMGIPGL |
| AA15381 | 104 | 1.05e-14 | QSLASNYPLE | ATGISWVPESIQDIFSYSDNMNIGSSQL | QNSPINNQIK |
| L1g080330.1 | 53 | 1.39e-13 | EIHSMTFPQE | SDVMSWGTDPFEDILQFHDNVPTQNDHV | EYNGSEVLGG |
| V3g55120.1 | 134 | 2.32e-13 | SSLRMLYPKV | SEEIYWVQEPVPGVFDYPASLNVSDQRN | LVVSQEMQDI |
| T9G416300.1 | 120 | 3.54e-13 | QSLSDNTPGA | HAEATWFPSSMDVLPVYTDNIAASDNQI | QSGSSAMTSD |
| Z8P12580_001 | 6 | 1.22e-12 | MFSRE | TTKINLSPDTLHDTLDYPDNNFSLSNQI | LGNSTMVSND |
| AE423505 | 29 | 7.89e-9 | EQGGGGNGLM | KNEVEWAQEGWTSLLNPSDGGGDGEESS | SKKGVVVPQF |

PSR Motif 7

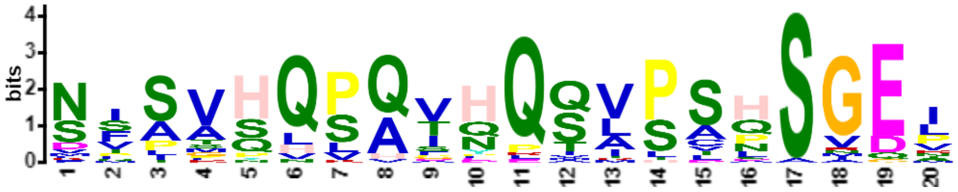


| Name | Start | *p*-value | Sites | | |
|---|---|---|---|---|---|
| Z5P12960_001 | 46 | 2.96e-20 | VVYPAAQSSQ | NMSVHQLQTHQSVPCHSGEI | SAVTGPLSTA |
| Y681141g002 | 166 | 1.28e-19 | SVQPATQASP | SISVHQLQIHQSVPSHSGEL | CAVSNSSPAA |
| Y803301g004 | 214 | 1.88e-19 | AMQSAAQASP | SISAHQPQIHQVVPSQSGEI | CTVSNSSSAA |
| Z4P30200_001 | 181 | 3.35e-16 | VVYPPAQSSS | NISKQQPQTDQSVPCHSGEV | CAVTGASSSA |
| B010973 | 204 | 3.35e-16 | AAYQVPKPPP | NFTVHQPQIHQQLPDPSGQL | CSVASPSSSX |
| O29637.m000753 | 229 | 4.06e-16 | MAYQVSKPPS | DISAHQPQVHQQLPAPSIDI | RPVLTPTSSV |
| AB10.1900 | 181 | 5.41e-16 | VVYSAAPVSS | NVSVPQSQSHQMVPSQSGEL | GAVNSPSTSA |
| N12G035600.1 | 229 | 5.95e-16 | MAYQVSKPSS | NVLSQQPQVHQQLPAPSGEI | HPVLTPTSST |
| V1g63530.1 | 198 | 7.91e-16 | SQSKAMIQTS | NSAVSQPAVNQSASSHSGEM | CNVASPPNGN |
| B012506 | 204 | 9.54e-16 | TAYQAPKPPQ | NFLVQQPQVHQQIPVLSGDL | SSVASPSSSV |
| U1459G0190 | 181 | 1.05e-15 | SQSKSMVQPS | NLAASQPAVNQSASSHSGEI | CPVASPPNSN |
| T2G118400.1 | 207 | 2.88e-15 | TQQVGPPAQS | SISVHQSAAQQTVSSQSGEP | MAVVAPSPTG |
| S02G121600.1 | 207 | 3.45e-15 | TQQVGQPVQS | SISVHQSATQQTVSSQSVEP | LAVVAPSPTA |
| U5938G0020 | 191 | 3.77e-15 | QSQVGPPAQS | SVSVHQSATQQIVLSQSGEP | SAVAAPSPTA |
| U1079G0490 | 193 | 4.51e-15 | QSQVGPPAQS | SIAVHQSVTQQTVSSQSGEP | SAVAAPSPTA |
| F28403001 | 146 | 9.95e-15 | MAYQVPKPSS | NFSANQPQVHPQLSAPSGEV | HNVVTPSSSV |
| X10145.1 | 110 | 2.15e-14 | VALPADQAPT | SFSVHHPQTHPSVPSNSNDL | SAVASPSSSA |
| W3g21240.1 | 180 | 2.55e-14 | SQSKSMAQPS | NSAASQPAFNQSTSSHSGDI | CPVTSPPPNN |
| AA40820 | 113 | 2.77e-14 | LKVFVLLSFL | VFSVSVVHVHQQVPLHSGER | SIMSSPSPSI |
| AA15381 | 189 | 2.77e-14 | LKVFVLLSFL | VFSVSVVHVHQQVPLHSGER | SIMSSPSPSI |
| R2G162409 | 207 | 3.88e-14 | TQPVGQPVQS | SNSVHQSATQQTVSSQSVEP | LAVVAPSPTA |
| X14587.1 | 215 | 4.97e-14 | GSNQVVCPPS | NLSMQQPAIPQSAPSNSGEL | CTVASPSPTA |
| Z2P10180_001 | 209 | 5.86e-14 | VVCPAAQSSP | DISMHQLQSHRSVPCHAGEV | CSITSPMSAT |
| NS107600.1 | 176 | 6.36e-14 | KMAYQVSKSS | SVPSHLPQVHQQLLAPSGEI | QPVLTPTSSA |
| T9G416300.1 | 195 | 2.91e-13 | SQSKAMMQPS | NSIASLLAVNQSASSHSGEI | CPVASPPNSS |
| L7g098250.1 | 227 | 3.97e-13 | VESISKSSSQ | FPAGHQSQDHQQLPALSGEN | HVGVAPSSSA |
| S01G384300.1 | 190 | 7.33e-13 | SHSKAMIQTS | NSATSLPAVNQSASSHSMEI | CPVASPPNSS |
| D06g008200.2 | 208 | 7.91e-13 | MQYQEQNQLS | NFPVHQVQPFQQIPTASVET | SAVVPASVET |
| C5G018400 | 223 | 9.92e-13 | PATYEPIASS | NSSLHQPQIQQQRPVDSGDS | NPAAVPTSSA |
| R2G006477 | 196 | 1.07e-12 | SHSKAMIQIS | NSATSLPAVNQSASSHSREI | CPVASPPNSS |
| V1g28920.4 | 208 | 1.80e-12 | QPQGGPPVQS | STSVHQSATEQIVTTQSVEP | CAVAAPSPSA |
| K19G167500.1 | 223 | 4.02e-12 | EPKVAKSSSQ | LPIGHQSQSHQQLPASSGEN | RVGVAPTSST |
| P11G0061900 | 62 | 1.34e-11 | YPAASHLLVQ | MTAPPQPQFYQQMPSNSGGV | SAANGSLSSG |
| M02G257800.1 | 248 | 2.31e-11 | MAYQVSKPSS | NTPVQHSQGHLQLPSLSAEI | RPVLTPTSSA |
| Y728141g003 | 434 | 2.84e-11 | VVYPAAQAST | DSSAYHSQIYESVHSHSGEQ | CAVGSLSSSA |
| AB01.763 | 157 | 1.14e-10 | KALCVADNSQ | KVPVQNKQVQQAIPSNSGQL | TVVSNPSSSC |

PSR Motif 8

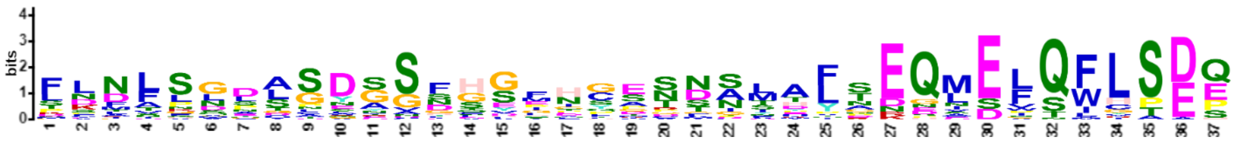

| Name | Start | *p*-value | Sites | | |
|---|---|---|---|---|---|
| H10.14 | 161 | 1.22e-32 | DDQSEALMRD | FLNIPGDTSDGSFHGMNCGNDSLALTEQMELQFLSEE | LDIAITDHGE |
| X11622.1 | 158 | 1.80e-32 | DEHSDELMKD | FLNLSGDASDGSFHGENCDSNSLAITEQMELQILSEQ | LGIAITDNEE |
| G027630t1 | 174 | 8.89e-31 | EDHSEIIMKD | FLNFPGDDCDGNFHGLHCESNNFTLTEQLELQFLSDE | LDIAIADHGE |
| M14G000700.1 | 106 | 1.06e-30 | EERSDGFMID | FLNLSGDASEGGFHGMNCTSDNLELTEQLELQFLSDE | LDIAITDHGE |
| F29458001 | 138 | 1.52e-30 | EGNSEDMMKG | FLNLSSDASDESFHVMNCASDNITFSEQLELQFLSDE | LDIAIADNGE |
| Y6550950g002 | 166 | 2.57e-30 | DEHSDDLMKD | FLNLSGDASDSSFHGENYDNNSIALGEQMELQMLSEQ | LGIAITDNGE |
| T4G270600.1 | 169 | 7.61e-29 | ALDEAEHSDD | LKDFLNLSGDASDGSFHGETNALAFDEQMEFQFLSEQ | LGIAITDNEE |
| W6g49040.1 | 171 | 6.14e-28 | HDEPEQSDDL | KDFLNLSGGDASDGSFHGENNAMAFAEQMEFQFLSEQ | LGIAITDNEE |
| U0176G0870 | 169 | 1.15e-27 | AHDEAEHSDD | FEDFLNLSGDASDGSFHGENNALAFAEQMEFQLLSEQ | LGVAVTDNEE |
| N06G178000.1 | 170 | 1.57e-27 | KETSDAFVKD | FLNLHGNASEGSFHGITSASDNLALTEQLELQFLSDE | LDIAITDHGE |
| S04G036500.1 | 163 | 1.83e-27 | GQSEAEPAAD | LKDFLNLSGDASGGSFHGESNGMAFREQMEFQFLSEQ | LGIAITDNEE |
| U0164G0530 | 152 | 3.93e-27 | DHDESEQSDD | LKDFLNLSGDASHGSFHGEDNASAFAEQMEFQFLSEQ | LGIAITDNEE |
| M07G003200.1 | 172 | 3.93e-27 | EEHSDAFMKD | FLNLSGNASEGSFHGMNYTGGNLELTEQLELQFLSDE | LEIAITDHGE |
| T1G092500.1 | 174 | 9.69e-27 | RSGGHDDAGD | LKDFLNLSGDVSEGSFHGESSAMAFSEQMEFQFLSEQ | LGIAITNNEE |
| V1g34470.1 | 167 | 1.13e-26 | AYDEAEHSDD | MKDFLNLSGDASDASYHGEDNAMAFTEQMEFQFLSEQ | LGIAITDNEE |
| B021120 | 130 | 1.13e-26 | EKRSEDLMKD | FLNFPGDASDGSFHGANFASDSLTFTEQLDLQILSAE | LDIAMTDNSE |
| O30169.m006598 | 219 | 2.36e-26 | EEHSDCLMKD | FVNFPGDASRSSFHGMTCASDNLVLADQLELQFLSDE | LDIAITDHGE |
| Z7P15740_001 | 155 | 3.65e-26 | DEHTDDLMKD | FLNLSGDASDGSTHGEIYGNNGIALSEQIELQLLSEQ | LGIAITDNGE |
| Z10P23260_001 | 163 | 2.70e-25 | DEHTDDLMKD | FLNLSGEVSDGSIHGENCGSSGLALNEQIELQILSEQ | LGIAITDNGE |
| W2g04640.1 | 166 | 2.76e-24 | RNSGHADDEH | TDDLKDFLNLSSDCSFHGKCSAMAYNEQMEFQFLSEQ | LGIAISNNEE |
| U0548G0730 | 169 | 2.25e-23 | SNGGHGDAEY | SDDLKDYLNLSGDGSFHGERNAMTFNEHMEFQFLSEQ | LVIAVTNNEE |
| J3859491 | 135 | 3.30e-23 | DGDNTLVKDF | FNLSGDACSDGAFHDLDCSNDSYCLSDQMELQFLSDE | LELAITDRSE |
| PHL6_AT3G13040 | 146 | 7.05e-23 | GNSGSLLKDF | LNLSGDACSDGDFHDFGCSNDSYCLSDQMELQFLSDE | LELAITDRAE |
| V3g03538.2 | 183 | 8.69e-20 | HGDAEHSDDL | KDFLNLSGDASGSGFHEGISNCMDFNEQMEFQFLSEQ | LGIAITDNEE |
| R2G010920 | 157 | 1.20e-19 | EAEHPAELKD | FLNLSGEASWRGGFHGESSGVAFRFGEQVEFQFLSEQ | LGIAITDHDE |
| J3849734 | 130 | 2.54e-19 | GDNNSSLVKD | FFNLSADACSNGGGYHDLDCSNDTLSDQLELQFLSDE | LELAITDRAE |
| S10G254300.1 | 174 | 1.21e-18 | NGLDEAEQSD | DLKDFLDLSGDASDGSFQENNALAYDEQMEFQFLSEQ | LGIAITDNEK |
| A025.314 | 131 | 2.47e-18 | HDTSENLIQD | LFNLSGNASDTGLCSENYPNDDMIVTEQFDWQIISDH | LDLAITDIGE |
| K19G167500.1 | 110 | 2.69e-15 | RNTHFISQSL | SNMASLPLSYSSNSEPIPSTTSTPYSNGNSVSWHTDS | LPSFLDFPAN |
| K03G166400.1 | 107 | 6.29e-15 | RNTHFISQPL | GNMASLPLSYSSNSEPIPSTTSTPYSNGNSVSWHTDS | LPSFLDFTAN |
| L7g098250.1 | 110 | 3.86e-14 | RSAHFISQSL | SNMASVPLPYSSNNGPVPSTTSTHYSNGNSASWHADP | LPSFLDFSAN |
| L7g115530.1 | 79 | 3.21e-13 | VSQHDRQYQD | SPFTSQALGDNVSSEIHSATFTSHLQENDDISWGPDP | LPDILGFPDI |
| K03G250000.1 | 99 | 6.83e-13 | VSQHDRQYQD | PPFVSQTIGDSVSSEIHSMTFISHPQENEDLSWGPDP | CQDILGFPEN |
| Y728141g003 | 321 | 1.07e-12 | VSASYISQPS | SSALSLSSTYSSYSGTCQPSMSNYPKESAEITWYSDS | LQGVLDYSDN |
| R2G162409 | 105 | 1.24e-12 | PNHNSYNSQV | PSTASSSTLNYGSQYGGFEPSITDFPRDIEPTWCPDP | VESILGYSGD |
| S02G121600.1 | 105 | 1.92e-12 | SNHNPYNSQP | PSTASSSTLNYGSQYGGFEPSITDFPRDVEPTWCPDP | VESILGYSGD |
| K13G126200.1 | 48 | 4.24e-12 | TNAHLISQSS | TNITQFPLSYSSNTGPPTSATPSHYSKESSVSWHTDS | LPSFLDFPEN |
| G003939t1 | 111 | 1.96e-11 | RNAPFISQSP | TDATALPLPQSSNSALPQSTISSHFNKESSGSWCTDP | GFLDFPVNTP |
| A131.37 | 164 | 1.18e-10 | CNNLENSMND | DDNVSPQDASGCNLVGQSCRGSVMTSEQADWQVVADP | TGLVITNNCE |
| X14587.1 | 96 | 2.04e-9 | NVSSFISQPS | NAGNFMGSANLSHTGIYHPSTSNYQGEPLESNWRPDS | VQGILDYSDN |

PSR Motif 9

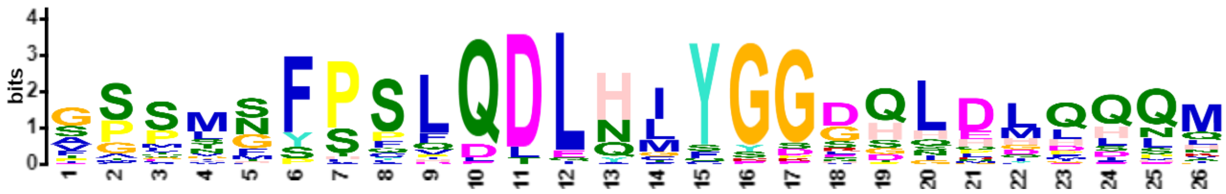


| Name | Start | *p*-value | Sites | | |
|---|---|---|---|---|---|
| U0123G0060 | 207 | 1.89e-29 | VLDVCSIEDI | GPSMGFPSLQDLHMYGGGHLDLQQQM | ERPMESFFAC |
| C1G159000 | 200 | 4.64e-28 | VSDMSNMKDL | SSSMSFPSLQDLHIYGGEQLDMQQQM | DRSSLDGFSP |
| U0095G1450 | 182 | 1.09e-26 | ILDVCSIKDI | GPSMGFHSLQDLHMYGGGHLDLQQQM | ERPMESFFAC |
| T1G069400.1 | 213 | 3.39e-26 | IVDVCSFKDI | GPSMGFASLQDLHMYGGGHLDLQHQM | ERPMEAFFAN |
| F36717001 | 193 | 4.65e-26 | VPDMGAMKDF | GSSLNFPSLQDLNIYGGDQLDLQQSM | DRSLDGFIQN |
| Z3P23640_001 | 183 | 5.44e-26 | VPDMGAVKEM | GSPVSYPSLQDLHIYGGDQLEMQHQM | ERPLDGLFPT |
| Y806121g001 | 194 | 1.43e-24 | VVDMAAMKDM | GSQMSYPSLQDLQLYGGDQLDMQQQM | DRPLDGFFPT |

| | | | | | |
|---|---|---|---|---|---|
| O30017.m000317 | 183 | 9.76e-24 | GVPDLGAMKD | FGPLNFPQFQDLNIYGGDQLDLQQNM | DRPSLDGFMP |
| X07475.1 | 143 | 6.02e-23 | VIDMVSMKEM | ASPMCFPSLQDLHLYGGDHLEIPQQI | ERPLDMFFPT |
| Z11P14430_001 | 190 | 6.83e-23 | VADMGAIKEM | GSPMSFPSLQDLHLCGGDQLDLQSQV | DGPLDGFYPI |
| B009935 | 205 | 1.44e-22 | LDVSAMKDFG | SPVNTFPSLQDLHIYGGEQLDLQSQM | DRTSLDGFMP |
| M10G174100.1 | 193 | 1.63e-22 | GIPGMGAMKE | FGTLNFPAFQDLNIYGGDQLDLQHNM | DRPSLDGFMP |
| A094.1 | 202 | 3.01e-22 | SSHQGLLDLG | ASCMNYPCLQDLHIYGGDQNPHHHQL | ELQSGGQIDS |
| U2141G0040 | 585 | 3.83e-22 | SLGNQAVIDI | GSSISFPSLQDLHLYEGRHLELQQQM | ERPMDSFLAF |
| S04G058700.1 | 223 | 4.32e-22 | MLDVCSLKDM | GPSMGFPSLQDLHMYGGGGHLDLQQQ | MERPMEAFFA |
| H32.94 | 199 | 6.96e-22 | GADMGTMKNF | VPPLNFSSFQDLNIYGGDQLDHHLQQ | NMDRSSSLDH |
| Y1141531g006 | 193 | 2.50e-21 | VIHVGAMKDM | GSPMSFPSLQDLHLYGGDDQLDMQQH | MDRPLDGFFQ |
| AB08.2515 | 193 | 3.52e-21 | SFKGIGNQDL | GSPMSFPSLQDLHIYSGDQLDGFFPN | DQTLSLGKKR |
| M08G081800.1 | 201 | 1.20e-20 | GVPDMGAMKD | FGPLNFPPFQDLNIYGSGQLDLLHNM | DRPSLDGFMS |
| R2G052544 | 224 | 2.86e-20 | MLDVCSLKDM | APSMGFPSLQDLHMYGGGGGHCLDLQ | QQMERPMEAF |
| W2g07770.1 | 187 | 3.00e-18 | LGNHQAAVLD | VCSMGFPSLQDLHMYGGAGGGHLDLQ | QQQPPASTME |
| T4G185800.1 | 198 | 1.94e-17 | SLGNQAVLDI | GSSTGFSSLQDLHFYGGSSHMDHLLH | QMERPMDSFL |
| EH01655.1 | 208 | 8.88e-17 | SAAKNFLPPP | PPENNFPYLQDLNIYGGGDHHQIISN | NKSGKSPLIW |
| U3413G0110 | 200 | 2.52e-16 | KSLGNQAVLD | IGSSMSSPPQDLQLYGGSHLELQQQM | ERPMDSFLAF |
| V3g05500.2 | 219 | 2.04e-15 | LDVCSIKDIG | AASMGFPSLQDLHLYGDLQQNQPIES | FFACSDGGIG |
| K15G263700.1 | 215 | 1.01e-14 | DMGVVMKEFG | SPLGFSSFQDLENIYGGDQIDLQQNM | EKPPLDHGFM |
| W6g45410.1 | 222 | 1.28e-14 | LGSQAVLDIG | SSMSFPSLQDDLQLYGGSHLDHLHQQ | HEQMEIRPSI |
| EM01502.1 | 204 | 1.74e-14 | MSCGLKDFGP | AMNNSFPSLQDLNLYGPTDQQLLDHH | HHNHMERSSS |
| F15900001 | 197 | 2.56e-14 | YTEVGSMKDF | SSSVNFPCLEDLHIYGERPNPYDSSK | APIIWSNDMQ |
| R2G081671 | 200 | 3.22e-14 | LGSQGVLDIG | TSSTSFSSVQDLQCFYGGSSHMDQLL | HQMERPMDGF |
| K08G163500.1 | 206 | 6.36e-14 | DMGVVMKEFG | SPLGFSSFQDLENIYGGNQIDLQQNM | EKPSLDHGFM |
| S10G220000.1 | 206 | 1.34e-13 | GSSQAVLDIG | TSSTSFSSVQDLQGFYGGSSHMDQLL | HQMERPMDGF |
| Z8P33560_001 | 207 | 2.59e-13 | ADMAAMKDLD | PSMCFPSPHDLQALYGGDQLEMHQQM | NASKHPFVWD |
| M11G023600.1 | 195 | 2.59e-13 | VSEMGNMKEI | VSASNFPPIQDLQTYGDHSHDGFLPT | DDSMSSCTIP |
| AC21_2850V3.1 | 423 | 7.50e-11 | QNGLGFMHGG | STSASYTSLQDLYCSSGDTLRGGLVK | AEMNLAERNH |
| V1g31837.1 | 209 | 9.06e-11 | YRSLGGASMD | VGSSMSFQDLTLYGSGSSHLDLQQQM | EIRPTMAPMD |
| Y728141g003 | 581 | 4.25e-9 | HEQLEPLKKS | EAGHHYPFLQTIHIQRKLQLQIEEQG | KYLQMMFEKQ |

### PSR Motif 10

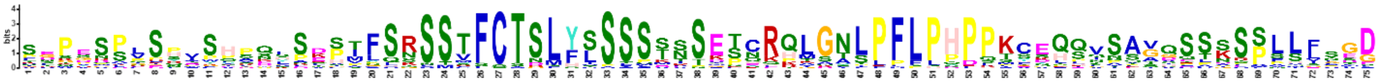

| Name | Start | *p*-value | Sites | | |
|---|---|---|---|---|---|
| U0176G0870 | 81 | 4.84e-74 | PFNVQKRSPE | SDPESPPSQISHPKFSDPIFSNSSTFCTSLFSSSSTNSEPCRQMGTLPFLPHPPKCEQQVSAGQSSSSSLLLNGD | IGNAHDEAEH |
| S10G254300.1 | 87 | 5.61e-70 | SVNLQKRSPE | SDPESPQSHVSHPKFSEPMFSNSSTFCTSLFSSSSTKTEPCHQMGTLPFLPHPPKCEQQVSAGQSSSSSLLLSGD | TGNGLDEAEQ |
| T4G270600.1 | 81 | 5.12e-69 | SVNLQKRSPE | TDPESPLSHISHPKFSDPILSNSSTFCTSLFSSSSKNTDPCRQMGTLPFLPHPPKCEQQVSAGQSSSSSLLFAGD | TGNALDEAEH |
| Z10P23260_001 | 71 | 2.40e-65 | PLHTKKSSSG | PEPGSSCSYVSHPQYPEHMFSRSSTFCTSLYSSSSTSSESCQKLQNLPFLPHPPKCKHQNSAVQSSNSPLQFSGD | ISATSGEDEH |
| M07G003200.1 | 81 | 2.76e-65 | TAQHQMCCLK | FGPDSPLSPTSHVQSSKSTFQRSSVFCTSLYLSSSSISETNRQLGNLPFLPHPPTYSHSVSATDSTKSPLLFSED | LSNQCDEEHS |
| W6g49040.1 | 79 | 3.39e-64 | ILPFQKRSSE | PEPESPLSHVSHPNVSEPVYSNSSTFCTSLFSSSSMETEPCRQLGTLPFLPHPPKCEQQVSAGHSSSSSLLVPGG | DGDIGNAHDE |
| M14G000700.1 | 15 | 1.87e-59 | TVQHQKYHPK | SGPDSPVSLAYHVQLSKSTFQRSSVFCTSLYLSSSSISETNRQLGNFPFLPHPPTYSQSVSATDSTKSPQLVSED | LSSPFDEERS |
| X11622.1 | 65 | 1.32e-58 | YLLNPKLSPE | SDSESPISHVSLPHYSEPIFSRSSMFCTSLFSSSSKSSDSCRRLSNLPFLPHPLKCEQQTSAAQDSDSPFHFTSD | INNTPNSEDE |
| AB01.3572 | 53 | 1.49e-58 | QFNHPNLIQE | PEMRSPLSNVPHPQCSENPFSRSSTFCTRLFLSSPSSSMTCRQLKNLPFLPHPPKSEQLVSAVQCSSSPSLFSGD | MCDVHTEGEH |
| Z7P15740_001 | 63 | 1.49e-58 | AQTELLKSSS | FKPGSPRSYVSHSQCSDHMFSRSSTFCTSLYSSSSAISESSRKLSSSPFLPHPPKCEKQNSAVQLSSSPLLFNGD | TSARSCEDEH |
| O30169.m006598 | 128 | 2.40e-58 | IVQPPKYFLK | SGPDMPLSPASHIQHSKSTFQRSSVFCTSLYLSSSSSSETNRQLGNLPFLPHPSAHAHSLSAIDSTKSPLLFTDD | ISNPYDEEHS |
| B021120 | 39 | 6.26e-58 | MFQLQKISPD | PEPSSPVSYESYPQYPTSVFSRSSTFCTDLYISSSTSSETHRHLGNLPFLPQPQTFDQPIATVQSSKSPLLLSGD | SSNQTEEKRS |
| V1g34470.1 | 79 | 1.82e-57 | PFRLQKRSPL | SDPESPLSHVSQPNFSDPMASNSSTFCTSLFSSSSTNSAPCQQKGALPFLPPPPKCEQQVIPGQSCSSSLHLTGD | IGNAYDEAEH |
| U0164G0530 | 64 | 9.37e-57 | ELIQSSSLPK | SLPFNLQKRSPESDPEKPIFSNSSTFCTSLFSSSSTNSVPCRQMGTLPFLPHPPKCEQQVSAGQSSSSSLLLNGD | IGNDHDESEQ |
| N06G178000.1 | 78 | 9.37e-57 | STVQPVKYSL | KSGPDSPPPVFHVQHSKSTFQRCSVFCTSLYLSSSSSSETNRPLGNLPFLPHPPTHSQSVSAVGSTKSPALFSWD | ISGFQVEKET |
| G027630t1 | 83 | 3.34e-56 | PFIRTESFKN | NLKSGPSSPISPSSHAKSAFSRSSVFCTSLYLSSSSTSETQRQLGNLPFLPHPPTCGQSISAVDSSKSPVVFSED | LHNPYNEDHS |
| H10.14 | 70 | 1.39e-53 | MIQPQKHFLK | SGPSSPVSPSPNGQHPRSTFSRSSLFCTSLYLSSSSKSETQRQLGNLPFLPNPPTYNHSTSAVESAKSPAIFSED | LGNPYDDDQS |
| F29458001 | 48 | 6.29e-53 | EVQPQKLCSK | SGPYSSVSSDTDAQYPKCTFSRSSVFCTSLYLSSSSSTETHRPLGNLPFLPHPSMSYQSISAVHSTKTPFLSGDS | SGLYDEGNSE |
| Y6550950g002 | 74 | 1.35e-51 | PFCLKKLCPE | SEPGSSLSNESHTQHFEHAFSGSSTFCASLYSSSSTSSESYRQLSNLPFLPHPQKCKLENSVVQSLNSPLLHVGD | TSNVHNEDEH |
| U0548G0730 | 81 | 2.46e-49 | PFELQKCSTD | SNPESSLSRISKADLADPILSKFSTFCTSLYSSSSTNSKSCRQTSASPFLPHPPKCEQQQNSAGQSSCPSSLLFG | ADLSNGGHGD |
| W2g04640.1 | 79 | 4.78e-47 | LPKILPFSTD | SNGESSLSRMSQAEFSDPILSSSSTFCTSLYTSSPMNSGSCRKTGYLPFLPQPPKCEQQQNSAGQSSSSLMLLDA | DLRNSGHADD |

| | | | | | |
|---|---|---|---|---|---|
| T1G092500.1 | 83 | 1.46e-43 | PFNLEKCSPG | SNPDSAVSHVSQAELSDPVSSSSSTFCTSMFSSFQTNSESCRQKGALPFLPHPPKCEQKLQQQISAGQSSSSSSL | LFGADLRSGG |
| S04G036500.1 | 72 | 1.02e-39 | LQKCSPDSTN | GTPVSRVSQAGLLSKDLVSPSSSSTFCTSMYSSSSTNSKSCRQADALPFLPHPPKSEHQQTSGGQSSSSSSLLFG | TDLNNGGQSE |
| A025.314 | 39 | 6.44e-37 | TNELWNPSPY | IETTNPGSSFSHSESSRHMLSQSSSFCTSLHVSSSSVPENNRHLANLPFLPDPLKGKLPASKASSLNSFLSISED | LKTESKEHDT |
| PHL6_AT3G13040 | 53 | 4.55e-34 | PEHRKTPFIR | SQSPDSPGQLWPKNSSQSTFSRSSTFCTNLYLSSSSTSETQKHLGNSLPFLPDPSSYTHSASGVESARSPSIFTE | DLGNQCDGGN |
| N18G144000.1 | 57 | 7.64e-34 | VNSITTQLQQ | KTNANPVPNHSPSSTSHSPGLKSTMFCTDLHLSSMKSSETNRPLGISPFLPHPHPSPPIVNLVIPSIESSLFMGG | AMTTESENDN |
| V3g03538.2 | 93 | 4.77e-33 | FELQKCSTTD | FNPGRSFSHVSQTDLSDPILSSSSTFCTSLYSSLSTNSKCRETGALPFLPHPRKLEQQHSSAGQSPNSSLLLAAD | PSNSGHGDAE |
| J3859491 | 44 | 1.38e-31 | NLQPENRNSP | FIRSHSPDSPWPKNSPQGTFSRSSTFCTNLYISSSSTSESQKHLGNTLPFLPDPSTNSQPPSAVESARSPSIFSE | DMSNPFDGDN |
| J3849734 | 41 | 8.69e-29 | FNFLNVQPET | TSKSPFIRSQSPDWPKNSTFSRSSTFCTNLYSSSSSANETQKHLGNSLPFLPDPSASASGVESARSPSVFSEDLG | NPFDGDNNSS |
| R2G010920 | 63 | 2.46e-27 | LLSARYDHRG | ACASQVELPDRISPSNASTFCTSMYSSSSSSSAAADSSKSCRRAGALPFLPHPPKREQQQQQSSSSPSLLVCTD | LGNGGQDEAE |
| G039819t1 | 97 | 2.18e-25 | EAPASAPQAS | VSQPRDLIPNPVPHNSYIICHKSTRLCTGFHSSTSESSNTEKHVGNSRCSSPLPMFDHLIPATNTPDSSLLFGEG | KTTTCDKEDF |
| A131.37 | 71 | 1.64e-24 | TREQPLTPLE | DLSVAAGETTCAYSFSLSPNLRNMCSQSSMFCPSLHSSHKEFQIGNIPFLPYPQISDHPVSSSQSLKFMLLGHGD | SSLYQYKEC |

### PSR Motif 11

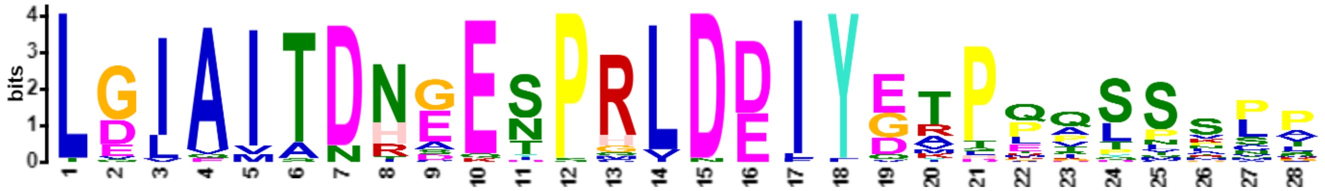


| Name | Start | *p*-value | Sites | | |
|---|---|---|---|---|---|
| S04G036500.1 | 200 | 4.14e-31 | QMEFQFLSEQ | LGIAITDNEESPRLDDIYDRPPQTSSCP | VLSCSDQDDL |
| W6g49040.1 | 208 | 6.26e-31 | QMEFQFLSEQ | LGIAITDNEESPRLDDIYGTPPQLSSLP | VSSCSNQSVQ |
| Z7P15740_001 | 192 | 1.21e-29 | QIELQLLSEQ | LGIAITDNGESPRLDDIYETPQVSSLPL | SSNHNQAVQL |
| U0164G0530 | 189 | 1.44e-29 | QMEFQFLSEQ | LGIAITDNEESPRLDDIYGAPPQLSSLP | VSSCANQGLQ |
| R2G010920 | 194 | 3.26e-29 | QVEFQFLSEQ | LGIAITDHDESPRLDDIYDRPPQTSSCP | VLSCSDQEEG |
| T4G270600.1 | 206 | 1.75e-28 | QMEFQFLSEQ | LGIAITDNEESPHLDDIYGTPPQLSSLP | VSSCSNQSIQ |
| M14G000700.1 | 143 | 2.34e-28 | QLELQFLSDE | LDIAITDHGENPRLDEIYGTPETSSKPV | TGFACYQNFP |
| T1G092500.1 | 211 | 2.70e-28 | QMEFQFLSEQ | LGIAITNNEESPRLDDIYDRPLQTSSCP | VPSYSGQEDL |
| Z10P23260_001 | 200 | 1.60e-27 | QIELQILSEQ | LGIAITDNGESPHLDDIYETPQVSSLPL | SANHNQTDQP |
| X11622.1 | 195 | 2.70e-27 | QMELQILSEQ | LGIAITDNEESPRLDDIYEKPQISPQPV | SSCQNQNPQP |
| O30169.m006598 | 256 | 2.70e-27 | QLELQFLSDE | LDIAITDHGENPRVDEIYETPEASSNPA | IGSTCNLNVA |
| F29458001 | 175 | 2.19e-26 | QLELQFLSDE | LDIAIADNGENPRLDEIYEMPQDSSTPA | MALGLTVNQN |
| U0176G0870 | 206 | 2.77e-26 | QMEFQLLSEQ | LGVAVTDNEESPRLDDIYGTPLQLSSLP | VSSCSNQGLQ |
| V3g03538.2 | 220 | 8.74e-26 | QMEFQFLSEQ | LGIAITDNEEIPRLDDIYGRPQQPLALP | LSSSSDQEDG |
| V1g34470.1 | 204 | 9.78e-26 | QMEFQFLSEQ | LGIAITDNEESPRLDDIYDTPPPQLSSL | PVSSCSNQSL |
| S10G254300.1 | 211 | 1.53e-25 | QMEFQFLSEQ | LGIAITDNEKSPHLDDIYGTPPQLSSLP | ISSCSTQSIQ |
| J3859491 | 172 | 6.23e-25 | QMELQFLSDE | LELAITDRSETPRLDEIYETPLASSSNP | VTGLSQTQRS |
| W2g04640.1 | 203 | 9.48e-25 | QMEFQFLSEQ | LGIAISNNEESPRLDDIYDRPPQLMSLP | VSSCSDQEDL |
| H10.14 | 198 | 1.05e-24 | QMELQFLSEE | LDIAITDHGETPRIDEIYEIPQASPKSA | LGFTCSQSFA |
| G027630t1 | 211 | 2.93e-24 | QLELQFLSDE | LDIAIADHGENPRLDEIYETPQKLNVAF | TCNQNSASVV |
| Y6550950g002 | 203 | 3.24e-24 | QMELQMLSEQ | LGIAITDNGESPRLDDIYETLAPSVPLS | SNCNQTSQPL |
| M07G003200.1 | 209 | 7.20e-24 | QLELQFLSDE | LEIAITDHGENPGLDEIYGTHETSSKPA | TGFACNQDSP |
| J3849734 | 167 | 3.37e-23 | QLELQFLSDE | LELAITDRAETPRLDEIYEKPMASPNPV | TVLSPSQRCD |
| PHL6_AT3G13040 | 183 | 3.37e-23 | QMELQFLSDE | LELAITDRAETPRLDEIYETPLASNPVT | RLSPSQSCVP |
| A025.314 | 168 | 7.11e-23 | QFDWQIISDH | LDLAITDIGENPGLDDIYGAPQISSVST | SGLECSPKHH |
| B021120 | 167 | 9.39e-23 | QLDLQILSAE | LDIAMTDNSENPRLDEIYEAPTPDSRLK | CKPFHQPIHP |
| N06G178000.1 | 207 | 8.79e-22 | QLELQFLSDE | LDIAITDHGENPRVDEIYETSESYTAMG | LTCNQNFASV |
| AB01.3572 | 181 | 3.18e-21 | QMDLQFLSEQ | LGIEMADNGESPCLDDLYDVPQVSCVPP | STSESNQNNQ |
| U0548G0730 | 206 | 2.11e-19 | HMEFQFLSEQ | LVIAVTNNEESPRLDDIYEYDRPPQHSS | LPVSSFSDQE |
| Q58g00470.1 | 135 | 8.91e-19 | QFNLQILSEQ | LGIQIADTDEYPQLNDIYQMPQTSSMPM | MTGSSENSQL |
| J3871004 | 142 | 8.53e-17 | QMELQFLSDE | LQLAITDRAQTPRLDEIYQVTGSSPGQN | CVPAAMPVLS |
| A131.37 | 201 | 4.18e-14 | QADWQVVADP | TGLVITNNCEDSMLDEFFVATKGSSSNT | IGFQGNHMDG |

PSR Motif 12

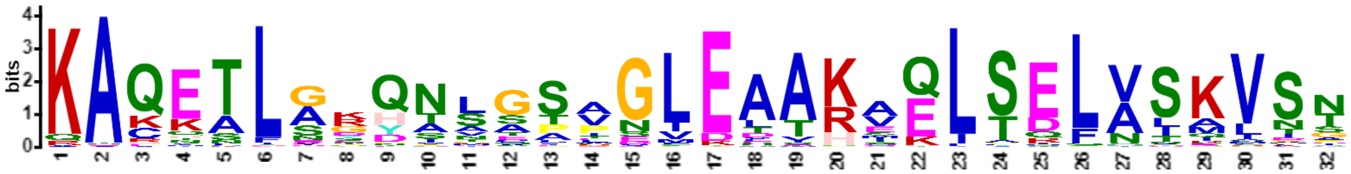


| Name | Start | *p*-value | Sites | | |
|---|---|---|---|---|---|
| G026448t1 | 254 | 6.15e-37 | QGKYLQAVLE | KAQETLGRQNLGSVGLEAAKVQLSELVSKVSN | QCLNSAFSDL |
| AB09.1410 | 173 | 6.50e-36 | QGKYLQSVLE | KAQETLGKQNLGSAGLEAAKVQLSELVSKVSN | ECFSSAFPGV |
| AB07.828 | 281 | 6.50e-36 | QGKYLQSVLE | KAQETLGKQNLGSAGLEAAKVQLSELVSKVSN | ECFSSAFPGV |
| Y771701g004 | 188 | 6.50e-36 | QGKYLQSVLE | KAQETLGKQNLGSAGLEAAKVQLSELVSKVSN | ECLNTTFPGL |
| Z5P07670_001 | 130 | 2.37e-35 | QGKYLQSVLE | KAQETLGKQNLGSPGLEAAKVQLSELVSKVSN | ECFSNAFPGL |
| B008138 | 184 | 4.79e-35 | QGKYLQSVLE | KAQETLGRQNLGSAGLEAAKVQLSELVSKVST | ECLNSTLSEL |
| B108138 | 184 | 4.79e-35 | QGKYLQSVLE | KAQETLGRQNLGSAGLEAAKVQLSELVSKVST | ECLNSTLSEL |
| M13G060200.1 | 196 | 2.22e-34 | QGKYLQAVLE | KAQETLGRQNLGTVGLEAAKVQLSELVSKVST | QCLNSTFSEL |
| O29950.m001149 | 189 | 5.01e-34 | QGKYLQSVLE | KAQETLGRQNLGSIGLEAAKVQLSELVSKVST | QCLNSAFSEL |
| H785.2 | 189 | 1.77e-33 | QGKYLQAVLE | KAQETLGRQNLGAVGLEAAKVQLSELVSKVST | QCLNSAFSEL |
| F33381001 | 184 | 1.77e-33 | QGKYLQAVLE | KAQETLGRQNLGAVGLEAAKVQLSELVSKVST | QCLHSAFSEL |
| M19G032700.2 | 184 | 1.10e-32 | QGKYLQSVLE | KAQETLGRQNLGTVGLEAAKVQLSELVSKVSS | KCLNSAFSEL |
| N09G142300.1 | 189 | 1.68e-32 | QGKYLQAVLE | KAQETLGRQNLGTVGLEAAKVQLSELVSKVSA | QCLNSAFSEL |
| N08G149600.1 | 190 | 2.07e-32 | QGKYLQAVLE | KAQETLGKQNLGTMGLEAAKVQLSELVSKVST | QCLNSAFSEL |
| EJ01031.1 | 197 | 1.78e-31 | QGKYLQSVLE | KAQETLGGQNMGTVGLEAAKVQLSDLVSKVST | QCFSSAFSDM |
| U2430G0200 | 191 | 3.09e-31 | QGKYLQSVLE | KAQETLAKQNAGSVGLETAKMQLSELVSKVST | ECLHNAFTGF |
| U0468G0380 | 338 | 3.09e-31 | QGKYLQSVLE | KAQETLAKQNAGSVGLETAKMQLSELVSKVST | ECLHHAFTGF |
| T9G419000.1 | 191 | 3.09e-31 | QGKYLQSVLE | KAQETLAKQNAGSVGLETAKMQLSELVSKVST | ECLQHAFTGF |
| W3g20900.1 | 191 | 3.70e-31 | QGKYLQSVLE | KAQETLAKQNAGSVGLETAKMELSELVSKVST | ECLQHAFSGF |
| L2g086450.1 | 193 | 7.51e-31 | QGKYLQSVLE | KAQETLGRQNLGIVGLEAAKVQLSELVSKVSS | QCLNSTFSEM |
| K13G316600.1 | 189 | 7.51e-31 | QGKYLQSVLE | KAQETLGRQNLGIVGLEAAKVQLSELVSKVSS | QCFNSAFTEL |
| D10g083340.1 | 179 | 7.51e-31 | QGKYLQAVLE | KAQETLGTQNLGTIGLEAAKVQLSDLVSKVSN | QCLNSAFSEI |
| PHL9_AT3G04030 | 188 | 1.49e-30 | QGKYLQSVLE | KAQETLGRQNLGAAGIEAAKVQLSELVSKVSA | EYPNSSFLEP |
| Y6550973g003 | 158 | 1.76e-30 | QGQYLQSVLE | KAQETLGKQNLGSTGLEAAKVQLSELVSKVSH | ECLNTAFPGL |
| K12G184700.1 | 189 | 5.56e-30 | QGKYLQSVLE | KAQETLGRQNLGVVGIEAAKVQLSELVSKVSS | QCLNSAFTEP |
| J3845853 | 188 | 6.52e-30 | QGKYLQAILE | KAQETLGRQNLGPAGIEATKAQLSELVSKVSS | EYPDASFLEL |
| PHL10_AT5G18240 | 188 | 1.43e-29 | QGKYLQSILE | KAQETLGRQNLGAAGIEATKAQLSELVSKVSA | DYPDSSFLEP |
| S01G386700.1 | 191 | 3.55e-29 | QGKYLQSVLE | KAQETLSKQNAGSVGVETAKMQLSELVSKVST | ECLQHSFTGF |
| R2G009060 | 196 | 3.55e-29 | QGKYLQSVLE | KAQETLSKQNAGSVGVETAKMQLSELVSKVST | ECLQHAFTGF |
| AD140s0042.1 | 196 | 7.41e-29 | QGKYLQSILE | KAKETLAGHTSSSPGLEAAHAELTELASKVSN | SDPLGGGHSF |
| AH181986g0010 | 188 | 9.89e-29 | QGKYLQSVLE | KARETLAGHNLGSVGLEAARAEISELASKVSS | ECLNNAFSSL |
| AD078s0062.1 | 191 | 1.32e-28 | QGKYLQSILE | KAKETLAGHASSSPGLEAAHAELTELASKVSN | DPLGQSFSWM |
| K11G183400.1 | 197 | 1.52e-28 | QGKYLQAVLE | KAQETLGRQNLGVVGLEAAKLQLSELVSKVSS | QCLNSAFSEL |
| K12G089100.1 | 192 | 1.75e-28 | QGKYLQAVLE | KAQETLGRQNLGAVGLEATKLQLSELVSKVSS | QCLNSAFSDR |
| Z6P32220_001 | 187 | 6.94e-28 | QGKYLQSVLE | KAQETLGKQNLGSAGLEVAKFQLSELVSKVSV | DSCLTSSEGS |
| J3828237 | 190 | 1.35e-27 | QGKYLQSVLE | KAQETLGKQNLGEAGLEAAKVHLSEFVSKVSA | EYPNTSFLEQ |
| L6g032990.1 | 186 | 4.92e-27 | QGKYLQSVLE | KAKETLGRQNLGAMGLDAAKVQLSELASRVST | ENLDSKFSEL |
| Y785961g002 | 181 | 6.33e-27 | QGKYLQSVLE | KAQETLGKPSVGSVGLEAAKVQLSELASKVSK | ECFGTAFPNL |
| D10g085620.1 | 171 | 7.17e-27 | QGKYLQSVLE | KAQETLGRQNMETVGLEAVKVQLSEFVSKASN | QCLNSPFTDI |
| Z8P21520_001 | 194 | 1.04e-26 | QGKYLQSVLE | KAQETLGKQNLGSPGVEDAKVQLCELVSKISS | ECFSNAFQGS |
| N03G148100.1 | 184 | 1.04e-26 | QGKYLQSILE | KACKALNDQAAASAGLEAAREELSELAIKVSN | ECQGIVPMDN |
| G020332t1 | 179 | 1.04e-26 | QGKYLQSILE | KACKALNDQAAASAGLEAAREELSELAIKVSN | DCQEMIPLDN |
| K16G152200.1 | 189 | 1.18e-26 | QGKYLQAVLE | KAQETLGRQNLGAEGVEAAKVQLSELASRVSP | QSLDSKFSEL |
| AC21_2850V3.1 | 189 | 1.33e-26 | QGKYLQAILQ | KAKETLASHTSESPGLEAAHAELTELASKVTT | VGYLSDFSNL |
| Mpo198s0044 | 184 | 1.33e-26 | QGKYLQSILE | KARETLAGHTAASPGLEAAHAELSDLASKVTT | ENLSPAFSLT |
| B000838 | 161 | 1.92e-26 | QGKYLQSVLK | KAQETLAGYNSSSLGLEAAKAELSQLASMVDN | RCPSSSFSGL |
| M17G054800.1 | 181 | 2.76e-26 | QGKYLQSILE | KACKALNDQAVATAGLEAAREELSELAIKVSN | ECAGIAPLDT |
| M01G314800.1 | 181 | 2.76e-26 | QGKYLQSILE | KACKALNDQAVATAGLEAAREELSELAIKVSN | ERAGIAPLDT |
| AD078s0059.1 | 192 | 2.76e-26 | QGKYLQSILE | KAKETLANHTAATPGLEAAHVELTELASKVIT | DPLGPSFPSL |
| Z2P01180_001 | 189 | 5.01e-26 | QGKYLQSVLE | KAQETLGKQHLGTPGLEAAKVHLSQLVFKVSN | ECFSNALTGL |
| V1g63690.1 | 190 | 5.01e-26 | QGKYLQSVLE | KAHETLAKQNTGSASLENAKMQLSELVSKVST | ECLHNAFTGF |
| AH21538g0020 | 473 | 6.34e-26 | QGKYLQSILE | KARETLAGYNVGSIGLEATRAELSELASKVKS | ECFNSTFSAL |

| | | | | | |
|---|---|---|---|---|---|
| O29904.m002952 | 177 | 7.13e-26 | QGKYLQSILE | KACKALNDQAAVSAGLEAAREELSELAIKVSN | ECQGIVPADN |
| AD173s0028.1 | 208 | 8.01e-26 | QGKYLQSILE | KAKETLAGHTSSSPGLEAAHAKLAELASKVTN | DPVLMGHGSF |
| H128.40 | 148 | 2.82e-25 | QGKYLQSVLK | KAQETLAGYSSSSVGIELAKAELSRLVSMVNT | GCPSSSFSEL |
| AA10677 | 156 | 3.16e-25 | QGKYLQSVLE | KAQEAFGKQNSGSAGLEASRTQLSELISKVSN | ECFTSTFPYL |
| AD088s0085.1 | 216 | 3.53e-25 | QGKYLQSILE | KAKETLAGHRSSSPGLEAAHAELTELASKVGN | DPSGCSFSLV |
| AC18_20920V3.2 | 189 | 3.53e-25 | QGKYLQSILE | KAKETLASHTSESPGLEAAHAELTELANKVTT | VGMIPLGFST |
| G011693t1 | 165 | 4.41e-25 | QGKYLQSVLK | KAQETLAGYSSSSVGVELAKAELSQLVSMVNT | GCTSSSFSEL |
| B022348 | 195 | 4.93e-25 | QGKYLQSILE | KACKALNEQTIASVGLEAARQELSELAIKVSN | DCLGMVPLDT |
| A119.73 | 184 | 4.93e-25 | QGKYLQSVLE | KAQETLAKQNPGSSGLEATRAQISELVSQVSA | ECLNSAFSGL |
| AD1140s0043.1 | 190 | 6.14e-25 | QGKYLQSILE | KAKETLANHTGVAPGLEAAHAELTELASKVNT | EPLDPSFSSL |
| K02G070900.1 | 189 | 6.86e-25 | QGKYLQAVLE | KAQETLGRQNIGAEGVEATKVQLSELASRVSP | QSLDSRFSEL |
| A089.17 | 172 | 7.65e-25 | QGKYLQSILE | KACKALADQTVASAGLEAARQELSALVIKVSN | GCLSAPSEFL |
| Z9P05030_001 | 187 | 1.32e-24 | QGRYLQLVLE | KAQERLEKQSLGSVGLEVAKVQLSELVTKVSG | ECFGNSFLGV |
| T3G199300.1 | 188 | 1.47e-24 | QGKYLQAVLE | QAQETLGKQNLGPANLEDAKIKISELVSQVSN | ECFSNAITDI |
| R234155.1 | 169 | 1.47e-24 | QGKYLQAVLE | QAQETLGKQNLGPANLEDAKIKISELVSQVSN | ECFSNAITDV |
| AF481.g073115 | 224 | 2.25e-24 | QGRYLQSILE | KAQQTLAGQTVTSDGLEVARAELSDLATKVSN | ECLNPSFSLV |
| S09G178000.1 | 188 | 2.78e-24 | QGKYLQAVLE | QAQETLGKQNLGPANLEDAKIKISELVSQVST | ECFSNAITDV |
| AA17183 | 186 | 4.23e-24 | QGKYLQTVLE | KARTTLEIQNTGSAGHEAAKEQLSELVSKVSN | ECFTSAFIGL |
| AC19_2940V3.3 | 193 | 4.23e-24 | QGKYLQSILE | KAKETLASHTMESPSLEAAHAELSELATKVTT | LGMFPSGFSN |
| J3831963 | 180 | 4.69e-24 | QGKYLQSILE | KACKAFDDQAAAFVGLEAAREELSELAIKVSN | SSQGTAVPFF |
| B019826 | 185 | 5.21e-24 | QGKYLQSVLE | KAQETLGRQNLGSVEFDAAKMQFPELVSKVSA | ECLNSTFPEL |
| M08G087600.1 | 166 | 5.78e-24 | QGKYLQTVLK | KAQETLAGYNSSSMGIELAKAELCRLVSMVNS | GCPSSSISEL |
| W5g40960.1 | 188 | 8.72e-24 | QGKYLQAVLE | QAQETLGKQNLGPASLEDAKIKISELVSQVSN | ECLSNAVTEI |
| N01G035100.1 | 177 | 8.72e-24 | QGKYLQSVLK | KAQETLSGYSSSSLGIELAKAELSRLVSMVNN | ECRSPSISEL |
| AC13_23440V3.1 | 184 | 1.07e-23 | QGKYLQTILE | KAKETLAGHTSASPDLKAAHAELTELASKVIG | DPGTFAASTS |
| F21072001 | 178 | 2.17e-23 | QGKYLQSILE | KACKALKDQAAATAGLEAAREELSELQIKVSN | DCEGMNPLET |
| PHL2_AT3G24120 | 184 | 2.65e-23 | QGKYLQSILE | KACKAFDEQAATFAGLEAAREELSELAIKVSN | SSQGTSVPYF |
| R2G701218 | 188 | 3.23e-23 | QGKYLQAVLE | QAQETLGKQNLGPANLEDAKIKISQLVSQVST | ECFSNAITDV |
| V2g21880.1 | 185 | 3.57e-23 | QGKYLQSVLE | QAQESLGKQNLGPANLEDAKIKISELVSQVSN | ECFSNAVTDI |
| F19971001 | 150 | 3.57e-23 | QGKYLQSVLK | KAQETLAGYNSSSVGVELAKAELTQLVSIFDT | GCPSSSFSEL |
| K01G049100.1 | 156 | 1.15e-22 | QGKYLQSVLK | KAQEALAGYNSSPVGIELTKAELSQLVTIINN | ACPSSPISEL |
| C3G342000 | 160 | 1.85e-22 | QGMYLQSILK | KAQETLSGYNSSSTELEAAKTEVSELVSMVET | GCLDSSFSEF |
| AC122_8217V3.1 | 193 | 1.85e-22 | QGKYLQSILE | KAKETLASHTNESPSLEAAHAELTKLATKVTT | VGNMMPSGFS |
| AC22_8210V3.1 | 193 | 1.85e-22 | QGKYLQSILE | KAKETLASHTNESPSLEAAHAELTKLATKVTT | VGNMMPSGFS |
| N15G053300.1 | 185 | 2.03e-22 | QGKYLQSILE | KACKALNDEAAASAVLEAAREELSELAIKVSG | KCQEIVPVAN |
| AC4_16540V3.1 | 189 | 2.45e-22 | QGKYLQSILE | KAKEALGSHIGASPGLETVHAELTELASKVNS | EPMNMCFPPL |
| AC12_10900V3.1 | 209 | 2.45e-22 | QGKYLQTILE | KAKEALGRQIGESPGLETVHAKLTELVSKVNI | EPMNMSFPPF |
| C2G398700 | 170 | 4.72e-22 | QGKYLQSILE | KACKALNEQPVASVGLEEARQELSELAIKVTN | DCLGAVPLDK |
| K02G108500.1 | 161 | 1.54e-21 | QGKYLQSVLK | KAQEALAGYNSSPVGIELTKAELSQLVTIIND | ACPSSPISEL |
| J3860014 | 181 | 2.21e-21 | QGKYLQSILE | KACKAFEEQAAMFTGLETAREELSELAIKVSN | SSQGATVPYF |
| AC10_13030V3.1 | 186 | 2.42e-21 | QGKYLQSILE | KAKETLADHTSTSPVLKLAHEELTELASKVID | YEIPKPAFES |
| X12613.1 | 188 | 2.89e-21 | QGKYLQAVLE | KAQETLGKQNLTSSGLEAAKIKLAEQVPEVSN | EFLNNGFPNS |
| AC3_14670V3.1 | 182 | 3.76e-21 | QGKYLQTILE | KAKETLAGHTSTSPHVKAAHDELTELASKVIS | DPGTFEVTLP |
| PHL3_AT4G13640 | 180 | 1.27e-20 | QGKYLQSILE | KACKAIEEQAVAFAGLEAAREELSELAIKASI | TNGCQGTTST |
| U0261G1090 | 241 | 2.50e-20 | QGKYLQAVLE | QAQETLGKQNSGPANPEDAKIKISEPASQVSN | ECFSNTMSET |
| AD1173s0030.1 | 192 | 6.76e-20 | QGKYLQSILE | KAKETLANHAGVVPELEAAHAKLTDLASTVIT | EPGGPSFPSL |
| L5g041350.1 | 155 | 8.65e-20 | QGKYLQSVLM | KAQEALSGYNSSPIGIKLTKDELSQLVTMINN | ACPSSPISDL |
| J3831359 | 173 | 1.30e-19 | QGKYLQSVLL | KAQHTLAGYTSSTLGMDFARTELSRLASMVNQ | SSSFSELTQV |
| EL01279.1 | 155 | 1.30e-19 | QGKYLQSVLR | KAQETISEYSSCSIEVEHAKAQLSQLASMVDS | TCPSSSFSVL |
| EE00086.1 | 154 | 1.41e-19 | QGKYLQSVLR | KAQETLSEYSSCSIEVEHAKAQLSQLMSMVDS | GCTSSSFSIL |
| AC3_23200V3.3 | 186 | 2.28e-19 | QGKYLQSILE | KAKETLADHTSASPVLKEVHEELTTLASKVIN | YEAPKPAFES |
| AH120020g0010 | 190 | 2.47e-19 | QGKYLQAILE | KAQQTLASEASTSAGIEATRAKLADLASKLPT | NSVRPNYSAM |
| K20G035300.1 | 161 | 3.39e-19 | QGKYLQSVLT | KAHEALARHSSSTTGVELAKFELSLLVSIINN | ACPSSPISEL |
| X02890.1 | 151 | 5.43e-19 | QGKYLQNVLR | KAQETLSAYSSNSIGTDVVKAELSELASAVGN | ECLNSPPPQP |
| A025.171 | 178 | 6.35e-19 | QGKYLQSILE | KARETLARHHMGSAGIEETRRELSELSAKVSD | GCRNPATFSL |
| W2g47190.1 | 171 | 6.86e-19 | QGKYLQTILE | KAQNNLSYDATGTANLEATRTQLTDFNLALSG | FMNNVSQVCE |
| J3852594 | 168 | 7.42e-19 | QGKYLQSVLM | KAQQTLAGYTSASLGMDFARSELSRLASMMNP | SSSFSEQTQV |
| T1G286400.1 | 166 | 8.02e-19 | QGKYLQTILE | KAQNNLSYDASGAANLEATRSQLTDFNLALSG | FMDNVSQVCE |
| U1295G0130 | 187 | 1.27e-18 | QGKYLQAILE | RAQKNLSYDASGTVNLEATRSQLTDFNLALSG | FMDNVTQACE |
| R233960.1 | 170 | 2.93e-18 | QGKYLQTILE | KAQSNLSYHATGAANLEATRSQLTDFNLALSG | FMDNVTQACE |
| PHL8_AT1G69580 | 180 | 3.16e-18 | QGKYLQSVLM | KAQQTLAGYSSSNLGMDFARTELSRLASMVNR | GCPSTSFSEL |
| K07G229800.1 | 161 | 4.59e-18 | QGKYLQSVLT | KAHEALARHSSSTTGMELAKAELYQLESIINN | ACPDSPLSEL |
| AF035.g025591 | 122 | 9.60e-18 | QGRYLQSILE | KAQQTLAGQVVASDGLDTAQAELSALATKFPN | KCITNLELAT |

| | | | | | |
|---|---|---|---|---|---|
| V3g52500.2 | 165 | 2.29e-17 | QGKYLQTILE | KAQKNLTYDSSAATNLEATRSQLTDFNLALSG | FMDDATQVCE |
| AA00348 | 188 | 2.64e-17 | QGKYLQSVLE | KAEITLENQNLCPVGSEAAKVQISELASNMKE | IPDPHNLKVN |
| B106931 | 167 | 2.84e-17 | QGKYLQAILE | KAQKSLSTDTKCPRSLEATRAQLTDFNLALSG | LMENMNQVRE |
| B006931 | 167 | 2.84e-17 | QGKYLQAILE | KAQKSLSTDTKCPRSLEATRAQLTDFNLALSG | LMENMNQVRE |
| EA00576.1 | 165 | 3.51e-17 | QGKYLQSILE | KAQHSLSTDINQSENLESTKAQLTDFNLALSN | FMQTINGDEI |
| C3G208000 | 163 | 4.05e-17 | QGKYLQTILE | KAQKSLSSDINCPENLEATRDQITDFNLALSG | LMENMTQVNE |
| S04G270600.1 | 168 | 4.34e-17 | QGKYLQTILE | KAQSNLSYDATGGANLEATRSQLTDFNLALSG | FMDNVTQVCE |
| D12g098370.1 | 192 | 6.17e-17 | QGKYLQTILE | KACKVLNYTSVESPDLDTAREQLSELAIKGAT | NNCDGIVPVS |
| EN00196.1 | 134 | 8.16e-17 | EGKYLQSILE | KAQHSLSPDINQSENLELTKSQLTDFNLALSN | LMQTINGDEI |
| EM00636.1 | 139 | 8.16e-17 | EGKYLQSILE | KAQHSLSPDINQSENLELTKSQLTDFNLALSN | LMQTINGDEI |
| J3830924 | 173 | 1.08e-16 | QGKYLQSVLM | KAQQTLAGYTSSTLGMDFSRTKLSRFASLVNP | SSSFSELTQV |
| Y738571g007 | 150 | 1.74e-16 | QGKYLQAILE | KAQKSLSFDMNNSGSLEATRAQLTDFNLALSG | LMDNVNQVCE |
| A010.492 | 166 | 6.69e-16 | QTRYLQTVLE | RAKTALSSYNLDSSGVEAFRAELSELALSVVI | HDYMDTAISK |
| EB00920.1 | 154 | 2.31e-15 | EGKYMQSILE | KTQHSLSPDINQSENLELTKSQLTDFNLALSN | FMQTKRETI* |
| X01685.1 | 204 | 1.19e-14 | GKYLQSVLEK | AHETLAKQNLGSAPGLEAAKAQLSELVSKVST | ECLHNAFTFE |
| Q13g00280.1 | 183 | 1.27e-14 | QGKYLHSVLE | KAQETLGYQNLGSAGLEAIKETFLVSNDDRLN | TAFQPSMVDL |
| AH8183372g0010 | 177 | 2.48e-14 | QGKYLQGILE | KAQQALIYQTSTPARLGAEPAELVEMVSNVAT | DCNETASFSR |
| F22645001 | 149 | 2.98e-14 | QGKYLQAVLD | KAQQSLSINVNCPGSLEAMRAQLTNFNMALSS | LTENTNEEDM |
| D09g091880.2 | 191 | 3.57e-14 | QGKYLQSILE | KACKAFNSQSLELNGLEMNREELSELAFINGP | SLPDIGPSFE |
| AG007.g003614 | 187 | 3.57e-14 | QSRYLQSILE | RAAAALSHQATAPVELTNVKAELADLVSDVTN | ESMVSSFSTF |
| AE149357 | 177 | 5.12e-14 | QSKYLQNILE | KARDAFVGHIPTSAELEAAHAELTELAKGTLE | GFEFTVPPLP |
| O29813.m001501 | 171 | 2.22e-13 | GKYLRSVLKK | AQETLSGYNPSSAMGIEIAKAELSRLVSMVNT | GCSSSSISEL |
| X17021.1 | 172 | 2.63e-13 | QGKYLQAILE | KAHRSLPLDVNGTRDFEAASAQLTDFNLALSG | LMDNVNQVCE |
| AF027.g023545 | 206 | 6.17e-13 | QGRYLQSILE | KAQLTLARQNDSATELEPECPESSDQTTKVSA | MDCSGSNYSR |
| AA34364 | 165 | 7.30e-13 | QEKYLQSFLD | RACNALVDPNLTEIGFDAIKQQFTELSNRNIN | GGIKYEPVKL |
| EG00679.1 | 110 | 7.72e-13 | GKYLQSILEK | ACKALDEQTVVPDGGLEAAREELSELAIKVAN | DCNNNNGLLS |
| U1153G0050 | 371 | 3.04e-12 | KYLQSMLEKA | QEALEKQNVAGTTVDLEAAKTQLSELVSKASA | KRLQNERSII |
| Y850271g001 | 303 | 1.17e-10 | QGKYLHSMLE | RACNALIDPNLGLIGLASIRCDLPELPNIETD | DCLGHPHVSL |

PSR Motif 13

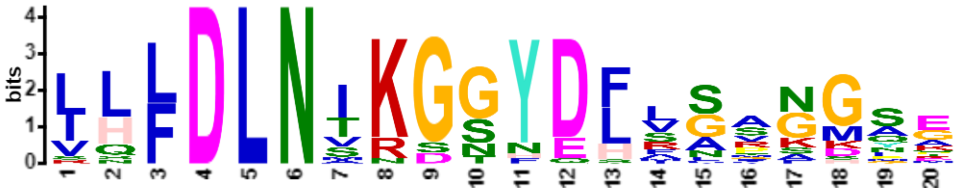


| Name | Start | *p*-value | Sites | | |
|---|---|---|---|---|---|
| M03G100100.1 | 254 | 4.37e-23 | DVKHHLQGDS | IHFDLNTKGNYDFVSANGSE | LELKMLSYRR |
| M01G133400.1 | 253 | 5.31e-22 | DVKHHLQGDS | IHFDLNTKGTYDFVAANGSE | LELKMLSYRR |
| B106931 | 256 | 3.19e-20 | EIKIKVEESS | LLFDLNVKGGYDFLGPKGSE | LDPNMNIQRI |
| B006931 | 256 | 3.19e-20 | EIKIKVEESS | LLFDLNVKGGYDFLGPKGSE | LDPNMNIQRI |
| O29827.m002675 | 237 | 1.26e-18 | NDFEHKIEGS | IHLDLNTKGSYDFAAVNGSQ | LELR |
| R2G125704 | 240 | 6.32e-18 | VKCTPKTEDS | LLLDLNIKGGYDLSSRGMQA | CELELKINQQ |
| K09G211400.1 | 236 | 6.32e-18 | DQKPKVEGGS | IQFDLNIKGSNDLVCAGGAE | MDANMISYRV |
| K01G009600.1 | 240 | 6.32e-18 | DQKPKVEGGS | IQFDLNIKGSNDLVSAGGAE | MDANMVSSYR |
| F22645001 | 232 | 1.78e-17 | DVKFREGEGF | RHFDLNTKGNYEFAGANGAD | FEAKMIAYRR |
| T7G223500.1 | 247 | 2.48e-17 | VKWTPKTEDL | LQLDLNIKGGYDLSSRGMQA | CEVDLKINQQ |
| U1295G0130 | 271 | 2.76e-17 | DVKCTPDEGL | LLLDLNIKGGYDHLSSSGMR | GGEADLKISQ |
| B021405 | 254 | 4.26e-17 | ENMIEGDEGS | SLFDLNVKGGYEFFGPKGSG | S |
| N17G009200.1 | 241 | 4.73e-17 | DLEHKVEAHP | IHLDLNTKGTYDSLSVNGST | IGTQHAFI* |
| U0225G0780 | 245 | 1.81e-16 | VKCTPKTEDS | LLLDLNIRGGYDLSSRGMPG | CELDL* |
| J3827790 | 248 | 5.82e-16 | DACVKPESGF | VHFDLNSKDGYDLLNSGKYG | IEMKPNVIAD |
| PHL11_AT5G45580 | 232 | 5.82e-16 | DACVKPESGF | VHFDLNSKSGYDLLNCGKYG | IEVKPNVIGD |
| T1G286400.1 | 250 | 7.03e-16 | DVKCTPDEGL | LLLDLNIRGGYDHRSAADLK | MNQHMR* |
| G016647t1 | 260 | 1.12e-15 | DVKLKVEGES | INFDLNTKDSFEFVAVNGNE | LQSHMFSYKR |
| V5g20520.1 | 248 | 2.32e-15 | VRCTPRTEDL | LLLDLNIKGGHDLSSTGMQX | |
| S04G270600.1 | 254 | 2.54e-15 | DVKCATDEGL | LLLDLNIRGGYDHRASADLK | MNQHMR* |

| | | | | | |
|---|---|---|---|---|---|
| V3g52500.2 | 249 | 6.13e-15 | DVKCTADEDL | LLLDLNIKGGYDHRLSSHGM | RRGDADLTVG |
| C3G208000 | 252 | 7.94e-15 | EVKVKVEGGS | LLLDLNMNGNYEFMGGNGNG | NEYDTNIHTH |
| Z6P01310_001 | 240 | 2.37e-14 | DSKLGNDAGT | VLLDLNAKGSYQLFGFFGSP | YAHLLCYRTI |
| D08g076010.2 | 329 | 2.54e-13 | IMNIKLEESS | VSFDLNSRSSYDFIGMNSAA | LEAKQFSNGR |

PSR Motif 14

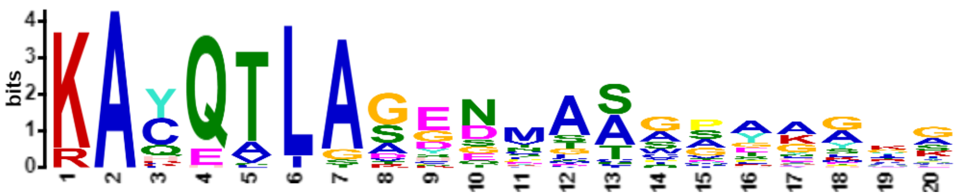


| Name | Start | *p*-value | Sites | | |
|---|---|---|---|---|---|
| O30017.m000317 | 151 | 4.56e-21 | HGKYMQNMLE | KAYQTLAGENMASGSYKGIG | NQGVPDLGAM |
| G033672t1 | 156 | 4.56e-21 | QGKYMQSILE | KACQTLAGENMAAGGYKGMG | NQGVPDMGAM |
| N03G077700.1 | 106 | 2.11e-19 | HGKYMQNMLE | KAYQTLASENMASGIYKGIG | NQGVPDMGGG |
| J3830485 | 165 | 7.87e-19 | QGKYMQSILE | RACQTLAGENMAAAVAAGGG | YKTNLGSTSL |
| Y1141531g006 | 160 | 1.92e-18 | QRKYMQTILE | KACQTLAGEGMASVSYKGHG | NQGVIHVGAM |
| F36717001 | 160 | 4.54e-18 | QGKYMQTILE | KACQTLAGENMALGNYKGIG | NQGVPDMGAM |
| Y806121g001 | 161 | 7.48e-18 | QGKYMQTILE | KACQTLAGEGMASGSYKALG | NQGVVDMAAM |
| K15G263700.1 | 175 | 1.97e-17 | QGKYMQSILE | KAYQTLAGENMASAATNLKG | AIVPHHQGIP |
| J3853188 | 165 | 2.69e-17 | QGKYMQSILE | RACQTLAGENMAAASGGGFK | GNLGSSSLSA |
| S04G058700.1 | 184 | 3.14e-17 | QGKYMQSILE | KAYQTIAAGDVAACPAAGYK | SLLGNHHQAM |
| T1G069400.1 | 177 | 5.78e-17 | QGKYMQSILE | KAYQTIASGDVAACPAGGYK | SLGNPAIVDV |
| U0123G0060 | 172 | 1.05e-16 | QGRYMQSILK | KAYQTLASGDGAACPAGYKS | LGNQAVLDVC |
| K08G163500.1 | 166 | 1.05e-16 | QGKYMQSILE | KAYQTLAGENMASAATNLKS | AIVPHHQGIP |
| C1G159000 | 167 | 1.05e-16 | QGKYMQNILE | KACQTLASDNMASGSYKSVN | NQEVSDMSNM |
| PHL14_AT1G79430 | 165 | 1.40e-16 | QGKYMQSILE | RACQTLAGENMAAATAAAAV | GGGYKGNLGS |
| H32.94 | 163 | 1.62e-16 | QGKYMQSILE | KACQTLAGENNIAAAAASYK | GLGNHCGADM |
| U3413G0110 | 172 | 2.86e-16 | QAKYMQSILE | KAYQALASDDCATWPGGYKS | LGNQAVLDIG |
| EM01502.1 | 163 | 3.78e-16 | HGKYMQTILE | KACQTLAGENNMAAAAAAAS | GSYNKAAVGP |
| W2g07770.1 | 153 | 4.59e-15 | QGKYMQSILE | KAYQTLAAGDVAAAVACGPA | GYKSLGNHQA |
| P5G0044100 | 164 | 2.20e-14 | QGKYMQTILE | KACQTLARENTATGGGEMCF | LKEMGSPLGF |
| K15G215000.1 | 166 | 2.77e-14 | QGKYMQSILE | KAYHTLAGENMATNMKGMGA | PLGTTEMGVM |
| Z3P23640_001 | 150 | 3.49e-14 | HGKYMQNILE | RAYQTLAAESMVSGGYKGQA | SRGVPDMGAV |
| R2G081671 | 170 | 8.55e-14 | QGKYMQSIVE | KAYQALGSSDCATWPAGYRT | LGSQGVLDIG |
| U2141G0040 | 556 | 1.06e-13 | QGKYMQSILE | KAHQALASGDCGTWPGGYKS | LGNQAVIDIG |
| S10G220000.1 | 175 | 3.43e-13 | QGKYMQSIVE | KAYQALGSSDCTTWPAGYRS | LGSSQAVLDI |
| R2G052544 | 184 | 3.43e-13 | QGKYMQSILE | KAYQTIATGDLAACSPVAAG | YKSLLGNPQA |
| A094.1 | 172 | 3.80e-13 | QGKYMQNILE | KAFQTIAGETMTSGSLKAGG | SSHQGLLDLG |
| V1g31837.1 | 180 | 1.04e-12 | QGKYMQSILE | KAYQTLASGGDCATWPAAGY | RSLGGASMDV |
| Z8P33560_001 | 170 | 2.28e-12 | QGKYMQSILE | KAYQTLAADGMTSGDHHHHH | HHHHQGLADM |
| AB08.2515 | 168 | 2.75e-12 | QGKYMQSILE | KACQTLSSDNMGASGSFKGI | GNQDLGSPMS |
| M10G174100.1 | 160 | 2.75e-12 | QGKYIQSLLE | KACQTLAGDQNLASGSYKGM | GNQGIPGMGA |
| AA11035 | 169 | 3.66e-12 | QGKYMQSILE | KAYQTLAGEEKSIPATASFK | FHFLPRPPFI |
| M08G081800.1 | 168 | 3.66e-12 | QGKYIQSLLE | KACQTLAGDQDLASGSYKGI | GNQGVPDMGA |
| T4G185800.1 | 169 | 5.82e-12 | QGKYMQSILE | KAHQALGTSDYATWPAGYRS | LGNQAVLDIG |
| AA17379 | 167 | 6.38e-12 | QGKYMQSILE | KACQTLAACENLQPSADAYK | AAFSGHGSVQ |
| X08334.1 | 170 | 8.38e-12 | QGKYMQTMLE | KALQTLAVESTGSGDYKEIG | NQGVIDMSSM |
| D05g007890.2 | 192 | 1.00e-11 | QGKYLQSVLK | KAQETLAGYGTSSGVELAKA | ELSQLVSMVN |
| EH01655.1 | 170 | 1.20e-11 | QGKYMQTILE | KACQTLSSGESNAAAAAASA | SSSGMYGFSA |
| U0008G1310 | 170 | 2.65e-11 | QGKYLQSVLR | RAQQVLADHNLASSPEATKA | ELSELVSEVE |
| F15900001 | 164 | 2.89e-11 | QGKYMQTILE | KACQTLTGKNGDCQSYHGVG | NQGYTEVGSM |
| B008492 | 147 | 3.43e-11 | QGKYLQSVLE | KAQETLAGCNSSCVGTRSCK | S |
| U1779G0420 | 272 | 3.73e-11 | QGRYLQSVLR | RAQQVLADHSPASSPEAAKA | ELSELVSAVE |
| W8g33750.1 | 175 | 7.30e-11 | QGRYLQSVLR | RAQQVLADHSLASSPEAATA | ELSELASAVD |
| AH21538g0020 | 216 | 1.52e-10 | QGKYLQSILE | KARETLAGYNVGSIGLEATR | AELSELASKV |

| | | | | | |
|---|---|---|---|---|---|
| V3g05500.2 | 188 | 2.65e-10 | QGKYMQSILE | KAYQSLGSGEPAAGYKSLGG | VLDVCSIKDI |
| AB03.7 | 155 | 3.62e-10 | QGKYLQSVLK | KAQETLAAYTFSSSAQAEAA | DAGAECPSLL |
| V1g17400.1 | 191 | 4.92e-10 | QGKYLHSVLE | KAQEALAKNQHAAGAGAGHE | AGKPPARQRL |
| X01094.1 | 109 | 1.80e-8 | QGKYLQSVLR | KAHETLAAYSSSSITELSEL | VSAAETLCPS |
| AG011.g005338 | 177 | 3.66e-8 | QEKYLQSILD | KAKQLLAVPETTTAASAPTP | NLLLDSTPAV |

### PSR Motif 15

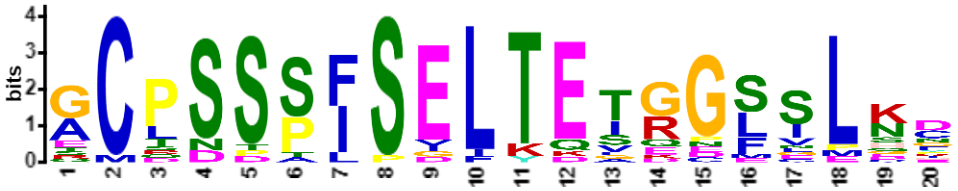


| Name | Start | *p*-value | Sites | | |
|---|---|---|---|---|---|
| K20G035300.1 | 193 | 1.84e-22 | LSLLVSIINN | ACPSSPISELTETRGLSLNC | GERKQDRGTM |
| M08G087600.1 | 198 | 2.90e-22 | LCRLVSMVNS | GCPSSSISELTETGGSILKD | IERTQMRNTV |
| K02G108500.1 | 193 | 6.08e-21 | LSQLVTIIND | ACPSSPISELTETRGLSLSC | GERKRDRGTM |
| K01G049100.1 | 188 | 6.08e-21 | LSQLVTIINN | ACPSSPISELTETRGLSLSC | GERKRDRGTM |
| F19971001 | 182 | 8.76e-21 | LTQLVSIFDT | GCPSSSFSELTETGGSGLKD | KERKPMRGTG |
| L4g081710.1 | 190 | 3.37e-20 | LSQLLSIINN | ACPSSPLSELTETRGFSLNF | GERKQNRGTM |
| K07G229800.1 | 193 | 1.18e-19 | LYQLESIINN | ACPDSPLSELTETRGLSLNC | GERKQDRGTM |
| N05G101800.1 | 209 | 2.24e-18 | LSRLVSMVNT | GCQSSSISELTEIGGSNLKD | TERKLMRGTV |
| G011693t1 | 197 | 4.83e-18 | LSQLVSMVNT | GCTSSSFSELTEGGGSSLKI | ERKPMRGTIC |
| H128.40 | 180 | 6.20e-18 | LSRLVSMVNT | GCPSSSFSELTEVGDSIMKD | VNQKGTRGTI |
| O29813.m001501 | 203 | 2.09e-17 | LSRLVSMVNT | GCSSSSISELTEIGNSILND | TTDKNQMIRR |
| L5g041350.1 | 187 | 2.09e-17 | LSQLVTMINN | ACPSSPISDLTESRGLSLNY | EERKHENGTS |
| EL01279.1 | 187 | 7.90e-16 | LSQLASMVDS | TCPSSSFSVLTESGGSLLKN | EGNKPLGHNG |
| N01G035100.1 | 209 | 8.76e-16 | LSRLVSMVNN | ECRSPSISELTEIEGSSLKG | TERKKMRGTV |
| B000838 | 193 | 1.97e-15 | LSQLASMVDN | RCPSSSFSGLTEAGGFSLQE | VGKQSMRGMD |
| C3G342000 | 192 | 3.55e-15 | VSELVSMVET | GCLDSSFSEFTKTGCFVLDN | EGKKLVRTTD |
| K11G183400.1 | 229 | 1.17e-13 | LSELVSKVSS | QCLNSAFSELKEIQGFSPHH | QKQTQTNNNQ |
| EE00086.1 | 186 | 1.39e-13 | LSQLMSMVDS | GCTSSSFSILTQSDGSVLEK | DERDKLLVHD |
| B019826 | 217 | 6.37e-13 | FPELVSKVSA | ECLNSTFPELKELQGLCLCP | QQTQKMQPTD |
| PHL8_AT1G69580 | 212 | 1.56e-12 | LSRLASMVNR | GCPSTSFSELTQVEEEEEGF | LWYKKPENRG |
| K15G215000.1 | 288 | 1.42e-9 | GDHKVQISPP | SMDSDPISEIYDTKPMILHG | ESVSDQNKFD |

### PSR Motif 16

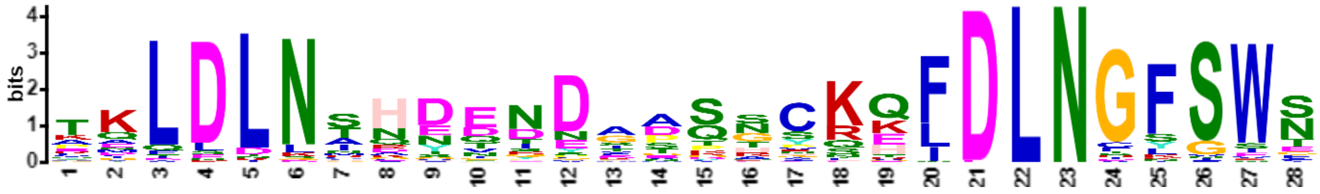


| Name | Start | *p*-value | Sites | | |
|---|---|---|---|---|---|
| N09G142300.1 | 393 | 3.19e-31 | STGYRLPYFA | TKLDLNSHDEIDAASNCKQLDLNGFSWN | * |
| M13G060200.1 | 400 | 2.19e-30 | SPGYRLSYFT | TKLDLNSHDEIDAASSCKQLDLNGFSWN | * |
| Y771701g004 | 388 | 5.42e-30 | SNGFGLQSVT | TQLDLNAHEENDGPPNCKQFDLNGFSWS | |
| D10g085620.1 | 373 | 5.42e-30 | SQEYKLSYFE | PKLDLNMHDETDAASSCKQFDLNGFSWS | * |
| H785.2 | 373 | 1.30e-29 | FLETKPMGGQ | TQLDLNARDENDAASSCKHFDLNGFSWS | * |
| V1g63690.1 | 323 | 1.99e-29 | TDGFSISCQT | TKLDLNINETNDGPQNCKKFDLNGFSWT | * |
| O29950.m001149 | 392 | 2.45e-29 | SQGYRLPYFA | TKLDLNSHEEIDAASSCKQLDLNGFSWN | |
| N08G149600.1 | 386 | 2.45e-29 | SPGYGLPYFA | TKLDLNSHEEIDAASSCKQLDLNGFSWN | * |

| | | | | | |
|---|---|---|---|---|---|
| K16G152200.1 | 388 | 3.71e-29 | SQDYRLANFD | VKLDLNSHDDNDASSHCQQFDLNGFSWN | C* |
| K02G070900.1 | 388 | 3.71e-29 | SQDYRLANFE | VKLDLNSHDDNDASSHCQQFDLNGFSWN | C* |
| F33381001 | 278 | 1.24e-28 | SHGYRLPCFG | AKLDLNAHDENDVTLSCKQFDLNGFSWN | * |
| U2430G0200 | 328 | 3.96e-28 | RDGFSMSCQT | AKLDLNINDTNDGPQHCKKFDLNGFSWA | * |
| G026448t1 | 451 | 5.77e-28 | DEVNRLPYFA | TKLDLNVHEENDAASSCKQFDLNGLSWN | * |
| M19G032700.2 | 388 | 6.96e-28 | SPGYRLSYYA | TKLDLNSHGEIDAASGCRQLDLNGFSWN | * |
| B008138 | 388 | 2.08e-27 | SKGFRLPHLT | PKLDLNSQEENDSTSTCKQFDLNGFSWT | |
| B108138 | 388 | 2.08e-27 | SKGFRLPHLT | PKLDLNSQEENDSTSTCKQFDLNGFSWT | |
| D10g083340.1 | 382 | 2.49e-27 | PQEYKLPYFA | PKLDLNTDDQTDAASNCKQLDLNGFSWN | * |
| A119.73 | 385 | 5.05e-27 | EVFDRLSNHT | AVLDLNAHDENDASSECKEFDLNGFSWS | * |
| AA17183 | 366 | 7.15e-27 | LNEFGMPCLT | PQLDLNTRDDNDAARNCNEFDLNGFSWN | |
| T3G199300.1 | 385 | 1.20e-26 | LLDFGHSCPG | KKLDLNTHNVDDTDQAYRHFDLNGFSWS | * |
| S09G178000.1 | 385 | 1.20e-26 | LVDFEHPCSG | KKLDLNTHNVDDTDQAYRHFDLNGFSWS | * |
| U0261G1090 | 441 | 1.42e-26 | LLDFEHPSSG | KKLDLNTHNIDDSDQGYRHFDLNGFSWS | * |
| PHL10_AT5G18240 | 375 | 2.35e-26 | EDCKLETHTR | TALDLNTHDENYGTTRPKQFDLNGFSWN | |
| U0468G0380 | 475 | 3.27e-26 | RDGFSMSCQA | TKLDLNINDTNDGPQHCKKFDLNGLSWI | * |
| Y6550973g003 | 360 | 4.55e-26 | PNEFGLSSMT | TQLDLNAHEENDGLPNSKQFDLNGFSWS | |
| V2g21880.1 | 377 | 6.30e-26 | YPSFERPNSG | KKLDLNTHNTDDSDQGFRHFDLNGFSWS | * |
| Y785961g002 | 384 | 7.41e-26 | LNEFALPCLT | SQLDLNSHDDNDPPPGCRQFDLNGFSWS | |
| W5g40960.1 | 384 | 3.08e-25 | LLDFDQQSSG | KNLDLNTHNIDDNDQGYRHFDLNGFSWS | * |
| R2G009060 | 332 | 3.60e-25 | RGSFSMTRKA | AKLDLNINDTTDGPQNCKKIDLNGFNWT | |
| EL01782.1 | 355 | 3.60e-25 | STELKLPFVS | TNLDLNTNDENNIASNCTKLDLNGFSWS | * |
| J3845853 | 365 | 6.65e-25 | EDCKVETRTS | TALDLNTHDETYGTTRPKQFDLNGFSWS | |
| W3g20900.1 | 327 | 9.01e-25 | RGGFSMSCQT | ANLDLNMNDTYDGPKHCKKFDLNGFSWA | * |
| R2G701218 | 387 | 1.90e-24 | LIDFENSCSG | KKLDLNTHNVDDTDQAYRHFDLNDFSWS | |
| R234155.1 | 366 | 2.96e-24 | LVDFEHPCSV | NKLDLNTHNVDDTNQAYRHFDLNGFSWG | |
| L6g032990.1 | 369 | 8.14e-24 | SQDYRLANFD | MKLDLNSHDDNGASSHSQKFDLNGFSWN | C* |
| X12613.1 | 327 | 1.43e-23 | SNDFGLACFG | AQLDLNAHDNNDSDSSNKEFDLNGLSWS | * |
| X01685.1 | 353 | 4.35e-23 | EARRNDPCLT | TQLDLNANYSSDAAPNCKKFDLNGISWT | * |
| AH181986g0010 | 423 | 1.71e-21 | VSFNKLSKLT | EELDLNANDDTSIASNCREFDLNGYGWT | YEH |
| Z9P05030_001 | 328 | 1.94e-21 | SDGFGLSCLT | TELELKIHEDDEGASGCKQFELNGFSWN | |
| J3828237 | 367 | 1.94e-21 | DHKLETQGTT | TELDLNTQVDNYCTTRPKQLDLNGFSWN | |
| Z6P32220_001 | 296 | 2.81e-21 | TDGFELPCLT | TDLELKIHEGSEGASSCKKFDLNGFSWN | |
| AA10677 | 348 | 1.64e-18 | PSECEQPSAT | TQLDLNACDEYNVSKNGNGNDLNGFSWS | |
| K07G229800.1 | 304 | 1.71e-16 | GNKLRKSEVS | EMLDLNCQYQRDIDSSVKEIDLNFSSSF | * |
| Q13g00280.1 | 315 | 2.08e-16 | KGFKLHIMPE | PQLDLNCRSQNDDVQTSKHFDLNGLFEL | ELLEK* |
| K20G035300.1 | 307 | 3.08e-16 | NKKLRKSEVS | QMLDLNSQYQRDIDSSVKEIDLNFSSSF | * |
| N05G101800.1 | 325 | 3.39e-16 | VKRSKSGDKL | RIFDLNSHYQNDFESGSKTLDLNCKGIE | QVNGQV* |
| Z2P01180_001 | 370 | 3.74e-16 | QSDSFGLAGH | TAQLDLNADEDDEGATNSKFDLNGFSWS | |
| L4g081710.1 | 312 | 1.57e-15 | GKKPRKSEFS | QMLDLNSKYERDIDSSSLEIDLNCSSSF | * |
| U1153G0050 | 509 | 3.01e-15 | GFSNMTLQTT | AELDLNINDGNCRPRNREKIDLNGSGRN | * |
| N01G035100.1 | 325 | 6.28e-15 | VKRSKIGDRL | RIFDLNSHCQNEFESGSKTIDLNCKGIE | QVNGQL* |
| L2g086450.1 | 362 | 9.04e-15 | IGEFQQRNTA | RLDLNSRGDNNEGATTCKQLDLNRFSWN | * |
| Z5P07670_001 | 265 | 1.30e-14 | QSNSFGMPGH | TAQLDLNADEDNEGDRDSKFDLNGFGWS | |
| B019826 | 388 | 2.89e-14 | DMSKGFRLPH | LTAKLDLNTQENDSTTHKQFDLNGFSWT | |
| Z8P21520_001 | 361 | 3.77e-14 | QSDRFGLLTH | TAQLDLNAHEDNEGGTSSRFDLNGFSWS | |
| B000838 | 323 | 6.36e-14 | GGQCTKLGLT | DKLDLNSQYQSEMESGCQEFDLNSRVEE | PIYVSKLFGA |
| AH120020g0010 | 348 | 2.30e-13 | SMLGNCRKVA | EGLDLNRKGEGSVPQQGRELDLNAYGWG | R |
| AF481.g073115 | 450 | 2.50e-13 | KLSTCNPKLG | KGLDLNMDSDGNKVGGIQEFDLNGYING | S* |
| EE00086.1 | 268 | 3.22e-13 | MHSSKRVQLL | EKIDLNSDTMNEFDQGRKVIDLNINGVE | FFNGNF* |
| EJ01031.1 | 387 | 2.69e-12 | KTKMPLFSTR | LDLNTGSENNAASSYKHHQLDLNGFSWS | * |
| C1G443600 | 359 | 4.32e-12 | ENKSKAFGLP | YLTSKLDLNVHDDNDTASQLDLNGFSWS | * |
| AD140s0042.1 | 461 | 7.47e-12 | RTHVKVEDSA | GSLDLNHGTPGLAMQRASDLDLNTYSWE | R* |
| S02G422000.1 | 325 | 9.42e-12 | GSSDEHGQQE | LDLSINGRSNPPKPRDSQRIDLNGSSWN | * |

PSR Motif 17

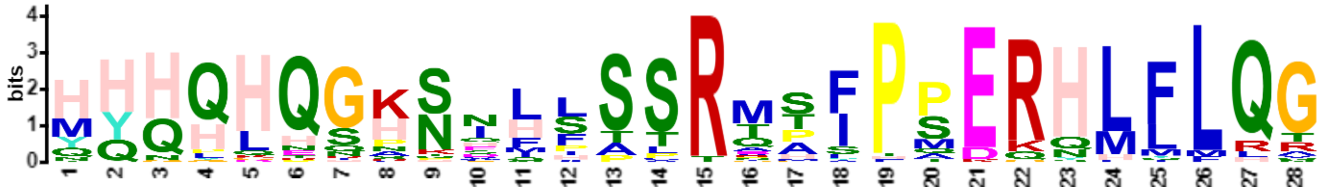


| Name | Start | *p*-value | Sites | | |
|---|---|---|---|---|---|
| G026448t1 | 68 | 6.42e-33 | SSFWRQKMYH | HHHQHQGKNIHPSSRMPIPPERHLFLQG | GNGPGDSGLV |
| M13G060200.1 | 3 | 1.45e-32 | MY | HHHQHQGKSIHSSSRMAIPPERHLFLQG | GNGPGDSGLV |
| O29950.m001149 | 3 | 4.98e-31 | MY | HHHQHQGKSVHSSSRMSIPPERHLFLQG | GNGPGDSGLV |
| N09G142300.1 | 3 | 9.42e-31 | MY | HHHQHQGKSIHSSSRMPIPPERHLLLQG | GNGPGDSGLV |
| K12G184700.1 | 3 | 4.28e-30 | MY | HHHQHQGKNIHSSSRMPIPSERHMFLQT | GNGSGDSGLV |
| K13G316600.1 | 3 | 1.35e-29 | MY | HHHQHQGKNIHSSSRMPIPSERHMFLQA | GNGSGDSGLV |
| L2g086450.1 | 2 | 7.86e-29 | M | YHHHHQGKNIHSSSRMSIPSERHMFLQT | GNGSSDSGLV |
| H785.2 | 3 | 1.02e-28 | MY | HHHQHPGKNMHLSSRMPIPPERHLFLQG | GNGPGDSGLV |
| Y785961g002 | 1 | 6.78e-28 | | MYHQHQGKNDLFSSRAAFPPERHLYLQG | GNAPGNSGLI |
| EJ01031.1 | 7 | 1.24e-27 | MYHHHH | HHHHHQGKNIQASTRMSIPSERHLFLQG | PTSSNNNGDQ |
| F33381001 | 3 | 5.10e-27 | MY | HHHHHQGKNIHPSSRTPITPERNLFLQG | GNGPGDSGLV |
| Y771701g004 | 3 | 5.72e-27 | MY | HHQHHQGPSNLLASRVSFPPERHLFLQG | GSVPGESGLV |
| M19G032700.2 | 3 | 8.06e-27 | MY | QHHQHQGKNIHSSSRNSIPPERHLFLQV | GNGPGDSGLV |
| AB07.828 | 95 | 1.77e-26 | RTASESGKMY | NHHHHQGSNNLVSSRTAFPSERHLFLQG | GTVQGESGLV |
| W5g40960.1 | 1 | 1.25e-25 | | MYHQHQGRSDLFTTRTSFPMERHLFLHG | GNTQGDSGLV |
| N08G149600.1 | 4 | 1.71e-25 | MYH | HHHQHRGKSVNSSSRMSIPPERYLFLQG | GNGPGDSGLV |
| X12613.1 | 1 | 3.19e-25 | | MYHQHQGHSDLFSPRAAFPQDRHLFLQG | GNVTGESGLV |
| K12G089100.1 | 3 | 3.19e-25 | MY | THQQHQGKNIHSSSRMPIPSERQMFLQT | GNGSGDSGLV |
| U0261G1090 | 54 | 4.81e-25 | QKVESNVKQK | MYHQHQGPSELFAARTTFPMERHLFLRG | GNTQGDSGLV |
| Z2P01180_001 | 4 | 5.89e-25 | MYH | HQHHHQGHNNILSCRTAFPAEKHLLLQG | GSIPEDSGLV |
| R2G009060 | 9 | 8.83e-25 | MSSCQMYH | HQQQLQSHSHFLSSRQTFPPERHMLLQG | GSIPAEPGLV |
| U0468G0380 | 151 | 1.32e-24 | VALSFHNMYH | HQQQLQSHSQFLSSRQTFPPERHLLLQG | GTIPAESGLV |
| T3G199300.1 | 1 | 2.90e-24 | | MYHQHQGPSELFTTRTSFPMEQHLFLRG | GNAQGDSGLV |
| S09G178000.1 | 1 | 2.90e-24 | | MYHQHQGPSELFTTRTSFPMEQHLFLRG | GNAQGDSGLV |
| S01G386700.1 | 4 | 3.53e-24 | MYH | HQQQLQSHSHFLSSRQTFPPERHMILQG | GSIPAESGLV |
| U2430G0200 | 4 | 1.02e-23 | MYH | HQQQLQSNSQLLSSRQTFPPERHLLLQG | GTIPAESGLV |
| B019826 | 3 | 1.23e-23 | MY | HHHHHHGKTSFPSSTMSIPPERHLFLQN | GNGRGDTGLI |
| R2G701218 | 1 | 1.79e-23 | | MYHQHQGLSELFTTRTSFPMEQHLFLRG | GNAQGDSGLV |
| T9G419000.1 | 4 | 1.97e-23 | MYH | HQQQLQSHSHFLSSRQTSPPERHLLLQG | GSIPAEPGLV |
| V1g63690.1 | 4 | 2.16e-23 | MYH | HQQQLQRHSQLLSSRQTFPSERHLLLQG | GIVPGESGLV |
| Z6P32220_001 | 3 | 4.13e-23 | MF | QHQPHQGHNNLLSPRTTFPSERQLFLQR | GNTTGEPGLV |
| Y6550973g003 | 2 | 5.96e-23 | M | YHHHLEGHSNLLASRTTFPPKKHLFLQG | GSVPGESGLV |
| W3g20900.1 | 4 | 2.51e-22 | MYH | HQQQLQSHNQLLPSRQSFPSERHLLMQG | GSVSGESGLV |
| Z9P05030_001 | 3 | 1.34e-20 | MY | QHQPHQSHNNLISSRIAAPSERQLFLQR | GSTTGESGLV |
| Z8P21520_001 | 2 | 9.84e-20 | M | YHHYHQSHSNIFSSRETFPADGQLLLQR | GGSAPEESGL |
| J3828237 | 3 | 1.69e-19 | MY | YQNQHQGKSILSSSRMHLPSERHHQFLR | GGNSLGGSGL |
| K16G152200.1 | 2 | 7.74e-19 | M | YYQQQQQAKNMHALRMHSPTERHMMLQG | GNGAGDPGLV |
| K11G183400.1 | 7 | 2.18e-18 | MYSTHQ | QHQGKNIHSSSSSSRMPIPSERHMFLQT | GNGSGDSGLV |
| D10g083340.1 | 1 | 2.72e-18 | | MYHHHHQAPNMHPSTRMSFPERHLFLQG | GNANGDSGLV |
| K02G070900.1 | 2 | 3.63e-18 | M | YYQQQQQAKNMHALRMHSPTERHMMMQG | GNGSGDSGLV |
| X01685.1 | 16 | 5.21e-18 | ANQPRQKMYN | NHHQGHNNNLLIPSRGAFPADKHLFLQG | GSVQGESGLV |
| C1G443600 | 3 | 2.48e-17 | MY | HHNHHNGKNSFPSTRMSIPQERSHMFLQ | GNGNACGGDA |
| P6G0056100 | 1 | 3.50e-17 | | MYGHHEGKHGILTSRAAFHQDRNLLLQG | ANAPGDAGLV |
| AA17183 | 1 | 4.31e-17 | | MYHLDHNSSNLISLRESYPSERNLFLQA | GNGPVDSGLV |
| R155434.2 | 4 | 1.00e-13 | MYH | QQQLHTHNQHLSSSRPGLPPEKQFLLHG | AGGGGGGGDA |

PSR Motif 18

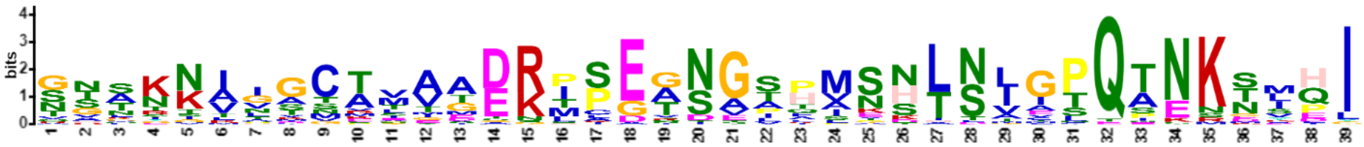

| Name | Start | *p*-value | Sites | | |
|---|---|---|---|---|---|
| S01G386700.1 | 112 | 2.86e-35 | SKNIHAQANG | GNAKNVVGCAMAMEKPPEGNGSPASHLNLGTQTNKSVHI | GEALQMQIEV |
| R2G009060 | 117 | 6.98e-34 | SKNIHAQANG | VNAKNVIGCTMAMDKPLEGNGSPASHLNLGTQTNKSVHI | GEALQMQIEV |
| W3g20900.1 | 112 | 5.28e-32 | SKNLHAQANA | GNVKNALVCTTATEKPSEANGSPVSHLNLGTQTNKSVHI | GEALQMQIEV |
| U0468G0380 | 259 | 8.97e-32 | SKNLHAQANV | GNARNAVDCAITAEKPSEGNGSPVSHLNLGTQTNKSMHI | GEALQMQIEV |
| U2430G0200 | 112 | 4.24e-31 | SKNLHAQANV | GNARNVVACTIATDKPSEGIGPPVSHLNLGTQTNKSMPI | GEALQMQIEV |
| T9G419000.1 | 112 | 7.04e-31 | SKNVHAQANG | GNAKNMVGCTMAMEKPPEGNSSPASHINLGTQTNKSVHI | GEALQMQIEV |
| H785.2 | 110 | 7.04e-31 | LSKNLHGQTN | SGTNKLGAVVMAGEKMPEANGTHMNNLSIGPQTNKSLHI | NETLQMQIEV |
| X01685.1 | 124 | 1.04e-29 | SKNLQTQSSN | GSTKNIVSCTVATDRTLEGNGAPISDLNIASQTKKTMQI | SEALQMQIEV |
| S09G178000.1 | 109 | 1.04e-29 | SKNLQAQANV | STSKNAIGCTSVADRIPGTSAATMSSTNVVPQAEKTIQI | GEALQMQIEV |
| R2G701218 | 109 | 4.80e-29 | SKNLQAQANV | STSKNAIGCTSIADRIPGTSAATMSSTNVVPQAEKTIQI | GEALQMQIEV |
| G026448t1 | 175 | 4.80e-29 | LSKNLHGQAN | NGSNKIGAVAMAGDRMSEANGTHVNNLSIGPQANNGLQI | GEALQMQIEV |
| T3G199300.1 | 109 | 1.07e-28 | SKNLQAQANV | STSKNAIGCTNIADRMPGTSAPTMSSTNAIPQAEKTIQI | GEALQMQIEV |
| U0261G1090 | 162 | 5.14e-28 | SKNLQAQANV | GTTKNTIGCEIVADRIPGISAPAMSNTNVIPQEEKTIQI | GEALQMQIEV |
| O29950.m001149 | 110 | 5.14e-28 | LSKNLHGQAN | SGSNKIGTGAVVGDRISETNVTHINNLSMGTQTNKGLHI | GEALQMQIEV |
| Z2P01180_001 | 110 | 5.74e-28 | RLSKNLQAQA | NGAESVIGCKLAAERTSEGNGSRASNTNIIPQSNKTYPI | NEALQMQIEV |
| R234155.1 | 90 | 7.98e-28 | SKNLQAQANA | STSKNAIGCTPVADRIPGTTAATMSSTNVLPQAEKTIQI | GEALQMQIQV |
| N09G142300.1 | 110 | 1.71e-27 | LSKNLHGQAV | SRSSKIGANAVAVDRMSEANVTHLNNLSIGTQTNKNLHI | SEALQMQIEV |
| T2G436900.1 | 107 | 7.55e-27 | SKNLQAQANA | TNAKNVLGCRTGTDKPCERNGSPASHLNTEPQINRSMHI | SEALQMQIEV |
| K13G316600.1 | 110 | 1.03e-26 | LSKNLHGQSN | NVTHKITTSATTGERLSETNGTHMNKLSLGPQANKDLHI | SEALQMQIEV |
| N08G149600.1 | 111 | 1.41e-26 | LSKNLHGQAI | SGSSKIGATTVESDRMSEANVIHINNLSIGSQTNKSLHI | GEALQMQIEV |
| K12G184700.1 | 110 | 2.61e-26 | LSKNLHGQSN | NVTYKITTSASTGERLSETNGTHMNKLSLGPQANKDLHI | SEALQMQIEV |
| Z8P21520_001 | 111 | 1.07e-25 | SKNLQAQANS | GSTKSVIGCTLAAERTIDVNGSFMVNTSTMPQSNKSFQI | NEALQMQIEV |
| Y771701g004 | 109 | 1.07e-25 | RLSKSLQAQP | NTGTTKNGCPLAADRTAEGNGSLMSNTTVASQTNKTMQI | SEALQMQIEV |
| W5g40960.1 | 109 | 3.17e-25 | SKNLQGQANV | GTTKNALGCTGVADRIPGTSALAMASASAIPQAEKTIQI | GEALQMQIEV |
| K02G070900.1 | 109 | 3.44e-24 | ISKNMHGQTN | TSNNKIASTMEAAARISEASGVQMKHLSIGLQTNKNSEI | NDALQMQIEV |
| S02G422000.1 | 108 | 8.69e-24 | SKNLQAQANA | VNAKNALSCRTGTDNPCEGSGSPPPHLNLEPQINRSMHI | SEALQMQIEV |
| Z5P07670_001 | 51 | 1.04e-23 | SKNLQAQASS | GSAKIATGCKLVAGRTAEGNGLLLGSTNIIPQSNKNIPI | NEALQMQIEV |
| X12613.1 | 109 | 4.82e-23 | SKNLQAQANI | ATTKNVIGCVAAADRTPGQSGLIVSNTSATCEAEKTMQI | SEALQMQIEV |
| B008138 | 94 | 5.27e-23 | KYRLSKNLHG | QANTGINKNAVAGDRISEASGAFISNTSLCSQTNKEYEL | KRISNIRSLQ |
| B108138 | 94 | 5.27e-23 | KYRLSKNLHG | QANTGINKNAVAGDRISEASGAFISNTSLCSQTNKEYEL | KRISNIRSLQ |
| V2g21880.1 | 106 | 7.50e-23 | RLSKNLQAQV | NVGTTKNGCAVVADSMPATSTPAMTNTNVIPQAEKTIQI | GEALQMQIEV |
| PHL9_AT3G04030 | 109 | 8.20e-23 | LSKNLNGQAN | NSFNKIGIMTMMEEKTPDADEIQSENLSIGPQPNKNSPI | GEALQMQIEV |
| K16G152200.1 | 109 | 9.77e-23 | ISKNMHGQTN | TSNNKIASTMEAATGISEASGVQMKHLSIGLQTNKNSEI | NDALQMQIEV |
| L6g032990.1 | 107 | 2.33e-22 | SRSMNGQTNT | GSSKIAPTSEVVTSRMSESSGIHMKDLNIGLQTNKNSDI | NEALNMQIEV |
| J3828237 | 111 | 3.52e-20 | LSKNLNGQAN | SGLNKIGMMTMMEEKSPDADEIQSETLSIGPQPNKNSPI | SEALHMQIEV |
| L2g086450.1 | 112 | 5.21e-20 | NLHGQSSSNV | THKINTHATSVSDERLSETNGTHMNKLTLGPQTNNNKDL | HISEALQMQI |
| W7g48596.1 | 118 | 8.30e-20 | KNLQSQANAS | RAQGVLGCSTTEIDKPCEGNGSPASHLDLETQTNSSMHI | NEALQMQIEV |
| PHL10_AT5G18240 | 109 | 3.71e-18 | LSKNLNGQAN | SSLNKTSVMTMVEENPPEVDESHSESLSIGPQPSMNLPI | SDALQMQIEV |
| K12G089100.1 | 112 | 1.33e-17 | KSLHGQSNNM | THKITINSGAATDERLRENNGTHMNSLNLAPQSNNKDLY | ISEALHMQIE |
| V1g63690.1 | 112 | 1.37e-16 | SKNLHAQANV | GNSRNVVGCTMATEKHSEGNGSPVSHHLGAQTNKSMHIG | EALQMQIEVQ |

PSR Motif 19

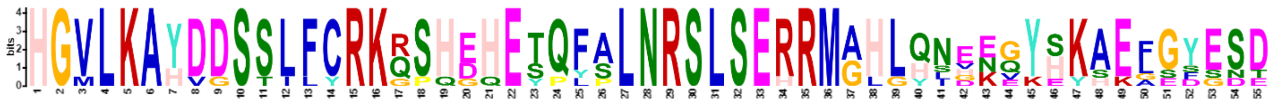

| Name | Start | *p*-value | Sites | | |
|---|---|---|---|---|---|
| S09G178000.1 | 269 | 1.40e-68 | GFIKEHRLQH | HGVLKAYDDSSLFCRKRSHEHETQFALNRSLSERRMAHLQNEEGYSKAEFGYESD | TEMAHEYAEP |
| R234155.1 | 250 | 1.34e-66 | GFIKEHKLQH | HGVLKAYDDSSLFCRKRSHEHETQFALNRSLSEHRMAHLQNEEGYHKAEFGYESD | TEMAHEYTAP |
| R2G701218 | 272 | 5.25e-62 | KEQRLQHQQH | HGVLKAHDGSSLFCRKRPHEHETQFALNRSLSERRMAHLQNDEQYSKAEFGYESD | TEIVHEYTAP |
| T3G199300.1 | 269 | 2.01e-57 | GFIKEHRLHH | HGVLKAHDDSSLFCRKGSHEHETPLALNRSLSERRMAHLQIENGYSKAELGYEND | TEMAPEYIGP |
| V2g21880.1 | 266 | 1.20e-53 | GYINEHRLHS | HGVLKAYDDSSILYRKQSHGHEYQFPLNRSLSERRMGHLHNVKEYHKAELGSESD | TEIQQEYITP |
| W5g40960.1 | 268 | 2.77e-46 | EGFKEHRLQN | HGVLKAYDDSTLFCRKQSQDQESQYSLNRSLSERRMGHLYSGKQYHKSEGSDSDT | EVLHEYITPQ |
| U0261G1090 | 322 | 8.82e-44 | GFIKEHRLQN | HGMLKAYVDSSLFCRKQSHDHESQFSLNRSLSERRMGLGHLHNVKEYHKAEFGSE | SDTEIPHEYI |

PSR Motif 20

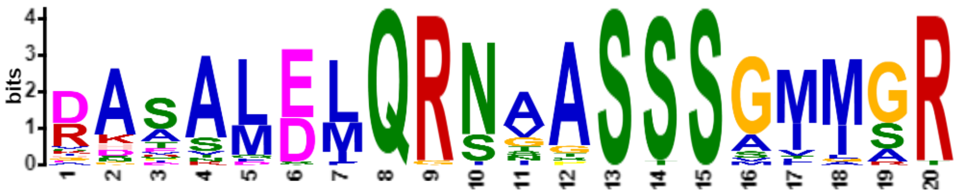


| Name | Start | *p*-value | Sites | | |
|---|---|---|---|---|---|
| Y1141531g006 | 92 | 2.69e-23 | HKEINDHSIK | DASALDMQRNAASSSGIMGR | TMNENVHITE |
| AB06.154 | 101 | 8.42e-22 | HKEFNDHAVK | DASALELQRNAASSSGFMGR | AINDRNVHLS |
| R2G052544 | 115 | 9.77e-22 | KEFNEHSVKD | AAAAMEMQRNAASSSGMMGR | SMNDRSVHMN |
| W2g07770.1 | 85 | 2.98e-21 | HKEFSEHSVK | EAAAMEMQRNAASSSGIMGR | SMNHDRNVND |
| Y806121g001 | 93 | 5.00e-21 | HKELNDHSVK | DASALEMQRNGASSSGIMSR | TMNENVHITE |
| M10G174100.1 | 101 | 7.33e-21 | HKDFNDHSIK | DASALDLQRSAASSSGMMSR | SMNEMQMEVQ |
| J3853188 | 106 | 7.33e-21 | YGDHSTKEGS | RASAMDIQRNVASSSGMMSR | NMNEMQMEVQ |
| PHL14_AT1G79430 | 106 | 7.33e-21 | YGDHSTKEGS | RASAMDIQRNVASSSGMMSR | NMNEMQMEVQ |
| M08G081800.1 | 100 | 2.50e-20 | HKEFNDHSIK | DASALDLQRSAASSSGMISR | SMNDNSHMIY |
| J3830485 | 106 | 2.50e-20 | YGDHSTKEGS | RASAMDIQRNVASSSGMISR | NMNEMQMEVQ |
| V1g31837.1 | 111 | 4.49e-20 | QHKEFGDHSS | VKEAMEMQRNAASSSGMMGR | SMNDRSAHMN |
| L6g444980.1 | 107 | 5.04e-20 | FNDHSIKDGM | RASALELQRNTASSSAMIGR | NMNEMQIEVQ |
| K08G163500.1 | 107 | 5.04e-20 | FNDHSIKDGM | RASALELQRNTASSSAMIGR | NMNEMQIEVQ |
| F36717001 | 101 | 7.10e-20 | HKEFNDHSIK | DASALELQRNIASSSGVMSR | NTNEMQMEVQ |
| B009935 | 101 | 4.03e-19 | KEFNDHAVKD | GDRALELQRNAASSSGIMGR | SMNENVRLTE |
| D12g017370.1 | 104 | 4.48e-19 | FNDHSVKDGD | RATSLELQRNSASSSGMIGR | NMNEMQMEVQ |
| K15G215000.1 | 107 | 7.50e-19 | FNDHSIKDGM | RASALELQRNIGSSSAMIGR | NMNEMQMEVQ |
| R2G081671 | 101 | 1.52e-18 | RLGKQHKELG | DHTAMEMQRSVASSSGMIAR | SMNDRSVNVN |
| A094.1 | 104 | 2.25e-18 | FNDHSSKESE | RAAVLELQRSGASSSGIMGR | DGNDNMQVRE |
| EM01502.1 | 104 | 1.24e-17 | HKDFHDHSMK | DDSSLELQRNNASSSGMLGR | SMNEMQMEVN |
| Z3P23640_001 | 91 | 1.48e-17 | PHKEFNDQSA | KNAALELQRNSASSSAIMGR | NMNDRNMHMN |
| P5G0044100 | 105 | 1.48e-17 | DHHSTKDATG | RAVSLDFQGNAASSSGMMGR | ATNEMQMEVQ |
| X08334.1 | 101 | 1.63e-17 | HKEFAEHSIK | DAATQELQRNAASSSSIMGR | NTNRNLHTNN |
| S10G220000.1 | 106 | 1.95e-17 | RLGKQHKEFG | DHTAMEMQRSVASSSGVIAR | SMNDRSVNVN |
| O30017.m000317 | 92 | 6.15e-17 | HKEFSDPSIK | DGPALDLQRSAASTSAMMGR | SMNEMQMEVQ |
| C1G159000 | 98 | 1.12e-16 | QPHKEFNDHS | IKEALDLQRNASSSSGLMGR | SMNDRNVHIS |
| N03G077700.1 | 38 | 2.82e-16 | HKEFNDHSIK | DAQALDLQRSGGSSSVMMSR | SMNDNSHMVD |
| X07475.1 | 38 | 3.91e-16 | HKEFNDHSVK | DAANMSMQRNAPSSSSIMGR | SMNENVHITE |
| EH01655.1 | 111 | 4.61e-16 | NDQYSMKDVE | RASSLELQRNNTSSSGIAAR | NMNEMQIEVN |
| AA17379 | 98 | 8.09e-16 | KQPHKDFNDH | SKDALDMQRIGASSSGTMGR | TMTDRNAQTA |
| AB08.2515 | 100 | 1.03e-15 | HKEFSDHSVK | DASAFELQRNAASSSMMGRT | MNDRNVHLSE |
| G033672t1 | 97 | 1.64e-15 | GKQPHKEFND | HSIKDDLQRNAASSSGMIAR | SMNEMQMEVQ |

PSR Motif 21

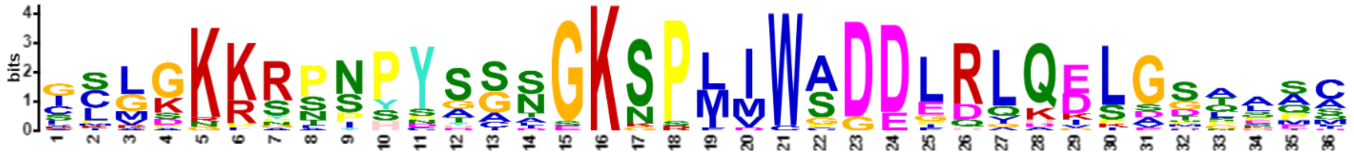

| Name | Start | *p*-value | Sites | | |
|---|---|---|---|---|---|
| O30017.m000317 | 223 | 1.37e-36 | LDGFMPNNDH | ICLGKKRSSPYSGSGKSPLIWSDDLRLQDLGSAPTC | LGPPDDLFKS |
| M08G081800.1 | 242 | 1.98e-36 | DGFMSNNHDD | ICLGKKRTNPYAGSGKSPLIWSDDLRLQDLGSGLSC | LGPQDDPLKG |
| Y1141531g006 | 233 | 2.01e-35 | LDGFFQANEG | VCLGKKRFNPYSSNGKSPLICADDLRLQELGSAAAC | MGAQEEPSKN |
| M10G174100.1 | 233 | 2.01e-35 | LDGFMPNNDN | ICLGKKRPSPYDGSGKSPLIWPDDLRLQDLGSGPAC | LEPQDDPFKG |
| G033672t1 | 228 | 7.83e-35 | LDAFMPNNDN | ICLGKKRASPYSGSGKSPLIWSDELRLQDLGTAASC | LGPQDDPFKS |
| Z3P23640_001 | 221 | 1.20e-32 | PLDGLFPTDD | SIMSKKRSNPYHSNGKGPLIWADDLRLQELGSTAAC | IVSQEEPPPS |
| K15G263700.1 | 257 | 3.37e-32 | HGFMPINETL | CLGKKRSNNPYSGSGKSPLIWSDDLRLQDLGGPASS | CLGPQDDPFK |
| K08G163500.1 | 248 | 6.00e-32 | HGFMPINESL | CLGKKRSNNPYSGSGKNPLIWSDDLRLQDLGGPASS | CLGPQDDPFK |
| X07475.1 | 183 | 7.98e-32 | DMFFPTNESI | AMGSKKRPYPYGSNGKSPMIWADDLRLQELGSQEEA | SKCDQLQIAP |
| AB08.2515 | 222 | 9.20e-32 | LDGFFPNDQT | LSLGKKRPNPYSSNEKSPIVWADDLRLQELGSAEPS | KSDQVLQLDS |
| B009935 | 248 | 2.25e-29 | FMPAAANDNI | CLGKKRPSPYSSSSGKNPLIWADDLRLQELGGTAAA | CLGPQEDAFK |
| C1G159000 | 240 | 3.03e-28 | LDGFSPINDN | ICLGRRRPNSYGSTSKNPIIWSDDIRLQELGMSVPC | LGSQDDSFKN |
| AB06.154 | 232 | 3.22e-27 | IDDFFPTNET | LCLGKKRSNPYGSNGKSPMDWADDLHIQELEPSKSQ | IDMEGEKKFE |
| L6g444980.1 | 227 | 1.16e-25 | MQINENLCLG | KKRPNPNTNPYSGNGKNPLMWSDDLRLQDLGTASSC | LDDPFKGDQI |
| H32.94 | 244 | 4.57e-24 | HGFMSNDDTL | CLGKKRSNPYSTCSGKSPLIWADDPRLQEPSCLGPQ | DHDLFKHSAH |
| D12g017370.1 | 253 | 1.25e-23 | LSNTSTDHNL | SLGKKRASPYNTSNGKSPFMWSDDFRLHELGSNNED | DHQIIQMERS |
| U0123G0060 | 248 | 1.69e-23 | SFFACNESSI | GSMGKKRPSPYAAAGKSPMMWGDDGDQAKSDHLQMA | PPMMDGGIDV |
| Z11P14430_001 | 217 | 2.28e-23 | DQLDLQSQVD | GPLDGFYPISDSILGKKRMIWADDLRLQELGSTSAC | VGPKEEPCCK |
| U0095G1450 | 223 | 2.52e-23 | SFFACNESSI | GSMGKKRPSPYAAAGKSPMMWGDDGDQDKSDQLQMA | PPMMDGSIDV |
| U2141G0040 | 626 | 1.09e-22 | SFLAFNESCT | GSVGKKSPNHYSSTGKSPMVWAGEEQAKISADQLQM | ASSMMMET* |
| U3413G0110 | 241 | 1.06e-21 | SFLAFNESCI | GSVGKKSPSHYSCIGKSPIVWAGEEQVKISADQLQI | APSMMMEAGD |
| S04G058700.1 | 263 | 1.36e-20 | MEAFFASCDI | GSLAKKRPISPYADGKSPMMWCDDEDGKGIVDQLQM | APSMMDAAGI |
| T4G185800.1 | 243 | 1.33e-19 | LTLGENFIGS | SSADKKGPNHCSSSGKSSMIWAGEEDQQVKSGTDQL | QMGSSTTMEG |

PSR Motif 22

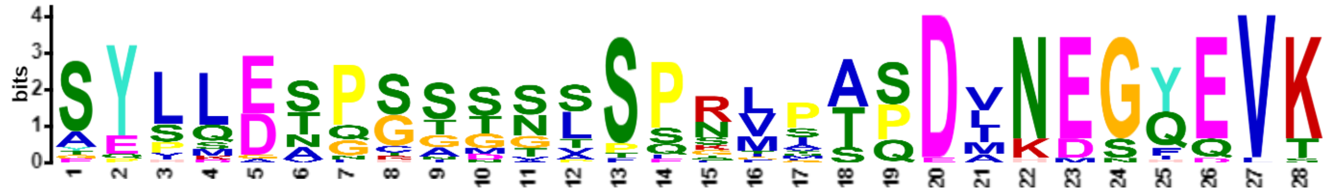

| Name | Start | *p*-value | Sites | | |
|---|---|---|---|---|---|
| Z1P16440_001 | 100 | 1.59e-29 | KEMTEQSKDA | SYLLENPGSSSLSPRVPTPDVNEGQEVK | EALRAQMEVQ |
| Z3P11980_001 | 100 | 4.58e-28 | KEMTEQSKDA | SYLLENPSSSVLSPRVPTPDVNEGQEVK | EALRAQMEVE |
| F07064001 | 111 | 1.59e-27 | GEAPKDGISA | SYLSESPGTSNSSPNLPTSDINEGYEVK | EALRVQMEVQ |
| Y674801g004 | 99 | 2.88e-27 | KEMTDQSKDA | SYPLENPSSSGLSPRLPTPDVNEGQEVK | EALRAQMEVQ |
| O29805.m001543 | 117 | 8.60e-26 | GEASKDGLSG | SYLLESPGAGSSSPNIVTSDMNEGYEVK | EALRVQMEVQ |
| M116G001200.1 | 116 | 2.32e-25 | SDTFKDGLSG | SYLLENPCTGNSSLNMTASDVNEGYEVK | EALRAQMEVQ |
| G030479t1 | 116 | 2.96e-25 | GEGPKDGMSA | SYLLESPGTNNSTPSLPSSDMNEGYEVK | EALRAQMEVQ |
| H127.186 | 188 | 4.77e-25 | GEAPKDGMSA | SYLLESPGISNSSPNLPSSDVHEGYEVK | EALRVQMEVQ |
| K10G196600.1 | 189 | 3.00e-24 | DEGLKDGMSA | SYLQESPGTDNSSPKLPASDANEGHEVK | EALRAQMEVQ |
| B0105317 | 106 | 7.22e-24 | KDLTEPSKDA | SYLLESPHNGTSSPGMSASDLNEGYEVK | EALRAQMEVQ |
| EM01478.1 | 110 | 9.97e-24 | GKEFEASKDG | GYLLDSPGASNSSQNLSASDINEGYEVK | EALRAQMEVQ |
| AB05.877 | 94 | 2.09e-23 | KEISEQSKDA | SYLLETPTSSALSPSVGTPDANEGQEVK | EALRAQMEVQ |
| Y1714681g012 | 136 | 4.78e-23 | KEMTDQSKDA | SYPSETLSSSGLSPGVPAPDVNEGQEVK | EALRAQMEVQ |
| T2G168800.1 | 204 | 2.08e-21 | KEGSEQSKDA | SYLLDAQSGMSVSPRVAAQDVKESQEVK | EALRAQMEVQ |

| | | | | | |
|---|---|---|---|---|---|
| R2G064197 | 107 | 3.00e-21 | KEGSEQSKDA | SYLLDAQSGMSVSPRVAAQDMKESQEVK | EALRAQMEVQ |
| D04g015290.2 | 115 | 3.00e-21 | DEASKDGLTA | TYSLESPCSGGTPQQLPASDLNEGFEVK | EALRAQMEVQ |
| R2G113742 | 117 | 1.25e-20 | KELTEQSKDA | SYLMEAQSGTTLSPRGSTPDVKESQEVK | EALRAQMEVQ |
| N15G053300.1 | 118 | 2.74e-20 | SKDVGIAASV | AESQDTGSSTSTSSRMIAQDINEGYQVT | EALRVQMEVQ |
| AA126160 | 94 | 3.86e-20 | LGKQSGKEMT | EQPKDVPGSSSLSPRMPTPDINENQEVK | EALRAQMEVQ |
| N03G148100.1 | 117 | 4.26e-19 | SKDVGIAASV | AESQDTGSSTSTSSRLIAQDLNDGYQVT | EALRVQMEVQ |
| M01G314800.1 | 114 | 4.62e-19 | SKDVGIAASV | AESQDTGSSTSASSRMIAQDLNDGYQVT | EALRVQMEVQ |
| EH01640.1 | 36 | 1.02e-18 | KEFGEASKDG | SYHLDSPRASTPPQNLLSSDMNMGYEVK | EALRAQMEVQ |
| W8g25799.1 | 104 | 1.29e-18 | KEMAEQSKDA | SYILGAQSGTNLSPTVPTPDLKESQELK | EALRAQMEVQ |
| EG00679.1 | 42 | 1.02e-17 | TENSKDASCI | AESQDTGSSTSASSKTMAQDINDGYQVT | EALRAQMEVQ |
| Q305g00050.1 | 103 | 6.83e-17 | ELGDQSKDSS | YPLDATNSSGSLSPLVQTPDANDGIEVK | EALRAQMEMQ |
| C1G374700 | 34 | 2.27e-15 | SEKTKTAHSA | SYVMDSPGTDGSFESFSSSELNEGFDVS | QALRKEIEGQ |

### PSR Motif 23

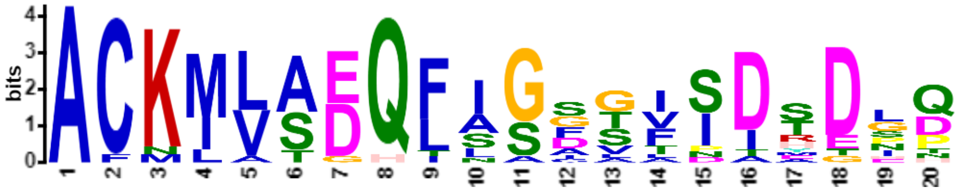

| Name | Start | *p*-value | Sites | | |
|---|---|---|---|---|---|
| L1g090670.1 | 181 | 8.92e-23 | ERRYMAMLER | ACKMLADQFIGDTTIDTDIQ | KFQELPSTEL |
| K10G196600.1 | 256 | 1.14e-22 | ERRYMGMLER | ACKMLADQFIGDVIIDRDGQ | KFQGLENKTS |
| K15G123000.1 | 179 | 3.60e-22 | ERRYMAMLER | ACKMLADQFIGATVIDTDSQ | KFQGIGSKAP |
| K20G193600.1 | 262 | 3.92e-21 | ERRYMGMLER | ACKMLADQFIGDVTIDMDGQ | KFQGLESKTS |
| O29805.m001543 | 184 | 5.62e-21 | EKRYLAMLER | ACKMLADQFLGGTVIDSDIQ | KDSGSKKKRS |
| K09G017300.1 | 180 | 5.62e-21 | ERRYMAMLER | ACKMLADQFISATVIDTDSQ | KFQGIGSKAP |
| U1931G0070 | 135 | 3.74e-18 | QKYIDTLLEK | ACKIVSEQLSGFSISDHDLP | DLASAGVICD |
| T2G168800.1 | 272 | 4.76e-18 | QKYIDSILES | ACKMVTEQFASSGFSISDPD | LPEISPGGVM |
| S02G161700.1 | 104 | 4.76e-18 | QKYIDSILGS | ACKMVTEQFASSGFSISDPD | LPEISPGGIM |
| W9g12750.1 | 171 | 5.77e-17 | QKYIDTLLEK | ACKIVSEQLASSGFSISDND | LPELSGGVMC |
| H127.186 | 255 | 8.84e-17 | ERRYLTMLER | ACKMLADQFLGGALSDSENQ | KCVGVAGKMA |
| W8g25799.1 | 172 | 9.98e-16 | QNYIDTLLEK | ACNIVSEQLNGFSISDHDLT | SAGVMLSSSD |
| V3g22297.1 | 174 | 1.61e-15 | QTYIDSLLEK | ACMLVSEQLSGFSISDYDLP | DLASAGFQIP |
| T6G120200.1 | 185 | 1.11e-14 | QKYIDAILDK | AFKIVSEQLSGISISDRDLP | DLASAGVMFS |
| G030479t1 | 183 | 7.44e-14 | ERRYMAMLER | ACKMLADHFIGGGADTETEN | LGFSSKVPRN |
| B0105317 | 174 | 4.48e-13 | RRYVESMLER | ACKLLAGQTISCLVPDAEGH | ELPGSAPKAS |
| Y1714681g012 | 204 | 7.78e-13 | HKYIDSLLEK | ACKIASEQIAASGFNATGHD | LPDLATGIIC |

### PSR Motif 24

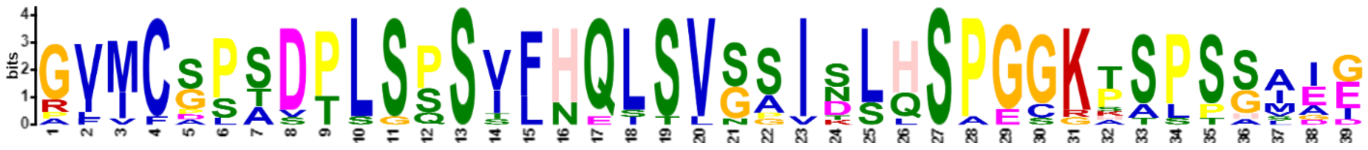

| Name | Start | *p*-value | Sites | | |
|---|---|---|---|---|---|
| Y1714681g012 | 230 | 9.98e-43 | TGHDLPDLAT | GIICSPSDPLSPSVFHQLSVGAISLHSPGGKTSPSSAIE | GYLFYHQKAP |
| Y674801g004 | 193 | 5.87e-42 | TGHDLPDLAT | GVMCSPSDPLSPSVFHQLSLGAISLHSPGGKTSPSSAIE | GQSFYPKAPE |
| T2G168800.1 | 299 | 1.56e-39 | DPDLPEISPG | GVMCGPTDTLSSSVFNQLSVSSIDSHSPGGKPSPSGIEG | PPMLLQKSPE |

| | | | | | |
|---|---|---|---|---|---|
| X17080.1 | 216 | 9.01e-39 | EHDLHDLATR | GVMCAPSDPLSPSIFHQLSVGSISLHSPGGRTAPTSAID | GQFFFQKPPE |
| S02G161700.1 | 131 | 1.65e-38 | DPDLPEISPG | GIMCGPTDTLSSSVFNQLSVSSIDSHSPGGKPSPSGMEG | PPLLLQKSPE |
| U1931G0070 | 160 | 2.73e-37 | DHDLPDLASA | GVICDPADPLSPSVFHQLSVSSISLQSPGGKASPSALDG | QLFFQKPPEL |
| W9g12750.1 | 197 | 1.17e-36 | SDNDLPELSG | GVMCGSADTLSSSIFHQLSVSPINLHSPEGKPTPSGIEG | QMILQKSPEL |
| R2G064197 | 203 | 1.38e-35 | NPDLPEISPG | GVMCGSTDTLGSSVLNQLSVSSIDSHSPGGKPSPSGMEG | PTLLQKSPEL |
| Z3P11980_001 | 194 | 3.51e-35 | TEQELSDMAT | RVICSPSDPLSQSILHQLSVNSINLQSPGCKTSSSSAIE | GQFFYQKPHE |
| Z1P16440_001 | 194 | 7.90e-33 | AEHELPDMTP | RVVCPPSSPLSPSILHQLSVSSINLQSPGCKTSPSSSAI | EGQLFCQKPP |
| T6G120200.1 | 210 | 8.02e-29 | DRDLPDLASA | GVMFSPADPLSPSVFHQLSVSAVSLLSPGGGKALPHVAI | DISQKPPELK |
| Z9P19070_001 | 195 | 2.32e-28 | SVGDLPNLAN | AFICSLSDPSSQSTFHESSVGSIKLHSAGGKHSPPSAIE | CQFNQQEPLK |
| AB101.3516 | 186 | 3.29e-25 | EQISSSGLNS | PLMCSSSVPLSPSSLHQITVGGITLQSPESKSSLSSIGE | GQFFYQKSPE |

HRS Motif 1

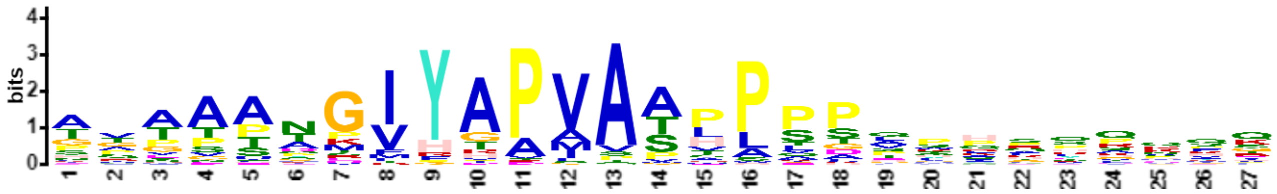


| Name | Start | *p*-value | Sites | | |
|---|---|---|---|---|---|
| N_06G071500.1 | 334 | 2.59e-20 | TTASGETSTL | AAAAANGIYAPVAAPPPTSHQMLQKQE | HTQSEQMQSE |
| K_01G086700.1 | 316 | 3.64e-20 | ASGEELTTVT | TTTAPTGIYAPVAAHPPAVTHTLPIMK | QKEHSHSEER |
| J_3871943 | 282 | 4.18e-19 | ATTSGTTTTT | TTTTTTGIYGAMAAPPPPQWPSHFNFT | PSIIVEEGSG |
| K_10G204200.1 | 311 | 9.03e-19 | ATSTASREVA | TVAAPAGIYAPVATHPIPVSHPPADSI | KNPQFKKVQL |
| M_010G128900.1 | 325 | 1.22e-18 | AATTTAGETS | TISAANGIYAPIAAPPPAVPQNRQHKQ | SEHSQSEGRG |
| F_1011942001 | 289 | 2.58e-18 | AATTSSGEAT | GVTTANGIYAPVASVPPSHPQGSTQRQ | QPMKPKKSQS |
| E_E00311.1 | 316 | 2.99e-18 | ATTTSGEAAS | GVANSTGIYAPVASLPPPFHHASKQRQ | HISDNDRGCS |
| Z_3P21090_001 | 284 | 7.11e-18 | VAAAAQPGNG | ACAPTNGIYAPVASLPSDPRFQQQLQT | KKQYQRSCSG |
| K_02G098800.1 | 311 | 1.44e-17 | SASGGELTTA | TPAAPTGIYAPVAAHLPAGTHTSPIMK | HSHSPSEEKN |
| O_28883.m000751 | 334 | 1.90e-17 | AATTASMETV | TTAAANGIYAPVAAPLGTIPKQQRAQS | QHLQSERRGS |
| M_008G117500.1 | 321 | 3.76e-17 | AATTASAETS | PISAANGIYAPIAAAPPTVPQKPHHVQ | YEPLQSEGGG |
| B_012707 | 294 | 4.30e-17 | AAVTATKREA | NGVAANGIYAPVAPQPVPESCSQQQKK | QKQSPQSSLQ |
| N_14G098400.1 | 317 | 5.62e-17 | TASGETSTIA | AAAAANGIYTPVAAPLSTAPPKLQKQL | LKHSEQMQLE |
| K_20G186500.1 | 309 | 6.43e-17 | ATSTASREVA | TVAAPARIYAPVATHPTPVSHPPDDSI | KKPQFKKVKL |
| 1AthHRS1_AT1G13300 | 281 | 1.41e-16 | GKTTGGATTS | STTTTTGIYGTMAAPPPPQWPSHSNYR | PSIIVDEGSG |
| E_L01092.1 | 280 | 3.04e-16 | TSVEESGSGG | GAAASKGIYAPVAAAAPPPPFREAKHK | PSSSHSSDGG |
| Y_30s1094441g001 | 294 | 3.04e-16 | AAAAVATAAQ | PANGAGGIYAPVASLPSDLRHQQQQVK | QQSHRSPTGP |
| R_2G016370 | 393 | 6.44e-16 | VHLAGDASGT | TTTAADKVYAPVATTLTPAPRPRPRPG | PERQSSSCSG |
| R_2G159119 | 399 | 8.24e-16 | VHLAGDASGT | TTTAADKVYAPVATTLTPAPRPRPRPR | PERQSSSCSG |
| L_5g054300.1 | 299 | 1.92e-15 | TKTTVSGELT | TVTTPTGIYAPVATHPSSTTTVIKPKS | KKFELSENSH |
| Z_10P19700_001 | 276 | 1.92e-15 | VAAAAQPANG | ARAPSNGMYAPVASHPSDSRYRQKQPQ | RSITLRWR |
| J_3836112 | 282 | 2.44e-15 | ITGGATTSGT | TTRSTTGIYGAMAAPPLPQWPSPSNFR | PSIIVEEKGS |
| T_1G146600.1 | 348 | 1.23e-14 | AQSQVQLAGD | ASGTANTVYAPVATLPSGTRQGQRQSS | RCSGGRRSGD |
| G_G011857t1 | 323 | 2.14e-14 | AATTASGETA | SVTPTNGIYTPVAAPLPKLPQPSGAIV | QRPQRSQSEE |
| J_3875351 | 315 | 2.39e-14 | ASHSKANAVA | SRETTTGIYGPMASPLPSEWPSHSNFG | GEISEEISRC |
| L_1g093080.1 | 313 | 4.60e-14 | ASSTTTGEMT | KVVAPSGIYAPVATHPQVATIKKPEFK | KFEHSISEER |
| AA_5036G0050 | 336 | 7.86e-14 | QPQVQLSAAD | ALGSADPVYAPVATLPSGLQPHSQKQS | SRYSEGRRSG |
| W_02g22020.1 | 346 | 1.33e-13 | HVQLAAAGNN | ASGSANPVYAPVAMLPAGLQPHSHRKQ | HQQQQQGQRH |
| D_05g009720.2 | 324 | 1.64e-13 | GAPAASGEAS | GVANSNGIYAPIATHPKGPLHDHVSGG | TLRQHNNKSS |
| P_8G0062400 | 341 | 2.23e-13 | AAVAHPPPLT | EGMVCPPIYAPVASLPMLNQHQQQQQQ | QQAKQPHRSP |
| S_8G036900.1 | 404 | 2.48e-13 | LASTGDASGT | PTATANKVYAPVATTLTTPGLLQPRPE | RQSSSCSGGR |
| C_3G398400 | 257 | 7.49e-13 | FQDRPTTGDA | AQLVTNGVYAPVAARPQSPLLTSQRQH | KQPSTVHPES |
| AA_1881G0390 | 303 | 2.40e-12 | QSHAAVHHLQ | HHPAAAMVHRAVAPPPPPAYKAAAMVG | SPESEGRGSG |
| 2AthHOH1_AT3G25790 | 293 | 2.64e-12 | HSTANAVNAV | ASGETTGIYGPMVSSLPSEWPRHSNFG | RKISEDRSRC |
| AA_2274G0070 | 388 | 4.65e-12 | QGYYQSPAAV | HHPAAAMVHRAVAPPPPPAYKAATRVV | SPESEGRGSG |
| M_006G155200.1 | 366 | 5.10e-12 | SHAPPPHFCA | TPPASQDFYTAAAAPPPSLHHHTLHHQ | LQLFRPTSQV |
| B_001193 | 305 | 5.60e-12 | ETKGELNGVS | GVTAHPTIYAPITAPSVPTVRSEQRQK | LSPRSPLQAE |
| Z_11P13410_001 | 283 | 5.60e-12 | YTVAAPPADA | PCTPAVGVFAPAASLPSDSRVRPQQSK | QSDRSPTGPQ |
| R_2G348238 | 400 | 6.14e-12 | SPAAVHHLQH | HPAAAAMVHRAAAAPPPPQQAAYSKAA | AMAGSPPGSE |
| M_018G074200.1 | 365 | 8.87e-12 | SHAPPPHFCA | APPVSQDFYTAAATPPPPLHHHTLYHQ | LHLYKPTAQA |
| H_1.213 | 299 | 9.72e-12 | ATITAATAPA | SGEGCNGIYAPVPVAALPTPAKERPQQ | TQSHRMQSEE |
| Q_112g00420.1 | 306 | 3.40e-11 | DTDAPPVTVS | DATPVSRIYAPVASVPLGSSQSNAKKT | DVNSQSGEGG |
| Z_8P18270_001 | 293 | 3.40e-11 | DYAAASPGNG | AYAPPDSIYAPLASVPSDSKLRNQQLH | QQSQRSIARP |
| T_5G124700.1 | 392 | 8.05e-11 | PPPAAVHHLQ | HHPAAAMVHHRAAAPPPPQAAAYKAAM | AGSPPESSEG |
| R_2G171468 | 133 | 1.13e-10 | ITSPPEPKLL | GGGAPTPIRAAVSAVAVPQPLPPPLFR | REDSSASSGL |
| V_3g10730.1 | 332 | 1.45e-10 | YAAAAAQQQA | NNNSPAVVYAPVATMPSSSSALHVPSK | KKQTTSRGSK |
| G_G037970t1 | 344 | 4.22e-10 | YATAAAAAYS | GAPTLYGTHHPAAAHAPPHFCAPPVPQ | EFYTAAATPA |
| SfaHRS3_1s0189.1 | 405 | 5.82e-10 | PDYVAAAAAA | AQTAAAGVYDPSVSHHSHQYSSYCKST | LPQDHFANFN |
| C_3G045700 | 273 | 1.28e-9 | YTPSTTNANA | PSADSEKIYVPVAATSQLTLSLRQQQQ | QQKHHKQSEL |
| E_H01532.1 | 378 | 1.61e-9 | LSPSFCGSQS | AQPGQQELYPAIAPPPPPHNHQLHRHA | IHHHQQQQQQ |

| | | | | | |
|---|---|---|---|---|---|
| V_2g04810.1 | 431 | 5.01e-9 | PAMAHGRAMA | PPAAAYKVHHPVAASPESEGRGSGGGG | GGGRERSESI |
| R_2G124540 | 131 | 5.01e-9 | SEITSPEPKL | LGGAPMPIRAVAAVPPLPPPFFRREDS | SAGSGLSLVP |
| CbrHRS1_322305 | 658 | 6.73e-9 | AAAAAAAAAG | AGTAAGGFGTPVASLAAAGVSQQRSHF | GLHASSLQTQ |
| MviHRS1_13691 | 471 | 8.39e-9 | QSEVAARAAA | AAAAAASMPIPGVGPPPPVPPPPGMPE | MGPHGMMVRH |
| J_3839360 | 308 | 1.49e-8 | ASHSTANAAT | AGETTIGIYGPLQAEWPSQSNFGRTIS | EERSRCSNKG |
| Z_4P03680_001 | 269 | 1.72e-8 | TDAAAHAAPV | IGVSETGRYADVSPLPSPLMILDRSYQ | QTDKHSKGRN |
| T_3G370000.1 | 89 | 3.09e-7 | ELAQHRRLAP | SPAPAPELFAPAPTSPAPAAKRKAAAP | EGGAVKAEAD |

HRS Motif 2

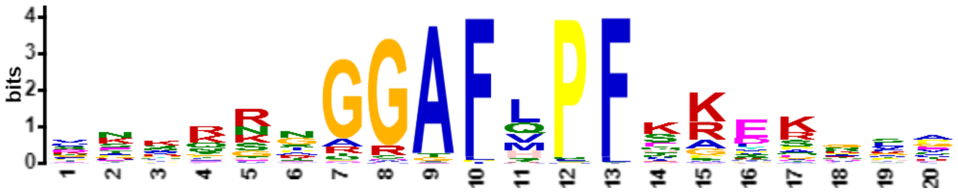


| Name | Start | *p*-value | Sites | | |
|---|---|---|---|---|---|
| F_1015462001 | 150 | 2.34e-20 | GFTVSPKLGF | DNKQRNGGAFLPFSKERNSC | PSPNLRNCEA |
| B_003614 | 148 | 7.99e-20 | CFSVSPKLAL | DNKQRNGGAFLPFSKEKSSC | ATPTLSVLPE |
| M_006G155200.1 | 150 | 2.04e-19 | GFNVSSKLIL | DTKQRNGGAFLPFSKERNLC | PSPTLALAAC |
| M_018G074200.1 | 150 | 2.04e-19 | GFNVSSKLAL | DTKQRNGGAFLPFSKERNLC | PSPTLALSST |
| N_06G060000.1 | 152 | 2.04e-19 | GFSVSPKISL | DTKQRNGGAFLPFSKERNLC | PSPILALAST |
| N_14G127500.1 | 151 | 2.04e-19 | GFNVSPKLGL | DTKQRNGGAFLPFSKERNLC | PSPTLALASA |
| O_29807.m000474 | 150 | 2.04e-19 | GFNVSPKLGL | DTKQRNGGAFLPFSKERNLC | PSPTLALAST |
| H_127.55 | 157 | 3.21e-19 | GFNVSPKLAL | DTKQRNGGAFLPFSKERNSC | PTPTPVRAGL |
| K_06G213400.1 | 157 | 4.02e-19 | GFSVSPKLAL | DNKHRNGGAFLPFSKERNSC | QGLRGLPELA |
| K_04G151000.1 | 154 | 4.02e-19 | GFSVSPNPAL | DNKHRNGGAFLPFSKERNSC | QGLRDLPEVA |
| B_006061 | 153 | 1.19e-18 | CFSVSPKLAL | DSKQRNGGAFLPFSKERNTC | ASPTLRGLPE |
| Y_30s1025391g002 | 152 | 2.74e-18 | AFDVSPKLAL | DTKQRNGGAFLPFSKEKNKG | MGSASRGLPE |
| K_17G119600.1 | 152 | 1.10e-17 | FSISPKLALD | NKQRNGGGAFLPFSKERNSC | QGSTLRPLPE |
| K_05G011500.1 | 152 | 1.10e-17 | FSISPKLALD | NKQRNGGGAFLPFSKERNSC | QGSTLRPLPE |
| Z_11P13410_001 | 126 | 1.10e-17 | KVEPPTKPIA | VNARRIGGAFHPFEREKHVA | APIPASSSTT |
| Z_6P05330_001 | 150 | 1.61e-17 | AFDVSPKLSL | DTKQRNGGAFLPFSKEKSKA | ARSASRALSE |
| AB_01.3080 | 114 | 4.07e-17 | KGDVVMRPIA | VNAKRIGGAFHPFEKEKVVV | KTPVVAAAPA |
| AB_10.37 | 114 | 4.07e-17 | KGDVVMKPIA | VNAKRIGGAFHPFEKEKVVV | KTPVVAVGPV |
| J_3853942 | 155 | 4.87e-17 | ASSPKLGHFD | VKQRNGGGAFLPFSKEKTLP | ELALSTEVKR |
| G_G037970t1 | 151 | 4.87e-17 | GFNVSPKLAL | DTKPRNGGAFLPFTKERNSC | PGSALQALPD |
| D_05g009720.2 | 163 | 4.87e-17 | LTPKKVSVVE | VKKNGSGGAFHPFKKEKNTV | AAVETTPALA |
| Z_7P00660_001 | 149 | 1.18e-16 | YGYHRNFFSE | CKNRNGGGAFVPFKRLSSFA | ASSKEEEKPP |
| G_G011857t1 | 153 | 1.40e-16 | QEGAGKSGSA | VEVKRNGGAFQPFHREKTVE | KSVPSVGKAN |
| AB_06.1057 | 171 | 5.25e-16 | KGDIVRRPVA | VNARKIGGAFHPFEKEKVGV | KAPAAAPASS |
| C_7G203800 | 152 | 6.16e-16 | SFSPKLSLDD | KKRNGGGGAFLPFTKERNTC | ASPNTLRVLS |
| B_006595 | 167 | 7.21e-16 | HSVPENAFQP | CKYRSRGGAFMPFKGYSCFP | ATVVKEDKDV |
| E_E00311.1 | 148 | 7.21e-16 | DSPRKVAVTE | VKRNGGGGAFHPFKKEKSTT | TTTVAVVAAP |
| O_28883.m000751 | 173 | 9.87e-16 | KEDLPRKAAV | TEVKRNGGAFQPFHKEKGIA | KTPPSVPASA |
| B_011780 | 168 | 1.15e-15 | HSVPENPFQS | CKYRSRGGAFMPFKGYTGFP | TTPLKEDKEV |
| X_31310.1 | 124 | 2.11e-15 | PVEPPRKPIA | VSAKKIGGAFLPFEREKQVP | LPLSSATPQI |
| 2PabHRS2_33594g0010 | 115 | 3.81e-15 | LFPTNSSLFS | DSKQRIGGAFLPFSRERQSN | PSVSARSNAV |
| MpoHRS1_0119s0041 | 193 | 3.81e-15 | STVTSPANLF | NTHKRTGGAFLPFIKEKPVA | TVASRVKVLS |
| AA_5036G0050 | 152 | 4.40e-15 | DKELPCKPVA | LNARKAGGAFHPFKKEKRTE | LPASSTTAAV |
| Z_3P21090_001 | 116 | 5.08e-15 | LKEPPKKPIA | VSAKRIGGAFQPFEREKLVV | PPPASRAVAA |
| AA_2274G0070 | 177 | 5.86e-15 | TSPEFSALDG | SAGQRNGGAFLPFSKDKAMA | ESAALPKLAL |
| AA_1881G0390 | 178 | 5.86e-15 | TSPKLSALDG | GSGQRNGGAFLPFSKDKAMA | ESAALPELAL |
| N_14G098400.1 | 148 | 6.75e-15 | KEDVSRKAVE | TEVNRNGGAFQPFQKEKTTG | KNVQTTAKNP |
| L_4g113140.1 | 151 | 8.93e-15 | DNNNNIGFNI | SPKHRNGGAFLPFSKERNNS | SCQGQGLPEL |
| K_01G086700.1 | 146 | 1.03e-14 | DVVVPRKTDV | VEVKRNGGAFQPFQREEKSG | DAKASINNDA |
| M_010G128900.1 | 158 | 1.18e-14 | DLPRKAAVTE | VKRNGAGGAFQPFHREKSVG | KSSNQAISKA |
| N_13G128300.1 | 162 | 1.18e-14 | QSTCENPVQL | CNYKSKGGAFMPFKALSVFE | GTERKEEKEV |
| F_1011942001 | 132 | 1.35e-14 | KEDTPKKIPS | MEVKKNGGAFHPFKRDKAVG | TNPTSAPSAA |

| | | | | | |
|---|---|---|---|---|---|
| N_06G071500.1 | 164 | 2.30e-14 | EEDVPRKAVV | TELKRNGGAFQPFQKEKTTG | KSIQTIAKTP |
| Z_10P27180_001 | 126 | 2.30e-14 | ADVIPKLALD | TTKQRSGGAFLPFSREKSRG | AHATSAEKVV |
| F_1031804001 | 168 | 2.63e-14 | RPATEDPFQP | CKSRTGGRAFLPFKGYSGFP | VATARKEDKD |
| L_5g054300.1 | 138 | 2.63e-14 | KEDVPRKTNV | VEVKRNGGAFQPFHKEEIAA | EKDNALESDK |
| R_2G016370 | 190 | 3.88e-14 | LLLPCRPVAL | NACRKSGGAFQPFEKEKKKE | DKEEKQRAEL |
| 1AtrHRS1_122.65 | 70 | 4.41e-14 | SFPSSQKPAF | NGKQRNGGAFLPFSKDRESR | KTLSDLALAS |
| M_008G117500.1 | 154 | 5.00e-14 | DLPRKAAVTE | VKRNGAGGAFQPFHREKSIG | KTSNQAIAKA |
| W_02g22020.1 | 161 | 6.43e-14 | TKDLPCKPVA | LNARKAGGAFQPFEKEKRAE | LPASSTTAAA |
| Z_8P18270_001 | 123 | 6.43e-14 | LKEPPKKPIA | VSLKKIGGAFQPFDREKHVA | PPAVVAVAPP |
| S_8G036900.1 | 190 | 7.29e-14 | LLLPCRPVAL | NAYRKPGGAFQPFEKDKDKE | KQRGELPLPA |
| J_3875404 | 149 | 9.32e-14 | ASSPKLGHVD | AKQRNSGGAFHPFTKEQHLP | ELALSTEVKR |
| Y_30s1094441g001 | 139 | 9.32e-14 | KAEPPKKPIA | VNAKKIGGAFRPFDREKHVP | APVPSAAAPA |
| Y_30s723941g001 | 123 | 9.32e-14 | KGDPPKRPIA | VNAKKIGGAFHPFERGKHAP | AAPAASSSTV |
| X_9781.1 | 147 | 9.32e-14 | VSPKLGLLDA | KQRNGGGGAFLPFSKEKAKG | GGGVTPRGLP |
| R_2G142748 | 119 | 1.19e-13 | LLLPCRPVAL | NACRKPGGAFQPFEKEKKKK | DKEEKQCAEL |
| R_2G159119 | 190 | 1.19e-13 | LLLPCRPVAL | NACRKPGGAFQPFEKEKKKK | DKEEKQRAEL |
| Q_30g00060.1 | 129 | 1.51e-13 | TQSNNEVPSS | PKRWNTGGAFLPFSKEKTVD | KASGCSDLAL |
| 1PabHRS1_14087g0010 | 152 | 2.15e-13 | SFPSNHKLFS | DSKHRMGGAFHPFSKERQVI | APRPVRSTVE |
| PpaHRS3_11430V3.1 | 201 | 2.15e-13 | RPLTSSRLLL | NSKDRQGGAFVPFNREKQLP | SPPFSSQSTG |
| T_5G124700.1 | 181 | 3.05e-13 | PIDTLGALDT | AAGQRNGGAFLPFAKDKAAA | EAAALPELAL |
| 3AthHOH3_AT1G25550 | 148 | 3.83e-13 | PQPNNKKPMV | IEVKRSAGAFQPFQKEKPKA | ADSQPLIKAI |
| P_27G0017800 | 154 | 3.83e-13 | QRTVTPPEAT | DDKLRTGGAFLPFSKERDGR | VRGGLPELAL |
| Z_6P09870_001 | 124 | 3.83e-13 | KVEPPKKPMA | VSLKKTGGAFQPFEREKHAT | TPPAATAAAS |
| K_02G098800.1 | 146 | 4.29e-13 | DVVPRKTNVV | EVKRNGGGAFQPFQREEKSG | VSKANNNNEA |
| B_001193 | 145 | 1.03e-12 | NQDLSRKPSV | IESKKGGGAFHPFQRDDKLT | APSITVIPSS |
| K_10G204200.1 | 154 | 1.15e-12 | VNKKVPGLEL | KRSGNGGGAFQPFHKEERGA | KTSESLSKAP |
| M_005G134600.1 | 147 | 1.15e-12 | RSTCENPIQL | GNHGNKGGAFVPFKALSGFE | RSKKKEEKEV |
| S_3G046800.1 | 186 | 1.58e-12 | DTLCAHDAAA | AGQRNGGGAFLPFAKDKTAS | AAEGAALPEL |
| G_G000558t1 | 159 | 1.75e-12 | EDLSEKPIEL | CNNKSWGGAFVPFKGQDDKE | ISGLSLMTPS |
| L_4g086835.1 | 182 | 1.94e-12 | TVAEDLFQSC | SSNRNGGSAFMPFSAYSSVP | VTTVTLSAPK |
| 5AthHOH4_AT2G03500 | 156 | 2.39e-12 | NSSPKLGHFD | AKQRNGSGAFLPFSKEQSLP | ELALSTEVKR |
| 3AzfiHRS4_114.g046051 | 201 | 3.24e-12 | AFSTSSKPFF | HSKNKSGGAFMPFNRDKLQA | VVQPRARPGA |
| R_2G348238 | 181 | 4.83e-12 | HGAAGAGHGQ | RNGGGGGGAFLPFAKDKTAS | AAEGAALPEL |
| B_012707 | 133 | 5.34e-12 | KQDLGPKLSV | VETKKSGGAFHPFERDTKRV | TKPTATSVAA |
| Z_4P23210_001 | 172 | 5.34e-12 | RQGKENFFSE | RKSRNGGGAFVPFRGLPPFA | DSCKREEEKP |
| H_1.213 | 138 | 7.15e-12 | KDVPKKTTTA | DQVKINGGAFQPFTKDQPTE | NCTSNKQPAG |
| L_1g093080.1 | 147 | 7.15e-12 | TKKAVPVLEL | KRNGCGGGAFHPFHKEDRVN | NTTSELLSKG |
| W_01g08160.1 | 172 | 8.67e-12 | DTSPKLITAL | DGGGGGGGAFLPFSKDNAMG | DGSAAAAAAL |
| O_29676.m001636 | 158 | 9.54e-12 | RSTCENPTQL | CNHRSKGGGFMPFKSTSGFE | KKEEKEVVSQ |
| V_3g10730.1 | 151 | 9.54e-12 | NKELPCKPVA | LNATKTRGAFHPFEKEKRAE | PELPASSTTA |
| Z_10P28540_001 | 166 | 1.05e-11 | QQKKENLFLE | SKSLSGGGAFVPFKGLSAVT | ANGEEGKPTV |
| PpaHRS1_4480V3.1 | 193 | 1.15e-11 | SSITSSQMLL | SAKQRPAGAFLPFIRDQPGA | SSFLSRPTPR |
| C_3G045700 | 160 | 1.15e-11 | QENSPRKEIS | ADVKENGGAFHPFHKTKPIK | TVASETVTIN |
| SfaHRS3_1s0189.1 | 208 | 1.27e-11 | GSIETSRLLL | SSKQRPTGAFLPFTRKKQLT | PPAVVSQSNP |
| PpaHRS2_8580V3.1 | 194 | 1.27e-11 | SWVTSSQMLL | GANQRPTGAFLPFIREQHGV | NSFLSRPTPR |
| N_12G093900.1 | 157 | 1.27e-11 | RSTCENPVQL | CNYRSKGGAFMAFKALPVFD | GTGRKEEKEV |
| V_2g04810.1 | 178 | 1.27e-11 | KERSEHPPLD | AGTRGNGGAFLPFTKEMITT | TADHSPAALL |
| N_15G143500.1 | 174 | 1.39e-11 | QYEKEDTFQV | CNSRSAARAFMPFKTYSGLS | RNDDNDNNSN |
| K_07G209500.1 | 179 | 1.68e-11 | SVAEDPFQTC | SNRNGGRRAFMPFSRYSSSS | SSVPVTTVGL |
| 2ScuHRS2_42.g012676 | 183 | 2.02e-11 | NSRSVFPSNC | CRIISSGGAFVPFNKHRQLP | AAVSQPDVQK |
| E_H00466.1 | 194 | 2.42e-11 | KEEESSESQS | IFRNREGGAFSPFKKPSGVG | VGVTIKEEKK |
| F_1003551001 | 159 | 2.65e-11 | NSGSENPIRP | CSYGNRAGAFVPFKVSSGFP | AKGDKEVVPV |
| T_1G146600.1 | 155 | 2.65e-11 | RKELLCKPVA | LNTRKAGDAFQPFVKEKRAE | MPASPTTAAA |
| 4AzfiHRS5_92.g042967 | 187 | 3.47e-11 | NSSSIMKPIP | ISCERRGGAFMPFLKTKTAS | TISAVEQQSP |
| J_3835011 | 141 | 3.47e-11 | RCVTQTPFQT | CNNLNQGGAFLPFKPPPAPL | PLMTPMMDCN |
| AA_2901G0070 | 127 | 3.47e-11 | QDKERSSSEV | KVNSHGGGAFMPFKAVGSGA | PAFAPPCFRK |
| 1ScuHRS1_185.g025023 | 221 | 3.80e-11 | AFSNAWKPVF | HSKTSSGGAFTPFNRDKPTV | IQPTARPGAL |
| Z_9P26570_001 | 165 | 4.96e-11 | KEKEDNLFLE | STSRNGRGAFVPFKGISGPK | IKSKEETKTT |
| 6AthHOH5_AT4G37180 | 154 | 5.41e-11 | RCVSQNPFQT | CNYPNQGGVFMPFNRPPPPP | PPAPLSLMTP |
| O_29780.m001324 | 176 | 5.41e-11 | QYANDDTFSA | CKSRSAARAFMPFKTYPDLS | RKEDKHSTQE |
| E_B00308.1 | 187 | 6.44e-11 | RGVMDDLSQS | GKIRTVGRAFVPFKGCANFP | VMTVGKEDKN |
| J_3852516 | 142 | 8.35e-11 | GKKAKVVEVK | PNNYNNCGGFQPFQREKKRE | TDLQPAAVRA |
| T_2G013500.1 | 130 | 8.35e-11 | EKQDKGRSSP | EAKSGGGGAFVPFKAVGSGA | PAFAPLCLRV |

| | | | | | |
|---|---|---|---|---|---|
| MenHRS1_2781 | 212 | 9.09e-11 | YQSPGTKAVL | VGSQRPTGAFLPFTRDRDRQ | VAGTGIGASG |
| K_02G135400.1 | 178 | 1.08e-10 | SVVEDPFQTC | SNINEGRRAFMPFSRYSSSS | SSVPVTTVGL |
| C_6G095900 | 170 | 1.17e-10 | DGSAITTETN | LPRNGGGGAFVPFKGFTGFL | TSTRMEEKER |
| Z_4P32500_001 | 156 | 1.17e-10 | KEELFNLFLE | CKSPGGDGAFLPFEALSTVS | ASSEEKKPAA |
| G_G006998t1 | 171 | 1.28e-10 | KRNDEDPFQG | CKNRGSARAFMPFKPNLGLA | VRKEEKEEIP |
| AB_07.3042 | 164 | 1.28e-10 | HQESETPSLE | SKSKSFGGAFVPFKSVERKE | EVKVTGGLPD |
| SfaHRS4_40s0035.1 | 201 | 1.51e-10 | SSIRSSRGLL | SSKQRSTGAFLPFTRKQVNV | PMLSQSDHVT |
| Z_10P19700_001 | 108 | 1.64e-10 | DTEPPKKPIV | AISKKIGSAFQPFQRQKYVV | PPPASSAATA |
| J_3831280 | 144 | 1.78e-10 | VAKKARVVEV | KPNSHNRGGFQPFQKEKKRV | FSETDLHPAV |
| SfaHRS1_10s0233.1 | 209 | 1.94e-10 | SVTSSRFELF | SSKQPPAGAFLPFVGEQQVV | TAAPPPSGST |
| Q_112g00420.1 | 150 | 1.94e-10 | EETELMKPIS | LDSGKERGAFHPFHRDRKIA | PHVADISLTV |
| Z_9P09630_001 | 159 | 2.10e-10 | DAGKESLNLE | SKNRSAGGAFVQFKGISALA | MKPKEEVLQQ |
| 3AtrHRS3_093.40 | 118 | 2.48e-10 | RNEDAGLNPK | LGTKKQRGAFQPFSREREGR | AEAVSGSGNE |
| K_20G186500.1 | 152 | 3.16e-10 | VSKIVCGVEL | KRSGSGGGAFQPLHKEEKSS | KPSESLSKTP |
| E_H01532.1 | 172 | 3.16e-10 | FKLGFTNEGK | KVINGGGGAFLPFSSSSSMN | CGEGFPELAL |
| M_004G144800.1 | 190 | 3.43e-10 | QYVNEDAFQA | CKGRTAARTFMPFKAYPGSS | RKEDDSNREE |
| N_17G094500.1 | 172 | 3.72e-10 | QFANEDTFQA | CKGRSPARTFLPFKTYSGLS | RKEDNDTSEE |
| P_2G0120600 | 160 | 4.03e-10 | KKGSERAAED | RSGGGGGGAFLPFKEMAGKE | KAAAVPLPPP |
| Z_6P00650_001 | 166 | 9.56e-10 | PRKKESLHLE | SKSLSGGSAFVPFKAPPALS | ASTKKQEKPS |
| E_L01092.1 | 128 | 1.03e-9 | SKQDSPRKSD | VVEVKKNGAFHPFKKAKSGG | GGPITGAAQP |
| D_02g090400.2 | 162 | 1.03e-9 | AKEKQYQLGA | CKLTSERGAFLPFQRQALKV | DKKCLAVKDL |
| Z_4P28210_001 | 161 | 1.12e-9 | HRQGKGKSLE | CKSRSSGVEFLPFKALSPLA | ASSEEEEEKP |
| M_007G039400.1 | 145 | 1.30e-9 | RSTCENPVQL | GDHSNKGEAFVPFKALSGFE | GSKRKEEKEV |
| KniHRS1_00369_0070 | 150 | 1.63e-9 | LRDGGEERAG | PQLFRTPGAFLPFSREQHSP | HVARPSSPRA |
| 2AtrHRS2_033.259 | 171 | 1.90e-9 | TDFGEKAYTI | GRYRNSGGAFVPLNGNGNGN | GLVRKEENDI |
| 2AzfiHRS3_92.g042991 | 107 | 1.90e-9 | SAKLSNGIIS | CNPERRRGAFVPFTMTRLRA | TNISPPDHDH |
| S_2G016300.1 | 168 | 1.90e-9 | SEKQDKGRSL | PEAKSRGGALLPFKAAVGSG | APAFASLCLR |
| CbrHRS1_322305 | 335 | 2.56e-9 | TAGNDISFLQ | PGRERPGGAFLPFVPNRDNK | ASTGVGTEKG |
| AA_0456G0940 | 128 | 3.19e-9 | EKQDKERSSS | EANSHGGGGFVPFKAVGSGA | PAFAPPCFRK |
| 4ScuHRS4_13.g005748 | 140 | 3.43e-9 | ENVYTDMEGM | KPVIGNGRAFLPFASLRKSI | LSSISAEHGV |
| SfaHRS2_4s0229.1 | 250 | 4.59e-9 | SSVTSSRLFF | SSTQRPAGAFLPFIRGLQQV | TAVAPLLSHS |
| SmoHRS2_66160 | 105 | 5.30e-9 | VEIYSKHAVP | GIFANSGGAFSPFVRAAPST | EEKPGLRSTS |
| PEQU_26724 | 117 | 5.30e-9 | TSPEESSKLR | KPVAKIGGAFQLLEKEKPCP | PPASEVLESS |
| E_N01205.1 | 176 | 8.14e-9 | KWGGVDLIKS | GNNRTEGRAFSSFKRCADLP | ALSLATPEIK |
| R_2G173882 | 126 | 9.37e-9 | KQDKERSSPE | ARSRAAGGAFLPLKAAVGSG | APAFAPLGLR |
| M_009G106600.1 | 179 | 1.24e-8 | QYGNEDAFQA | GIGRTAARTFMPFKACSGLS | SKEEKLPVPS |
| R_2G100176 | 141 | 1.63e-8 | ESEKQDKGRC | SPEARSRGALLPFKADVGSG | APAFAPLFLR |
| H_6.397 | 157 | 2.30e-8 | SFAEEHPVQP | SNHSSRGGAIGPFKGKSGFQ | EKGVPSLSLM |
| E_K00828.1 | 158 | 2.30e-8 | RCDQEGEGNG | SKFRHKEGSFMPFKKSYDVA | MVEEERGGNP |
| Z_6P34680_001 | 162 | 2.46e-8 | DRQEEEENLC | LESKSGADTFVPFKGMAALV | MSSKQVTPTA |
| PEQU_27343 | 155 | 2.82e-8 | SHGRKDCDAL | FLESNTGRAFLPFKAMAAPR | KEEEKLSISL |
| Z_2P13380_001 | 162 | 6.65e-8 | RRQEEQSNLE | CKNRRSGAALSLFKGLPPLA | ARSMTEDKPT |
| C_3G398400 | 106 | 8.62e-8 | KDKPSQNTIG | IEPNRLKGAFRPFQRDEGHR | ASTATSTSTT |
| 5ScuHRS5_10.g004815 | 289 | 9.80e-8 | IDVTVELRSS | SWRPRGGGAFSPFPSSSKQL | CSSSSNGTGS |
| 4AthHOH2_AT1G68670 | 144 | 9.80e-8 | EERVAKKAKV | VEVKPKSGAFQPFQKRVLET | DLQPAVKVAS |
| P_8G0062400 | 169 | 1.11e-7 | DETSRQTIVA | ADRNKVTSAFHPFLPQKRSP | PPATGTVEVA |
| J_3862509 | 128 | 1.26e-7 | DRCVTQTCSN | NNNSNQGGAILSFNVPPRPP | PLSLRTPILT |
| Q_208g00340.1 | 137 | 1.35e-7 | STTSEATVTP | KPATESNGTFLPFYKVERSS | SSTSNKGNLP |
| SmoHRS1_402040 | 152 | 2.36e-7 | EFETRKALRS | STASADSGAFAPFVRDKKWD | QAELCLSSSI |
| J_3864545 | 138 | 2.51e-7 | STNEKEDRSV | THKPNQGWAFTPFNLPPPPP | PLTLRTPPSE |
| CatHRS1_Chlat1_8216 | 233 | 2.67e-7 | VVVPQPHRPA | VVLASHGGAFTPFLPAREPD | PNRSPPRSQS |
| 7AzfiHRS2_208.g057980 | 175 | 3.42e-7 | DKHRTEISLV | FPKDRRQGAFSPFLPKDGMR | TSSNVESGEN |
| W_07g02800.2 | 131 | 2.94e-6 | PEKQDKVRIS | SEAKSNGGAFVGSGAPAFAR | PKQSLMRKED |
| AA_3297G0100 | 114 | 3.11e-6 | EKEQESESTS | PESKLRGGAPVPLRAVGCGA | PALPPYFGRE |
| S_1G078900.1 | 128 | 6.67e-6 | DKEHQSEITS | PEPKLLGGAPMPIRAVAAVP | PLPPPFFRRE |

HRS Motif 3

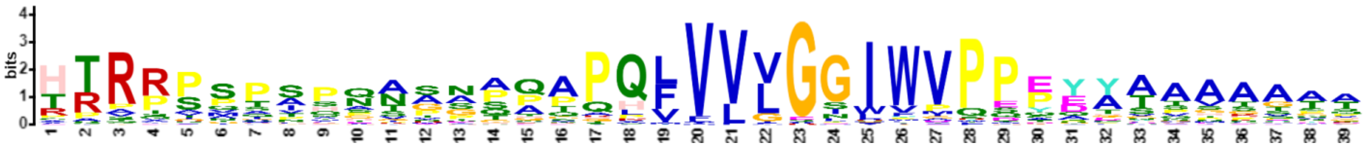

| Name | Start | *p*-value | Sites | | |
|---|---|---|---|---|---|
| M_018G074200.1 | 304 | 1.28e-38 | VKSHLQKYRL | HTRRPSPSPQAAGAPPPQLVVLGGIWVPPEYATAAAAAH | SGGPTLYGAH |
| B_003614 | 300 | 2.04e-38 | VKSHLQKYRL | HTRRPSPSPQVAGTPAPQLVVLGGIWVPPEYATAAAAAA | AHPAAASTIY |
| N_14G127500.1 | 313 | 3.23e-38 | VKSHLQKYRL | HTRRPSPSPQAPGAPTPQLVVLGGIWVPPEYATAAAAAH | SGAPTLYGTH |
| O_28883.m000751 | 289 | 1.92e-37 | KSHLQKYRLH | TRRPSPTIHNNSNPQAPQFVVVGGIWVPPPEYAAVAATT | ASMETVTTAA |
| M_008G117500.1 | 276 | 5.62e-37 | KSHLQKYRLH | TRRPSPTIHNNSNQQAPQFVVVGGIWVPPPEYAAVAATT | ASAETSPISA |
| Z_6P05330_001 | 262 | 6.95e-37 | VKSHLQKYRL | HTRRPAPAPQAAAAAAPQLVVLGGIWVPPEYATSAAAAA | AAGPAIYGAH |
| N_14G098400.1 | 269 | 1.31e-36 | KSHLQKYRLH | TRRPSPPIHNNSNPQAPQFVVVGGIWVQPPEYAAVAATT | ASGETSTIAA |
| 1PabHRS1_14087g0010 | 315 | 2.98e-36 | VKSHLQKYRL | HTRRPSPSPPSSNPQAPHLVVLGGIWVPPEYAAHAAAAA | QQGQGGLYNP |
| O_29807.m000474 | 306 | 5.49e-36 | VKSHLQKYRL | HTRRPSPSPQAPGAPTPQLVVLGGIWVPPEYATAAHGGT | PTLYGTHHHP |
| B_006061 | 303 | 1.00e-35 | VKSHLQKYRL | HTRRPSPSPQAAGTPAPQLVVLSGIWVPPEYATAAAAAA | HPAAPTIYSA |
| 2PabHRS2_33594g0010 | 287 | 2.68e-35 | VKSHLQKYRL | HTRRPSPSPPTPASQAPQLVVLGGIWVPPEYAAHAAVAA | HQGPGLYGTL |
| G_G011857t1 | 278 | 2.68e-35 | KSHLQKYRLH | TRRPSPTVHNNANAQAPQFVVVGGIWVPPPEYAPMAATT | ASGETASVTP |
| M_006G155200.1 | 305 | 3.26e-35 | VKSHLQKYRL | HTRRPSPSPHAAGAPAPHLVVLGSIWVPPEYATAAAAAH | SGGAALYGAH |
| H_127.55 | 314 | 7.05e-35 | VKSHLQKYRL | HTRRPSPSPQATGAPAPQLVVLGGIWVPPEYATAHGGTP | TLYGTHHHPA |
| G_G037970t1 | 304 | 1.03e-34 | VKSHLQKYRL | HTRRPSPSPQATGAPTPPLVVLGGIWVPPEYATAAAAAY | SGAPTLYGTH |
| S_3G046800.1 | 331 | 1.25e-34 | VKSHLQKYRL | HTRRPMPAPPAPATAAPQLVVLGGIWVPPEYATQAAGQA | IYGAHPATQP |
| E_H01532.1 | 312 | 1.50e-34 | VKSHLQKYRL | HTRRPSPSPQTSVAAGPQVVVLGGIWVPPEYAAAAAAHH | HSGAQAAALY |
| X_9781.1 | 297 | 1.81e-34 | VKSHLQKYRL | HTRRPIPAPPASAAAAPQLVVLGGIWVPPEYATTAAGPA | IYGAHPGPAH |
| T_5G124700.1 | 320 | 2.64e-34 | VKSHLQKYRL | HTRRPMPTPPAPATAAPQLVVLGGIWVPPEYATQAAGQA | IYGAHPATQP |
| AA_2274G0070 | 321 | 3.17e-34 | VKSHLQKYRL | HTRRPMTAPPTAAPAAPHLVVLGGIWVPPEYATQAAAPT | IYGAHPATQP |
| F_1011942001 | 244 | 3.82e-34 | KSHLQKYRLH | TRRPNPAIQHNGNPQAPQFVVVGGIWVPPPEYTAVAATT | SSGEATGVTT |
| R_2G348238 | 329 | 9.51e-34 | VKSHLQKYRL | HTRRPMPAPPAPATAAPQLVVLGGIWVPPEYASQAAGQA | IYGAHPATQP |
| M_010G128900.1 | 280 | 2.33e-33 | KSHLQKYRLH | TRRPSPTIHTNSSQQAPQFVVVGGIWVPPTEYAAVAATT | TAGETSTISA |
| C_7G203800 | 294 | 1.88e-32 | VKSHLQKYRL | HTRRPSPSHQAAGNQAPQLVVLGSIWVPPEYATAAHGGG | PTIYSAHPAS |
| W_01g08160.1 | 321 | 7.23e-32 | VKSHLQKYRL | HTRRPMPSPAPPTAATPQLVVLGGIWVPPEYATQAAGPA | IYGAHPATQP |
| J_3875404 | 278 | 1.01e-31 | VKSHLQKYRL | HTRRPSPSPQTSGGQGPHLVVLGGIWVPPEYTTAHGGTP | TLYHHQVHHH |
| E_L01092.1 | 232 | 1.40e-31 | VKSHLQKYRL | HTRRPSTTIHNNNTQAPQFVVVGGIWMQPEYAAMATTAT | SVEESGSGGG |
| Q_208g00340.1 | 266 | 1.94e-31 | VKSHLQKYRL | HTRRPNPLMQAPPPGAPQLVVLGGIWVPPEYAAAAGATF | YGTHPSAHYC |
| B_012707 | 252 | 1.55e-30 | KSHLQKYRLH | TRRPSPAIHSNGNQQTPQFVVVGGLWVPPPDYAAVTATK | REANGVAANG |
| 5AthHOH4_AT2G03500 | 288 | 1.55e-30 | VKSHLQKYRL | HTRRPSPSPQTSGGPGPHLVVLGGIWVPPEYTSAHGGTP | TLYHHQVHHH |
| B_001193 | 259 | 2.47e-30 | KSHLQKYRLH | TRRPSPAVQNNGSPQPPQFVVVGGIWVPPHDYAAATETK | GELNGVSGVT |
| D_05g009720.2 | 277 | 3.36e-30 | SHLQKYRLHT | RRPSPSSIHNNNNQQPPQFVVVGGIWVPPPEYASMAAGA | PAASGEASGV |
| V_2g04810.1 | 333 | 4.57e-30 | VKSHLQKYRL | HTRRPMQAPAAPPAGAPQLVVLGGIWMPPDYAAQAAQAA | AGPAAIYGAH |
| H_1.213 | 257 | 3.24e-29 | KSHLQKYRLH | TRRPTQNVQTNGNSQAPQFVVVGGIWVPPQEYATITAAT | APASGEGCNG |
| SfaHRS2_4s0229.1 | 422 | 4.35e-29 | KSHLQKYRLH | TRRPSPSPQSVSQQPPPQLVVLGGIWVPPEYAAVAAAAA | QPTSGVYNPS |
| MpoHRS1_0119s0041 | 361 | 4.35e-29 | VKSHLQKYRL | HTRRPSPAPQAPQQQAPQLVVLGGIWVPPEYTSGGMYHD | PSSSVSQQGH |
| N_06G071500.1 | 287 | 5.01e-28 | SHLQKYRLHT | RRPSPAIHNNSNPQAAPQFVVVGGIWVPPPEYAAVATTT | ASGETSTLAA |
| SfaHRS3_1s0189.1 | 366 | 4.55e-27 | VKSHLQKYRL | HTQRPSPPPQSASAHAPQLVVLGTIWGPDPDYVAAAAAA | AQTAAAGVYD |
| W_02g22020.1 | 296 | 2.25e-26 | KYRLHTRRPS | STGQSSAAAGVPAPPAPQFVVVGSIWVPPPEYAAAAAAQ | QHVQLAAAGN |
| P_27G0017800 | 291 | 2.25e-26 | KSHLQKYRLH | TRRPISAPQASAAAAPPQLVVLGGIWVPPEYTASSTTAK | AGQLFYGGHP |
| 1AtrHRS1_122.65 | 207 | 2.92e-26 | VKSHLQKYRL | HTRRPSPTPQAVPTPAPQLVVLGGIWVPQPDFSNPSPNV | NPSNPSQTLY |
| T_1G146600.1 | 294 | 5.60e-26 | KYRLHTRRPN | STTVVQSTSTSAAQPAPQFVVVGGIWVPPAEYATAAAAA | SAAAAAQSQV |
| C_3G045700 | 232 | 1.06e-25 | KSHLQKYRLH | TRRPGPSMHGNGNTPPTQFVVVGGIWVPPQDYTPSTTNA | NAPSADSEKI |
| R_2G016370 | 339 | 2.28e-25 | LHTRRPNSAA | AAVQSGGTSVVAPPAAPQFVVVGGIWVPPPEYAAAAAAQ | PQQVHVHLAG |
| AA_5036G0050 | 283 | 2.28e-25 | HLQKYRLHTR | RPSSTVQSSAAVAPPVPQFVVVGGIWVPPTQYAAAAAAA | AAAAQPQVQL |
| S_8G036900.1 | 338 | 2.93e-25 | HTRRPNSTTA | VAVQSGGTSVVAPPTAPQFVVVGGIWVPPPEYAAAAAAA | AAAAAAAAAS |
| E_E00311.1 | 269 | 4.83e-25 | LQKYRLHTRR | PNNQSMQNNNNNNQQAPQFVVVGGIWMQPEYATPMAAAT | TTSGEAASGV |
| J_3831280 | 268 | 6.19e-25 | VKSHLQKYRL | HTRRPATAQGNGNSQQPQFVVVGGIWVPSPQDFPPPSDV | ANKGDGVYAQ |
| SfaHRS4_40s0035.1 | 358 | 7.01e-25 | VKSHLQKYRL | HTRRPSPPPQTASAPTPQLVVLGSIWGSDPDYMAGSTAQ | TAVEVHDPSP |
| K_01G086700.1 | 266 | 1.87e-24 | VKSHLQKFRL | HTRRSPIIHNNASSQAGPLFLVGNIFVQPPEYAAVATTS | TASGEELTTV |
| Z_10P27180_001 | 254 | 3.03e-24 | SHLQKYRLHT | RRPIPAPHAAAGAGAPPQLVVLGGIWVPPEYATSAGPAM | YGAHAAPVPR |
| 1AthHRS1_AT1G13300 | 237 | 6.20e-24 | KSHLQKYRLH | TRRPRQTVPNNGNSQTQHFVVVGGLWVPQSDYSTGKTTG | GATTSSTTTT |
| 1ScuHRS1_185.g025023 | 368 | 6.99e-24 | VKSHLQKYRL | HTRRPNASSAAMSTHPPQVVVLGSIWVPPPDQSPALYGS | LQPRSRIYRS |
| Z_1P23220_001 | 195 | 6.99e-24 | KSHLQKYRLH | ARRRRSTAVENPSAGVPQLVVLGGIWIPPPGYGVAAATH | PVVNEPRNGT |
| J_3871943 | 232 | 7.86e-24 | KSHLQKYRLH | TRRPSQTVPNNGNSQKQHFVVVGGLWVPQSDYSTGKTTG | RATTSGTTTT |
| P_8G0062400 | 294 | 1.26e-23 | HLQKYRLHSR | RPSPAMSHSNAANAPPHQVVFLGGLWVPPPEYAVAAAAA | VAHPPPLTEG |
| R_2G159119 | 341 | 1.26e-23 | LHTRRPNSAA | AAVQSGGTSVVAPPAAPQFVVVGGIWVPPPEYAAAVAVA | AAAAAQPQQV |

| | | | | | |
|---|---|---|---|---|---|
| Z_11P13410_001 | 242 | 1.42e-23 | QKYRLHNRRP | STAVHSSSSNTTNPPIPQFVVVSGIWVPPPDYTVAAPPA | DAPCTPAVGV |
| KniHRS1_00369_0070 | 333 | 2.01e-23 | KSHLQKYRLH | TRRPSPIPTSGASPGPGQVVVLGSIWVPPEYNAAASSAA | ALYDAATAAA |
| N_06G060000.1 | 313 | 2.26e-23 | VKSHLQKYRL | HTRRPSPSPQATGAPARLVVLGGIWVPPEYATAAAAAHT | GAPTLYGAHP |
| PpaHRS2_8580V3.1 | 350 | 2.54e-23 | KSHLQKYRLH | TRRPSSSPQPAGAGQSPKLVVLGGIWVPPQYAASGSQPS | SGVYDPSIAH |
| 2AthHOH1_AT3G25790 | 252 | 3.20e-23 | KSHLQKYRLH | ARRPSQTTPNNRNSQTQHFVVVGGIWVPQTNHSTANAVN | AVASGETTGI |
| PpaHRS1_4480V3.1 | 349 | 4.03e-23 | KSHLQKYRLH | TRRPSSSPQSGGMGQSPQLVVLGGIWVPPQYAASGSQPS | PGFYNPSAAH |
| J_3853942 | 278 | 4.52e-23 | VKSHLQKYRL | HTRRPSPSPQTSGGQGPHLVVIGAISAPPEYTTAHGGTP | TLYHHQVQNH |
| J_3875351 | 276 | 5.07e-23 | KSHLQKYRLH | TRRPSQTIPNNRDSQTQHFVVVGGIWVPQASHSKANAVA | SRETTTGIYG |
| K_05G011500.1 | 303 | 6.37e-23 | VKSHLQKYRL | HTRRPSPSPQAGAAAPQLVVLGGIWVPPEYATAAHTATP | TLYGAHPTSH |
| K_10G204200.1 | 266 | 6.37e-23 | KSHLQKYRLH | TRRPSPMVHNSSNPQAAPFVLVGNIFVQSPEYAAVATST | ASREVATVAA |
| Q_30g00060.1 | 274 | 6.37e-23 | KSHLQKYRLH | TRPLPAPAQHAVASGHPKLVVLGGIWVPPEYATSATVYS | THPSSATQFC |
| J_3839360 | 269 | 1.26e-22 | KSHLQKYRLH | TRRPGQTIPNKRDSQTQHFVVVGGIWVPQASHSTANAAT | AGETTIGIYG |
| Y_30s1025391g002 | 252 | 1.26e-22 | VKSHLQKYRL | HTRRPIPAPQASVAAAPQLVVLGSIWVPPELGSPESEMR | SGGERSESIE |
| K_17G119600.1 | 304 | 1.76e-22 | VKSHLQKYRL | HTRRPSPSPQAVGAAPQLVVLGGIWVPPEYATAAHTATP | TLYGAHPTSH |
| L_4g113140.1 | 290 | 1.97e-22 | VKSHLQKYRL | HTRRPSPSAQNGAPAAQLVVLGGIWVPPEYATAAHAGGT | PTLYGGHPTS |
| Y_30s1094441g001 | 252 | 2.75e-22 | HLQKYRLHSR | RPSPAVQSSGTSSPPTPQYVVVGEIWVRPPDYAAAAVAT | AAQPANGAGG |
| L_5g054300.1 | 253 | 5.34e-22 | VKSHLQKFRL | HTRRSPIIHNNSNSHTAPMFLVGNIYVQPQEYAAVATKT | TVSGELTTVT |
| D_12g006800.1 | 294 | 5.96e-22 | VKSHLQKYRL | HTRRPSPSPQATTAPPHLVVLGGIWVPPEYAAAAAHGGA | PPTATFYGPH |
| K_20G186500.1 | 264 | 1.42e-21 | KSHLQKYRLH | TRRPIPMVHNSSDPQAAPFVLVGNIFVQSPEYAAVATST | ASREVATVAA |
| 4ScuHRS4_13.g005748 | 271 | 2.19e-21 | VKSHLQKYRL | HTRRPNQNFHNMPSHTPQMVLIGGIWVPEYTEHAAGVSQ | RNSADQHNYH |
| SmoHRS1_402040 | 284 | 3.73e-21 | KSHLQKYRLH | TRRPATVAPGTAAQKSPQVVVLGSIWVPPEYAVFDHHQA | APSMQSTPLQ |
| K_02G098800.1 | 266 | 4.15e-21 | VKSHLQKFRL | HTRRSPIIHNSASSQAGSLFLVGNIFVQPPEYATSSASG | GELTTATPAA |
| J_3836112 | 236 | 6.33e-21 | KSHLQKYRLH | TRRPSQTVSNNGNSQAQHLVVVEGIWVPKSDHSTGRITG | GATTSGTTTR |
| 3AtrHRS3_093.40 | 224 | 1.80e-20 | SHLQKYRLHT | RTPAAVPASQPINPQAPQFVVVGGIWGLPEYAVPTPSPV | TCGAIYATPI |
| J_3852516 | 276 | 1.99e-20 | HLQKYRLHTR | RPATTVTAQGNGNSQQPQFVVVGGIWVPSPQDVPPPSDV | ANNGGGAYAP |
| K_06G213400.1 | 309 | 4.07e-20 | VKSHLQKYRL | HTRRPSPSLHTGGPTPQLVVLGGIWVPPEYATTTAAMTN | SGDPTLYGPH |
| AB_01.3080 | 241 | 4.99e-20 | LQKYRLHTRR | PTASAVQNSTNSSPQQPQIVLVGGIWMPPPQYNATQNAP | LNTVFAPVAS |
| F_1015462001 | 250 | 6.76e-20 | VKSHLQKYRL | HTRRPSPSPQATGAAAPQLVVVGLGGIWVPPEYATAAHT | GGTTLYGAHP |
| MenHRS1_2781 | 467 | 1.36e-19 | YRLHTRRPSP | VPHATAAAAAAAAQPQPQLVVLGGMWVPPTYAAVTSGAQ | HQSPSGYDLA |
| PEQU_26724 | 216 | 2.72e-19 | LQKYRLHTRR | LPPTIQSSSRISPQIPPQFVLVGGILVPPPEYAVAASSA | EQSSAAFNGL |
| K_04G151000.1 | 307 | 5.40e-19 | VKSHLQKYRL | HTRRPSPSLQTGAPTPQLVVFGGIWVPPEYARAAAHSGG | PTLCGPHPTS |
| AB_06.1057 | 287 | 5.40e-19 | LQKYRLHIKR | PTTPTVHNNSNSNAQQPQIVLVGSLWMPPPNYTTEAEST | PSNEIYTPVA |
| 4AthHOH2_AT1G68670 | 274 | 1.28e-18 | LQKYRLHTRR | PAATSVAAQSTGNQQQPQFVVVGGIWVPSSQDFPPPSDV | ANKGGVYAPV |
| Z_8P18270_001 | 253 | 1.71e-18 | QKYRLHTRRP | TPTVQSSSSSTSSPPATQFVLVGGILVSPPDYAAASPGN | GAYAPPDSIY |
| Z_10P19700_001 | 228 | 6.15e-17 | LQKYRLHGRR | RSPAVQCSSNGSLAVSPQVVLVGGIVVPSPDYNMADAAV | AAAAQPANGA |
| 5AzfiHRS6_15.g013719 | 303 | 8.77e-17 | KSHLQKYRLH | TRRPNGCSGERTKAQTPQLVVLGGIWVTPPSQSSSEDPA | FAATTTHSLD |
| Z_3P21090_001 | 241 | 9.57e-17 | IKSHLQKYRL | HTRRPSPAVQSSSHPAPQFVLLVPPPDYAAAAAVAAAAQ | PGNGACAPTN |
| CbrHRS1_322305 | 590 | 1.05e-16 | VKSHLQKYRL | HTRRPSPPQNISLPHPPQVFVLGSFYLPDYGSSPTLAQQ | PPIVATTPAG |
| V_3g10730.1 | 290 | 2.73e-16 | QKYRLHTTRR | PSSTVAQSNNAAQAAAPQFVVVGSCIWVPPQEYAAAAAQ | QQANNNSPAV |
| Z_4P03680_001 | 222 | 4.97e-16 | VKSHLQKYRL | HARRSCPVEEISTSSNIQFVVVRGIWIPTPDCAILTSTD | AAAHAAPVIG |
| 3AzfiHRS4_114.g046051 | 360 | 5.41e-16 | VKSHLQKYRL | HTRRPSPSPAVSAHTPQVVLVGGIWVPPEYALHNSTHNS | IQNGIDQSPA |
| J_3829805 | 260 | 1.16e-15 | HLQKYRLHTR | RPATPTLTNGCENPQQQQLIVVEGIWVPSKDAVNNRVYA | PVAVQPPPRS |
| L_1g093080.1 | 268 | 3.69e-15 | SHLQKYRLHT | RRPSSTNNESANSQTAAPFVLVGNIFVQQQEYGGVASST | TTGEMTKVVA |
| 3ScuHRS3_71.g016842 | 684 | 8.31e-15 | MRRPHNMSLG | SIQTDNPIVEKGNTNGHHLVVFGHIWVPPEQYQQPTNVH | TLSHHSHPIN |
| 3AthHOH3_AT1G25550 | 267 | 9.76e-15 | HLQKYRLHTR | RPATPVVRTGGENPQQRQFMVMEGIWVPSHDTTNNRVYA | PVATQPPQSS |
| M_009G106600.1 | 316 | 2.74e-14 | VKSHLQKYRL | HTRRVTPATAAAPANQSVVVLGGLWMTQDQYGDSSKATS | SQSGSPQGPL |
| 4AzfiHRS5_92.g042967 | 329 | 9.46e-14 | HLQKYRLHTR | RQVDGSFGKTTMSHDVPQLVVLGGIWVPQGYGNEGSKQQ | KLASNLPSDS |
| M_004G144800.1 | 331 | 1.49e-13 | VKSHLQKYRL | HTRRVPPATASAPANQSAIVLGGLWMAQDQYGDSSKANS | SKSGSPQGPL |
| C_6G095900 | 316 | 3.40e-13 | VKSHLQKYRL | HTRKFPANASSPGPENQKLALFGGLWAPEDQCGASKAST | SQSESPQGPL |
| PpaHRS3_11430V3.1 | 350 | 5.70e-13 | VKSHLQKYRL | HTRRPSPSPPSPSHAPQLVVVGWVRPEFTAVEGAASQAS | TGDNPLPGIS |
| Z_4P28210_001 | 267 | 6.14e-13 | SHLQKYRLHT | ERVHAQRLPNAVATADRAVVVLGGLWDREEQYTSSSQQS | FSQSGSPMSP |
| Z_6P09870_001 | 237 | 4.87e-12 | VKSHLQKYRL | HTRRPNPADQSSSSRPQFVLVGGLVVPPASPPGGNGACA | PPNGIYAPVA |

HRS Motif 4

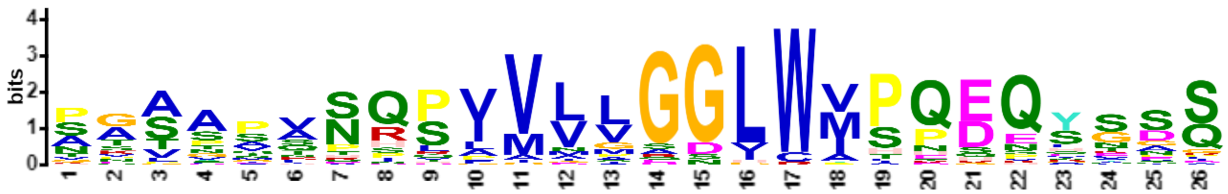

| Name | Start | *p*-value | Sites | | |
|---|---|---|---|---|---|
| N_17G094500.1 | 318 | 9.12e-25 | YRLHTRRMPP | ATAASANQSVVVLGGLWMSQDQYGDS | SKTTSSQSGS |
| T_2G013500.1 | 295 | 1.57e-24 | QKYRLHNRRA | PGSAVVSQPIVLVGGLWIPQEQSSSQ | SGSPQGPLHF |
| B_006595 | 318 | 4.49e-24 | KYRLHTRRLP | AVSPPANQPVVVLGGLWMPQDQYGPS | PKPSTSQSGS |
| R_2G173882 | 282 | 2.77e-23 | QKYRLHNRRA | PGSGVVSQPIVLVGGLWIPQEQSSSQ | SGSPHGPLHF |
| AA_3297G0100 | 270 | 3.24e-23 | QKYRLHNRRL | PGAAPASQPIVLLGDLWVPQEQSSSQ | SGSPQGPLQF |
| B_011780 | 287 | 3.80e-23 | KYRLHTRRLP | AASSPANQSVVVLGGLWMPQDQFGAS | SKPSSASQSG |
| F_1031804001 | 319 | 4.44e-23 | RLHTRRMPTT | SAAPPSNQPVVVLGSLWMPQDQYGDS | SKASSSQSAS |
| K_07G209500.1 | 329 | 5.20e-23 | LQKYRLHTRR | VPAASSNQPVVVLGGLWMSQDQYNDS | SKVSSSGSGS |
| K_02G135400.1 | 331 | 5.20e-23 | LQKYRLHTRR | VPAASSNQPVVVLGGLWMSQDQYNDS | SKVSSSGSGS |
| S_2G016300.1 | 329 | 7.08e-23 | QKYRLHNRRA | PGSGVVSQPIMLVGGLWIPQEQSSSQ | SGSPHGPLHF |
| T_9G080000.1 | 282 | 9.63e-23 | QKYRLHNRRS | PGVAPVSQSIMLVGGLWVPQEQTSSQ | SGSPQGPLQF |
| N_15G143500.1 | 325 | 2.37e-22 | YRLHTRRMPP | ATAASANQSVVVLGGLWMSKDQYGDS | MKATSSQSGS |
| S_1G078900.1 | 274 | 8.69e-22 | QKYRLHNRRS | PGVAPVSQPIMLVGGLWAPQEQSSSG | SPQGPLQFSG |
| L_4g086835.1 | 334 | 8.79e-21 | QKYRLHTRRV | PAASGTDQSVVVLGGLWMPQEHYNDS | SKGSSTASGS |
| R_2G100176 | 305 | 8.59e-20 | QKYRLHNRRA | PGSGVVRQPIVLVGGLWIPQEQGSPQ | SGSPHGPLHH |
| PEQU_27343 | 310 | 5.13e-19 | YRLHTRRIPN | SSATQVNRPVVMVGGLWVSQDQQNAS | QSGSPQGPLQ |
| AA_1535G0300 | 279 | 1.44e-18 | QKYRLHNRRS | PGAAPASQQIVLLGDLWVTQEESSSQ | SGSPQGPLQF |
| H_62.51 | 325 | 3.90e-18 | RLHTRRVPPT | NATSTTNSSAMMLGGLWVSQDQYVDS | SKPSNSQSGS |
| Z_6P34680_001 | 309 | 1.15e-17 | KYRLHTRKMP | NASPPASRPVVVLGGLWVPPENHTAP | PQKSVSQSGS |
| Z_9P09630_001 | 253 | 3.26e-17 | KYRLHSRKLP | NASASFSRPPMVLGGLWVPPENHTVS | PQQSDSQSGS |
| W_03g55590.1 | 284 | 6.02e-17 | QKYRLHNRKS | PGTASASHSIVLVGDLWASQEVSCSQ | SGSPQGPLQL |
| R_2G171468 | 292 | 9.02e-17 | RLHNRRSSPG | AAAAPVSQSIMLVDGVWAAQEQSSGS | QSGSRQGPLQ |
| V_1g07630.1 | 284 | 1.64e-16 | QKYRLHNQRS | SGSSSSSHSIVLVGDHWPPQEQSSSQ | SRSPEAEGPL |
| R_2G124540 | 271 | 2.42e-16 | KYRLHNRRSP | GVVAPVSQSVMLAGGLWAPPHQEQSS | SQSGSPQGPL |
| Z_10P28540_001 | 318 | 4.33e-16 | KYRLHARRMP | NSSDAVNRRVAVAGGVWVPEEQYSNS | SQQSVSQSGS |
| D_01g108300.2 | 323 | 1.23e-15 | YRLHTRRIPN | SQTQPANQSGVALGNLWMSQGQYGES | SKQSSSQSGS |
| O_29780.m001324 | 326 | 2.57e-15 | RLHTRRMPSA | TAAATNHSLVVLGGGLRMSQDQYGDD | SKAAGSQSGS |
| G_G006998t1 | 328 | 8.27e-15 | RLHTRRLPPS | TTTPANQSVVVLGSGLWISQDQYGES | SKGSSSQSGS |
| AA_0456G0940 | 291 | 4.67e-14 | YRLHNQRAPS | SAVVGQPILFVNPAGLWIPPEQSSSQ | SGSPQGPLHF |
| Z_9P26570_001 | 323 | 6.54e-14 | KYRLHTRKMP | NATSAISQPVMVVRGLCVPDENYTTL | PRSASHSGSP |
| K_20G009800.1 | 306 | 8.41e-14 | YRLHTQRVPV | ATAANFSRSAVDLGGLWMHNESLKGG | SSGSPQGPLQ |
| K_07G178500.1 | 306 | 9.93e-14 | YRLHTQRVPV | AKAANSNRSAVALGGLWMHNESLKGR | SSGSPQGPLQ |
| AA_2901G0070 | 289 | 1.92e-13 | YRLHNRRAPS | SAVVSQPILVGNPAGLWIPPEQSCSQ | SGSPQGPLHF |
| 2AtrHRS2_033.259 | 302 | 3.66e-13 | TRRPNVSVSS | ANSNQQNPPLVLLSNLWVPHDKLYNQ | TSSTASGSSS |
| W_07g02800.2 | 289 | 3.66e-13 | KYRLHNRPRV | PSSTIVNQPIVLMQGLCYIPQEQSSS | QSGSPEGPLH |
| Z_6P00650_001 | 314 | 5.90e-13 | HLQKYRLRMT | SSSNAKNRPAAIGGGGWVHEEQHSAS | LQQSVPQSGS |
| Z_7P00660_001 | 270 | 6.38e-13 | QKYRLHARRV | HNASATADRPVVGEGLWIPQKQYAGS | SRRSASRSGS |
| V_1g58830.1 | 289 | 1.90e-12 | YRLHTSRMVS | GGPLMHHRPVVLSGGLWMLPSEESSS | LSGSPPGPLS |
| 7AzfiHRS2_208.g057980 | 335 | 5.46e-12 | RLNAQRPNRS | SKLNETSPSLVLLGGVWVTHDPNGIQ | HDEREHIYNN |
| Q_112g00420.1 | 269 | 6.83e-12 | YRLHTKRPNS | TTGSCQTPQFILMGGIWVPPSEYASA | ADTDAPPVTV |
| E_N01205.1 | 307 | 1.06e-11 | SHLQKYRLLA | RKLAPKTNTLVGIGSLWMPQQQKESS | KKSNSQFGSP |
| 8AzfiHRS8_08.g011579 | 466 | 3.37e-11 | TRRPIVSSSC | SEAYQQPHHLVLFGGIWVPTSPVEQS | SSPHQQSTSM |
| 2AzfiHRS3_92.g042991 | 268 | 3.88e-11 | NATRQVKMGS | SETTMLPHDVVVMGGICVPNHECLNI | RQQRNSLNFS |

HRS Motif 5

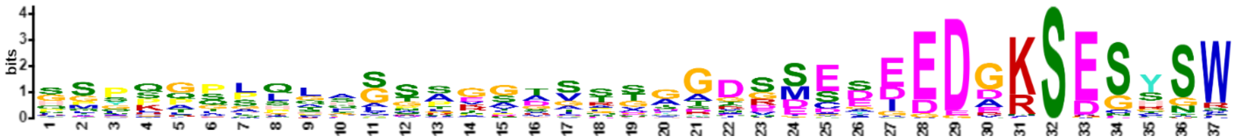

| Name | Start | *p*-value | Sites | | |
|---|---|---|---|---|---|
| N_17G094500.1 | 352 | 2.70e-34 | DSSKTTSSQS | GSPQGPLQLAGNTGGTSTTGGDSMEDDEDAKSEGYSW | KSHIHRSGKD |
| G_G006998t1 | 362 | 1.42e-33 | ESSKGSSSQS | GSPQGPLQLAANTGGTSTTGGDSMEDDEDAKSESYSW | KSHIHKPGKD |
| N_15G143500.1 | 359 | 2.69e-33 | DSMKATSSQS | GSPQGPLQFAGNTGGTSTTGGDSMEDDEDAKSEGYSW | KSHIHRSVKD |
| O_29780.m001324 | 360 | 3.69e-33 | DDSKAAGSQS | GSPQGPLQLAGNTGATSTTGGDSMEDDEDAKSEGYSW | KSQIHRSGKV |
| M_004G144800.1 | 373 | 1.72e-32 | DSSKANSSKS | GSPQGPLQLAVNTGGTSTTGGDSMEDDEDAKSEGYSW | KSHIHRSGKD |
| M_009G106600.1 | 358 | 1.72e-32 | DSSKATSSQS | GSPQGPLQLAVNTGGTSTTGGDSMEDDEDAKSEGYSW | KSHSHRSGKD |
| B_011780 | 322 | 3.11e-32 | SSKPSSASQS | GSPQGPLQLAGTAGGISTTGGDSMEDDEDRRSESYSW | KGHLHKSGED |

| | | | | | |
|---|---|---|---|---|---|
| B_006595 | 352 | 3.05e-31 | PSPKPSTSQS | GSPQGPLQLAGAAGGLSTTGGDSMEDDDDGRSESYSW | KGHHQKSGED |
| F_1031804001 | 353 | 1.20e-29 | DSSKASSSQS | ASPQGPLQLGGAAGGTSTTGGDSMEDDEDEKSESYGW | KSQVHKSVKE |
| F_1003551001 | 334 | 1.53e-29 | DVSKKNISQS | GSPQGPLHLAGSAKGVSGTGGDSMEYDEDDKSEGHSW | KDQLHKPGED |
| N_14G127500.1 | 408 | 2.44e-27 | HLHSTLHHQL | HMYKATSQGHSSPESDVRGTGDRSESIEDGKSESSSW | KAESGENGGE |
| G_G037970t1 | 396 | 1.62e-26 | LHHHAIHHQL | HMYKANSQAHSSPESDVRGAGDRSESIEDGKSESSSW | KGESGDNGGA |
| D_01g108300.2 | 357 | 2.98e-26 | ESSKQSSSQS | GSPQGPLHLAGSCGGTSTTVGDSMDEEDDVKSENHSW | KNHVHTSGKV |
| M_007G039400.1 | 303 | 4.45e-26 | DPSTGNISQS | NSPKGSLHASGSAKATSNTGGDSMEAEDDDKSESHSW | NGALHKPGEA |
| G_G000558t1 | 326 | 3.75e-25 | EHMKANISQS | GSPQGPLLATGSAKDMSSTGGDSMEAEEDEKSDGHSW | RGGVHKQGEI |
| H_62.51 | 358 | 4.53e-25 | VDSSKPSNSQ | SGSPQGPLQLTYTGGTSTTGGDSMEDDGDAKSESYSW | KGHIHKPGKD |
| M_005G134600.1 | 315 | 4.53e-25 | DPVMHNISES | NSPKAPLHGSSSAKAASNSGGDSMEAEDDDKSESHSW | NGVLHHPGEV |
| F_1015462001 | 332 | 5.46e-25 | PHHHSLHHQL | HMYKASPQTQSSPDSDMRGAGDRSESIEDGKSESSSW | KGESGENGGE |
| K_05G011500.1 | 391 | 5.46e-25 | PQHHNALHHL | HMYKAASHGHGSPESDVPGGGERSESIEDGKSESSSW | KGESGENEGE |
| B_006061 | 384 | 1.37e-24 | PTHQLHHSLH | HVYKASSQTHSSPESDGRPAGDGSESIEDGKSESSSW | NKGDSGDQNG |
| K_04G151000.1 | 396 | 1.97e-24 | HENILHHHHF | HMYKTAPQTRSSPVSDVRSGGDRSETIEDGKSESGSW | KAENGEKKGL |
| M_018G074200.1 | 393 | 6.85e-24 | LHHHTLYHQL | HLYKPTAQAHSSPESDVRGTRDRSESIEDGKSESSSW | KGESGENDGG |
| AA_2901G0070 | 315 | 1.15e-23 | WIPPEQSCSQ | SGSPQGPLHFSTSGIAVSSAATVSCEEEDGRSESYGW | K* |
| L_4g086835.1 | 369 | 1.37e-23 | SSKGSSTASG | SPQSPLHLATGSRGGTSPTEGDSMEDDDDEKSESYSW | KSHIHRHGKV |
| K_17G119600.1 | 392 | 1.93e-23 | PPHHNALHHL | HMYKAAPHGQGSPESDVPSNGERSESIEDGKSESSSW | KGENEGERKG |
| O_29807.m000474 | 405 | 2.28e-23 | HHHTLHHQLH | MYNKATSQAHSSPESDIRGTGDRSESIEDGKSESSSW | KAESGENGGE |
| AA_0456G0940 | 317 | 2.28e-23 | WIPPEQSSSQ | SGSPQGPLHFSTSGIAVSSAATVSCEDEDGRSESYGW | K* |
| B_003614 | 387 | 2.70e-23 | QLHHSLHPQQ | HIYKASSQTHSSPESDGRPAGDGSESIEDGKSENSSW | KGDSGENGGD |
| R_2G100176 | 332 | 2.70e-23 | IPQEQGSPQS | GSPHGPLHHLSTSVAAVSSAATASCEEEDGRSESYGW | K |
| O_29676.m001636 | 321 | 3.78e-23 | DSSKQSNSKS | SSPQGPLHGCGSAKGMSSTGGDSMEVEDDDRSVSNSW | NGRQHKQAGE |
| C_7G203800 | 377 | 4.47e-23 | IHHPHPHHQP | HVYKTSSKTHSSPESDTQGAGDRSESIEDGKSESSSW | KGDSGENGGE |
| T_2G013500.1 | 321 | 5.27e-23 | WIPQEQSSSQ | SGSPQGPLHFSTSGLAVSSAATVSCEEEDGRSESYGW | K* |
| N_13G128300.1 | 333 | 7.34e-23 | DPSKPSISES | NSPQGPFHACGYAKGISSTGGNSGEAEDDDKSESHSW | TGRLHKAGEV |
| S_2G016300.1 | 356 | 1.02e-22 | IPQEQSSSQS | GSPHGPLHFSTASGIAVSSAATVSCEEEDGRSESYGW | K* |
| K_02G135400.1 | 366 | 1.20e-22 | SSKVSSSGSG | SPQSPLHLAAGSRGGTSPTEGDSMEDDEDARSESFSW | KSHMHKAGKV |
| K_07G209500.1 | 364 | 1.41e-22 | SSKVSSSGSG | SPQSPLHLAAGSRGGTSPTEGDSIEDDEDARSESYSW | KSHMNKPGKV |
| R_2G173882 | 308 | 1.65e-22 | WIPQEQSSSQ | SGSPHGPLHFSTSGIAVSSAATVSCEEEDGRSESYGW | K |
| AA_0185G0030 | 309 | 3.13e-22 | TTSQHSTSQS | GSPQGPLRLTVSSRGAGDSCEGGEEEEEDGKSESYSW | EMENGAKAPS |
| R_2G124540 | 299 | 5.01e-22 | PPHQEQSSSQ | SGSPQGPLQFSGSGVAATVGGDSSSSDEDDKSEGYSR | KYV |
| H_127.55 | 405 | 5.64e-21 | HHPTLHHQLL | MYKAASAQTHSSPESDIRGAADRSESIEDGKSESSSW | KGDSGGENGN |
| Z_10P28540_001 | 351 | 2.73e-20 | SNSSQQSVSQ | SGSPQSPLQLAGATPVVSVTAGDSCEEEDGKSESFSW | K |
| AA_1535G0300 | 305 | 9.49e-20 | WVTQEESSSQ | SGSPQGPLQFSGSGVAVSVATVGDSYEEDDRSEGYSR | K* |
| AA_3297G0100 | 296 | 2.43e-19 | WVPQEQSSSQ | SGSPQGPLQFSGSGVAVSVDTVGDSYEEDDRSEGYSR | K* |
| M_006G155200.1 | 394 | 2.77e-19 | LHHHTLHHQL | QLFRPTSQVHSSPESDFRGSRDRSESIEDGKSESSSW | KGESGENDGG |
| C_6G095900 | 357 | 6.08e-19 | CGASKASTSQ | SESPQGPLQLMTSGGSTTGDDSMEDDEEDGKSESYNW | KGHA* |
| K_06G213400.1 | 401 | 7.87e-19 | HDYTLHHHHF | HMYKTAAQTQNSPESDIRSSGDRSESIVDGKSESGKW | NGESGEKKGL |
| W_03g55590.1 | 311 | 1.49e-18 | ASQEVSCSQS | GSPQGPLQLSGSGVAVSAATAGDSCCEDDDKSEGYVR | K* |
| N_12G093900.1 | 324 | 4.57e-18 | DPSKPTISQS | NSPQGPFHGCGSLKDVSSTGGDSIEAEDDDVSESHSX | |
| 2PabHRS2_33594g0010 | 388 | 9.48e-18 | QTPSQSLSSP | QGPLQLTGQSSGARGTSVDGGREESVGEDGKSESSSW | KGEEIAKNVE |
| H_6.397 | 321 | 9.48e-18 | NHSGDSSKPS | ISPSGSPQGPLLSHGADSNGADSMETEEDEKSDGRSW | RGRIHRPEEI |
| Z_7P00660_001 | 302 | 3.49e-17 | YAGSSRRSAS | RSGSPQSPLHLAASAHALSITAGDSSEEDGKSESCSW | K |
| W_07g02800.2 | 318 | 4.94e-17 | PQEQSSSQSG | SPEGPLHFSGSGMAGGGSSAATVSCEEEDGRSESYGW | K* |
| V_4g03160.1 | 329 | 9.80e-17 | QYTASQHSTS | KSGGSPMQLTGSSRATAGDSCDGEEEEEDGRSASYGW | GMQLNGTMAS |
| 1PabHRS1_14087g0010 | 427 | 1.23e-16 | QTPSQSQNSP | QGPLQCNGHSSGARGTSADAGREESVGEDAKSDSSSW | KDEDSGETTE |
| PEQU_27343 | 336 | 1.23e-16 | WVSQDQQNAS | QSGSPQGPLQLTCAAQALSMTGGDSCTDDGKSESYSW | K |
| Z_6P34680_001 | 342 | 1.72e-16 | TAPPQKSVSQ | SGSPQSPLQLADARRAISPTAGDSCEEEGGKSECHNW | R |
| K_17G178500.1 | 305 | 2.40e-16 | AQDKCGDDKS | GSPQGPLFLGGSGKGLSSSGRNSMDTEGDEESDCRNW | KATTIVTFVI |
| MpoHRS1_0119s0041 | 442 | 5.17e-16 | HRSQSHSSPQ | GPLQSTSQLSSGARQTSAEMYREDSPGEDGKSESSSW | KGEDDSRGRS |
| Z_4P23210_001 | 335 | 5.76e-16 | HTCSSQQSVS | QSNSPQSPLQLAGSALALSVTAGESLEDDGKSESYSW | K |
| W_12g39640.1 | 347 | 1.36e-15 | STSQSGSPQG | PLQLTVSSSHAVSVTAGDSCDGGEEEEEDGKSGSYSW | EMQNGARASS |
| T_9G080000.1 | 311 | 1.51e-15 | QEQTSSQSGS | PQGPLQFSGSGMAVSAATVGGDGSSSDEDDKSDESYS | RK* |
| V_1g07630.1 | 312 | 1.86e-15 | PQEQSSSQSR | SPEAEGPLQFSSSGVAVSAATVSDSSEEDDRSDGHSR | K* |
| S_1G078900.1 | 303 | 4.26e-15 | QEQSSSGSPQ | GPLQFSGSGVAISTATVGGDSSGSSSDEDDKSEGYSR | KCV* |
| N_06G060000.1 | 409 | 4.72e-15 | MHHHHHTLHQ | LHMYKATSQAHSSPESDVRGGDRSESIEDGKSESSSW | KSGENGGDRK |
| D_12g006800.1 | 390 | 1.06e-14 | HPGSHMYKPS | PKAHSHSSPESDARGTGQHGDRSSESIEDGKSESYSS | ENGGERKVVV |
| SfaHRS1_10s0233.1 | 462 | 2.13e-14 | EPQKSSLLSQ | SSPQGPLQLSSQLSGGAQELSNEDSGGGDVKSDSTSW | KSCQLDVIEG |
| Z_2P13380_001 | 329 | 2.35e-14 | PVIWVREEQY | TSSSQQSISQAGSPTSPLQLTAADSSEEDGKSESYNW | K |
| 7AthHOH6_AT1G49560 | 291 | 2.86e-14 | GGESLKRSNA | QSDSPQGPLQLPSTTTTTGGDSSMEDVEDAKSESFQL | ERLRSP |
| SfaHRS2_4s0229.1 | 518 | 3.47e-14 | EPQKSTSVSQ | SSPQGPLQLTSQLSGGAQEPSLDDTGGEDVKSDSTSW | KGHQQEDEGK |
| S_8G147101.1 | 353 | 3.47e-14 | QSQSGSPQGP | LQLTTVSSRAMSATAGDSCDGDGDEAEGGGRSESYGW | EMQQQQQQH |

| | | | | | |
|---|---|---|---|---|---|
| E_H01532.1 | 427 | 3.83e-14 | QLHLYNKPPP | LTQRGRNISPETDVRGGGGAGDPSESIEDGKSESGSW | KADSGGGENG |
| D_02g090400.2 | 317 | 3.83e-14 | DENGESSKNN | GTQSGSPEGPLHFTGSGSAKGVSVNEEEDNKSESYNW | NGQLHKSIEG |
| SfaHRS3_1s0189.1 | 467 | 1.20e-13 | CPQSQQAVSQ | SQTSPQAPLQVTNPVSSGVGYQYDSGGEEGKSDSTSW | KGRPLETGGG |
| Z_6P00650_001 | 342 | 1.20e-13 | HEEQHSASLQ | QSVPQSGSPESPLRLPVASTTGDSCEEEDRKSESHNC | V |
| PpaHRS1_4480V3.1 | 451 | 1.32e-13 | TSQSQSSPQG | PLQVTSQLSGATQGPSSAAYHDESAGEEDAKSESASW | KCDNPKGEEL |
| P_2G0120600 | 328 | 3.05e-13 | LGGLWAPPEN | FPLLKQATGSPQGPLHLANAGEDSGEEEDGRSESYSW | KGHLQRPAGE |
| E_B00308.1 | 352 | 5.76e-13 | SQEQCGESSK | QNNCQSGSPEGPLQLGGSSVGYSSEYEDDDKSESHTH | LSVRDS* |
| L_5g017980.1 | 287 | 9.01e-13 | SEGNFSESVS | PQGPLTPLLLGGSAQGLSSHGRNSMDAEDEQSDCRNW | KSD* |
| AA_1128G0130 | 327 | 1.29e-12 | HLQTSGSSQS | HPGSPEARLQLSESSRAVSVTAGDSYEEDRRSESCSV | ETQHRTTS* |
| J_3835011 | 270 | 3.07e-12 | RETQSLISLS | RSGSPQSPLVGRGLFNNNGSHISEDDEEEEKSDGRSW | RGESDKKRQV |
| PpaHRS2_8580V3.1 | 458 | 3.35e-12 | TSQSQNSPQG | PLHVTSQLSSATQGPSSEVYHDESPGEEEAKSEGTSW | QREDLKSEEP |
| E_H00466.1 | 396 | 5.58e-12 | SPQQHLRDDM | AGGSTSAAQVGSATTTEEGESIEEADEDDQKSKSCSW | KGKSAEHPF* |
| R_2G171468 | 322 | 7.19e-12 | EQSSGSQSGS | RQGPLQFSRAGMAVGGGDDSSSSDDDEDDDKSEDGYS | LKCV |
| 1AtrHRS1_122.65 | 287 | 9.24e-12 | YAPTHRTYCD | QMALAHMNPSHPNGSVIEQRGGSESIEEDGKSESGSA | ENEVRDAGPA |
| T_3G370000.1 | 341 | 1.00e-11 | SQHSTSQSGS | PQGPLQLTVSSRAMSATAGDSCDGDEAEGGRSESYSF | GMQQQHGTKA |
| AA_0930G0300 | 334 | 1.19e-11 | SGSSQSRPGS | PEARLQLSESSGRVVSVTAGDSYEADEEDRRSENCSV | EAQHRTPS* |
| J_3862509 | 243 | 2.48e-11 | PAKILTASDQ | HGLSRSDSPQGPLVDRGLFSNNGHSSEEEEKSDGRSW | KSESRRKRQG |
| K_11G058600.1 | 290 | 1.03e-10 | MSQSGSPQGP | LTPLILGGGGGGSAKGLSSPGQNSVDGEDEQSDCRNW | KGGLHHHQLE |
| 2AtrHRS2_033.259 | 350 | 2.40e-10 | FQLVKKTASS | QSGSPQGPFQYPGKARTSVNGGDSGGEEDGLSDEETN | * |

HRS Motif 6

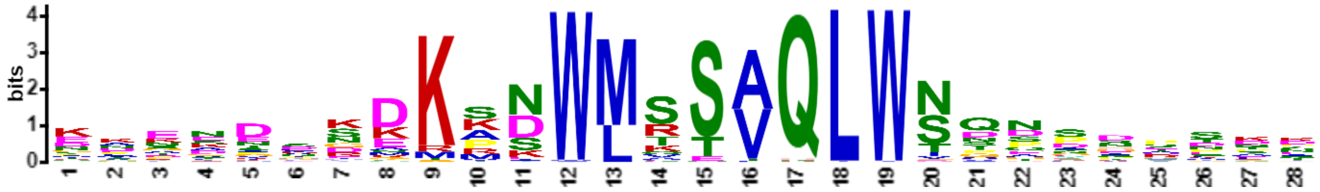


| Name | Start | *p*-value | Sites | | |
|---|---|---|---|---|---|
| N_15G143500.1 | 116 | 8.76e-26 | DDDDEQDGRI | NKEKDSRDKKNWMSSVQLWNTNDHPSTN | YIFDKKQSLK |
| N_17G094500.1 | 114 | 8.76e-26 | DDDDEQDGPI | KKEKDSKDKKNWMSSVQLWNSNDHHSTD | YIFDQKQNLK |
| O_29676.m001636 | 98 | 7.06e-25 | KGNSCRNEER | ELENDMIDKKNWMSSVQLWNNNNHTNNN | NFDSENQESK |
| AB_07.3042 | 104 | 7.06e-25 | KSKLDEERGV | KSEKDCKDKMNWMSSAQLWSDNYSDDND | HINNNDIKDS |
| Z_4P28210_001 | 109 | 2.17e-24 | KSKVEDDGGV | KVETDCRDKMNWMSSVQLWSDNYSENDD | EEKIVSNEQD |
| Z_2P13380_001 | 110 | 4.91e-24 | KSKVGDDGGI | KVESDCKDKMNWMSSVQLWSDNYIENND | DDKAIAKEKD |
| B_006595 | 109 | 1.08e-23 | KRNSDEDGAP | KKENDCRDKKNWMSTAQLWSSNGYSNKD | NCYDRNKKST |
| 2AtrHRS2_033.259 | 116 | 2.33e-23 | KAQEERENEA | KAERDCKDKMNWMSSAQLWSDNSDNSEE | KKEKLDEYEA |
| O_29780.m001324 | 117 | 2.33e-23 | DDDGDQDGPI | KKEKDTKDKKNWMSSVQLWNTDDHPSTH | NILETKQNLQ |
| Z_7P00660_001 | 91 | 2.33e-23 | KSKVEEDGRV | KAETDCKDKKNWMSSAQLWSDNHSDNDD | DDKAHANIIS |
| F_1031804001 | 109 | 2.99e-23 | KKDCDEEGGA | KEEKECKDKKNWMSSVQLWNSDDSASTD | HIYDKKQDSK |
| K_07G209500.1 | 116 | 7.98e-23 | SEEEEENDDD | KDDNECRDKRNWMSSVQLWNNNTTTTTT | NNNNNASDRK |
| L_4g086835.1 | 118 | 1.01e-22 | ENKNNDRDKN | NNNECSKDKKNWMSSVQLWNNNTTTSNN | NVSDHHHHHK |
| Z_6P00650_001 | 109 | 5.11e-22 | KSKFEEKGAV | NPEADCKEKRNWMSSAQLWSDNSNRSNY | EENENEKEVL |
| M_004G144800.1 | 131 | 2.33e-21 | HGDDRESDGL | IKEKDSKDKKNWMSSVQLWITDDHHPST | DYLFDAKQSF |
| K_02G135400.1 | 116 | 3.54e-21 | EEEEEEENDD | DDDNECRDKRNWMSSVQLWNNNTTTATT | NNNNPSDRKQ |
| M_018G074200.1 | 100 | 3.54e-21 | PLKTPTSEAL | EKTTNISDKANWMTTAQLWSQDSNESKP | QTTLTSPKQT |
| N_13G128300.1 | 106 | 4.35e-21 | KENSGGDGGG | NMGNDLSDKKNWMSSVQLWNTNNINSDS | KQHDSKSETK |
| K_20G009800.1 | 111 | 5.34e-21 | KKECDQRKEI | EKEKECRDKKNWMSSFQLWNNDDKADNN | NNAYECDKKH |
| E_N01205.1 | 135 | 7.99e-21 | PLKKSLPKKE | DDKIDSVDKMNWMSSVQLWNSDNHPNTE | NKKKWGGVDL |
| M_007G039400.1 | 90 | 9.76e-21 | ELNDLIPLKG | NSNEDGNDKKNWMSSVQLWNTNNHLDCK | KQDSKSEPKQ |
| M_009G106600.1 | 120 | 1.19e-20 | IDDHHENDGL | IEEKDSKDKKNWMSSVQLWNADDHHPST | DYLFDPKQNL |
| AA_1128G0130 | 111 | 1.19e-20 | KVEDKAGAVM | KLEADANDKMDWMSSAQLWSCGNHSDTT | DTSKDGSKGR |
| G_G006998t1 | 126 | 1.45e-20 | TTKKDKDPNN | NNYNINKDKKNWMSSVQLWNTDDDDYRS | TDHKLDTKRN |
| N_14G098400.1 | 111 | 1.45e-20 | RNKNSSTNKD | KNSSDHKKKSDWLRSAQLWNQSPDPTPK | EDVSRKAVET |
| M_006G155200.1 | 100 | 1.76e-20 | PLKTSTSETP | EKTSNISDKANWMTTAQLWSQESNETKP | QTTLTSPILT |
| Z_10P28540_001 | 109 | 1.76e-20 | KEKFEEEGRE | KVEKDCEAKMNWMSSAQLWSDNPNRNNC | KDNKNEKKVT |
| W_12g39640.1 | 139 | 2.58e-20 | SAASAAAEGV | KAEADANDKRSWMSSAQLWSCGSHTSTS | TSNGGSVKKQ |
| 3AthHOH3_AT1G25550 | 114 | 5.46e-20 | EAEKTEMMTN | ENNDGDKKKSDWLRSVQLWNQSPDPQPN | NKKPMVIEVK |
| M_005G134600.1 | 90 | 5.46e-20 | ELNALVPLKG | DSNEDGNDKKKWMSSVQLWNTNNNINLD | CKNQDTRSEP |
| H_1.213 | 100 | 7.87e-20 | EDEQESHKAS | TNTNTDKKKSDWLRSVQLWNQSPDPPPK | KDVPKKTTTA |
| N_06G071500.1 | 127 | 7.87e-20 | NNSGSSNNKD | KNSSDHRKKSDWLRSVQLWNQSPDPSPE | EDVPRKAVVT |

| | | | | | |
|---|---|---|---|---|---|
| K_06G213400.1 | 105 | 1.13e-19 | LKHPNSQESA | EKTSNISDKANWMTSAQLWSQASEGTKP | QSTITSLPKE |
| K_04G151000.1 | 103 | 1.13e-19 | IKHPNSQEST | EKTSNISDKANWMTSAQLWSQASEGTKP | QSTITSPKNG |
| K_01G086700.1 | 108 | 1.13e-19 | DEDEEHFHKH | KKTATDKKKSDWLRSVQLWNPNPPPTKE | DVVVPRKTDV |
| Z_9P26570_001 | 108 | 1.13e-19 | KRKREDEGGL | KPEADCKDKMSWMSSAQLWSVNSREDKS | DGDRNVTEER |
| H_62.51 | 116 | 1.61e-19 | EEDEEDRAAA | RKKEDCRDKKNWMSSVQLWNSEDLPSST | VNHRSFDPKQ |
| N_14G127500.1 | 100 | 1.61e-19 | PLKNHASETL | DRSSNISDKANWMTTAQLWSQDSNETKP | QTTLTTSPKE |
| N_12G093900.1 | 101 | 1.61e-19 | VSVKEDYSGA | DGGNDMRDKKNWMSSVQLWNTGDVNSDS | KQHDSKSETK |
| AA_0185G0030 | 113 | 1.91e-19 | RKAEEKVGGV | KTEVDANDKRSWMSSAQLWSCGSHNSTS | NSNGGSVKKQ |
| M_010G128900.1 | 119 | 2.28e-19 | DNKRNKSNSS | ISNNDHKKKSDWLRSVQLWNQSPDPPQK | QDLPRKAAVT |
| Z_6P34680_001 | 109 | 2.28e-19 | RSRCEEEGGA | KLEVDCKDKKKWMSSAQLWSDYSSDDNR | NDDDQSVADE |
| S_8G147101.1 | 131 | 2.71e-19 | AQDGAAVKAD | DAEADANDKRSWMSSAQLWSCGSHDDST | ANTNGVAAAA |
| AA_0930G0300 | 114 | 2.71e-19 | KVEDKAGAVM | KPEVDANDKTDWMSSAQLWSCGNHSDTT | DTSRAGSKGR |
| O_29807.m000474 | 100 | 3.80e-19 | PLKNSTSETL | DKSSNMSDKANWMTTAQLWSQESNEIKP | QTTFNSPKET |
| P_2G0120600 | 108 | 3.80e-19 | AKLEEAEGGV | RDEKDFRDKVNWMSSAQLWSDNYSANTS | STTSTTVASS |
| H_127.55 | 102 | 4.50e-19 | PLKNPSSEST | EKLNTISDKANWMTSAQLWSQAGNETKP | QPTTIISSSL |
| K_17G119600.1 | 101 | 5.32e-19 | MKHLASDQSS | EKATNMSDKANWMTSAQLWSQASEGTKQ | QPTITTPKES |
| B_011780 | 109 | 6.28e-19 | KRNSDEDVRA | KKEKESRDKKNWMSSAQLWSSSGYSNQD | TSYDRNQKSV |
| F_1015462001 | 100 | 6.28e-19 | PLKHSTPKGS | EKTSNMSDKANWMTSAQLWSPAGDETKQ | QSTLTSSKEP |
| C_3G045700 | 111 | 7.40e-19 | DKSNIDESNV | NNNNNIGKKVDWLRSVQLWNQNPEVPPP | SQQQEYHQQQ |
| H_6.397 | 104 | 8.71e-19 | KRISEEDGAA | NRERDGCDKKNWMSSVQLWSNNVNPDPE | QHSISRLKST |
| E_B00308.1 | 124 | 1.02e-18 | DKVEGNNKEK | DKDVSSKEKMNWMSSVQLWNSDANNQNP | DTDFNNNKLS |
| P_0G0029600 | 49 | 1.20e-18 | NRKFEEAGRS | KDEEDRREKANWMSSAQLWSDTSSTSNS | GDSKNRSRQR |
| Z_4P32500_001 | 105 | 1.20e-18 | LLPVESRVEE | DGEVKVDDKMNWMSSAQLWSDNYGENDI | SHENHEVMIC |
| K_02G098800.1 | 108 | 1.41e-18 | DDDEEHSHKH | TKTTTDKKKSDWLRSVQLWNPNPPPTKE | DVVPRKTNVV |
| G_G037970t1 | 101 | 1.65e-18 | PLKNSSSENS | EKSQNISDKANWMTTAQLWSQAGNETKP | QSSITSPKET |
| G_G000558t1 | 105 | 1.65e-18 | KRNTEENRRA | NMEDDGGDMKNWMSSVQLWNSNFNNVDH | NKKANAVQEL |
| K_05G011500.1 | 100 | 1.94e-18 | PMKHLASESS | EKATNMSDKANWMTSAQLWSQASSEGTK | QQPPITTLKE |
| B_012707 | 96 | 2.26e-18 | SEEEEELNKP | NNKDRNDKKADWLKSVQLWNQAPDPPLK | QDLGPKLSVV |
| N_06G060000.1 | 101 | 2.26e-18 | PLKNSTSETL | DNSSNISDKANWMTTAQLWSQESNETKL | QTTLNTFPKK |
| F_1011942001 | 95 | 2.64e-18 | QSHQPNDNKD | KNNDKSGKKSDWLRSVQLWNQTPDPPVK | EDTPKKIPSM |
| O_28883.m000751 | 134 | 3.08e-18 | DSHKPNKSIR | DINNDQKKKSDWLRSVQLWNQSSPDSEP | PKEDLPRKAA |
| Q_9g01150.1 | 118 | 3.08e-18 | LDQNGATQNS | EKETRDRNKMDWMSSAQLWSDNYSDEAI | SNNNHDETNN |
| Z_9P09630_001 | 105 | 3.59e-18 | RRKCEEEAGV | KLEADYEDKKNWMSSAQLWGVNSSENND | EDDKSITDER |
| T_3G370000.1 | 121 | 4.18e-18 | KAAAPEGGAV | KAEADANDKRSWMSSAQLWSCGGGHDGS | SSSTATNNGA |
| F_1003551001 | 105 | 4.85e-18 | KGRSEGDEGA | EMGKDSSDKRSWMSCAQLWSCNRSSDNN | NKKSVLEFQQ |
| K_07G178500.1 | 111 | 4.85e-18 | KKECDQREES | EKEKECRDKKSWMSSFQLWNTDDKADIN | NNAYECDKKQ |
| W_07g02800.2 | 89 | 4.85e-18 | VEKKGGDRKE | DEEDAAGDKSNWMSTAQLWTGNSGGPDA | AAADPEKQDK |
| T_9G080000.1 | 90 | 4.85e-18 | AAKRGEDLEL | EMKADDGDKKKWMSTAQLWVDSDAKSKS | EKEKRSEMTS |
| K_01G183700.1 | 116 | 6.54e-18 | QDPPLKTSSG | GNENENSEKKNWMSSAQLWSTQKSKSRN | EEDDRSVPAN |
| D_01g108300.2 | 111 | 7.57e-18 | SSREDTKAEI | TKDKDSREKMSWMSSVQLWNSESHCENT | DEAINKQQSK |
| Z_4P23210_001 | 108 | 7.57e-18 | KSKVEEGGGI | KEETEYKDKRNWMSSAKLWNDNCSQNNN | DSKKHDKIKF |
| 3AtrHRS3_093.40 | 82 | 8.76e-18 | TTQAPAGEKK | RKPEDQNDKPDWLRSVQLWNSDADPSRN | EDAGLNPKLG |
| Z_10P27180_001 | 80 | 8.76e-18 | QQLQERVPLK | YMNVDGSDKANWMVSAQLWSPPDDAAKQ | QAAPPPKEAD |
| B_001193 | 108 | 1.01e-17 | DEESNKPKNE | DINEKPGKKADWLKSVQLWNQDPDPSLN | QDLSRKPSVI |
| C_6G095900 | 116 | 1.01e-17 | ENFVPSKTNS | SSEIDLRDKKNWMSSAQLWTSNGYSSKT | DDEKPKSIVD |
| J_3852516 | 97 | 1.01e-17 | GEHESPTPEE | IGNNVDKKKSDWLRSVQLWNPSPDTNEV | SNPERVVGKK |
| AA_0456G0940 | 88 | 1.01e-17 | KKKVHREEED | KEDGDAGDKSKWMSTAQLWTGDSGRDDA | DSEKQDKERS |
| K_20G186500.1 | 114 | 1.17e-17 | QHSHKQRVSK | ENNNSDKRKSDWLRSVQLWNPDPPAEED | VSKIVCGVEL |
| K_10G204200.1 | 116 | 1.17e-17 | QHSHKQRVSK | ENNNSDKRKSDWLRSVQLWNPDPPAEED | VNKKVPGLEL |
| E_L01092.1 | 92 | 1.17e-17 | QDQSEEIRNI | GNFDINSKKSDWLKSVQLWNQTPDPPSK | QDSPRKSDVV |
| D_05g009720.2 | 123 | 1.17e-17 | SFNNNNNNST | SSKDAKNKKSDWLRSVQLWNQTSDPTPK | EELTPKKVSV |
| K_11G058600.1 | 89 | 1.35e-17 | QDPPLKTSSG | GNKNESSEKMNWMSSAQLWSTQKTKSRN | EEDDRSVPAN |
| G_G011857t1 | 114 | 1.35e-17 | KDHDTNTKEK | NVAAADKKKSDWLRSVQLWNNNQSPDPP | LQEGAGKSGS |
| D_02g090400.2 | 111 | 1.35e-17 | KGNSDEIGRV | KKSNDLSDKKNWMSSAQLWSTPVQYESF | NLQNLKRSGV |
| AB_01.3080 | 79 | 1.35e-17 | SDEENNHKNS | GNSGGSDKKPDWMRSVQLWNQQPEPKGD | VVMRPIAVNA |
| 1AthHRS1_AT1G13300 | 113 | 2.06e-17 | DDEHGNHDPD | NDSEDKNTKSDWLKSVQLWNQPDHPLLP | KEERLQQETM |
| J_3836112 | 111 | 2.06e-17 | DDEHGNHDPD | NDYEDKNMKSDWLKSVQLWNQPDPLLPK | EEGTQEKMVD |
| L_1g093080.1 | 108 | 2.06e-17 | YDQHFHHKQQ | KLSLDNKKKSDWLRSVQLWNSDPSSEED | VTKKAVPVLE |
| AA_2901G0070 | 85 | 2.06e-17 | EAEKKKGDRG | EEEDGAGDKSKWMSTAQLWTGNSGRDDA | DSEKQDKERS |
| L_5g017980.1 | 89 | 2.37e-17 | KEKVRLMKMD | DLKENCDNKKNWMSSAQLWTNETKSKNE | GDDDRTVLHK |
| J_3871943 | 112 | 2.72e-17 | EDEQGNHDPE | NDSEDKNMKSDWLKSVQLWNQPDPLLPK | EEQSQQMMET |
| V_1g07630.1 | 87 | 2.72e-17 | AGKRSEVRDA | EAKAEDNDKRKWMSTAQLWVDNRGSDSD | SVVQKEQKKE |
| J_3864545 | 100 | 3.58e-17 | ASDGKLDVGE | GAKVESDNKKNWMSSAQLWISNPNSQLQ | STNEKEDRSV |
| PEQU_27343 | 96 | 3.58e-17 | DEYKEEEKAM | NLEKESKEKMEWMSSAQLWSDNSSHESI | DGKEDSQEVF |
| C_7G203800 | 102 | 4.68e-17 | PLKHSSSEGS | EKTSSISEKSSWMVSAQLWSQSNDGNKQ | QTITSSPKET |
| J_3835011 | 100 | 4.68e-17 | ASNGKLDVGQ | GSKLENDYKKNWMSSAQLWISNSNSTNK | EEDRCVTQTP |
| 1AtrHRS1_122.65 | 38 | 5.35e-17 | PIKRSSSEGS | ETTNMTSDKANWMTSAQLWSDQSFPSSQ | KPAFNGKQRN |

| | | | | | |
|---|---|---|---|---|---|
| E_K00828.1 | 106 | 5.35e-17 | FKSEGGNGRA | KASNDSRERKNWMSSVQLWTTPVQYETN | YLTRNQDSIL |
| J_3843187 | 79 | 6.97e-17 | DYEHGTHDPD | NDSEDKNMKSDWLKSVQLWNQPEPILPK | EERLQPKIIE |
| M_008G117500.1 | 115 | 6.97e-17 | NSKRNKTSIS | SGNNDHKKKSDWLRSVQLWNHSPDLPQK | QDLPRKAAVT |
| E_H01532.1 | 111 | 9.05e-17 | NNSCNNNISD | KEANLSADKANWMTSAQLWNQESGEEEE | EEEETKPPPS |
| S_1G078900.1 | 89 | 9.05e-17 | EAAKRGGGDA | DAKADDGDKRKWMSTAQLWVDSEAKSDE | SDKEHQSEIT |
| E_E00311.1 | 109 | 1.33e-16 | EEEQESKKLK | NDDDSSNKKSDWLRSVQLWNNQTPDLSK | EDSPRKVAVT |
| V_4g03160.1 | 131 | 1.33e-16 | SSPSPAAKRK | QAEADASDKRSWMSSAQLWTCGSHSNGG | IRKQQAQKLS |
| R_2G171468 | 87 | 1.33e-16 | AAKRGEYDDG | AATVDDGDKRKWMSTAQLWVDSDAKSDE | SDKEQQSEIT |
| R_2G124540 | 87 | 1.33e-16 | DEAAKRGGDA | EAKADDGDKRKWMSTAQLWLDSDAKSDE | SDKEQLSEIT |
| Z_3P21090_001 | 80 | 1.51e-16 | SFKSPSSGAV | AAAIGSDRKPDWLRSVQLWNQEPDTSLK | EPPKKPIAVS |
| T_2G013500.1 | 89 | 1.71e-16 | KARSDRKAAA | EEDGAAGDKSKWMSTAQLWTGDSGREDD | SESEKQDKGR |
| X_31310.1 | 87 | 1.94e-16 | EEESALHESK | KRESSCDRKPDWLQSVQLWNQHPDLTFP | VEPPRKPIAV |
| AA_1535G0300 | 79 | 2.49e-16 | AGKRSGNGDA | DAKAEDGDKRKWMSTAQLWVDSRGSDAD | SEKEQESETT |
| B_003614 | 97 | 2.82e-16 | PLKHSGSEGS | EKAASISEKVSWMTTAQLWSQSSDGINK | EQATVASAKE |
| 4AthHOH2_AT1G68670 | 106 | 2.82e-16 | EEEEDGEHES | SPELVNNKKSDWLRSVQLWNHSPDLNPK | EERVAKKAKV |
| Q_33g00510.1 | 99 | 3.18e-16 | PIKDDKEMMV | AEKDCKDNKMNWMTSVQLWKNDGDDTNI | CSGQNKVHRQ |
| AA_3297G0100 | 75 | 3.18e-16 | AGKRRGDGDA | EAKAEDGDKRKWMSTAQLWIDSRVSDAD | SEKEQESEST |
| B_006061 | 101 | 3.59e-16 | LKHSASDQGL | EKASTASEMASWMTTAQLWSQSSDGSKQ | QPATAASAKE |
| D_12g006800.1 | 117 | 3.59e-16 | ELGDEKLVAN | NTSTIVDNKANWMTSAQLWSPPSDHQET | KQQVQISTTC |
| Y_30s1094441g001 | 101 | 3.59e-16 | IEEEKGGSME | MDQERSDNKPDWLRSVQLWSQDPDHSLP | KAEPPKKPIA |
| S_2G016300.1 | 129 | 4.57e-16 | SDRSLPAAEE | DEDGAAGDKSKWMSTAQLWTGDSGREDA | ESEKQDKGRS |
| AA_2274G0070 | 124 | 6.53e-16 | LKNIGIDEAD | KPGNPPSEKASWMVSAQLWNGPAADTAA | NGPQTPKEHS |
| AA_1881G0390 | 125 | 6.53e-16 | LKNIGIDAAD | KPGNPPSEKASWMVSAQLWNGPAADTAA | KGPQTPKEHS |
| R_2G173882 | 85 | 8.25e-16 | MKEEAGKAVA | EEDGAAGDKSKWMSTAQLWTGDCGREDA | EPEKQDKERS |
| W_03g55590.1 | 90 | 9.26e-16 | GKRRKDGDDA | EAKAEDGDKTKWMSTAQLWVDSRGSDAD | SENDRRSGST |
| 2AthHOH1_AT3G25790 | 117 | 1.17e-15 | DEEHQSHETD | IDFDDKNMKSEWLKSVQLWNQSDAVVSN | NRQDRSQEKT |
| L_4g113140.1 | 103 | 1.17e-15 | QLTSSETLEK | TTNNNVCDMANWMTSAQLWSQTSELGTK | QQQNSTKENN |
| S_3G046800.1 | 128 | 1.31e-15 | NIGIDAAAAD | KMGNPPSEKASWMESAQLWNGPAAATAA | DMAAKGPQTP |
| 1PabHRS1_14087g0010 | 92 | 1.64e-15 | NKLEENHVAK | KARFNNGDKPNWMISANLWNPDPETIDS | RKGGGISMEE |
| Y_30s723941g001 | 85 | 1.64e-15 | TEEEKGGFME | MEQERSEKKPDWLRSVQLWNQETDHSLP | KGDPPKRPIA |
| L_5g054300.1 | 102 | 1.83e-15 | ENDEEDDDDE | EQHSSHKMKSDWLRSVQLWNPNPSSAKE | DVPRKTNVVE |
| V_1g58830.1 | 83 | 2.29e-15 | KVESNGGDHR | DEAEGSGDKSSWMSSAQLWIGGGSSKAP | EKEGRSSAPE |
| Q_208g00340.1 | 105 | 2.86e-15 | NSNNGDGVGT | GTGISNMEKSSWMTSVQLWTNDSTTSEA | TVTPKPATES |
| P_9G0028800 | 91 | 2.86e-15 | LKEEMTQRRS | KKKQEDDGRMNWMSSTQLWCNNSSTTTA | STGGRENKSS |
| Y_30s1025391g002 | 101 | 2.86e-15 | LKHASIDGSE | KAPNAPSEKASWMVSAQLWSAANDATRN | QPMAAPPLKE |
| Z_6P05330_001 | 101 | 2.86e-15 | LKHMNMDGSD | KDPSAHSEKASWMVSAQLWSPPVDAAKQ | QPVPPPKETE |
| 5AthHOH4_AT2G03500 | 104 | 3.96e-15 | VLEEFIPLRN | QPEKTNNKGSNWMTTAQLWSQSETKPKN | IDSTTDQSLP |
| AB_10.37 | 79 | 4.42e-15 | SDEENNHKNS | GNSGGSDKKPDCMRSVQLWNQQPEPKGD | VVMKPIAVNA |
| R_2G348238 | 128 | 4.42e-15 | KNIGIDAAAD | KMGNPTSEKASWMESAQLWNGPAATADV | AARGPQTPKE |
| Z_8P18270_001 | 87 | 5.47e-15 | EEEKGETRKS | AAAIGWDRKPEWLRSVQLWNQEPDTDLK | EPPKKPIAVS |
| R_2G100176 | 103 | 5.47e-15 | ASAAADDEEE | EADGATADKSKWMSTAQLWTGDSGREDA | ESEKQDKGRC |
| Z_11P13410_001 | 89 | 6.09e-15 | SVDGAAKKSG | SAATRADEKPDWLRSVQLWNQEADAFGK | VEPPTKPIAV |
| J_3875351 | 117 | 6.77e-15 | DVEHESHDTG | IDFVDKNMKSEWLKSVQLWNQPEAVPVS | KSERSQQETQ |
| J_3853942 | 104 | 8.36e-15 | VLEEFIPLRN | TPEKENNKGNNWMTTAQLWSQPETKPKN | IDPTTDQSPK |
| J_3831280 | 103 | 8.36e-15 | NREHESPREV | DKSDVDSKKSDWLRSAHLWNHHSQDPDM | TVVVAKKARV |
| J_3839360 | 116 | 9.28e-15 | DDEHESNETG | LDCDDKNMKSEWLKSVQLWNQPDSVLSK | KLERSQQETE |
| T_5G124700.1 | 124 | 1.03e-14 | KKIGIDAAAD | KMGNQPSEKASWMESAQLWNGPGAALAA | ADTAAKGPQT |
| C_3G398400 | 70 | 1.14e-14 | VELISSTETR | KLYSPKEKKVDWLKSAQLWNHDFSDQKD | KPSQNTIGIE |
| AB_06.1057 | 135 | 1.27e-14 | EENNNNHGNG | WIGDGFDKKPDWLRSVQLWNPQAEPPKG | DIVRRPVAVN |
| W_01g08160.1 | 115 | 1.27e-14 | VVAADKAAAA | GGNSVSSEKASWMVSAQLWNAPASASAA | DTAAKGPQTP |
| X_9781.1 | 101 | 1.91e-14 | VKQMSVDGLE | TPNGPSSEKASWMISAQLWSNPAESKPQ | QAPKGGGDVS |
| 6AthHOH5_AT4G37180 | 108 | 2.33e-14 | MASNGKFDDV | ERAKPETDKKSWMSSAQLWISNPNSQFR | STNEEEEDRC |
| P_18G0031200 | 103 | 3.15e-14 | NSSTPEADDE | ESGDTAVRKRDWLKSVQLWTPHPSEEDN | RGPWIPVSEK |
| Q_30g00060.1 | 100 | 4.24e-14 | VKKNCTGDDK | SDNSEPLEKANWLASVQLWTQSNNEVPS | SPKRWNTGGA |
| AA_5036G0050 | 114 | 4.24e-14 | EEAETPERRS | PPAENKKATPNWLQSVQLWSQEPQQSSP | DKELPCKPVA |
| V_3g10730.1 | 113 | 5.69e-14 | KKDESERARP | PTPETKKAMPDWLQSVQLWSQEPQQSCP | NKELPCKPVA |
| P_8G0062400 | 129 | 6.27e-14 | EEDSFSTSKR | KNQDFSEKKPDWLRSVQLWSPLPDPHHD | EDDETSRQTI |
| K_17G178500.1 | 107 | 8.38e-14 | TNSEANGSLM | IVGNESSDTKNWMNSVQLWNVETKQRNE | EGDLFVPSNP |
| T_1G146600.1 | 115 | 8.38e-14 | KAAETHGRQS | PPTEANKAMPDWLQSVQLWSQEPQQQPS | SPRKELLCKP |
| R_2G142748 | 74 | 1.01e-13 | TAAASSSRRL | PHPETKKAMPDWLQSVQLWSNQQQPSAS | PPQHQDELLL |
| R_2G159119 | 145 | 1.01e-13 | TAAASSSRPL | PHPETKKAMPDWLQSVQLWSNQQQPSAS | PPQHQDELLL |
| V_2g04810.1 | 135 | 1.48e-13 | SKNIDAADQN | QQAASENPKASWMVSAQLWNGPPPSSAD | APQTPKERSE |
| R_2G016370 | 145 | 1.96e-13 | TAAASSSRPL | PHPETKKAMPDWLQSVQLWSNQQQPSVS | PPQHQDELLL |
| MpoHRS1_0119s0041 | 134 | 2.15e-13 | EEVEKVSKRS | DEGRDADHRPSWMAEAQLWSQPSTRSDA | WKEKESSTED |
| S_8G036900.1 | 146 | 2.15e-13 | ALASSRRPLL | PQPETKKAMPDWLQSVQLWSNQQQPPPQ | QQHQDELLLP |
| J_3862509 | 84 | 5.89e-13 | AIERLKEEAS | SVKLEIDNKKNWMNSTQLWISKTNPQLP | STNGEEDRCV |
| SfaHRS1_10s0233.1 | 143 | 6.45e-13 | DHDQEGDGSE | CPGVVDIEKMTWMTEAQLWTQQLEPSSE | LHCENDRMSM |

| | | | | | |
|---|---|---|---|---|---|
| Z_6P09870_001 | 87 | 9.21e-13 | SEGSGEAKKA | AMVGRLETKPDWLRSVQLWDQQPDTVLK | VEPPKKPMAV |
| J_3875404 | 98 | 2.03e-12 | VTRPVLEEFI | PLRNQPEKANNWMTTAQLWSQPETKPKS | IDQTTDQSPK |
| SmoHRS2_66160 | 67 | 5.17e-12 | KTLEESKRDE | KACDPDLDKPSWMAQVQLWSEAKQIKTQ | VEIYSKHAVP |
| P_27G0017800 | 116 | 6.12e-12 | VAAEAPEKSA | AADDVLSEKTSWMTTAQLWGTAEGAAAV | QRTVTPPEAT |
| Q_178g00480.1 | 95 | 2.88e-11 | KLNEDDDDDE | GDRLEKMSRCSWLKSAQLWNNDKKTLSS | PNKNENKFLL |
| SfaHRS3_1s0189.1 | 146 | 8.60e-11 | GDEEGDRQVE | RTRGIDIGRPAWMAEAQLWTQHAAIAEA | RNEKDLSPED |
| PEQU_26724 | 79 | 8.60e-11 | PTSSSSPSDK | SLKLAGDSKPDWLRSAQLWAPSTVAGVE | TSPEESSKLR |
| SfaHRS2_4s0229.1 | 185 | 2.87e-10 | ELGDQSDSCL | ATRGIDIGRKAWMAEAQIWTQQSEPMEV | HHEKEISSPE |
| PpaHRS3_11430V3.1 | 142 | 6.42e-10 | QNDAGNNSQQ | HEDKVVIGRPAWMKETQLWTKQSESIDY | EKASPSRGSR |
| SfaHRS4_40s0035.1 | 139 | 8.56e-10 | NDLQGDQPVE | QPRGIDIGRPALMAEAQLWTQQVATSAA | HIERGALPEE |
| G_G019009t1 | 117 | 9.19e-10 | YEEEQTDVSA | AEKYDCKGKSLMRSSPQFWNSNCNNKKQ | KTFIEFIQAP |
| Q_112g00420.1 | 113 | 3.46e-9 | GLEQQHGGEG | GGKSSENKTPHWMQSVHLSIRTPNSSDE | ETELMKPISL |
| 1ScuHRS1_185.g025023 | 129 | 5.37e-8 | PVNRAPTCTG | RNAEEVQQRPNWMKEAGLVDTHTGSNER | TVKVLKSTSR |

### HRS Motif 7

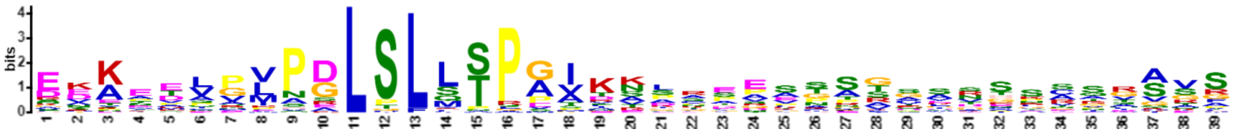


| Name | Start | *p*-value | Sites | | |
|---|---|---|---|---|---|
| G_G006998t1 | 194 | 5.06e-29 | KPNLGLAVRK | EEKEEIPVHGLTLLTPGIKNLKEESGSTGSRTSCSRAVS | SSAPNAQSNF |
| F_1031804001 | 194 | 5.71e-28 | SGFPVATARK | EDKDELPVHGLSLLTPGIKNPREESGSSGSKTSCSRGVS | SSAPNLQPNL |
| O_29780.m001324 | 201 | 3.31e-26 | YPDLSRKEDK | HSTQELPVPGLSLLTPGIKNLREESTSTGSRVSCSRAVS | SSTPNPQSNL |
| M_004G144800.1 | 215 | 1.25e-25 | YPGSSRKEDD | SNREELPVPALSLLTPGIKSIKEESNSTGSRSSCSRSVS | TTAPNSESNL |
| N_15G143500.1 | 201 | 4.52e-25 | GLSRNDDNDN | NSNGELPVPGLSLLTPGIKNFREESSSTSSRISCSRAVS | SSAPNPHSNL |
| M_009G106600.1 | 199 | 1.10e-24 | MPFKACSGLS | SKEEKLPVPSLSLSTPGIKSLKEESNSTGSRSSCSRSVS | TISGPNSDSN |
| N_17G094500.1 | 197 | 1.57e-24 | YSGLSRKEDN | DTSEELPVPGLSLLTPGIKNLRAESGSTRISCSRAVSSS | APNPQPNLRN |
| B_011780 | 193 | 2.53e-22 | YTGFPTTPLK | EDKEVLPVPALSLFTPGIKTPRHEPNPSCLNSKNSLATA | TTNFEEAKEM |
| H_62.51 | 201 | 8.48e-22 | KGFSGFPGRK | EDKEELPVQGLSLSTPGIKSLRDELNSGSRTSCSQAVSS | SATISQSNFR |
| AA_0456G0940 | 159 | 4.43e-20 | AFAPPCFRKY | DKANDVGMPDLSLLSPAPIKSTPAATTGAADESRRQVVG | FAQAAARTAA |
| K_07G178500.1 | 193 | 1.12e-19 | MPFSTYPASK | EVKEDCVFNGLSLQTPGTAVKNTREGSGCRTSSCRVVSS | APSPLRQPQS |
| L_4g086835.1 | 212 | 1.66e-19 | VTTVTLSAPK | EEKEEPVRNRLSFLTPEVKSLREGFGSRGSRSSSNRAVS | SSSPPTVQPS |
| AA_2901G0070 | 158 | 2.77e-19 | AFAPPCFRKD | DKAYDVGMPDLSLLSPAPTKSTPSAATGAADESRRQVVG | FAQAAARAAA |
| K_20G009800.1 | 193 | 1.79e-18 | IPFSSYPATK | EEKEDCVANGLSLQTPGTAVKSTREGSGCRTSSCRVGSS | APSPLHQPQS |
| B_006595 | 192 | 2.02e-18 | YSCFPATVVK | EDKDVLPISALSLLTPGIRTHSIEASPGYLNSKSIISSR | VINSSSTKHH |
| S_2G016300.1 | 200 | 2.58e-18 | AFASLCLRTD | DKAARVGMPDLSLLSPPATKSAAEESRRQVVGLAQAAAR | AAAMAPAAPA |
| AA_3297G0100 | 145 | 2.90e-18 | ALPPYFGREE | RVASTVGLPGLPLMSPAVKRPISPVTAGDDHRHAVTARF | AAAMPPSGPV |
| Z_9P26570_001 | 190 | 9.44e-18 | ISGPKIKSKE | ETKTTVMLPDLSLLSPAGNSASSPVSATVEDHPVGGSGS | KGVGRAPVSA |
| T_2G013500.1 | 161 | 1.06e-17 | AFAPLCLRVD | DKAADAGMPDLSLLSPPAIKSAPAPAGAAEESRRQVVGF | AQAAARAVAM |
| W_07g02800.2 | 160 | 2.36e-17 | RPKQSLMRKE | DMAYDVRMPDLSLLSPPASAAAADESRRQVVGFSQAAAR | AAAMAASGPA |
| AA_0185G0030 | 175 | 2.95e-17 | PTFAKSPEKP | EKAAMPPVPDLSLSSPEIDTPCPVAAPSANSSAITDAGA | QLQQPVQRKA |
| AA_1535G0300 | 151 | 7.17e-17 | PPAPYFRRED | KVASTLGLPGLPLMSPAVKRPLSPVTAGDDHRHVVTARF | AAAMPPSGPR |
| R_2G173882 | 159 | 8.00e-17 | FAPLGLRMDD | KAAARAGMPDLCLLSPPATKSAAEESRRQVVGFAQAATR | AAAATAPAAP |
| T_9G080000.1 | 158 | 1.11e-16 | PPPPCFRGDD | NAASTVGLPGLSLLPPAAKTSISPAPAVDEHRQNATARF | SAPMSPSGPA |
| F_1003551001 | 181 | 2.34e-16 | FKVSSGFPAK | GDKEVVPVAGLSLMTPMVEVDPNNSNSKCGNSGETGSGS | VSTLLTDSMR |
| Q_33g00510.1 | 141 | 1.66e-15 | NKVHRQSDVS | EKVDNQPLPDLSLLTPGIVKDDSNKCFFSATNPSTRKEH | LASLFPSPNT |
| D_01g108300.2 | 198 | 1.83e-15 | GYSGFSVTAV | RKDNKDELPGLSLHTPGITKLREDTVTSGLNSKHSGSRG | GLSSVAGCQS |
| E_B00308.1 | 213 | 2.02e-15 | ANFPVMTVGK | EDKNVLTVPKLSLCTPEIKNSMDAITSISFIPKSSSSKL | GSLSSTNIRS |
| T_3G370000.1 | 190 | 3.65e-15 | TLARSPDDAA | EKPTAVPVPELTLSSPAIDAACPAAPSATSSAVTDGGGA | AQRQHQQQQR |
| K_02G135400.1 | 214 | 7.19e-15 | TVGLGAAASK | EEKEESVMNRLSLLTPSSVKEGCGSRGSRSSSNRAVSSS | SPPMVQPSLR |
| Z_10P28540_001 | 190 | 9.57e-15 | GLSAVTANGE | EGKPTVALPDLSLQAPAMDNGALTLSAVTEGHFVSGSGS | KGAGRAPTSA |
| AA_0930G0300 | 178 | 1.85e-14 | PTLATASEKA | GNATPPVLAELSLSSPAIDRACPVSPNANANCNAIDQAR | AQGQPAAQRK |
| K_07G209500.1 | 214 | 2.23e-14 | TTVGLGAASK | EEKEESVRNRLSLLTPSVKEGCGSRGSRSSSNRAVSSSP | PTAQPGLRAT |
| V_4g03160.1 | 192 | 2.23e-14 | TMAKSPERPE | AAAMAVVPADLSLSSPAATADAAPSSNSSAVTTDAGAQS | AQQQRKARRC |
| Z_4P23210_001 | 198 | 2.23e-14 | PPFADSCKRE | EEKPAAALPDLSLQSPAIRNHHVSGCASRAVGNAPEMTP | ATVGAHVSLQ |
| Z_2P13380_001 | 187 | 3.52e-14 | LPPLAARSMT | EDKPTASLPELSLQSAVIKSNPDVVTPVTVDHRGGSAKG | ARKSPELFRA |
| AB_07.3042 | 184 | 6.07e-14 | VPFKSVERKE | EVKVTGGLPDLSLVSRPVSAAPEDHRSGVISTPRPIARC | QLSXXXXTSA |
| P_2G0120600 | 181 | 6.07e-14 | PFKEMAGKEK | AAAVPLPPPDLSLLSPATKASSSGEHQRSGGSISLTRVS | SRGTGMASPA |
| PEQU_27343 | 177 | 8.67e-14 | FKAMAAPRKE | EEKLSISLPDLSLMPLPIKVPCSAPITTVAAEEDNRSSG | LNSKAVTIAS |

| | | | | | |
|---|---|---|---|---|---|
| Z_6P34680_001 | 186 | 2.27e-13 | GMAALVMSSK | QVTPTAGLSDLSLRSPAAGSASFPVSVVAGNHPGSGSVS | KCVGRAPTST |
| C_6G095900 | 198 | 3.20e-13 | FLTSTRMEEK | EREIHQLEASLSLLTPGVRKPRTEPTSCLMNSKTTGGSR | GLPTSSAEKT |
| 2AtrHRS2_033.259 | 193 | 1.43e-12 | LNGNGNGNGL | VRKEENDIVDLSLSNGGIKELGHVSPKGSCRGNAGSSQS | ALQRKARRCW |
| W_12g39640.1 | 197 | 2.52e-12 | LATLPAFAKS | LEKADAAVPDLSLSSRVAMADAPACPAAPSATSSAVTDV | AVAQRQQAVQ |
| K_17G178500.1 | 162 | 7.55e-12 | NKSVSKTVMK | DNKKMSQVPSLGLMSPAVLELNHRKTESGYGHGSSMIIT | SSVEIKGHHQ |
| V_1g58830.1 | 149 | 1.29e-11 | GGPAFLPVSL | RKEAPLRIPDLPFLSSGSLKINSPAPAAAASASAGLQVA | GFGLDAARTA |

### HRS Motif 8

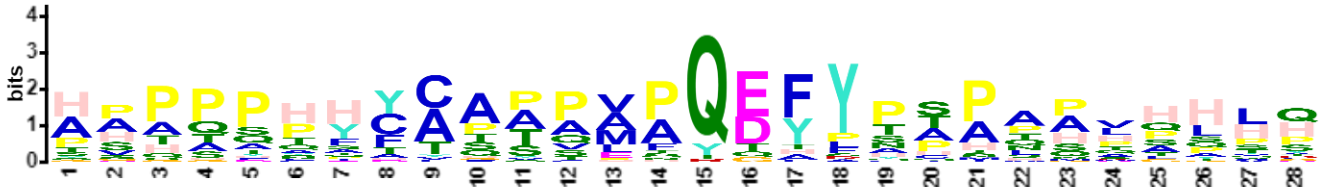


| Name | Start | *p*-value | Sites | | |
|---|---|---|---|---|---|
| Z_6P05330_001 | 310 | 2.82e-26 | AAAGPAIYGA | HAAPAHYCAATPVPQEFYPPPPPVAHHH | HLHPPLHRGA |
| K_17G119600.1 | 352 | 8.19e-25 | PTLYGAHPTS | HAPPPHYCAATPMPQDFYTTAPPQPLLP | PPPPHHNALH |
| K_05G011500.1 | 351 | 1.70e-24 | PTLYGAHPTS | HAPPPHYCAATPMPQDFYTAAPPQPLLP | PPPQHHNALH |
| B_003614 | 353 | 4.71e-23 | AASTIYGAHP | ASHPSTHYCAPAVAQEFYPTPPHHQLHH | SLHPQQHIYK |
| S_3G046800.1 | 374 | 3.32e-22 | QAAGQAIYGA | HPATQPHYTAAVAAQEYYPSPAAVHHLQ | HHPAAAAAAM |
| N_06G060000.1 | 364 | 4.85e-22 | PTLYGAHPAS | HAPPPHFCAAPPMPQDFYTTAAAAAPSS | PQPTHHQMHH |
| K_04G151000.1 | 356 | 1.68e-21 | PTLCGPHPTS | HVPPPHYCAPTPMPQEFYNSAPSLSLPS | PAHENILHHH |
| T_5G124700.1 | 364 | 1.68e-21 | AAGQAIYGAH | PATQPHYTAAVAAAQEYYPPPAAVHHLQ | HHPAAAMVHH |
| N_14G127500.1 | 365 | 3.09e-21 | PTLYGTHPAS | HAPPPHFCAAPPVHQDFYTAAAATPSPQ | QPPHHHLHST |
| B_006061 | 354 | 9.04e-21 | AAPTIYSAHP | ASHASTHYCAPAVAQEFYPTPTHQLHHS | LHHVYKASSQ |
| O_29807.m000474 | 357 | 9.04e-21 | LYGTHHHPVS | HAPPPHFCAAPPVPQDFYTAAAAAGATT | PPLPPPHHQL |
| F_1015462001 | 299 | 2.58e-20 | GGTTLYGAHP | PSHTSPHYCGPPVPQDFYAAPAQPHHHS | LHHQLHMYKA |
| AA_1881G0390 | 275 | 5.74e-20 | QAGAPAIYGA | HPATQPHYTAAVAAQEYYQSHAAVHHLQ | HHPAAAMVHR |
| K_06G213400.1 | 361 | 8.05e-20 | PTLYGPHPTS | QVAPPHYCAATQMPQEFYNSALPLSLLP | PPHDYTLHHH |
| X_9781.1 | 339 | 8.05e-20 | TTAAGPAIYG | AHPGPAHYCTAPVPQDFYPTATGVAPLP | HHQIHPALHR |
| W_01g08160.1 | 364 | 8.93e-19 | QAAGPAIYGA | HPATQPHYTAAVAAQEYYHHHHHHLQHH | PAAAALVHHR |
| P_27G0017800 | 337 | 7.16e-18 | TAKAGQLFYG | GHPAAAHYCAAAVQQEFYSPPLQPPHYH | HGTASAARAA |
| R_2G348238 | 371 | 8.77e-18 | SQAAGQAIYG | AHPATQPHYTAAAAQEYYPSPAAVHHLQ | HHPAAAAMVH |
| 5AthHOH4_AT2G03500 | 341 | 1.03e-16 | HQVHHHHTNT | AGPPPPHFCSSQEFYTTPPPPQPLHHHH | FQTFNGSSGG |
| H_127.55 | 364 | 1.25e-16 | LYGTHHHPAA | PHAPTHFCAPPPVPQEFYSSTSPAAAPP | PALHHPTLHH |
| C_7G203800 | 338 | 1.68e-15 | AHGGGPTIYS | AHPASLATTHYCAPQEFYPAPSSHQLHH | TIHHPHPHHQ |
| J_3853942 | 331 | 2.41e-15 | HQVQNHQGNA | AVQPPPHFCSSQEFYTARPPPQQLHHHH | FQTCNGSSAD |
| 1AtrHRS1_122.65 | 259 | 3.45e-15 | NPSQTLYTTT | PPTQPHHFCQPTMAQDHLYAPTHRTYCD | QMALAHMNPS |
| J_3875404 | 331 | 1.28e-14 | HQVHHHNGNA | ATQPPQHFCSSQEFYTTPPTPQPLHHQH | FQTFNGSSAG |
| Q_208g00340.1 | 307 | 1.52e-14 | AAAAGATFYG | THPSAHYCATATANQQYPPHHLHHSHNR | QTFNNRSSVM |
| D_12g006800.1 | 347 | 5.42e-14 | TFYGPHSTSH | APSPHYIAAPQALAQEFYNTPQPLQTLH | HQQLYHPGSH |
| L_4g113140.1 | 349 | 7.56e-14 | HHLTLPHYCT | APGQDQYYTTAPPPQQLLPPPHHMHHVY | KTTPHGQGSP |
| PpaHRS2_8580V3.1 | 398 | 1.35e-13 | SSGVYDPSIA | HPPQPTEFCPTSLPQDYFACINNDTSSV | SLQMHRQPIF |
| 2PabHRS2_33594g0010 | 340 | 2.02e-13 | GLYGTLSNTT | SSHSQSHYCQSPMPQEHYSQMNMSSQLQ | LPQPAYCEQQ |
| V_2g04810.1 | 384 | 3.85e-13 | PAAIYGAHPA | TQTHYTAAVSSAAAQEYYHSAAAHHHHL | QQQQHHHPHP |
| SfaHRS1_10s0233.1 | 414 | 1.58e-12 | QAAAQPTSGV | YDPSPPSYCQPSHPQDYFTCNMSTGSSG | GQLKQHPTTF |
| PpaHRS1_4480V3.1 | 397 | 1.61e-11 | SPGFYNPSAA | HSPQPNQFCPTSLPQGYSCISNSTQGAQ | LEMHREPIFE |
| Q_30g00060.1 | 317 | 3.83e-11 | SATVYSTHPS | SATQFCTTAATPLSRGFYSSPTAPQPPQ | ASNAHSHCSV |
| S_8G147101.1 | 91 | 2.67e-10 | ELAQAQHRRH | PPPPPPSDLFAPATTEAAAAPAPPPQTQ | AQAQDGAAVK |

HRS Motif 9

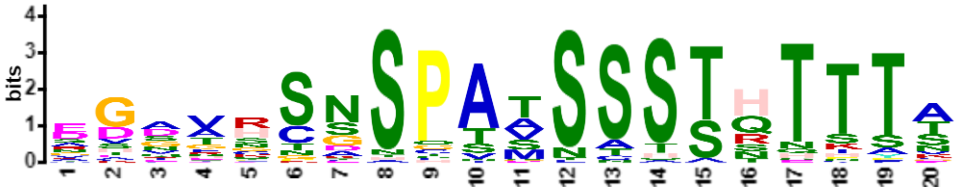


| Name | Start | *p*-value | Sites | | |
|---|---|---|---|---|---|
| D_05g009720.2 | 377 | 1.51e-22 | ERDRSHSHSD | CGGVHSNSPATSSSTHTTTA | SPAY* |
| N_14G098400.1 | 360 | 2.23e-21 | MQLEERGSHS | DGGFRSNSPATSSSTHTTTN | SPSF* |
| N_06G071500.1 | 377 | 4.92e-21 | MQSEERGSHS | EDGVRSNSPATSSSTHTTTN | SPLF* |
| F_1011942001 | 333 | 1.77e-20 | SQSEERGSHS | EGGVQSNSPATSSSTHTTTT | SPVF* |
| E_E00311.1 | 355 | 5.70e-20 | SDNDRGCSHS | DGGVHSNSPATSSSTHTSTA | SPAIY* |
| L_1g093080.1 | 354 | 2.30e-19 | SISEERGNNS | EGAVHSNSPTSSSSTHTTTN | ASTGY* |
| P_8G0062400 | 397 | 3.19e-18 | GTREDQRGVD | DGATRSNSPATSSSSHTTSA | SHFNRHNVAQ |
| H_1.213 | 342 | 5.31e-18 | QSEERGCSHS | EGRVHSSSPSTSSSTHTTTA | SPVF* |
| G_G011857t1 | 365 | 8.72e-18 | SQSEERGSHS | EGRVHSNSPSTSSSTHTTTD | SPLF* |
| M_008G117500.1 | 362 | 1.11e-17 | QSEGGGSHSE | GGAHSNNSPATSSSTHTTTT | * |
| M_010G128900.1 | 367 | 1.11e-17 | SEGRGSHGER | GGAHSNNSPATSSSTHTTTT | SPVF* |
| O_28883.m000751 | 370 | 1.26e-17 | SQHLQSERRG | SHSERSNSPATSSSTHTTTN | SPVF |
| K_02G098800.1 | 348 | 2.27e-17 | HSHSPSEEKN | NHSVHSNSPASSSSTHTITT | SPPA* |
| K_20G186500.1 | 359 | 3.61e-17 | SNSDDRANHS | EGAVHSNSPTSSSSTHTSTS | PGY* |
| K_10G204200.1 | 361 | 3.61e-17 | SISDERANHS | EGAVHSNSPTSSSSTHTSTS | LGY* |
| J_3831280 | 334 | 1.88e-16 | APQSPKRSVE | RSSGRCNSPAASSSTNTTTT | SASPVS |
| B_012707 | 340 | 2.87e-16 | QSEGRSSGQR | DDDAHSNSPSASSSTHTTAA | ASPSL |
| Y_30s1094441g001 | 341 | 2.87e-16 | LHSQGRCSGD | DGAANSASPTTSSSSQTTTA | SPPF |
| K_01G086700.1 | 354 | 1.74e-15 | KEHSHSEERP | NHSVLSNSPASSSSTHTITT | SPPVPN* |
| J_3875351 | 354 | 3.08e-15 | ISEEISRCSD | KGIVRCSSPAMSSSTRTKTK | DAKMS |
| T_1G146600.1 | 382 | 5.39e-15 | QSSRCSGGRR | SGDASSDSPAVSSSSHTTSA | * |
| L_5g054300.1 | 339 | 8.50e-15 | ELSENSHSVE | RVVAHSNSPASSCSTHTPTT | SRC* |
| AA_5036G0050 | 374 | 1.45e-14 | RYSEGRRSGD | TGDACSASPAVSSSSQTTSA | |
| 2AthHOH1_AT3G25790 | 332 | 1.89e-14 | ISEDRSRCSN | NGFFRCSSPAMSCSTRTKTK | DAKIIS |
| W_02g22020.1 | 393 | 2.07e-14 | SGSEGRRSGD | AGDGSSSSPAVSSSSQTTSA | * |
| J_3839360 | 343 | 2.93e-14 | ISEERSRCSN | KGIIRCSSPAMSSSTRTKTK | DAKLS |
| Z_11P13410_001 | 335 | 5.31e-14 | RSEESNSSGN | AAAFNSLSPAASSSSQTTTA | SHLEITCKSF |
| S_8G036900.1 | 442 | 6.83e-14 | QSSSCSGGRR | SGDACSGSPAVSSSSSQTTS | A* |
| J_3852516 | 345 | 1.43e-13 | PPQSPKRSVE | RSSGQCNSQAASSSTNTTTS | SPVS |
| Z_8P18270_001 | 348 | 1.55e-13 | GEDDNSMRDD | GDATNSASPTTSATSQTTTA | SPLF |
| J_3836112 | 324 | 5.51e-13 | EEKGSGSPSE | EVVFRCSSPAMSSSTRNHYV | KNI |
| AB_01.3080 | 326 | 5.51e-13 | RCEGDDNSED | AEQTNSDSSATSSSSRTTTA | SAPLVRVC* |
| Z_6P09870_001 | 317 | 5.96e-13 | GGRCSEDVNS | MGDDDSASPTTSASSQTTTA | SPPF |
| J_3871943 | 323 | 6.43e-13 | VEEGSGSHSD | EVVVRCSSPAMSSSTRNCFS | KPTHIFKSNP |
| 1AthHRS1_AT1G13300 | 322 | 9.42e-13 | VDEGSGSHSE | GVVVRCSSPAMSSSTRNHYV | KNN |
| V_3g10730.1 | 377 | 9.42e-13 | SKHSEDQRSG | DASSSSGSPAVSSSSQTTSA | * |
| R_2G016370 | 433 | 9.42e-13 | QSSSCSGARR | NGDACSGSPAVSSSSSQTAS | A |
| Z_3P21090_001 | 330 | 1.85e-12 | GGEDSTGGDD | DDATNTESLATSASSQTTTA | SPPCKLRSGT |
| 4AthHOH2_AT1G68670 | 331 | 1.99e-12 | PVAVAQSPKR | SLERSCNSPAASSSTNTNTS | TPVS |
| C_3G045700 | 329 | 4.43e-12 | KSSLAEHSTR | EKNARSSSPAVTSSSTETTT | ASPNF* |
| R_2G159119 | 439 | 3.30e-11 | QSSSCSGARR | SGDACSGSPAVLSSSSHTAS | A |
| K_05G011500.1 | 220 | 6.83e-11 | KENSGSDGAV | VDQGKGGSPVASSHAQTTTT | TTSAQTHRKA |
| K_17G119600.1 | 221 | 2.02e-10 | KENSGSDGTV | VDQGKGGSPVASSHATTTTT | TSSAQTHRKA |
| L_4g113140.1 | 208 | 2.15e-10 | EKGKTNSGNE | VDNQGKGSPVASSQTQTTSN | NSNQTHRKAR |
| C_3G398400 | 306 | 2.29e-10 | RGSQAQHCME | DEGTHSDHQSSSSSTQSTTV | SVSLV* |
| 3AthHOH3_AT1G25550 | 320 | 2.60e-10 | QPPQSSTSGE | RSNRGCKSPATSSTTTHTPH | LLPLS |
| PEQU_26724 | 302 | 2.76e-10 | EDDNNNPADH | ESETKSDSCNMSSASQTTTL | PLL |
| AB_06.1057 | 357 | 8.26e-10 | DTSPERDDNV | EDSERPKSHSSNTSSQTTTA | SLP* |
| N_06G060000.1 | 233 | 1.99e-9 | GTGNSSSDGQ | ATNTGTGNCTNNTTTSTTTS | SQAHRKARRC |
| O_29807.m000474 | 226 | 8.13e-9 | HGKGNGNLAS | CDGQPTNTGTGNNSTNTSTT | TQTHRKARRC |
| N_14G127500.1 | 233 | 1.01e-8 | GTGNSSSDGQ | ATNTGTSNGADNTTTSTTTG | SQTHRKARRC |

| | | | | | |
|---|---|---|---|---|---|
| M_006G155200.1 | 226 | 1.73e-8 | GKGAGNSSSS | DGQATNTATIASASTNTSTS | QTHRKARRCW |

### HRS Motif 10

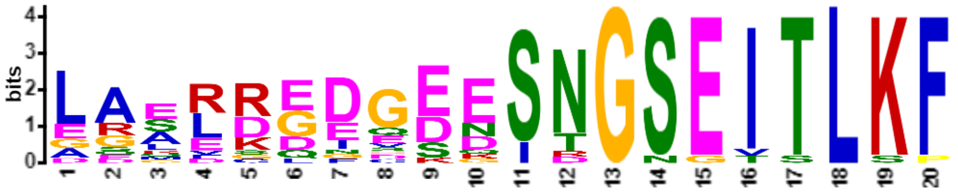


| Name | Start | *p*-value | Sites | | |
|---|---|---|---|---|---|
| H_127.55 | 459 | 2.19e-23 | NGNSGDQRKG | LAALREDGEESNGSEITLKF | * |
| O_29807.m000474 | 455 | 1.02e-22 | SGENGGERKG | LAAFREDGEESNGSEITLKF | |
| B_006061 | 436 | 4.35e-22 | GDQNGGERKG | LASLRDDGDESNGSEITLKF | |
| M_018G074200.1 | 444 | 6.14e-22 | GENDGGERRG | LAALREDCEESNGSEITLKF | * |
| G_G037970t1 | 448 | 5.03e-21 | DNGGAGERKG | LAALREEGEESNGSEITLKF | * |
| K_05G011500.1 | 435 | 1.85e-19 | SSWKGESGEN | EGERKGIGEESNGSEITLKF | * |
| 5AthHOH4_AT2G03500 | 413 | 1.48e-18 | GGGERKGLAA | LREECEDHSNINGSEITLKF | |
| L_4g113140.1 | 415 | 5.56e-18 | SSWKEGSSED | EGERKGFVEESNGSEITLKF | * |
| K_06G213400.1 | 448 | 1.21e-17 | NGESGEKKGL | AALRDQEGEESTGSEITLKF | * |
| K_04G151000.1 | 443 | 1.77e-17 | KAENGEKKGL | AALRDQEGDESTGSEITLKF | * |
| J_3875404 | 404 | 1.95e-17 | GGGEGKGLAA | LRQEGEDQSNINGSEITLKF | |
| N_06G060000.1 | 457 | 4.10e-17 | SGENGGDRKG | LASLREDGDDSNGSEISLKP | LRY* |
| J_3853942 | 403 | 2.05e-16 | GGGERKGLAA | LREESGNQSNINGSEITLKF | |
| 1AtrHRS1_122.65 | 363 | 4.07e-16 | DMDEDGDGDD | DDGDDGDEDDSTGSEITLKF | * |
| K_17G119600.1 | 433 | 5.25e-16 | SESSSWKGEN | EGERKGISKGSNGSEITLKF | * |
| B_003614 | 438 | 7.34e-16 | GENGGDRKGL | GSMRDDGDDQSNGSEITLKF | |
| E_H01532.1 | 474 | 2.31e-15 | KADSGGGENG | GERRRLDGEESNGSGVTLKF | * |
| 2PabHRS2_33594g0010 | 449 | 2.94e-15 | GRGLSLRKQP | LQSVDGDEEDSRGSETTLKF | |
| C_7G203800 | 427 | 1.30e-14 | SGENGGERRG | LASLRDEGEKSDGNEITLSF | * |

### HRS Motif 11

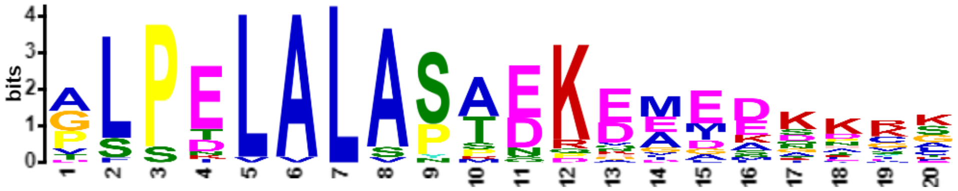


| Name | Start | *p*-value | Sites | | |
|---|---|---|---|---|---|
| B_006061 | 179 | 1.16e-22 | RNTCASPTLR | GLPELALASTDKEMEDKKCS | ESDNAVSVSC |
| B_003614 | 174 | 4.52e-19 | KSSCATPTLS | VLPELALASTNKEMEDKKRS | ESDNGVSVSC |
| K_05G011500.1 | 178 | 2.72e-18 | RNSCQGSTLR | PLPELVLASAEKEMEDKKRA | EVEIKGVSCQ |
| G_G037970t1 | 177 | 4.33e-18 | RNSCPGSALQ | ALPDLALASANKDMEDKRCS | DTENGMSCQR |
| K_17G119600.1 | 178 | 5.82e-18 | RNSCQGSTLR | PLPELALAYAEKEMEDKKLR | PEVEIKGVSC |
| K_06G213400.1 | 181 | 7.01e-17 | KERNSCQGLR | GLPELALASPEKEIEENKCE | LEAEKCSKRE |
| X_9781.1 | 174 | 1.03e-16 | AKGGGGVTPR | GLPELALASADKGEEEKKGG | VAVMENGGFG |
| R_2G348238 | 206 | 2.44e-16 | DKTASAAEGA | ALPELALAPADKDAADADRK | PYLDAAAGSN |
| T_5G124700.1 | 204 | 6.99e-16 | AKDKAAAEAA | ALPELALAPAEKDAAETDRK | PYLDAAGANG |
| S_3G046800.1 | 211 | 8.76e-16 | DKTASAAEGA | ALPELALAPADKDAGDAERK | PYLDASSNNG |
| L_4g113140.1 | 176 | 1.52e-15 | ERNNSSCQGQ | GLPELALASTQKEEDKKHVG | EAEKGKTNSG |
| O_29807.m000474 | 170 | 3.58e-15 | LPFSKERNLC | PSPTLALASTDKELMEDKKC | LETENGLSCS |
| AA_1881G0390 | 201 | 4.88e-15 | SKDKAMAESA | ALPELALAPAEKDAIAGAGA | EVDKKPYHDA |

| | | | | | |
|---|---|---|---|---|---|
| P_27G0017800 | 177 | 1.78e-14 | SKERDGRVRG | GLPELALAPMEKEDEKRPYS | ENGGKGGAAA |
| N_06G060000.1 | 172 | 3.79e-14 | LPFSKERNLC | PSPILALASTEPEMEDQKCL | ETENGFSCPK |
| H_127.55 | 185 | 6.00e-14 | SCPTPTPVRA | GLPELALVSREKDGMDQDSK | FSDQMGESGM |
| AA_2274G0070 | 200 | 6.00e-14 | SKDKAMAESA | ALPKLALAPAEKDAIAGAGG | EVDKMQYHDA |
| K_04G151000.1 | 178 | 6.57e-14 | KERNSCQGLR | DLPEVALASSEKEMEKKCEL | ESEKCSKREN |
| M_018G074200.1 | 170 | 7.88e-14 | LPFSKERNLC | PSPTLALSSTDKHMEFDHKK | CSEAENGFSC |
| 1AtrHRS1_122.65 | 91 | 1.03e-13 | PFSKDRESRK | TLSDLALASTEKEEIESPKF | TEHEASGANK |
| W_01g08160.1 | 200 | 1.03e-13 | MGDGSAAAAA | ALPELALAPAEKAADAITIA | AGEVDKKPYA |
| N_14G127500.1 | 171 | 1.01e-12 | LPFSKERNLC | PSPTLALASADQQEMENKKC | LEIENGFSCS |
| E_H01532.1 | 195 | 7.87e-11 | SSSSSMNCGE | GFPELALANSSRNEEMKNSS | SSSEVVKNIC |
| C_7G203800 | 179 | 9.01e-11 | NTCASPNTLR | VLSDLALASTDDKELNDNKK | CLESDKVKGG |
| Z_6P05330_001 | 176 | 1.10e-10 | KSKAARSASR | ALSELALASPEKVGGSSSAT | EGQAAAPPTH |
| 1PabHRS1_14087g0010 | 183 | 1.18e-10 | PRPVRSTVER | TLPNLALSSAEREVDSSLVG | NETVCLDATT |

### HRS Motif 12

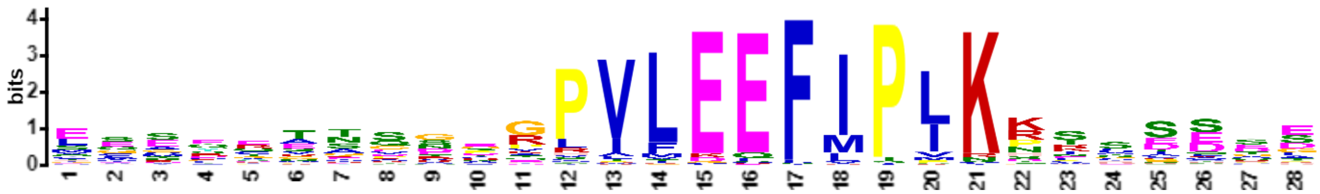


| Name | Start | *p*-value | Sites | | |
|---|---|---|---|---|---|
| K_20G186500.1 | 68 | 1.38e-23 | TSEYNLNGHS | ECSEQTTSTEGPVLEEFIPIKKRASSSS | PCCDEDDEQH |
| L_1g093080.1 | 61 | 1.26e-22 | TTEYNLNGQS | ECSEQTTSTDGPVLEEFIPIKKRASSYS | QEVFDDVEDY |
| N_06G060000.1 | 73 | 4.00e-22 | TNAVETSRQQ | LQSYRANQGPRPVLEEFIPLKNSTSETL | DNSSNISDKA |
| B_003614 | 69 | 5.00e-22 | TNAMEALRQQ | LQTYRTNQGPRPVLEEFIPLKHSGSEGS | EKAASISEKV |
| F_1011942001 | 55 | 1.21e-21 | GTTQEYFHGQ | SECSEQTSSDGPVLEEFIPIKKTSDDED | EQQSHQPNDN |
| K_10G204200.1 | 68 | 1.50e-21 | ASEYNLNGHS | ECSEQTTSTEGPVLEEFIPIKKMASSSP | FCDEEDEDDE |
| B_011780 | 79 | 3.48e-21 | LNDAIVSLKE | EGMQCTTSNVRPVMEEFIPLKRNSDEDV | RAKKEKESRD |
| B_012707 | 61 | 5.25e-21 | ATSTEYFPGP | TECSEQTSSEGPVLEEFIPLKRSSSSEE | EEELNKPNNK |
| G_G011857t1 | 66 | 5.25e-21 | STTTDYMQGQ | SECSEQTSSDGPVLEEFIPIKRSSDCSE | EDDEQESRKS |
| M_010G128900.1 | 62 | 6.43e-21 | TTEDHNMHGQ | SECSEQTSSEGPVLEEFIPIKRTHSSDD | EENDNNHDDD |
| B_006061 | 72 | 7.87e-21 | TNAMEASRQQ | LQTYRTNQVSRPVLEEFIPLKHSASDQG | LEKASTASEM |
| H_127.55 | 74 | 1.73e-20 | TNAVEASRQQ | LQAYRANQGPRPVLEEFIPLKNPSSEST | EKLNTISDKA |
| N_17G094500.1 | 79 | 1.73e-20 | LNDAILFLKA | ESTQCAASNNQPVLEEFIPLKTNCDDDD | DEQDGPIKKE |
| 4AthHOH2_AT1G68670 | 64 | 2.55e-20 | SGTTTTTSEQ | CSEQTTSVCGGPVFEEFIPIKKISSLCE | EVQEEEEEDG |
| M_008G117500.1 | 62 | 2.55e-20 | TTTEYNMHGQ | SECSEQTSSEGPVLEEFIPIKRTHSYDE | NDNENDDHQE |
| Y_30s1094441g001 | 65 | 3.08e-20 | AFILGSIRVQ | MGSSEETVTDGPVLEEFIPLKPTSSSIE | EEKGGSMEMD |
| B_006595 | 79 | 4.49e-20 | LDDAIITLKE | HAMQCTTSDVRPVMEEFIPLKRNSDEDG | APKKENDCRD |
| B_001193 | 70 | 4.49e-20 | ATTTEYFPGQ | SECSEETSSEAPVLEEFIPLKRCFSSEE | DEESNKPKNE |
| N_15G143500.1 | 79 | 4.49e-20 | LNDAILFLKA | ESIQYAASNNPPILEEFIPLKKNCDDDD | DDDEQDGRIN |
| Y_30s1025391g002 | 72 | 6.50e-20 | NNAVETYRQQ | LETYQTNQAPRPVLEEFIPLKHASIDGS | EKAPNAPSEK |
| Z_6P05330_001 | 72 | 7.81e-20 | NNAIECYKQQ | LETYQTNQGPRPVLEEFIPLKHMNMDGS | DKDPSAHSEK |
| N_14G127500.1 | 72 | 9.36e-20 | TNAMETSRQQ | IQAYRANQGPRPVLEEFIPLKNHASETL | DRSSNISDKA |
| C_3G045700 | 70 | 1.91e-19 | VYSSERSDQC | EEKQEETSSEGPVLEEFIPMKRSLSCEE | EEEDKSNIDE |
| Y_30s723941g001 | 49 | 2.70e-19 | TQAIESIRMQ | IWSSEETVTDGPVLEEFIPLKPTSSSTE | EEKGGFMEME |
| X_31310.1 | 49 | 3.21e-19 | TQTIETVKQQ | MESENVRSDDGPVLEEFIPLKPSLSSTS | EEESALHESK |
| K_01G086700.1 | 66 | 4.52e-19 | TVAEYNLNGQ | SECSEQTSTDGPVFEEFIPLKKRASQDS | VEEEDEDEEH |
| N_06G071500.1 | 63 | 4.52e-19 | TTEYMHGQSE | CSEQTLSEGTRPVLEEFIPIKRTHSSSD | NDNDDDDEND |
| G_G037970t1 | 73 | 5.35e-19 | TNAVEASRQQ | LLACRANHGSRPVLEEFMPLKNSSSENS | EKSQNISDKA |
| N_14G098400.1 | 63 | 8.80e-19 | TTDYLHGQSE | SSEQTSSEGTKPVLEEFIPIKKTNSSSD | NDNEEQYLHK |
| E_E00311.1 | 64 | 1.04e-18 | CKQQLHGQLS | ECSEQTSSDVVPVLEEFIPIKRAFSSFH | SDEDDEEEEE |
| AA_5036G0050 | 59 | 1.68e-18 | IEGMRSQMDG | VGSEETVSDHGPVLEEFIPLKPSLSLSS | SEEDSTHAAA |
| 1AtrHRS1_122.65 | 10 | 3.68e-18 | MEASRQQLA | SCSELTRSSSKPVLEEFIPIKRSSSEGS | ETTNMTSDKA |
| H_1.213 | 63 | 3.68e-18 | TADYMHVQSE | CSEQTMSEVGPPVLEEFIPIKRTSSPDE | DEQESHKAST |
| V_3g10730.1 | 60 | 4.30e-18 | EGMKSQMDAV | VGSEETVSDHGPVLEEFIPLKPSLSLCS | SEEESTHAVA |
| L_4g086835.1 | 80 | 6.77e-18 | LNDAILVLKE | ELEKCTSKNSVPVLEEFIPLKKEIDQSE | ENKNNDRDKN |
| O_29807.m000474 | 72 | 6.77e-18 | TDAVETSRQQ | LQAYRANQGPRRILEEFMPLKNSTSETL | DKSSNMSDKA |

| | | | | | |
|---|---|---|---|---|---|
| D_01g108300.2 | 79 | 9.13e-18 | VKDAIVALRE | ESMQYRKSRTEPVLEEFIPLKKSSREDT | KAEITKDKDS |
| M_006G155200.1 | 72 | 1.64e-17 | TSAVETSRQK | LQAYRGNQVPIPVLEEFIPLKTSTSETP | EKTSNISDKA |
| J_3831280 | 66 | 1.89e-17 | LSVTSTTSEQ | YSEQTASVCGGPVLEEFIPIKKSNNEEN | REHESPREVD |
| K_07G178500.1 | 81 | 2.90e-17 | LNDAISALKV | ESEKCMACKSEPVLEEFIPLKKECDQRE | ESEKEKECRD |
| T_1G146600.1 | 58 | 2.90e-17 | IEGMKSQMHG | VGSEGTVSDHGPVLEEFMPLKPSLSLSS | DEHESADDAA |
| K_17G119600.1 | 72 | 3.34e-17 | TNAMEASRQQ | LQAYKVNHGTKPVLEEFIPMKHLASDQS | SEKATNMSDK |
| M_018G074200.1 | 72 | 3.85e-17 | TNAVETSRQQ | LQAYRANQVPTPVLEEFIPLKTPTSEAL | EKTTNISDKA |
| Z_11P13410_001 | 49 | 4.42e-17 | THAIDSARRQ | MNGHATMSEDGPILEEFIPLKPSSSSSE | DRSVDGAAKK |
| G_G006998t1 | 79 | 5.07e-17 | LNDAIVALKE | ESMQCVTRNVEPVLEEFIPLKNNKKETK | HSEEDGASIT |
| AB_06.1057 | 97 | 5.07e-17 | QAIESCKKPE | VGGSEEISSEGPILEEFIPLRPSCSSSD | EENNNNHGNG |
| L_5g054300.1 | 62 | 5.82e-17 | CKQQLFGTQS | ECSEQTSTDEGLVFEEFIPIKKRALSPD | CDENDEEDDD |
| X_9781.1 | 72 | 5.82e-17 | NNAVEAYRQQ | LEAYQTSQGPRPVLEEFIPVKQMSVDGL | ETPNGPSSEK |
| P_8G0062400 | 72 | 7.64e-17 | RMAGAGCRSE | VDEEETSCEGGPILEEFIPIKRRSSPGA | DQGEEDGHRQ |
| F_1015462001 | 72 | 1.14e-16 | TDAVEASRQQ | LQSYRANQGARPFLEEFIPLKHSTPKGS | EKTSNMSDKA |
| M_004G144800.1 | 95 | 1.31e-16 | NDAIQVVREE | LMQCGTSNNQQPVLEEFIPLKKKIDDHG | DDRESDGLIK |
| J_3852516 | 61 | 1.49e-16 | LSSTSTTSEQ | CSEQTTSVSGGPVLEEFIPTKKIEENGE | HESPTPEEIG |
| Z_6P09870_001 | 49 | 1.70e-16 | VTRAIECVRE | QMGDDESVNDAPVLEEFIPLKPSLASTS | SEGSGEAKKA |
| S_8G036900.1 | 68 | 1.94e-16 | SHMDSVVVGS | EEETVSDHGGGPVLEEFMPLKPTLSSSS | SEDDDDDEHD |
| D_05g009720.2 | 69 | 2.21e-16 | EYNLNAQSTE | CSDDEHTSSDVPILEEFIPLKSTFSHED | EDEDENDEEN |
| Z_3P21090_001 | 48 | 2.21e-16 | VTHAIESVRQ | QMSDGERVSNGPVLEEFIPLKPSFKSPS | SGAVAAAIGS |
| K_05G011500.1 | 72 | 3.24e-16 | TNAMEASRQQ | LQAFKVNHGAKPVLEEFIPMKHLASESS | EKATNMSDKA |
| E_H01532.1 | 77 | 3.24e-16 | TNAMEASRQQ | LQSQRTNQEERPILEQFIPLKNTSNNSC | NNNISDKEAN |
| 1AthHRS1_AT1G13300 | 69 | 4.18e-16 | MTTENMYGQP | ECSEQTTGECGPVLEQFLTIKDSSTSNE | EEDEEFDDEH |
| K_07G209500.1 | 79 | 4.18e-16 | DAISVLKVES | QKCCRVARDSPPVLEEFIPLKKELGDQS | EEEEENDDDK |
| E_L01092.1 | 51 | 4.74e-16 | QQLTHEKSGC | NSEQTSNDDVVPVLEEFMPIKRAADSDS | DQDQDQSEEI |
| R_2G016370 | 64 | 6.89e-16 | MRSHMDSVVV | GSEETVSDHGGPVLEEFMPLKPTTLSSS | SSQPQSQDDH |
| R_2G159119 | 64 | 6.89e-16 | MRSHMDSVVV | GSEETVSDHGGPVLEEFMPLKPTTLSSS | SSQPHYQDEH |
| O_29780.m001324 | 79 | 7.80e-16 | LNDAILFLKS | ELMQCAALDHQPILEEFMPLKKHSDRDD | DDDGDQDGPI |
| Z_4P28210_001 | 79 | 7.80e-16 | LREVIERLRM | EIERCRCESFGHVFEEFMPLKSKVEDDG | GVKVETDCRD |
| H_62.51 | 80 | 8.82e-16 | LKDAIIALKE | ESMRYAAKNALPVLEEFIPLKKRDRDEE | DEEDRAAARK |
| 3AthHOH3_AT1G25550 | 70 | 8.82e-16 | SSEHVGGQSE | CSERTTSECGGAVFEEFMPIKWSSASSD | ETDKDEEAEK |
| K_20G009800.1 | 81 | 9.97e-16 | LNDAISALKV | ESEKCRACKSEPVFEEFIPLKKECDQRK | EIEKEKECRD |
| 2AthHOH1_AT3G25790 | 70 | 1.13e-15 | TSTDNLYGQS | ECSEQTTGECGRILDLFIPIKHSSTSIE | EEVDDKDDDD |
| K_04G151000.1 | 74 | 1.13e-15 | TNAVEASRQQ | LQAFRSNQGTRPVREEFMPIKHPNSQES | TEKTSNISDK |
| AB_01.3080 | 43 | 1.13e-15 | ENCKKKNDEV | VGRGEEIASEGPILEEFIPLRPSCSSSD | EENNHKNSGN |
| AB_10.37 | 43 | 1.13e-15 | ESCKKKKHEV | VGRGEEIASEGPILEEFIPLRPSCSSSD | EENNHKNSGN |
| E_B00308.1 | 84 | 1.43e-15 | KEEIGKMKEE | MAELEKCNAEPPVLEEFIPLKKICTDEK | DDDKVEGNNK |
| W_02g22020.1 | 59 | 1.43e-15 | GMRSQMDAAG | SEETVSDQGPPPVLEEFIPLKPSLSLSS | SEEESTHADA |
| F_1031804001 | 79 | 1.62e-15 | LSDAILALKE | ELLRSKASNVQPVLEEFIPLKKDCDEEG | GAKEEKECKD |
| Z_8P18270_001 | 48 | 1.82e-15 | VTQAIESVRH | RAGEEEGVYEGPVLEEFIPLKPSPTSTL | SEEEKGETRK |
| J_3836112 | 69 | 2.60e-15 | TATENMYGQS | ECSEQTTGECTPVLEQFLTIKDSSTSNE | EEELDDEHGN |
| J_3875351 | 70 | 2.92e-15 | TTTDTLYGQP | ECSEQTTGECGPVLDLFLPIKQSSASTD | EEEEEEEVDD |
| K_02G135400.1 | 77 | 3.28e-15 | NDAISVLKAE | SQKCRVARDSAPVLEEFIPLKKERGDQS | EEEEEEEEND |
| C_7G203800 | 74 | 4.64e-15 | NAIETYRQQL | QSYRGNQGTNKPVLEEFIPLKHSSSEGS | EKTSSISEKS |
| Z_7P00660_001 | 61 | 4.64e-15 | LSEVIEGLNK | EIQRCRGDRFGCVFEEFIPMKSKVEEDG | RVKAETDCKD |
| L_4g113140.1 | 72 | 5.21e-15 | TNAMEASKQQ | LQAFRSNQGAKPILEEFIPVKQLTSSET | LEKTTNNNVC |
| T_5G124700.1 | 94 | 6.54e-15 | AYGQQLEAYQ | MGSLQGAPARPLVLEEFIPLKKIGIDAA | ADKMGNQPSE |
| P_27G0017800 | 82 | 6.54e-15 | NNAMEAYRQQ | LDMHHTNRSPRPVLEEFIPLKNPSVAAE | APEKSAAADD |
| 2PabHRS2_33594g0010 | 32 | 8.20e-15 | VQAPRPVQQA | GFQFEEEQYNRPVLEELIPIKKRCDESG | DESPENNQND |
| K_06G213400.1 | 75 | 1.15e-14 | TNAVEASRQQ | LQAFRSNQGTRPVLEEFMPILKHPNSQE | SAEKTSNISD |
| Q_112g00420.1 | 56 | 1.99e-14 | TIESYRQQIG | CEGDVATSSDVPVLEEFVPINRCSSVEE | EESSHENESK |
| AA_2274G0070 | 95 | 2.22e-14 | RQQLEAYQMG | SQGGAAAARRPLVLEEFIPLKNIGIDEA | DKPGNPPSEK |
| F_1003551001 | 75 | 2.47e-14 | LNDAILRLRE | EAMQCTESEGGHVTEEFIPLKGRSEGDE | GAEMGKDSSD |
| E_N01205.1 | 107 | 2.75e-14 | SIAAIAKIKE | ELIQCRNSITEPIIEEFIPLKKSLPKKE | DDKIDSVDKM |
| M_009G106600.1 | 87 | 3.06e-14 | NDAIQVLREE | PMQRGASKNQQPVLEEFIPLKKNIDDHH | ENDGLIEEKD |
| R_2G348238 | 98 | 3.79e-14 | SYRQQLEAYQ | MGSLQGAPARPLVLEEFMPLKNIGIDAA | ADKMGNPTSE |
| S_3G046800.1 | 97 | 3.79e-14 | SYRQQLEAYQ | MGSLQGAPARPLVLEEFMPLKNIGIDAA | AADKMGNPPS |
| O_28883.m000751 | 71 | 5.21e-14 | SECSEQTTST | DGTANGTGTRSLVLEEFIPIKRINSSSH | NDNDNDDDNE |
| PEQU_26724 | 48 | 5.21e-14 | ITQAIEIRRQ | EMADDEYSGEVPVVEEFISLKPTSSSSP | SDKSLKLAGD |
| E_H00466.1 | 112 | 6.42e-14 | LKDAIERLKK | ESLKWEERERVPVMEEFIPLKSESSYDE | GRAKPSNISQ |
| 1PabHRS1_14087g0010 | 54 | 1.32e-13 | ADCERASLHE | DVQNAPNTNGRPVLEEFIPIKKSYQATE | NKLEENHVAK |
| Z_2P13380_001 | 80 | 1.32e-13 | LSEVIEELRR | EIDRCHGESFGCIVEEFIPIKSKVGDDG | GIKVESDCKD |
| J_3871943 | 67 | 1.79e-13 | TATDNVYGQS | ECSEQTTGQCTPVLEQFLTIKDSSPFNE | EEVEEEFEDE |

| | | | | | |
|---|---|---|---|---|---|
| D_12g006800.1 | 80 | 2.42e-13 | MEASRQQLHS | HRENNHIGQMRPVLEEFIPLKNNNASVE | LGDEKLVANN |
| 3AtrHRS3_093.40 | 46 | 2.95e-13 | LELSNEEVPA | RSGYGETSGEDPPFEEFFPLKRSTSETT | QAPAGEKKRK |
| AA_1881G0390 | 96 | 3.59e-13 | YRQQLEAYQM | ESQGGAAAARPLVLEQFIPLKNIGIDAA | DKPGNPPSEK |
| Q_208g00340.1 | 63 | 7.10e-13 | NNAMEAYKEQ | LESYRGEKRTSVLLEEFIPIKKNYDEIQ | KVSSNSNNGD |
| Q_178g00480.1 | 59 | 1.14e-12 | AAAMPILVVD | EEEERCVDDSTPILKEFIPLKPTVLKKL | NEDDDDDEGD |
| K_02G098800.1 | 66 | 3.18e-12 | TVAEYNLNAQ | SECSEQTSTDGPVFEEFIIPIKKKASQA | SVEEDDDEEH |
| P_18G0031200 | 71 | 4.18e-12 | FPPRVDFRSE | HEDEELISDGAPALEEFIPMKLNSSTPE | ADDEESGDTA |
| P_0G0029600 | 19 | 4.18e-12 | LHAIEDLEEE | LARCGGGGAAHPKLEEFIPINRKFEEAG | RSKDEEDRRE |
| Z_9P09630_001 | 75 | 4.58e-12 | LTDVIEGLKK | ELERCRGQKLANAFEEFIPIRRKCEEEA | GVKLEADYED |
| Z_9P26570_001 | 78 | 5.01e-12 | LTDLIEGLKQ | ELKQCRDGRPTHVFEEVIPVKRKREDEG | GLKPEADCKD |
| J_3839360 | 70 | 5.48e-12 | ATMENLCGQS | ECSEQTTGECGGILDLFRPIKHSSTSLE | GEEEEVDADD |
| W_01g08160.1 | 79 | 5.48e-12 | LEAYQMGSQH | SAAAAAAARAPLVLEEFIPVKNIGIDVV | AADKAAAAGG |
| PpaHRS3_11430V3.1 | 99 | 1.02e-11 | ASLQTSVLDA | QDSDYERPKRTLVLENFMPLKRRWEKQL | KEKIFQNDAG |
| P_2G0120600 | 77 | 2.65e-11 | DTIEGVKEEL | AGCRDGGGGGRPALEEFMPVNAKLEEAE | GGVRDEKDFR |
| PpaHRS2_8580V3.1 | 102 | 2.88e-11 | KANSMDSGKD | QHGVSSGPGCRHTLEEFMPLKRRWERDQ | PSVSSDGERG |
| Z_6P00650_001 | 79 | 5.20e-11 | TDVIDGLKKE | LGQCKMGESCARVLGEFMPKKSKFEEKG | AVNPEADCKE |
| Z_1P23220_001 | 51 | 6.15e-11 | QTIENYMQLM | MMEDTVRTSNETVLKEFISLKPCSGASE | ENKDPDPVPE |
| Z_4P23210_001 | 78 | 6.68e-11 | LGEVIEGLNK | EIERFRCERFGRAFEEFVPVKSKVEEGG | GIKEETEYKD |
| Z_10P28540_001 | 79 | 7.25e-11 | LTDVIEGLEK | ELEKCRGERCARVFGESIPIKEKFEEEG | REKVEKDCEA |
| V_2g04810.1 | 105 | 1.01e-10 | QLEACQMGSS | HGGTAGPGRAPLVLEEFIPLSKNIDAAD | QNQQAASENP |
| SfaHRS4_40s0035.1 | 95 | 1.09e-10 | TPQPLQLRCS | TPEKSSNGNNMLVLKEFMPLKKRQCQSL | RKEEVGNDLQ |
| SfaHRS3_1s0189.1 | 100 | 1.19e-10 | LRCSTPEKSQ | SNVSSPNDNNKLVLKEFMPLKKRRLERL | QGGDQLLGGD |
| J_3843187 | 35 | 1.29e-10 | ATYNLYRRPE | CSEQTIGECRHVVLEKFLTIKDSSPSNE | EEDEEFDYEH |
| D_02g090400.2 | 80 | 2.64e-10 | KHAIERLKAE | ALLYKEKDKSPVMMEEFIPLKKGNSDEI | GRVKKSNDLS |
| H_6.397 | 74 | 5.35e-10 | LTDVIARLEE | EKVQYAELNDRSAVEEILPLKRISEEDG | AANRERDGCD |
| Q_33g00510.1 | 71 | 5.78e-10 | HVIDVLNDDL | MKRGAQSSNARPILEQFLPIKDDKEMMV | AEKDCKDNKM |
| PpaHRS1_4480V3.1 | 101 | 1.15e-9 | RSGSMDSGRN | QHVMPSSPGSRPTLEDFMPFKRRWEQRD | QPSESSDVEE |
| SmoHRS2_66160 | 37 | 1.80e-9 | EGNCSYRDEA | IGHDSVTAKSDSVLEEFIPPKTLEESKR | DEKACDPDLD |
| J_3829805 | 85 | 2.42e-9 | HIQPEGSERT | TSECEGRDGCIALCEEFVPIKSCERVKD | DEAENNIDKK |
| Z_10P19700_001 | 48 | 4.34e-9 | VTRAIESVRH | VMGDDEKVNHGPVTEELIPPMSEQRSEA | KKSAAVGSEM |
| G_G019009t1 | 83 | 5.01e-9 | ALGTLKDEFM | SIKNGMESETGRPLMEFLAIKRKHYEEE | QTDVSAAEKY |
| 4ScuHRS4_13.g005748 | 67 | 2.33e-8 | NEGLLNSALQ | QETDIRSRSTQPLLDMFLPSKKLVADSS | LQVNYSKEND |
| 1ScuHRS1_185.g025023 | 101 | 9.40e-8 | TWSHIQQKSN | SDDADSKPESRIIIEEFLPVNRAPTCTG | RNAEEVQQRP |
| SfaHRS2_4s0229.1 | 130 | 1.38e-7 | LVQDERDTGT | TVSSPAASGGHQRGLEFIPLKRLSETQQ | LRDDEGDDDD |

### HRS Motif 13

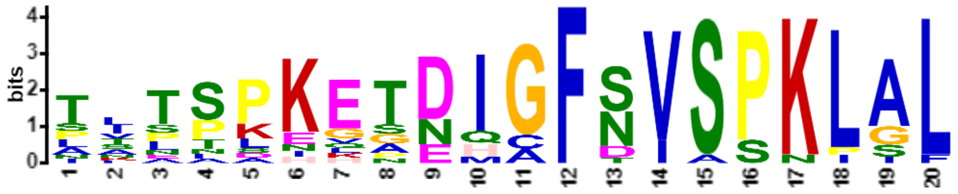


| Name | Start | *p*-value | Sites | | |
|---|---|---|---|---|---|
| G_G037970t1 | 131 | 4.64e-22 | QAGNETKPQS | SITSPKETEIGFNVSPKLAL | DTKPRNGGAF |
| K_17G119600.1 | 131 | 1.39e-21 | QASEGTKQQP | TITTPKESDIGFSISPKLAL | DNKQRNGGGA |
| M_018G074200.1 | 130 | 2.33e-21 | QDSNESKPQT | TLTSPKQTDIGFNVSSKLAL | DTKQRNGGAF |
| O_29807.m000474 | 130 | 7.02e-21 | QESNEIKPQT | TFNSPKETNIGFNVSPKLGL | DTKQRNGGAF |
| N_14G127500.1 | 131 | 1.12e-20 | DSNETKPQTT | LTTSPKETNIGFNVSPKLGL | DTKQRNGGAF |
| H_127.55 | 137 | 1.27e-20 | TKPQPTTIIS | SSLSPKETDIGFNVSPKLAL | DTKQRNGGAF |
| K_05G011500.1 | 131 | 8.23e-19 | ASSEGTKQQP | PITTLKESDIGFSISPKLAL | DNKQRNGGGA |
| M_006G155200.1 | 130 | 1.11e-18 | QESNETKPQT | TLTSPILTDIGFNVSSKLIL | DTKQRNGGAF |
| B_003614 | 128 | 2.63e-17 | SSDGINKEQA | TVASAKEGDHCFSVSPKLAL | DNKQRNGGAF |
| F_1015462001 | 130 | 3.43e-17 | PAGDETKQQS | TLTSSKEPDIGFTVSPKLGF | DNKQRNGGAF |
| N_06G060000.1 | 132 | 5.30e-17 | ESNETKLQTT | LNTFPKKTNIGFSVSPKISL | DTKQRNGGAF |
| K_06G213400.1 | 137 | 1.74e-16 | SEGTKPQSTI | TSLPKEGADIGFSVSPKLAL | DNKHRNGGAF |
| Y_30s1025391g002 | 132 | 3.90e-15 | ANDATRNQPM | AAPPLKETEHAFDVSPKLAL | DTKQRNGGAF |
| Z_6P05330_001 | 130 | 3.90e-15 | SPPVDAAKQQ | PVPPPKETEQAFDVSPKLSL | DTKQRNGGAF |

| | | | | | |
|---|---|---|---|---|---|
| B_006061 | 133 | 1.03e-13 | SDGSKQQPAT | AASAKEVGDQCFSVSPKLAL | DSKQRNGGAF |
| D_12g006800.1 | 155 | 1.71e-12 | KQQVQISTTC | NKENDHHNNIGFSIASKLAL | DNNNSSCPSP |
| K_04G151000.1 | 134 | 1.83e-12 | ASEGTKPQST | ITSPKNGADMGFSVSPNPAL | DNKHRNGGAF |

### HRS Motif 14

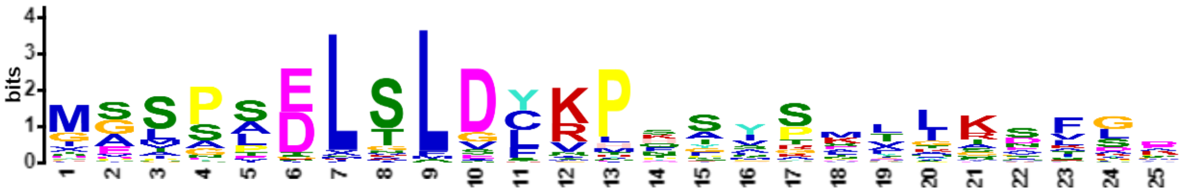


| Name | Start | *p*-value | Sites | | |
|---|---|---|---|---|---|
| H_127.55 | 1 | 5.31e-28 | | MASPSELSLDCKPHSYSMLLKSFGD | QTTADHAQKL |
| G_G037970t1 | 1 | 3.90e-27 | | MASPSELTLDCKPHSYSMLLKSFGD | QQIDQTQKLE |
| N_06G060000.1 | 1 | 3.90e-27 | | MASPSELTLDCKPHSYSMLLKSFGD | QNDHQTQKLE |
| O_29807.m000474 | 1 | 3.90e-27 | | MASPSELTLDCKPHSYSMLLKSFGD | QNDHTQKLEE |
| M_018G074200.1 | 1 | 5.10e-27 | | MASPSELSLDCKPHSYSMLLKSFGE | QNDQTQKLEE |
| N_14G127500.1 | 1 | 1.53e-25 | | MASPSELTLDCRPQSYSMLLKSFGD | QNDHTLKIEE |
| F_1015462001 | 1 | 2.97e-25 | | MASPSELSLDCKPHSYSLLLKSFGD | QPDQTQKLED |
| 5AthHOH4_AT2G03500 | 1 | 2.97e-25 | | MASSSELSLDCKPQSYSMLLKSFGD | NFQSDPTTHK |
| B_006061 | 1 | 1.60e-24 | | MGSPAELSLDCKPHSYSLLLKSFGE | QTDQTQKLEE |
| J_3875404 | 1 | 1.60e-24 | | MASSSELSLDCKPQSYSMLLKSFGE | NFQSDQTTQK |
| K_05G011500.1 | 1 | 1.60e-24 | | MASPSELSLDCKPQSYSLLLKSFGD | QTDHSYKLEE |
| M_006G155200.1 | 1 | 2.92e-24 | | MASPSELSLDCKPHSYSMLLKSYGH | QNDETDKLEE |
| K_17G119600.1 | 1 | 4.33e-24 | | MESPSELSLDCKPQSYSLLLKSFGD | QTDQTYKLEE |
| B_003614 | 1 | 1.64e-23 | | MGSPAELSLDYKPNSYSLLLKTFGD | QTQNLEEILS |
| L_4g113140.1 | 1 | 6.98e-23 | | MASSSELGLDCKPHSYSMLLKSFGE | QSDQSYKLEE |
| J_3853942 | 1 | 1.40e-22 | | MASSSELSLDCKPQRYSMLLKSFGD | NFQSDQTTQK |
| Y_30s1025391g002 | 1 | 2.18e-20 | | MGSPSDLSLDYKPNSYPMIQKPLGE | QPDQTQKIQE |
| Z_6P05330_001 | 1 | 1.37e-19 | | MASPAELSLDYKPNSYTMIQKQIGE | QPDQTQKIQD |
| X_9781.1 | 1 | 3.16e-18 | | MASPSDLSLDYKPNNYPIIQKPVGE | HPNQTQKIQE |
| K_02G135400.1 | 2 | 8.46e-18 | M | GSVPAELSLDLRPTFIPKTITDFLR | HLSDNNNPPA |
| N_17G094500.1 | 2 | 8.46e-18 | M | GSLPPELSLDFRPTYVPKTISDFLN | EVSIIGDISE |
| N_15G143500.1 | 2 | 9.55e-18 | M | GSVPPDLSLDFRPTYVPKTISDFLK | EVSLIGDVSE |
| D_01g108300.2 | 2 | 1.74e-17 | M | GSIPAELSLDCRPSFIPKSITDFLR | QLSLIRNVPD |
| G_G006998t1 | 2 | 3.95e-17 | M | GSVPPELSLDFRPTFVPKTISNFLK | EVSMVGNVSD |
| E_H01532.1 | 2 | 3.95e-17 | M | MTSPSELSFDCKPQIYSMLPKPFGE | QILSCDQQTQ |
| K_04G151000.1 | 1 | 9.83e-17 | | MSSQVELSMDYKPYSYSTLLKSYAD | ETETDQTHKL |
| O_29780.m001324 | 2 | 1.54e-16 | M | GSLPPELSLDFRPSYVPKTISDILN | EVSLIRNVSD |
| F_1031804001 | 2 | 2.67e-16 | M | GSIPPELSLDLRPSYMPKTINDFLS | GISTIGDVSE |
| O_29676.m001636 | 1 | 3.70e-16 | | MELSLDLSLVYVPKTISEYLKEVSK | VKDSSLKLSK |
| M_005G134600.1 | 1 | 9.67e-16 | | MELSLDLSLVYVPKAISECLKEVSM | VKDGSQKLPN |
| P_27G0017800 | 11 | 1.63e-15 | MERESSQKAW | MGSPSELTLDFRPHGHSAALRAIAD | QSDQTQNLED |
| L_4g086835.1 | 3 | 3.02e-15 | MG | CVVPAELSLDLRPTFVPKTIANFLF | HLSTIQTTSD |
| Z_10P27180_001 | 1 | 9.06e-15 | | MGSPPELSLDYKLDNYTAVAKAAGA | QPEQNQKIQE |
| B_011780 | 2 | 9.99e-15 | M | GSTSPELSLDFKPTYVPQTISGFLR | ELSTIGDISQ |
| G_G000558t1 | 1 | 2.39e-14 | | MELSLDLGLAYVPKSISEFLKEVSN | IKNGFQRLSK |
| N_13G128300.1 | 1 | 2.63e-14 | | MELSLDLSLVYVPKTISEYLMEISG | IKDSRRKLSK |
| H_6.397 | 1 | 6.12e-14 | | MDLSLDLSLVYVPKTMSEFMRDISK | TKNISAKVSK |
| B_006595 | 2 | 6.71e-14 | M | GSTSPELSLDFKPWQAPKTICCFLR | ELSAIEDIPE |
| MpoHRS1_0119s0041 | 1 | 7.36e-14 | | MGSPADLTLGCRPQSSGNEFKAVST | SGDQIERVRK |
| M_007G039400.1 | 1 | 2.00e-13 | | MELSLDLSLVCEPKTINECLKEVSM | VKDGSQKLTK |
| K_20G009800.1 | 4 | 2.61e-13 | MGL | VVPEEELSLDLRPSFVPKTITDFLC | HLSTTPNASH |
| K_07G178500.1 | 4 | 5.27e-13 | MGL | VIPEEELSLDLRPSFVPKTITDFLC | HLSTTPNASV |
| K_11G058600.1 | 1 | 6.82e-13 | | MELSLDLSLGFVPKPLSLFFADVSA | NRDKVATLDG |
| G_G019009t1 | 3 | 1.24e-12 | MA | SSSSLELNLSLKPSYVPKTIANLLK | DLSHVDTASD |
| R_2G100176 | 8 | 2.05e-12 | MGLDVME | IGMGADLSLDLRHFASKAVRQSKDD | TPAPDMDACI |

| | | | | | |
|---|---|---|---|---|---|
| R_2G348238 | 2 | 2.23e-12 | M | ASSPSDLTLDYKPNGNGAAYAVTTP | KPPQETLVVD |
| S_2G016300.1 | 42 | 3.10e-12 | IATMGLDVVE | IGMGADLSLDLRHFATKAVRQSKDD | APAPDMDACI |
| S_3G046800.1 | 2 | 3.65e-12 | M | ASSPSDLTLDYKPNGNAAAYAVTIP | KPQQEPLVDG |
| AA_0456G0940 | 8 | 4.65e-12 | MGLDVAE | IGLGTDLSLDLKMFASRSVGRVKDA | PASAMDDCIR |
| N_12G093900.1 | 1 | 6.42e-12 | | MELSLDLGSVYVPKTITEYLREVSK | VKDSSQKLSK |
| AA_2901G0070 | 8 | 6.96e-12 | MGLDVAE | IGVGTDLSLDLKMFAARSVGRVKDA | PASAMDDCIR |
| R_2G173882 | 8 | 8.83e-12 | MGLDVVE | IGMGADLSLDLRHFASKAVRQSKDY | APAPAPDMDA |
| C_7G203800 | 2 | 9.56e-12 | M | VSITSEVCQDSKPHSYSMLLKSFVD | HPEQTQKLEE |
| AA_1881G0390 | 2 | 9.56e-12 | M | DSSPSDLTLDYKPNGGGGTAAGAYP | MIIPKQAPPI |
| AA_2274G0070 | 2 | 1.12e-11 | M | DSSPSDLTLDYKPNGGGGTVAGAYP | MIIGKQAPLV |
| T_5G124700.1 | 2 | 1.21e-11 | M | ASSASDLTLDYKPNGNSNGGGAYAV | IPKQQQETLV |
| 2AtrHRS2_033.259 | 1 | 2.09e-11 | | MGSPSELSLDYKPPCKGEERFSKPS | GANERFFKAV |
| W_03g55590.1 | 8 | 2.43e-11 | MGLDVGE | IGMGLDLSLDLKMFAARSAVRMAAA | AAKEATGVEA |
| T_2G013500.1 | 8 | 4.81e-11 | MGLDVLE | IGMGSDLSLDLRYFASKAVRQARDA | PASDVDACIR |
| W_07g02800.2 | 9 | 6.02e-11 | MGLDVGEI | GVGAADLSLDLKMFAAKSFGRVRGK | DTTTTAMGDC |
| O_29790.m000831 | 11 | 7.52e-11 | MASSSNARSQ | FDLALGLKLSYVPKSIPNLIKDLSR | DESSDEVKIL |
| K_07G209500.1 | 3 | 8.71e-11 | MG | SVAAEELNLDLRSSFVPKTITDFLR | HLSANNNHPA |
| V_2g04810.1 | 2 | 1.09e-10 | M | DSSPSDLTLDYTPNGNGNGGGGGGH | PMTPKQAPLV |
| K_01G183700.1 | 25 | 1.56e-10 | QRKTPKSNII | MEPSLDLRLGFVPKPLSLFFGDVSG | NRDKCDKVVT |
| G_G019003t1 | 2 | 2.57e-10 | M | NPTYVDLSLSLKPSYVPKSISNLLE | DLSKIDNESD |
| PpaHRS2_8580V3.1 | 1 | 4.20e-10 | | MGSPIDLTLGCNTQAPSSNGRYNDH | RNSSACLEDC |
| SfaHRS3_1s0189.1 | 1 | 5.17e-10 | | MASPPDLSLGCQTQAPSSASTITSS | EVSNPLVNNQ |
| K_06G213400.1 | 2 | 5.17e-10 | M | PSQAELSMMDYKPYSYSTLLKSFLD | QTETDQTYKL |
| 1ScuHRS1_185.g025023 | 1 | 7.80e-10 | | MGSPTSLVLECRHSSSTTTAANSGS | CRNFSNTADD |
| D_02g090400.2 | 6 | 8.93e-10 | MGSNS | KEMNIDLNFVYVPKLISDVLTEVSA | MDDISKKLKK |
| N_01G103800.1 | 59 | 1.09e-9 | VFADEALRCL | YAFGLGLKLSYVPQSLPNAIKDLSI | GESNEVKLLL |
| V_1g07630.1 | 8 | 1.25e-9 | MGLDVGE | IGAPLDLGLDLKLFVARTAGRLAAA | KEAPSMDACI |
| H_62.51 | 2 | 3.57e-9 | M | GSIPPELSLDFRPSSFVPKTISDFL | KEVSRIGSVS |
| Z_10P28540_001 | 3 | 3.57e-9 | MG | SSAAADVSLDLKLFTARTVAGFLKE | ALVMEIGDGR |
| M_009G106600.1 | 2 | 4.33e-9 | M | GSLSSELSLDFARPSTAKALPFLPK | TIADFLKEVS |
| SfaHRS1_10s0233.1 | 1 | 6.34e-9 | | MESPVDLTLGSAHTASSSYTNATNT | VNGLESSLSF |
| D_12g006800.1 | 4 | 7.65e-9 | MNM | ASSNSELSLECKPHQSYSMLLKSFG | EKNVDQTQNL |
| Z_6P00650_001 | 2 | 7.65e-9 | M | GSAAADASLELRLFAARSVTRSLKE | APAMEKGDGR |
| Z_6P34680_001 | 2 | 7.65e-9 | M | GSAAAEFGLDFKLCAMRTVGGFLKE | AAAAVQSSDG |
| MenHRS1_2781 | 622 | 1.11e-8 | DGGDGRAAER | RPSPLEAALDYKAPNLDMTLGSRGE | YTPPRLKQDE |
| 3AzfiHRS4_114.g046051 | 1 | 2.45e-8 | | MGSPTTLVLECRRPSAAYTATTSTA | NSGCRSSNFT |
| PpaHRS3_11430V3.1 | 1 | 2.45e-8 | | MGSPANLALGPQVSTITSTIETALN | CTVSADGLEA |
| L_5g017980.1 | 5 | 2.45e-8 | MEQQ | QQLNLDLTLASVPKTVSHFLNNVVQ | TKDMSQKLSM |
| SfaHRS2_4s0229.1 | 1 | 3.11e-8 | | MGSPADLTLGTSQAPSTNATTNTAA | ATTTTTVLSK |
| 2ScuHRS2_42.g012676 | 1 | 1.93e-7 | | MGSPTSLILGRNPSSCSIRTRNERG | CLVNLHLGDG |
| C_6G095900 | 16 | 3.90e-7 | LDLKPTTTIT | TSSSSTITTNYMPESIHVFLGEVSL | IGNVSEKLSK |
| M_004G144800.1 | 21 | 7.32e-7 | DFVRPSSTTK | TFTSAIKTFTFLPETITDFLKEVSM | IGDAAVKGLK |
| J_3862509 | 4 | 1.74e-6 | MVQ | TDTDQRMSLDLNLYSLAKPLSQFLN | DVSKIKDNHS |
| PpaHRS1_4480V3.1 | 224 | 3.28e-6 | SFLSRPTPRT | VASAADLSLSLGERTTTPSSRNLRP | SDSEVGSIDA |

### HRS Motif 15

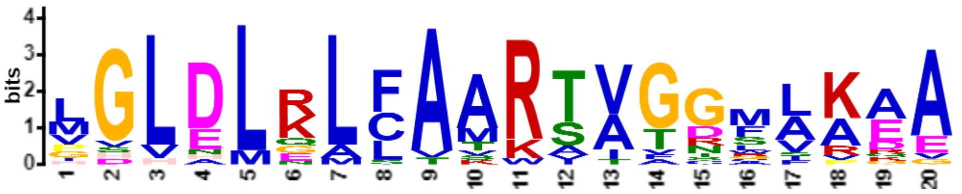


| **Name** | **Start** | ***p*-value** | **Sites** | | |
|---|---|---|---|---|---|
| S_1G078900.1 | 14 | 1.12e-21 | DVGEIGMGLD | LGLDLRLFAARTAGGMAAAA | AKGAPAGIES |
| T_9G080000.1 | 14 | 3.50e-20 | DVAEIGMGLD | LGLDLKLFAARSAGGMAAAA | AKGAPAGIEA |
| R_2G171468 | 14 | 1.19e-19 | DVGEIGMGLD | LGLDLRLFAARSAGGMAKGA | APAGIQSCIR |
| Z_4P23210_001 | 7 | 1.33e-19 | MAVAEE | MGLEMKLCAMRTVGGFLKEA | SAIECGDGGR |

| | | | | | |
|---|---|---|---|---|---|
| AA_3297G0100 | 1 | 2.04e-19 | | MGLDLKLFAARSAGRMAAAA | KDAPAVDACI |
| R_2G124540 | 14 | 2.52e-19 | DVGGIGMGLD | LGLDLGLFAARSAGGMAAAA | KGAPAEIESC |
| Z_9P09630_001 | 4 | 1.30e-17 | MAG | IGLELRLCASRTVGGFVKQA | AAVVETTDRG |
| Z_4P32500_001 | 11 | 1.42e-17 | MIEPAAMGSE | VGLALQLCAMRTVGGFVKDA | ATESAGRAAR |
| AA_1128G0130 | 10 | 1.54e-16 | MGTSHSEAV | PGLDLGLFAAKTVGNWLKEG | VASASSEQEE |
| AA_1535G0300 | 5 | 5.52e-16 | MGLD | VGLNLKLFAAWSAGRMAAAA | KEAPAVDACI |
| Z_9P26570_001 | 8 | 1.39e-15 | MGSAVAD | VVLDLNLCAVRAVGGSLKEA | AALESGDGRV |
| Z_4P28210_001 | 8 | 2.36e-15 | MGSAVFE | MGLELELCATRVVGDFVKEV | SAIESGGSGR |
| AA_0185G0030 | 11 | 1.35e-14 | MGSSVQKETV | GDVDLRLLAARTVTDSLRAA | LSRSSAGEKA |
| AA_0930G0300 | 12 | 1.56e-14 | GTSTSHSEAI | PGLDLRLLVAKTIGNLLKEE | GVASGSSEQE |
| W_12g39640.1 | 23 | 2.72e-14 | EAAAAAAFVL | GGVDMRMLAARTATGALARA | GGGEAAAAAA |
| PEQU_27343 | 12 | 3.35e-14 | GDKMGLLAPE | LGLDLKLFTKRTIFSCLKDE | IQAGSRAAKL |
| V_1g58830.1 | 14 | 6.18e-14 | EAGESAMGGD | LSLDLQAFAARTVAGRIPAA | SREDVLRKLE |
| AB_07.3042 | 5 | 6.09e-13 | MAPD | LGLDLKLSATKIVTTLSKEA | SSIQDDDKKI |
| Z_2P13380_001 | 8 | 2.53e-12 | MGSALAE | MGLELELCAMRTTVCGFVKE | ASAIESVGGG |
| T_3G370000.1 | 9 | 2.78e-11 | MMARKEDH | HHHHLRALAARAVTDSLRAA | ASRATDADRA |
